# UCL-MetIsoLib: A Public High-Resolution Tandem Mass Spectrometry Library for HILIC-Based Isomer-Resolved Profiling of Glycolysis, Central Carbon Metabolism, and Beyond in Urine, Plasma, Tissues, Cells, and Patient-Derived Organoids

**DOI:** 10.1101/2025.07.04.663184

**Authors:** Kristian Serafimov, Maria Virginia Giolito, Sébastien Ibanez, Ophélie Renoult, Sarah Srhir, Paula Varon Rugeles, Florine Laloux-Morris, Cyril Corbet, Julien Pierrard, Marc Van den Eynde, Michael Lämmerhofer, Olivier Feron

## Abstract

We present UCL-MetIsoLib, a publicly accessible high-resolution tandem mass spectrometry (HRMS/MS) library developed for HILIC-based, ion-pairing free, isomer-resolved metabolomics using a bioinert UHPLC system and the Acquity Premier BEH Amide column. The platform integrates two complementary methods operating under distinct chromatographic conditions (pH 3.5, ESI+; pH 11.0, ESI−), enabling broad metabolic coverage. A total of 334 metabolites are annotated in the library structure, with thiol derivatization incorporated into the extraction protocol to mitigate redox-driven artifacts. Metabolite identification is supported by 245 authentic reference standards and curated according to MSI Level 1 and Level 2 criteria. Validation followed FDA guidelines for bioanalytical method validation and was performed across five biological matrices—urine, plasma, tissues, cultured cells, and patient-derived colorectal organoids—with a U-^13^C, U-^15^N-labeled Amino Acid Mixture used as an isotope labeled internal standard. The method demonstrated high precision (<15% RSD intra-/inter-day) and recovery (85–115% across all QC levels). To demonstrate biological applicability, UCL-MetIsoLib was applied to a case study comparing healthy and colorectal cancer-derived organoids. The method enabled confident annotation of metabolite isomers, including key glycolytic intermediates such as DHAP and GA3P, as well as sugar phosphates from the glycolysis and pentose phosphate pathways. Metabolic alterations were observed in tumor organoids, including accumulation of nucleotide derivatives and shifts in central carbon metabolism. These findings emphasize the value of isomer-resolved spectral libraries in detecting biologically meaningful differences that are often missed in conventional untargeted metabolomics workflows.

---

High-resolution mass spectrometry (HRMS) has become a cornerstone of untargeted metabolomics, enabling the detection of thousands of features in complex biological matrices^1,2^. However, the confident identification of these metabolites remains a major bottleneck, particularly for structurally related compounds and isomers that cannot be distinguished based on m/z values alone^3,7,8^. The annotation challenge is further compounded by the diversity of sample types encountered in translational research — including biofluids, tissues, cell cultures, and emerging patient-derived models such as organoids — each introducing unique matrix effects and analytical variability^3,9^. While public MS/MS spectral libraries such as HMDB, MassBank, and GNPS have accelerated metabolite annotation efforts^4,5,6^, they often lack critical contextual information such as chromatographic retention times, ionization mode specificity, and validation across biological matrices^6,7^. Moreover, these resources typically do not resolve isomeric compounds, many of which share near-identical fragmentation patterns, leading to ambiguous or incorrect annotations in data-dependent acquisition (DDA) workflows^2,8^. There is a pressing need for high-quality, matrix-aware MS/MS reference libraries that enable isomer-resolved metabolite identification using reproducible and accessible analytical platforms^3,7,9^. In this context, hydrophilic interaction liquid chromatography (HILIC) has emerged as the preferred separation technique for polar and semi-polar metabolites. Hydrophilic interaction liquid chromatography (HILIC) has emerged as the preferred separation mode in metabolomics.^10–14^ This preference is due to its ability to retain hydrophilic analytes, without the need for ion-pairing reagents, which are known to contaminate MS systems^15,16^, and its strong compatibility of HILIC with ESI-MS. Various stationary phases have been described in the literature, with zwitterionic sulfobetaine-coated^17–20^ and amide-linked^21–23^ phases being the most commonly used in metabolomics studies. Undoubtedly, HILIC presents certain analytical challenges, specifically the frequently encountered issue of shifting retention times, for which a solution has recently been proposed.^24^ However, the most significant challenge in HILIC metabolomics studies remains the accurate separation of molecular isomers and potential metabolites that may interconvert in the ion source during ionization process as a result of in-source fragmentation e.g. Fumarate/Malate, AMP/ADP/ATP (attributable to other nucleotides as well). There are numerous isomeric species among multiple biochemical pathways that are of critical importance in every biological sample e.g. sugar phosphates and their metabolites related to the glycolysis and pentose phosphate pathway^25–27^, nitrous oxide cascade related metabolites such as asymmetric and symmetric dimethylarginine (ADMA/SDMA)^28^, sugars^29^, organic acids^30,31^ and many more. Further complicating untargeted metabolomics workflows are additional isomeric or structurally related metabolite pairs. Citrate and isocitrate, key intermediates of the TCA cycle, are difficult to resolve chromatographically due to their nearly identical structures. Amino acid isomers such as leucine and isoleucine exhibit near-identical MS/MS fragmentation patterns, complicating confident annotation. Methylmalonate and succinate represent another pair of structurally related metabolites relevant to mitochondrial and vitamin B12 metabolism, which are challenging to separate under standard HILIC conditions. Other notable examples include choline and betaine, compounds central to methyl group metabolism and osmoregulation, and nucleotide sugars like UDP-glucose and UDP-galactose, which are critical in glycosylation pathways yet difficult to resolve without tailored chromatographic strategies. Even redox cofactors such as NAD^+^, NADH, NADP^+^, and NADPH can exhibit overlap due to similar retention characteristics and fragment spectra. In many of these cases, tandem mass spectrometry alone does not suffice to solve the issue of assay specificity, as the obtained product ions are not characteristic of the individual metabolite, with the only differences being in the MS^2^-fragment ratios^32^. Additionally, the choice of chromatographic hardware plays a crucial role in metabolomics analysis. Conventional liquid chromatography systems often incorporate stainless steel components, which can present challenges when analyzing polar and labile metabolites. Stainless steel surfaces are prone to adsorbing certain analytes, especially phosphate- and carboxylate-containing metabolites, leading to reduced recovery, peak tailing, and overall signal suppression. These interactions can compromise data reproducibility and metabolite coverage, particularly for metabolomics workflows aimed at capturing a broad range of small, polar compounds. To mitigate these issues, the implementation of biocompatible LC systems—utilizing inert materials such as PEEK (polyether ether ketone) and various alloys for fluidic pathways (e.g. MP35N, as used in this study) - has gained attention. These systems minimize non-specific analyte adsorption and metal-induced degradation, ensuring improved metabolite recovery and enhanced chromatographic performance. Considering all of the above, we introduce UCL-MetIsoLib, a publicly available HRMS/MS library developed at the IREC Metabolomics Facility, specifically designed for HILIC-based isomer-resolved metabolomics.

The library comprises 334 curated metabolites, acquired under optimized dual-polarity conditions, and includes both MSI Level 1 and Level 2 annotations supported by 245 authentic standards and in silico spectral verification. The method was analytically validated across five biological matrices—urine, plasma, tissues, cultured cells, and patient-derived organoids—in accordance with FDA bioanalytical guidelines, making it directly applicable to both experimental and clinical metabolomics workflows. To demonstrate its utility, we applied UCL-MetIsoLib to a case study comparing healthy and metastatic colorectal cancer organoids. All curated MS/MS spectra are provided as .msp files in the Supplementary Material, making UCL-MetIsoLib a ready-to-use, open-access resource for the metabolomics community (**Figure 1**).

**Figure 1.**
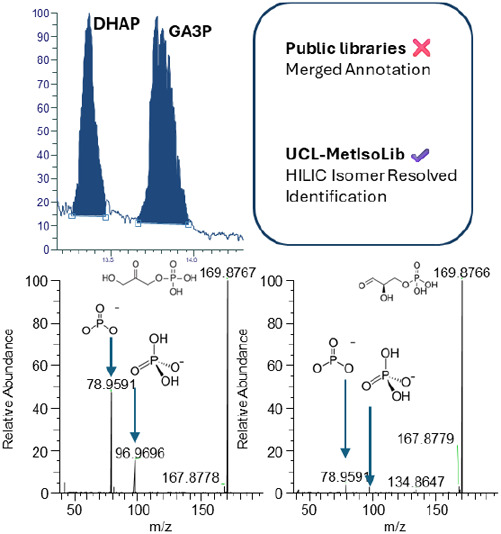
Isomer-resolved annotation of DHAP and GA3P using UCL-MetIsoLib. HILIC-based chromatographic separation of the isomeric sugar phosphates dihydroxyacetone phosphate (DHAP) and glyceraldehyde 3-phosphate (GA3P) demonstrates baseline resolution using the validated dual-polarity method. Despite nearly identical MS/MS fragmentation patterns in negative mode — both producing diagnostic phosphate-related ions at m/z 78.9591 (PO_3_^−^) and 96.9696 (H_2_PO_4_^−^) — UCL-MetIsoLib enables confident differentiation based on retention time.

## Experimental

### Materials

Formic acid, acetic acid and acetonitrile of Ultra LC-MS grade were supplied by Fisher Scientific (Brussels, Belgium). Ammonium hydroxide solution (Suprapur® quality 25% NH_3_ basis), NIST Plasma and N-Ethyl-Maleimide was purchased from Sigma-Aldrich (Merck, Hoeilaart, Netherlands). Uniformly U-^13^C, U-^15^N-labeled Amino Acid Mixture was obtained from Buchem BV (Apeldoorn, Netherlands). All standards used in this study were provided by Sigma-Aldrich (Merck, Hoeilaart, Netherlands). PFA bottles for mobile phases were provided by AHF Analysentechnik (Tübingen, Germany).

### Organoid culture

The colorectal cancer (CRC) organoid model was generated from colon cancer tissue obtained from a patient undergoing surgery at Cliniques Universitaires St Luc in Brussels (ethics committee approval ONCO-2015-02 updated on 13-05-2019 following the principles of the Declaration of Helsinki). The CRC-N organoid model was derived from an adjacent healthy colon of a patient undergoing surgery (ethics committee approvals B4032020000038 & B4032023000027) in accordance with the Declaration of Helsinki. Signed informed consent was acquired from the patients before sample collection, and all personal and clinical data remained strictly confidential for the investigators. All CRC organoid cultures were plated in 50-μl domes of Cultrex Basement Membrane Extract, Type 2 (#3532-010-02, Bio-Techne) and maintained in Advanced DMEM/F 12 (Thermo Fisher Scientific) supplemented with 1 % Glutamax (Thermo Fisher Scientific), 1% HEPES (Thermo Fisher Scientific), 1% penicillin-streptomycin (Thermo Fisher Scientific), 1% B 27 without vitamin A (Thermo Fisher Scientific), 1% N 2 supplement (Thermo Fisher Scientific), 50 ng/mL EGF (STEMCELL Technologies), 10 mM nicotinamide (Sigma-Aldrich), and 1.25 mM N-acetyl cysteine (Sigma-Aldrich). CRC-N medium was further supplemented with 10 nM gastrin (Sigma), 0.5 nM hWNT 3 surrogate (Thermo Fisher), 250 ng/ml Rspondin-1, 100 ng/ml Noggin, 500 nM A-8301, and 10 μM SB202190 (STEMCELL technologies). Organoids were typically passaged every 7–10 days, and the medium was changed every 2–3 days. Eight wells of 50 μl were pooled for metabolomic studies. Organoids were recovered using 500 μl Gentle Cell Dissociation Reagent (STEMCELL Technologies). The Cultrex dome was scraped and collected using a P 1000, then incubated on ice before being centrifuged for 5 minutes at 400 g at 4 °C. The pellet was washed three times with cold PBS and constantly kept on ice. The organoids were transferred to a pre-weighted 1.5-ml tube, centrifuged, and then snap-frozen in liquid nitrogen. After freezing, the tubes containing the organoid pellets were weighed to calculate the organoid mass for normalization. Pellets were stored at -80°C.

### Cell culture

The human CRC cell line (HCT-116) was purchased from ATCC (Manassas, Virginia, USA). The cells were cultured in DMEM (61965.026, Gibco) supplemented with 10% heat-inactivated FBS (F7524, Merck and 1% penicillin/streptomycin (15140-122, Gibco). The cells were regularly tested for mycoplasma contamination. After reaching 80% confluence, cells were washed with cold PBS, frozen by liquid nitrogen contact and scraped.

### Animal Experiments

Rj:NMRI-Foxn1nu/nu 5-week-old female mice were purchased from Janvier and housed in room with 12 h light/12 h dark cycles. Experimental procedure was authorized by the local ethical committee (reference 2020-032). After accommodation, mice were injected subcutaneously with 2 × 10^6^ FaDu cells in 0.9% NaCl. Mice were weighed once per week and tumors were measured 3 times a week. Mice were sacrificed when tumors reached 600 mm^3^, and tumors were collected and and immediately snap-frozen in liquid nitrogen.

### Sample preparation and metabolite extraction

Cell, tissue and organoid samples were prepared in homogenization tubes. 1 mL of extraction solvent was added (50/50 *v/v* MeOH/H_2_O with the MeOH fraction containing 20 mM of N-Ethylmaleimide (NEM) for derivatization), 20 μL internal standard solution of U-^13^C, U-^15^N-labeled Amino Acid Mixture and 0.15 g 1.0 mm zirconia/glass beads for homogenization of cells and organoids, for tissues, the same amount of zirconia/glass beads was used but in a diameter of 2.8 mm. After 10 minutes, the derivatization reaction was stopped by the addition of 10 μL concentrated formic acid. Derivatization and extraction have been performed as previously described^12,34^. Thereafter the samples were homogenized at 4 °C for 10 cycles (10 s per cycle, 5000 rpm, pause 30s for cells and organoids and 15 s per cycle, 6800 rpm, pause 30s for tissues) with Precellys Evolution high performance cell lyser (Bertin Technologies, France). The samples were then spun down for 10 min (20000 g at 4 °C), the supernatant removed and spun down once more for 10 min (20000 g at 4 °C). The supernatant was then removed and evaporated to dryness with a high-performance evaporator (Genevac EZ-2) (Genevac, Ipswich, UK). Reconstitution was performed in 50 μL ACN/H_2_O (50/50; v/v). After reconstituting, the sample was briefly vortexed for 10 seconds and spun down for 10 min (20000 g at 4 °C). The supernatant was taken and placed on a Whatman® Puradisc 4 syringe filter (PTFE, 0.2 um) fixed inside a 1.5 mL Eppendorf tube. By centrifuging for 2 minutes at 10000g at 4 °C, sample filtration was performed for extra purification and the filtered sample transferred to UHPLC vials equipped with silanized microinserts for sample analysis. In the case of urine and plasma, the steps were identical with no homogenization required. Sample amounts this protocol is based on are 100 μl of plasma, 500 μL of urine, 50 mg of tumor tissue, 3 million cells and 50 mg of patient derived colorectal organoid matrix.

### Untargeted HILIC-HR-MS analysis

LC-MS analysis was performed on a ThermoFisher (Dilbeek, Belgium) Vanquish Horizon system equipped with a binary pump, autosampler and thermostated column compartment. Chromatographic separation was performed on a Waters (Antwerp, Belgium) Acquity Premier VanGuard-FIT BEH Amide column (100 × 2.1 mm, 1.7 μm), equipped with a VanGuard-FIT BEH Amide pre-column (5 × 2.1 mm, 1.7 μm). Silanized microinserts were used during sample analysis to limit the interaction between phosphorylated compounds and glass surfaces. For metabolite analysis in ESI^+^ mode, a stock solution of 500 mM ammonium hydroxide was prepared and adjusted to a pH of 3.5 with formic acid. A following 1:10 dilution with water (mobile phase A) and with acetonitrile (mobile phase B) resulted in the final mobile phase conditions used during sample analysis in positive ionization mode. The chromatographic gradient was as follows: (0.0 min, 100% B; 20 min 70% B; 21 min 70% B; 21.01 min 100% B; 25 min 95% B) and a flow rate of 0.2 mL min^-1^ was used. The chromatographic separation in ESI^-^ mode differed. A stock solution of 1000 mM ammonium hydroxide was prepared and adjusted to a pH of 11.0 with formic acid. After a subsequent dilution of 1:10 with water (mobile phase A) and with acetonitrile (mobile phase B) the desired mobile phase conditions were achieved. Gradient elution followed a 2-step pattern and was performed as follows: (0.0 min, 95% B; 20 min 85% B; 25 min 60% B; 25.01 min 95% B; 30 min 95% B) and was carried out at a flow rate of 0.5 mL min^-1^. During both methods, a constant column temperature of 60 °C was maintained throughout the entire analytical run and in both cases the injection volume was 3 μL. The autosampler was kept at 4°C. Ion source parameters were as follows: In negative mode: sheathe gas (50, nitrogen), aux gas (10, nitrogen), sweep gas (0 psi, nitrogen), ion transfer tube temperature (325°C), ion source voltage -3000 V (negative mode), vaporizer temperature (350°C). In positive mode - sheathe gas (35, nitrogen), aux gas (7, nitrogen), sweep gas (0 psi, nitrogen), ion transfer tube temperature (325°C), ion source voltage +3500 V (negative mode), vaporizer temperature (275°C). Analytes were recorded via a full scan with a mass resolving power of 11150 over a mass range from 70 to 900 *m*/*z* (scan time: 100 ms, RF lens: 70 %). Normalized AGC Target was set to 500% (ESI^+^) and 100% (ESI^-^), Maximum Injection Time 100 ms and 1 Microscans were additional settings. To obtain MS/MS fragment spectra, data-dependent acquisition was carried out at a resolving power of 11150 over a mass range from 70 to 900 *m/z* and an isolation window of 2 *m/z*. RF lens was set to 70%, TopN was adjusted 10, Microscans set to 1, AGC Target was set to 100% (ESI^+^) and 50% (ESI^-^) with a Maximum Injection Time of 50 ms. Collision Energy was Nomalized to 30/50/70%. Dynamic Exclusion, Targeted Mass and Apex Detection were utilized as scan filters. Dynamic Exclusion Mode was operated in the custom setting (exlude after 1 hits for 10 seconds), Targeted Mass involved the utilization of an in-house metabolite inclusion list and Apex Detection was set to 30%. Blank solvents (mobile phase A and B) were injected in the beginning of the chromatographic batch to ensure proper column and system equilibration. Samples were first analyzed in negative and subsequently in positive ionization mode in randomized order. Data processing comprising peak finding, blank subtraction, feature alignment, normalization, MS/MS spectral deconvolution and score-based metabolite identification was performed with MS-DIAL software (version 4.9.221218)^34^. A similarity score of 80% was chosen as threshold for reliable metabolite identification. All metabolite annotations were supported by retention time matching, precursor mass, and MS^2^ spectral similarity against authentic standards and a curated in-house spectral library. Identification levels were assigned following the MSI guidelines (Level 1 corresponding to confirmed identity by authentic reference standard and Level 2 based on spectral MS/MS identification). An annotated metabolite list from the PDO metabolomics analysis with Log2 Fold Change, p-Values, Retention Times, MS/MS Fragments, and Precursor m/z is provided as part of the manuscript in the supplementary information under **Table S-1**. Normalization of metabolite intensities was performed using two complementary approaches: (1) Consideration of the biological variance of the material i.e. normalization to the wet weight of the organoid pellet, weight of the specific tissue, cell number or creatinine concentration in the case of urine (determined via LC-MS standard addition) to account for differences in biological input material, and (2) feature-specific normalization using a U-^13^C, U-^15^N-labeled Amino Acid Mixture as internal standard. The uniformly labeled extract provided broad coverage of isotopically matched internal standards across the metabolome, enabling correction for matrix effects, extraction variability, and instrument-related signal drift. Metabolites detected in less than 20 % of samples were filtered out. Missing values for the remaining metabolites were imputed using the Random Forest method in Python. Metabolites with an p value <0.05 and a fold-change greater than ±1.0 were considered significantly regulared. For peak finding an intensity threshold of 500 counts per second (cps) was selected. All further statistical processing and visualization was conducted with Python.

### Method validation

To assess method performance, PRM acquisiton was performed for a total of 245 authentic chemical standards with the previously mentioned ion source and chromatographic parameters. Determination of Matrix effect (ME), extraction recovery (ER), and process efficiency (PE) followed the Matuszewski protocol.^35^ ME was evaluated at a concentration level of 100 ng mL^-1^ in a pre- and post-extraction spiking experiment with organoid cell matrix after as follows:

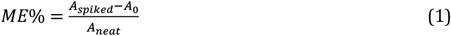

With *A*_*spiked*_ being the area of the post-extraction spiked organoid extract samples, *A*_0_ the peak area of the analyte endogenously present in the cell extract (previously determined by LC-MS standard addition quantification) and *A*_*neat*_ the peak area of the selected metabolite in neat solution. In the same manner, ER and PE were evaluated at the same concentration level via pre- and post-extraction spiked organoid extract samples and pre-extraction spiked versus neat standard solution, respectively. Calibration functions were generated by plotting normalized peak areas (peak area nominal divided by the peak area of the closest eluting isotope-labeled internal standard) versus the concentration of standard solutions (1, 50, 100, 500 and 1000 ng mL^-1^). Matrix matched calibration was performed as well by spiking a series of 5 concentration levels (1, 50, 100, 500 and 1000 ng mL^-1^) to the biological matrix (organoid matrix). Weighted linear regression (1/x^2^) was used for both external and matrix-matched calibration curves. Linearity was evaluated by the determination of the regression coefficients (r^2^) of the matrix matched calibration functions for each metabolite and by evaluating the back-calculation values of the concentrations of the non-zero calibrators regarding their nominal concentrations (spiked concentrations; endogenous amount as pre-determined via LC-MS and standard addition corrected), which should be in at least 75% of calibrator levels ±15%. Further evaluation and validation of the developed method, including intra- and inter-run accuracy and precision was carried out in organoid extract samples according to the FDA guidelines for bioanalytical method validation at 4 different concentration levels (QC_LLOQ_/QC_low_/QC_mid_/QC_high_ 10 ng mL^-1^/50 ng mL^-1^/ 500 ng mL^-1^/ 1000 ng mL^-1^). For the determination of LoDs and LoQs, the external calibration function was used for the individual metabolites as follows:

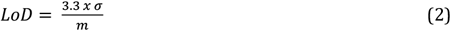

With σ being the standard error of the calibration function’s slope and m the slope value itself. Analogous to the LoQ determination, the following equation was used:

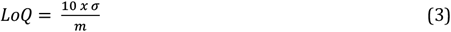

Individual metabolite accuracies (ACC) were calculated with quality control samples obtained by spiking organoid extract samples at four different concentration levels and corrected for endogenously present levels as:

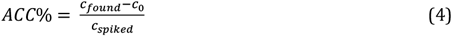

With *c*_*spiked*_being the concentration spiked to the organoid extract, *c*_0_the endogenously present amount (determined by LC-MS standard addition quantification) and *c*_*found*_the concentration of the target metabolite found in the spiked sample.

### Data processing

Xcalibur was used for data acquisition and system control. Freestyle was used for data evaluation. Skyline was employed for peak integration, linear regression (1/x^2^ as weight), and concentration calculation. Statistical analysis and visualization were performed with Python. MS-DIAL was used for metabolite curation and chromatographic batch alignment.

## Results and discussion

### Library Curation

The UCL-MetIsoLib library comprises 334 curated metabolites, distributed across two complementary acquisition modes using HILIC-based separation under distinct pH conditions: pH 3.5 for positive mode (ESI^+^) and pH 11.0 for negative mode (ESI^−^). All data were acquired on a high-resolution Orbitrap system using normalized collision energies (NCE) of 30, 50, and 70, capturing diverse and informative fragmentation profiles. Chromatographic separation was performed on a bioinert UHPLC system equipped with an Acquity Premier BEH Amide column, supporting high retention stability and resolution of polar and isomeric compounds. Spectra were curated from internally generated datasets, with manual inspection of all entries to ensure spectral quality, fragmentation logic, and chromatographic consistency. Annotation followed MSI Level 1 or Level 2 guidelines, supported by 245 authentic reference standards, while Level 2 annotations were cross validated through in silico fragmentation. Fragment curation and library export were supported by custom in-house Python scripts, which systematically filtered and consolidated spectral data. For each metabolite, multiple adduct forms were detected (e.g., [M+H]^+^, [M+Na], [M+NH_4_]^+^), and redundant spectra were collapsed to a single consensus entry. This was achieved by evaluating precursor mass proximity, retention time alignment, and shared high-intensity fragment ions across adduct types. Fragment selection was based on reproducibility across collision energies and ranked using a weighted scoring system favoring signal intensity, *m/z* coverage, and adduct relevance. This ensured that the final .msp files included only the most informative and non-redundant fragmentation patterns for each compound. Spectral similarity between experimental MS/MS and in silico predictions was assessed using a cosine similarity metric, calculated via vector normalization of *m/z* and intensity pairs:

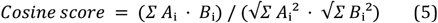

where *A*_i_ and *B*_i_ represent the intensities of the *i*-th matched fragment ion in the experimental and predicted spectra, respectively. Only matches exceeding a cosine score threshold of ≥ 0.7 were retained for Level 2 annotation. In-house Python scripts also evaluated dot product and reverse dot product scores to rank adduct-specific spectra, ensuring selection of the most representative fragmentation pattern per metabolite. Notable ionization trends were observed across compound classes: nucleotides, phosphorylated metabolites, and CoA derivatives consistently showed stronger signals in negative mode, while amines, amino acids, and small polyamines preferentially ionized in positive mode. This dual-mode acquisition strategy ensures complementary coverage, expanding the range of detectable metabolites while reducing redundancy between modes. Finalized .msp files for both ionization modes are provided in the Supplementary Material and include curated metadata for each compound: precursor m/z, retention time, collision energy, polarity, HMDB ontology, SMILES, and InChI codes. These resources are ready-to-use in untargeted metabolomics workflows, spectral matching pipelines, and method development applications.

### Method Performance

Method validation was conducted in accordance with FDA Bio-analytical Method Validation guidelines, assessing linearity, limits of detection and quantification (LoD/LoQ), precision, accuracy, recovery, process efficiency and matrix effects across five biological matrices: urine, plasma, tissues, cultured cells, and patient-derived colorectal organoids. A U-^13^C, U-^15^N-labeled Amino Acid Mixture was used as a universal internal standard across all matrices, ensuring matrix-matched normalization and controlling for extraction variability. Linearity was evaluated using 5-point calibration curves ranging from 500 pg/mL to 1 μg/mL, covering the expected physiological range of most endogenous metabolites. Quantification was based on MS^1^ signal intensities, as MS^2^ detection exhibited partial saturation at higher concentrations due to DDA acquisition limitations. According to FDA criteria, ≥85% of calibration points were required to fall within 85–115% of nominal values, which was achieved for all compounds across all matrices (**Table S7-S11**). Limits of detection (LoD) and quantification (LoQ) were calculated using signal-to-noise ratios of 3 and 10, respectively. LoD and LoQ distributions demonstrated matrix-dependent trends, with the highest detection limits observed in tissue homogenates, followed by plasma, organoids, urine, and cells, which consistently produced the lowest background and highest sensitivity. These distributions are illustrated in **Figure 2**.

**Figure 2:**
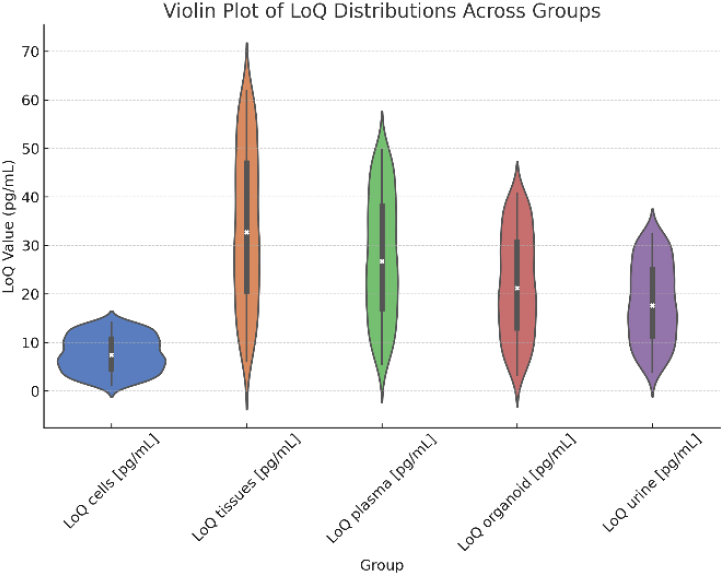
LoQ distribution comparison between 5 different types of biological matrix. Values in pg/mL.

Matrix effects were assessed by comparing post-extraction spiked standards to neat solutions and data is provided in **Table S2-S6**. The most pronounced signal suppression was observed in tissues and plasma, particularly for nucleotides and polar phosphorylated compounds. Organoids and urine exhibited intermediate matrix effects, while cultured cells displayed minimal ion suppression, reflecting their lower matrix complexity. Precision, evaluated across intra- and inter-day replicate injections (n = 5 per QC level), remained below 15% relative standard deviation (RSD) for all compounds (**Table S12-S21**). Inter- and intra-day accuracy values across low, medium, and high concentrations ranged between 85–115%, meeting FDA bioanalytical criteria for all matrices and ionization modes **(Table S22-S31**). In addition to baseline resolution of critical isomeric metabolite pairs such as citrate/isocitrate, fumarate/malate, isoleucine/leucine, and ADMA/SDMA, the optimized HILIC-MS workflow provided excellent separation of key sugar phosphates from central carbon metabolism. Specifically, hexose monophosphates, including glucose-1-phosphate (G1P), glucose-6-phosphate (G6P), fructose-6-phosphate (F6P), galactose-1-phosphate (Gal1P), and galactose-6-phosphate (Gal6P)—were successfully resolved (**Figure 3**). Despite their structural similarity and identical precursor ions, each metabolite eluted distinctly, enabling unbiased quantification across both glycolytic and galactose metabolism pathways. Furthermore, the pentose phosphate pathway intermediates ribose-5-phosphate (Ri5P), xylulose-5-phosphate (X5P), ribulose-1-phosphate (Ru1P), and ribulose-5-phosphate (Ru5P) were separated, supporting confident annotation of pentose phosphate pathway dynamics (**Figure 3**). Additional glycolytic intermediates, including glyceraldehyde-3-phosphate (GA3P) and dihydroxyacetone phosphate (DHAP), were also well resolved (**Figure 3**), eliminating ambiguity in the interpretation of lower glycolysis flux. Importantly, 2-phosphoglycerate (2-PG) and 3-phosphoglycerate (3-PG) were baseline-separated (**Figure 1, Figure 3**), overcoming one of the common chromatographic challenges in untargeted metabolomics. The successful separation of this cluster of sugar phosphate isomers underscores the method’s robustness in handling polar and structurally similar metabolites from central carbon metabolism. These findings are particularly relevant for biological models such as tumor-derived organoids, where the precise resolution of glycolytic and pentose phosphate intermediates is crucial for the accurate interpretation of metabolic pathway fluxes. The sample preparation protocol incorporated thiol derivatization, which effectively stabilized redox-sensitive metabolites and minimized oxidative degradation during processing and analysis. Together, these results demonstrate the method’s robustness, reproducibility, and broad applicability for quantitative metabolomics across a range of complex biological matrices.

**Figure 3.**
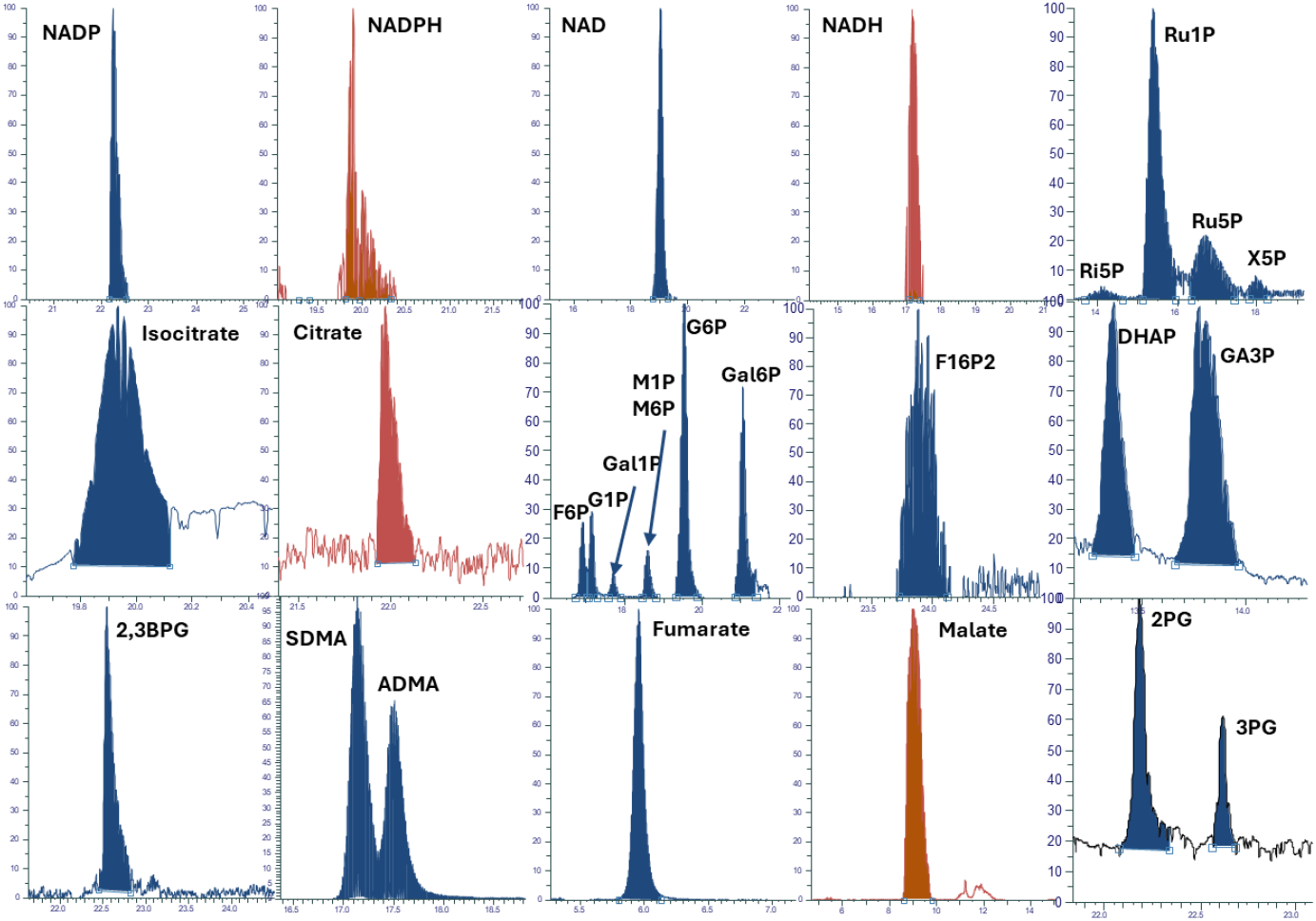
Chromatographic performance of chosen critical metabolites detected in human colorectal organoid samples.

### Application

Post method validation, the current untargeted metabolomic workflow was applied to patient-derived colorectal organoids. Two groups of organoids were analyzed for further in-depth metabolic profiling – healthy and primary tumour organoids. Statistical analysis with the PCA (**Figure S1**) showed clear grouping of the individual samples. Volcano plots were generated, illustrating the two-sided comparison between healthy and primary tumour organoids (**Figure 4**) in terms of up- and downregulated metabolites, based on p-values and log2 fold changes. The resulting p-values and log2 fold changes from the two-sided T-Test analysis are provided in **Table S1**. Based on these results, boxplots of the significantly up- and downregulated metabolites were generated and presented in **Figure S2**.

**Figure 4:**
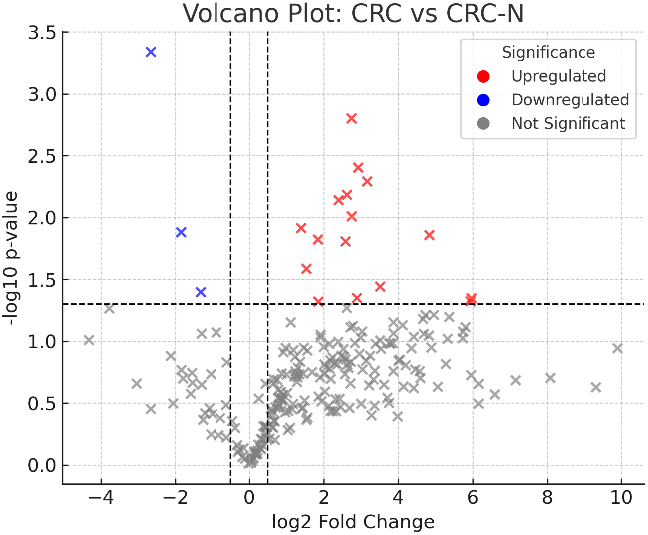
Volcano plot comparison between CRC organoid and CRC-N (healthy colorectal organoid). Significant up/downregulated metabolites marked in red/blue (p value < 0.05 and log2FC > ±1.0).

Metabolomic profiling between colorectal cancer organoids (CRC) and healthy colon organoids (CRC-N) revealed significant remodeling of central carbon and nucleotide metabolism, consistent with tumor-specific metabolic reprogramming. Key TCA cycle and glycolytic intermediates were significantly elevated in CRC organoids. Citrate, a key metabolite in the TCA cycle, was upregulated, indicating enhanced mitochondrial activity. This suggests active oxidative metabolism, not only for ATP generation but also for supplying biosynthetic precursors (e.g., citrate export for fatty acid synthesis). 2-phosphoglycerate, a glycolytic intermediate upstream of phosphoenolpyruvate, was also increased, suggesting an augmented glycolytic flux. The presence of elevated 3-hydroxybutyrate reflects increased β-oxidation of fatty acids, potentially serving to fuel the TCA cycle or acting as a stress-adaptive energy source. Elevated levels of 2-hydroxyglutarate may reflect aberrant isocitrate dehydrogenase (IDH) activity, such as mutations in IDH1 or IDH2, or indicate broader mitochondrial metabolic dysregulation. Several nucleosides were significantly downregulated in CRC organoids. Adenosine and inosine levels were both suppressed, suggesting disruption in purine metabolism, particularly the nucleotide salvage pathway. These pathways are essential for maintaining nucleotide pools in rapidly dividing cells. However, tumor cells often rely on de novo nucleotide biosynthesis to meet the demands of high proliferation rates. The observed drop in salvage nucleosides may thus reflect a metabolic switch toward more energy-intensive biosynthetic routes, or alternatively, altered extracellular nucleotide turnover and purinergic signaling, both of which play roles in cell cycle progression, inflammation, and immune evasion. Additionally, creatinine,, a byproduct of creatine metabolism involved in cellular energy buffering via the phosphocreatine system, was decreased, pointing to altered phosphagen dynamics. Together, these alterations highlight a CRC-specific metabolic signature characterized by enhanced glycolysis, increased TCA activity, intensified lipid oxidation, and reprogrammed nucleotide metabolism, all supporting biosynthetic demands and rapid cell division.

## Conclusion

We present UCL-MetIsoLib, a validated, high-resolution tandem mass spectrometry (HRMS/MS) library specifically designed for isomer-resolved metabolomics using dual-mode HILIC separation. The method employs a bioinert UHPLC system with an Acquity Premier BEH Amide column and has been analytically validated across five biologically relevant matrices—urine, plasma, tissues, cultured cells, and patient-derived organoids—in accordance with FDA bioanalytical guidelines. The library comprises 334 meticulously curated metabolites, with MSI Level 1 and Level 2 annotations, supported by 245 authentic standards and in silico spectral verification. All spectra were manually curated and processed using custom in-house Python scripts to remove redundant adducts, eliminate noise, and generate high-quality .msp files for both ionization modes. Notably, the method achieves robust isomer resolution through polarity-specific fragmentation and retention time filtering, features often lacking in public spectral repositories. To the best of our knowledge, this is the only methodological platform that combines this level of comprehensiveness and selectivity within a HILIC-based metabolomics framework.

## ASSOCIATED CONTENT

### Supporting Information

The Supporting Information is available free of charge on the ACS Publications website. Information and data on chromatographic parameters such as ME, PE, ER. Assay validation data values i.e. precision, accuracy, calibration, LoD, LoQ. Annotated peak list of metabolites in organoid study. Fold change and statistical values for significant metabolites between both organoid models. Figures related to the multivariate statistical analysis of both organoid models. File type docx. All curated MS/MS spectral data and associated metadata are available as Supplementary Material in .msp format. Raw vendor files were used to generate composite consensus spectra and are not representative of individual sample injections.

## AUTHOR INFORMATION

## Insert Table of Contents artwork here

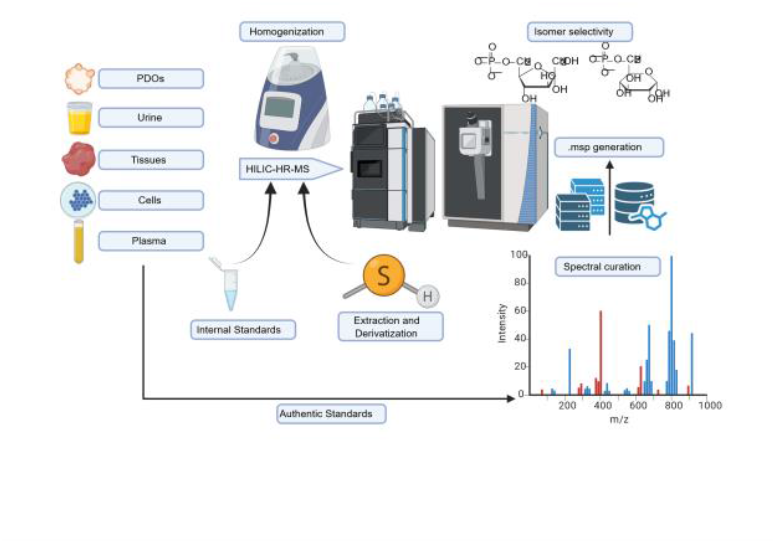

