## Supplementary Info for "UCL-MetIsoLib: A Public High-Resolution Tandem Mass Spectrometry Library for HILIC-Based Isomer-Resolved Profiling of Glycolysis, Central Carbon Metabolism, and Beyond in Urine, Plasma, Tissues, Cells, and Patient-Derived Organoids"

**Dr. Kristian Serafimov**

IREC Metabolomics Core

Pole of Pharmacology and Therapeutics

Institut de Recherche Expérimentale et Clinique (IREC)

Université catholique de Louvain

1200 Brussels, Belgium

E-mail: [kristian.serafimov](mailto:kristian.serafimov)@uclouvain.be

**Content**

**Table S1** | Full Metabolite Annotation Table with Log2 Fold Changes, Raw and FDR-Adjusted p-Values from Differential Analysis Between Healthy and Tumor-Derived Colorectal Organoids. Includes retention times, molecular formulas, precursor m/z values, adduct types, MS/MS fragment ions based for identification and ontology classifications where available.	5

[**Figure S2** | Boxplots showing normalized intensities [cps] of the significantly regulated metabolites (p < 0.05, |log₂ fold change| > 1.0) between healthy and tumor-derived colorectal organoids (n = 3). Each plot displays relative abundance based on normalized signal intensities.	166](#_Toc198131377)

**Table S1**| Full Metabolite Annotation Table with Log2 Fold Changes, p-Values from Differential Analysis Between Healthy and Tumor-Derived Colorectal Organoids. Includes retention times, molecular formulas, precursor m/z values, adduct types, MS/MS fragment ions based for identification and ontology classifications where available.

| **Metabolite** | **Sum Formula** | | **Precursor m/z** | | **Adduct** | **tR [min]** | **Ontology** | **MSI ID Level** | **Log2 FC** | **p value** |
| --- | --- | --- | --- | --- | --- | --- | --- | --- | --- | --- |
| 1-Methylguanine | | C6H7N5O | | 166.0723 | [M+H]+ | 4.23 | HMDB0003282 | 2 | 1.49 | 3.02E-01 |
| 1-Methylnicotinamide | | C7H9N2O | | 137.0709 | [M+H]+ | 6.4 | HMDB0000699 | 2 | -3.77 | 5.44E-02 |
| 2-Hydroxyisocaproic acid | | C6H12O3 | | 131.0714 | [M-H]- | 0.6 | HMDB0000665 | 2 | 0.43 | 5.56E-01 |
| 2-Phenylglycine | | C8H9NO2 | | 152.0706 | [M+H]+ | 11.77 | HMDB0002210 | 2 | -0.89 | 8.50E-02 |
| 4-amino-butanal | | C4H9NO | | 88.0757 | [M+H]+ | 4.95 | HMDB0001080 | 2 | 0.98 | 1.14E-01 |
| 4-aspartyl-phosphate | | C4H8NO7P | | 211.9966 | [M-H]- | 15.02 | HMDB0012250 | 2 | -1.08 | 3.49E-01 |
| 4-Hydroxyphenylacetylglycine | | C10H11NO4 | | 210.0761 | [M+H]+ | 12.86 | HMDB0000735 | 2 | 0.13 | 8.83E-01 |
| 4-Oxoglutaramate | | C5H7NO4 | | 146.0448 | [M-H]- | 7.62 |  | 2 | 0.97 | 3.76E-01 |
| 5-Hydroxyindolepyruvate | | C11H9NO4 | | 220.0604 | [M-H]- | 11.93 |  | 2 | 2.24 | 3.67E-01 |
| 5-Hydroxylysine | | C6H14N2O3 | | 163.1077 | [M+H]+ | 20.69 | HMDB0000450 | 2 | -1.84 | 1.71E-01 |
| 5-Phosphoribosylamine | | C5H12NO7P | | 228.0279 | [M-H]- | 15.69 | HMDB0001128 | 2 | 4.36 | 1.14E-01 |
| Biopterin | | C9H11N5O3 | | 238.0395 | [M+H]+ | 6.61 | HMDB0000633 | 2 | 0.36 | 6.26E-01 |
| CDP-choline | | C14H26N4O11P2 | | 487.1001 | [M-H]- | 15.32 | HMDB0001413 | 2 | 4.82 | 9.02E-02 |
| Cysteamine | | C8H14N2O2S | | 203.0849 | [M+H]+ | 4.42 | HMDB0002991 | 2 | -0.46 | 4.44E-01 |
| Cytidine 3',5'-cyclic monophosphate | | C9H12N3O7P | | 304.034 | [M-H]- | 2.95 | HMDB0000052 | 2 | 0.05 | 9.62E-01 |
| dADP | | C10H15N5O9P2 | | 410.0272 | [M-H]- | 19.09 | HMDB0001508 | 2 | 0.66 | 2.03E-01 |
| Diethanolamine | | C4H11NO2 | | 106.0863 | [M+H]+ | 8.46 | HMDB0062263 | 2 | 0.44 | 2.21E-01 |
| Dihydrobiopterin | | C9H13N5O3 | | 240.1091 | [M+H]+ | 6.29 | HMDB0000038 | 2 | -0.26 | 7.53E-01 |
| Dihydroneopterin triphosphate | | C9H16N5O13P3 | | 493.9885 | [M-H]- | 14.12 | HMDB0000980 | 2 | -1.54 | 2.17E-01 |
| Ectoine | | C6H10N2O2 | | 143.0815 | [M+H]+ | 10.3 | HMDB0240650 | 2 | 0.3 | 7.33E-01 |
| Guanidinoacetic acid | | C3H7N3O2 | | 116.0353 | [M-H]- | 6.15 | HMDB0000128 | 2 | 0.26 | 2.92E-01 |
| Guanidinobutanoic acid | | C5H11N3O2 | | 146.0924 | [M+H]+ | 10.6 | HMDB0003464 | 2 | -2.12 | 1.32E-01 |
| Hercynine | | C9H15N3O2 | | 198.1237 | [M+H]+ | 13.77 | HMDB0029422 | 2 | 3.29 | 3.98E-01 |
| Homoarginine | | C7H16N4O2 | | 189.1346 | [M+H]+ | 17.33 | HMDB0000670 | 2 | 3.4 | 1.05E-01 |
| Hydantoin-5-propionate | | C6H8N2O4 | | 173.0557 | [M+H]+ | 17.3 | HMDB0001212 | 2 | 5.81 | 7.70E-02 |
| Indole-3-acetamide | | C10H10N2O | | 175.0864 | [M+H]+ | 2.28 | HMDB0029739 | 2 | 0.79 | 2.47E-01 |
| N1-Acetylspermidine | | C9H21N3O | | 188.1757 | [M+H]+ | 17.37 | HMDB0001276 | 2 | 5.37 | 6.37E-02 |
| N1-Acetylspermine | | C12H28N4O | | 245.2336 | [M+H]+ | 21.09 | HMDB0001186 | 2 | 9.32 | 2.36E-01 |
| N8-Acetylspermidine | | C9H21N3O | | 188.1757 | [M+H]+ | 17.88 | HMDB0002189 | 2 | 6.59 | 2.70E-01 |
| N-Acetyl-Aspartyl-Glutamate | | C11H16N2O8 | | 305.0979 | [M+H]+ | 13.51 | HMDB0000279 | 2 | 3.87 | 1.75E-01 |
| N-Acetyl-Aspartyl-Glutamyl-Glutamate | | C16H23N3O11 | | 434.1405 | [M-H]- | 13.52 |  | 2 | 0.78 | 2.89E-01 |
| N-Acetyl-Putrescine | | C6H14N2O | | 131.1179 | [M+H]+ | 7.92 | HMDB0002064 | 2 | 3.14 | 1.44E-01 |
| N-Acyl-Homoserine-lactone | | C6H9NO3 | | 144.0655 | [M-H]- | 6.3 |  | 2 | 0.22 | 7.66E-01 |
| N-Carbamoyl-Proline | | C6H10N2O3 | | 159.0764 | [M+H]+ | 14.43 | HMDB0000226 | 2 | 0.88 | 3.02E-01 |
| N-Carbamoyl-Sarcosine | | C4H8N2O3 | | 133.0608 | [M+H]+ | 15.78 | HMDB0012265 | 2 | 4.42 | 9.22E-02 |
| N-Methyl-Aspartic Acid | | C5H9NO4 | | 148.0604 | [M+H]+ | 14.02 | HMDB0001161 | 2 | 3.86 | 1.04E-01 |
| Orotidine 5'-monophosphate | | C10H13N2O11P | | 367.0184 | [M-H]- | 24.83 | HMDB0000218 | 2 | 4.15 | 2.33E-01 |
| Phosphodimethylethanolamine | | C4H12NO4P | | 168.0431 | [M-H]- | 20.01 | HMDB0060244 | 2 | -0.18 | 8.65E-01 |
| Phosphorylcholine | | C5H14NO4P | | 184.0733 | [M+H]+ | 15.8 | HMDB0001565 | 2 | 2.39 | 1.01E-01 |
| Putrescine | | C4H12N2 | | 89.1073 | [M+H]+ | 19.39 | HMDB0001414 | 2 | 3.62 | 2.25E-01 |
| S-Adenosylmethioninamine | | C14H23N6O3S | | 355.1552 | [M+H]+ | 21.22 | HMDB0000988 | 2 | 2.34 | 3.69E-01 |
| Selenohomocysteine | | C4H9NO2Se | | 183.9871 | [M+H]+ | 7.39 | HMDB0004119 | 2 | 1.56 | 1.19E-01 |
| S-Methyl-Thioinosine | | C11H14N4O4S | | 299.0809 | [M+H]+ | 17.36 |  | 2 | 4.73 | 6.17E-02 |
| Spermidine | | C7H19N3 | | 146.1652 | [M+H]+ | 18.78 | HMDB0001257 | 2 | 5.06 | 2.34E-01 |
| Spermine | | C10H26N4 | | 203.223 | [M+H]+ | 22.7 | HMDB0001256 | 2 | 2.69 | 3.47E-01 |
| Tetrahydroneopterin | | C10H16N2O4S | | 258.1197 | [M+H]+ | 9.15 | HMDB0000942 | 2 | 0.58 | 3.52E-01 |
| Trigonelleine | | C4H8N2O3 | | 138.055 | [M+H]+ | 13.86 | HMDB0001123 | 2 | 1.87 | 2.81E-01 |
| Trimethyllysine | | C9H21N2O2 | | 189.1597 | [M+H]+ | 17.87 | HMDB0001325 | 2 | 3 | 9.35E-02 |
| Tyramine | | C8H11NO | | 138.0913 | [M+H]+ | 2.91 | HMDB0000306 | 2 | -0.6 | 1.48E-01 |
| 1-Methylhistidine | | C7H11N3O2 | | 170.0924 | [M+H]+ | 14.79 | HMDB0000001 | 1 | 2.72 | 7.72E-02 |
| 2-Aminoadipic Acid | | C6H11NO4 | | 162.0761 | [M+H]+ | 7.53 | HMDB0000510 | 1 | 0.08 | 9.26E-01 |
| 2-Deoxyadenosine | | C10H13N5O3 | | 252.1091 | [M+H]+ | 3.24 | HMDB0000101 | 1 | -0.62 | 3.29E-01 |
| 2'-Deoxyadenosine 5'-monophosphate | | C10H14N5O6P | | 330.0609 | [M-H]- | 12 | HMDB0000905 | 1 | 4.84 | 1.39E-02 |
| 2'-deoxyadenosine 5'-triphosphate | | C10H16N5O12P3 | | 489.9936 | [M-H]- | 22.94 | HMDB0001532 | 1 | 0.05 | 9.42E-01 |
| 2'-Deoxycytidine-5'-monophosphate | | C9H14N3O7P | | 306.0497 | [M-H]- | 14.87 | HMDB0001202 | 1 | 7.15 | 2.07E-01 |
| 2-Deoxyguanosine | | C10H13N5O4 | | 268.1038 | [M+H]+ | 6.43 | HMDB0000085 | 1 | 1.19 | 1.18E-01 |
| 2-hydroxybutyric acid | | C4H8O3 | | 103.0401 | [M-H]- | 1.4 | HMDB0000008 | 1 | 0.45 | 4.99E-01 |
| 2-hydroxyglutaric acid | | C5H8O5 | | 147.0299 | [M-H]- | 9.73 | HMDB0059655 | 1 | 2.62 | 6.55E-03 |
| 2-ketobutyric acid | | C4H6O3 | | 101.0244 | [M-H]- | 0.61 | HMDB0000005 | 1 | 0.66 | 2.19E-01 |
| 2-oxoadipic acid | | C6H8O5 | | 159.0299 | [M-H]- | 4.96 | HMDB0000225 | 1 | 0.22 | 8.16E-01 |
| 2-phosphoglycerate | | C3H7O7P | | 184.9857 | [M-H]- | 20.42 | HMDB0000362 | 1 | 3.52 | 3.62E-02 |
| 3-hydroxy-3-methyl-2-oxobutanoate | | C5H8O4 | | 131.035 | [M-H]- | 0.76 | HMDB0006833 | 1 | 0.18 | 8.22E-01 |
| 3-hydroxybutyric acid | | C4H8O3 | | 103.0401 | [M-H]- | 1.7 | HMDB0000011 | 1 | 2.75 | 1.58E-03 |
| 3-Methylhistidine | | C7H11N3O2 | | 170.0924 | [M+H]+ | 15.9 | HMDB0000479 | 1 | 3.79 | 1.05E-01 |
| 3-Phosphoglyceraldehyde | | C3H7O6P | | 168.9907 | [M-H]- | 15.28 | HMDB0001473 | 1 | 3.99 | 4.06E-01 |
| 3-phosphoglycerate | | C3H7O7P | | 21.45 | [M-H]- | 21.45 | HMDB0000807 | 1 | 2.63 | 1.55E-01 |
| 4-amino-butyric acid | | C4H9NO2 | | 104.0706 | [M+H]+ | 11.03 | HMDB0000112 | 1 | 1.94 | 3.07E-01 |
| 4-Hydroxybenzoic acid | | C7H6O3 | | 137.0244 | [M-H]- | 2.54 | HMDB0000500 | 1 | 1.1 | 5.01E-01 |
| 4-methyl-2-oxopentanoate | | C6H10O3 | | 129.0557 | [M-H]- | 0.57 | HMDB0000695 | 1 | -0.11 | 8.91E-01 |
| 5-Hydroxy-Tryptophan | | C11H12N2O3 | | 221.0921 | [M+H]+ | 8.94 | HMDB0000472 | 1 | -0.38 | 5.04E-01 |
| 6-phosphogluconate | | C6H13O10P | | 275.0174 | [M-H]- | 21.81 | HMDB0001316 | 1 | 0.75 | 2.05E-01 |
| Acetylcarnitine | | C9H18NO4 | | 204.123 | [M+H]+ | 8.97 | HMDB0240773 | 1 | 0.88 | 1.83E-01 |
| Aconitic Acid | | C6H6O6 | | 173.0092 | [M-H]- | 13.52 | HMDB0247961 | 1 | 3.24 | 1.31E-01 |
| Adenine | | C5H5N5 | | 136.0618 | [M-H]- | 3.52 | HMDB0000034 | 1 | 0.32 | 6.18E-01 |
| Adenosine | | C10H13N5O4 | | 268.104 | [M+H]+ | 3.84 | HMDB0000050 | 1 | -1.84 | 1.32E-02 |
| Adenosine 3',5'-cyclic monophosphate | | C10H12N5O6P | | 328.0452 | [M-H]- | 1.53 | HMDB0000058 | 1 | 1.22 | 1.93E-01 |
| Adenosine Monophosphate | | C10H14N5O7P | | 346.0558 | [M-H]- | 13.14 | HMDB0000045 | 1 | 0.55 | 6.14E-01 |
| Adenosine-5'-diphosphate | | C10H15N5O10P2 | | 426.0221 | [M-H]- | 21.93 | HMDB0001341 | 1 | 2.59 | 1.56E-02 |
| Adenosine-5'-triphosphate | | C10H16N5O13P3 | | 505.9885 | [M-H]- | 24.26 | HMDB0000538 | 1 | 1.54 | 2.60E-02 |
| Adenylosuccinate | | C14H18N5O11P | | 462.0668 | [M-H]- | 22.8 | HMDB0000536 | 1 | 5.32 | 9.57E-02 |
| Alanine | | C3H7NO2 | | 90.055 | [M+H]+ | 11.69 | HMDB0000161 | 1 | 2.73 | 1.47E-01 |
| alpha-ketoglutaric acid | | C5H6O5 | | 145.0142 | [M-H]- | 5.53 | HMDB0000208 | 1 | 3.22 | 2.28E-01 |
| Anserine | | C10H16N4O3 | | 241.1295 | [M+H]+ | 16.26 | HMDB0000194 | 1 | -1.29 | 2.25E-01 |
| Arginine | | C6H14N4O2 | | 175.119 | [M+H]+ | 18.77 | HMDB0000517 | 1 | 2.72 | 2.20E-01 |
| Argininosuccinic Acid | | C10H18N4O6 | | 289.1154 | [M-H]- | 17.83 | HMDB0000052 | 1 | 6.16 | 2.22E-01 |
| Asparagine | | C4H8N2O3 | | 133.0605 | [M+H]+ | 13.89 | HMDB0000168 | 1 | 2.55 | 1.35E-01 |
| Aspartic acid | | C4H7NO4 | | 134.0448 | [M+H]+ | 15.89 | HMDB0000191 | 1 | 4.69 | 8.73E-02 |
| Asymmetric dimethylarginine | | C8H18N4O2 | | 203.1503 | [M-H]- | 16.31 | HMDB0001539 | 1 | 2.07 | 1.63E-01 |
| Azelaic acid | | C9H16O4 | | 187.0976 | [M-H]- | 3.9 | HMDB0000784 | 1 | 0.62 | 4.92E-01 |
| beta-Alanine | | C3H7NO2 | | 90.055 | [M+H]+ | 12.23 | HMDB0000056 | 1 | 2.84 | 1.04E-01 |
| Betaine | | C5H12NO2 | | 118.0863 | [M+H]+ | 8.21 | HMDB0000043 | 1 | -0.01 | 9.76E-01 |
| Biotin | | C10H16N2O3S | | 245.0954 | [M+H]+ | 2.34 | HMDB0000030 | 1 | 1.35 | 1.85E-01 |
| Carnitine | | C7H15NO3 | | 162.1125 | [M+H]+ | 11.07 | HMDB0000062 | 1 | 2.36 | 1.54E-01 |
| Carnosine | | C9H14N4O3 | | 227.1139 | [M+H]+ | 19.45 | HMDB0000033 | 1 | 3.04 | 1.69E-01 |
| cGMP | | C10H12N5O7P | | 344.0402 | [M-H]- | 3.31 | HMDB0001314 | 1 | 2.45 | 3.40E-01 |
| Choline | | C5H13NO | | 104.107 | [M+H]+ | 5.37 | HMDB0000097 | 1 | 1.3 | 1.64E-01 |
| Cinnamic acid | | C9H8O2 | | 147.0452 | [M-H]- | 1.59 | HMDB0000930 | 1 | 2.18 | 3.67E-01 |
| Citraconic acid | | C5H6O4 | | 129.0193 | [M-H]- | 1.68 | HMDB0000634 | 1 | -1.2 | 3.82E-01 |
| Citramalic acid | | C5H8O5 | | 147.0299 | [M-H]- | 22.26 | HMDB0000426 | 1 | 1.86 | 4.80E-02 |
| Citric Acid | | C6H8O7 | | 191.0197 | [M-H]- | 22.13 | HMDB0000094 | 1 | 2.4 | 7.24E-03 |
| Citrulline | | C6H13N3O3 | | 176.103 | [M+H]+ | 14.46 | HMDB0000904 | 1 | 1.04 | 5.24E-01 |
| Creatine | | C4H9N3O2 | | 132.0768 | [M+H]+ | 11.98 | HMDB0000064 | 1 | 0.69 | 6.30E-01 |
| Creatinine | | C4H7N3O | | 114.0662 | [M+H]+ | 4.45 | HMDB0000562 | 1 | -2.66 | 4.60E-04 |
| Cystathionine | | C7H14N2O4S | | 223.0747 | [M+H]+ | 19.64 | HMDB0000099 | 1 | 8.08 | 1.98E-01 |
| Cysteine | | C9H14N2O4S | | 247.0747 | [M+H]+ | 7.63 | HMDB0000574 | 1 | -0.15 | 7.34E-01 |
| Cysteinylglycine | | C11H17N3O5S | | 304.0962 | [M+H]+ | 8.58 | HMDB0000078 | 1 | -1.79 | 2.00E-01 |
| Cystine | | C6H12N2O4S2 | | 241.0311 | [M+H]+ | 19.74 | HMDB0000192 | 1 | 2.57 | 1.85E-01 |
| Cytidine | | C9H13N3O5 | | 244.0928 | [M+H]+ | 7.12 | HMDB0000089 | 1 | 1.25 | 1.86E-01 |
| Cytidine 5'-diphosphate | | C9H15N3O11P2 | | 402.0109 | [M-H]- | 22.87 | HMDB0001546 | 1 | 1.92 | 9.35E-02 |
| Cytidine 5'-triphosphate | | C9H16N3O14P3 | | 481.9772 | [M-H]- | 23.8 | HMDB0000082 | 1 | -3.04 | 2.20E-01 |
| Cytidine monophosphate | | C9H14N3O8P | | 322.0446 | [M-H]- | 16.68 | HMDB0000095 | 1 | 9.91 | 1.14E-01 |
| Cytosine | | C4H5N3O | | 112.0505 | [M+H]+ | 7.42 | HMDB0000630 | 1 | 1.25 | 1.76E-01 |
| Dihydroxyacetonephosphate | | C3H7O6P | | 168.9907 | [M-H]- | 18.28 | HMDB0247961 | 1 | 4.89 | 1.13E-01 |
| Dimethylglycine | | C4H9NO2 | | 104.0706 | [M+H]+ | 10.3 | HMDB0000092 | 1 | 0.48 | 4.97E-01 |
| Erythrose-4-phosphate | | C4H9O7P | | 199.0013 | [M-H]- | 5.89 | HMDB0001321 | 1 | 1.12 | 7.05E-02 |
| FAD | | C27H33N9O15P2 | | 789.1499 | [M-H]- | 10.46 | HMDB0001248 | 1 | 0.85 | 2.58E-01 |
| Folic acid | | C19H19N7O6 | | 442.147 | [M+H]+ | 12.77 | HMDB0000121 | 1 | 0.91 | 2.72E-01 |
| Fructose-6-phosphate | | C6H13O9P | | 259.0224 | [M-H]- | 16.34 | HMDB0000124 | 1 | 2.24 | 1.36E-01 |
| Fumaric Acid | | C4H4O4 | | 115.0037 | [M-H]- | 7.77 | HMDB0000134 | 1 | 2.89 | 4.48E-02 |
| Galactosamine | | C6H13NO5 | | 180.0866 | [M+H]+ | 15.47 | HMDB0001015 | 1 | 0.18 | 8.19E-01 |
| Galactose-1-phosphate | | C6H13O9P | | 259.0224 | [M-H]- | 17.44 | HMDB0000645 | 1 | -0.83 | 5.76E-01 |
| Galactose-6-Phosphate | | C6H13O9P | | 259.0224 | [M-H]- | 20.89 |  | 1 | 0.8 | 3.25E-01 |
| gamma-Glutamylcysteine | | C14H21N3O7S | | 376.1173 | [M+H]+ | 13.42 | HMDB0001049 | 1 | 1.45 | 1.83E-01 |
| Glucosamine | | C6H13NO5 | | 180.0866 | [M+H]+ | 14.71 | HMDB0001514 | 1 | 0.67 | 4.38E-01 |
| Glucosamine-6-Phosphate | | C6H14NO8P | | 258.0384 | [M-H]- | 10.4 | HMDB0001254 | 1 | 4.57 | 1.75E-01 |
| Glucose | | C6H12O6 | | 179.0561 | [M-H]- | 2.25 | HMDB0304632 | 1 | -2.66 | 3.53E-01 |
| Glucose-1-phosphate | | C6H13O9P | | 259.0224 | [M-H]- | 16.75 | HMDB0001586 | 1 | -0.3 | 7.87E-01 |
| Glucose-6-Phosphate | | C6H13O9P | | 259.0224 | [M-H]- | 19.2 | HMDB0001401 | 1 | 0.45 | 6.05E-01 |
| Glucuronic acid | | C6H10O7 | | 193.0354 | [M-H]- | 4.88 | HMDB0000127 | 1 | 4.6 | 1.99E-01 |
| Glutamic acid | | C5H9NO4 | | 148.0604 | [M+H]+ | 14.96 | HMDB0000148 | 1 | 3.54 | 1.15E-01 |
| Glutamine | | C5H10N2O3 | | 147.0764 | [M+H]+ | 13.46 | HMDB0000641 | 1 | 4.04 | 1.41E-01 |
| Glutathione | | C14H21N3O7S | | 433.1388 | [M+H]+ | 13.13 | HMDB0001049 | 1 | 3.51 | 9.86E-02 |
| Glutathionylspermine | | C20H41N7O5S | | 492.2963 | [M-H]- | 20.87 |  | 1 | 2.19 | 2.02E-01 |
| Glycerophosphocholine | | C8H20NO6P | | 258.1101 | [M+H]+ | 14.02 | HMDB0000086 | 1 | 3.87 | 7.05E-02 |
| Glycine | | C2H5NO2 | | 76.0393 | [M+H]+ | 14.02 | HMDB0000123 | 1 | 2.96 | 1.27E-01 |
| Glycolic acid | | C2H4O3 | | 75.0088 | [M-H]- | 2.35 | HMDB0000115 | 1 | 0.64 | 5.03E-01 |
| Guanine | | C5H5N5O | | 152.0567 | [M+H]+ | 6.22 | HMDB0000132 | 1 | 2 | 1.11E-01 |
| Guanosine | | C10H13N5O5 | | 284.0989 | [M+H]+ | 8.43 | HMDB0000133 | 1 | 0.08 | 8.97E-01 |
| Guanosine 5'-diphosphate | | C10H15N5O11P2 | | 442.0171 | [M-H]- | 24.35 | HMDB0001201 | 1 | 1.23 | 1.34E-01 |
| Guanosine-5'-monophosphate | | C10H14N5O8P | | 362.0507 | [M-H]- | 18.2 | HMDB0001397 | 1 | 5.28 | 1.53E-01 |
| Hippuric acid | | C9H9NO3 | | 178.051 | [M-H]- | 0.9 | HMDB0000714 | 1 | 0.34 | 6.87E-01 |
| Histamine | | C5H9N3 | | 112.0869 | [M+H]+ | 18.76 | HMDB0000870 | 1 | 2.15 | 1.05E-01 |
| Histidine | | C6H9N3O2 | | 156.0768 | [M+H]+ | 17.14 | HMDB0000177 | 1 | 2.48 | 1.32E-01 |
| Homoalanine | | C4H9NO2 | | 104.0706 | [M+H]+ | 10.35 | HMDB0245013 | 1 | 1.18 | 3.22E-01 |
| Homocysteine | | C10H16N2O4S | | 261.0904 | [M+H]+ | 7.57 | HMDB0000742 | 1 | 2.02 | 1.54E-01 |
| Homogentisic acid | | C8H8O4 | | 167.035 | [M-H]- | 0.64 | HMDB0000130 | 1 | 5.97 | 4.50E-02 |
| Homoleucine | | C7H15NO2 | | 146.1176 | [M+H]+ | 6.8 | HMDB0003464 | 1 | 1.92 | 8.87E-02 |
| Homovanillic acid | | C9H10O4 | | 181.0506 | [M-H]- | 0.95 | HMDB0000118 | 1 | 1.02 | 2.63E-01 |
| Hypotaurine | | C2H7NO2S | | 110.027 | [M+H]+ | 11.5 | HMDB0000965 | 1 | 1.54 | 4.33E-01 |
| Hypoxanthine | | C5H4N4O | | 137.0458 | [M+H]+ | 3.84 | HMDB0000157 | 1 | 0.15 | 7.79E-01 |
| Inosine | | C10H12N4O5 | | 269.088 | [M+H]+ | 5.69 | HMDB0000195 | 1 | -1.32 | 4.02E-02 |
| Inosine 5'-monophosphate | | C10H13N4O8P | | 347.0398 | [M-H]- | 15.96 | HMDB0000175 | 1 | 5.73 | 9.48E-02 |
| Isocitric Acid | | C6H8O7 | | 191.0197 | [M-H]- | 18.66 | HMDB0000193 | 1 | 1.37 | 1.47E-01 |
| Isoleucine | | C6H13NO2 | | 132.1019 | [M+H]+ | 8.43 | HMDB0000172 | 1 | 1.29 | 2.14E-01 |
| Lactic Acid | | C3H6O3 | | 89.0244 | [M-H]- | 1.32 | HMDB0000190 | 1 | -0.62 | 6.01E-01 |
| Leucine | | C6H13NO2 | | 132.1019 | [M+H]+ | 7.84 | HMDB0000687 | 1 | 0.89 | 2.99E-01 |
| Levulinic acid | | C5H8O3 | | 115.0401 | [M-H]- | 0.58 | HMDB0000720 | 1 | 0.45 | 4.86E-01 |
| Lysine | | C6H14N2O2 | | 147.1128 | [M+H]+ | 19.27 | HMDB0000182 | 1 | 3.76 | 1.04E-01 |
| Malic Acid | | C4H6O5 | | 133.0142 | [M-H]- | 9.92 | HMDB0000691 | 1 | 2.62 | 5.38E-02 |
| Mannose-1 with Mannose-6-Phosphate | | C6H13O9P | | 259.0224 | [M-H]- | 18.07 |  | 1 | 0.69 | 4.45E-01 |
| Mesaconic acid | | C5H6O4 | | 129.0193 | [M-H]- | 8.97 | HMDB0000749 | 1 | 2.16 | 1.67E-01 |
| Methionine | | C5H11NO2S | | 150.0583 | [M+H]+ | 9.06 | HMDB0000696 | 1 | 0.77 | 3.44E-01 |
| Methionine sulfoxide | | C5H11NO3S | | 166.0532 | [M+H]+ | 13.3 | HMDB0250776 | 1 | 3.75 | 3.16E-01 |
| Methylmalonic Acid | | C4H6O4 | | 117.0193 | [M-H]- | 6.09 | HMDB0000202 | 1 | 0.88 | 2.92E-01 |
| N-Acetyl-Galactosamine | | C6H13NO5 | | 180.0866 | [M+H]+ | 15.47 | HMDB0001015 | 1 | -1.29 | 8.69E-02 |
| N-Acetyl-Leucine | | C8H15NO3 | | 174.1125 | [M+H]+ | 6.84 | HMDB0011756 | 1 | -0.78 | 4.19E-01 |
| N-Acetyl-Methionine | | C7H13NO3S | | 192.0689 | [M+H]+ | 7.73 | HMDB0011745 | 1 | 2.92 | 3.95E-03 |
| N-Acetylneuraminic acid | | C11H19NO9 | | 310.1133 | [M+H]+ | 14.01 | HMDB0000230 | 1 | 4.96 | 6.13E-02 |
| N-Acetyl-Proline | | C5H9NO2 | | 116.0706 | [M+H]+ | 10.3 | HMDB0000162 | 1 | 1 | 1.86E-01 |
| N-Acetyl-Serine | | C5H9NO4 | | 148.0604 | [M+H]+ | 10.52 | HMDB0002931 | 1 | 2.32 | 2.94E-01 |
| N-Acetyl-Serotonin | | C12H14N2O2 | | 219.1128 | [M+H]+ | 2.72 | HMDB0001238 | 1 | 0.61 | 3.99E-01 |
| NAD | | C21H27N7O14P2 | | 664.1164 | [M+H]+ | 18.6 | HMDB0000902 | 1 | 5.74 | 8.39E-02 |
| NADH | | C21H29N7O14P2 | | 666.132 | [M+H]+ | 17.06 | HMDB0001487 | 1 | 2.29 | 1.51E-01 |
| NADP | | C21H28N7O17P3 | | 742.0682 | [M-H]- | 22.36 | HMDB0000217 | 1 | 1.3 | 1.96E-01 |
| NADPH | | C21H30N7O17P3 | | 744.0838 | [M-H]- | 21.16 | HMDB0000221 | 1 | 3.37 | 1.73E-01 |
| N-alpha-Acetyl-lysine | | C8H16N2O3 | | 189.1234 | [M+H]+ | 11.57 | HMDB0000446 | 1 | 1.43 | 3.44E-01 |
| N-alpha-Acetyl-Ornithine | | C7H14N2O3 | | 175.1077 | [M+H]+ | 12.45 | HMDB0003357 | 1 | 2.14 | 3.35E-01 |
| N-Epsilon-Acetyl-lysine | | C8H16N2O3 | | 189.1234 | [M+H]+ | 11.81 | HMDB0000206 | 1 | 3.71 | 2.87E-01 |
| Nicotinamide | | C6H6N2O | | 123.0553 | [M+H]+ | 1.72 | HMDB0001406 | 1 | 0.18 | 7.93E-01 |
| O-Phosphoethanolamine | | C2H8NO4P | | 140.0118 | [M-H]- | 18.13 | HMDB0000224 | 1 | 2.75 | 9.78E-03 |
| Ophthalmic acid | | C11H19N3O6 | | 290.1342 | [M+H]+ | 16.21 | HMDB0005765 | 1 | 4.67 | 6.62E-02 |
| Ornithine | | C5H12N2O2 | | 133.0972 | [M+H]+ | 19.59 | HMDB0000214 | 1 | 1.42 | 3.51E-01 |
| Oxaloacetate | | C4H4O5 | | 130.9986 | [M-H]- | 10.53 | HMDB0000223 | 1 | 4.46 | 1.70E-01 |
| Pantothenic acid | | C9H17NO5 | | 220.1179 | [M+H]+ | 2.89 | HMDB0000210 | 1 | -1.04 | 5.67E-01 |
| Phenylalanine | | C9H11NO2 | | 166.0863 | [M+H]+ | 7.76 | HMDB0000159 | 1 | 0.9 | 2.95E-01 |
| Phenylpyruvate | | C9H8O3 | | 163.0401 | [M-H]- | 0.6 | HMDB0000205 | 1 | 0.32 | 7.46E-01 |
| Phosphoenolpyruvate | | C3H5O6P | | 166.9751 | [M-H]- | 19.37 | HMDB0000263 | 1 | 2.53 | 1.16E-01 |
| Pipecolic acid | | C6H11NO2 | | 130.0863 | [M+H]+ | 9.94 | HMDB0000070 | 1 | 0.14 | 8.07E-01 |
| Proline | | C5H9NO2 | | 116.0706 | [M+H]+ | 10.3 | HMDB0000162 | 1 | 1.69 | 1.78E-01 |
| Pyridoxal | | C8H9NO3 | | 168.0655 | [M+H]+ | 2 | HMDB0001545 | 1 | 0.92 | 1.23E-01 |
| Pyridoxamine | | C8H12N2O2 | | 169.0972 | [M+H]+ | 2.31 | HMDB0001431 | 1 | 4.42 | 2.40E-01 |
| Pyridoxine | | C8H11NO3 | | 170.0812 | [M+H]+ | 2.95 | HMDB0000239 | 1 | 1.16 | 3.56E-01 |
| Pyroglutamic acid | | C5H7NO3 | | 130.0499 | [M-H]- | 9.55 | HMDB0000267 | 1 | 4.01 | 1.44E-01 |
| Pyrrolysine | | C12H21N3O3 | | 256.1656 | [M+H]+ | 14.93 | HMDB0002275 | 1 | 2.67 | 1.65E-01 |
| Pyruvic Acid | | C3H4O3 | | 87.0088 | [M-H]- | 0.71 | HMDB0000243 | 1 | -1.25 | 4.33E-01 |
| Riboflavin | | C17H20N4O6 | | 377.1456 | [M+H]+ | 5.11 | HMDB0000244 | 1 | -4.32 | 9.80E-02 |
| Ribose-5-Phosphate | | C5H11O8P | | 229.0119 | [M-H]- | 14.45 | HMDB0001548 | 1 | -0.32 | 6.94E-01 |
| Ribulose-1-Phosphate | | C5H11O8P | | 229.0119 | [M-H]- | 15.99 |  | 1 | 1.47 | 1.13E-01 |
| Ribulose-5-Phosphate | | C5H11O8P | | 229.0119 | [M-H]- | 16.57 | HMDB0000618 | 1 | -1 | 3.88E-01 |
| Saccharopine | | C11H20N2O6 | | 277.1394 | [M+H]+ | 19.44 | HMDB0000279 | 1 | 3.03 | 8.36E-02 |
| S-Adenosylhomocysteine | | C14H20N6O5S | | 385.1289 | [M+H]+ | 14.42 | HMDB0000939 | 1 | 3.88 | 8.66E-02 |
| S-Adenosylmethionine | | C15H23N6O5S | | 399.1445 | [M+H]+ | 20.55 | HMDB0001185 | 1 | 5.95 | 4.76E-02 |
| Sarcosine | | C3H7NO2 | | 90.055 | [M+H]+ | 10.96 | HMDB0000271 | 1 | 0.62 | 3.69E-01 |
| Sedoheptulose-7-phosphate | | C7H15O10P | | 289.033 | [M-H]- | 17.17 | HMDB0258206 | 1 | 3.34 | 2.05E-01 |
| Selenomethionine | | C5H11NO2Se | | 198.0028 | [M+H]+ | 6.73 | HMDB0003966 | 1 | 3.01 | 3.21E-01 |
| Serine | | C3H7NO3 | | 106.0499 | [M+H]+ | 13.29 | HMDB0062263 | 1 | 2.18 | 1.43E-01 |
| Succinate | | C4H6O4 | | 117.0193 | [M-H]- | 8.64 | HMDB0000254 | 1 | 0.82 | 3.37E-01 |
| Sucrose | | C12H22O11 | | 341.1089 | [M-H]- | 4.19 | HMDB0041627 | 1 | 3.37 | 3.33E-01 |
| Symmetric dimethylarginine | | C8H18N4O2 | | 203.1503 | [M+H]+ | 16.1 | HMDB0003334 | 1 | 2.59 | 1.40E-01 |
| Taurine | | C2H7NO3S | | 126.0219 | [M+H]+ | 10.73 | HMDB0000251 | 1 | 2.02 | 1.84E-01 |
| Thiamine | | C12H17N4OS | | 265.1117 | [M+H]+ | 7.7 | HMDB0000235 | 1 | 1.05 | 3.67E-01 |
| Thiamine monophosphate | | C12H18N4O4PS | | 345.078 | [M+H]+ | 18.08 | HMDB0002666 | 1 | -1.04 | 1.85E-01 |
| Threonic acid | | C4H8O5 | | 135.0299 | [M-H]- | 2.26 | HMDB0000943 | 1 | 3.17 | 5.11E-03 |
| Threonine | | C4H9NO3 | | 120.0655 | [M+H]+ | 12.25 | HMDB0000167 | 1 | 4.2 | 1.57E-01 |
| Thymidine | | C10H14N2O5 | | 243.0975 | [M+H]+ | 2.42 | HMDB0000273 | 1 | -1.55 | 1.81E-01 |
| Thymidine Monophosphate | | C10H15N2O8P | | 321.0493 | [M-H]- | 10.05 | HMDB0001202 | 1 | 1.9 | 1.96E-01 |
| Thymine | | C5H6N2O2 | | 127.0502 | [M+H]+ | 2.04 | HMDB0000262 | 1 | 2.79 | 7.52E-02 |
| Trimethylamine N-oxide | | C3H9NO | | 76.0757 | [M+H]+ | 5.84 | HMDB0000925 | 1 | 0.38 | 4.86E-01 |
| Tryptophan | | C11H12N2O2 | | 205.0972 | [M+H]+ | 8 | HMDB0000929 | 1 | 0.25 | 6.88E-01 |
| Tyrosine | | C9H11NO3 | | 182.0812 | [M+H]+ | 10.52 | HMDB0000158 | 1 | 1.56 | 4.15E-01 |
| Uracil | | C4H4N2O2 | | 113.0345 | [M+H]+ | 3.52 | HMDB0000300 | 1 | 0.83 | 2.87E-01 |
| Uridine 5'-diphosphate | | C9H14N2O12P2 | | 402.9949 | [M-H]- | 22.65 | HMDB0000295 | 1 | 1.4 | 1.22E-02 |
| Uridine Monophosphate | | C9H14N3O8P | | 323.0286 | [M-H]- | 13.75 | HMDB0000288 | 1 | 4.12 | 7.44E-02 |
| Uridine-5-Diphosphogalactose | | C15H24N2O17P2 | | 14.83 | [M-H]- | 14.83 | HMDB0000302 | 1 | 6.16 | 3.20E-01 |
| Urocanic acid | | C6H6N2O2 | | 139.0502 | [M+H]+ | 6.14 | HMDB0000301 | 1 | -2.06 | 3.19E-01 |
| UTP | | C9H15N2O15P3 | | 482.9613 | [M-H]- | 23.17 | HMDB0000285 | 1 | 1.85 |  |
| Valine | | C5H11NO2 | | 118.0863 | [M+H]+ | 8.21 | HMDB0000883 | 1 | 2.03 |  |
| XMP | | C10H13N4O9P | | 363.0347 | [M-H]- | 20.1 | HMDB0001554 | 1 | 2.43 |  |
| Xylulose-5-Phosphate | | C5H11O8P | | 229.0119 | [M-H]- | 18.05 | HMDB0000868 | 1 | -1.58 |  |

### **Table S2** | Matrix effect (ME), Process Efficiency (PE) and Extraction Recovery (RE) of metabolites without internal standard correction (Colorectal organoid extract as matrix, n = 5). The concentration of the QC sample used was 100 ng/mL.

| Metabolite | ME [%] | RE [%] | PE [%] |
| --- | --- | --- | --- |
| 1-Methylhistidine | 103.6 ± 0.7 | 88.6 ± 3.1 | 91.8 ± 2.5 |
| 2,3-Bisphosphoglycerate | 103.7 ± 1.3 | 85.1 ± 2.1 | 88.2 ± 0.8 |
| 2-Aminoadipic Acid | 98.4 ± 0.4 | 91.5 ± 2.8 | 90.0 ± 2.3 |
| 2-Deoxyadenosine | 98.4 ± 3.1 | 82.0 ± 1.8 | 80.7 ± 2.2 |
| 2'-Deoxyadenosine 5'-monophosphate | 104.0 ± 3.0 | 81.9 ± 3.3 | 85.2 ± 2.0 |
| 2'-deoxyadenosine 5'-triphosphate | 88.5 ± 2.3 | 86.7 ± 1.9 | 76.7 ± 1.0 |
| 2-Deoxycytidine | 94.9 ± 0.7 | 80.1 ± 3.4 | 76.0 ± 1.5 |
| 2'-Deoxycytidine-5'-monophosphate | 98.6 ± 1.5 | 82.2 ± 2.4 | 81.0 ± 1.1 |
| 2-Deoxyguanosine | 96.0 ± 1.3 | 88.4 ± 1.4 | 84.9 ± 1.8 |
| 2-Deoxyinosine | 90.4 ± 0.7 | 77.2 ± 0.4 | 69.8 ± 2.0 |
| 2-hydroxybutyric acid | 90.2 ± 2.2 | 80.7 ± 1.5 | 72.8 ± 3.3 |
| 2-hydroxyglutaric acid | 89.3 ± 0.7 | 87.3 ± 3.0 | 78.0 ± 3.4 |
| 2-isopropylmalic acid | 91.9 ± 2.3 | 92.2 ± 1.6 | 84.7 ± 3.5 |
| 2-ketobutyric acid | 92.1 ± 1.5 | 79.1 ± 1.2 | 72.9 ± 3.2 |
| 2-oxoadipic acid | 93.4 ± 2.5 | 82.9 ± 1.9 | 77.4 ± 2.9 |
| 2-phosphoglycerate | 103.6 ± 2.7 | 76.2 ± 1.7 | 78.9 ± 2.8 |
| 3-amino-isobutyric acid | 89.9 ± 0.9 | 86.5 ± 0.9 | 77.8 ± 3.6 |
| 3-hydroxy-3-methyl-2-oxobutanoate | 93.1 ± 3.1 | 76.2 ± 0.7 | 70.9 ± 2.9 |
| 3-hydroxybutyric acid | 104.0 ± 3.5 | 75.2 ± 1.9 | 78.2 ± 1.8 |
| 3-Hydroxy-Proline | 88.1 ± 1.8 | 81.8 ± 2.0 | 72.1 ± 0.6 |
| 3-isopropylmalic acid | 102.5 ± 1.0 | 85.3 ± 0.6 | 87.4 ± 1.1 |
| 3-methyl-2-oxopentanoate | 89.3 ± 2.1 | 76.4 ± 1.8 | 68.2 ± 0.9 |
| 3-Methylhistidine | 94.0 ± 3.2 | 81.2 ± 2.0 | 76.3 ± 1.0 |
| 3-Phosphoglyceraldehyde | 102.8 ± 3.1 | 87.8 ± 3.3 | 90.3 ± 1.3 |
| 3-phosphoglycerate | 98.4 ± 0.7 | 81.3 ± 1.6 | 80.0 ± 0.5 |
| 4-amino-butyric acid | 101.7 ± 2.2 | 92.6 ± 3.6 | 94.2 ± 1.3 |
| 4-Hydroxybenzoic acid | 104.0 ± 0.8 | 83.6 ± 1.7 | 86.9 ± 1.4 |
| 4-Hydroxyglutamate | 97.0 ± 2.4 | 85.6 ± 1.6 | 83.0 ± 1.3 |
| 4-Hydroxy-Proline | 95.6 ± 1.2 | 84.5 ± 2.5 | 80.8 ± 1.5 |
| 5-Aminolevulinic acid | 96.5 ± 1.1 | 83.1 ± 1.4 | 80.2 ± 2.7 |
| 5-Aminovaleric acid | 103.4 ± 2.7 | 84.0 ± 0.5 | 86.9 ± 2.8 |
| 5-Hydroxy-Tryptophan | 89.8 ± 1.4 | 77.6 ± 2.3 | 69.7 ± 1.8 |
| 5-Methoxytryptophan | 92.4 ± 3.3 | 89.7 ± 0.8 | 82.9 ± 2.8 |
| 5-Methylcytosine | 96.6 ± 3.0 | 93.2 ± 1.9 | 90.0 ± 2.8 |
| 6-phosphogluconate | 88.1 ± 1.4 | 77.3 ± 1.9 | 68.1 ± 3.6 |
| Acetoacetate | 103.1 ± 1.7 | 80.6 ± 2.1 | 83.1 ± 1.6 |
| Acetylcarnitine | 99.7 ± 1.0 | 78.8 ± 2.4 | 78.6 ± 1.0 |
| Acetyl-CoA | 93.6 ± 0.6 | 81.1 ± 3.3 | 75.9 ± 1.5 |
| Aconitic Acid | 93.5 ± 1.7 | 76.9 ± 3.4 | 71.9 ± 0.5 |
| Adenine | 91.8 ± 0.8 | 89.1 ± 2.0 | 81.8 ± 1.2 |
| Adenosine | 87.9 ± 1.2 | 94.4 ± 3.2 | 83.0 ± 2.2 |
| Adenosine 3',5'-cyclic monophosphate | 90.4 ± 2.0 | 94.6 ± 2.5 | 85.5 ± 1.3 |
| Adenosine Monophosphate | 91.1 ± 2.0 | 83.8 ± 1.3 | 76.3 ± 2.6 |
| Adenosine-5'-diphosphate | 101.0 ± 3.0 | 86.5 ± 3.1 | 87.4 ± 2.4 |
| Adenosine-5'-triphosphate | 100.3 ± 0.6 | 93.6 ± 1.4 | 93.9 ± 1.2 |
| Adenylosuccinate | 95.5 ± 0.9 | 75.9 ± 3.5 | 72.5 ± 2.0 |
| Alanine | 98.8 ± 2.2 | 94.5 ± 2.4 | 93.4 ± 0.5 |
| Allantoin | 95.0 ± 1.1 | 75.3 ± 0.4 | 71.5 ± 1.8 |
| alpha-ketoglutaric acid | 91.1 ± 0.6 | 91.9 ± 1.2 | 83.7 ± 3.2 |
| Anserine | 97.7 ± 1.3 | 82.2 ± 2.1 | 80.3 ± 2.9 |
| Trigonelleine | 90.3 ± 2.5 | 86.1 ± 1.1 | 77.7 ± 3.3 |
| Arginine | 95.6 ± 2.7 | 78.8 ± 1.4 | 75.3 ± 0.5 |
| Argininosuccinic Acid | 95.5 ± 2.4 | 82.4 ± 2.4 | 78.7 ± 1.6 |
| Ascorbic acid | 101.9 ± 1.2 | 90.8 ± 1.8 | 92.5 ± 2.2 |
| Asparagine | 96.7 ± 1.2 | 89.3 ± 0.7 | 86.4 ± 0.8 |
| Aspartic acid | 96.5 ± 3.4 | 86.3 ± 2.5 | 83.3 ± 0.4 |
| Asymmetric dimethylarginine | 102.0 ± 3.0 | 77.0 ± 2.5 | 78.5 ± 2.1 |
| Azelaic acid | 97.2 ± 2.2 | 77.2 ± 1.8 | 75.0 ± 3.2 |
| beta-Alanine | 102.1 ± 1.6 | 76.2 ± 1.0 | 77.8 ± 0.7 |
| Betaine | 101.2 ± 0.9 | 95.0 ± 3.1 | 96.1 ± 1.7 |
| beta-Ureidopropionate | 97.4 ± 3.6 | 93.8 ± 2.9 | 91.4 ± 3.1 |
| Biotin | 88.0 ± 1.9 | 75.6 ± 1.6 | 66.5 ± 1.7 |
| Cadaverine | 103.8 ± 2.4 | 88.1 ± 2.7 | 91.4 ± 3.3 |
| Carnitine | 90.8 ± 2.3 | 77.7 ± 1.0 | 70.6 ± 0.8 |
| Carnosine | 100.0 ± 3.2 | 93.4 ± 3.4 | 93.4 ± 2.8 |
| cGMP | 100.9 ± 0.8 | 78.2 ± 1.2 | 78.9 ± 1.2 |
| Choline | 103.1 ± 1.2 | 80.1 ± 2.0 | 82.6 ± 0.7 |
| Cinnamic acid | 95.6 ± 2.8 | 90.3 ± 2.8 | 86.3 ± 1.8 |
| Citraconic acid | 97.4 ± 0.5 | 86.3 ± 3.6 | 84.1 ± 0.7 |
| Citramalic acid | 88.6 ± 2.4 | 80.7 ± 0.4 | 71.5 ± 1.9 |
| Citric Acid | 102.5 ± 1.8 | 75.7 ± 0.9 | 77.6 ± 0.7 |
| Citrulline | 99.8 ± 1.6 | 76.2 ± 2.6 | 76.0 ± 0.7 |
| Creatine | 100.3 ± 2.4 | 85.8 ± 1.1 | 86.1 ± 1.3 |
| Creatinine | 97.4 ± 2.5 | 89.1 ± 0.7 | 86.8 ± 2.0 |
| Cystathionine | 96.3 ± 2.1 | 86.2 ± 3.2 | 83.0 ± 2.6 |
| Cysteamine | 102.5 ± 2.0 | 75.5 ± 2.4 | 77.4 ± 1.1 |
| Cysteine | 97.2 ± 1.6 | 93.5 ± 2.3 | 90.9 ± 3.6 |
| Cysteinylglycine | 92.1 ± 1.8 | 93.9 ± 0.4 | 86.5 ± 3.4 |
| Cystine | 100.6 ± 0.4 | 75.7 ± 2.8 | 76.2 ± 3.4 |
| Cytidine | 90.5 ± 1.7 | 77.6 ± 0.4 | 70.2 ± 1.5 |
| Cytidine 5'-diphosphate | 96.4 ± 2.5 | 81.8 ± 1.8 | 78.9 ± 3.0 |
| Cytidine 5'-triphosphate | 88.1 ± 3.4 | 85.6 ± 0.8 | 75.4 ± 2.3 |
| Cytidine monophosphate | 91.3 ± 0.9 | 90.6 ± 2.3 | 82.7 ± 1.3 |
| Cytosine | 91.1 ± 2.0 | 91.0 ± 0.5 | 82.9 ± 2.6 |
| Dihydroxyacetonephosphate | 98.3 ± 3.3 | 77.3 ± 0.4 | 76.0 ± 2.9 |
| Dimethylglycine | 97.0 ± 2.7 | 94.8 ± 1.7 | 92.0 ± 0.9 |
| Erythrose-4-phosphate | 90.1 ± 1.8 | 77.3 ± 2.6 | 69.6 ± 1.4 |
| Ethanolamine | 99.0 ± 0.7 | 83.1 ± 1.0 | 82.3 ± 0.8 |
| FAD | 89.0 ± 2.7 | 93.3 ± 1.4 | 83.0 ± 2.9 |
| Folic acid | 102.5 ± 1.0 | 83.3 ± 0.6 | 85.4 ± 1.8 |
| Fructose | 91.5 ± 2.3 | 89.4 ± 2.4 | 81.8 ± 3.0 |
| Fructose-1,6-bisphosphate | 97.7 ± 2.9 | 77.4 ± 1.0 | 75.6 ± 3.4 |
| Fructose-6-Phosphate | 89.4 ± 3.1 | 76.2 ± 1.7 | 68.1 ± 0.8 |
| Fumaric Acid | 92.9 ± 0.5 | 81.7 ± 3.0 | 75.9 ± 3.2 |
| Galactosamine | 104.0 ± 1.2 | 87.6 ± 0.8 | 91.1 ± 3.0 |
| Galactose-1-phosphate | 93.1 ± 2.2 | 89.4 ± 2.2 | 83.2 ± 2.7 |
| Galactose-6-phosphate | 101.7 ± 1.8 | 87.2 ± 0.5 | 88.7 ± 3.4 |
| gamma-Glutamylcysteine | 89.4 ± 2.4 | 93.8 ± 0.8 | 83.9 ± 2.2 |
| Glucaric Acid | 98.2 ± 1.5 | 81.7 ± 2.7 | 80.2 ± 2.5 |
| Gluconic acid | 88.8 ± 1.6 | 85.5 ± 2.4 | 75.9 ± 3.5 |
| Gluconolactone | 96.1 ± 3.1 | 89.9 ± 2.0 | 86.4 ± 1.4 |
| Glucosamine | 90.9 ± 3.2 | 90.6 ± 2.4 | 82.4 ± 2.4 |
| Glucosamine-6-Phosphate | 97.4 ± 2.2 | 77.6 ± 1.6 | 75.6 ± 3.4 |
| Glucose | 99.6 ± 3.5 | 86.5 ± 2.8 | 86.2 ± 2.0 |
| Glucose-1-phosphate | 89.2 ± 2.5 | 80.1 ± 3.0 | 71.4 ± 1.1 |
| Glucose-6-phosphate | 96.3 ± 2.7 | 81.0 ± 1.3 | 78.0 ± 3.2 |
| Glucuronolactone | 88.0 ± 3.1 | 77.9 ± 2.8 | 68.6 ± 1.3 |
| Glutamic acid | 88.5 ± 1.7 | 90.3 ± 1.6 | 79.9 ± 2.4 |
| Glutamine | 98.9 ± 1.1 | 81.5 ± 2.4 | 80.6 ± 0.5 |
| Glutathione (GSSG) | 98.6 ± 1.7 | 78.6 ± 1.3 | 77.5 ± 1.2 |
| Glutathione | 89.3 ± 2.2 | 91.8 ± 3.0 | 82.0 ± 2.2 |
| Glycerophosphocholine | 102.0 ± 3.3 | 77.6 ± 1.1 | 79.2 ± 1.8 |
| Glycine | 89.0 ± 1.1 | 78.1 ± 3.4 | 69.5 ± 3.5 |
| Glycolic acid | 94.2 ± 1.0 | 88.2 ± 2.9 | 83.1 ± 3.5 |
| Guanine | 101.0 ± 1.5 | 75.8 ± 1.3 | 76.6 ± 2.6 |
| Guanosine | 100.9 ± 2.2 | 78.1 ± 2.6 | 78.8 ± 2.6 |
| Guanosine 5'-diphosphate | 89.2 ± 2.7 | 86.0 ± 3.2 | 76.7 ± 3.2 |
| Guanosine-5'-monophosphate | 99.1 ± 2.8 | 80.2 ± 1.4 | 79.5 ± 1.0 |
| Hippuric acid | 92.3 ± 2.1 | 86.2 ± 0.8 | 79.6 ± 3.4 |
| Histamine | 93.0 ± 2.6 | 86.9 ± 2.2 | 80.8 ± 3.4 |
| Histidine | 91.2 ± 2.2 | 92.7 ± 2.8 | 84.5 ± 0.7 |
| Histidinol | 98.1 ± 2.2 | 89.5 ± 2.3 | 87.8 ± 2.9 |
| Homoalanine | 93.1 ± 2.8 | 81.4 ± 0.8 | 75.8 ± 1.3 |
| Homocitrulline | 88.1 ± 0.4 | 89.0 ± 2.3 | 78.4 ± 0.6 |
| Homocysteine | 98.3 ± 2.4 | 86.4 ± 3.0 | 85.0 ± 0.9 |
| Homocystine | 98.2 ± 1.2 | 88.5 ± 1.8 | 86.9 ± 3.0 |
| Homogentisic acid | 101.7 ± 2.8 | 76.9 ± 1.3 | 78.2 ± 3.1 |
| Homoleucine | 102.3 ± 2.2 | 77.5 ± 3.5 | 79.3 ± 1.1 |
| Homophenylalanine | 100.8 ± 2.8 | 91.6 ± 2.6 | 92.4 ± 2.1 |
| Homoserine | 100.2 ± 2.7 | 82.0 ± 0.5 | 82.2 ± 2.5 |
| Homovanillic acid | 95.2 ± 0.9 | 77.6 ± 2.9 | 73.9 ± 0.8 |
| Hypotaurine | 93.1 ± 2.0 | 91.5 ± 3.0 | 85.2 ± 0.9 |
| Hypoxanthine | 99.5 ± 2.8 | 88.8 ± 0.6 | 88.3 ± 1.8 |
| Inosine | 103.5 ± 2.0 | 82.1 ± 0.5 | 85.0 ± 3.4 |
| Inosine 5'-monophosphate | 93.5 ± 0.9 | 77.0 ± 3.1 | 72.0 ± 1.1 |
| Inositol | 88.4 ± 2.6 | 77.7 ± 0.5 | 68.7 ± 2.4 |
| Isocitric Acid | 95.7 ± 2.5 | 92.8 ± 2.4 | 88.8 ± 2.1 |
| Isoleucine | 101.4 ± 2.9 | 83.5 ± 2.5 | 84.6 ± 0.4 |
| Itaconic acid | 91.2 ± 2.0 | 75.1 ± 1.3 | 68.5 ± 2.2 |
| ITP | 98.2 ± 0.7 | 87.0 ± 3.6 | 85.4 ± 1.2 |
| Kynurenic acid | 98.5 ± 1.0 | 91.2 ± 3.0 | 89.9 ± 3.2 |
| Kynurenine | 89.0 ± 0.6 | 76.5 ± 2.3 | 68.1 ± 3.0 |
| Lactic Acid | 92.2 ± 2.9 | 80.0 ± 3.1 | 73.8 ± 2.9 |
| Lactose | 88.6 ± 2.5 | 91.7 ± 1.9 | 81.2 ± 3.5 |
| Leucine | 103.8 ± 2.7 | 91.1 ± 3.3 | 94.6 ± 1.8 |
| Levulinic acid | 93.3 ± 3.5 | 79.5 ± 3.5 | 74.2 ± 0.9 |
| Lysine | 92.5 ± 2.4 | 80.8 ± 1.6 | 74.7 ± 3.4 |
| Malic Acid | 88.7 ± 1.9 | 91.7 ± 0.7 | 81.3 ± 2.4 |
| Malonic acid | 99.2 ± 3.3 | 75.5 ± 2.8 | 74.9 ± 2.3 |
| Mannose | 93.0 ± 2.7 | 82.0 ± 1.8 | 76.3 ± 2.4 |
| Mannose-1 with Mannose-6-phosphate | 100.3 ± 1.9 | 81.2 ± 2.6 | 81.4 ± 1.8 |
| Mesaconic acid | 89.9 ± 1.5 | 90.2 ± 1.3 | 81.1 ± 0.5 |
| Methionine | 102.3 ± 1.0 | 93.3 ± 2.6 | 95.5 ± 0.8 |
| Methionine sulfone | 95.0 ± 1.1 | 90.5 ± 1.3 | 85.9 ± 3.0 |
| Methionine sulfoxide | 96.3 ± 1.9 | 78.3 ± 2.7 | 75.4 ± 1.8 |
| Methylmalonate | 88.2 ± 3.3 | 75.8 ± 2.1 | 66.9 ± 3.4 |
| N-Acetyl-Alanine | 95.1 ± 3.3 | 94.3 ± 3.0 | 89.7 ± 0.6 |
| N-Acetyl-Aspartic Acid | 89.6 ± 0.6 | 86.7 ± 0.5 | 77.7 ± 0.7 |
| N-Acetyl-Citrulline | 101.6 ± 2.9 | 77.6 ± 1.7 | 78.8 ± 1.9 |
| N-Acetyl-Galactosamine | 90.3 ± 1.2 | 78.1 ± 1.9 | 70.5 ± 0.4 |
| N-Acetyl-Glucosamine | 92.7 ± 1.3 | 76.2 ± 1.7 | 70.6 ± 2.2 |
| N-Acetyl-Glutamine | 89.5 ± 0.6 | 78.8 ± 2.5 | 70.5 ± 0.9 |
| N-Acetyl-Glycine | 103.3 ± 2.6 | 84.5 ± 2.7 | 87.3 ± 1.3 |
| N-Acetyl-Leucine | 103.5 ± 2.7 | 84.9 ± 2.5 | 87.9 ± 3.5 |
| N-Acetyl-Methionine | 100.6 ± 2.6 | 90.2 ± 3.1 | 90.7 ± 3.3 |
| N-Acetylneuraminic acid | 97.8 ± 3.1 | 87.3 ± 0.6 | 85.3 ± 1.1 |
| N-Acetyl-Phenylalanine | 100.3 ± 1.9 | 88.7 ± 1.8 | 89.0 ± 2.4 |
| N-Acetyl-Proline | 88.8 ± 3.4 | 79.9 ± 1.7 | 70.9 ± 2.7 |
| N-Acetyl-Serine | 101.9 ± 1.1 | 93.3 ± 1.4 | 95.1 ± 1.7 |
| N-Acetyl-Serotonin | 93.0 ± 2.8 | 77.1 ± 1.0 | 71.7 ± 3.6 |
| N-Acetyl-Tryptophan | 93.0 ± 1.3 | 91.9 ± 1.7 | 85.5 ± 3.0 |
| N-Acetyl-Tyrosine | 97.0 ± 1.1 | 79.7 ± 3.6 | 77.3 ± 3.5 |
| NAD | 97.9 ± 3.4 | 77.2 ± 1.4 | 75.5 ± 0.9 |
| NADH | 97.1 ± 3.1 | 89.7 ± 3.0 | 87.1 ± 2.3 |
| NADP | 97.7 ± 3.2 | 84.9 ± 2.1 | 83.0 ± 0.7 |
| NADPH | 96.2 ± 0.5 | 82.9 ± 2.0 | 79.8 ± 2.9 |
| N-alpha-Acetyl-lysine | 92.8 ± 2.4 | 75.4 ± 3.6 | 69.9 ± 3.2 |
| N-alpha-Acetyl-Ornithine | 91.8 ± 3.1 | 85.1 ± 1.4 | 78.1 ± 2.9 |
| N-epsilon-Acetyl-lysine | 96.5 ± 2.1 | 78.7 ± 0.9 | 75.9 ± 2.8 |
| N-epsilon-Acetyl-Ornithine | 89.5 ± 1.5 | 76.1 ± 3.0 | 68.1 ± 1.1 |
| Nicotinamide | 97.2 ± 1.1 | 84.9 ± 0.9 | 82.5 ± 3.2 |
| Nicotinic acid | 95.1 ± 1.1 | 86.9 ± 0.6 | 82.6 ± 1.3 |
| N-Methyl-Aspartic Acid | 89.0 ± 3.3 | 91.4 ± 3.6 | 81.3 ± 1.8 |
| O-Acetyl-Serine | 90.2 ± 1.3 | 87.5 ± 2.1 | 78.9 ± 1.8 |
| O-Phosphoethanolamine | 89.3 ± 2.8 | 91.7 ± 1.0 | 81.9 ± 1.4 |
| O-Phospho-Serine | 95.1 ± 0.9 | 83.7 ± 2.3 | 79.6 ± 0.9 |
| Ophthalmic acid | 88.2 ± 1.7 | 78.2 ± 2.3 | 69.0 ± 1.0 |
| Ornithine | 90.1 ± 2.6 | 92.7 ± 3.1 | 83.5 ± 2.6 |
| Orotic acid | 98.0 ± 1.2 | 93.7 ± 3.2 | 91.8 ± 0.6 |
| Oxaloacetate | 88.3 ± 0.6 | 88.6 ± 0.3 | 78.3 ± 2.6 |
| Pantothenic acid | 92.5 ± 1.9 | 93.8 ± 0.8 | 86.8 ± 2.8 |
| p-Coumaric Acid | 99.5 ± 1.3 | 83.8 ± 2.7 | 83.3 ± 2.9 |
| Phenylalanine | 99.0 ± 3.3 | 90.4 ± 1.4 | 89.5 ± 1.4 |
| Phenylpyruvate | 98.9 ± 0.9 | 87.3 ± 1.3 | 86.4 ± 1.6 |
| Phosphocreatine | 99.4 ± 2.0 | 94.7 ± 3.2 | 94.1 ± 2.2 |
| Phosphoenolpyruvate | 96.4 ± 3.3 | 76.9 ± 1.1 | 74.1 ± 0.6 |
| Phosphoribosylpyrophosphate | 95.5 ± 2.5 | 83.3 ± 2.8 | 79.5 ± 2.4 |
| Picolinate | 92.4 ± 2.5 | 83.5 ± 1.2 | 77.2 ± 3.4 |
| Pipecolic acid | 100.1 ± 2.9 | 88.6 ± 3.2 | 88.6 ± 0.9 |
| Proline | 101.2 ± 1.5 | 94.2 ± 1.2 | 95.3 ± 3.4 |
| Pyridoxal | 95.4 ± 3.1 | 89.0 ± 3.0 | 84.9 ± 3.2 |
| Pyridoxal-5-phosphate | 96.2 ± 0.8 | 92.7 ± 1.0 | 89.2 ± 0.7 |
| Pyridoxamine | 89.9 ± 2.7 | 90.6 ± 2.2 | 81.4 ± 1.0 |
| Pyridoxine | 103.8 ± 2.1 | 91.0 ± 1.2 | 94.5 ± 2.8 |
| Pyroglutamic acid | 90.4 ± 2.3 | 87.8 ± 1.0 | 79.3 ± 0.3 |
| Pyrrolysine | 102.6 ± 2.0 | 81.8 ± 2.6 | 84.0 ± 3.5 |
| Pyruvic Acid | 93.1 ± 2.2 | 80.7 ± 1.2 | 75.1 ± 3.0 |
| Riboflavin | 100.8 ± 0.7 | 75.3 ± 2.4 | 75.9 ± 0.5 |
| Ribose | 94.1 ± 1.1 | 84.7 ± 3.0 | 79.7 ± 1.2 |
| Ribose-5-phosphate | 99.4 ± 2.6 | 87.2 ± 2.5 | 86.7 ± 1.6 |
| Ribulose-1-phosphate | 98.4 ± 2.5 | 84.4 ± 1.1 | 83.1 ± 3.3 |
| Ribulose-5-phosphate | 99.5 ± 2.5 | 81.1 ± 3.5 | 80.7 ± 1.9 |
| S-Adenosylmethionine | 98.1 ± 2.7 | 79.9 ± 2.6 | 78.4 ± 2.4 |
| Sarcosine | 91.7 ± 3.5 | 76.3 ± 1.4 | 69.9 ± 1.7 |
| Sedoheptulose-7-phosphate | 93.0 ± 0.5 | 80.4 ± 3.1 | 74.8 ± 3.2 |
| Selenocystine | 90.6 ± 1.3 | 82.6 ± 1.7 | 74.9 ± 1.6 |
| Selenomethionine | 100.7 ± 0.8 | 84.7 ± 2.3 | 85.3 ± 0.8 |
| Serine | 92.8 ± 0.7 | 88.8 ± 1.1 | 82.4 ± 3.1 |
| Serotonin | 93.7 ± 0.9 | 94.8 ± 3.2 | 88.9 ± 0.4 |
| Sorbitol | 99.7 ± 0.4 | 82.8 ± 2.0 | 82.5 ± 3.4 |
| Spermidine | 96.2 ± 3.5 | 87.6 ± 1.2 | 84.3 ± 3.0 |
| Spermine | 97.6 ± 3.2 | 90.9 ± 3.3 | 88.7 ± 2.3 |
| Succinic Acid | 94.4 ± 2.7 | 87.6 ± 0.9 | 82.7 ± 3.0 |
| Sucrose | 94.2 ± 3.0 | 77.2 ± 1.0 | 72.7 ± 0.4 |
| Symmetric dimethylarginine | 95.7 ± 1.0 | 82.5 ± 3.4 | 79.0 ± 0.3 |
| Taurine | 100.8 ± 3.2 | 78.1 ± 3.5 | 78.8 ± 1.3 |
| Thiamine | 89.7 ± 3.4 | 92.6 ± 2.9 | 83.0 ± 0.9 |
| Thiamine monophosphate | 93.2 ± 3.1 | 84.9 ± 3.1 | 79.1 ± 2.3 |
| Thiamine pyrophosphate | 98.9 ± 3.1 | 78.3 ± 2.3 | 77.5 ± 1.4 |
| Threonic acid | 91.9 ± 0.4 | 84.1 ± 0.6 | 77.3 ± 3.1 |
| Threonine | 96.7 ± 1.1 | 78.2 ± 2.5 | 75.6 ± 1.1 |
| Thymidine | 95.0 ± 3.5 | 89.8 ± 1.3 | 85.3 ± 0.9 |
| Thymidine Monophosphate | 94.5 ± 3.4 | 87.7 ± 1.8 | 82.9 ± 1.1 |
| Thymine | 101.8 ± 1.4 | 78.8 ± 2.6 | 80.2 ± 3.4 |
| Trimethylamine N-oxide | 95.7 ± 3.3 | 81.5 ± 1.4 | 78.0 ± 0.4 |
| Tryptophan | 103.3 ± 2.7 | 85.1 ± 1.0 | 87.9 ± 2.5 |
| Tyrosine | 99.0 ± 3.4 | 89.6 ± 3.2 | 88.7 ± 3.4 |
| Uracil | 102.0 ± 0.7 | 81.6 ± 1.3 | 83.2 ± 2.4 |
| Uric Acid | 89.2 ± 1.2 | 90.9 ± 0.8 | 81.1 ± 1.7 |
| Uridine | 96.6 ± 2.6 | 81.5 ± 0.7 | 78.7 ± 2.4 |
| Uridine 5'-diphosphate | 89.3 ± 1.6 | 91.3 ± 1.3 | 81.5 ± 1.0 |
| Uridine Monophosphate | 94.2 ± 2.2 | 91.1 ± 3.4 | 85.8 ± 2.4 |
| Uridine-5-Diphosphogalactose | 99.4 ± 2.1 | 86.1 ± 3.4 | 85.5 ± 1.3 |
| Uridine-5-Diphosphoglucose | 91.2 ± 1.4 | 82.0 ± 3.2 | 74.8 ± 0.5 |
| Urocanic acid | 90.7 ± 2.9 | 85.7 ± 0.7 | 77.8 ± 3.5 |
| UTP | 95.4 ± 2.2 | 88.2 ± 1.3 | 84.2 ± 2.1 |
| Valine | 89.4 ± 2.5 | 84.3 ± 1.5 | 75.4 ± 2.3 |
| Xanthine | 92.2 ± 1.0 | 83.1 ± 0.3 | 76.6 ± 1.8 |
| Xanthosine | 96.1 ± 0.9 | 84.1 ± 1.9 | 80.8 ± 2.4 |
| Xanthurenic acid | 100.1 ± 0.8 | 76.2 ± 2.4 | 76.3 ± 3.6 |
| XMP | 101.8 ± 0.7 | 82.7 ± 0.8 | 84.2 ± 2.2 |
| Xylulose-5-phosphate | 90.6 ± 2.2 | 86.0 ± 0.8 | 77.9 ± 1.3 |

### **Table S3** | Matrix effect (ME), Process Efficiency (PE) and Extraction Recovery (RE) of metabolites without internal standard correction (Tumor tissue extract as matrix, n = 5). The concentration of the QC sample used was 100 ng/mL.

| Metabolite | ME [%] | | RE [%] | | PE [%] |
| --- | --- | --- | --- | --- | --- |
| 1-Methylhistidine | | 103.0 ± 3.4 | 84.1 ± 3.1 | 86.6 ± 3.0 | |
| 2,3-Bisphosphoglycerate | | 103.6 ± 2.8 | 89.9 ± 1.9 | 93.1 ± 2.8 | |
| 2-Aminoadipic Acid | | 103.7 ± 0.9 | 93.1 ± 3.3 | 96.5 ± 2.4 | |
| 2-Deoxyadenosine | | 98.4 ± 2.3 | 78.6 ± 1.8 | 77.3 ± 2.3 | |
| 2'-Deoxyadenosine 5'-monophosphate | | 98.4 ± 0.9 | 86.4 ± 1.8 | 85.0 ± 0.8 | |
| 2'-deoxyadenosine 5'-triphosphate | | 104.0 ± 2.8 | 83.0 ± 1.6 | 86.3 ± 2.2 | |
| 2-Deoxycytidine | | 88.5 ± 0.9 | 94.6 ± 2.0 | 83.7 ± 1.1 | |
| 2'-Deoxycytidine-5'-monophosphate | | 94.9 ± 1.6 | 89.7 ± 1.7 | 85.1 ± 0.8 | |
| 2-Deoxyguanosine | | 98.6 ± 3.2 | 82.0 ± 1.8 | 80.9 ± 2.0 | |
| 2-Deoxyinosine | | 96.0 ± 3.2 | 76.5 ± 2.9 | 73.4 ± 1.2 | |
| 2-hydroxybutyric acid | | 90.4 ± 2.5 | 77.8 ± 3.3 | 70.3 ± 2.6 | |
| 2-hydroxyglutaric acid | | 90.2 ± 2.1 | 80.0 ± 0.7 | 72.2 ± 3.2 | |
| 2-isopropylmalic acid | | 89.3 ± 0.5 | 79.2 ± 3.4 | 70.7 ± 2.2 | |
| 2-ketobutyric acid | | 91.9 ± 2.1 | 88.2 ± 3.2 | 81.1 ± 2.4 | |
| 2-oxoadipic acid | | 92.1 ± 2.6 | 79.3 ± 2.4 | 73.0 ± 0.6 | |
| 2-phosphoglycerate | | 93.4 ± 2.7 | 87.1 ± 0.9 | 81.4 ± 1.7 | |
| 3-amino-isobutyric acid | | 103.6 ± 3.0 | 94.3 ± 3.2 | 97.7 ± 0.7 | |
| 3-hydroxy-3-methyl-2-oxobutanoate | | 89.9 ± 3.1 | 92.9 ± 0.9 | 83.5 ± 1.2 | |
| 3-hydroxybutyric acid | | 93.1 ± 2.3 | 93.5 ± 2.4 | 87.0 ± 1.0 | |
| 3-Hydroxy-Proline | | 104.0 ± 2.3 | 86.5 ± 2.4 | 90.0 ± 1.6 | |
| 3-isopropylmalic acid | | 88.1 ± 3.2 | 77.4 ± 0.5 | 68.2 ± 1.6 | |
| 3-methyl-2-oxopentanoate | | 102.5 ± 0.8 | 91.4 ± 0.6 | 93.7 ± 2.6 | |
| 3-Methylhistidine | | 89.3 ± 2.1 | 86.0 ± 2.7 | 76.8 ± 3.2 | |
| 3-Phosphoglyceraldehyde | | 94.0 ± 0.7 | 93.4 ± 1.8 | 87.8 ± 3.2 | |
| 3-phosphoglycerate | | 102.8 ± 3.3 | 94.5 ± 3.2 | 97.1 ± 1.4 | |
| 4-amino-butyric acid | | 98.4 ± 1.9 | 88.0 ± 3.1 | 86.6 ± 2.3 | |
| 4-Hydroxybenzoic acid | | 101.7 ± 0.6 | 83.6 ± 3.3 | 85.0 ± 1.8 | |
| 4-Hydroxyglutamate | | 104.0 ± 3.4 | 86.1 ± 3.5 | 89.5 ± 1.0 | |
| 4-Hydroxy-Proline | | 97.0 ± 3.5 | 87.2 ± 0.5 | 84.6 ± 2.3 | |
| 5-Aminolevulinic acid | | 95.6 ± 3.3 | 83.2 ± 2.0 | 79.5 ± 2.6 | |
| 5-Aminovaleric acid | | 96.5 ± 2.4 | 75.8 ± 0.6 | 73.1 ± 3.4 | |
| 5-Hydroxy-Tryptophan | | 103.4 ± 3.1 | 85.2 ± 2.4 | 88.1 ± 2.5 | |
| 5-Methoxytryptophan | | 89.8 ± 1.0 | 78.7 ± 3.0 | 70.7 ± 3.1 | |
| 5-Methylcytosine | | 92.4 ± 1.5 | 79.9 ± 3.5 | 73.8 ± 3.2 | |
| 6-phosphogluconate | | 96.6 ± 0.9 | 92.3 ± 0.6 | 89.2 ± 1.9 | |
| Acetoacetate | | 88.1 ± 1.8 | 75.2 ± 2.0 | 66.3 ± 1.1 | |
| Acetylcarnitine | | 103.1 ± 1.0 | 86.2 ± 2.5 | 88.9 ± 2.3 | |
| Acetyl-CoA | | 99.7 ± 2.3 | 85.8 ± 2.2 | 85.5 ± 1.5 | |
| Aconitic Acid | | 93.6 ± 1.4 | 75.7 ± 3.4 | 70.9 ± 2.4 | |
| Adenine | | 93.5 ± 2.5 | 80.1 ± 1.1 | 74.9 ± 0.9 | |
| Adenosine | | 91.8 ± 0.8 | 81.6 ± 0.7 | 74.9 ± 2.8 | |
| Adenosine 3',5'-cyclic monophosphate | | 87.9 ± 1.8 | 76.5 ± 1.5 | 67.2 ± 0.4 | |
| Adenosine Monophosphate | | 90.4 ± 3.1 | 85.5 ± 1.4 | 77.3 ± 3.0 | |
| Adenosine-5'-diphosphate | | 91.1 ± 1.9 | 89.1 ± 1.5 | 81.2 ± 2.9 | |
| Adenosine-5'-triphosphate | | 101.0 ± 0.9 | 79.2 ± 3.4 | 80.0 ± 3.2 | |
| Adenylosuccinate | | 100.3 ± 2.7 | 82.7 ± 2.9 | 82.9 ± 2.2 | |
| Alanine | | 95.5 ± 2.8 | 82.0 ± 1.5 | 78.3 ± 1.9 | |
| Allantoin | | 98.8 ± 2.3 | 79.5 ± 0.8 | 78.5 ± 2.0 | |
| alpha-ketoglutaric acid | | 95.0 ± 3.5 | 83.9 ± 3.2 | 79.7 ± 1.4 | |
| Anserine | | 91.1 ± 3.5 | 85.0 ± 0.9 | 77.4 ± 2.1 | |
| Trigonelleine | | 97.7 ± 2.4 | 77.4 ± 0.4 | 75.6 ± 2.1 | |
| Arginine | | 90.3 ± 0.3 | 82.3 ± 2.5 | 74.3 ± 3.3 | |
| Argininosuccinic Acid | | 95.6 ± 0.4 | 85.5 ± 1.5 | 81.7 ± 1.0 | |
| Ascorbic acid | | 95.5 ± 2.8 | 90.5 ± 0.3 | 86.4 ± 1.0 | |
| Asparagine | | 101.9 ± 0.8 | 78.7 ± 2.2 | 80.2 ± 1.6 | |
| Aspartic acid | | 96.7 ± 1.9 | 82.0 ± 2.8 | 79.3 ± 0.7 | |
| Asymmetric dimethylarginine | | 96.5 ± 3.1 | 78.3 ± 2.9 | 75.6 ± 0.5 | |
| Azelaic acid | | 102.0 ± 3.1 | 78.9 ± 1.2 | 80.5 ± 1.1 | |
| beta-Alanine | | 97.2 ± 0.7 | 84.4 ± 1.4 | 82.0 ± 3.2 | |
| Betaine | | 102.1 ± 3.1 | 93.0 ± 2.0 | 95.0 ± 1.7 | |
| beta-Ureidopropionate | | 101.2 ± 1.5 | 91.3 ± 2.5 | 92.4 ± 3.6 | |
| Biotin | | 97.4 ± 3.5 | 77.5 ± 2.6 | 75.5 ± 1.8 | |
| Cadaverine | | 88.0 ± 1.8 | 79.7 ± 0.7 | 70.1 ± 2.3 | |
| Carnitine | | 103.8 ± 0.9 | 93.6 ± 2.7 | 97.2 ± 2.2 | |
| Carnosine | | 90.8 ± 1.3 | 94.7 ± 0.5 | 86.0 ± 2.5 | |
| cGMP | | 100.0 ± 3.5 | 86.9 ± 1.2 | 86.9 ± 2.1 | |
| Choline | | 100.9 ± 3.5 | 80.6 ± 3.2 | 81.3 ± 0.6 | |
| Cinnamic acid | | 103.1 ± 0.9 | 85.3 ± 1.4 | 87.9 ± 0.4 | |
| Citraconic acid | | 95.6 ± 1.7 | 94.4 ± 3.2 | 90.2 ± 0.6 | |
| Citramalic acid | | 97.4 ± 3.5 | 89.2 ± 1.5 | 86.9 ± 1.1 | |
| Citric Acid | | 88.6 ± 1.1 | 92.5 ± 3.0 | 82.0 ± 0.3 | |
| Citrulline | | 102.5 ± 3.0 | 85.7 ± 2.0 | 87.8 ± 1.3 | |
| Creatine | | 99.8 ± 2.2 | 83.8 ± 0.3 | 83.6 ± 3.1 | |
| Creatinine | | 100.3 ± 3.0 | 81.6 ± 3.3 | 81.8 ± 2.8 | |
| Cystathionine | | 97.4 ± 0.7 | 75.4 ± 0.7 | 73.4 ± 1.8 | |
| Cysteamine | | 96.3 ± 0.5 | 79.9 ± 1.4 | 76.9 ± 2.8 | |
| Cysteine | | 102.5 ± 2.3 | 89.9 ± 0.8 | 92.1 ± 1.3 | |
| Cysteinylglycine | | 97.2 ± 2.7 | 88.7 ± 1.0 | 86.2 ± 1.2 | |
| Cystine | | 92.1 ± 2.8 | 80.9 ± 0.6 | 74.5 ± 3.6 | |
| Cytidine | | 100.6 ± 1.3 | 76.6 ± 3.5 | 77.1 ± 0.3 | |
| Cytidine 5'-diphosphate | | 90.5 ± 0.9 | 93.8 ± 0.3 | 84.9 ± 2.1 | |
| Cytidine 5'-triphosphate | | 96.4 ± 3.5 | 89.5 ± 0.6 | 86.3 ± 2.5 | |
| Cytidine monophosphate | | 88.1 ± 1.9 | 77.4 ± 0.8 | 68.2 ± 1.2 | |
| Cytosine | | 91.3 ± 1.3 | 88.5 ± 1.4 | 80.8 ± 0.5 | |
| Dihydroxyacetonephosphate | | 91.1 ± 2.7 | 83.5 ± 3.2 | 76.1 ± 2.9 | |
| Dimethylglycine | | 98.3 ± 1.9 | 84.7 ± 2.2 | 83.3 ± 1.5 | |
| Erythrose-4-phosphate | | 97.0 ± 1.4 | 83.0 ± 3.0 | 80.5 ± 0.6 | |
| Ethanolamine | | 90.1 ± 2.1 | 88.4 ± 1.4 | 79.6 ± 1.6 | |
| FAD | | 99.0 ± 0.6 | 89.6 ± 3.1 | 88.7 ± 1.1 | |
| Folic acid | | 89.0 ± 2.6 | 85.5 ± 0.7 | 76.1 ± 1.0 | |
| Fructose | | 102.5 ± 3.2 | 77.6 ± 0.8 | 79.5 ± 3.6 | |
| Fructose-1,6-bisphosphate | | 91.5 ± 3.6 | 78.8 ± 1.1 | 72.1 ± 3.5 | |
| Fructose-6-Phosphate | | 97.7 ± 0.5 | 75.4 ± 2.3 | 73.7 ± 0.5 | |
| Fumaric Acid | | 89.4 ± 1.4 | 93.4 ± 3.2 | 83.5 ± 0.9 | |
| Galactosamine | | 92.9 ± 3.0 | 76.4 ± 3.6 | 71.0 ± 0.8 | |
| Galactose-1-phosphate | | 104.0 ± 3.5 | 86.3 ± 1.7 | 89.8 ± 1.2 | |
| Galactose-6-phosphate | | 93.1 ± 2.8 | 87.6 ± 2.9 | 81.6 ± 1.2 | |
| gamma-Glutamylcysteine | | 101.7 ± 1.0 | 79.9 ± 1.2 | 81.3 ± 2.8 | |
| Glucaric Acid | | 89.4 ± 1.2 | 85.2 ± 1.5 | 76.2 ± 1.6 | |
| Gluconic acid | | 98.2 ± 2.3 | 86.7 ± 2.8 | 85.1 ± 2.1 | |
| Gluconolactone | | 88.8 ± 0.8 | 87.8 ± 2.2 | 78.0 ± 3.0 | |
| Glucosamine | | 96.1 ± 2.6 | 75.3 ± 3.1 | 72.4 ± 1.4 | |
| Glucosamine-6-Phosphate | | 90.9 ± 1.3 | 77.1 ± 2.7 | 70.1 ± 0.5 | |
| Glucose | | 97.4 ± 3.4 | 86.8 ± 2.5 | 84.5 ± 0.4 | |
| Glucose-1-phosphate | | 99.6 ± 3.1 | 85.5 ± 2.7 | 85.2 ± 2.6 | |
| Glucose-6-phosphate | | 89.2 ± 2.2 | 90.1 ± 3.2 | 80.4 ± 2.5 | |
| Glucuronolactone | | 96.3 ± 0.7 | 88.6 ± 0.8 | 85.3 ± 2.7 | |
| Glutamic acid | | 88.0 ± 2.2 | 92.0 ± 2.1 | 81.0 ± 2.0 | |
| Glutamine | | 88.5 ± 2.7 | 81.8 ± 0.8 | 72.4 ± 2.9 | |
| Glutathione (GSSG) | | 98.9 ± 2.8 | 81.2 ± 1.7 | 80.3 ± 1.9 | |
| Glutathione | | 98.6 ± 1.1 | 94.1 ± 2.2 | 92.8 ± 2.6 | |
| Glycerophosphocholine | | 89.3 ± 3.5 | 78.3 ± 2.8 | 69.9 ± 1.6 | |
| Glycine | | 102.0 ± 2.9 | 91.5 ± 3.6 | 93.3 ± 0.6 | |
| Glycolic acid | | 89.0 ± 3.5 | 83.9 ± 2.6 | 74.7 ± 1.0 | |
| Guanine | | 94.2 ± 0.8 | 89.7 ± 2.5 | 84.5 ± 1.2 | |
| Guanosine | | 101.0 ± 2.6 | 90.3 ± 3.3 | 91.2 ± 1.4 | |
| Guanosine 5'-diphosphate | | 100.9 ± 0.5 | 94.4 ± 2.5 | 95.2 ± 2.0 | |
| Guanosine-5'-monophosphate | | 89.2 ± 1.3 | 83.7 ± 1.6 | 74.7 ± 2.7 | |
| Hippuric acid | | 99.1 ± 1.6 | 90.1 ± 1.8 | 89.3 ± 0.4 | |
| Histamine | | 92.3 ± 2.9 | 83.2 ± 3.4 | 76.8 ± 1.6 | |
| Histidine | | 93.0 ± 3.0 | 81.1 ± 0.6 | 75.4 ± 1.8 | |
| Histidinol | | 91.2 ± 1.8 | 80.1 ± 1.2 | 73.1 ± 3.5 | |
| Homoalanine | | 98.1 ± 0.5 | 89.4 ± 1.2 | 87.7 ± 1.4 | |
| Homocitrulline | | 93.1 ± 1.1 | 88.2 ± 0.6 | 82.1 ± 1.8 | |
| Homocysteine | | 88.7 ± 2.8 | 87.6 ± 0.9 | 77.7 ± 2.0 | |
| Homocystine | | 98.0 ± 0.4 | 79.2 ± 3.4 | 77.6 ± 1.4 | |
| Homogentisic acid | | 95.4 ± 3.2 | 76.4 ± 2.7 | 72.9 ± 2.4 | |
| Homoleucine | | 95.7 ± 1.8 | 92.9 ± 2.4 | 88.9 ± 3.3 | |
| Homophenylalanine | | 94.4 ± 2.7 | 91.5 ± 3.4 | 86.4 ± 1.4 | |
| Homoserine | | 99.4 ± 1.7 | 91.6 ± 0.4 | 91.1 ± 1.0 | |
| Homovanillic acid | | 91.5 ± 3.1 | 75.7 ± 3.0 | 69.3 ± 3.1 | |
| Hypotaurine | | 88.0 ± 1.9 | 88.5 ± 2.3 | 77.9 ± 1.1 | |
| Hypoxanthine | | 87.5 ± 0.3 | 92.6 ± 2.3 | 81.0 ± 3.5 | |
| Inosine | | 89.2 ± 0.4 | 75.5 ± 1.4 | 67.3 ± 2.0 | |
| Inosine 5'-monophosphate | | 85.2 ± 3.4 | 89.8 ± 1.7 | 76.5 ± 2.2 | |
| Inositol | | 88.0 ± 2.6 | 92.2 ± 1.0 | 81.1 ± 2.0 | |
| Isocitric Acid | | 99.2 ± 1.6 | 76.8 ± 3.3 | 76.2 ± 3.0 | |
| Isoleucine | | 91.3 ± 1.3 | 84.4 ± 2.6 | 77.1 ± 0.5 | |
| Itaconic acid | | 100.8 ± 0.5 | 89.3 ± 1.3 | 90.0 ± 1.9 | |
| ITP | | 103.9 ± 2.0 | 85.2 ± 2.3 | 88.5 ± 1.6 | |
| Kynurenic acid | | 96.5 ± 2.2 | 79.1 ± 1.6 | 76.3 ± 3.3 | |
| Kynurenine | | 86.6 ± 1.3 | 75.4 ± 2.9 | 65.3 ± 2.1 | |
| Lactic Acid | | 88.4 ± 2.6 | 79.1 ± 0.6 | 69.9 ± 2.5 | |
| Lactose | | 103.5 ± 2.9 | 79.7 ± 2.0 | 82.5 ± 0.4 | |
| Leucine | | 90.4 ± 1.4 | 77.3 ± 1.7 | 69.9 ± 1.2 | |
| Levulinic acid | | 90.1 ± 0.9 | 76.8 ± 3.5 | 69.2 ± 1.8 | |
| Lysine | | 88.6 ± 2.9 | 88.6 ± 1.3 | 78.5 ± 3.5 | |
| Malic Acid | | 93.7 ± 1.5 | 81.8 ± 2.7 | 76.6 ± 2.7 | |
| Malonic acid | | 102.6 ± 0.5 | 80.6 ± 1.9 | 82.7 ± 2.7 | |
| Mannose | | 100.7 ± 3.0 | 78.1 ± 2.6 | 78.6 ± 1.2 | |
| Mannose-1 with Mannose-6-phosphate | | 93.2 ± 0.5 | 89.1 ± 1.7 | 83.0 ± 2.9 | |
| Mesaconic acid | | 101.6 ± 1.3 | 83.2 ± 1.9 | 84.5 ± 1.5 | |
| Methionine | | 97.1 ± 3.4 | 80.1 ± 2.5 | 77.8 ± 3.3 | |
| Methionine sulfone | | 91.6 ± 3.4 | 80.2 ± 1.9 | 73.5 ± 0.4 | |
| Methionine sulfoxide | | 96.7 ± 2.1 | 83.0 ± 2.2 | 80.3 ± 1.5 | |
| Methylmalonate | | 97.8 ± 1.1 | 78.1 ± 2.4 | 76.4 ± 1.5 | |
| N-Acetyl-Alanine | | 98.4 ± 0.4 | 86.5 ± 3.2 | 85.1 ± 1.6 | |
| N-Acetyl-Aspartic Acid | | 98.5 ± 0.7 | 80.2 ± 0.7 | 79.0 ± 3.5 | |
| N-Acetyl-Citrulline | | 87.4 ± 1.9 | 90.8 ± 2.4 | 79.4 ± 3.4 | |
| N-Acetyl-Galactosamine | | 88.4 ± 3.0 | 77.9 ± 2.4 | 68.9 ± 2.8 | |
| N-Acetyl-Glucosamine | | 94.3 ± 1.0 | 88.0 ± 2.7 | 83.0 ± 3.0 | |
| N-Acetyl-Glutamine | | 94.1 ± 3.6 | 91.9 ± 2.5 | 86.5 ± 0.7 | |
| N-Acetyl-Glycine | | 94.4 ± 1.7 | 78.8 ± 0.7 | 74.4 ± 3.3 | |
| N-Acetyl-Leucine | | 88.3 ± 1.6 | 84.5 ± 0.9 | 74.6 ± 1.3 | |
| N-Acetyl-Methionine | | 89.1 ± 2.2 | 90.5 ± 1.1 | 80.6 ± 2.3 | |
| N-Acetylneuraminic acid | | 94.3 ± 3.1 | 82.3 ± 0.3 | 77.6 ± 3.0 | |
| N-Acetyl-Phenylalanine | | 86.1 ± 2.8 | 80.7 ± 1.1 | 69.5 ± 0.6 | |
| N-Acetyl-Proline | | 103.7 ± 0.6 | 80.5 ± 2.8 | 83.5 ± 1.8 | |
| N-Acetyl-Serine | | 88.2 ± 2.7 | 85.9 ± 1.4 | 75.8 ± 1.4 | |
| N-Acetyl-Serotonin | | 93.9 ± 1.2 | 90.4 ± 2.6 | 84.9 ± 3.5 | |
| N-Acetyl-Tryptophan | | 104.3 ± 0.9 | 90.7 ± 1.4 | 94.6 ± 1.2 | |
| N-Acetyl-Tyrosine | | 104.5 ± 3.6 | 75.8 ± 2.1 | 79.2 ± 0.4 | |
| NAD | | 100.6 ± 0.6 | 89.2 ± 1.3 | 89.7 ± 0.8 | |
| NADH | | 93.8 ± 1.2 | 91.6 ± 0.9 | 85.9 ± 2.0 | |
| NADP | | 91.5 ± 1.6 | 76.4 ± 0.7 | 69.9 ± 2.0 | |
| NADPH | | 94.5 ± 3.1 | 90.1 ± 3.4 | 85.1 ± 1.4 | |
| N-alpha-Acetyl-lysine | | 104.0 ± 1.8 | 86.5 ± 1.5 | 90.0 ± 2.8 | |
| N-alpha-Acetyl-Ornithine | | 93.9 ± 1.1 | 91.2 ± 2.5 | 85.6 ± 2.1 | |
| N-epsilon-Acetyl-lysine | | 94.8 ± 1.3 | 89.0 ± 2.5 | 84.4 ± 1.0 | |
| N-epsilon-Acetyl-Ornithine | | 86.1 ± 1.9 | 82.6 ± 0.8 | 71.1 ± 0.8 | |
| Nicotinamide | | 104.8 ± 0.7 | 75.2 ± 1.2 | 78.8 ± 1.0 | |
| Nicotinic acid | | 101.2 ± 1.7 | 93.2 ± 3.4 | 94.3 ± 1.0 | |
| N-Methyl-Aspartic Acid | | 88.0 ± 3.5 | 91.4 ± 0.9 | 80.4 ± 2.0 | |
| O-Acetyl-Serine | | 102.3 ± 1.6 | 85.2 ± 3.4 | 87.2 ± 1.4 | |
| O-Phosphoethanolamine | | 89.9 ± 2.4 | 81.9 ± 0.5 | 73.6 ± 3.1 | |
| O-Phospho-Serine | | 86.5 ± 2.5 | 78.1 ± 3.2 | 67.6 ± 0.3 | |
| Ophthalmic acid | | 93.3 ± 2.4 | 76.4 ± 2.4 | 71.3 ± 2.2 | |
| Ornithine | | 87.0 ± 3.5 | 80.2 ± 2.7 | 69.8 ± 2.2 | |
| Orotic acid | | 98.9 ± 0.5 | 85.6 ± 1.3 | 84.7 ± 3.3 | |
| Oxaloacetate | | 89.6 ± 0.6 | 83.3 ± 2.2 | 74.6 ± 1.0 | |
| Pantothenic acid | | 90.7 ± 3.0 | 82.6 ± 1.6 | 74.9 ± 1.1 | |
| p-Coumaric Acid | | 94.2 ± 3.1 | 87.9 ± 3.3 | 82.8 ± 2.0 | |
| Phenylalanine | | 88.5 ± 2.8 | 89.7 ± 3.1 | 79.4 ± 1.7 | |
| Phenylpyruvate | | 95.1 ± 2.2 | 85.6 ± 0.4 | 81.4 ± 1.1 | |
| Phosphocreatine | | 104.9 ± 1.5 | 85.6 ± 1.5 | 89.8 ± 0.7 | |
| Phosphoenolpyruvate | | 86.5 ± 0.7 | 93.3 ± 0.8 | 80.7 ± 1.2 | |
| Phosphoribosylpyrophosphate | | 98.5 ± 2.9 | 75.9 ± 1.0 | 74.8 ± 2.6 | |
| Picolinate | | 98.7 ± 0.9 | 84.8 ± 0.6 | 83.7 ± 1.9 | |
| Pipecolic acid | | 92.5 ± 2.2 | 87.3 ± 1.5 | 80.8 ± 3.4 | |
| Proline | | 91.1 ± 1.1 | 88.4 ± 1.9 | 80.5 ± 2.2 | |
| Pyridoxal | | 92.3 ± 0.3 | 83.7 ± 2.3 | 77.3 ± 2.5 | |
| Pyridoxal-5-phosphate | | 87.5 ± 1.7 | 81.1 ± 3.2 | 71.0 ± 3.0 | |
| Pyridoxamine | | 86.9 ± 0.6 | 82.7 ± 1.4 | 71.9 ± 2.5 | |
| Pyridoxine | | 96.6 ± 2.7 | 75.5 ± 1.6 | 72.9 ± 2.9 | |
| Pyroglutamic acid | | 87.5 ± 1.4 | 93.5 ± 1.6 | 81.8 ± 2.3 | |
| Pyrrolysine | | 85.4 ± 2.4 | 88.9 ± 3.4 | 75.9 ± 0.4 | |
| Pyruvic Acid | | 91.3 ± 1.8 | 80.9 ± 1.1 | 73.9 ± 1.3 | |
| Riboflavin | | 100.6 ± 0.5 | 77.9 ± 2.6 | 78.4 ± 2.3 | |
| Ribose | | 100.8 ± 1.4 | 78.6 ± 2.7 | 79.2 ± 0.7 | |
| Ribose-5-phosphate | | 103.3 ± 1.6 | 79.1 ± 1.9 | 81.7 ± 1.0 | |
| Ribulose-1-phosphate | | 91.3 ± 2.8 | 93.8 ± 1.9 | 85.6 ± 1.4 | |
| Ribulose-5-phosphate | | 100.7 ± 2.7 | 89.5 ± 0.4 | 90.1 ± 1.8 | |
| S-Adenosylmethionine | | 99.2 ± 2.3 | 93.7 ± 2.9 | 93.0 ± 1.1 | |
| Sarcosine | | 87.5 ± 2.3 | 94.7 ± 1.3 | 82.9 ± 1.8 | |
| Sedoheptulose-7-phosphate | | 87.3 ± 2.3 | 82.4 ± 1.4 | 71.9 ± 2.9 | |
| Selenocystine | | 103.2 ± 0.5 | 89.1 ± 1.7 | 92.0 ± 3.0 | |
| Selenomethionine | | 91.4 ± 1.5 | 94.0 ± 3.3 | 85.9 ± 1.1 | |
| Serine | | 95.4 ± 2.0 | 77.1 ± 1.9 | 73.6 ± 2.4 | |
| Serotonin | | 94.4 ± 0.4 | 89.5 ± 3.3 | 84.5 ± 1.0 | |
| Sorbitol | | 104.3 ± 0.5 | 76.8 ± 0.8 | 80.1 ± 2.6 | |
| Spermidine | | 100.8 ± 3.5 | 75.4 ± 1.2 | 76.0 ± 2.2 | |
| Spermine | | 103.6 ± 0.8 | 76.0 ± 1.4 | 78.7 ± 1.2 | |
| Succinic Acid | | 88.3 ± 3.5 | 78.5 ± 1.9 | 69.3 ± 3.4 | |
| Sucrose | | 99.9 ± 2.8 | 79.4 ± 2.5 | 79.3 ± 2.7 | |
| Symmetric dimethylarginine | | 97.2 ± 1.6 | 85.7 ± 1.7 | 83.3 ± 1.0 | |
| Taurine | | 93.0 ± 2.2 | 93.8 ± 1.9 | 87.2 ± 1.9 | |
| Thiamine | | 105.0 ± 2.1 | 88.0 ± 3.5 | 92.4 ± 3.6 | |
| Thiamine monophosphate | | 97.8 ± 2.6 | 93.6 ± 1.0 | 91.5 ± 3.4 | |
| Thiamine pyrophosphate | | 88.7 ± 3.5 | 79.6 ± 2.8 | 70.6 ± 0.7 | |
| Threonic acid | | 91.1 ± 2.6 | 83.9 ± 3.6 | 76.4 ± 3.5 | |
| Threonine | | 94.6 ± 0.9 | 90.0 ± 0.4 | 85.1 ± 1.2 | |
| Thymidine | | 99.0 ± 0.5 | 84.6 ± 2.2 | 83.8 ± 0.7 | |
| Thymidine Monophosphate | | 97.2 ± 2.3 | 89.3 ± 1.4 | 86.8 ± 3.5 | |
| Thymine | | 88.7 ± 0.7 | 88.1 ± 1.7 | 78.1 ± 1.7 | |
| Trimethylamine N-oxide | | 93.0 ± 1.6 | 75.8 ± 1.0 | 70.5 ± 0.9 | |
| Tryptophan | | 90.2 ± 1.4 | 76.6 ± 1.1 | 69.1 ± 3.4 | |
| Tyrosine | | 85.7 ± 1.1 | 93.7 ± 2.7 | 80.3 ± 0.8 | |
| Uracil | | 100.3 ± 0.6 | 87.8 ± 1.8 | 88.1 ± 1.3 | |
| Uric Acid | | 85.1 ± 3.3 | 90.1 ± 3.2 | 76.7 ± 0.9 | |
| Uridine | | 87.3 ± 0.9 | 91.0 ± 2.7 | 79.4 ± 2.4 | |
| Uridine 5'-diphosphate | | 96.2 ± 3.1 | 75.9 ± 0.6 | 73.0 ± 0.8 | |
| Uridine Monophosphate | | 97.1 ± 1.3 | 93.9 ± 2.5 | 91.2 ± 1.0 | |
| Uridine-5-Diphosphogalactose | | 104.0 ± 1.7 | 75.6 ± 0.8 | 78.6 ± 1.6 | |
| Uridine-5-Diphosphoglucose | | 101.2 ± 1.6 | 78.2 ± 3.0 | 79.1 ± 1.1 | |
| Urocanic acid | | 99.8 ± 3.3 | 92.2 ± 2.7 | 92.0 ± 2.1 | |
| UTP | | 101.4 ± 2.8 | 85.7 ± 1.3 | 86.9 ± 0.9 | |
| Valine | | 85.7 ± 2.5 | 82.6 ± 1.8 | 70.8 ± 1.0 | |
| Xanthine | | 91.6 ± 0.9 | 87.3 ± 3.3 | 80.0 ± 2.2 | |
| Xanthosine | | 89.5 ± 1.2 | 86.5 ± 3.4 | 77.4 ± 1.9 | |
| Xanthurenic acid | | 102.4 ± 1.8 | 78.4 ± 0.4 | 80.3 ± 1.4 | |
| XMP | | 104.0 ± 3.3 | 82.5 ± 0.8 | 85.8 ± 3.4 | |
| Xylulose-5-phosphate | | 104.4 ± 1.9 | 78.2 ± 0.4 | 81.6 ± 2.3 | |

### **Table S4** | Matrix effect (ME), Process Efficiency (PE) and Extraction Recovery (RE) of metabolites without internal standard correction (Plasma extract as matrix, n = 5). The concentration of the QC sample used was 100 ng/mL.

| Metabolite | ME [%] | | | RE [%] | | | | | PE [%] |
| --- | --- | --- | --- | --- | --- | --- | --- | --- | --- |
| 1-Methylhistidine | | | 93.0 ± 0.8 | | | 94.7 ± 1.3 | | 88.1 ± 3.2 | |
| 2,3-Bisphosphoglycerate | | | 94.1 ± 2.1 | | | 92.5 ± 3.3 | | 87.0 ± 2.3 | |
| 2-Aminoadipic Acid | | | 99.7 ± 1.4 | | | 87.4 ± 2.6 | | 87.1 ± 0.8 | |
| 2-Deoxyadenosine | | | 101.9 ± 1.8 | | | 87.2 ± 0.9 | | 88.9 ± 1.3 | |
| 2'-Deoxyadenosine 5'-monophosphate | | | 92.0 ± 2.6 | | | 83.0 ± 1.7 | | 76.4 ± 2.4 | |
| 2'-deoxyadenosine 5'-triphosphate | | | 92.4 ± 3.4 | | | 88.2 ± 1.7 | | 81.5 ± 2.9 | |
| 2-Deoxycytidine | | | 90.4 ± 3.6 | | | 78.6 ± 1.9 | | 71.1 ± 2.5 | |
| 2'-Deoxycytidine-5'-monophosphate | | | 98.1 ± 0.8 | | | 89.5 ± 3.5 | | 87.8 ± 2.0 | |
| 2-Deoxyguanosine | | | 103.5 ± 0.5 | | | 79.6 ± 2.8 | | 82.4 ± 2.7 | |
| 2-Deoxyinosine | | | 98.0 ± 0.5 | | | 83.1 ± 2.2 | | 81.4 ± 1.0 | |
| 2-hydroxybutyric acid | | | 101.2 ± 1.9 | | | 81.5 ± 0.9 | | 82.5 ± 2.9 | |
| 2-hydroxyglutaric acid | | | 95.9 ± 2.8 | | | 75.5 ± 1.0 | | 72.4 ± 0.3 | |
| 2-isopropylmalic acid | | | 101.6 ± 2.2 | | | 83.3 ± 1.8 | | 84.6 ± 3.6 | |
| 2-ketobutyric acid | | | 97.3 ± 1.2 | | | 92.9 ± 0.7 | | 90.4 ± 1.4 | |
| 2-oxoadipic acid | | | 101.6 ± 3.0 | | | 93.2 ± 3.4 | | 94.7 ± 2.6 | |
| 2-phosphoglycerate | | | 89.4 ± 1.8 | | | 76.9 ± 1.6 | | 68.7 ± 3.0 | |
| 3-amino-isobutyric acid | | | 97.3 ± 1.6 | | | 87.1 ± 2.7 | | 84.7 ± 1.9 | |
| 3-hydroxy-3-methyl-2-oxobutanoate | | | 91.2 ± 1.4 | | | 86.9 ± 0.3 | | 79.3 ± 0.6 | |
| 3-hydroxybutyric acid | | | 96.8 ± 2.5 | | | 78.3 ± 0.6 | | 75.8 ± 3.4 | |
| 3-Hydroxy-Proline | | | 100.2 ± 3.3 | | | 90.0 ± 3.3 | | 90.2 ± 2.7 | |
| 3-isopropylmalic acid | | | 101.6 ± 3.3 | | | 91.4 ± 0.4 | | 92.9 ± 1.2 | |
| 3-methyl-2-oxopentanoate | | | 94.0 ± 1.6 | | | 77.9 ± 2.0 | | 73.2 ± 1.1 | |
| 3-Methylhistidine | | | 100.5 ± 1.5 | | | 93.3 ± 1.2 | | 93.8 ± 2.8 | |
| 3-Phosphoglyceraldehyde | | | 88.7 ± 2.1 | | | 83.7 ± 2.6 | | 74.2 ± 3.1 | |
| 3-phosphoglycerate | | | 98.7 ± 3.0 | | | 94.3 ± 1.6 | | 93.1 ± 3.4 | |
| 4-amino-butyric acid | | | 94.1 ± 3.1 | | | 76.5 ± 2.6 | | 72.0 ± 1.8 | |
| 4-Hydroxybenzoic acid | | | 91.3 ± 0.5 | | | 84.1 ± 3.5 | | 76.8 ± 1.6 | |
| 4-Hydroxyglutamate | | | 100.3 ± 2.8 | | | 87.9 ± 1.9 | | 88.2 ± 1.9 | |
| 4-Hydroxy-Proline | | | 103.6 ± 3.3 | | | 87.7 ± 2.9 | | 90.9 ± 2.5 | |
| 5-Aminolevulinic acid | | | 103.0 ± 3.3 | | | 86.7 ± 1.7 | | 89.3 ± 0.4 | |
| 5-Aminovaleric acid | | | 102.4 ± 1.1 | | | 84.5 ± 2.1 | | 86.5 ± 1.4 | |
| 5-Hydroxy-Tryptophan | | | 95.9 ± 1.9 | | | 94.1 ± 2.7 | | 90.2 ± 1.7 | |
| 5-Methoxytryptophan | | | 102.5 ± 1.6 | | | 76.1 ± 0.4 | | 78.0 ± 1.4 | |
| 5-Methylcytosine | | | 97.6 ± 3.5 | | | 83.2 ± 1.0 | | 81.2 ± 2.9 | |
| 6-phosphogluconate | | | 98.4 ± 0.6 | | | 93.6 ± 2.8 | | 92.1 ± 0.8 | |
| Acetoacetate | | | 89.3 ± 3.5 | | | 81.2 ± 1.8 | | 72.5 ± 3.0 | |
| Acetylcarnitine | | | 88.8 ± 2.3 | | | 78.4 ± 1.7 | | 69.6 ± 0.6 | |
| Acetyl-CoA | | | 93.8 ± 0.4 | | | 77.4 ± 1.2 | | 72.6 ± 3.1 | |
| Aconitic Acid | | | 96.4 ± 1.8 | | | 89.6 ± 1.9 | | 86.4 ± 0.5 | |
| Adenine | | | 99.3 ± 0.4 | | | 76.2 ± 2.3 | | 75.7 ± 2.5 | |
| Adenosine | | | 101.9 ± 0.8 | | | 89.8 ± 2.3 | | 91.5 ± 3.0 | |
| Adenosine 3',5'-cyclic monophosphate | | | 97.7 ± 2.1 | | | 84.3 ± 3.3 | | 82.4 ± 3.4 | |
| Adenosine Monophosphate | | | 99.9 ± 1.9 | | | 88.9 ± 2.1 | | 88.8 ± 0.6 | |
| Adenosine-5'-diphosphate | | | 99.2 ± 2.7 | | | 76.3 ± 1.4 | | 75.7 ± 3.0 | |
| Adenosine-5'-triphosphate | | | 92.4 ± 2.7 | | | 90.4 ± 3.5 | | 83.5 ± 3.4 | |
| Adenylosuccinate | | | 100.7 ± 3.5 | | | 87.3 ± 2.6 | | 87.9 ± 2.0 | |
| Alanine | | | 92.4 ± 1.0 | | | 81.3 ± 2.8 | | 75.1 ± 0.9 | |
| Allantoin | | | 98.6 ± 2.4 | | | 83.3 ± 0.3 | | 82.1 ± 1.1 | |
| alpha-ketoglutaric acid | | | 89.1 ± 2.5 | | | 93.1 ± 2.7 | | 83.0 ± 3.1 | |
| Anserine | | | 97.2 ± 1.8 | | | 83.6 ± 0.4 | | 81.3 ± 2.3 | |
| Trigonelleine | | | 97.6 ± 0.6 | | | 90.3 ± 3.3 | | 88.1 ± 2.4 | |
| Arginine | | | 102.1 ± 1.9 | | | 85.9 ± 3.3 | | 87.7 ± 2.0 | |
| Argininosuccinic Acid | | | 102.0 ± 0.8 | | | 86.8 ± 1.3 | | 88.5 ± 1.1 | |
| Ascorbic acid | | | 102.0 ± 2.5 | | | 90.4 ± 2.7 | | 92.2 ± 1.1 | |
| Asparagine | | | 95.7 ± 2.5 | | | 83.7 ± 0.6 | | 80.1 ± 1.4 | |
| Aspartic acid | | | 89.2 ± 1.7 | | | 89.5 ± 0.6 | | 79.8 ± 2.9 | |
| Asymmetric dimethylarginine | | | 90.2 ± 2.0 | | | 88.3 ± 2.5 | | 79.6 ± 0.7 | |
| Azelaic acid | | | 92.2 ± 0.6 | | | 77.9 ± 2.2 | | 71.8 ± 1.0 | |
| beta-Alanine | | | 99.1 ± 0.5 | | | 79.2 ± 2.3 | | 78.5 ± 2.0 | |
| Betaine | | | 88.8 ± 0.7 | | | 89.3 ± 1.3 | | 79.3 ± 2.5 | |
| beta-Ureidopropionate | | | 89.4 ± 2.4 | | | 86.5 ± 3.0 | | 77.3 ± 0.9 | |
| Biotin | | | 103.1 ± 1.4 | | | 89.9 ± 2.6 | | 92.7 ± 1.5 | |
| Cadaverine | | | 101.2 ± 2.4 | | | 76.5 ± 1.5 | | 77.4 ± 1.0 | |
| Carnitine | | | 102.3 ± 1.9 | | | 86.6 ± 1.8 | | 88.6 ± 1.8 | |
| Carnosine | | | 97.5 ± 3.3 | | | 75.4 ± 0.5 | | 73.5 ± 0.7 | |
| cGMP | | | 101.7 ± 1.6 | | | 80.2 ± 0.7 | | 81.6 ± 2.1 | |
| Choline | | | 103.5 ± 3.3 | | | 79.5 ± 2.7 | | 82.3 ± 1.1 | |
| Cinnamic acid | | | 92.7 ± 0.7 | | | 82.1 ± 1.5 | | 76.1 ± 2.6 | |
| Citraconic acid | | | 94.3 ± 2.3 | | | 87.3 ± 3.0 | | 82.3 ± 2.1 | |
| Citramalic acid | | | 91.7 ± 1.6 | | | 92.2 ± 1.9 | | 84.5 ± 2.3 | |
| Citric Acid | | | 89.9 ± 2.4 | | | 86.3 ± 3.4 | | 77.6 ± 2.4 | |
| Citrulline | | | 101.2 ± 2.5 | | | 77.3 ± 3.0 | | 78.2 ± 1.8 | |
| Creatine | | | 98.1 ± 1.7 | | | 92.4 ± 1.2 | | 90.6 ± 3.2 | |
| Creatinine | | | 90.6 ± 0.7 | | | 85.2 ± 1.8 | | 77.2 ± 2.3 | |
| Cystathionine | | | 98.3 ± 2.0 | | | 78.5 ± 0.5 | | 77.2 ± 1.4 | |
| Cysteamine | | | 103.5 ± 2.1 | | | 78.3 ± 2.4 | | 81.0 ± 2.2 | |
| Cysteine | | | 95.9 ± 3.5 | | | 86.0 ± 0.6 | | 82.5 ± 1.3 | |
| Cysteinylglycine | | | 90.7 ± 2.1 | | | 81.5 ± 1.6 | | 73.9 ± 1.8 | |
| Cystine | | | 90.2 ± 2.1 | | | 83.1 ± 2.7 | | 75.0 ± 2.1 | |
| Cytidine | | | 90.5 ± 3.2 | | | 92.0 ± 2.7 | | 83.3 ± 0.5 | |
| Cytidine 5'-diphosphate | | | 91.3 ± 0.4 | | | 83.8 ± 3.0 | | 76.5 ± 0.9 | |
| Cytidine 5'-triphosphate | | | 94.8 ± 0.9 | | | 77.3 ± 3.4 | | 73.3 ± 2.5 | |
| Cytidine monophosphate | | | 92.9 ± 0.4 | | | 87.7 ± 0.5 | | 81.5 ± 0.7 | |
| Cytosine | | | 98.3 ± 2.4 | | | 88.8 ± 1.5 | | 87.3 ± 3.3 | |
| Dihydroxyacetonephosphate | | | 90.4 ± 3.5 | | | 84.0 ± 0.9 | | 75.9 ± 1.0 | |
| Dimethylglycine | | | 96.5 ± 2.2 | | | 77.2 ± 3.3 | | 74.5 ± 2.6 | |
| Erythrose-4-phosphate | | | 103.5 ± 0.8 | | | 80.0 ± 1.5 | | 82.8 ± 3.0 | |
| Ethanolamine | | | 97.8 ± 2.1 | | | 78.4 ± 2.4 | | 76.7 ± 0.3 | |
| FAD | | | 93.7 ± 1.0 | | | 87.3 ± 3.3 | | 81.8 ± 2.5 | |
| Folic acid | | | 100.4 ± 1.2 | | | 86.9 ± 2.1 | | 87.2 ± 0.7 | |
| Fructose | | | 100.6 ± 0.9 | | | 88.5 ± 3.3 | | 89.0 ± 2.3 | |
| Fructose-1,6-bisphosphate | | | 101.3 ± 2.0 | | | 75.6 ± 0.8 | | 76.6 ± 3.3 | |
| Fructose-6-Phosphate | | | 92.5 ± 3.5 | | | 75.1 ± 0.6 | | 69.5 ± 3.6 | |
| Fumaric Acid | | | 95.9 ± 1.7 | | | 87.7 ± 2.6 | | 84.1 ± 3.3 | |
| Galactosamine | | | 91.0 ± 0.3 | | | 88.4 ± 0.5 | | 80.4 ± 3.2 | |
| Galactose-1-phosphate | | | 94.2 ± 1.2 | | | 87.5 ± 1.9 | | 82.4 ± 1.9 | |
| Galactose-6-phosphate | | | 99.4 ± 3.6 | | | 88.4 ± 2.8 | | 87.9 ± 1.8 | |
| gamma-Glutamylcysteine | | | 94.2 ± 3.3 | | | 83.1 ± 3.0 | | 78.3 ± 2.5 | |
| Glucaric Acid | | | 99.2 ± 0.6 | | | 84.1 ± 2.1 | | 83.4 ± 3.5 | |
| Gluconic acid | | | 99.3 ± 3.2 | | | 81.8 ± 1.0 | | 81.2 ± 1.8 | |
| Gluconolactone | | | 102.4 ± 2.4 | | | 92.9 ± 3.0 | | 95.1 ± 0.7 | |
| Glucosamine | | | 94.6 ± 0.5 | | | 85.6 ± 0.9 | | 81.0 ± 1.4 | |
| Glucosamine-6-Phosphate | | | 96.5 ± 0.7 | | | 85.1 ± 2.5 | | 82.1 ± 1.4 | |
| Glucose | | | 89.4 ± 2.5 | | | 76.8 ± 2.8 | | 68.7 ± 0.9 | |
| Glucose-1-phosphate | | | 89.6 ± 3.2 | | | 78.6 ± 2.7 | | 70.4 ± 0.7 | |
| Glucose-6-phosphate | | | 98.2 ± 3.4 | | | 82.2 ± 3.3 | | 80.7 ± 1.1 | |
| Glucuronolactone | | | 96.7 ± 3.5 | | | 78.3 ± 1.4 | | 75.7 ± 3.1 | |
| Glutamic acid | | | 100.9 ± 0.9 | | | 94.0 ± 1.9 | | 94.8 ± 0.4 | |
| Glutamine | | | 99.3 ± 3.5 | | | 85.2 ± 0.7 | | 84.6 ± 2.1 | |
| Glutathione (GSSG) | | | 103.2 ± 1.3 | | | 87.5 ± 0.6 | | 90.3 ± 0.9 | |
| Glutathione | | | 103.6 ± 0.6 | | | 76.9 ± 2.2 | | 79.7 ± 3.5 | |
| Glycerophosphocholine | | | 103.1 ± 1.0 | | | 90.8 ± 1.2 | | 93.6 ± 1.8 | |
| Glycine | | | 96.6 ± 1.0 | | | 92.9 ± 3.4 | | 89.7 ± 2.0 | |
| Glycolic acid | | | 102.9 ± 3.3 | | | 88.3 ± 2.0 | | 90.9 ± 1.1 | |
| Guanine | | | 90.5 ± 1.9 | | | 86.4 ± 3.3 | | 78.2 ± 2.1 | |
| Guanosine | | | 90.8 ± 1.6 | | | 86.6 ± 1.5 | | 78.6 ± 2.9 | |
| Guanosine 5'-diphosphate | | | 100.2 ± 1.0 | | | 88.2 ± 2.8 | | 88.4 ± 2.7 | |
| Guanosine-5'-monophosphate | | | 95.5 ± 3.0 | | | 92.8 ± 2.5 | | 88.6 ± 3.2 | |
| Hippuric acid | | | 100.3 ± 3.3 | | | 90.8 ± 0.3 | | 91.1 ± 2.5 | |
| Histamine | | | 89.4 ± 2.9 | | | 77.6 ± 2.3 | | 69.4 ± 3.2 | |
| Histidine | | | 103.7 ± 3.4 | | | 84.1 ± 2.3 | | 87.2 ± 2.9 | |
| Histidinol | | | 99.8 ± 2.8 | | | 87.9 ± 0.6 | | 87.7 ± 0.5 | |
| Homoalanine | | | 96.8 ± 3.0 | | | 77.2 ± 2.1 | | 74.7 ± 2.1 | |
| Homocitrulline | | | 102.1 ± 2.6 | | | 77.7 ± 3.0 | | 79.3 ± 3.2 | |
| Homocysteine | | | 99.2 ± 0.9 | | | 82.8 ± 2.6 | | 82.1 ± 1.7 | |
| Homocystine | | | 101.3 ± 2.4 | | | 82.9 ± 0.6 | | 84.0 ± 0.6 | |
| Homogentisic acid | | | 98.5 ± 1.6 | | | 79.4 ± 2.2 | | 78.2 ± 1.1 | |
| Homoleucine | | | 95.8 ± 2.6 | | | 87.6 ± 1.7 | | 83.9 ± 0.6 | |
| Homophenylalanine | | | 99.6 ± 1.4 | | | 86.4 ± 3.0 | | 86.1 ± 0.7 | |
| Homoserine | | | 97.3 ± 0.6 | | | 86.1 ± 0.9 | | 83.8 ± 2.4 | |
| Homovanillic acid | | | 90.6 ± 2.2 | | | 82.6 ± 2.6 | | 74.8 ± 2.3 | |
| Hypotaurine | | | 102.2 ± 0.3 | | | 89.8 ± 3.6 | | 91.8 ± 0.5 | |
| Hypoxanthine | | | 92.0 ± 1.9 | | | 77.5 ± 1.7 | | 71.3 ± 2.8 | |
| Inosine | | | 99.4 ± 1.4 | | | 83.9 ± 1.0 | | 83.4 ± 2.2 | |
| Inosine 5'-monophosphate | | | 98.0 ± 3.6 | | | 85.1 ± 0.4 | | 83.4 ± 2.6 | |
| Inositol | | | 103.3 ± 1.7 | | | 78.9 ± 0.7 | | 81.5 ± 1.3 | |
| Isocitric Acid | | | 101.2 ± 1.3 | | | 87.1 ± 3.6 | | 88.2 ± 1.4 | |
| Isoleucine | | | 89.9 ± 3.4 | | | 92.2 ± 2.0 | | 82.9 ± 1.3 | |
| Itaconic acid | | | 93.7 ± 2.9 | | | 90.4 ± 3.0 | | 84.7 ± 1.0 | |
| ITP | | | 91.5 ± 0.5 | | | 85.4 ± 1.5 | | 78.1 ± 2.3 | |
| Kynurenic acid | | | 93.3 ± 2.1 | | | 78.6 ± 2.1 | | 73.3 ± 0.9 | |
| Kynurenine | | | 98.4 ± 2.2 | | | 77.0 ± 1.7 | | 75.8 ± 1.4 | |
| Lactic Acid | | | 101.2 ± 1.0 | | | 92.9 ± 3.2 | | 94.0 ± 0.9 | |
| Lactose | | | 94.7 ± 0.4 | | | 79.2 ± 0.5 | | 75.0 ± 1.4 | |
| Leucine | | | 97.5 ± 0.9 | | | 91.6 ± 1.2 | | 89.3 ± 2.3 | |
| Levulinic acid | | | 100.5 ± 1.3 | | | 86.8 ± 2.2 | | 87.2 ± 1.0 | |
| Lysine | | | 97.1 ± 2.4 | | | 86.8 ± 3.5 | | 84.3 ± 0.6 | |
| Malic Acid | | | 99.4 ± 3.5 | | | 88.9 ± 3.0 | | 88.4 ± 1.7 | |
| Malonic acid | | | 95.9 ± 1.1 | | | 89.3 ± 1.0 | | 85.7 ± 1.0 | |
| Mannose | | | 94.4 ± 0.5 | | | 83.4 ± 2.8 | | 78.7 ± 0.6 | |
| Mannose-1 with Mannose-6-phosphate | | | 89.5 ± 1.4 | | | 78.3 ± 2.4 | | 70.1 ± 2.9 | |
| Mesaconic acid | | | 100.8 ± 0.4 | | | 81.2 ± 2.9 | | 81.9 ± 1.0 | |
| Methionine | | | 91.4 ± 3.2 | | | 83.9 ± 0.9 | | 76.7 ± 1.0 | |
| Methionine sulfone | | | 103.1 ± 0.9 | | | 93.7 ± 2.9 | | 96.6 ± 2.0 | |
| Methionine sulfoxide | | | 103.3 ± 1.2 | | | 78.6 ± 2.8 | | 81.2 ± 2.4 | |
| Methylmalonate | | | 92.5 ± 1.1 | | | 77.4 ± 2.6 | | 71.6 ± 2.5 | |
| N-Acetyl-Alanine | | | 102.8 ± 2.0 | | | 82.2 ± 2.3 | | 84.5 ± 3.0 | |
| N-Acetyl-Aspartic Acid | | | 96.6 ± 2.4 | | | 90.5 ± 3.1 | | 87.5 ± 1.6 | |
| N-Acetyl-Citrulline | | | 102.6 ± 2.4 | | | 84.3 ± 3.5 | | 86.5 ± 0.7 | |
| N-Acetyl-Galactosamine | | | 91.2 ± 2.6 | | | 90.1 ± 3.1 | | 82.2 ± 2.9 | |
| N-Acetyl-Glucosamine | | | 98.3 ± 2.3 | | | 76.5 ± 2.9 | | 75.2 ± 1.6 | |
| N-Acetyl-Glutamine | | | 103.0 ± 1.5 | | | 92.3 ± 0.9 | | 95.1 ± 1.3 | |
| N-Acetyl-Glycine | | | 91.8 ± 1.0 | | | 87.3 ± 3.1 | | 80.2 ± 1.7 | |
| N-Acetyl-Leucine | | | 94.6 ± 2.1 | | | 77.1 ± 1.7 | | 72.9 ± 1.6 | |
| N-Acetyl-Methionine | | | 98.7 ± 2.7 | | | 84.2 ± 2.5 | | 83.1 ± 0.5 | |
| N-Acetylneuraminic acid | | | 99.8 ± 3.0 | | | 85.0 ± 2.8 | | 84.8 ± 2.1 | |
| N-Acetyl-Phenylalanine | | | 99.8 ± 2.9 | | | 90.5 ± 0.8 | | 90.3 ± 1.1 | |
| N-Acetyl-Proline | | | 99.4 ± 1.6 | | | 81.5 ± 0.5 | | 81.0 ± 1.3 | |
| N-Acetyl-Serine | | | 93.0 ± 1.8 | | | 76.8 ± 2.5 | | 71.4 ± 1.5 | |
| N-Acetyl-Serotonin | | | 99.1 ± 0.8 | | | 89.3 ± 0.5 | | 88.5 ± 2.2 | |
| N-Acetyl-Tryptophan | | | 90.9 ± 0.5 | | | 82.8 ± 1.1 | | 75.3 ± 1.8 | |
| N-Acetyl-Tyrosine | | | 91.4 ± 2.3 | | | 94.5 ± 0.3 | | 86.4 ± 1.1 | |
| NAD | | | 90.0 ± 2.3 | | | 83.2 ± 0.7 | | 74.8 ± 2.7 | |
| NADH | | | 101.6 ± 3.5 | | | 79.6 ± 2.3 | | 80.9 ± 0.3 | |
| NADP | | | 93.6 ± 0.9 | | | 80.2 ± 1.7 | | 75.0 ± 2.5 | |
| NADPH | | | 101.0 ± 1.4 | | | 80.5 ± 1.5 | | 81.3 ± 1.8 | |
| N-alpha-Acetyl-lysine | | | 102.6 ± 3.1 | | | 83.8 ± 0.8 | | 85.9 ± 1.8 | |
| N-alpha-Acetyl-Ornithine | | | 102.2 ± 2.7 | | | 76.2 ± 0.7 | | 77.9 ± 3.0 | |
| N-epsilon-Acetyl-lysine | | | 90.1 ± 0.4 | | | 91.7 ± 1.2 | | 82.7 ± 2.5 | |
| N-epsilon-Acetyl-Ornithine | | | 92.1 ± 0.9 | | | 87.4 ± 1.6 | | 80.5 ± 2.5 | |
| Nicotinamide | | | 94.3 ± 2.6 | | | 90.3 ± 3.4 | | 85.2 ± 1.8 | |
| Nicotinic acid | | | 101.2 ± 2.3 | | | 76.0 ± 1.9 | | 76.9 ± 3.1 | |
| N-Methyl-Aspartic Acid | | | 96.4 ± 1.2 | | | 94.1 ± 0.5 | | 90.8 ± 2.6 | |
| O-Acetyl-Serine | | | 100.3 ± 0.4 | | | 86.0 ± 3.0 | | 86.2 ± 3.5 | |
| O-Phosphoethanolamine | | | 103.0 ± 2.7 | | | 80.1 ± 2.8 | | 82.5 ± 3.4 | |
| O-Phospho-Serine | | | 101.6 ± 0.9 | | | 77.7 ± 0.9 | | 78.9 ± 3.3 | |
| Ophthalmic acid | | | 90.8 ± 1.1 | | | 85.5 ± 2.9 | | 77.6 ± 2.1 | |
| Ornithine | | | 100.4 ± 0.9 | | | 82.4 ± 3.1 | | 82.7 ± 0.8 | |
| Orotic acid | | | 90.5 ± 3.2 | | | 80.3 ± 2.7 | | 72.7 ± 2.5 | |
| Oxaloacetate | | | 98.1 ± 2.2 | | | 92.1 ± 3.6 | | 90.4 ± 2.7 | |
| Pantothenic acid | | | 89.3 ± 2.4 | | | 83.3 ± 2.3 | | 74.4 ± 2.8 | |
| p-Coumaric Acid | | | 92.2 ± 0.7 | | | 76.4 ± 1.3 | | 70.5 ± 0.6 | |
| Phenylalanine | | | 94.8 ± 2.0 | | | 88.3 ± 1.8 | | 83.7 ± 3.4 | |
| Phenylpyruvate | | | 99.8 ± 1.8 | | | 93.5 ± 1.5 | | 93.3 ± 1.2 | |
| Phosphocreatine | | | 102.4 ± 2.1 | | | 82.6 ± 3.4 | | 84.6 ± 2.4 | |
| Phosphoenolpyruvate | | | 94.5 ± 1.4 | | | 83.5 ± 3.4 | | 78.9 ± 2.4 | |
| Phosphoribosylpyrophosphate | | | 102.1 ± 2.4 | | | 94.9 ± 0.5 | | 96.9 ± 2.5 | |
| Picolinate | | | 89.7 ± 1.9 | | | 93.3 ± 0.5 | | 83.6 ± 1.0 | |
| Pipecolic acid | | | 101.0 ± 1.7 | | | 80.7 ± 3.0 | | 81.5 ± 1.7 | |
| Proline | | | 91.6 ± 1.0 | | | 91.0 ± 1.9 | | 83.4 ± 3.3 | |
| Pyridoxal | | | 97.1 ± 0.5 | | | 86.8 ± 2.0 | | 84.3 ± 2.8 | |
| Pyridoxal-5-phosphate | | | 91.3 ± 0.7 | | | 77.8 ± 0.6 | | 71.0 ± 3.0 | |
| Pyridoxamine | | | 97.8 ± 0.9 | | | 82.3 ± 3.2 | | 80.5 ± 0.7 | |
| Pyridoxine | | | 101.1 ± 0.9 | | | 75.9 ± 2.7 | | 76.7 ± 1.6 | |
| Pyroglutamic acid | | | 90.1 ± 1.7 | | | 92.0 ± 2.2 | | 82.9 ± 2.1 | |
| Pyrrolysine | | | 90.5 ± 2.0 | | | 81.4 ± 1.4 | | 73.7 ± 3.1 | |
| Pyruvic Acid | | | 100.6 ± 0.6 | | | 86.4 ± 0.5 | | 86.9 ± 1.7 | |
| Riboflavin | | | 92.0 ± 1.1 | | | 93.6 ± 3.2 | | 86.1 ± 0.9 | |
| Ribose | | | 92.4 ± 1.2 | | | 89.4 ± 2.6 | | 82.6 ± 1.6 | |
| Ribose-5-phosphate | | | 96.7 ± 3.0 | | | 78.9 ± 1.8 | | 76.3 ± 0.3 | |
| Ribulose-1-phosphate | | | 91.4 ± 3.5 | | | 92.8 ± 2.9 | | 84.8 ± 1.9 | |
| Ribulose-5-phosphate | | | 99.5 ± 2.3 | | | 87.3 ± 1.4 | | 86.8 ± 0.9 | |
| S-Adenosylmethionine | | | 93.7 ± 0.6 | | | 78.8 ± 1.0 | | 73.8 ± 3.6 | |
| Sarcosine | | | 93.7 ± 1.7 | | | 89.0 ± 1.1 | | 83.4 ± 1.5 | |
| Sedoheptulose-7-phosphate | | | 103.5 ± 3.4 | | | 78.6 ± 3.2 | | 81.3 ± 1.6 | |
| Selenocystine | | | 102.7 ± 3.3 | | | 89.9 ± 0.4 | | 92.3 ± 1.7 | |
| Selenomethionine | | | 97.3 ± 0.7 | | | 86.7 ± 0.9 | | 84.3 ± 2.0 | |
| Serine | | | 94.1 ± 0.4 | | | 94.9 ± 0.6 | | 89.3 ± 1.0 | |
| Serotonin | | | 95.5 ± 0.6 | | | 75.4 ± 2.1 | | 72.0 ± 1.0 | |
| Sorbitol | | | 102.8 ± 2.6 | | | 81.5 ± 1.4 | | 83.8 ± 1.1 | |
| Spermidine | | | 90.9 ± 3.2 | | | 82.8 ± 2.9 | | 75.2 ± 2.8 | |
| Spermine | | | 95.5 ± 1.7 | | | 87.3 ± 3.2 | | 83.4 ± 0.5 | |
| Succinic Acid | | | 98.1 ± 1.1 | | | 78.4 ± 1.2 | | 76.9 ± 2.2 | |
| Sucrose | | | 90.8 ± 1.4 | | | 81.7 ± 1.6 | | 74.2 ± 1.4 | |
| Symmetric dimethylarginine | | | 98.9 ± 1.2 | | | 84.3 ± 3.1 | | 83.4 ± 1.5 | |
| Taurine | | | 89.7 ± 2.1 | | | 77.2 ± 1.9 | | 69.2 ± 0.6 | |
| Thiamine | | | 98.1 ± 2.9 | | | 80.7 ± 0.8 | | 79.2 ± 1.5 | |
| Thiamine monophosphate | | | 99.3 ± 2.9 | | | 93.8 ± 1.1 | | 93.2 ± 2.5 | |
| Thiamine pyrophosphate | | | 100.9 ± 2.1 | | | 76.4 ± 1.9 | | 77.1 ± 2.7 | |
| Threonic acid | | | 91.5 ± 2.1 | | | 90.4 ± 1.2 | | 82.7 ± 2.2 | |
| Threonine | | | 98.1 ± 0.8 | | | 93.4 ± 3.1 | | 91.6 ± 1.4 | |
| Thymidine | | | 99.9 ± 2.0 | | | 94.4 ± 1.8 | | 94.3 ± 2.9 | |
| Thymidine Monophosphate | | | 103.6 ± 2.1 | | | 92.8 ± 1.7 | | 96.2 ± 1.0 | |
| Thymine | | | 93.9 ± 1.5 | | | 83.8 ± 1.5 | | 78.6 ± 3.3 | |
| Trimethylamine N-oxide | | | 103.3 ± 0.7 | | | 80.0 ± 2.2 | | 82.6 ± 2.6 | |
| Tryptophan | | | 95.0 ± 1.8 | | | 94.6 ± 1.4 | | 89.8 ± 1.7 | |
| Tyrosine | | | 100.1 ± 3.2 | | | 85.7 ± 3.4 | | 85.8 ± 1.0 | |
| Uracil | | | 97.9 ± 2.5 | | | 80.6 ± 0.5 | | 78.9 ± 1.3 | |
| Uric Acid | | | 94.6 ± 1.9 | | | 91.6 ± 1.4 | | 86.7 ± 2.7 | |
| Uridine | | | 98.7 ± 1.5 | | | 92.3 ± 1.5 | | 91.1 ± 2.0 | |
| Uridine 5'-diphosphate | | | 97.8 ± 1.7 | | | 79.0 ± 1.8 | | 77.3 ± 1.8 | |
| Uridine Monophosphate | | | 102.4 ± 0.9 | | | 80.8 ± 2.3 | | 82.7 ± 2.2 | |
| Uridine-5-Diphosphogalactose | | | 89.6 ± 3.0 | | | 79.8 ± 2.8 | | 71.5 ± 3.6 | |
| Uridine-5-Diphosphoglucose | | | 89.3 ± 2.3 | | | 86.2 ± 2.8 | | 77.0 ± 2.8 | |
| Urocanic acid | | | 102.2 ± 0.9 | | | 93.5 ± 3.4 | | 95.5 ± 2.5 | |
| UTP | | | 90.4 ± 1.1 | | | 88.9 ± 0.5 | | 80.4 ± 3.6 | |
| Valine | | | 100.3 ± 1.0 | | | 76.3 ± 1.8 | | 76.5 ± 2.9 | |
| Xanthine | | | 99.7 ± 1.4 | | | 88.9 ± 0.4 | | 88.6 ± 0.5 | |
| Xanthosine | | | 102.7 ± 2.5 | | | 75.6 ± 1.8 | | 77.6 ± 2.5 | |
| Xanthurenic acid | | | 102.4 ± 2.0 | | | 89.9 ± 2.9 | | 92.0 ± 2.1 | |
| XMP | | | 98.0 ± 2.7 | | | 92.8 ± 1.6 | | 90.9 ± 1.7 | |
| Xylulose-5-phosphate | | | 101.7 ± 0.8 | | | 84.1 ± 1.7 | | 85.5 ± 2.1 | |

### **Table S5** | Matrix effect (ME), Process Efficiency (PE) and Extraction Recovery (RE) of metabolites without internal standard correction (Cell extract as matrix, n = 5). The concentration of the QC sample used was 100 ng/mL.

| Metabolite | ME [%] | RE [%] | PE [%] |
| --- | --- | --- | --- |
| 1-Methylhistidine | 104.4 ± 3.3 | 87.0 ± 2.9 | 90.8 ± 0.6 |
| 2,3-Bisphosphoglycerate | 91.8 ± 1.7 | 78.6 ± 2.5 | 72.2 ± 1.3 |
| 2-Aminoadipic Acid | 91.6 ± 1.6 | 83.1 ± 3.3 | 76.1 ± 3.4 |
| 2-Deoxyadenosine | 89.3 ± 2.1 | 87.1 ± 1.6 | 77.8 ± 2.0 |
| 2'-Deoxyadenosine 5'-monophosphate | 103.9 ± 0.6 | 94.8 ± 0.7 | 98.5 ± 0.5 |
| 2'-deoxyadenosine 5'-triphosphate | 91.3 ± 2.6 | 84.4 ± 1.5 | 77.1 ± 2.4 |
| 2-Deoxycytidine | 103.6 ± 2.3 | 82.6 ± 3.4 | 85.6 ± 0.3 |
| 2'-Deoxycytidine-5'-monophosphate | 101.7 ± 1.8 | 92.6 ± 1.1 | 94.2 ± 2.4 |
| 2-Deoxyguanosine | 90.3 ± 0.5 | 92.7 ± 3.3 | 83.7 ± 1.8 |
| 2-Deoxyinosine | 100.8 ± 0.6 | 92.0 ± 3.0 | 92.7 ± 0.8 |
| 2-hydroxybutyric acid | 100.2 ± 0.4 | 75.8 ± 3.5 | 76.0 ± 1.3 |
| 2-hydroxyglutaric acid | 89.4 ± 3.3 | 82.3 ± 2.0 | 73.6 ± 1.9 |
| 2-isopropylmalic acid | 100.3 ± 0.8 | 93.7 ± 2.7 | 94.0 ± 2.1 |
| 2-ketobutyric acid | 90.4 ± 1.5 | 79.1 ± 0.3 | 71.5 ± 2.2 |
| 2-oxoadipic acid | 97.6 ± 1.7 | 84.4 ± 1.5 | 82.4 ± 1.5 |
| 2-phosphoglycerate | 102.1 ± 0.4 | 84.7 ± 3.3 | 86.5 ± 1.5 |
| 3-amino-isobutyric acid | 90.0 ± 0.3 | 89.5 ± 2.0 | 80.6 ± 0.5 |
| 3-hydroxy-3-methyl-2-oxobutanoate | 87.4 ± 2.3 | 77.4 ± 0.7 | 67.6 ± 2.1 |
| 3-hydroxybutyric acid | 96.7 ± 3.5 | 80.8 ± 1.7 | 78.1 ± 2.9 |
| 3-Hydroxy-Proline | 95.7 ± 3.1 | 80.5 ± 2.2 | 77.0 ± 0.5 |
| 3-isopropylmalic acid | 88.8 ± 2.8 | 78.5 ± 0.5 | 69.7 ± 1.5 |
| 3-methyl-2-oxopentanoate | 88.6 ± 3.0 | 94.8 ± 1.8 | 84.0 ± 0.6 |
| 3-Methylhistidine | 94.6 ± 2.9 | 88.8 ± 2.7 | 84.0 ± 1.4 |
| 3-Phosphoglyceraldehyde | 92.0 ± 3.1 | 88.8 ± 1.8 | 81.7 ± 0.5 |
| 3-phosphoglycerate | 98.8 ± 0.9 | 86.2 ± 3.2 | 85.2 ± 3.1 |
| 4-amino-butyric acid | 99.9 ± 1.5 | 81.0 ± 3.2 | 80.9 ± 3.2 |
| 4-Hydroxybenzoic acid | 99.6 ± 0.9 | 90.8 ± 0.5 | 90.4 ± 0.7 |
| 4-Hydroxyglutamate | 95.5 ± 0.3 | 93.5 ± 3.3 | 89.3 ± 3.0 |
| 4-Hydroxy-Proline | 103.8 ± 1.8 | 83.5 ± 1.8 | 86.7 ± 3.6 |
| 5-Aminolevulinic acid | 99.8 ± 3.6 | 93.1 ± 3.0 | 92.9 ± 3.0 |
| 5-Aminovaleric acid | 95.9 ± 1.5 | 76.4 ± 0.9 | 73.3 ± 0.8 |
| 5-Hydroxy-Tryptophan | 94.4 ± 3.2 | 77.1 ± 3.3 | 72.8 ± 0.7 |
| 5-Methoxytryptophan | 98.5 ± 1.8 | 84.5 ± 1.4 | 83.2 ± 1.3 |
| 5-Methylcytosine | 99.5 ± 1.3 | 77.9 ± 0.6 | 77.5 ± 1.4 |
| 6-phosphogluconate | 87.2 ± 2.1 | 89.4 ± 0.8 | 78.0 ± 2.9 |
| Acetoacetate | 100.1 ± 2.5 | 81.0 ± 3.1 | 81.1 ± 0.3 |
| Acetylcarnitine | 102.0 ± 3.0 | 76.2 ± 3.4 | 77.7 ± 2.6 |
| Acetyl-CoA | 96.9 ± 1.5 | 93.2 ± 2.0 | 90.3 ± 1.9 |
| Aconitic Acid | 96.6 ± 2.6 | 88.7 ± 2.8 | 85.7 ± 1.8 |
| Adenine | 101.7 ± 0.9 | 79.1 ± 1.2 | 80.4 ± 2.5 |
| Adenosine | 89.1 ± 0.4 | 88.2 ± 1.0 | 78.6 ± 2.8 |
| Adenosine 3',5'-cyclic monophosphate | 93.2 ± 0.6 | 77.4 ± 2.3 | 72.1 ± 1.4 |
| Adenosine Monophosphate | 98.0 ± 0.8 | 89.3 ± 2.7 | 87.5 ± 1.9 |
| Adenosine-5'-diphosphate | 104.0 ± 1.7 | 78.8 ± 3.5 | 82.0 ± 2.9 |
| Adenosine-5'-triphosphate | 87.4 ± 1.8 | 79.1 ± 2.3 | 69.1 ± 3.1 |
| Adenylosuccinate | 95.3 ± 0.3 | 75.2 ± 1.4 | 71.7 ± 2.9 |
| Alanine | 91.2 ± 2.3 | 94.9 ± 1.0 | 86.5 ± 2.1 |
| Allantoin | 96.3 ± 2.9 | 75.8 ± 1.1 | 73.0 ± 2.7 |
| alpha-ketoglutaric acid | 96.3 ± 2.3 | 91.5 ± 1.9 | 88.1 ± 1.4 |
| Anserine | 87.8 ± 0.4 | 80.8 ± 2.1 | 70.9 ± 1.9 |
| Trigonelleine | 93.6 ± 2.4 | 93.7 ± 2.8 | 87.7 ± 1.5 |
| Arginine | 90.2 ± 1.6 | 80.3 ± 2.4 | 72.4 ± 2.3 |
| Argininosuccinic Acid | 89.1 ± 2.6 | 92.2 ± 1.7 | 82.2 ± 0.7 |
| Ascorbic acid | 92.2 ± 1.1 | 94.1 ± 1.7 | 86.8 ± 0.4 |
| Asparagine | 95.5 ± 0.8 | 89.8 ± 0.6 | 85.8 ± 0.4 |
| Aspartic acid | 96.3 ± 3.3 | 79.5 ± 3.5 | 76.6 ± 1.8 |
| Asymmetric dimethylarginine | 103.5 ± 1.5 | 84.7 ± 0.8 | 87.7 ± 2.8 |
| Azelaic acid | 95.2 ± 2.7 | 77.9 ± 0.4 | 74.2 ± 1.3 |
| beta-Alanine | 99.7 ± 0.5 | 82.2 ± 2.2 | 82.0 ± 2.7 |
| Betaine | 91.7 ± 2.3 | 94.9 ± 1.6 | 87.0 ± 0.7 |
| beta-Ureidopropionate | 87.2 ± 3.0 | 84.8 ± 2.5 | 73.9 ± 1.1 |
| Biotin | 94.2 ± 1.1 | 91.7 ± 3.0 | 86.4 ± 2.1 |
| Cadaverine | 89.3 ± 0.5 | 82.7 ± 1.7 | 73.9 ± 1.1 |
| Carnitine | 90.1 ± 0.4 | 94.5 ± 2.0 | 85.1 ± 1.0 |
| Carnosine | 96.7 ± 3.0 | 92.4 ± 1.7 | 89.4 ± 2.8 |
| cGMP | 97.1 ± 3.5 | 90.5 ± 1.3 | 87.9 ± 1.1 |
| Choline | 99.0 ± 0.8 | 87.8 ± 3.2 | 86.9 ± 2.3 |
| Cinnamic acid | 100.5 ± 2.7 | 91.9 ± 2.9 | 92.4 ± 1.8 |
| Citraconic acid | 93.2 ± 2.0 | 75.4 ± 2.6 | 70.3 ± 2.1 |
| Citramalic acid | 96.5 ± 2.6 | 84.7 ± 3.5 | 81.7 ± 2.5 |
| Citric Acid | 102.5 ± 2.0 | 79.8 ± 3.4 | 81.8 ± 3.5 |
| Citrulline | 88.1 ± 0.7 | 90.7 ± 2.7 | 79.9 ± 0.6 |
| Creatine | 101.4 ± 1.4 | 77.4 ± 2.0 | 78.5 ± 1.9 |
| Creatinine | 98.3 ± 3.6 | 85.9 ± 0.6 | 84.4 ± 1.5 |
| Cystathionine | 90.0 ± 1.3 | 90.3 ± 3.4 | 81.3 ± 2.0 |
| Cysteamine | 103.3 ± 1.2 | 82.8 ± 1.1 | 85.5 ± 2.7 |
| Cysteine | 93.9 ± 2.2 | 75.6 ± 0.8 | 71.0 ± 3.0 |
| Cysteinylglycine | 89.2 ± 3.3 | 89.6 ± 1.4 | 79.9 ± 1.9 |
| Cystine | 97.9 ± 1.1 | 88.7 ± 2.4 | 86.8 ± 2.1 |
| Cytidine | 96.3 ± 2.7 | 81.1 ± 1.5 | 78.1 ± 2.2 |
| Cytidine 5'-diphosphate | 90.5 ± 3.1 | 86.8 ± 1.2 | 78.6 ± 2.8 |
| Cytidine 5'-triphosphate | 103.8 ± 2.1 | 91.8 ± 2.7 | 95.3 ± 1.5 |
| Cytidine monophosphate | 99.7 ± 0.5 | 80.6 ± 2.2 | 80.4 ± 0.4 |
| Cytosine | 86.7 ± 0.7 | 87.6 ± 3.5 | 75.9 ± 1.7 |
| Dihydroxyacetonephosphate | 98.1 ± 2.8 | 78.1 ± 3.6 | 76.6 ± 1.3 |
| Dimethylglycine | 88.9 ± 0.8 | 93.9 ± 1.9 | 83.5 ± 0.3 |
| Erythrose-4-phosphate | 96.2 ± 2.8 | 78.0 ± 2.6 | 75.0 ± 3.2 |
| Ethanolamine | 95.8 ± 1.0 | 76.5 ± 2.6 | 73.3 ± 3.2 |
| FAD | 91.8 ± 3.2 | 84.2 ± 0.3 | 77.3 ± 3.0 |
| Folic acid | 98.0 ± 1.8 | 89.4 ± 2.6 | 87.6 ± 2.0 |
| Fructose | 97.0 ± 1.6 | 83.9 ± 0.8 | 81.4 ± 0.7 |
| Fructose-1,6-bisphosphate | 94.3 ± 3.0 | 93.0 ± 2.7 | 87.7 ± 0.4 |
| Fructose-6-Phosphate | 92.5 ± 0.9 | 82.6 ± 1.0 | 76.4 ± 2.3 |
| Fumaric Acid | 98.6 ± 2.3 | 79.2 ± 3.1 | 78.1 ± 1.7 |
| Galactosamine | 97.2 ± 1.0 | 77.1 ± 0.7 | 74.9 ± 3.4 |
| Galactose-1-phosphate | 89.3 ± 2.7 | 88.9 ± 1.8 | 79.4 ± 0.3 |
| Galactose-6-phosphate | 92.7 ± 2.4 | 88.2 ± 2.4 | 81.8 ± 3.5 |
| gamma-Glutamylcysteine | 101.8 ± 1.2 | 92.0 ± 1.9 | 93.7 ± 1.8 |
| Glucaric Acid | 103.3 ± 3.4 | 82.7 ± 2.4 | 85.4 ± 1.1 |
| Gluconic acid | 103.3 ± 0.4 | 77.1 ± 3.4 | 79.6 ± 0.9 |
| Gluconolactone | 95.4 ± 1.7 | 89.8 ± 2.0 | 85.7 ± 2.0 |
| Glucosamine | 99.9 ± 2.2 | 85.5 ± 3.6 | 85.4 ± 0.5 |
| Glucosamine-6-Phosphate | 91.3 ± 0.5 | 86.0 ± 1.2 | 78.5 ± 0.9 |
| Glucose | 88.4 ± 0.8 | 94.4 ± 1.0 | 83.4 ± 3.0 |
| Glucose-1-phosphate | 93.4 ± 1.0 | 81.9 ± 2.5 | 76.5 ± 2.6 |
| Glucose-6-phosphate | 102.6 ± 0.9 | 89.2 ± 3.3 | 91.5 ± 3.5 |
| Glucuronolactone | 89.4 ± 3.4 | 75.7 ± 2.3 | 67.7 ± 2.1 |
| Glutamic acid | 94.3 ± 1.4 | 90.0 ± 1.2 | 84.9 ± 1.9 |
| Glutamine | 104.0 ± 2.0 | 92.9 ± 0.3 | 96.6 ± 0.8 |
| Glutathione (GSSG) | 102.3 ± 0.8 | 84.8 ± 2.3 | 86.8 ± 2.3 |
| Glutathione | 91.4 ± 0.5 | 84.2 ± 3.4 | 77.0 ± 2.9 |
| Glycerophosphocholine | 97.0 ± 3.0 | 83.5 ± 3.5 | 81.0 ± 3.1 |
| Glycine | 98.5 ± 0.6 | 86.9 ± 1.1 | 85.6 ± 1.2 |
| Glycolic acid | 102.2 ± 0.9 | 94.9 ± 0.4 | 97.0 ± 3.1 |
| Guanine | 96.7 ± 2.6 | 94.3 ± 2.8 | 91.2 ± 1.3 |
| Guanosine | 96.5 ± 2.4 | 93.5 ± 1.1 | 90.2 ± 3.0 |
| Guanosine 5'-diphosphate | 87.2 ± 0.8 | 90.2 ± 1.4 | 78.7 ± 1.6 |
| Guanosine-5'-monophosphate | 99.4 ± 2.7 | 77.9 ± 1.6 | 77.4 ± 1.9 |
| Hippuric acid | 89.8 ± 2.7 | 85.4 ± 1.2 | 76.7 ± 2.2 |
| Histamine | 93.8 ± 3.5 | 89.8 ± 0.4 | 84.2 ± 3.3 |
| Histidine | 94.0 ± 0.5 | 85.6 ± 3.1 | 80.5 ± 1.9 |
| Histidinol | 96.1 ± 2.7 | 92.7 ± 1.4 | 89.1 ± 2.6 |
| Homoalanine | 100.3 ± 3.5 | 90.8 ± 1.7 | 91.1 ± 1.1 |
| Homocitrulline | 103.6 ± 0.9 | 78.4 ± 0.5 | 81.2 ± 2.1 |
| Homocysteine | 96.5 ± 2.8 | 91.2 ± 1.2 | 88.0 ± 2.6 |
| Homocystine | 95.2 ± 2.9 | 91.4 ± 3.0 | 87.0 ± 1.4 |
| Homogentisic acid | 101.7 ± 2.0 | 75.3 ± 3.6 | 76.6 ± 2.2 |
| Homoleucine | 101.2 ± 0.8 | 94.8 ± 1.0 | 96.0 ± 2.1 |
| Homophenylalanine | 101.9 ± 0.4 | 85.9 ± 0.9 | 87.5 ± 2.7 |
| Homoserine | 94.6 ± 0.6 | 94.3 ± 3.2 | 89.2 ± 0.9 |
| Homovanillic acid | 94.4 ± 1.3 | 82.2 ± 2.7 | 77.6 ± 2.4 |
| Hypotaurine | 97.7 ± 0.8 | 79.0 ± 3.6 | 77.2 ± 3.6 |
| Hypoxanthine | 92.4 ± 1.1 | 93.2 ± 3.0 | 86.1 ± 1.0 |
| Inosine | 102.0 ± 2.0 | 86.6 ± 2.3 | 88.3 ± 1.8 |
| Inosine 5'-monophosphate | 92.0 ± 1.0 | 90.2 ± 2.6 | 82.9 ± 0.4 |
| Inositol | 91.6 ± 2.9 | 93.9 ± 2.8 | 86.0 ± 2.4 |
| Isocitric Acid | 91.4 ± 2.2 | 77.1 ± 2.7 | 70.5 ± 1.1 |
| Isoleucine | 92.2 ± 1.7 | 85.6 ± 0.6 | 79.0 ± 2.4 |
| Itaconic acid | 104.1 ± 2.8 | 90.0 ± 2.7 | 93.7 ± 0.9 |
| ITP | 97.2 ± 0.6 | 77.1 ± 3.5 | 75.0 ± 3.2 |
| Kynurenic acid | 100.0 ± 0.5 | 94.9 ± 1.6 | 94.9 ± 0.6 |
| Kynurenine | 94.9 ± 2.7 | 75.2 ± 0.9 | 71.4 ± 2.1 |
| Lactic Acid | 99.8 ± 0.9 | 75.8 ± 1.3 | 75.7 ± 3.1 |
| Lactose | 87.7 ± 1.1 | 83.9 ± 1.6 | 73.6 ± 2.5 |
| Leucine | 89.8 ± 2.4 | 91.3 ± 0.5 | 82.0 ± 2.1 |
| Levulinic acid | 93.0 ± 2.8 | 94.7 ± 2.6 | 88.1 ± 2.9 |
| Lysine | 97.1 ± 2.5 | 76.8 ± 1.4 | 74.6 ± 0.5 |
| Malic Acid | 92.9 ± 1.2 | 89.5 ± 2.8 | 83.2 ± 0.7 |
| Malonic acid | 102.9 ± 1.5 | 75.4 ± 2.2 | 77.6 ± 3.3 |
| Mannose | 94.6 ± 0.7 | 75.5 ± 1.4 | 71.4 ± 0.6 |
| Mannose-1 with Mannose-6-phosphate | 92.2 ± 2.6 | 89.2 ± 2.9 | 82.3 ± 0.6 |
| Mesaconic acid | 91.5 ± 2.6 | 77.5 ± 1.5 | 70.9 ± 0.7 |
| Methionine | 88.8 ± 2.5 | 93.5 ± 0.5 | 83.0 ± 1.5 |
| Methionine sulfone | 90.0 ± 2.2 | 89.5 ± 2.6 | 80.6 ± 0.6 |
| Methionine sulfoxide | 89.7 ± 1.2 | 90.1 ± 0.8 | 80.8 ± 2.8 |
| Methylmalonate | 86.9 ± 0.5 | 91.9 ± 2.1 | 79.9 ± 1.1 |
| N-Acetyl-Alanine | 93.2 ± 1.3 | 91.9 ± 1.3 | 85.6 ± 1.4 |
| N-Acetyl-Aspartic Acid | 99.0 ± 3.0 | 78.7 ± 1.1 | 77.9 ± 2.4 |
| N-Acetyl-Citrulline | 100.2 ± 2.3 | 78.5 ± 3.4 | 78.7 ± 2.1 |
| N-Acetyl-Galactosamine | 90.4 ± 3.4 | 79.2 ± 0.5 | 71.6 ± 1.0 |
| N-Acetyl-Glucosamine | 99.9 ± 1.0 | 80.7 ± 1.6 | 80.6 ± 2.4 |
| N-Acetyl-Glutamine | 100.3 ± 2.2 | 89.5 ± 2.0 | 89.8 ± 3.3 |
| N-Acetyl-Glycine | 94.0 ± 1.8 | 82.8 ± 1.7 | 77.8 ± 2.0 |
| N-Acetyl-Leucine | 95.6 ± 0.4 | 93.2 ± 3.2 | 89.1 ± 0.4 |
| N-Acetyl-Methionine | 93.2 ± 1.3 | 77.5 ± 2.5 | 72.2 ± 1.7 |
| N-Acetylneuraminic acid | 102.6 ± 3.3 | 83.8 ± 2.8 | 86.0 ± 1.7 |
| N-Acetyl-Phenylalanine | 90.6 ± 2.8 | 87.1 ± 2.7 | 78.9 ± 1.0 |
| N-Acetyl-Proline | 91.2 ± 0.4 | 82.8 ± 0.9 | 75.5 ± 2.9 |
| N-Acetyl-Serine | 95.3 ± 2.3 | 88.9 ± 1.4 | 84.7 ± 2.2 |
| N-Acetyl-Serotonin | 99.3 ± 1.6 | 89.4 ± 1.1 | 88.8 ± 0.9 |
| N-Acetyl-Tryptophan | 102.9 ± 3.2 | 80.3 ± 1.7 | 82.7 ± 1.2 |
| N-Acetyl-Tyrosine | 93.3 ± 1.4 | 90.9 ± 3.6 | 84.8 ± 2.8 |
| NAD | 95.4 ± 2.0 | 92.6 ± 1.9 | 88.3 ± 1.1 |
| NADH | 99.2 ± 1.3 | 80.3 ± 1.9 | 79.6 ± 2.3 |
| NADP | 102.4 ± 1.3 | 76.7 ± 0.7 | 78.5 ± 3.0 |
| NADPH | 89.0 ± 2.2 | 79.7 ± 0.4 | 70.9 ± 0.4 |
| N-alpha-Acetyl-lysine | 87.4 ± 0.8 | 78.8 ± 3.2 | 68.9 ± 1.9 |
| N-alpha-Acetyl-Ornithine | 99.5 ± 0.9 | 85.2 ± 1.9 | 84.7 ± 2.0 |
| N-epsilon-Acetyl-lysine | 96.2 ± 2.6 | 89.9 ± 3.1 | 86.5 ± 1.0 |
| N-epsilon-Acetyl-Ornithine | 100.8 ± 3.5 | 93.4 ± 2.3 | 94.1 ± 1.4 |
| Nicotinamide | 97.8 ± 1.4 | 91.3 ± 1.0 | 89.3 ± 1.1 |
| Nicotinic acid | 97.2 ± 2.2 | 82.9 ± 2.1 | 80.6 ± 0.8 |
| N-Methyl-Aspartic Acid | 88.0 ± 2.8 | 78.9 ± 1.8 | 69.4 ± 1.0 |
| O-Acetyl-Serine | 94.4 ± 1.7 | 93.9 ± 2.1 | 88.6 ± 3.0 |
| O-Phosphoethanolamine | 97.7 ± 3.1 | 79.9 ± 1.9 | 78.1 ± 1.9 |
| O-Phospho-Serine | 87.3 ± 0.7 | 83.1 ± 1.2 | 72.6 ± 1.1 |
| Ophthalmic acid | 90.3 ± 2.9 | 76.5 ± 3.4 | 69.1 ± 3.2 |
| Ornithine | 99.7 ± 2.6 | 81.6 ± 1.0 | 81.4 ± 1.0 |
| Orotic acid | 98.0 ± 0.6 | 89.2 ± 1.1 | 87.4 ± 1.5 |
| Oxaloacetate | 101.3 ± 2.8 | 79.2 ± 0.7 | 80.2 ± 1.7 |
| Pantothenic acid | 99.9 ± 3.4 | 94.4 ± 3.6 | 94.3 ± 2.6 |
| p-Coumaric Acid | 99.4 ± 1.8 | 90.4 ± 1.6 | 89.8 ± 0.8 |
| Phenylalanine | 99.4 ± 1.5 | 89.1 ± 2.7 | 88.5 ± 0.5 |
| Phenylpyruvate | 89.1 ± 2.1 | 86.8 ± 3.2 | 77.4 ± 1.8 |
| Phosphocreatine | 95.2 ± 2.3 | 81.3 ± 3.0 | 77.4 ± 1.5 |
| Phosphoenolpyruvate | 90.5 ± 2.3 | 89.7 ± 3.0 | 81.2 ± 1.1 |
| Phosphoribosylpyrophosphate | 97.9 ± 1.7 | 94.2 ± 2.3 | 92.2 ± 0.5 |
| Picolinate | 93.1 ± 2.7 | 93.0 ± 0.5 | 86.6 ± 2.7 |
| Pipecolic acid | 91.7 ± 3.1 | 87.4 ± 0.4 | 80.1 ± 3.6 |
| Proline | 95.6 ± 1.3 | 88.4 ± 2.7 | 84.6 ± 1.4 |
| Pyridoxal | 103.7 ± 1.3 | 85.2 ± 0.8 | 88.3 ± 1.3 |
| Pyridoxal-5-phosphate | 99.6 ± 1.6 | 83.7 ± 3.2 | 83.3 ± 2.5 |
| Pyridoxamine | 96.6 ± 3.3 | 77.5 ± 3.6 | 74.8 ± 1.1 |
| Pyridoxine | 94.0 ± 1.1 | 80.4 ± 1.1 | 75.6 ± 3.6 |
| Pyroglutamic acid | 99.8 ± 3.2 | 91.5 ± 2.4 | 91.3 ± 1.0 |
| Pyrrolysine | 97.7 ± 3.1 | 93.2 ± 1.2 | 91.1 ± 3.4 |
| Pyruvic Acid | 96.1 ± 2.8 | 86.6 ± 2.8 | 83.2 ± 2.8 |
| Riboflavin | 90.3 ± 3.6 | 81.0 ± 1.9 | 73.1 ± 0.6 |
| Ribose | 94.2 ± 3.5 | 90.3 ± 2.3 | 85.0 ± 0.6 |
| Ribose-5-phosphate | 99.7 ± 2.6 | 75.2 ± 0.3 | 74.9 ± 1.6 |
| Ribulose-1-phosphate | 99.5 ± 2.9 | 84.5 ± 1.4 | 84.0 ± 1.6 |
| Ribulose-5-phosphate | 95.1 ± 3.6 | 91.0 ± 3.0 | 86.5 ± 2.3 |
| S-Adenosylmethionine | 92.1 ± 2.8 | 82.3 ± 0.8 | 75.8 ± 1.6 |
| Sarcosine | 96.9 ± 0.5 | 76.8 ± 0.6 | 74.4 ± 0.4 |
| Sedoheptulose-7-phosphate | 94.8 ± 0.9 | 89.6 ± 2.8 | 84.9 ± 0.6 |
| Selenocystine | 90.0 ± 1.1 | 89.6 ± 2.9 | 80.7 ± 2.9 |
| Selenomethionine | 88.8 ± 0.4 | 92.6 ± 1.4 | 82.2 ± 0.7 |
| Serine | 97.0 ± 3.0 | 85.9 ± 3.5 | 83.3 ± 2.7 |
| Serotonin | 100.0 ± 1.8 | 83.9 ± 3.3 | 83.9 ± 0.7 |
| Sorbitol | 98.2 ± 1.8 | 84.2 ± 0.9 | 82.7 ± 0.6 |
| Spermidine | 91.6 ± 0.7 | 80.7 ± 1.3 | 73.9 ± 1.0 |
| Spermine | 104.0 ± 2.3 | 75.9 ± 0.5 | 78.9 ± 0.5 |
| Succinic Acid | 98.4 ± 2.1 | 86.3 ± 0.9 | 84.9 ± 1.1 |
| Sucrose | 95.0 ± 0.8 | 78.2 ± 1.8 | 74.3 ± 1.7 |
| Symmetric dimethylarginine | 88.9 ± 1.4 | 79.6 ± 3.1 | 70.7 ± 0.6 |
| Taurine | 97.4 ± 3.5 | 79.2 ± 1.9 | 77.1 ± 2.1 |
| Thiamine | 99.6 ± 1.1 | 92.8 ± 0.6 | 92.4 ± 0.8 |
| Thiamine monophosphate | 97.5 ± 2.4 | 85.0 ± 3.4 | 82.8 ± 1.1 |
| Thiamine pyrophosphate | 99.6 ± 2.0 | 91.7 ± 2.8 | 91.3 ± 2.4 |
| Threonic acid | 89.5 ± 2.8 | 94.7 ± 2.7 | 84.7 ± 2.7 |
| Threonine | 96.1 ± 2.1 | 80.7 ± 1.3 | 77.5 ± 3.3 |
| Thymidine | 89.4 ± 3.4 | 86.4 ± 0.4 | 77.3 ± 3.2 |
| Thymidine Monophosphate | 99.9 ± 1.1 | 78.3 ± 2.9 | 78.2 ± 2.8 |
| Thymine | 87.0 ± 3.2 | 88.5 ± 1.0 | 77.0 ± 1.2 |
| Trimethylamine N-oxide | 103.1 ± 0.8 | 94.9 ± 1.7 | 97.8 ± 3.5 |
| Tryptophan | 90.3 ± 1.2 | 75.5 ± 1.8 | 68.2 ± 0.8 |
| Tyrosine | 89.8 ± 1.7 | 90.9 ± 0.5 | 81.7 ± 2.9 |
| Uracil | 94.9 ± 2.3 | 94.6 ± 1.7 | 89.8 ± 3.1 |
| Uric Acid | 102.5 ± 2.1 | 92.0 ± 3.1 | 94.3 ± 0.5 |
| Uridine | 90.4 ± 3.3 | 89.9 ± 0.5 | 81.3 ± 3.1 |
| Uridine 5'-diphosphate | 88.3 ± 1.2 | 81.0 ± 2.2 | 71.5 ± 1.4 |
| Uridine Monophosphate | 88.9 ± 1.0 | 85.7 ± 0.6 | 76.2 ± 1.2 |
| Uridine-5-Diphosphogalactose | 97.5 ± 0.9 | 93.1 ± 2.2 | 90.7 ± 1.6 |
| Uridine-5-Diphosphoglucose | 101.8 ± 2.9 | 83.7 ± 3.1 | 85.2 ± 2.1 |
| Urocanic acid | 103.7 ± 1.1 | 82.0 ± 1.1 | 85.0 ± 0.9 |
| UTP | 98.0 ± 3.2 | 85.8 ± 1.8 | 84.1 ± 1.9 |
| Valine | 91.0 ± 1.7 | 91.6 ± 2.9 | 83.3 ± 3.4 |
| Xanthine | 102.3 ± 1.7 | 91.5 ± 2.9 | 93.6 ± 2.3 |
| Xanthosine | 100.1 ± 3.1 | 80.7 ± 1.6 | 80.8 ± 3.3 |
| Xanthurenic acid | 95.3 ± 2.2 | 83.9 ± 2.8 | 79.9 ± 1.6 |
| XMP | 98.3 ± 2.8 | 94.0 ± 0.6 | 92.4 ± 1.3 |
| Xylulose-5-phosphate | 94.6 ± 0.9 | 87.5 ± 3.3 | 82.8 ± 0.8 |

### **Table S6** | Matrix effect (ME), Process Efficiency (PE) and Extraction Recovery (RE) of metabolites without internal standard correction (Urine extract as matrix, n = 5). The concentration of the QC sample used was 100 ng/mL.

| Metabolite | ME [%] | RE [%] | PE [%] |
| --- | --- | --- | --- |
| 1-Methylhistidine | 104.4 ± 3.3 | 87.0 ± 2.9 | 90.8 ± 0.6 |
| 2,3-Bisphosphoglycerate | 91.8 ± 1.7 | 78.6 ± 2.5 | 72.2 ± 1.3 |
| 2-Aminoadipic Acid | 91.6 ± 1.6 | 83.1 ± 3.3 | 76.1 ± 3.4 |
| 2-Deoxyadenosine | 89.3 ± 2.1 | 87.1 ± 1.6 | 77.8 ± 2.0 |
| 2'-Deoxyadenosine 5'-monophosphate | 103.9 ± 0.6 | 94.8 ± 0.7 | 98.5 ± 0.5 |
| 2'-deoxyadenosine 5'-triphosphate | 91.3 ± 2.6 | 84.4 ± 1.5 | 77.1 ± 2.4 |
| 2-Deoxycytidine | 103.6 ± 2.3 | 82.6 ± 3.4 | 85.6 ± 0.3 |
| 2'-Deoxycytidine-5'-monophosphate | 101.7 ± 1.8 | 92.6 ± 1.1 | 94.2 ± 2.4 |
| 2-Deoxyguanosine | 90.3 ± 0.5 | 92.7 ± 3.3 | 83.7 ± 1.8 |
| 2-Deoxyinosine | 100.8 ± 0.6 | 92.0 ± 3.0 | 92.7 ± 0.8 |
| 2-hydroxybutyric acid | 100.2 ± 0.4 | 75.8 ± 3.5 | 76.0 ± 1.3 |
| 2-hydroxyglutaric acid | 89.4 ± 3.3 | 82.3 ± 2.0 | 73.6 ± 1.9 |
| 2-isopropylmalic acid | 100.3 ± 0.8 | 93.7 ± 2.7 | 94.0 ± 2.1 |
| 2-ketobutyric acid | 90.4 ± 1.5 | 79.1 ± 0.3 | 71.5 ± 2.2 |
| 2-oxoadipic acid | 97.6 ± 1.7 | 84.4 ± 1.5 | 82.4 ± 1.5 |
| 2-phosphoglycerate | 102.1 ± 0.4 | 84.7 ± 3.3 | 86.5 ± 1.5 |
| 3-amino-isobutyric acid | 90.0 ± 0.3 | 89.5 ± 2.0 | 80.6 ± 0.5 |
| 3-hydroxy-3-methyl-2-oxobutanoate | 87.4 ± 2.3 | 77.4 ± 0.7 | 67.6 ± 2.1 |
| 3-hydroxybutyric acid | 96.7 ± 3.5 | 80.8 ± 1.7 | 78.1 ± 2.9 |
| 3-Hydroxy-Proline | 95.7 ± 3.1 | 80.5 ± 2.2 | 77.0 ± 0.5 |
| 3-isopropylmalic acid | 88.8 ± 2.8 | 78.5 ± 0.5 | 69.7 ± 1.5 |
| 3-methyl-2-oxopentanoate | 88.6 ± 3.0 | 94.8 ± 1.8 | 84.0 ± 0.6 |
| 3-Methylhistidine | 94.6 ± 2.9 | 88.8 ± 2.7 | 84.0 ± 1.4 |
| 3-Phosphoglyceraldehyde | 92.0 ± 3.1 | 88.8 ± 1.8 | 81.7 ± 0.5 |
| 3-phosphoglycerate | 98.8 ± 0.9 | 86.2 ± 3.2 | 85.2 ± 3.1 |
| 4-amino-butyric acid | 99.9 ± 1.5 | 81.0 ± 3.2 | 80.9 ± 3.2 |
| 4-Hydroxybenzoic acid | 99.6 ± 0.9 | 90.8 ± 0.5 | 90.4 ± 0.7 |
| 4-Hydroxyglutamate | 95.5 ± 0.3 | 93.5 ± 3.3 | 89.3 ± 3.0 |
| 4-Hydroxy-Proline | 103.8 ± 1.8 | 83.5 ± 1.8 | 86.7 ± 3.6 |
| 5-Aminolevulinic acid | 99.8 ± 3.6 | 93.1 ± 3.0 | 92.9 ± 3.0 |
| 5-Aminovaleric acid | 95.9 ± 1.5 | 76.4 ± 0.9 | 73.3 ± 0.8 |
| 5-Hydroxy-Tryptophan | 94.4 ± 3.2 | 77.1 ± 3.3 | 72.8 ± 0.7 |
| 5-Methoxytryptophan | 98.5 ± 1.8 | 84.5 ± 1.4 | 83.2 ± 1.3 |
| 5-Methylcytosine | 99.5 ± 1.3 | 77.9 ± 0.6 | 77.5 ± 1.4 |
| 6-phosphogluconate | 87.2 ± 2.1 | 89.4 ± 0.8 | 78.0 ± 2.9 |
| Acetoacetate | 100.1 ± 2.5 | 81.0 ± 3.1 | 81.1 ± 0.3 |
| Acetylcarnitine | 102.0 ± 3.0 | 76.2 ± 3.4 | 77.7 ± 2.6 |
| Acetyl-CoA | 96.9 ± 1.5 | 93.2 ± 2.0 | 90.3 ± 1.9 |
| Aconitic Acid | 96.6 ± 2.6 | 88.7 ± 2.8 | 85.7 ± 1.8 |
| Adenine | 101.7 ± 0.9 | 79.1 ± 1.2 | 80.4 ± 2.5 |
| Adenosine | 89.1 ± 0.4 | 88.2 ± 1.0 | 78.6 ± 2.8 |
| Adenosine 3',5'-cyclic monophosphate | 93.2 ± 0.6 | 77.4 ± 2.3 | 72.1 ± 1.4 |
| Adenosine Monophosphate | 98.0 ± 0.8 | 89.3 ± 2.7 | 87.5 ± 1.9 |
| Adenosine-5'-diphosphate | 104.0 ± 1.7 | 78.8 ± 3.5 | 82.0 ± 2.9 |
| Adenosine-5'-triphosphate | 87.4 ± 1.8 | 79.1 ± 2.3 | 69.1 ± 3.1 |
| Adenylosuccinate | 95.3 ± 0.3 | 75.2 ± 1.4 | 71.7 ± 2.9 |
| Alanine | 91.2 ± 2.3 | 94.9 ± 1.0 | 86.5 ± 2.1 |
| Allantoin | 96.3 ± 2.9 | 75.8 ± 1.1 | 73.0 ± 2.7 |
| alpha-ketoglutaric acid | 96.3 ± 2.3 | 91.5 ± 1.9 | 88.1 ± 1.4 |
| Anserine | 87.8 ± 0.4 | 80.8 ± 2.1 | 70.9 ± 1.9 |
| Trigonelleine | 93.6 ± 2.4 | 93.7 ± 2.8 | 87.7 ± 1.5 |
| Arginine | 90.2 ± 1.6 | 80.3 ± 2.4 | 72.4 ± 2.3 |
| Argininosuccinic Acid | 89.1 ± 2.6 | 92.2 ± 1.7 | 82.2 ± 0.7 |
| Ascorbic acid | 92.2 ± 1.1 | 94.1 ± 1.7 | 86.8 ± 0.4 |
| Asparagine | 95.5 ± 0.8 | 89.8 ± 0.6 | 85.8 ± 0.4 |
| Aspartic acid | 96.3 ± 3.3 | 79.5 ± 3.5 | 76.6 ± 1.8 |
| Asymmetric dimethylarginine | 103.5 ± 1.5 | 84.7 ± 0.8 | 87.7 ± 2.8 |
| Azelaic acid | 95.2 ± 2.7 | 77.9 ± 0.4 | 74.2 ± 1.3 |
| beta-Alanine | 99.7 ± 0.5 | 82.2 ± 2.2 | 82.0 ± 2.7 |
| Betaine | 91.7 ± 2.3 | 94.9 ± 1.6 | 87.0 ± 0.7 |
| beta-Ureidopropionate | 87.2 ± 3.0 | 84.8 ± 2.5 | 73.9 ± 1.1 |
| Biotin | 94.2 ± 1.1 | 91.7 ± 3.0 | 86.4 ± 2.1 |
| Cadaverine | 89.3 ± 0.5 | 82.7 ± 1.7 | 73.9 ± 1.1 |
| Carnitine | 90.1 ± 0.4 | 94.5 ± 2.0 | 85.1 ± 1.0 |
| Carnosine | 96.7 ± 3.0 | 92.4 ± 1.7 | 89.4 ± 2.8 |
| cGMP | 97.1 ± 3.5 | 90.5 ± 1.3 | 87.9 ± 1.1 |
| Choline | 99.0 ± 0.8 | 87.8 ± 3.2 | 86.9 ± 2.3 |
| Cinnamic acid | 100.5 ± 2.7 | 91.9 ± 2.9 | 92.4 ± 1.8 |
| Citraconic acid | 93.2 ± 2.0 | 75.4 ± 2.6 | 70.3 ± 2.1 |
| Citramalic acid | 96.5 ± 2.6 | 84.7 ± 3.5 | 81.7 ± 2.5 |
| Citric Acid | 102.5 ± 2.0 | 79.8 ± 3.4 | 81.8 ± 3.5 |
| Citrulline | 88.1 ± 0.7 | 90.7 ± 2.7 | 79.9 ± 0.6 |
| Creatine | 101.4 ± 1.4 | 77.4 ± 2.0 | 78.5 ± 1.9 |
| Creatinine | 98.3 ± 3.6 | 85.9 ± 0.6 | 84.4 ± 1.5 |
| Cystathionine | 90.0 ± 1.3 | 90.3 ± 3.4 | 81.3 ± 2.0 |
| Cysteamine | 103.3 ± 1.2 | 82.8 ± 1.1 | 85.5 ± 2.7 |
| Cysteine | 93.9 ± 2.2 | 75.6 ± 0.8 | 71.0 ± 3.0 |
| Cysteinylglycine | 89.2 ± 3.3 | 89.6 ± 1.4 | 79.9 ± 1.9 |
| Cystine | 97.9 ± 1.1 | 88.7 ± 2.4 | 86.8 ± 2.1 |
| Cytidine | 96.3 ± 2.7 | 81.1 ± 1.5 | 78.1 ± 2.2 |
| Cytidine 5'-diphosphate | 90.5 ± 3.1 | 86.8 ± 1.2 | 78.6 ± 2.8 |
| Cytidine 5'-triphosphate | 103.8 ± 2.1 | 91.8 ± 2.7 | 95.3 ± 1.5 |
| Cytidine monophosphate | 99.7 ± 0.5 | 80.6 ± 2.2 | 80.4 ± 0.4 |
| Cytosine | 86.7 ± 0.7 | 87.6 ± 3.5 | 75.9 ± 1.7 |
| Dihydroxyacetonephosphate | 98.1 ± 2.8 | 78.1 ± 3.6 | 76.6 ± 1.3 |
| Dimethylglycine | 88.9 ± 0.8 | 93.9 ± 1.9 | 83.5 ± 0.3 |
| Erythrose-4-phosphate | 96.2 ± 2.8 | 78.0 ± 2.6 | 75.0 ± 3.2 |
| Ethanolamine | 95.8 ± 1.0 | 76.5 ± 2.6 | 73.3 ± 3.2 |
| FAD | 91.8 ± 3.2 | 84.2 ± 0.3 | 77.3 ± 3.0 |
| Folic acid | 98.0 ± 1.8 | 89.4 ± 2.6 | 87.6 ± 2.0 |
| Fructose | 97.0 ± 1.6 | 83.9 ± 0.8 | 81.4 ± 0.7 |
| Fructose-1,6-bisphosphate | 94.3 ± 3.0 | 93.0 ± 2.7 | 87.7 ± 0.4 |
| Fructose-6-Phosphate | 92.5 ± 0.9 | 82.6 ± 1.0 | 76.4 ± 2.3 |
| Fumaric Acid | 98.6 ± 2.3 | 79.2 ± 3.1 | 78.1 ± 1.7 |
| Galactosamine | 97.2 ± 1.0 | 77.1 ± 0.7 | 74.9 ± 3.4 |
| Galactose-1-phosphate | 89.3 ± 2.7 | 88.9 ± 1.8 | 79.4 ± 0.3 |
| Galactose-6-phosphate | 92.7 ± 2.4 | 88.2 ± 2.4 | 81.8 ± 3.5 |
| gamma-Glutamylcysteine | 101.8 ± 1.2 | 92.0 ± 1.9 | 93.7 ± 1.8 |
| Glucaric Acid | 103.3 ± 3.4 | 82.7 ± 2.4 | 85.4 ± 1.1 |
| Gluconic acid | 103.3 ± 0.4 | 77.1 ± 3.4 | 79.6 ± 0.9 |
| Gluconolactone | 95.4 ± 1.7 | 89.8 ± 2.0 | 85.7 ± 2.0 |
| Glucosamine | 99.9 ± 2.2 | 85.5 ± 3.6 | 85.4 ± 0.5 |
| Glucosamine-6-Phosphate | 91.3 ± 0.5 | 86.0 ± 1.2 | 78.5 ± 0.9 |
| Glucose | 88.4 ± 0.8 | 94.4 ± 1.0 | 83.4 ± 3.0 |
| Glucose-1-phosphate | 93.4 ± 1.0 | 81.9 ± 2.5 | 76.5 ± 2.6 |
| Glucose-6-phosphate | 102.6 ± 0.9 | 89.2 ± 3.3 | 91.5 ± 3.5 |
| Glucuronolactone | 89.4 ± 3.4 | 75.7 ± 2.3 | 67.7 ± 2.1 |
| Glutamic acid | 94.3 ± 1.4 | 90.0 ± 1.2 | 84.9 ± 1.9 |
| Glutamine | 104.0 ± 2.0 | 92.9 ± 0.3 | 96.6 ± 0.8 |
| Glutathione (GSSG) | 102.3 ± 0.8 | 84.8 ± 2.3 | 86.8 ± 2.3 |
| Glutathione | 91.4 ± 0.5 | 84.2 ± 3.4 | 77.0 ± 2.9 |
| Glycerophosphocholine | 97.0 ± 3.0 | 83.5 ± 3.5 | 81.0 ± 3.1 |
| Glycine | 98.5 ± 0.6 | 86.9 ± 1.1 | 85.6 ± 1.2 |
| Glycolic acid | 102.2 ± 0.9 | 94.9 ± 0.4 | 97.0 ± 3.1 |
| Guanine | 96.7 ± 2.6 | 94.3 ± 2.8 | 91.2 ± 1.3 |
| Guanosine | 96.5 ± 2.4 | 93.5 ± 1.1 | 90.2 ± 3.0 |
| Guanosine 5'-diphosphate | 87.2 ± 0.8 | 90.2 ± 1.4 | 78.7 ± 1.6 |
| Guanosine-5'-monophosphate | 99.4 ± 2.7 | 77.9 ± 1.6 | 77.4 ± 1.9 |
| Hippuric acid | 89.8 ± 2.7 | 85.4 ± 1.2 | 76.7 ± 2.2 |
| Histamine | 93.8 ± 3.5 | 89.8 ± 0.4 | 84.2 ± 3.3 |
| Histidine | 94.0 ± 0.5 | 85.6 ± 3.1 | 80.5 ± 1.9 |
| Histidinol | 96.1 ± 2.7 | 92.7 ± 1.4 | 89.1 ± 2.6 |
| Homoalanine | 100.3 ± 3.5 | 90.8 ± 1.7 | 91.1 ± 1.1 |
| Homocitrulline | 103.6 ± 0.9 | 78.4 ± 0.5 | 81.2 ± 2.1 |
| Homocysteine | 96.5 ± 2.8 | 91.2 ± 1.2 | 88.0 ± 2.6 |
| Homocystine | 95.2 ± 2.9 | 91.4 ± 3.0 | 87.0 ± 1.4 |
| Homogentisic acid | 101.7 ± 2.0 | 75.3 ± 3.6 | 76.6 ± 2.2 |
| Homoleucine | 101.2 ± 0.8 | 94.8 ± 1.0 | 96.0 ± 2.1 |
| Homophenylalanine | 101.9 ± 0.4 | 85.9 ± 0.9 | 87.5 ± 2.7 |
| Homoserine | 94.6 ± 0.6 | 94.3 ± 3.2 | 89.2 ± 0.9 |
| Homovanillic acid | 94.4 ± 1.3 | 82.2 ± 2.7 | 77.6 ± 2.4 |
| Hypotaurine | 97.7 ± 0.8 | 79.0 ± 3.6 | 77.2 ± 3.6 |
| Hypoxanthine | 92.4 ± 1.1 | 93.2 ± 3.0 | 86.1 ± 1.0 |
| Inosine | 102.0 ± 2.0 | 86.6 ± 2.3 | 88.3 ± 1.8 |
| Inosine 5'-monophosphate | 92.0 ± 1.0 | 90.2 ± 2.6 | 82.9 ± 0.4 |
| Inositol | 91.6 ± 2.9 | 93.9 ± 2.8 | 86.0 ± 2.4 |
| Isocitric Acid | 91.4 ± 2.2 | 77.1 ± 2.7 | 70.5 ± 1.1 |
| Isoleucine | 92.2 ± 1.7 | 85.6 ± 0.6 | 79.0 ± 2.4 |
| Itaconic acid | 104.1 ± 2.8 | 90.0 ± 2.7 | 93.7 ± 0.9 |
| ITP | 97.2 ± 0.6 | 77.1 ± 3.5 | 75.0 ± 3.2 |
| Kynurenic acid | 100.0 ± 0.5 | 94.9 ± 1.6 | 94.9 ± 0.6 |
| Kynurenine | 94.9 ± 2.7 | 75.2 ± 0.9 | 71.4 ± 2.1 |
| Lactic Acid | 99.8 ± 0.9 | 75.8 ± 1.3 | 75.7 ± 3.1 |
| Lactose | 87.7 ± 1.1 | 83.9 ± 1.6 | 73.6 ± 2.5 |
| Leucine | 89.8 ± 2.4 | 91.3 ± 0.5 | 82.0 ± 2.1 |
| Levulinic acid | 93.0 ± 2.8 | 94.7 ± 2.6 | 88.1 ± 2.9 |
| Lysine | 97.1 ± 2.5 | 76.8 ± 1.4 | 74.6 ± 0.5 |
| Malic Acid | 92.9 ± 1.2 | 89.5 ± 2.8 | 83.2 ± 0.7 |
| Malonic acid | 102.9 ± 1.5 | 75.4 ± 2.2 | 77.6 ± 3.3 |
| Mannose | 94.6 ± 0.7 | 75.5 ± 1.4 | 71.4 ± 0.6 |
| Mannose-1 with Mannose-6-phosphate | 92.2 ± 2.6 | 89.2 ± 2.9 | 82.3 ± 0.6 |
| Mesaconic acid | 91.5 ± 2.6 | 77.5 ± 1.5 | 70.9 ± 0.7 |
| Methionine | 88.8 ± 2.5 | 93.5 ± 0.5 | 83.0 ± 1.5 |
| Methionine sulfone | 90.0 ± 2.2 | 89.5 ± 2.6 | 80.6 ± 0.6 |
| Methionine sulfoxide | 89.7 ± 1.2 | 90.1 ± 0.8 | 80.8 ± 2.8 |
| Methylmalonate | 86.9 ± 0.5 | 91.9 ± 2.1 | 79.9 ± 1.1 |
| N-Acetyl-Alanine | 93.2 ± 1.3 | 91.9 ± 1.3 | 85.6 ± 1.4 |
| N-Acetyl-Aspartic Acid | 99.0 ± 3.0 | 78.7 ± 1.1 | 77.9 ± 2.4 |
| N-Acetyl-Citrulline | 100.2 ± 2.3 | 78.5 ± 3.4 | 78.7 ± 2.1 |
| N-Acetyl-Galactosamine | 90.4 ± 3.4 | 79.2 ± 0.5 | 71.6 ± 1.0 |
| N-Acetyl-Glucosamine | 99.9 ± 1.0 | 80.7 ± 1.6 | 80.6 ± 2.4 |
| N-Acetyl-Glutamine | 100.3 ± 2.2 | 89.5 ± 2.0 | 89.8 ± 3.3 |
| N-Acetyl-Glycine | 94.0 ± 1.8 | 82.8 ± 1.7 | 77.8 ± 2.0 |
| N-Acetyl-Leucine | 95.6 ± 0.4 | 93.2 ± 3.2 | 89.1 ± 0.4 |
| N-Acetyl-Methionine | 93.2 ± 1.3 | 77.5 ± 2.5 | 72.2 ± 1.7 |
| N-Acetylneuraminic acid | 102.6 ± 3.3 | 83.8 ± 2.8 | 86.0 ± 1.7 |
| N-Acetyl-Phenylalanine | 90.6 ± 2.8 | 87.1 ± 2.7 | 78.9 ± 1.0 |
| N-Acetyl-Proline | 91.2 ± 0.4 | 82.8 ± 0.9 | 75.5 ± 2.9 |
| N-Acetyl-Serine | 95.3 ± 2.3 | 88.9 ± 1.4 | 84.7 ± 2.2 |
| N-Acetyl-Serotonin | 99.3 ± 1.6 | 89.4 ± 1.1 | 88.8 ± 0.9 |
| N-Acetyl-Tryptophan | 102.9 ± 3.2 | 80.3 ± 1.7 | 82.7 ± 1.2 |
| N-Acetyl-Tyrosine | 93.3 ± 1.4 | 90.9 ± 3.6 | 84.8 ± 2.8 |
| NAD | 95.4 ± 2.0 | 92.6 ± 1.9 | 88.3 ± 1.1 |
| NADH | 99.2 ± 1.3 | 80.3 ± 1.9 | 79.6 ± 2.3 |
| NADP | 102.4 ± 1.3 | 76.7 ± 0.7 | 78.5 ± 3.0 |
| NADPH | 89.0 ± 2.2 | 79.7 ± 0.4 | 70.9 ± 0.4 |
| N-alpha-Acetyl-lysine | 87.4 ± 0.8 | 78.8 ± 3.2 | 68.9 ± 1.9 |
| N-alpha-Acetyl-Ornithine | 99.5 ± 0.9 | 85.2 ± 1.9 | 84.7 ± 2.0 |
| N-epsilon-Acetyl-lysine | 96.2 ± 2.6 | 89.9 ± 3.1 | 86.5 ± 1.0 |
| N-epsilon-Acetyl-Ornithine | 100.8 ± 3.5 | 93.4 ± 2.3 | 94.1 ± 1.4 |
| Nicotinamide | 97.8 ± 1.4 | 91.3 ± 1.0 | 89.3 ± 1.1 |
| Nicotinic acid | 97.2 ± 2.2 | 82.9 ± 2.1 | 80.6 ± 0.8 |
| N-Methyl-Aspartic Acid | 88.0 ± 2.8 | 78.9 ± 1.8 | 69.4 ± 1.0 |
| O-Acetyl-Serine | 94.4 ± 1.7 | 93.9 ± 2.1 | 88.6 ± 3.0 |
| O-Phosphoethanolamine | 97.7 ± 3.1 | 79.9 ± 1.9 | 78.1 ± 1.9 |
| O-Phospho-Serine | 87.3 ± 0.7 | 83.1 ± 1.2 | 72.6 ± 1.1 |
| Ophthalmic acid | 90.3 ± 2.9 | 76.5 ± 3.4 | 69.1 ± 3.2 |
| Ornithine | 99.7 ± 2.6 | 81.6 ± 1.0 | 81.4 ± 1.0 |
| Orotic acid | 98.0 ± 0.6 | 89.2 ± 1.1 | 87.4 ± 1.5 |
| Oxaloacetate | 101.3 ± 2.8 | 79.2 ± 0.7 | 80.2 ± 1.7 |
| Pantothenic acid | 99.9 ± 3.4 | 94.4 ± 3.6 | 94.3 ± 2.6 |
| p-Coumaric Acid | 99.4 ± 1.8 | 90.4 ± 1.6 | 89.8 ± 0.8 |
| Phenylalanine | 99.4 ± 1.5 | 89.1 ± 2.7 | 88.5 ± 0.5 |
| Phenylpyruvate | 89.1 ± 2.1 | 86.8 ± 3.2 | 77.4 ± 1.8 |
| Phosphocreatine | 95.2 ± 2.3 | 81.3 ± 3.0 | 77.4 ± 1.5 |
| Phosphoenolpyruvate | 90.5 ± 2.3 | 89.7 ± 3.0 | 81.2 ± 1.1 |
| Phosphoribosylpyrophosphate | 97.9 ± 1.7 | 94.2 ± 2.3 | 92.2 ± 0.5 |
| Picolinate | 93.1 ± 2.7 | 93.0 ± 0.5 | 86.6 ± 2.7 |
| Pipecolic acid | 91.7 ± 3.1 | 87.4 ± 0.4 | 80.1 ± 3.6 |
| Proline | 95.6 ± 1.3 | 88.4 ± 2.7 | 84.6 ± 1.4 |
| Pyridoxal | 103.7 ± 1.3 | 85.2 ± 0.8 | 88.3 ± 1.3 |
| Pyridoxal-5-phosphate | 99.6 ± 1.6 | 83.7 ± 3.2 | 83.3 ± 2.5 |
| Pyridoxamine | 96.6 ± 3.3 | 77.5 ± 3.6 | 74.8 ± 1.1 |
| Pyridoxine | 94.0 ± 1.1 | 80.4 ± 1.1 | 75.6 ± 3.6 |
| Pyroglutamic acid | 99.8 ± 3.2 | 91.5 ± 2.4 | 91.3 ± 1.0 |
| Pyrrolysine | 97.7 ± 3.1 | 93.2 ± 1.2 | 91.1 ± 3.4 |
| Pyruvic Acid | 96.1 ± 2.8 | 86.6 ± 2.8 | 83.2 ± 2.8 |
| Riboflavin | 90.3 ± 3.6 | 81.0 ± 1.9 | 73.1 ± 0.6 |
| Ribose | 94.2 ± 3.5 | 90.3 ± 2.3 | 85.0 ± 0.6 |
| Ribose-5-phosphate | 99.7 ± 2.6 | 75.2 ± 0.3 | 74.9 ± 1.6 |
| Ribulose-1-phosphate | 99.5 ± 2.9 | 84.5 ± 1.4 | 84.0 ± 1.6 |
| Ribulose-5-phosphate | 95.1 ± 3.6 | 91.0 ± 3.0 | 86.5 ± 2.3 |
| S-Adenosylmethionine | 92.1 ± 2.8 | 82.3 ± 0.8 | 75.8 ± 1.6 |
| Sarcosine | 96.9 ± 0.5 | 76.8 ± 0.6 | 74.4 ± 0.4 |
| Sedoheptulose-7-phosphate | 94.8 ± 0.9 | 89.6 ± 2.8 | 84.9 ± 0.6 |
| Selenocystine | 90.0 ± 1.1 | 89.6 ± 2.9 | 80.7 ± 2.9 |
| Selenomethionine | 88.8 ± 0.4 | 92.6 ± 1.4 | 82.2 ± 0.7 |
| Serine | 97.0 ± 3.0 | 85.9 ± 3.5 | 83.3 ± 2.7 |
| Serotonin | 100.0 ± 1.8 | 83.9 ± 3.3 | 83.9 ± 0.7 |
| Sorbitol | 98.2 ± 1.8 | 84.2 ± 0.9 | 82.7 ± 0.6 |
| Spermidine | 91.6 ± 0.7 | 80.7 ± 1.3 | 73.9 ± 1.0 |
| Spermine | 104.0 ± 2.3 | 75.9 ± 0.5 | 78.9 ± 0.5 |
| Succinic Acid | 98.4 ± 2.1 | 86.3 ± 0.9 | 84.9 ± 1.1 |
| Sucrose | 95.0 ± 0.8 | 78.2 ± 1.8 | 74.3 ± 1.7 |
| Symmetric dimethylarginine | 88.9 ± 1.4 | 79.6 ± 3.1 | 70.7 ± 0.6 |
| Taurine | 97.4 ± 3.5 | 79.2 ± 1.9 | 77.1 ± 2.1 |
| Thiamine | 99.6 ± 1.1 | 92.8 ± 0.6 | 92.4 ± 0.8 |
| Thiamine monophosphate | 97.5 ± 2.4 | 85.0 ± 3.4 | 82.8 ± 1.1 |
| Thiamine pyrophosphate | 99.6 ± 2.0 | 91.7 ± 2.8 | 91.3 ± 2.4 |
| Threonic acid | 89.5 ± 2.8 | 94.7 ± 2.7 | 84.7 ± 2.7 |
| Threonine | 96.1 ± 2.1 | 80.7 ± 1.3 | 77.5 ± 3.3 |
| Thymidine | 89.4 ± 3.4 | 86.4 ± 0.4 | 77.3 ± 3.2 |
| Thymidine Monophosphate | 99.9 ± 1.1 | 78.3 ± 2.9 | 78.2 ± 2.8 |
| Thymine | 87.0 ± 3.2 | 88.5 ± 1.0 | 77.0 ± 1.2 |
| Trimethylamine N-oxide | 103.1 ± 0.8 | 94.9 ± 1.7 | 97.8 ± 3.5 |
| Tryptophan | 90.3 ± 1.2 | 75.5 ± 1.8 | 68.2 ± 0.8 |
| Tyrosine | 89.8 ± 1.7 | 90.9 ± 0.5 | 81.7 ± 2.9 |
| Uracil | 94.9 ± 2.3 | 94.6 ± 1.7 | 89.8 ± 3.1 |
| Uric Acid | 102.5 ± 2.1 | 92.0 ± 3.1 | 94.3 ± 0.5 |
| Uridine | 90.4 ± 3.3 | 89.9 ± 0.5 | 81.3 ± 3.1 |
| Uridine 5'-diphosphate | 88.3 ± 1.2 | 81.0 ± 2.2 | 71.5 ± 1.4 |
| Uridine Monophosphate | 88.9 ± 1.0 | 85.7 ± 0.6 | 76.2 ± 1.2 |
| Uridine-5-Diphosphogalactose | 97.5 ± 0.9 | 93.1 ± 2.2 | 90.7 ± 1.6 |
| Uridine-5-Diphosphoglucose | 101.8 ± 2.9 | 83.7 ± 3.1 | 85.2 ± 2.1 |
| Urocanic acid | 103.7 ± 1.1 | 82.0 ± 1.1 | 85.0 ± 0.9 |
| UTP | 98.0 ± 3.2 | 85.8 ± 1.8 | 84.1 ± 1.9 |
| Valine | 91.0 ± 1.7 | 91.6 ± 2.9 | 83.3 ± 3.4 |
| Xanthine | 102.3 ± 1.7 | 91.5 ± 2.9 | 93.6 ± 2.3 |
| Xanthosine | 100.1 ± 3.1 | 80.7 ± 1.6 | 80.8 ± 3.3 |
| Xanthurenic acid | 95.3 ± 2.2 | 83.9 ± 2.8 | 79.9 ± 1.6 |
| XMP | 98.3 ± 2.8 | 94.0 ± 0.6 | 92.4 ± 1.3 |
| Xylulose-5-phosphate | 94.6 ± 0.9 | 87.5 ± 3.3 | 82.8 ± 0.8 |

### **Table S7** | Calibration values for final method conditions (r^2^-value) and percent found of nominal calibrator concentration (Colorectal cancer organoid as matrix).

| Calibrator concentration accuracy  [%] | 10 ng/mL | 25 ng/mL | 50 ng/mL | 100 ng/mL | 500 ng/mL | 1000 ng/mL | r^2^-Value  matrix-matched calibration |
| --- | --- | --- | --- | --- | --- | --- | --- |
| 1-Methylguanine | 89.3 ± 2.2 | 93.3 ± 0.8 | 104.6 ± 2.9 | 108.5 ± 1.8 | 105.1 ± 0.7 | 113.2 ± 3.0 | 0.9996 |
| 1-Methylhistidine | 104.2 ± 2.6 | 102.3 ± 1.5 | 106.5 ± 0.9 | 96.0 ± 2.8 | 106.6 ± 0.4 | 101.3 ± 0.8 | 0.9994 |
| 1-Methylnicotinamide | 85.5 ± 0.6 | 88.0 ± 1.1 | 101.9 ± 1.5 | 95.2 ± 2.7 | 100.6 ± 2.8 | 91.4 ± 2.5 | 0.9992 |
| 2-Aminoadipic Acid | 94.5 ± 2.6 | 91.8 ± 2.5 | 94.8 ± 2.4 | 112.7 ± 0.4 | 98.5 ± 0.3 | 109.9 ± 3.0 | 0.9996 |
| 2-Deoxyadenosine | 85.0 ± 0.3 | 109.1 ± 1.2 | 107.5 ± 1.5 | 93.5 ± 2.5 | 95.2 ± 2.2 | 97.2 ± 1.8 | 0.9996 |
| 2'-Deoxyadenosine 5'-monophosphate | 101.0 ± 1.2 | 85.7 ± 2.1 | 105.9 ± 0.6 | 92.7 ± 0.7 | 90.9 ± 2.3 | 89.1 ± 1.6 | 0.9992 |
| 2'-deoxyadenosine 5'-triphosphate | 97.5 ± 1.1 | 114.6 ± 0.4 | 106.0 ± 1.0 | 88.4 ± 1.3 | 91.7 ± 3.2 | 105.0 ± 2.7 | 0.9995 |
| 2'-Deoxycytidine-5'-monophosphate | 91.2 ± 1.7 | 89.9 ± 0.7 | 100.9 ± 0.8 | 104.7 ± 2.2 | 109.2 ± 0.7 | 91.9 ± 0.9 | 0.9993 |
| 2-Deoxyguanosine | 90.3 ± 2.7 | 94.5 ± 1.6 | 99.4 ± 2.3 | 106.9 ± 2.6 | 86.0 ± 1.0 | 94.7 ± 2.8 | 0.9996 |
| 2-hydroxybutyric acid | 100.6 ± 0.9 | 88.9 ± 0.5 | 86.9 ± 0.8 | 89.5 ± 3.1 | 109.4 ± 2.9 | 90.0 ± 2.7 | 0.9991 |
| 2-hydroxyglutaric acid | 113.1 ± 2.2 | 96.2 ± 3.1 | 90.1 ± 0.6 | 87.7 ± 1.8 | 94.5 ± 0.9 | 104.5 ± 1.1 | 0.9995 |
| 2-Hydroxyisocaproic acid | 88.0 ± 3.1 | 90.5 ± 1.4 | 114.6 ± 1.3 | 109.5 ± 3.1 | 109.8 ± 2.4 | 101.7 ± 0.6 | 0.9998 |
| 2-ketobutyric acid | 98.6 ± 3.0 | 97.8 ± 1.4 | 105.6 ± 2.2 | 109.6 ± 2.2 | 93.5 ± 1.3 | 104.6 ± 3.0 | 0.9991 |
| 2-oxoadipic acid | 86.5 ± 0.5 | 93.5 ± 2.2 | 110.3 ± 1.8 | 90.6 ± 0.4 | 96.3 ± 2.2 | 92.8 ± 0.8 | 0.9995 |
| 2-Phenylglycine | 106.5 ± 2.6 | 90.1 ± 0.9 | 107.5 ± 0.8 | 104.2 ± 1.1 | 104.4 ± 2.4 | 91.9 ± 2.5 | 0.9994 |
| 2-phosphoglycerate | 112.1 ± 1.2 | 100.2 ± 1.1 | 100.8 ± 2.5 | 96.0 ± 3.2 | 96.0 ± 3.1 | 108.5 ± 0.6 | 0.9992 |
| 3-hydroxy-3-methyl-2-oxobutanoate | 104.3 ± 2.6 | 102.1 ± 0.7 | 95.7 ± 2.5 | 90.5 ± 0.9 | 113.9 ± 1.1 | 103.4 ± 1.6 | 0.9997 |
| 3-hydroxybutyric acid | 100.9 ± 2.4 | 95.5 ± 1.1 | 86.2 ± 3.1 | 96.9 ± 0.9 | 102.6 ± 2.4 | 110.1 ± 2.9 | 0.9993 |
| 3-Methylhistidine | 108.2 ± 2.1 | 93.3 ± 1.4 | 93.9 ± 1.8 | 109.3 ± 1.4 | 103.4 ± 1.8 | 107.6 ± 0.6 | 0.9999 |
| 3-Phosphoglyceraldehyde | 109.2 ± 0.8 | 108.0 ± 2.8 | 99.5 ± 2.9 | 108.2 ± 2.2 | 90.5 ± 1.4 | 85.9 ± 1.1 | 0.9992 |
| 3-phosphoglycerate | 89.4 ± 2.5 | 92.8 ± 0.7 | 89.1 ± 1.0 | 112.4 ± 2.1 | 105.7 ± 0.3 | 108.4 ± 0.4 | 0.9996 |
| 4-amino-butyric acid | 109.6 ± 2.5 | 99.6 ± 0.8 | 96.1 ± 2.3 | 95.7 ± 0.5 | 103.2 ± 2.0 | 93.6 ± 1.2 | 0.9997 |
| 4-aspartyl-phosphate | 87.9 ± 1.9 | 92.7 ± 0.9 | 107.8 ± 1.4 | 95.5 ± 1.3 | 85.5 ± 2.6 | 106.3 ± 1.2 | 0.9997 |
| 4-Hydroxybenzoic acid | 107.8 ± 1.5 | 114.2 ± 2.6 | 110.0 ± 0.9 | 107.1 ± 3.0 | 105.1 ± 2.4 | 107.0 ± 1.6 | 0.9998 |
| 4-Hydroxyphenylacetylglycine | 100.4 ± 3.2 | 93.5 ± 2.3 | 86.0 ± 1.6 | 89.3 ± 1.6 | 99.5 ± 1.5 | 91.6 ± 1.9 | 0.9995 |
| 4-methyl-2-oxopentanoate | 97.0 ± 1.2 | 108.8 ± 0.8 | 113.9 ± 2.5 | 108.4 ± 2.9 | 107.5 ± 0.5 | 108.6 ± 2.7 | 0.9992 |
| 4-Oxoglutaramate | 109.4 ± 0.9 | 86.6 ± 1.3 | 97.6 ± 1.6 | 98.3 ± 1.5 | 94.6 ± 2.6 | 92.8 ± 2.0 | 0.9999 |
| 5-Hydroxyindolepyruvate | 108.7 ± 0.8 | 100.4 ± 0.8 | 90.0 ± 1.5 | 113.9 ± 2.2 | 92.6 ± 1.9 | 94.6 ± 2.7 | 0.9995 |
| 5-Hydroxylysine | 102.1 ± 1.0 | 99.8 ± 2.3 | 85.5 ± 2.5 | 103.4 ± 0.3 | 87.6 ± 0.8 | 104.3 ± 2.2 | 0.9998 |
| 5-Hydroxy-Tryptophan | 108.9 ± 2.1 | 85.8 ± 2.5 | 85.6 ± 0.9 | 113.0 ± 2.5 | 86.7 ± 1.0 | 92.3 ± 3.1 | 0.9998 |
| 5-Phosphoribosylamine | 109.1 ± 1.8 | 88.2 ± 3.1 | 102.3 ± 2.1 | 94.8 ± 1.9 | 95.5 ± 1.1 | 113.4 ± 2.1 | 0.9997 |
| 6-phosphogluconate | 95.4 ± 1.2 | 96.5 ± 2.5 | 87.4 ± 0.7 | 114.7 ± 1.8 | 109.8 ± 2.5 | 110.0 ± 2.8 | 0.9997 |
| Acetylcarnitine | 96.5 ± 3.0 | 97.4 ± 3.1 | 113.5 ± 1.1 | 93.4 ± 1.8 | 87.5 ± 1.8 | 112.0 ± 1.4 | 0.9999 |
| Aconitic Acid | 102.6 ± 1.6 | 90.8 ± 0.7 | 89.1 ± 1.1 | 85.3 ± 1.3 | 111.5 ± 1.1 | 105.5 ± 1.9 | 0.9997 |
| Adenine | 89.9 ± 1.5 | 97.6 ± 2.2 | 97.3 ± 0.5 | 99.1 ± 2.5 | 102.6 ± 2.1 | 88.1 ± 2.4 | 0.9998 |
| Adenosine | 89.5 ± 1.9 | 90.6 ± 2.9 | 103.7 ± 0.4 | 97.9 ± 2.4 | 86.6 ± 1.2 | 113.7 ± 2.0 | 0.9993 |
| Adenosine 3',5'-cyclic monophosphate | 103.8 ± 1.9 | 111.9 ± 1.1 | 114.5 ± 0.3 | 85.0 ± 2.6 | 110.9 ± 2.1 | 92.6 ± 2.7 | 0.9999 |
| Adenosine Monophosphate | 113.1 ± 2.8 | 107.5 ± 1.7 | 92.4 ± 0.6 | 90.3 ± 1.2 | 92.7 ± 0.5 | 102.7 ± 1.3 | 0.9994 |
| Adenosine-5'-diphosphate | 104.6 ± 1.1 | 86.6 ± 2.4 | 112.8 ± 0.5 | 101.8 ± 3.1 | 98.3 ± 2.8 | 102.3 ± 1.3 | 0.9994 |
| Adenosine-5'-triphosphate | 98.1 ± 2.5 | 101.5 ± 2.7 | 103.9 ± 2.6 | 95.6 ± 1.5 | 106.7 ± 2.3 | 88.1 ± 2.4 | 0.9994 |
| Adenylosuccinate | 90.3 ± 3.1 | 94.0 ± 2.1 | 87.9 ± 1.0 | 93.5 ± 3.0 | 94.2 ± 1.0 | 89.2 ± 1.4 | 0.9992 |
| Alanine | 99.6 ± 2.3 | 92.9 ± 3.0 | 105.7 ± 3.0 | 96.1 ± 0.5 | 99.2 ± 1.6 | 90.0 ± 0.6 | 0.9995 |
| alpha-ketoglutaric acid | 100.1 ± 0.3 | 114.1 ± 1.2 | 102.5 ± 1.0 | 88.0 ± 0.3 | 101.4 ± 2.7 | 96.4 ± 2.5 | 0.9993 |
| Anserine | 103.3 ± 3.0 | 92.8 ± 1.0 | 114.3 ± 2.3 | 108.2 ± 2.5 | 109.2 ± 2.6 | 104.8 ± 1.7 | 0.9995 |
| Arginine | 102.9 ± 2.7 | 89.5 ± 2.5 | 103.4 ± 1.0 | 101.5 ± 1.3 | 111.4 ± 2.6 | 110.3 ± 0.8 | 0.9994 |
| Argininosuccinic Acid | 112.1 ± 0.7 | 108.4 ± 2.5 | 106.4 ± 1.1 | 113.7 ± 2.6 | 110.4 ± 1.8 | 107.8 ± 0.8 | 0.9993 |
| Asparagine | 85.5 ± 2.5 | 98.4 ± 1.4 | 89.4 ± 2.4 | 85.1 ± 3.0 | 98.3 ± 1.5 | 89.5 ± 0.5 | 0.9997 |
| Aspartic acid | 88.3 ± 1.0 | 103.5 ± 3.0 | 108.1 ± 0.8 | 87.9 ± 2.2 | 98.7 ± 2.8 | 100.9 ± 1.7 | 0.9993 |
| Asymmetric dimethylarginine | 98.5 ± 2.1 | 108.8 ± 3.2 | 112.9 ± 2.2 | 107.1 ± 3.2 | 94.5 ± 2.5 | 94.0 ± 0.5 | 0.9998 |
| Azelaic acid | 86.1 ± 2.4 | 96.9 ± 1.3 | 107.0 ± 1.8 | 96.8 ± 0.8 | 96.2 ± 2.4 | 85.9 ± 0.7 | 0.9993 |
| beta-Alanine | 86.0 ± 0.6 | 99.9 ± 2.8 | 111.2 ± 3.0 | 95.2 ± 0.5 | 93.0 ± 2.5 | 112.8 ± 2.8 | 0.9991 |
| Betaine | 107.9 ± 1.6 | 92.2 ± 2.0 | 95.1 ± 0.8 | 99.9 ± 2.2 | 113.6 ± 1.1 | 90.1 ± 0.6 | 0.9996 |
| Biopterin | 112.3 ± 0.9 | 92.1 ± 3.0 | 102.2 ± 1.7 | 88.5 ± 2.5 | 103.6 ± 0.5 | 106.3 ± 0.3 | 0.9992 |
| Biotin | 110.0 ± 0.4 | 98.1 ± 0.4 | 104.7 ± 1.4 | 104.4 ± 0.6 | 92.3 ± 3.2 | 104.2 ± 2.2 | 0.9991 |
| Carnitine | 104.0 ± 1.8 | 92.5 ± 0.7 | 90.8 ± 2.5 | 90.3 ± 2.1 | 97.0 ± 1.5 | 108.1 ± 0.7 | 0.9992 |
| Carnosine | 110.7 ± 0.4 | 85.7 ± 1.6 | 97.8 ± 2.0 | 114.3 ± 1.4 | 102.7 ± 0.9 | 111.8 ± 1.7 | 0.9998 |
| CDP-choline | 92.9 ± 0.8 | 109.5 ± 0.6 | 109.2 ± 1.7 | 105.9 ± 3.1 | 114.2 ± 0.5 | 95.1 ± 1.6 | 0.9994 |
| cGMP | 89.3 ± 0.4 | 114.7 ± 2.1 | 110.3 ± 2.0 | 89.5 ± 1.7 | 100.4 ± 2.2 | 93.5 ± 0.7 | 0.9991 |
| Choline | 104.7 ± 2.3 | 98.9 ± 1.5 | 85.3 ± 2.6 | 96.4 ± 1.0 | 113.8 ± 0.3 | 99.7 ± 0.9 | 0.9992 |
| Cinnamic acid | 108.4 ± 1.2 | 112.3 ± 2.8 | 94.8 ± 0.7 | 100.9 ± 2.0 | 114.7 ± 1.2 | 92.8 ± 2.7 | 0.9998 |
| Citraconic acid | 85.0 ± 1.9 | 98.0 ± 3.0 | 105.7 ± 1.6 | 99.4 ± 2.3 | 105.4 ± 1.7 | 112.5 ± 1.2 | 0.9995 |
| Citramalic acid | 110.4 ± 2.9 | 86.7 ± 2.4 | 98.2 ± 1.8 | 103.7 ± 0.9 | 86.7 ± 3.0 | 86.0 ± 3.2 | 0.9992 |
| Citric Acid | 106.0 ± 2.1 | 111.3 ± 2.3 | 96.3 ± 1.1 | 89.3 ± 0.4 | 85.1 ± 2.9 | 92.8 ± 0.4 | 0.9997 |
| Citrulline | 98.5 ± 0.6 | 89.1 ± 1.1 | 102.9 ± 2.9 | 92.1 ± 1.5 | 102.3 ± 0.7 | 105.8 ± 2.1 | 0.9993 |
| Creatine | 89.4 ± 0.4 | 113.7 ± 2.1 | 111.5 ± 1.9 | 105.8 ± 1.1 | 90.0 ± 2.8 | 86.4 ± 2.6 | 0.9994 |
| Creatinine | 110.9 ± 2.4 | 93.3 ± 2.5 | 93.5 ± 2.1 | 93.7 ± 1.9 | 97.6 ± 2.7 | 94.2 ± 0.4 | 0.9993 |
| Cystathionine | 95.4 ± 2.1 | 91.2 ± 2.3 | 113.0 ± 0.8 | 114.7 ± 2.0 | 107.1 ± 1.8 | 112.9 ± 3.2 | 0.9996 |
| Cysteamine | 105.6 ± 2.9 | 104.5 ± 3.1 | 92.0 ± 0.8 | 88.6 ± 1.0 | 106.9 ± 2.8 | 100.1 ± 1.6 | 0.9994 |
| Cysteine | 88.0 ± 0.4 | 94.2 ± 3.2 | 103.1 ± 1.8 | 108.6 ± 2.2 | 87.2 ± 1.8 | 103.5 ± 0.8 | 0.9992 |
| Cysteinylglycine | 92.1 ± 1.9 | 110.6 ± 2.2 | 113.4 ± 1.5 | 105.9 ± 1.4 | 107.3 ± 1.3 | 86.7 ± 0.7 | 0.9994 |
| Cystine | 104.7 ± 1.2 | 97.1 ± 0.4 | 103.8 ± 2.4 | 101.8 ± 2.2 | 112.9 ± 1.1 | 89.1 ± 0.7 | 0.9996 |
| Cytidine | 110.7 ± 2.8 | 105.7 ± 1.9 | 98.6 ± 0.8 | 114.4 ± 1.4 | 95.6 ± 3.0 | 110.2 ± 2.2 | 0.9994 |
| Cytidine 3',5'-cyclic monophosphate | 89.3 ± 1.1 | 107.2 ± 3.1 | 112.6 ± 1.2 | 85.0 ± 0.8 | 95.3 ± 0.8 | 109.5 ± 3.2 | 0.9992 |
| Cytidine 5'-diphosphate | 96.2 ± 1.1 | 102.5 ± 2.4 | 89.1 ± 0.6 | 99.0 ± 3.1 | 97.2 ± 3.1 | 109.0 ± 0.8 | 0.9994 |
| Cytidine 5'-triphosphate | 93.2 ± 2.5 | 100.8 ± 0.8 | 110.3 ± 3.0 | 98.5 ± 0.3 | 104.4 ± 2.8 | 113.2 ± 3.2 | 0.9998 |
| Cytidine monophosphate | 104.0 ± 2.1 | 101.8 ± 3.0 | 97.2 ± 1.0 | 88.0 ± 2.3 | 114.3 ± 0.6 | 88.2 ± 1.6 | 0.9999 |
| Cytosine | 114.4 ± 2.4 | 113.2 ± 1.0 | 90.5 ± 1.4 | 95.1 ± 2.1 | 100.7 ± 2.8 | 105.4 ± 2.8 | 0.9994 |
| dADP | 96.3 ± 0.5 | 108.8 ± 1.6 | 104.6 ± 2.7 | 106.5 ± 0.8 | 108.1 ± 2.9 | 107.7 ± 0.8 | 0.9999 |
| Diethanolamine | 87.7 ± 0.8 | 92.9 ± 1.8 | 99.7 ± 0.6 | 111.2 ± 1.1 | 92.6 ± 2.0 | 88.9 ± 1.9 | 0.9994 |
| Dihydrobiopterin | 92.1 ± 3.1 | 112.8 ± 0.6 | 99.0 ± 1.0 | 105.6 ± 2.5 | 102.6 ± 1.9 | 93.9 ± 3.0 | 0.9994 |
| Dihydroneopterin triphosphate | 104.3 ± 2.5 | 112.2 ± 0.3 | 106.3 ± 0.7 | 112.6 ± 0.3 | 86.0 ± 1.5 | 113.7 ± 1.4 | 0.9994 |
| Dihydroxyacetonephosphate | 107.0 ± 0.4 | 95.7 ± 0.4 | 106.9 ± 2.7 | 102.9 ± 2.6 | 87.2 ± 2.5 | 113.1 ± 0.3 | 0.9994 |
| Dimethylglycine | 88.6 ± 2.2 | 93.7 ± 2.5 | 114.6 ± 3.2 | 86.0 ± 2.0 | 101.3 ± 2.2 | 108.6 ± 2.9 | 0.9997 |
| Ectoine | 109.7 ± 2.4 | 87.3 ± 1.3 | 92.4 ± 2.9 | 88.3 ± 0.5 | 99.1 ± 0.9 | 90.3 ± 0.5 | 0.9998 |
| Erythrose-4-phosphate | 109.8 ± 1.7 | 106.4 ± 0.4 | 95.5 ± 0.5 | 91.7 ± 1.2 | 85.3 ± 1.3 | 102.4 ± 0.7 | 0.9995 |
| FAD | 112.0 ± 2.8 | 110.5 ± 1.2 | 98.5 ± 2.6 | 104.3 ± 2.0 | 114.8 ± 0.6 | 94.5 ± 1.9 | 0.9996 |
| Folic acid | 93.1 ± 0.8 | 94.3 ± 1.7 | 92.6 ± 2.6 | 94.1 ± 1.1 | 94.1 ± 1.9 | 105.7 ± 1.1 | 0.9995 |
| Fructose-6-phosphate | 95.6 ± 2.5 | 100.5 ± 3.1 | 103.2 ± 2.2 | 86.0 ± 2.2 | 87.5 ± 2.0 | 107.6 ± 0.6 | 0.9997 |
| Fumaric Acid | 113.8 ± 3.2 | 112.7 ± 0.6 | 103.5 ± 3.0 | 113.2 ± 1.0 | 100.9 ± 2.0 | 99.4 ± 2.3 | 0.9991 |
| Galactosamine | 101.5 ± 1.3 | 112.7 ± 2.2 | 87.9 ± 2.6 | 99.6 ± 2.1 | 86.7 ± 0.8 | 87.0 ± 2.8 | 0.9999 |
| Galactose-1-phosphate | 99.2 ± 2.4 | 108.2 ± 1.7 | 111.6 ± 2.3 | 96.4 ± 3.0 | 108.1 ± 0.5 | 89.2 ± 2.8 | 0.9995 |
| Galactose-6-Phosphate | 110.0 ± 2.9 | 110.1 ± 1.5 | 91.7 ± 2.9 | 102.3 ± 2.0 | 109.4 ± 0.6 | 95.0 ± 2.7 | 0.9996 |
| gamma-Glutamylcysteine | 105.0 ± 2.3 | 105.4 ± 1.9 | 104.1 ± 1.9 | 85.2 ± 2.1 | 86.8 ± 1.6 | 107.3 ± 2.7 | 0.9993 |
| Glucosamine | 113.9 ± 0.4 | 89.3 ± 3.1 | 101.9 ± 2.0 | 108.7 ± 1.4 | 99.5 ± 2.8 | 110.7 ± 2.7 | 0.9991 |
| Glucosamine-6-Phosphate | 90.3 ± 0.7 | 90.5 ± 2.9 | 90.2 ± 0.6 | 99.7 ± 1.3 | 111.1 ± 2.8 | 108.7 ± 1.8 | 0.9995 |
| Glucose | 110.8 ± 1.8 | 113.1 ± 2.1 | 107.6 ± 1.1 | 98.6 ± 2.9 | 87.8 ± 0.4 | 113.7 ± 2.0 | 0.9994 |
| Glucose-1-phosphate | 112.9 ± 1.3 | 87.9 ± 0.3 | 109.6 ± 0.8 | 89.3 ± 2.8 | 112.6 ± 1.9 | 114.2 ± 1.6 | 0.9998 |
| Glucose-6-Phosphate | 107.7 ± 1.5 | 90.1 ± 2.3 | 107.9 ± 0.6 | 88.8 ± 3.1 | 86.5 ± 2.3 | 109.0 ± 1.3 | 0.9997 |
| Glucuronic acid | 105.7 ± 0.4 | 87.9 ± 2.7 | 91.8 ± 1.2 | 91.7 ± 0.6 | 109.8 ± 2.6 | 87.4 ± 2.1 | 0.9998 |
| Glutamic acid | 96.5 ± 2.1 | 106.8 ± 1.9 | 95.5 ± 1.1 | 114.5 ± 0.6 | 93.4 ± 2.2 | 93.9 ± 1.6 | 0.9994 |
| Glutamine | 112.3 ± 1.0 | 91.2 ± 0.5 | 112.2 ± 1.0 | 88.6 ± 2.5 | 114.5 ± 2.5 | 97.5 ± 2.2 | 0.9992 |
| Glutathione | 101.2 ± 1.1 | 88.6 ± 1.5 | 109.6 ± 1.3 | 91.1 ± 1.5 | 88.8 ± 1.6 | 103.1 ± 2.9 | 0.9991 |
| Glutathionylspermine | 100.5 ± 2.4 | 114.0 ± 2.8 | 86.7 ± 1.8 | 88.3 ± 2.1 | 88.1 ± 1.5 | 88.0 ± 0.7 | 0.9997 |
| Glycerophosphocholine | 85.8 ± 1.8 | 95.8 ± 2.6 | 105.5 ± 1.6 | 114.4 ± 3.2 | 105.2 ± 3.1 | 88.7 ± 0.7 | 0.9997 |
| Glycine | 105.0 ± 1.3 | 107.1 ± 2.0 | 101.0 ± 1.6 | 97.8 ± 2.0 | 99.5 ± 0.6 | 96.5 ± 0.6 | 0.9996 |
| Glycolic acid | 98.9 ± 1.8 | 86.3 ± 0.7 | 86.7 ± 2.8 | 94.9 ± 2.2 | 109.5 ± 0.6 | 97.6 ± 0.4 | 0.9994 |
| Guanidinoacetic acid | 101.7 ± 2.9 | 112.2 ± 0.7 | 106.0 ± 1.7 | 113.5 ± 0.9 | 100.2 ± 0.7 | 96.9 ± 2.5 | 0.9992 |
| Guanidinobutanoic acid | 89.6 ± 0.4 | 88.1 ± 1.4 | 108.7 ± 0.6 | 99.0 ± 0.7 | 111.6 ± 2.4 | 96.5 ± 2.2 | 0.9997 |
| Guanine | 103.3 ± 0.9 | 92.8 ± 3.1 | 88.3 ± 1.2 | 85.4 ± 3.0 | 90.3 ± 2.9 | 114.9 ± 0.5 | 0.9995 |
| Guanosine | 100.1 ± 3.1 | 91.8 ± 2.1 | 95.4 ± 2.4 | 90.9 ± 2.0 | 111.4 ± 2.2 | 102.6 ± 1.3 | 0.9992 |
| Guanosine 5'-diphosphate | 114.0 ± 3.2 | 99.8 ± 2.2 | 90.6 ± 1.7 | 106.6 ± 1.5 | 92.4 ± 1.8 | 95.5 ± 0.6 | 0.9995 |
| Guanosine-5'-monophosphate | 102.2 ± 1.8 | 101.5 ± 2.4 | 104.5 ± 0.4 | 114.0 ± 1.9 | 91.7 ± 1.7 | 103.0 ± 0.9 | 0.9991 |
| Hercynine | 113.1 ± 2.2 | 102.2 ± 2.8 | 114.7 ± 2.8 | 93.1 ± 3.0 | 96.0 ± 1.2 | 110.5 ± 2.6 | 0.9993 |
| Hippuric acid | 97.0 ± 3.2 | 96.0 ± 1.3 | 101.3 ± 1.8 | 108.7 ± 0.8 | 112.7 ± 3.0 | 86.3 ± 2.6 | 0.9992 |
| Histamine | 105.7 ± 0.8 | 102.7 ± 1.3 | 97.6 ± 2.8 | 111.9 ± 2.2 | 110.7 ± 1.5 | 97.1 ± 2.8 | 0.9993 |
| Histidine | 104.0 ± 1.4 | 94.2 ± 1.9 | 109.7 ± 2.7 | 105.8 ± 1.5 | 96.4 ± 1.5 | 105.8 ± 2.2 | 0.9991 |
| Homoalanine | 109.5 ± 3.0 | 105.8 ± 2.1 | 114.5 ± 2.6 | 88.0 ± 0.5 | 100.5 ± 2.5 | 114.1 ± 2.9 | 0.9994 |
| Homoarginine | 111.7 ± 1.5 | 94.4 ± 2.2 | 105.2 ± 0.9 | 89.7 ± 1.2 | 105.0 ± 3.0 | 91.6 ± 0.9 | 0.9996 |
| Homocysteine | 107.9 ± 2.0 | 104.6 ± 1.7 | 103.2 ± 1.5 | 104.2 ± 3.1 | 100.1 ± 2.7 | 86.3 ± 2.4 | 0.9992 |
| Homogentisic acid | 100.5 ± 2.4 | 106.5 ± 1.4 | 96.3 ± 0.4 | 110.9 ± 2.2 | 98.7 ± 2.6 | 111.5 ± 2.1 | 0.9992 |
| Homoleucine | 100.0 ± 3.0 | 100.7 ± 1.3 | 105.4 ± 1.7 | 87.4 ± 2.2 | 110.8 ± 3.2 | 113.5 ± 1.5 | 0.9994 |
| Homovanillic acid | 104.5 ± 2.6 | 101.0 ± 0.3 | 98.3 ± 0.8 | 110.2 ± 0.6 | 98.2 ± 2.7 | 113.4 ± 2.1 | 0.9991 |
| Hydantoin-5-propionate | 97.2 ± 2.1 | 95.7 ± 2.2 | 106.1 ± 0.4 | 100.6 ± 1.1 | 110.4 ± 2.2 | 98.3 ± 3.0 | 0.9995 |
| Hypotaurine | 102.3 ± 1.3 | 113.0 ± 2.6 | 104.1 ± 1.7 | 112.0 ± 3.2 | 90.3 ± 1.0 | 113.8 ± 1.0 | 0.9997 |
| Hypoxanthine | 114.2 ± 2.2 | 95.0 ± 0.7 | 109.8 ± 1.7 | 110.1 ± 0.9 | 90.7 ± 0.8 | 110.2 ± 1.5 | 0.9997 |
| Indole-3-acetamide | 97.7 ± 1.8 | 112.2 ± 2.4 | 112.8 ± 1.0 | 103.0 ± 0.9 | 113.9 ± 2.5 | 93.2 ± 1.6 | 0.9996 |
| Inosine | 108.4 ± 0.4 | 90.1 ± 0.5 | 91.4 ± 1.8 | 109.9 ± 3.1 | 101.9 ± 0.4 | 99.2 ± 1.7 | 0.9991 |
| Inosine 5'-monophosphate | 106.2 ± 1.9 | 85.2 ± 1.8 | 106.4 ± 1.2 | 112.2 ± 1.5 | 105.5 ± 1.5 | 91.1 ± 1.8 | 0.9994 |
| Isocitric Acid | 100.9 ± 1.6 | 112.3 ± 1.7 | 94.4 ± 0.5 | 113.8 ± 2.9 | 94.5 ± 2.4 | 92.9 ± 2.7 | 0.9993 |
| Isoleucine | 95.1 ± 2.0 | 97.2 ± 1.7 | 90.7 ± 1.1 | 101.8 ± 2.5 | 98.0 ± 2.3 | 87.5 ± 0.5 | 0.9993 |
| Lactic Acid | 112.6 ± 1.8 | 104.3 ± 2.2 | 105.0 ± 1.9 | 112.4 ± 2.4 | 114.7 ± 1.1 | 112.7 ± 3.2 | 0.9992 |
| Leucine | 112.3 ± 1.3 | 113.8 ± 1.2 | 105.6 ± 2.2 | 99.7 ± 2.7 | 107.1 ± 3.2 | 90.9 ± 2.6 | 0.9996 |
| Levulinic acid | 95.0 ± 1.3 | 107.8 ± 1.9 | 89.9 ± 0.8 | 101.8 ± 2.2 | 101.9 ± 1.4 | 100.2 ± 2.4 | 0.9996 |
| Lysine | 102.7 ± 1.5 | 107.7 ± 0.3 | 103.3 ± 2.2 | 113.2 ± 0.9 | 105.7 ± 1.5 | 92.2 ± 0.9 | 0.9992 |
| Malic Acid | 90.8 ± 2.2 | 92.6 ± 0.3 | 88.8 ± 1.9 | 87.9 ± 2.7 | 110.3 ± 0.9 | 114.3 ± 1.4 | 0.9998 |
| Mannose-1 with Mannose-6-Phosphate | 103.3 ± 2.2 | 93.4 ± 1.8 | 101.1 ± 2.7 | 101.4 ± 2.6 | 109.2 ± 2.6 | 103.2 ± 0.9 | 0.9997 |
| Mesaconic acid | 93.2 ± 0.5 | 90.2 ± 1.3 | 95.5 ± 1.6 | 89.3 ± 2.6 | 114.0 ± 1.2 | 88.6 ± 2.1 | 0.9992 |
| Methionine | 88.1 ± 2.7 | 90.5 ± 2.4 | 97.9 ± 3.0 | 100.3 ± 2.3 | 97.1 ± 1.2 | 104.8 ± 1.6 | 0.9994 |
| Methionine sulfoxide | 111.1 ± 2.8 | 111.2 ± 1.6 | 101.9 ± 2.5 | 113.0 ± 0.9 | 85.8 ± 2.3 | 86.8 ± 2.2 | 0.9993 |
| Methylmalonic Acid | 114.0 ± 0.9 | 100.9 ± 3.2 | 105.1 ± 2.6 | 93.1 ± 2.4 | 96.4 ± 1.1 | 98.1 ± 2.7 | 0.9992 |
| N1-Acetylspermidine | 99.4 ± 2.3 | 93.0 ± 0.5 | 91.0 ± 1.5 | 93.5 ± 0.8 | 101.6 ± 1.1 | 88.8 ± 1.3 | 0.9997 |
| N1-Acetylspermine | 89.1 ± 1.5 | 105.9 ± 0.9 | 98.0 ± 2.9 | 108.3 ± 1.8 | 93.5 ± 2.5 | 102.7 ± 0.9 | 0.9999 |
| N8-Acetylspermidine | 110.5 ± 0.8 | 92.4 ± 2.4 | 106.7 ± 1.7 | 111.4 ± 0.7 | 110.0 ± 3.1 | 113.7 ± 2.9 | 0.9993 |
| N-Acetyl-Aspartyl-Glutamate | 85.4 ± 1.3 | 100.6 ± 2.8 | 100.8 ± 0.7 | 95.6 ± 1.2 | 114.9 ± 1.0 | 86.0 ± 0.6 | 0.9998 |
| N-Acetyl-Aspartyl-Glutamyl-Glutamate | 105.5 ± 2.7 | 106.2 ± 0.8 | 98.4 ± 2.0 | 86.1 ± 2.2 | 97.2 ± 2.8 | 110.3 ± 1.6 | 0.9998 |
| N-Acetyl-Galactosamine | 94.7 ± 3.0 | 99.3 ± 1.9 | 108.1 ± 2.3 | 100.0 ± 1.8 | 102.6 ± 2.7 | 104.4 ± 0.8 | 0.9991 |
| N-Acetyl-Leucine | 85.8 ± 2.6 | 88.3 ± 1.6 | 108.3 ± 1.9 | 96.1 ± 1.7 | 93.0 ± 0.8 | 107.5 ± 0.7 | 0.9994 |
| N-Acetyl-Methionine | 100.9 ± 0.7 | 87.8 ± 1.4 | 108.2 ± 2.5 | 113.1 ± 1.0 | 113.6 ± 2.1 | 94.3 ± 0.3 | 0.9997 |
| N-Acetylneuraminic acid | 88.9 ± 2.1 | 96.3 ± 2.7 | 115.0 ± 1.1 | 99.5 ± 3.0 | 102.4 ± 0.3 | 90.7 ± 0.5 | 0.9993 |
| N-Acetyl-Proline | 92.0 ± 2.5 | 97.0 ± 3.2 | 109.2 ± 3.0 | 86.1 ± 1.8 | 91.0 ± 0.9 | 105.0 ± 2.1 | 0.9992 |
| N-Acetyl-Putrescine | 108.6 ± 1.3 | 102.4 ± 2.2 | 102.7 ± 0.4 | 113.7 ± 3.0 | 96.2 ± 2.1 | 107.2 ± 3.1 | 0.9995 |
| N-Acetyl-Serine | 109.4 ± 1.6 | 100.8 ± 2.2 | 91.4 ± 0.5 | 86.6 ± 2.7 | 113.3 ± 1.4 | 107.9 ± 3.1 | 0.9992 |
| N-Acetyl-Serotonin | 86.1 ± 2.1 | 106.4 ± 2.2 | 97.9 ± 0.5 | 97.4 ± 3.0 | 113.0 ± 1.6 | 104.3 ± 2.7 | 0.9995 |
| N-Acyl-Homoserine-lactone | 109.2 ± 2.5 | 106.1 ± 3.0 | 93.4 ± 2.8 | 114.6 ± 1.4 | 106.3 ± 3.1 | 89.3 ± 2.4 | 0.9994 |
| NAD | 109.2 ± 2.0 | 88.7 ± 2.1 | 96.4 ± 0.9 | 111.9 ± 2.2 | 89.8 ± 0.3 | 109.7 ± 1.9 | 0.9992 |
| NADH | 111.9 ± 2.1 | 113.5 ± 1.7 | 91.9 ± 2.0 | 101.5 ± 2.9 | 112.7 ± 2.1 | 88.3 ± 2.3 | 0.9998 |
| NADP | 92.1 ± 0.4 | 114.8 ± 2.5 | 100.3 ± 2.1 | 103.3 ± 2.2 | 111.1 ± 0.7 | 114.6 ± 0.4 | 0.9994 |
| NADPH | 112.1 ± 0.6 | 91.2 ± 0.6 | 109.2 ± 2.7 | 85.9 ± 0.8 | 97.3 ± 1.6 | 110.6 ± 1.8 | 0.9992 |
| N-alpha-Acetyl-lysine | 103.7 ± 0.5 | 110.0 ± 2.3 | 85.1 ± 3.0 | 105.7 ± 1.6 | 109.0 ± 2.7 | 114.7 ± 2.0 | 0.9995 |
| N-alpha-Acetyl-Ornithine | 92.1 ± 1.6 | 88.0 ± 2.3 | 95.7 ± 2.3 | 114.2 ± 3.0 | 95.1 ± 0.9 | 108.4 ± 0.6 | 0.9997 |
| N-Carbamoyl-Proline | 112.2 ± 0.5 | 91.0 ± 1.4 | 100.2 ± 1.3 | 100.2 ± 0.6 | 110.2 ± 1.5 | 105.6 ± 2.2 | 0.9998 |
| N-Carbamoyl-Sarcosine | 89.8 ± 2.5 | 100.3 ± 2.9 | 101.0 ± 2.2 | 90.1 ± 0.9 | 107.0 ± 1.6 | 86.0 ± 2.2 | 0.9995 |
| N-Epsilon-Acetyl-lysine | 114.4 ± 2.2 | 90.2 ± 2.9 | 88.1 ± 1.5 | 105.4 ± 2.1 | 96.6 ± 3.0 | 96.6 ± 1.4 | 0.9991 |
| Nicotinamide | 106.2 ± 0.7 | 94.6 ± 0.5 | 93.2 ± 2.0 | 85.3 ± 0.6 | 107.7 ± 0.4 | 91.7 ± 1.6 | 0.9996 |
| N-Methyl-Aspartic Acid | 93.6 ± 2.1 | 87.4 ± 3.2 | 108.2 ± 3.0 | 108.3 ± 2.2 | 112.2 ± 2.5 | 105.9 ± 1.5 | 0.9992 |
| O-Phosphoethanolamine | 107.6 ± 0.9 | 86.6 ± 2.8 | 114.0 ± 0.8 | 98.0 ± 2.0 | 97.2 ± 0.9 | 107.9 ± 2.5 | 0.9992 |
| Ophthalmic acid | 107.5 ± 2.0 | 92.2 ± 2.1 | 104.1 ± 2.3 | 98.7 ± 3.0 | 111.4 ± 0.9 | 105.2 ± 1.7 | 0.9991 |
| Ornithine | 95.0 ± 2.5 | 94.0 ± 1.3 | 90.1 ± 1.7 | 98.7 ± 1.0 | 112.4 ± 2.5 | 94.0 ± 1.4 | 0.9997 |
| Orotidine 5'-monophosphate | 99.3 ± 2.9 | 111.0 ± 0.7 | 114.5 ± 2.4 | 85.8 ± 0.7 | 101.1 ± 0.7 | 109.4 ± 1.0 | 0.9999 |
| Oxaloacetate | 110.1 ± 1.2 | 98.0 ± 2.3 | 97.7 ± 2.8 | 98.9 ± 2.0 | 103.6 ± 1.6 | 94.6 ± 1.3 | 0.9995 |
| Pantothenic acid | 107.9 ± 1.2 | 111.9 ± 0.6 | 106.1 ± 1.6 | 103.4 ± 3.1 | 95.6 ± 2.1 | 96.0 ± 0.8 | 0.9996 |
| Phenylalanine | 89.7 ± 2.4 | 109.2 ± 1.1 | 107.6 ± 3.0 | 95.5 ± 0.8 | 110.9 ± 0.6 | 100.3 ± 1.6 | 0.9999 |
| Phenylpyruvate | 113.0 ± 0.9 | 101.0 ± 1.3 | 101.0 ± 1.6 | 100.4 ± 2.9 | 108.3 ± 1.0 | 85.9 ± 0.5 | 0.9994 |
| Phosphodimethylethanolamine | 102.6 ± 2.8 | 104.4 ± 1.0 | 109.4 ± 2.4 | 98.1 ± 1.6 | 98.3 ± 1.5 | 90.9 ± 2.4 | 0.9994 |
| Phosphoenolpyruvate | 98.7 ± 2.5 | 114.4 ± 2.8 | 112.5 ± 0.3 | 93.9 ± 1.5 | 89.8 ± 0.8 | 103.7 ± 1.8 | 0.9996 |
| Phosphorylcholine | 95.1 ± 2.4 | 94.6 ± 1.3 | 86.0 ± 1.4 | 112.7 ± 0.6 | 96.9 ± 1.1 | 97.1 ± 1.3 | 0.9993 |
| Pipecolic acid | 108.8 ± 3.0 | 98.6 ± 2.8 | 105.0 ± 2.6 | 106.9 ± 2.9 | 104.4 ± 2.7 | 97.6 ± 1.4 | 0.9996 |
| Proline | 89.1 ± 3.2 | 102.0 ± 1.8 | 97.0 ± 2.1 | 86.2 ± 1.9 | 101.2 ± 2.6 | 95.5 ± 1.3 | 0.9998 |
| Putrescine | 87.9 ± 3.0 | 95.1 ± 0.5 | 110.9 ± 2.5 | 94.9 ± 1.9 | 101.6 ± 2.4 | 89.9 ± 3.2 | 0.9996 |
| Pyridoxal | 97.8 ± 2.8 | 88.2 ± 3.1 | 85.6 ± 2.7 | 94.8 ± 2.6 | 90.5 ± 0.6 | 106.4 ± 2.7 | 0.9996 |
| Pyridoxamine | 97.9 ± 2.0 | 87.5 ± 1.8 | 97.2 ± 2.7 | 89.0 ± 2.3 | 109.3 ± 0.5 | 96.6 ± 0.5 | 0.9997 |
| Pyridoxine | 86.2 ± 1.8 | 108.6 ± 0.9 | 102.0 ± 2.6 | 114.7 ± 2.7 | 107.5 ± 0.5 | 99.6 ± 2.6 | 0.9994 |
| Pyroglutamic acid | 103.9 ± 1.8 | 86.6 ± 3.1 | 87.3 ± 2.2 | 114.5 ± 2.3 | 93.3 ± 2.4 | 93.0 ± 3.0 | 0.9992 |
| Pyrrolysine | 88.0 ± 2.3 | 93.5 ± 0.6 | 90.8 ± 0.8 | 107.2 ± 2.4 | 88.9 ± 2.7 | 104.8 ± 2.2 | 0.9996 |
| Pyruvic Acid | 86.6 ± 2.1 | 107.3 ± 0.7 | 114.8 ± 2.1 | 88.1 ± 2.9 | 97.1 ± 3.1 | 85.6 ± 1.3 | 0.9991 |
| Riboflavin | 108.9 ± 1.5 | 86.2 ± 3.0 | 93.0 ± 3.0 | 92.3 ± 0.9 | 91.7 ± 1.9 | 99.4 ± 1.9 | 0.9999 |
| Ribose-5-Phosphate | 103.7 ± 1.6 | 108.0 ± 1.2 | 106.5 ± 0.6 | 103.4 ± 0.3 | 112.9 ± 0.4 | 103.2 ± 2.3 | 0.9996 |
| Ribulose-1-Phosphate | 100.2 ± 0.3 | 102.6 ± 1.1 | 94.9 ± 1.4 | 109.6 ± 2.4 | 94.8 ± 1.7 | 100.2 ± 1.1 | 0.9997 |
| Ribulose-5-Phosphate | 94.2 ± 3.0 | 107.9 ± 0.7 | 113.9 ± 1.4 | 86.2 ± 0.3 | 107.5 ± 0.5 | 112.0 ± 0.7 | 0.9995 |
| Saccharopine | 109.0 ± 2.6 | 90.5 ± 0.5 | 106.2 ± 1.7 | 96.0 ± 1.5 | 111.0 ± 1.5 | 87.4 ± 2.8 | 0.9996 |
| S-Adenosylhomocysteine | 96.7 ± 2.2 | 107.4 ± 2.2 | 95.4 ± 1.9 | 104.1 ± 3.2 | 93.0 ± 1.7 | 92.1 ± 1.5 | 0.9997 |
| S-Adenosylmethioninamine | 106.8 ± 1.9 | 95.5 ± 2.0 | 102.3 ± 1.2 | 113.2 ± 0.5 | 112.4 ± 1.0 | 113.9 ± 2.0 | 0.9995 |
| S-Adenosylmethionine | 101.4 ± 2.5 | 103.9 ± 1.0 | 109.5 ± 0.6 | 98.6 ± 1.3 | 103.8 ± 2.1 | 90.6 ± 1.3 | 0.9998 |
| Sarcosine | 113.0 ± 0.9 | 91.4 ± 3.0 | 99.7 ± 1.1 | 112.1 ± 0.8 | 90.8 ± 0.4 | 102.1 ± 2.0 | 0.9992 |
| Sedoheptulose-7-phosphate | 96.8 ± 3.2 | 88.2 ± 1.0 | 109.7 ± 2.2 | 106.0 ± 3.1 | 114.7 ± 1.2 | 103.5 ± 1.1 | 0.9998 |
| Selenohomocysteine | 99.3 ± 1.8 | 100.6 ± 1.0 | 92.6 ± 3.1 | 111.1 ± 3.1 | 108.1 ± 2.5 | 85.9 ± 2.5 | 0.9998 |
| Selenomethionine | 109.4 ± 2.1 | 102.1 ± 0.4 | 110.4 ± 2.6 | 87.3 ± 0.4 | 88.4 ± 2.6 | 89.9 ± 2.7 | 0.9995 |
| Serine | 95.0 ± 0.4 | 98.8 ± 1.7 | 86.2 ± 3.0 | 92.8 ± 1.4 | 90.2 ± 2.3 | 89.6 ± 2.6 | 0.9998 |
| S-Methyl-Thioinosine | 86.9 ± 1.8 | 111.7 ± 1.4 | 104.5 ± 1.4 | 89.7 ± 0.9 | 86.7 ± 3.0 | 98.8 ± 0.5 | 0.9994 |
| Spermidine | 97.8 ± 1.6 | 103.3 ± 3.1 | 94.1 ± 0.6 | 106.8 ± 2.2 | 95.6 ± 2.5 | 99.5 ± 1.9 | 0.9998 |
| Spermine | 111.7 ± 0.4 | 97.4 ± 1.3 | 97.0 ± 2.3 | 104.0 ± 2.2 | 96.2 ± 0.5 | 93.1 ± 2.5 | 0.9997 |
| Succinate | 89.6 ± 2.8 | 101.7 ± 2.3 | 96.1 ± 1.4 | 113.3 ± 1.7 | 98.9 ± 3.1 | 95.0 ± 2.0 | 0.9995 |
| Sucrose | 104.4 ± 1.8 | 87.2 ± 1.9 | 108.1 ± 1.7 | 90.7 ± 2.4 | 108.2 ± 2.9 | 94.4 ± 2.2 | 0.9992 |
| Symmetric dimethylarginine | 111.4 ± 2.6 | 85.6 ± 3.1 | 98.9 ± 1.3 | 114.2 ± 0.8 | 86.7 ± 1.4 | 101.4 ± 2.2 | 0.9994 |
| Taurine | 90.4 ± 3.1 | 91.0 ± 2.6 | 107.7 ± 1.8 | 114.8 ± 2.8 | 88.8 ± 2.7 | 99.7 ± 3.2 | 0.9996 |
| Tetrahydroneopterin | 86.3 ± 2.7 | 104.3 ± 2.0 | 103.1 ± 1.9 | 102.1 ± 1.6 | 97.5 ± 0.7 | 105.5 ± 1.8 | 0.9994 |
| Thiamine | 85.6 ± 0.9 | 96.3 ± 2.2 | 95.3 ± 2.5 | 102.8 ± 0.3 | 97.1 ± 3.2 | 93.0 ± 1.6 | 0.9993 |
| Thiamine monophosphate | 110.0 ± 2.7 | 89.2 ± 2.5 | 97.8 ± 2.5 | 100.3 ± 0.4 | 108.4 ± 1.7 | 108.4 ± 1.9 | 0.9996 |
| Threonic acid | 106.4 ± 0.6 | 108.7 ± 1.2 | 87.9 ± 1.5 | 107.6 ± 1.1 | 98.5 ± 0.5 | 106.4 ± 0.4 | 0.9992 |
| Threonine | 109.4 ± 0.5 | 107.3 ± 1.0 | 105.5 ± 2.8 | 105.1 ± 3.1 | 111.3 ± 2.5 | 105.4 ± 0.4 | 0.9997 |
| Thymidine | 86.8 ± 2.1 | 94.1 ± 2.9 | 88.9 ± 2.8 | 85.1 ± 2.5 | 109.0 ± 2.4 | 107.0 ± 1.9 | 0.9997 |
| Thymidine Monophosphate | 85.2 ± 2.4 | 95.4 ± 1.8 | 98.8 ± 1.0 | 86.1 ± 0.9 | 93.3 ± 2.3 | 110.0 ± 1.3 | 0.9993 |
| Thymine | 114.4 ± 2.8 | 85.8 ± 1.4 | 100.7 ± 1.1 | 103.5 ± 0.7 | 107.4 ± 0.6 | 112.1 ± 2.3 | 0.9996 |
| Trigonelleine | 93.0 ± 2.4 | 87.1 ± 1.3 | 98.3 ± 1.3 | 98.1 ± 2.8 | 87.8 ± 1.8 | 85.1 ± 3.1 | 0.9993 |
| Trimethylamine N-oxide | 85.6 ± 1.8 | 98.3 ± 0.6 | 92.5 ± 2.0 | 109.8 ± 3.2 | 98.7 ± 2.8 | 106.5 ± 0.8 | 0.9994 |
| Trimethyllysine | 93.5 ± 2.3 | 102.8 ± 2.8 | 86.4 ± 3.1 | 109.5 ± 2.7 | 90.1 ± 2.2 | 101.7 ± 0.8 | 0.9992 |
| Tryptophan | 86.9 ± 2.6 | 98.1 ± 1.4 | 98.9 ± 2.6 | 94.7 ± 0.8 | 113.3 ± 2.3 | 103.5 ± 2.5 | 0.9993 |
| Tyramine | 95.0 ± 0.9 | 87.4 ± 2.4 | 114.1 ± 1.6 | 103.8 ± 2.9 | 103.6 ± 0.9 | 105.8 ± 3.1 | 0.9992 |
| Tyrosine | 86.0 ± 1.6 | 110.3 ± 1.1 | 106.8 ± 0.6 | 112.0 ± 3.1 | 93.3 ± 3.1 | 86.7 ± 0.3 | 0.9997 |
| Uracil | 91.2 ± 0.5 | 101.8 ± 1.3 | 103.9 ± 1.7 | 99.4 ± 2.5 | 107.8 ± 0.9 | 111.9 ± 1.4 | 0.9994 |
| Uridine 5'-diphosphate | 94.0 ± 2.8 | 97.8 ± 2.3 | 89.0 ± 2.1 | 88.5 ± 0.7 | 99.3 ± 1.4 | 112.8 ± 2.8 | 0.9994 |
| Uridine Monophosphate | 93.6 ± 1.8 | 105.1 ± 0.3 | 114.5 ± 1.3 | 89.2 ± 1.5 | 114.5 ± 2.7 | 113.6 ± 0.5 | 0.9999 |
| Uridine-5-Diphosphogalactose | 93.1 ± 2.3 | 107.2 ± 3.2 | 89.6 ± 1.9 | 102.2 ± 1.7 | 85.8 ± 3.1 | 86.8 ± 2.8 | 0.9993 |
| Urocanic acid | 111.2 ± 1.4 | 101.7 ± 1.8 | 105.1 ± 0.8 | 110.4 ± 3.1 | 98.5 ± 1.0 | 96.6 ± 1.3 | 0.9991 |
| UTP | 90.0 ± 1.5 | 86.7 ± 2.9 | 88.9 ± 1.3 | 97.1 ± 1.2 | 104.7 ± 0.6 | 88.3 ± 0.6 | 0.9996 |
| Valine | 96.9 ± 3.0 | 107.3 ± 3.1 | 99.1 ± 1.2 | 111.3 ± 1.5 | 86.4 ± 3.0 | 105.5 ± 2.8 | 0.9996 |
| XMP | 104.5 ± 1.2 | 97.3 ± 0.4 | 101.3 ± 2.3 | 90.5 ± 2.2 | 85.9 ± 2.0 | 112.4 ± 0.5 | 0.9998 |
| Xylulose-5-Phosphate | 107.6 ± 2.5 | 88.1 ± 1.9 | 109.7 ± 2.2 | 112.3 ± 2.0 | 109.9 ± 1.6 | 93.1 ± 2.6 | 0.9998 |

### **Table S8** | Calibration values for final method conditions (r^2^-value) and percent found of nominal calibrator concentration (Tumor tissue extract as matrix).

| Calibrator concentration accuracy  [%] | 10 ng/mL | 25 ng/mL | 50 ng/mL | 100 ng/mL | 500 ng/mL | 1000 ng/mL | r^2^-Value  matrix-matched calibration |
| --- | --- | --- | --- | --- | --- | --- | --- |
| 1-Methylguanine | 93.9 ± 0.6 | 94.7 ± 3.0 | 98.3 ± 0.5 | 89.1 ± 0.8 | 111.9 ± 2.4 | 85.3 ± 1.3 | 0.9997 |
| 1-Methylhistidine | 106.2 ± 2.0 | 98.6 ± 1.5 | 105.6 ± 1.5 | 102.6 ± 2.8 | 104.8 ± 2.2 | 92.8 ± 1.5 | 0.9995 |
| 1-Methylnicotinamide | 105.6 ± 2.9 | 92.2 ± 2.5 | 114.0 ± 2.5 | 91.4 ± 0.8 | 109.7 ± 1.1 | 92.4 ± 1.5 | 0.9997 |
| 2-Aminoadipic Acid | 98.2 ± 2.4 | 106.5 ± 2.8 | 112.4 ± 1.2 | 95.3 ± 1.9 | 114.4 ± 0.7 | 89.2 ± 0.6 | 0.9994 |
| 2-Deoxyadenosine | 103.5 ± 2.6 | 107.7 ± 2.7 | 114.1 ± 1.0 | 87.3 ± 1.9 | 92.6 ± 3.1 | 102.9 ± 1.3 | 0.9992 |
| 2'-Deoxyadenosine 5'-monophosphate | 110.0 ± 0.6 | 96.0 ± 1.8 | 93.9 ± 1.7 | 105.2 ± 1.3 | 98.0 ± 2.4 | 102.7 ± 1.2 | 0.9997 |
| 2'-deoxyadenosine 5'-triphosphate | 96.5 ± 2.8 | 97.1 ± 0.4 | 111.6 ± 2.3 | 87.7 ± 2.8 | 102.6 ± 2.1 | 96.8 ± 0.6 | 0.9994 |
| 2'-Deoxycytidine-5'-monophosphate | 114.6 ± 0.4 | 112.9 ± 1.8 | 106.4 ± 0.5 | 99.6 ± 0.5 | 112.0 ± 2.6 | 86.5 ± 2.8 | 0.9994 |
| 2-Deoxyguanosine | 99.6 ± 2.8 | 111.0 ± 2.7 | 86.3 ± 0.9 | 110.2 ± 2.7 | 108.4 ± 2.5 | 113.4 ± 0.9 | 0.9995 |
| 2-hydroxybutyric acid | 89.9 ± 2.1 | 103.6 ± 2.7 | 88.4 ± 1.5 | 96.9 ± 3.1 | 111.5 ± 3.1 | 100.2 ± 0.7 | 0.9996 |
| 2-hydroxyglutaric acid | 104.2 ± 2.4 | 92.8 ± 1.6 | 109.8 ± 2.6 | 90.7 ± 1.8 | 110.1 ± 2.3 | 97.3 ± 0.6 | 0.9999 |
| 2-Hydroxyisocaproic acid | 86.9 ± 1.2 | 94.2 ± 2.8 | 93.9 ± 1.0 | 97.4 ± 0.8 | 90.7 ± 0.8 | 95.3 ± 0.4 | 0.9993 |
| 2-ketobutyric acid | 86.4 ± 1.2 | 109.1 ± 0.9 | 114.6 ± 2.3 | 89.6 ± 2.5 | 99.7 ± 2.9 | 105.0 ± 2.4 | 0.9997 |
| 2-oxoadipic acid | 110.2 ± 1.7 | 87.4 ± 1.6 | 106.9 ± 3.2 | 93.3 ± 0.5 | 107.5 ± 2.4 | 93.7 ± 3.1 | 0.9994 |
| 2-Phenylglycine | 90.2 ± 3.1 | 95.3 ± 0.8 | 90.7 ± 1.9 | 99.8 ± 2.6 | 113.0 ± 3.1 | 104.5 ± 3.0 | 0.9999 |
| 2-phosphoglycerate | 89.9 ± 1.2 | 86.5 ± 1.5 | 106.3 ± 1.5 | 94.3 ± 2.9 | 100.1 ± 3.1 | 87.5 ± 1.4 | 0.9997 |
| 3-hydroxy-3-methyl-2-oxobutanoate | 97.4 ± 3.0 | 89.9 ± 0.7 | 113.7 ± 3.0 | 86.5 ± 0.6 | 99.6 ± 1.3 | 91.9 ± 1.9 | 0.9992 |
| 3-hydroxybutyric acid | 97.1 ± 1.2 | 98.8 ± 1.6 | 102.0 ± 1.5 | 103.8 ± 2.6 | 89.4 ± 2.9 | 114.7 ± 0.8 | 0.9998 |
| 3-Methylhistidine | 85.3 ± 2.5 | 85.3 ± 0.4 | 102.3 ± 2.0 | 96.7 ± 1.7 | 94.6 ± 1.5 | 100.6 ± 0.9 | 0.9993 |
| 3-Phosphoglyceraldehyde | 87.5 ± 2.5 | 101.8 ± 2.3 | 113.1 ± 1.0 | 97.5 ± 2.2 | 110.8 ± 0.6 | 109.9 ± 0.8 | 0.9999 |
| 3-phosphoglycerate | 88.1 ± 1.7 | 105.7 ± 1.0 | 91.8 ± 0.7 | 98.5 ± 1.5 | 92.7 ± 1.9 | 95.3 ± 3.0 | 0.9998 |
| 4-amino-butyric acid | 92.9 ± 1.5 | 91.1 ± 2.0 | 98.6 ± 1.8 | 101.9 ± 1.2 | 113.9 ± 2.1 | 109.6 ± 1.0 | 0.9993 |
| 4-aspartyl-phosphate | 90.9 ± 0.4 | 113.6 ± 2.2 | 106.8 ± 1.1 | 86.2 ± 1.2 | 108.9 ± 1.7 | 99.0 ± 0.3 | 0.9993 |
| 4-Hydroxybenzoic acid | 112.9 ± 0.7 | 103.0 ± 2.0 | 112.0 ± 0.5 | 88.7 ± 2.5 | 106.3 ± 2.0 | 106.8 ± 2.9 | 0.9995 |
| 4-Hydroxyphenylacetylglycine | 108.9 ± 2.2 | 111.8 ± 0.5 | 109.2 ± 2.0 | 95.8 ± 0.5 | 91.8 ± 1.8 | 97.8 ± 0.8 | 0.9995 |
| 4-methyl-2-oxopentanoate | 107.9 ± 0.7 | 91.6 ± 2.3 | 111.3 ± 2.8 | 85.8 ± 2.9 | 100.5 ± 0.3 | 86.8 ± 3.2 | 0.9992 |
| 4-Oxoglutaramate | 112.9 ± 1.2 | 96.5 ± 1.2 | 107.1 ± 2.3 | 102.2 ± 2.5 | 96.8 ± 2.3 | 106.8 ± 3.1 | 0.9992 |
| 5-Hydroxyindolepyruvate | 89.8 ± 2.7 | 87.4 ± 2.1 | 95.7 ± 2.8 | 87.9 ± 3.1 | 105.8 ± 0.7 | 88.4 ± 2.1 | 0.9995 |
| 5-Hydroxylysine | 85.9 ± 2.9 | 93.0 ± 0.9 | 112.9 ± 0.6 | 89.9 ± 1.3 | 96.4 ± 2.6 | 113.8 ± 3.1 | 0.9997 |
| 5-Hydroxy-Tryptophan | 103.3 ± 2.9 | 98.5 ± 0.6 | 85.7 ± 1.7 | 94.9 ± 1.7 | 109.7 ± 1.1 | 101.6 ± 1.4 | 0.9996 |
| 5-Phosphoribosylamine | 90.5 ± 1.8 | 91.7 ± 0.6 | 92.7 ± 0.5 | 99.7 ± 1.7 | 103.2 ± 0.8 | 105.2 ± 1.6 | 0.9992 |
| 6-phosphogluconate | 109.9 ± 2.1 | 87.2 ± 2.9 | 108.7 ± 0.5 | 86.9 ± 2.5 | 101.2 ± 2.9 | 97.9 ± 0.9 | 0.9992 |
| Acetylcarnitine | 90.7 ± 1.4 | 96.0 ± 0.8 | 112.7 ± 1.9 | 95.1 ± 0.7 | 94.6 ± 0.5 | 85.5 ± 1.6 | 0.9994 |
| Aconitic Acid | 101.4 ± 1.3 | 92.9 ± 2.3 | 92.9 ± 2.0 | 102.9 ± 3.0 | 88.7 ± 1.1 | 87.1 ± 1.9 | 0.9998 |
| Adenine | 111.2 ± 0.4 | 90.7 ± 1.2 | 103.8 ± 0.8 | 105.5 ± 0.5 | 95.1 ± 1.0 | 90.9 ± 2.2 | 0.9992 |
| Adenosine | 101.6 ± 2.8 | 104.3 ± 1.5 | 111.7 ± 2.5 | 104.8 ± 1.2 | 110.1 ± 2.4 | 87.2 ± 1.5 | 0.9994 |
| Adenosine 3',5'-cyclic monophosphate | 86.3 ± 1.8 | 108.9 ± 1.1 | 111.5 ± 2.1 | 100.4 ± 0.4 | 110.8 ± 0.5 | 96.2 ± 2.1 | 0.9993 |
| Adenosine Monophosphate | 111.0 ± 2.5 | 114.5 ± 2.9 | 94.3 ± 1.3 | 86.3 ± 1.2 | 111.4 ± 3.1 | 105.8 ± 1.7 | 0.9995 |
| Adenosine-5'-diphosphate | 98.7 ± 2.6 | 94.6 ± 0.8 | 109.5 ± 0.7 | 113.4 ± 1.4 | 114.9 ± 2.2 | 102.3 ± 2.5 | 0.9992 |
| Adenosine-5'-triphosphate | 90.7 ± 0.4 | 92.7 ± 0.8 | 93.2 ± 0.5 | 105.2 ± 2.8 | 88.8 ± 1.8 | 96.0 ± 2.6 | 0.9996 |
| Adenylosuccinate | 112.7 ± 2.1 | 107.0 ± 3.0 | 113.8 ± 1.5 | 110.1 ± 1.0 | 97.8 ± 1.4 | 93.2 ± 1.4 | 0.9993 |
| Alanine | 114.4 ± 3.0 | 103.2 ± 0.6 | 97.4 ± 2.5 | 106.1 ± 1.6 | 102.3 ± 2.5 | 85.4 ± 2.9 | 0.9998 |
| alpha-ketoglutaric acid | 90.8 ± 3.2 | 108.8 ± 1.8 | 96.0 ± 2.7 | 111.0 ± 1.0 | 98.1 ± 0.8 | 100.0 ± 0.7 | 0.9999 |
| Anserine | 112.9 ± 1.2 | 108.6 ± 1.8 | 107.7 ± 3.1 | 89.4 ± 0.6 | 98.2 ± 1.9 | 102.7 ± 1.3 | 0.9995 |
| Arginine | 104.5 ± 2.8 | 92.9 ± 2.9 | 105.3 ± 0.7 | 96.7 ± 0.6 | 113.5 ± 1.2 | 95.1 ± 0.7 | 0.9998 |
| Argininosuccinic Acid | 104.3 ± 1.0 | 93.2 ± 0.7 | 99.7 ± 1.7 | 102.2 ± 2.4 | 102.2 ± 2.7 | 95.0 ± 3.0 | 0.9994 |
| Asparagine | 105.1 ± 0.8 | 88.2 ± 1.3 | 97.9 ± 2.5 | 108.1 ± 2.8 | 106.3 ± 0.5 | 100.2 ± 2.7 | 0.9995 |
| Aspartic acid | 85.6 ± 1.4 | 102.7 ± 1.4 | 86.2 ± 2.1 | 113.4 ± 0.8 | 106.7 ± 0.4 | 98.8 ± 3.1 | 0.9998 |
| Asymmetric dimethylarginine | 101.1 ± 1.3 | 97.4 ± 1.9 | 86.8 ± 1.2 | 87.3 ± 0.9 | 104.1 ± 2.9 | 114.7 ± 3.1 | 0.9999 |
| Azelaic acid | 104.8 ± 0.4 | 114.0 ± 1.0 | 102.8 ± 1.1 | 104.3 ± 0.8 | 97.8 ± 1.1 | 97.6 ± 1.8 | 0.9997 |
| beta-Alanine | 92.6 ± 1.4 | 94.3 ± 1.6 | 106.7 ± 2.6 | 102.9 ± 1.4 | 95.3 ± 1.9 | 109.9 ± 0.8 | 0.9996 |
| Betaine | 112.0 ± 0.8 | 104.7 ± 2.6 | 97.6 ± 0.3 | 107.9 ± 0.5 | 89.5 ± 2.3 | 106.8 ± 2.8 | 0.9994 |
| Biopterin | 98.4 ± 2.7 | 101.4 ± 2.9 | 102.5 ± 1.0 | 112.0 ± 1.7 | 104.4 ± 1.2 | 109.3 ± 2.0 | 0.9991 |
| Biotin | 88.9 ± 1.1 | 108.8 ± 3.1 | 94.0 ± 0.4 | 88.1 ± 3.2 | 99.5 ± 1.3 | 105.0 ± 1.8 | 0.9993 |
| Carnitine | 91.7 ± 1.9 | 88.4 ± 3.2 | 112.6 ± 0.7 | 112.9 ± 1.3 | 91.3 ± 1.3 | 89.2 ± 0.8 | 0.9999 |
| Carnosine | 98.2 ± 2.7 | 88.8 ± 2.3 | 107.3 ± 2.7 | 86.4 ± 2.6 | 90.8 ± 2.4 | 109.0 ± 2.0 | 0.9992 |
| CDP-choline | 104.8 ± 2.1 | 85.5 ± 2.4 | 89.8 ± 1.6 | 87.3 ± 2.5 | 107.5 ± 1.1 | 87.9 ± 2.5 | 0.9998 |
| cGMP | 89.7 ± 1.8 | 106.0 ± 2.7 | 105.4 ± 3.0 | 93.0 ± 1.1 | 95.2 ± 0.8 | 104.9 ± 1.6 | 0.9996 |
| Choline | 93.8 ± 1.8 | 113.1 ± 0.6 | 100.0 ± 0.9 | 109.7 ± 2.0 | 111.3 ± 1.9 | 99.4 ± 2.7 | 0.9997 |
| Cinnamic acid | 85.3 ± 2.9 | 106.9 ± 1.9 | 97.7 ± 2.1 | 112.2 ± 1.2 | 90.9 ± 0.7 | 109.2 ± 0.9 | 0.9992 |
| Citraconic acid | 98.8 ± 0.6 | 87.0 ± 2.8 | 85.9 ± 3.0 | 86.2 ± 2.6 | 87.7 ± 1.6 | 95.4 ± 0.6 | 0.9993 |
| Citramalic acid | 86.0 ± 0.3 | 86.2 ± 2.0 | 85.6 ± 1.7 | 95.7 ± 0.6 | 113.4 ± 1.6 | 111.3 ± 1.9 | 0.9996 |
| Citric Acid | 110.3 ± 1.4 | 97.6 ± 1.9 | 101.0 ± 0.7 | 86.5 ± 1.5 | 97.2 ± 0.6 | 95.6 ± 1.1 | 0.9993 |
| Citrulline | 97.0 ± 2.8 | 110.8 ± 2.8 | 108.7 ± 2.8 | 85.0 ± 1.9 | 114.2 ± 0.8 | 110.6 ± 2.4 | 0.9991 |
| Creatine | 100.7 ± 2.6 | 88.5 ± 3.0 | 105.1 ± 1.1 | 94.7 ± 2.2 | 113.1 ± 2.9 | 89.6 ± 0.9 | 0.9995 |
| Creatinine | 100.9 ± 1.9 | 91.2 ± 1.5 | 113.5 ± 1.6 | 106.9 ± 2.6 | 109.4 ± 2.8 | 90.6 ± 2.3 | 0.9997 |
| Cystathionine | 102.5 ± 2.2 | 108.6 ± 1.4 | 85.4 ± 1.6 | 104.5 ± 2.1 | 114.4 ± 0.7 | 95.2 ± 0.7 | 0.9994 |
| Cysteamine | 101.8 ± 1.3 | 90.5 ± 2.7 | 90.0 ± 1.9 | 98.3 ± 2.3 | 97.2 ± 0.9 | 109.1 ± 2.5 | 0.9998 |
| Cysteine | 89.2 ± 0.5 | 113.0 ± 0.4 | 88.6 ± 2.4 | 113.9 ± 3.0 | 106.0 ± 0.9 | 113.5 ± 1.4 | 0.9991 |
| Cysteinylglycine | 94.4 ± 1.2 | 109.7 ± 2.6 | 107.8 ± 2.1 | 110.0 ± 1.9 | 113.1 ± 1.5 | 104.4 ± 2.4 | 0.9994 |
| Cystine | 99.8 ± 1.8 | 110.9 ± 0.4 | 97.9 ± 2.1 | 90.7 ± 3.0 | 93.3 ± 0.4 | 110.9 ± 1.0 | 0.9993 |
| Cytidine | 90.4 ± 0.7 | 104.9 ± 1.4 | 111.3 ± 0.6 | 97.3 ± 2.2 | 95.8 ± 3.0 | 99.0 ± 0.5 | 0.9993 |
| Cytidine 3',5'-cyclic monophosphate | 104.9 ± 1.0 | 113.1 ± 2.2 | 112.8 ± 0.8 | 92.3 ± 0.7 | 107.1 ± 3.1 | 114.2 ± 1.8 | 0.9991 |
| Cytidine 5'-diphosphate | 111.0 ± 2.2 | 100.7 ± 2.3 | 94.8 ± 1.7 | 99.7 ± 1.4 | 103.5 ± 0.3 | 86.7 ± 0.7 | 0.9996 |
| Cytidine 5'-triphosphate | 88.7 ± 3.0 | 97.3 ± 2.7 | 94.0 ± 2.2 | 85.9 ± 3.1 | 103.4 ± 0.7 | 110.8 ± 1.8 | 0.9996 |
| Cytidine monophosphate | 91.7 ± 1.7 | 91.6 ± 1.9 | 106.2 ± 3.1 | 96.9 ± 2.4 | 111.3 ± 2.3 | 88.1 ± 0.9 | 0.9995 |
| Cytosine | 111.2 ± 2.9 | 114.8 ± 3.2 | 103.1 ± 1.5 | 91.3 ± 3.2 | 111.5 ± 1.4 | 102.7 ± 0.5 | 0.9993 |
| dADP | 90.7 ± 2.8 | 108.1 ± 0.6 | 86.1 ± 1.6 | 98.6 ± 2.2 | 86.1 ± 3.1 | 99.8 ± 1.5 | 0.9996 |
| Diethanolamine | 99.8 ± 1.9 | 100.3 ± 0.8 | 102.2 ± 1.4 | 105.4 ± 1.7 | 88.1 ± 2.2 | 102.3 ± 1.2 | 0.9994 |
| Dihydrobiopterin | 97.9 ± 3.1 | 91.5 ± 0.5 | 109.7 ± 0.3 | 113.8 ± 0.5 | 97.6 ± 2.5 | 98.2 ± 2.6 | 0.9993 |
| Dihydroneopterin triphosphate | 103.2 ± 1.4 | 103.8 ± 2.5 | 100.0 ± 1.9 | 86.6 ± 2.4 | 101.4 ± 1.9 | 102.0 ± 1.2 | 0.9992 |
| Dihydroxyacetonephosphate | 89.6 ± 3.0 | 96.8 ± 1.7 | 87.6 ± 2.9 | 106.8 ± 1.4 | 97.5 ± 2.9 | 86.4 ± 0.7 | 0.9992 |
| Dimethylglycine | 86.9 ± 1.1 | 87.3 ± 2.6 | 102.5 ± 2.7 | 93.8 ± 1.6 | 99.4 ± 1.1 | 92.0 ± 1.9 | 0.9992 |
| Ectoine | 105.2 ± 1.3 | 113.1 ± 1.0 | 110.0 ± 3.0 | 106.0 ± 2.3 | 97.5 ± 2.2 | 89.7 ± 2.4 | 0.9992 |
| Erythrose-4-phosphate | 113.3 ± 2.3 | 113.5 ± 3.1 | 104.7 ± 1.9 | 88.6 ± 0.8 | 94.7 ± 1.6 | 113.2 ± 1.0 | 0.9994 |
| FAD | 85.3 ± 0.5 | 110.3 ± 2.0 | 109.6 ± 1.3 | 95.3 ± 1.6 | 86.2 ± 1.8 | 101.7 ± 2.7 | 0.9993 |
| Folic acid | 92.8 ± 1.7 | 105.1 ± 0.9 | 92.3 ± 2.0 | 109.0 ± 1.7 | 94.8 ± 3.0 | 93.9 ± 1.2 | 0.9998 |
| Fructose-6-phosphate | 90.7 ± 2.4 | 85.3 ± 2.3 | 100.6 ± 0.8 | 112.9 ± 1.0 | 97.9 ± 2.7 | 96.8 ± 1.4 | 0.9992 |
| Fumaric Acid | 90.4 ± 2.2 | 112.6 ± 0.8 | 104.5 ± 0.5 | 112.0 ± 1.6 | 108.1 ± 1.5 | 94.9 ± 0.7 | 0.9997 |
| Galactosamine | 103.6 ± 1.8 | 93.2 ± 0.5 | 113.0 ± 3.1 | 108.1 ± 2.9 | 102.4 ± 0.8 | 112.0 ± 1.6 | 0.9992 |
| Galactose-1-phosphate | 111.7 ± 2.8 | 113.8 ± 3.0 | 108.6 ± 2.1 | 102.3 ± 2.2 | 110.0 ± 2.0 | 98.0 ± 1.8 | 0.9994 |
| Galactose-6-Phosphate | 93.7 ± 1.1 | 96.3 ± 3.2 | 104.9 ± 1.2 | 97.3 ± 2.8 | 113.7 ± 0.4 | 111.9 ± 2.0 | 0.9996 |
| gamma-Glutamylcysteine | 104.5 ± 2.8 | 97.3 ± 0.9 | 92.1 ± 2.0 | 112.8 ± 2.7 | 86.0 ± 2.0 | 100.2 ± 2.9 | 0.9993 |
| Glucosamine | 92.7 ± 0.7 | 108.6 ± 0.4 | 98.6 ± 1.5 | 113.6 ± 1.9 | 108.5 ± 1.7 | 100.8 ± 0.4 | 0.9995 |
| Glucosamine-6-Phosphate | 90.6 ± 2.9 | 89.9 ± 0.4 | 114.5 ± 3.0 | 109.1 ± 2.4 | 102.7 ± 0.9 | 102.7 ± 0.7 | 0.9994 |
| Glucose | 110.7 ± 2.0 | 111.0 ± 2.6 | 105.3 ± 2.8 | 99.7 ± 1.6 | 102.1 ± 1.1 | 103.1 ± 0.6 | 0.9997 |
| Glucose-1-phosphate | 99.1 ± 1.3 | 110.3 ± 2.2 | 96.7 ± 0.4 | 86.3 ± 1.9 | 114.0 ± 1.4 | 110.7 ± 1.8 | 0.9997 |
| Glucose-6-Phosphate | 114.8 ± 1.7 | 110.0 ± 2.6 | 100.2 ± 2.9 | 86.8 ± 1.0 | 103.5 ± 0.7 | 112.9 ± 0.4 | 0.9996 |
| Glucuronic acid | 91.1 ± 1.8 | 90.4 ± 2.4 | 107.5 ± 1.5 | 87.2 ± 0.8 | 102.0 ± 2.3 | 98.0 ± 3.1 | 0.9996 |
| Glutamic acid | 104.1 ± 2.4 | 111.8 ± 2.3 | 92.9 ± 1.6 | 102.2 ± 2.1 | 90.5 ± 2.3 | 110.9 ± 2.5 | 0.9995 |
| Glutamine | 92.9 ± 1.7 | 103.4 ± 1.6 | 111.7 ± 3.0 | 90.9 ± 2.0 | 102.9 ± 1.0 | 88.9 ± 1.9 | 0.9997 |
| Glutathione | 99.8 ± 1.3 | 108.8 ± 0.7 | 110.5 ± 0.9 | 96.8 ± 0.4 | 90.8 ± 2.6 | 114.4 ± 1.7 | 0.9996 |
| Glutathionylspermine | 94.4 ± 3.0 | 87.5 ± 3.0 | 106.4 ± 2.4 | 113.8 ± 2.6 | 103.0 ± 0.4 | 112.7 ± 1.7 | 0.9996 |
| Glycerophosphocholine | 104.1 ± 0.9 | 95.0 ± 1.0 | 94.4 ± 2.7 | 99.4 ± 1.2 | 91.2 ± 2.7 | 112.8 ± 2.3 | 0.9991 |
| Glycine | 97.5 ± 0.3 | 112.8 ± 1.6 | 89.5 ± 2.5 | 92.6 ± 0.8 | 97.5 ± 1.4 | 91.8 ± 2.1 | 0.9998 |
| Glycolic acid | 99.8 ± 2.8 | 89.5 ± 0.5 | 104.1 ± 1.7 | 99.4 ± 1.6 | 87.6 ± 3.1 | 99.8 ± 0.8 | 0.9993 |
| Guanidinoacetic acid | 103.6 ± 0.5 | 88.8 ± 0.5 | 102.9 ± 1.0 | 110.7 ± 0.7 | 107.6 ± 2.0 | 105.5 ± 0.5 | 0.9992 |
| Guanidinobutanoic acid | 96.6 ± 1.5 | 100.2 ± 1.5 | 88.0 ± 1.6 | 85.4 ± 0.6 | 112.4 ± 3.2 | 87.4 ± 0.5 | 0.9997 |
| Guanine | 101.7 ± 0.9 | 108.2 ± 2.9 | 93.8 ± 1.8 | 90.4 ± 1.4 | 85.4 ± 1.7 | 101.9 ± 1.2 | 0.9993 |
| Guanosine | 90.0 ± 1.9 | 99.9 ± 1.9 | 110.3 ± 1.5 | 91.6 ± 1.6 | 104.0 ± 1.2 | 112.8 ± 1.1 | 0.9993 |
| Guanosine 5'-diphosphate | 92.5 ± 1.3 | 105.5 ± 0.4 | 107.3 ± 0.8 | 100.4 ± 0.5 | 104.8 ± 0.9 | 88.4 ± 1.0 | 0.9999 |
| Guanosine-5'-monophosphate | 100.8 ± 0.8 | 94.5 ± 1.8 | 100.3 ± 1.6 | 98.7 ± 1.8 | 85.4 ± 3.0 | 99.7 ± 2.8 | 0.9999 |
| Hercynine | 104.7 ± 1.2 | 102.2 ± 2.2 | 100.0 ± 2.4 | 95.9 ± 1.0 | 106.9 ± 1.3 | 111.9 ± 2.3 | 0.9999 |
| Hippuric acid | 85.8 ± 1.1 | 113.6 ± 0.8 | 99.6 ± 0.5 | 114.4 ± 1.1 | 111.7 ± 0.4 | 85.4 ± 2.9 | 0.9996 |
| Histamine | 96.9 ± 1.9 | 115.0 ± 1.4 | 95.5 ± 0.5 | 94.2 ± 3.1 | 85.2 ± 3.1 | 99.9 ± 1.8 | 0.9991 |
| Histidine | 90.4 ± 1.4 | 90.5 ± 1.9 | 90.9 ± 1.7 | 113.1 ± 1.1 | 99.0 ± 0.8 | 93.5 ± 1.6 | 0.9998 |
| Homoalanine | 106.9 ± 1.8 | 87.5 ± 1.2 | 103.8 ± 1.2 | 105.3 ± 2.5 | 99.0 ± 1.4 | 106.0 ± 3.0 | 0.9994 |
| Homoarginine | 101.2 ± 2.1 | 95.3 ± 0.3 | 110.7 ± 0.6 | 88.8 ± 0.3 | 92.4 ± 1.7 | 114.1 ± 1.8 | 0.9998 |
| Homocysteine | 112.1 ± 2.0 | 99.5 ± 0.6 | 101.5 ± 1.2 | 92.5 ± 1.2 | 93.3 ± 2.2 | 93.6 ± 2.8 | 0.9994 |
| Homogentisic acid | 109.1 ± 1.9 | 88.2 ± 1.1 | 85.1 ± 2.5 | 113.9 ± 1.1 | 101.3 ± 0.8 | 111.9 ± 2.6 | 0.9997 |
| Homoleucine | 113.4 ± 1.1 | 90.7 ± 3.0 | 101.5 ± 1.0 | 107.2 ± 1.5 | 103.5 ± 2.5 | 109.3 ± 2.0 | 0.9997 |
| Homovanillic acid | 87.8 ± 2.1 | 97.1 ± 2.0 | 94.7 ± 2.4 | 101.1 ± 2.0 | 88.2 ± 1.6 | 87.2 ± 1.7 | 0.9996 |
| Hydantoin-5-propionate | 112.3 ± 1.4 | 95.7 ± 1.9 | 113.3 ± 2.2 | 114.6 ± 1.3 | 93.3 ± 1.5 | 104.9 ± 3.0 | 0.9991 |
| Hypotaurine | 99.0 ± 1.9 | 100.0 ± 2.0 | 95.8 ± 1.7 | 87.3 ± 0.4 | 112.4 ± 1.8 | 104.7 ± 0.8 | 0.9999 |
| Hypoxanthine | 105.5 ± 1.6 | 100.3 ± 2.8 | 113.8 ± 2.2 | 114.1 ± 0.6 | 93.0 ± 2.6 | 112.8 ± 1.3 | 0.9994 |
| Indole-3-acetamide | 101.5 ± 0.7 | 100.4 ± 0.4 | 92.0 ± 2.9 | 96.1 ± 3.2 | 114.5 ± 1.5 | 114.8 ± 1.9 | 0.9996 |
| Inosine | 91.5 ± 2.8 | 114.3 ± 0.6 | 106.7 ± 1.7 | 97.2 ± 2.5 | 111.3 ± 1.8 | 93.4 ± 1.5 | 0.9994 |
| Inosine 5'-monophosphate | 104.4 ± 3.0 | 109.4 ± 2.5 | 106.1 ± 2.0 | 105.2 ± 0.8 | 96.8 ± 1.9 | 85.5 ± 3.0 | 0.9997 |
| Isocitric Acid | 107.4 ± 0.5 | 105.3 ± 2.1 | 103.0 ± 1.5 | 97.4 ± 1.9 | 86.2 ± 2.0 | 86.0 ± 2.1 | 0.9992 |
| Isoleucine | 103.1 ± 3.1 | 96.5 ± 1.9 | 95.8 ± 3.0 | 110.0 ± 1.5 | 103.0 ± 1.8 | 87.6 ± 1.2 | 0.9995 |
| Lactic Acid | 86.0 ± 1.2 | 98.8 ± 1.7 | 104.8 ± 3.1 | 94.5 ± 2.2 | 99.6 ± 2.3 | 105.0 ± 1.4 | 0.9994 |
| Leucine | 103.2 ± 2.3 | 97.4 ± 2.2 | 111.6 ± 1.6 | 89.3 ± 0.9 | 106.0 ± 0.4 | 89.8 ± 0.6 | 0.9991 |
| Levulinic acid | 109.5 ± 2.6 | 89.0 ± 0.5 | 114.5 ± 0.6 | 112.2 ± 0.8 | 97.9 ± 1.8 | 88.9 ± 1.2 | 0.9996 |
| Lysine | 106.3 ± 0.7 | 110.5 ± 2.5 | 112.8 ± 2.4 | 105.5 ± 3.2 | 87.4 ± 1.5 | 106.4 ± 1.9 | 0.9999 |
| Malic Acid | 98.0 ± 2.1 | 85.7 ± 0.6 | 93.0 ± 2.9 | 110.4 ± 1.4 | 85.3 ± 1.8 | 98.6 ± 2.5 | 0.9997 |
| Mannose-1 with Mannose-6-Phosphate | 95.3 ± 1.3 | 108.8 ± 1.3 | 95.1 ± 1.2 | 106.8 ± 1.1 | 107.0 ± 1.6 | 87.6 ± 1.6 | 0.9992 |
| Mesaconic acid | 93.2 ± 1.2 | 105.6 ± 1.5 | 108.5 ± 0.7 | 91.3 ± 1.0 | 108.2 ± 1.5 | 104.2 ± 2.9 | 0.9997 |
| Methionine | 90.2 ± 2.0 | 107.2 ± 2.7 | 110.5 ± 2.7 | 86.4 ± 1.0 | 105.2 ± 3.0 | 98.8 ± 0.6 | 0.9994 |
| Methionine sulfoxide | 111.5 ± 1.3 | 92.4 ± 0.8 | 95.4 ± 2.5 | 114.5 ± 3.2 | 109.0 ± 2.6 | 101.9 ± 0.4 | 0.9997 |
| Methylmalonic Acid | 113.9 ± 1.3 | 101.4 ± 1.7 | 96.4 ± 2.4 | 85.6 ± 0.7 | 111.1 ± 2.6 | 109.7 ± 2.9 | 0.9994 |
| N1-Acetylspermidine | 93.1 ± 0.8 | 89.2 ± 2.5 | 112.8 ± 0.6 | 86.3 ± 1.0 | 99.2 ± 0.5 | 106.1 ± 3.0 | 0.9991 |
| N1-Acetylspermine | 95.2 ± 1.8 | 90.9 ± 2.3 | 95.1 ± 1.7 | 96.2 ± 1.4 | 106.8 ± 2.4 | 91.9 ± 0.4 | 0.9995 |
| N8-Acetylspermidine | 106.4 ± 1.5 | 91.6 ± 1.7 | 106.4 ± 1.0 | 106.3 ± 0.7 | 99.9 ± 2.6 | 85.6 ± 2.6 | 0.9995 |
| N-Acetyl-Aspartyl-Glutamate | 109.1 ± 2.6 | 95.1 ± 0.4 | 87.4 ± 0.6 | 106.2 ± 2.6 | 107.7 ± 1.2 | 97.8 ± 2.1 | 0.9995 |
| N-Acetyl-Aspartyl-Glutamyl-Glutamate | 112.5 ± 1.8 | 112.9 ± 0.5 | 105.5 ± 1.9 | 110.3 ± 1.8 | 92.3 ± 2.8 | 89.1 ± 2.1 | 0.9994 |
| N-Acetyl-Galactosamine | 92.9 ± 2.8 | 98.1 ± 0.9 | 100.6 ± 3.2 | 111.4 ± 0.9 | 88.3 ± 0.9 | 85.3 ± 1.9 | 0.9993 |
| N-Acetyl-Leucine | 110.8 ± 1.9 | 103.9 ± 2.1 | 98.6 ± 1.7 | 106.9 ± 1.1 | 99.2 ± 2.8 | 95.3 ± 2.7 | 0.9997 |
| N-Acetyl-Methionine | 85.3 ± 1.1 | 99.6 ± 2.6 | 107.9 ± 2.2 | 93.0 ± 3.0 | 93.9 ± 2.4 | 90.7 ± 2.3 | 0.9998 |
| N-Acetylneuraminic acid | 95.1 ± 0.4 | 106.1 ± 2.6 | 86.6 ± 0.5 | 94.5 ± 3.1 | 106.5 ± 2.2 | 103.7 ± 2.9 | 0.9991 |
| N-Acetyl-Proline | 103.9 ± 0.6 | 92.6 ± 1.0 | 99.5 ± 2.7 | 101.5 ± 2.4 | 114.2 ± 1.2 | 101.4 ± 2.6 | 0.9994 |
| N-Acetyl-Putrescine | 114.4 ± 0.7 | 102.0 ± 0.4 | 85.0 ± 2.9 | 90.4 ± 1.5 | 101.9 ± 1.4 | 107.8 ± 1.9 | 0.9998 |
| N-Acetyl-Serine | 102.1 ± 0.9 | 103.3 ± 2.1 | 112.0 ± 2.9 | 97.9 ± 1.4 | 110.3 ± 3.1 | 114.2 ± 2.4 | 0.9993 |
| N-Acetyl-Serotonin | 107.6 ± 2.8 | 103.7 ± 1.7 | 113.7 ± 2.5 | 110.0 ± 2.7 | 92.2 ± 0.3 | 90.9 ± 2.8 | 0.9991 |
| N-Acyl-Homoserine-lactone | 111.3 ± 2.0 | 87.1 ± 0.6 | 105.7 ± 1.7 | 91.1 ± 2.1 | 85.2 ± 1.9 | 101.3 ± 1.9 | 0.9998 |
| NAD | 90.7 ± 0.8 | 104.0 ± 2.1 | 88.2 ± 0.4 | 114.3 ± 0.5 | 103.2 ± 0.8 | 87.1 ± 1.3 | 0.9992 |
| NADH | 99.5 ± 2.0 | 88.2 ± 3.1 | 87.0 ± 0.9 | 111.9 ± 1.9 | 103.0 ± 1.1 | 98.0 ± 2.1 | 0.9997 |
| NADP | 91.8 ± 1.6 | 92.2 ± 1.9 | 100.2 ± 3.0 | 114.9 ± 2.8 | 114.2 ± 0.3 | 112.5 ± 2.9 | 0.9997 |
| NADPH | 107.2 ± 1.1 | 94.6 ± 2.0 | 92.9 ± 1.0 | 109.8 ± 1.8 | 93.2 ± 0.7 | 101.5 ± 2.1 | 0.9993 |
| N-alpha-Acetyl-lysine | 105.9 ± 3.2 | 97.4 ± 0.9 | 92.7 ± 3.1 | 105.6 ± 2.1 | 102.8 ± 2.8 | 87.2 ± 3.1 | 0.9994 |
| N-alpha-Acetyl-Ornithine | 95.9 ± 2.4 | 97.2 ± 2.4 | 92.4 ± 2.1 | 110.6 ± 2.3 | 103.8 ± 2.6 | 102.6 ± 2.0 | 0.9997 |
| N-Carbamoyl-Proline | 86.9 ± 2.1 | 107.0 ± 1.7 | 99.0 ± 1.3 | 114.0 ± 1.6 | 109.9 ± 1.5 | 113.6 ± 1.0 | 0.9998 |
| N-Carbamoyl-Sarcosine | 90.3 ± 1.5 | 98.5 ± 1.3 | 90.0 ± 1.6 | 90.4 ± 1.7 | 88.2 ± 2.0 | 109.9 ± 2.6 | 0.9997 |
| N-Epsilon-Acetyl-lysine | 92.5 ± 2.7 | 93.3 ± 1.6 | 88.9 ± 1.0 | 107.3 ± 2.0 | 106.7 ± 3.1 | 105.0 ± 2.6 | 0.9996 |
| Nicotinamide | 99.9 ± 1.0 | 113.6 ± 1.2 | 90.1 ± 1.3 | 98.6 ± 0.4 | 92.7 ± 2.5 | 97.4 ± 0.9 | 0.9991 |
| N-Methyl-Aspartic Acid | 89.4 ± 1.1 | 111.0 ± 2.9 | 97.4 ± 1.6 | 113.5 ± 1.1 | 85.5 ± 2.0 | 87.2 ± 1.1 | 0.9998 |
| O-Phosphoethanolamine | 98.5 ± 1.3 | 101.6 ± 2.8 | 111.7 ± 2.0 | 106.0 ± 2.8 | 112.3 ± 1.4 | 107.6 ± 0.6 | 0.9994 |
| Ophthalmic acid | 90.6 ± 3.0 | 104.9 ± 1.6 | 102.1 ± 1.4 | 94.9 ± 2.1 | 107.3 ± 1.8 | 108.2 ± 1.1 | 0.9997 |
| Ornithine | 97.8 ± 2.7 | 107.4 ± 1.1 | 99.4 ± 1.8 | 100.3 ± 0.5 | 91.8 ± 2.9 | 96.9 ± 2.1 | 0.9993 |
| Orotidine 5'-monophosphate | 87.2 ± 1.1 | 106.3 ± 2.6 | 88.6 ± 2.1 | 105.6 ± 0.3 | 94.2 ± 1.2 | 114.0 ± 2.2 | 0.9996 |
| Oxaloacetate | 96.3 ± 0.8 | 89.6 ± 1.8 | 109.0 ± 0.8 | 95.0 ± 3.2 | 107.2 ± 1.9 | 106.8 ± 1.0 | 0.9993 |
| Pantothenic acid | 106.5 ± 1.8 | 95.3 ± 0.3 | 112.3 ± 1.2 | 93.7 ± 1.9 | 110.2 ± 0.3 | 93.8 ± 2.1 | 0.9991 |
| Phenylalanine | 107.4 ± 2.8 | 110.8 ± 2.1 | 107.1 ± 2.1 | 94.9 ± 2.0 | 114.0 ± 2.0 | 96.5 ± 3.2 | 0.9997 |
| Phenylpyruvate | 111.8 ± 0.4 | 114.5 ± 2.9 | 102.4 ± 3.2 | 110.7 ± 3.0 | 103.8 ± 1.4 | 111.4 ± 0.7 | 0.9991 |
| Phosphodimethylethanolamine | 96.6 ± 2.2 | 95.1 ± 2.4 | 103.5 ± 1.4 | 92.9 ± 2.2 | 102.4 ± 1.1 | 86.6 ± 2.3 | 0.9996 |
| Phosphoenolpyruvate | 114.7 ± 1.9 | 109.6 ± 1.2 | 98.5 ± 1.3 | 97.4 ± 0.3 | 88.6 ± 3.1 | 103.6 ± 2.4 | 0.9993 |
| Phosphorylcholine | 94.5 ± 0.4 | 104.5 ± 0.8 | 112.3 ± 1.6 | 103.2 ± 2.9 | 98.9 ± 2.7 | 106.8 ± 3.0 | 0.9996 |
| Pipecolic acid | 102.5 ± 1.5 | 93.1 ± 3.1 | 103.3 ± 1.7 | 106.9 ± 2.2 | 108.4 ± 1.7 | 85.3 ± 0.8 | 0.9993 |
| Proline | 98.4 ± 1.8 | 99.5 ± 2.0 | 102.0 ± 0.9 | 96.0 ± 2.0 | 112.6 ± 2.5 | 85.2 ± 0.7 | 0.9996 |
| Putrescine | 88.6 ± 1.0 | 114.6 ± 1.8 | 105.4 ± 2.1 | 111.2 ± 2.1 | 114.6 ± 2.4 | 85.9 ± 0.8 | 0.9997 |
| Pyridoxal | 86.6 ± 2.8 | 100.4 ± 3.2 | 106.3 ± 3.1 | 91.5 ± 2.1 | 112.6 ± 0.8 | 112.9 ± 1.0 | 0.9998 |
| Pyridoxamine | 90.7 ± 0.4 | 87.8 ± 0.8 | 96.6 ± 0.9 | 88.7 ± 2.0 | 100.5 ± 0.7 | 109.6 ± 2.9 | 0.9991 |
| Pyridoxine | 86.0 ± 1.6 | 101.0 ± 1.5 | 95.7 ± 2.5 | 104.1 ± 2.8 | 90.7 ± 2.9 | 97.0 ± 1.1 | 0.9993 |
| Pyroglutamic acid | 110.0 ± 2.7 | 92.4 ± 2.4 | 86.8 ± 1.3 | 109.9 ± 0.9 | 108.3 ± 1.9 | 86.1 ± 1.7 | 0.9996 |
| Pyrrolysine | 92.0 ± 1.2 | 109.2 ± 1.1 | 109.5 ± 0.7 | 113.0 ± 0.8 | 89.5 ± 2.4 | 106.9 ± 2.9 | 0.9993 |
| Pyruvic Acid | 104.3 ± 1.8 | 105.6 ± 2.8 | 106.8 ± 1.7 | 102.1 ± 1.9 | 105.8 ± 2.1 | 90.7 ± 1.6 | 0.9996 |
| Riboflavin | 86.1 ± 2.4 | 107.0 ± 2.4 | 107.6 ± 1.1 | 94.1 ± 0.9 | 96.0 ± 1.7 | 105.2 ± 2.5 | 0.9999 |
| Ribose-5-Phosphate | 101.9 ± 0.6 | 92.7 ± 1.9 | 97.2 ± 3.0 | 110.5 ± 0.7 | 108.7 ± 3.0 | 86.9 ± 0.6 | 0.9992 |
| Ribulose-1-Phosphate | 99.5 ± 1.0 | 101.8 ± 0.6 | 110.1 ± 0.5 | 96.0 ± 1.9 | 110.0 ± 2.2 | 88.3 ± 1.4 | 0.9996 |
| Ribulose-5-Phosphate | 88.8 ± 0.5 | 92.1 ± 2.2 | 114.6 ± 2.3 | 85.7 ± 0.7 | 100.1 ± 0.8 | 93.7 ± 3.1 | 0.9991 |
| Saccharopine | 98.0 ± 1.8 | 112.1 ± 0.6 | 92.8 ± 2.8 | 90.0 ± 1.6 | 88.8 ± 2.5 | 113.6 ± 2.9 | 0.9994 |
| S-Adenosylhomocysteine | 110.9 ± 1.5 | 105.2 ± 2.2 | 87.0 ± 2.0 | 103.9 ± 0.4 | 108.5 ± 1.0 | 112.5 ± 0.6 | 0.9998 |
| S-Adenosylmethioninamine | 85.5 ± 2.0 | 98.4 ± 0.8 | 103.7 ± 1.8 | 94.9 ± 2.3 | 91.1 ± 0.4 | 107.7 ± 2.8 | 0.9995 |
| S-Adenosylmethionine | 106.4 ± 1.8 | 113.6 ± 2.7 | 95.6 ± 0.9 | 111.3 ± 1.3 | 110.9 ± 0.4 | 114.2 ± 2.0 | 0.9996 |
| Sarcosine | 111.6 ± 1.2 | 95.4 ± 3.2 | 114.0 ± 3.1 | 90.3 ± 2.7 | 88.5 ± 0.8 | 93.4 ± 2.6 | 0.9995 |
| Sedoheptulose-7-phosphate | 96.7 ± 2.4 | 93.9 ± 1.7 | 85.4 ± 2.9 | 102.8 ± 3.0 | 99.2 ± 2.3 | 85.7 ± 2.5 | 0.9997 |
| Selenohomocysteine | 91.8 ± 2.0 | 98.3 ± 1.6 | 88.1 ± 3.1 | 85.2 ± 1.4 | 106.4 ± 1.7 | 95.9 ± 1.0 | 0.9994 |
| Selenomethionine | 94.4 ± 1.3 | 99.9 ± 0.3 | 99.5 ± 2.9 | 85.3 ± 2.4 | 105.4 ± 0.8 | 109.4 ± 1.5 | 0.9998 |
| Serine | 96.8 ± 0.5 | 102.6 ± 2.9 | 90.7 ± 0.9 | 86.4 ± 1.4 | 114.5 ± 2.5 | 86.1 ± 0.9 | 0.9994 |
| S-Methyl-Thioinosine | 105.1 ± 2.8 | 98.7 ± 0.5 | 108.5 ± 0.5 | 92.4 ± 1.6 | 104.1 ± 1.9 | 95.5 ± 1.2 | 0.9993 |
| Spermidine | 88.5 ± 1.3 | 114.7 ± 0.8 | 94.1 ± 0.9 | 95.6 ± 2.8 | 110.5 ± 1.3 | 89.3 ± 2.0 | 0.9997 |
| Spermine | 97.8 ± 1.0 | 102.1 ± 1.6 | 95.6 ± 2.8 | 97.6 ± 2.9 | 108.0 ± 0.5 | 103.3 ± 2.8 | 0.9997 |
| Succinate | 114.0 ± 1.8 | 94.5 ± 0.6 | 101.5 ± 1.8 | 112.0 ± 1.9 | 101.2 ± 2.1 | 101.3 ± 3.1 | 0.9994 |
| Sucrose | 110.3 ± 1.8 | 88.6 ± 2.2 | 85.3 ± 1.9 | 93.4 ± 0.8 | 91.8 ± 2.6 | 100.5 ± 0.8 | 0.9998 |
| Symmetric dimethylarginine | 86.8 ± 1.5 | 94.5 ± 0.6 | 106.9 ± 2.0 | 101.7 ± 0.6 | 95.5 ± 1.3 | 99.0 ± 1.0 | 0.9995 |
| Taurine | 98.5 ± 1.5 | 87.8 ± 2.8 | 95.6 ± 1.6 | 92.8 ± 1.8 | 91.8 ± 2.0 | 89.9 ± 0.4 | 0.9998 |
| Tetrahydroneopterin | 104.2 ± 0.8 | 113.8 ± 1.5 | 97.6 ± 3.1 | 93.3 ± 2.2 | 88.7 ± 0.5 | 113.8 ± 1.0 | 0.9997 |
| Thiamine | 92.6 ± 2.3 | 104.2 ± 1.0 | 109.4 ± 3.2 | 98.1 ± 1.3 | 87.6 ± 0.3 | 91.7 ± 2.3 | 0.9996 |
| Thiamine monophosphate | 108.7 ± 1.4 | 111.5 ± 3.0 | 114.9 ± 2.2 | 88.2 ± 1.8 | 104.9 ± 1.4 | 102.0 ± 3.0 | 0.9996 |
| Threonic acid | 88.2 ± 1.9 | 109.6 ± 1.8 | 99.4 ± 0.4 | 106.3 ± 1.6 | 93.2 ± 1.2 | 96.2 ± 2.0 | 0.9995 |
| Threonine | 114.8 ± 2.3 | 93.7 ± 2.2 | 104.4 ± 1.6 | 96.4 ± 1.9 | 110.6 ± 1.7 | 107.9 ± 2.2 | 0.9995 |
| Thymidine | 107.8 ± 1.8 | 111.3 ± 0.7 | 101.6 ± 1.5 | 100.9 ± 1.0 | 103.3 ± 2.5 | 96.2 ± 0.8 | 0.9994 |
| Thymidine Monophosphate | 100.3 ± 2.5 | 86.8 ± 0.7 | 113.5 ± 2.2 | 102.2 ± 2.7 | 98.0 ± 1.3 | 92.3 ± 2.7 | 0.9994 |
| Thymine | 112.3 ± 2.7 | 98.9 ± 2.9 | 107.1 ± 1.3 | 114.0 ± 1.1 | 106.6 ± 1.9 | 89.1 ± 1.7 | 0.9999 |
| Trigonelleine | 114.8 ± 1.6 | 87.5 ± 0.8 | 112.1 ± 1.6 | 101.2 ± 1.9 | 86.4 ± 0.8 | 95.9 ± 1.3 | 0.9992 |
| Trimethylamine N-oxide | 85.1 ± 1.0 | 95.1 ± 0.4 | 86.3 ± 3.0 | 96.6 ± 1.0 | 103.3 ± 0.6 | 86.1 ± 2.8 | 0.9996 |
| Trimethyllysine | 112.2 ± 1.6 | 97.9 ± 1.0 | 99.5 ± 0.5 | 87.3 ± 0.5 | 85.7 ± 2.6 | 89.6 ± 2.3 | 0.9999 |
| Tryptophan | 99.4 ± 1.2 | 96.0 ± 0.8 | 93.3 ± 1.2 | 113.2 ± 1.2 | 99.1 ± 2.0 | 100.0 ± 1.8 | 0.9995 |
| Tyramine | 103.5 ± 0.5 | 94.8 ± 1.0 | 111.1 ± 2.8 | 92.7 ± 2.5 | 94.4 ± 1.4 | 110.3 ± 0.6 | 0.9994 |
| Tyrosine | 104.5 ± 2.0 | 97.1 ± 2.2 | 113.3 ± 2.0 | 108.2 ± 2.9 | 96.4 ± 0.8 | 100.8 ± 0.9 | 0.9995 |
| Uracil | 105.3 ± 0.6 | 88.9 ± 0.4 | 103.5 ± 1.9 | 112.5 ± 3.1 | 100.2 ± 1.1 | 100.3 ± 1.3 | 0.9992 |
| Uridine 5'-diphosphate | 108.1 ± 2.2 | 105.3 ± 0.9 | 87.7 ± 0.6 | 89.6 ± 2.8 | 85.9 ± 2.9 | 104.2 ± 3.0 | 0.9996 |
| Uridine Monophosphate | 87.5 ± 0.8 | 106.9 ± 1.4 | 110.5 ± 2.1 | 93.9 ± 3.0 | 96.3 ± 0.5 | 93.5 ± 0.7 | 0.9996 |
| Uridine-5-Diphosphogalactose | 100.6 ± 2.0 | 88.0 ± 1.5 | 93.6 ± 1.8 | 111.4 ± 1.2 | 103.4 ± 3.0 | 87.5 ± 2.8 | 0.9992 |
| Urocanic acid | 100.9 ± 2.6 | 96.5 ± 2.4 | 96.5 ± 2.2 | 113.8 ± 1.0 | 113.1 ± 2.4 | 99.8 ± 1.4 | 0.9992 |
| UTP | 105.6 ± 0.8 | 105.9 ± 3.1 | 95.1 ± 2.4 | 106.1 ± 2.5 | 99.4 ± 1.2 | 103.4 ± 2.4 | 0.9991 |
| Valine | 113.1 ± 1.1 | 99.4 ± 2.2 | 114.6 ± 2.1 | 97.9 ± 2.0 | 85.3 ± 0.4 | 98.9 ± 1.4 | 0.9994 |
| XMP | 107.8 ± 2.3 | 110.0 ± 1.5 | 96.3 ± 2.9 | 104.6 ± 1.9 | 113.1 ± 1.3 | 113.1 ± 2.8 | 0.9995 |
| Xylulose-5-Phosphate | 91.7 ± 1.6 | 96.2 ± 0.7 | 109.5 ± 2.7 | 104.9 ± 2.8 | 86.4 ± 1.3 | 87.8 ± 1.6 | 0.9992 |

### **Table S9** | Calibration values for final method conditions (r^2^-value) and percent found of nominal calibrator concentration (Plasma extract as matrix).

| Calibrator concentration accuracy  [%] | 10 ng/mL | 25 ng/mL | 50 ng/mL | 100 ng/mL | 500 ng/mL | 1000 ng/mL | r^2^-Value  matrix-matched calibration |
| --- | --- | --- | --- | --- | --- | --- | --- |
| 1-Methylguanine | 106.0 ± 1.1 | 109.2 ± 1.3 | 103.7 ± 2.7 | 101.7 ± 2.3 | 92.9 ± 2.0 | 98.6 ± 1.5 | 0.9992 |
| 1-Methylhistidine | 99.9 ± 2.7 | 100.9 ± 0.8 | 110.7 ± 1.8 | 94.3 ± 2.2 | 106.7 ± 0.6 | 87.4 ± 0.3 | 0.9995 |
| 1-Methylnicotinamide | 110.0 ± 0.6 | 100.0 ± 1.4 | 114.6 ± 0.7 | 112.4 ± 2.2 | 111.1 ± 0.4 | 109.2 ± 2.5 | 0.9995 |
| 2-Aminoadipic Acid | 109.6 ± 3.1 | 100.7 ± 0.6 | 99.4 ± 1.4 | 89.0 ± 0.7 | 102.8 ± 1.5 | 113.9 ± 0.9 | 0.9998 |
| 2-Deoxyadenosine | 111.6 ± 1.2 | 113.7 ± 3.0 | 109.7 ± 0.9 | 95.7 ± 1.9 | 90.7 ± 1.0 | 100.2 ± 1.6 | 0.9996 |
| 2'-Deoxyadenosine 5'-monophosphate | 101.6 ± 1.5 | 89.7 ± 2.7 | 100.2 ± 2.7 | 97.9 ± 1.7 | 89.2 ± 2.9 | 109.3 ± 0.6 | 0.9996 |
| 2'-deoxyadenosine 5'-triphosphate | 112.5 ± 2.4 | 114.7 ± 1.0 | 103.6 ± 2.0 | 108.6 ± 2.6 | 93.7 ± 3.2 | 107.0 ± 2.7 | 0.9995 |
| 2'-Deoxycytidine-5'-monophosphate | 93.4 ± 1.6 | 97.3 ± 0.6 | 104.6 ± 1.1 | 95.9 ± 1.2 | 102.2 ± 1.2 | 114.7 ± 2.5 | 0.9998 |
| 2-Deoxyguanosine | 91.3 ± 2.7 | 111.7 ± 0.7 | 99.4 ± 0.5 | 100.2 ± 2.8 | 109.5 ± 1.4 | 89.5 ± 0.9 | 0.9992 |
| 2-hydroxybutyric acid | 95.5 ± 0.7 | 92.2 ± 1.3 | 106.8 ± 1.1 | 103.5 ± 2.6 | 103.7 ± 1.9 | 102.8 ± 0.5 | 0.9993 |
| 2-hydroxyglutaric acid | 101.9 ± 1.3 | 104.4 ± 2.1 | 94.4 ± 1.0 | 92.1 ± 1.0 | 112.7 ± 2.7 | 94.9 ± 2.3 | 0.9994 |
| 2-Hydroxyisocaproic acid | 88.9 ± 1.8 | 92.5 ± 1.3 | 113.2 ± 0.5 | 95.0 ± 1.0 | 110.2 ± 0.7 | 109.5 ± 2.8 | 0.9992 |
| 2-ketobutyric acid | 88.9 ± 2.4 | 97.6 ± 1.0 | 101.3 ± 1.2 | 108.7 ± 2.7 | 104.6 ± 1.4 | 90.4 ± 0.9 | 0.9998 |
| 2-oxoadipic acid | 90.9 ± 1.5 | 85.3 ± 2.9 | 107.3 ± 1.8 | 86.9 ± 1.1 | 103.4 ± 1.0 | 113.1 ± 2.7 | 0.9999 |
| 2-Phenylglycine | 105.2 ± 0.7 | 93.6 ± 2.0 | 101.7 ± 2.5 | 87.3 ± 1.7 | 104.5 ± 1.8 | 114.8 ± 2.1 | 0.9997 |
| 2-phosphoglycerate | 104.3 ± 3.1 | 89.2 ± 2.1 | 86.8 ± 2.4 | 97.3 ± 2.5 | 90.0 ± 0.6 | 113.6 ± 2.2 | 0.9996 |
| 3-hydroxy-3-methyl-2-oxobutanoate | 94.8 ± 1.6 | 96.4 ± 1.3 | 87.8 ± 2.3 | 112.8 ± 0.7 | 107.2 ± 1.3 | 104.6 ± 1.6 | 0.9992 |
| 3-hydroxybutyric acid | 103.2 ± 1.3 | 112.0 ± 1.7 | 111.4 ± 3.2 | 107.3 ± 2.5 | 113.7 ± 2.0 | 96.4 ± 2.2 | 0.9998 |
| 3-Methylhistidine | 89.4 ± 1.5 | 92.5 ± 2.7 | 102.9 ± 1.3 | 91.4 ± 2.6 | 92.4 ± 3.0 | 113.9 ± 2.0 | 0.9992 |
| 3-Phosphoglyceraldehyde | 102.2 ± 3.1 | 111.9 ± 1.3 | 101.7 ± 2.5 | 92.1 ± 1.2 | 85.1 ± 2.8 | 99.6 ± 2.3 | 0.9995 |
| 3-phosphoglycerate | 92.0 ± 2.3 | 103.3 ± 2.3 | 104.2 ± 0.6 | 94.8 ± 1.8 | 99.8 ± 2.7 | 87.3 ± 0.5 | 0.9995 |
| 4-amino-butyric acid | 114.7 ± 1.0 | 94.1 ± 2.5 | 86.5 ± 2.9 | 86.4 ± 2.6 | 113.4 ± 1.9 | 88.6 ± 1.3 | 0.9993 |
| 4-aspartyl-phosphate | 109.0 ± 1.0 | 104.4 ± 3.0 | 86.4 ± 2.7 | 89.3 ± 2.5 | 90.7 ± 0.5 | 90.5 ± 3.1 | 0.9995 |
| 4-Hydroxybenzoic acid | 96.5 ± 1.9 | 111.8 ± 1.5 | 108.3 ± 2.9 | 107.3 ± 1.3 | 97.5 ± 0.5 | 110.8 ± 1.6 | 0.9992 |
| 4-Hydroxyphenylacetylglycine | 90.0 ± 1.5 | 113.1 ± 3.1 | 111.7 ± 0.7 | 94.3 ± 0.8 | 88.3 ± 0.9 | 90.5 ± 0.4 | 0.9997 |
| 4-methyl-2-oxopentanoate | 112.6 ± 1.5 | 102.0 ± 1.2 | 104.7 ± 0.7 | 94.2 ± 1.7 | 103.4 ± 1.7 | 113.6 ± 1.3 | 0.9991 |
| 4-Oxoglutaramate | 85.9 ± 1.7 | 111.7 ± 1.1 | 93.9 ± 3.1 | 87.3 ± 0.3 | 103.9 ± 2.2 | 102.5 ± 0.4 | 0.9997 |
| 5-Hydroxyindolepyruvate | 96.5 ± 0.3 | 95.9 ± 2.7 | 94.9 ± 1.1 | 105.8 ± 2.2 | 110.5 ± 1.3 | 113.5 ± 0.7 | 0.9994 |
| 5-Hydroxylysine | 96.7 ± 0.4 | 89.9 ± 0.8 | 103.6 ± 2.0 | 106.9 ± 1.2 | 104.2 ± 2.7 | 87.0 ± 2.7 | 0.9993 |
| 5-Hydroxy-Tryptophan | 99.8 ± 2.4 | 86.2 ± 2.6 | 100.6 ± 2.9 | 99.8 ± 2.5 | 104.5 ± 1.5 | 107.0 ± 2.9 | 0.9997 |
| 5-Phosphoribosylamine | 87.2 ± 0.7 | 89.8 ± 2.1 | 85.1 ± 1.3 | 88.6 ± 0.9 | 100.1 ± 0.9 | 95.5 ± 1.3 | 0.9992 |
| 6-phosphogluconate | 113.4 ± 1.5 | 110.1 ± 1.6 | 91.5 ± 3.1 | 89.7 ± 0.4 | 107.7 ± 1.4 | 111.0 ± 2.2 | 0.9994 |
| Acetylcarnitine | 85.2 ± 1.0 | 93.6 ± 0.8 | 113.8 ± 1.7 | 93.4 ± 2.8 | 107.4 ± 2.6 | 96.7 ± 1.7 | 0.9993 |
| Aconitic Acid | 110.3 ± 1.5 | 103.5 ± 2.3 | 97.0 ± 1.6 | 103.5 ± 2.1 | 87.2 ± 2.3 | 96.1 ± 0.5 | 0.9994 |
| Adenine | 90.8 ± 0.5 | 95.5 ± 2.7 | 105.1 ± 3.1 | 88.9 ± 2.9 | 94.6 ± 1.0 | 94.8 ± 2.0 | 0.9994 |
| Adenosine | 95.1 ± 0.7 | 98.8 ± 1.5 | 102.0 ± 0.7 | 90.0 ± 3.0 | 86.1 ± 1.2 | 112.5 ± 1.4 | 0.9992 |
| Adenosine 3',5'-cyclic monophosphate | 112.4 ± 2.6 | 86.8 ± 1.4 | 110.6 ± 2.5 | 91.9 ± 1.3 | 95.4 ± 1.1 | 87.8 ± 2.5 | 0.9991 |
| Adenosine Monophosphate | 110.4 ± 2.7 | 93.8 ± 0.6 | 89.0 ± 2.1 | 92.4 ± 0.6 | 110.4 ± 1.3 | 88.0 ± 2.4 | 0.9998 |
| Adenosine-5'-diphosphate | 88.7 ± 2.1 | 99.7 ± 1.0 | 104.2 ± 2.2 | 95.4 ± 2.5 | 103.2 ± 1.6 | 113.4 ± 0.3 | 0.9994 |
| Adenosine-5'-triphosphate | 88.3 ± 1.4 | 111.3 ± 0.9 | 91.0 ± 0.8 | 87.2 ± 2.2 | 112.2 ± 0.7 | 95.0 ± 0.4 | 0.9994 |
| Adenylosuccinate | 86.3 ± 2.9 | 95.4 ± 0.6 | 108.3 ± 3.2 | 90.4 ± 0.5 | 93.9 ± 0.3 | 92.1 ± 0.6 | 0.9993 |
| Alanine | 106.2 ± 1.9 | 92.8 ± 0.4 | 105.2 ± 0.8 | 97.6 ± 2.3 | 104.4 ± 1.7 | 95.1 ± 2.9 | 0.9998 |
| alpha-ketoglutaric acid | 91.7 ± 2.2 | 109.1 ± 1.9 | 114.5 ± 1.7 | 102.1 ± 0.8 | 101.1 ± 1.5 | 102.7 ± 3.1 | 0.9997 |
| Anserine | 109.7 ± 1.1 | 89.9 ± 2.8 | 105.3 ± 2.2 | 91.5 ± 1.8 | 112.6 ± 0.6 | 105.5 ± 2.0 | 0.9996 |
| Arginine | 88.7 ± 1.3 | 99.2 ± 0.8 | 105.5 ± 3.0 | 88.5 ± 0.8 | 110.9 ± 1.1 | 112.1 ± 1.0 | 0.9995 |
| Argininosuccinic Acid | 106.4 ± 1.2 | 111.7 ± 1.1 | 105.5 ± 2.8 | 89.6 ± 1.4 | 101.2 ± 1.2 | 95.2 ± 1.3 | 0.9996 |
| Asparagine | 114.0 ± 2.1 | 104.8 ± 1.5 | 114.5 ± 2.9 | 100.7 ± 0.7 | 109.1 ± 1.9 | 100.4 ± 2.4 | 0.9993 |
| Aspartic acid | 111.4 ± 2.5 | 98.3 ± 0.9 | 86.5 ± 1.9 | 93.0 ± 1.3 | 98.4 ± 3.1 | 85.7 ± 0.4 | 0.9992 |
| Asymmetric dimethylarginine | 97.5 ± 1.1 | 110.6 ± 1.6 | 110.2 ± 1.2 | 111.4 ± 0.6 | 90.1 ± 0.4 | 96.0 ± 2.9 | 0.9996 |
| Azelaic acid | 88.4 ± 2.0 | 95.1 ± 1.4 | 110.3 ± 2.5 | 100.8 ± 2.2 | 85.8 ± 1.4 | 109.6 ± 2.9 | 0.9998 |
| beta-Alanine | 93.8 ± 3.2 | 114.3 ± 1.8 | 94.4 ± 2.5 | 85.5 ± 1.7 | 111.2 ± 2.6 | 92.9 ± 0.8 | 0.9993 |
| Betaine | 93.2 ± 0.6 | 108.4 ± 0.6 | 94.8 ± 1.7 | 107.5 ± 2.8 | 86.3 ± 1.0 | 107.2 ± 0.5 | 0.9992 |
| Biopterin | 107.7 ± 1.5 | 89.7 ± 2.7 | 96.0 ± 1.9 | 93.7 ± 2.2 | 92.8 ± 3.1 | 114.2 ± 1.1 | 0.9992 |
| Biotin | 112.4 ± 0.7 | 114.6 ± 2.6 | 86.8 ± 1.3 | 102.9 ± 1.8 | 113.0 ± 0.4 | 100.6 ± 2.4 | 0.9999 |
| Carnitine | 91.4 ± 1.3 | 102.0 ± 1.0 | 104.4 ± 0.7 | 96.4 ± 1.8 | 103.4 ± 1.8 | 110.4 ± 2.2 | 0.9998 |
| Carnosine | 102.3 ± 2.9 | 87.6 ± 1.7 | 107.7 ± 2.5 | 96.6 ± 0.7 | 111.8 ± 1.8 | 112.7 ± 3.1 | 0.9993 |
| CDP-choline | 106.0 ± 1.5 | 106.5 ± 0.3 | 93.2 ± 1.8 | 104.7 ± 2.2 | 89.5 ± 0.5 | 89.5 ± 3.1 | 0.9994 |
| cGMP | 106.8 ± 1.5 | 103.3 ± 1.9 | 91.7 ± 3.1 | 92.9 ± 3.2 | 93.2 ± 0.9 | 90.7 ± 1.5 | 0.9991 |
| Choline | 85.2 ± 1.1 | 107.0 ± 1.5 | 103.9 ± 1.9 | 100.6 ± 0.9 | 97.1 ± 1.1 | 103.5 ± 1.6 | 0.9999 |
| Cinnamic acid | 107.7 ± 1.3 | 90.5 ± 0.4 | 97.5 ± 2.6 | 113.8 ± 0.6 | 99.2 ± 0.9 | 104.2 ± 1.9 | 0.9994 |
| Citraconic acid | 113.5 ± 1.0 | 89.9 ± 0.4 | 111.3 ± 2.0 | 94.2 ± 2.8 | 97.3 ± 0.4 | 112.3 ± 0.9 | 0.9995 |
| Citramalic acid | 91.4 ± 1.4 | 90.8 ± 3.0 | 91.4 ± 2.7 | 90.8 ± 0.9 | 107.1 ± 2.3 | 91.7 ± 0.6 | 0.9996 |
| Citric Acid | 104.1 ± 0.5 | 98.7 ± 0.4 | 95.9 ± 1.4 | 90.4 ± 2.0 | 98.7 ± 3.1 | 98.4 ± 1.4 | 0.9998 |
| Citrulline | 110.3 ± 0.9 | 97.0 ± 1.0 | 92.1 ± 2.5 | 105.2 ± 1.1 | 103.0 ± 0.3 | 95.4 ± 0.9 | 0.9992 |
| Creatine | 107.6 ± 3.0 | 96.6 ± 2.5 | 112.0 ± 0.5 | 96.0 ± 2.7 | 106.8 ± 2.5 | 85.2 ± 2.4 | 0.9992 |
| Creatinine | 105.1 ± 0.9 | 88.9 ± 2.4 | 92.2 ± 0.7 | 93.1 ± 2.3 | 114.8 ± 1.8 | 89.2 ± 2.6 | 0.9998 |
| Cystathionine | 104.8 ± 0.4 | 108.9 ± 3.1 | 112.3 ± 2.1 | 104.7 ± 1.4 | 102.5 ± 1.8 | 111.8 ± 1.6 | 0.9997 |
| Cysteamine | 91.2 ± 2.0 | 89.3 ± 2.1 | 110.4 ± 1.3 | 99.2 ± 1.8 | 87.6 ± 3.0 | 106.0 ± 2.9 | 0.9994 |
| Cysteine | 106.8 ± 1.3 | 113.8 ± 1.1 | 112.6 ± 3.0 | 113.8 ± 1.6 | 114.3 ± 1.2 | 102.7 ± 0.5 | 0.9995 |
| Cysteinylglycine | 95.7 ± 0.8 | 100.7 ± 1.9 | 105.5 ± 0.8 | 91.5 ± 1.7 | 113.5 ± 0.6 | 102.4 ± 0.6 | 0.9994 |
| Cystine | 110.8 ± 0.8 | 86.5 ± 2.1 | 87.6 ± 2.8 | 100.9 ± 1.1 | 114.2 ± 0.4 | 99.6 ± 1.7 | 0.9994 |
| Cytidine | 88.8 ± 0.6 | 107.6 ± 2.4 | 106.9 ± 0.4 | 88.8 ± 2.4 | 91.2 ± 0.8 | 97.2 ± 2.4 | 0.9996 |
| Cytidine 3',5'-cyclic monophosphate | 112.0 ± 1.5 | 112.1 ± 0.5 | 95.8 ± 1.3 | 94.6 ± 1.8 | 110.6 ± 0.5 | 110.9 ± 1.2 | 0.9996 |
| Cytidine 5'-diphosphate | 111.9 ± 1.9 | 112.9 ± 2.1 | 87.1 ± 1.4 | 90.0 ± 3.1 | 98.8 ± 1.4 | 102.8 ± 2.6 | 0.9997 |
| Cytidine 5'-triphosphate | 111.9 ± 2.6 | 89.2 ± 0.7 | 100.8 ± 0.5 | 93.7 ± 2.2 | 104.4 ± 2.8 | 108.5 ± 2.8 | 0.9994 |
| Cytidine monophosphate | 93.7 ± 0.4 | 98.9 ± 2.6 | 88.5 ± 2.8 | 110.5 ± 2.1 | 90.3 ± 1.3 | 108.1 ± 1.0 | 0.9994 |
| Cytosine | 105.2 ± 2.7 | 107.4 ± 0.4 | 104.8 ± 0.4 | 111.3 ± 2.5 | 105.1 ± 1.9 | 108.0 ± 2.1 | 0.9992 |
| dADP | 97.9 ± 1.1 | 101.4 ± 1.2 | 105.0 ± 2.8 | 102.0 ± 2.7 | 98.0 ± 1.4 | 93.8 ± 2.2 | 0.9992 |
| Diethanolamine | 89.8 ± 1.4 | 110.8 ± 1.5 | 101.7 ± 2.8 | 112.5 ± 2.9 | 102.3 ± 1.0 | 101.8 ± 0.5 | 0.9997 |
| Dihydrobiopterin | 100.3 ± 1.8 | 90.8 ± 1.9 | 99.7 ± 2.2 | 112.6 ± 1.2 | 114.9 ± 2.1 | 107.5 ± 2.7 | 0.9998 |
| Dihydroneopterin triphosphate | 85.0 ± 2.4 | 91.6 ± 2.9 | 94.6 ± 0.6 | 111.5 ± 3.1 | 94.7 ± 1.4 | 105.6 ± 0.9 | 0.9993 |
| Dihydroxyacetonephosphate | 108.6 ± 0.3 | 114.0 ± 2.6 | 93.0 ± 1.4 | 89.7 ± 3.2 | 98.3 ± 1.2 | 112.9 ± 1.4 | 0.9991 |
| Dimethylglycine | 102.2 ± 1.7 | 101.6 ± 2.8 | 99.9 ± 0.8 | 96.7 ± 2.7 | 94.8 ± 1.1 | 104.1 ± 3.0 | 0.9993 |
| Ectoine | 95.6 ± 1.2 | 92.0 ± 2.6 | 86.7 ± 2.0 | 98.9 ± 1.5 | 113.8 ± 1.8 | 100.2 ± 1.0 | 0.9994 |
| Erythrose-4-phosphate | 100.8 ± 2.3 | 90.8 ± 2.7 | 93.3 ± 1.6 | 97.8 ± 1.8 | 113.7 ± 2.0 | 91.4 ± 0.7 | 0.9993 |
| FAD | 108.4 ± 1.4 | 101.9 ± 1.3 | 96.4 ± 1.4 | 108.2 ± 1.4 | 108.5 ± 1.9 | 97.3 ± 2.2 | 0.9998 |
| Folic acid | 86.9 ± 0.8 | 91.1 ± 0.4 | 90.4 ± 1.8 | 92.9 ± 2.0 | 87.4 ± 3.0 | 103.9 ± 1.6 | 0.9995 |
| Fructose-6-phosphate | 114.5 ± 1.0 | 93.4 ± 2.8 | 92.5 ± 2.5 | 102.1 ± 1.0 | 103.0 ± 2.1 | 95.8 ± 2.7 | 0.9997 |
| Fumaric Acid | 87.5 ± 2.7 | 111.3 ± 1.3 | 106.5 ± 1.3 | 106.3 ± 3.1 | 97.2 ± 0.7 | 99.7 ± 2.2 | 0.9997 |
| Galactosamine | 85.4 ± 2.4 | 107.9 ± 3.0 | 95.9 ± 0.8 | 99.0 ± 2.3 | 91.5 ± 1.4 | 97.4 ± 0.4 | 0.9994 |
| Galactose-1-phosphate | 89.5 ± 2.4 | 108.8 ± 0.8 | 87.5 ± 1.6 | 87.4 ± 1.4 | 97.8 ± 1.8 | 89.6 ± 0.9 | 0.9995 |
| Galactose-6-Phosphate | 92.8 ± 2.2 | 92.5 ± 1.9 | 112.3 ± 0.6 | 94.6 ± 0.7 | 86.5 ± 1.0 | 114.0 ± 2.7 | 0.9993 |
| gamma-Glutamylcysteine | 101.9 ± 1.6 | 103.8 ± 1.5 | 114.8 ± 0.7 | 110.6 ± 2.7 | 108.5 ± 0.5 | 89.7 ± 0.3 | 0.9998 |
| Glucosamine | 112.6 ± 1.7 | 110.6 ± 1.6 | 99.1 ± 0.3 | 93.8 ± 2.3 | 111.9 ± 0.4 | 96.8 ± 2.2 | 0.9992 |
| Glucosamine-6-Phosphate | 113.5 ± 1.1 | 105.9 ± 1.0 | 94.6 ± 2.8 | 87.8 ± 3.1 | 108.7 ± 0.7 | 109.8 ± 1.7 | 0.9999 |
| Glucose | 107.5 ± 1.9 | 111.6 ± 1.9 | 92.8 ± 1.3 | 98.8 ± 0.6 | 98.2 ± 0.6 | 109.4 ± 3.2 | 0.9992 |
| Glucose-1-phosphate | 108.1 ± 0.9 | 109.7 ± 1.6 | 102.4 ± 1.2 | 89.0 ± 3.0 | 96.2 ± 1.5 | 111.7 ± 1.3 | 0.9993 |
| Glucose-6-Phosphate | 102.7 ± 2.9 | 86.3 ± 2.2 | 106.9 ± 1.3 | 86.1 ± 2.6 | 97.3 ± 2.8 | 113.2 ± 2.3 | 0.9998 |
| Glucuronic acid | 110.6 ± 2.7 | 90.2 ± 1.4 | 110.9 ± 2.8 | 92.4 ± 0.8 | 107.3 ± 1.3 | 103.1 ± 2.6 | 0.9993 |
| Glutamic acid | 106.4 ± 1.3 | 96.5 ± 0.8 | 108.9 ± 3.0 | 106.4 ± 2.3 | 102.9 ± 1.1 | 100.7 ± 1.5 | 0.9997 |
| Glutamine | 89.5 ± 0.6 | 105.0 ± 1.1 | 108.6 ± 2.9 | 92.6 ± 1.3 | 108.2 ± 1.9 | 96.1 ± 0.8 | 0.9994 |
| Glutathione | 95.2 ± 1.5 | 112.7 ± 0.3 | 85.7 ± 2.8 | 111.8 ± 1.6 | 98.3 ± 1.2 | 111.0 ± 1.5 | 0.9999 |
| Glutathionylspermine | 111.1 ± 1.2 | 87.1 ± 2.6 | 115.0 ± 0.8 | 86.1 ± 1.4 | 90.4 ± 0.4 | 85.6 ± 2.2 | 0.9995 |
| Glycerophosphocholine | 92.2 ± 3.0 | 93.0 ± 2.8 | 106.3 ± 0.4 | 104.7 ± 1.2 | 89.2 ± 2.5 | 93.5 ± 1.9 | 0.9997 |
| Glycine | 91.3 ± 0.4 | 89.0 ± 2.2 | 85.5 ± 2.7 | 109.8 ± 3.0 | 101.2 ± 0.5 | 88.0 ± 1.3 | 0.9992 |
| Glycolic acid | 95.9 ± 0.4 | 104.7 ± 1.1 | 109.9 ± 1.1 | 114.8 ± 1.7 | 99.5 ± 0.8 | 87.8 ± 2.6 | 0.9996 |
| Guanidinoacetic acid | 91.2 ± 1.8 | 108.6 ± 3.0 | 105.3 ± 0.4 | 102.5 ± 2.6 | 110.6 ± 2.0 | 97.2 ± 2.9 | 0.9998 |
| Guanidinobutanoic acid | 96.6 ± 2.0 | 108.1 ± 1.9 | 103.4 ± 2.7 | 110.5 ± 0.4 | 85.3 ± 0.7 | 101.9 ± 3.2 | 0.9996 |
| Guanine | 114.7 ± 1.9 | 114.0 ± 2.7 | 88.3 ± 2.8 | 105.2 ± 1.0 | 86.1 ± 2.5 | 99.7 ± 1.2 | 0.9998 |
| Guanosine | 102.2 ± 0.6 | 87.4 ± 1.6 | 105.1 ± 2.3 | 99.6 ± 1.2 | 92.0 ± 2.8 | 98.2 ± 0.7 | 0.9998 |
| Guanosine 5'-diphosphate | 96.2 ± 1.4 | 111.8 ± 1.9 | 88.6 ± 1.4 | 85.1 ± 1.7 | 89.4 ± 0.6 | 93.4 ± 2.3 | 0.9996 |
| Guanosine-5'-monophosphate | 88.0 ± 0.5 | 110.7 ± 1.3 | 91.2 ± 2.7 | 107.4 ± 2.5 | 88.8 ± 0.7 | 91.9 ± 0.7 | 0.9998 |
| Hercynine | 94.6 ± 2.2 | 85.3 ± 0.9 | 97.4 ± 1.2 | 105.2 ± 2.1 | 114.6 ± 2.2 | 91.3 ± 1.7 | 0.9994 |
| Hippuric acid | 104.5 ± 2.3 | 87.3 ± 2.5 | 111.5 ± 1.2 | 85.2 ± 2.0 | 104.4 ± 3.0 | 86.2 ± 3.0 | 0.9996 |
| Histamine | 111.3 ± 0.9 | 95.7 ± 0.4 | 89.8 ± 1.3 | 87.4 ± 0.4 | 113.8 ± 2.5 | 112.9 ± 1.6 | 0.9996 |
| Histidine | 103.6 ± 0.4 | 99.0 ± 0.5 | 88.6 ± 0.7 | 86.6 ± 1.1 | 99.9 ± 0.8 | 107.8 ± 2.2 | 0.9997 |
| Homoalanine | 92.2 ± 2.0 | 88.6 ± 2.3 | 103.2 ± 0.8 | 93.3 ± 1.7 | 104.8 ± 0.7 | 109.8 ± 1.9 | 0.9996 |
| Homoarginine | 112.0 ± 0.4 | 111.4 ± 3.1 | 96.6 ± 1.4 | 101.0 ± 2.5 | 97.2 ± 1.0 | 100.5 ± 1.1 | 0.9993 |
| Homocysteine | 85.6 ± 1.0 | 89.9 ± 2.4 | 109.1 ± 0.6 | 95.3 ± 2.6 | 104.4 ± 0.4 | 86.1 ± 2.4 | 0.9998 |
| Homogentisic acid | 111.4 ± 3.2 | 103.2 ± 2.3 | 110.1 ± 1.1 | 95.7 ± 0.3 | 93.8 ± 1.1 | 111.3 ± 0.8 | 0.9991 |
| Homoleucine | 106.5 ± 3.1 | 96.4 ± 1.7 | 106.2 ± 1.7 | 100.9 ± 1.1 | 85.6 ± 2.4 | 105.0 ± 1.9 | 0.9994 |
| Homovanillic acid | 86.3 ± 2.7 | 109.7 ± 1.7 | 88.1 ± 2.5 | 97.7 ± 0.5 | 97.8 ± 1.4 | 101.1 ± 2.2 | 0.9994 |
| Hydantoin-5-propionate | 114.6 ± 1.4 | 113.0 ± 1.7 | 113.3 ± 1.4 | 106.3 ± 3.0 | 111.0 ± 1.9 | 86.1 ± 2.3 | 0.9998 |
| Hypotaurine | 92.7 ± 3.1 | 111.2 ± 2.1 | 93.7 ± 1.9 | 107.2 ± 2.9 | 87.0 ± 2.0 | 100.6 ± 0.8 | 0.9991 |
| Hypoxanthine | 109.7 ± 1.8 | 107.4 ± 0.4 | 105.3 ± 1.4 | 100.1 ± 1.4 | 96.9 ± 1.6 | 106.1 ± 1.7 | 0.9993 |
| Indole-3-acetamide | 114.4 ± 2.9 | 108.3 ± 1.1 | 94.7 ± 2.7 | 107.0 ± 2.0 | 90.7 ± 2.7 | 99.3 ± 2.1 | 0.9995 |
| Inosine | 107.6 ± 1.6 | 95.6 ± 2.7 | 87.9 ± 3.1 | 105.3 ± 0.9 | 88.2 ± 1.8 | 114.3 ± 1.7 | 0.9997 |
| Inosine 5'-monophosphate | 106.5 ± 0.4 | 103.1 ± 1.1 | 98.2 ± 2.7 | 94.5 ± 1.9 | 98.8 ± 1.5 | 105.8 ± 2.1 | 0.9995 |
| Isocitric Acid | 97.0 ± 1.7 | 94.3 ± 0.5 | 87.3 ± 2.8 | 95.4 ± 0.6 | 114.4 ± 0.9 | 94.8 ± 0.6 | 0.9997 |
| Isoleucine | 85.5 ± 2.9 | 114.2 ± 1.4 | 93.6 ± 2.5 | 86.9 ± 2.5 | 88.9 ± 0.5 | 105.2 ± 1.7 | 0.9994 |
| Lactic Acid | 86.7 ± 2.8 | 86.2 ± 1.2 | 101.4 ± 0.6 | 96.6 ± 2.6 | 92.9 ± 2.1 | 86.4 ± 1.8 | 0.9992 |
| Leucine | 108.1 ± 1.7 | 90.5 ± 0.4 | 114.7 ± 1.8 | 94.5 ± 2.0 | 94.6 ± 2.0 | 111.0 ± 2.3 | 0.9994 |
| Levulinic acid | 104.0 ± 2.5 | 97.3 ± 2.8 | 94.5 ± 0.7 | 110.3 ± 2.7 | 112.9 ± 1.0 | 103.7 ± 2.5 | 0.9995 |
| Lysine | 113.9 ± 1.8 | 86.4 ± 3.0 | 109.4 ± 2.4 | 110.8 ± 2.4 | 88.8 ± 1.3 | 85.5 ± 1.3 | 0.9994 |
| Malic Acid | 104.2 ± 0.4 | 92.7 ± 2.3 | 91.5 ± 1.1 | 102.3 ± 2.1 | 92.5 ± 1.5 | 101.2 ± 2.5 | 0.9999 |
| Mannose-1 with Mannose-6-Phosphate | 104.4 ± 2.8 | 102.4 ± 2.2 | 114.2 ± 0.7 | 99.3 ± 2.9 | 97.5 ± 1.6 | 112.2 ± 2.5 | 0.9998 |
| Mesaconic acid | 99.5 ± 1.7 | 101.1 ± 3.1 | 111.3 ± 0.8 | 108.5 ± 0.6 | 106.7 ± 1.4 | 102.9 ± 3.1 | 0.9999 |
| Methionine | 104.6 ± 2.5 | 97.8 ± 2.7 | 100.1 ± 1.9 | 96.4 ± 0.9 | 114.5 ± 1.3 | 95.4 ± 1.6 | 0.9995 |
| Methionine sulfoxide | 105.7 ± 1.9 | 104.6 ± 1.4 | 114.1 ± 0.5 | 112.6 ± 2.2 | 97.6 ± 1.9 | 93.5 ± 2.2 | 0.9998 |
| Methylmalonic Acid | 93.5 ± 3.1 | 86.6 ± 2.5 | 114.9 ± 3.0 | 96.2 ± 1.7 | 99.5 ± 2.4 | 108.3 ± 1.6 | 0.9998 |
| N1-Acetylspermidine | 112.6 ± 1.6 | 95.1 ± 1.3 | 98.0 ± 2.2 | 97.0 ± 0.4 | 91.4 ± 0.9 | 101.5 ± 1.7 | 0.9998 |
| N1-Acetylspermine | 87.2 ± 0.8 | 99.5 ± 0.8 | 93.9 ± 2.5 | 99.6 ± 0.4 | 96.0 ± 2.2 | 110.0 ± 1.4 | 0.9995 |
| N8-Acetylspermidine | 94.9 ± 0.4 | 109.0 ± 2.1 | 96.4 ± 2.6 | 108.3 ± 0.6 | 89.6 ± 1.2 | 112.4 ± 1.8 | 0.9995 |
| N-Acetyl-Aspartyl-Glutamate | 97.0 ± 2.6 | 93.0 ± 1.9 | 98.2 ± 2.1 | 106.3 ± 2.6 | 111.3 ± 1.5 | 100.2 ± 0.6 | 0.9991 |
| N-Acetyl-Aspartyl-Glutamyl-Glutamate | 86.9 ± 1.1 | 103.9 ± 1.0 | 86.7 ± 0.9 | 99.3 ± 0.3 | 99.4 ± 2.3 | 106.3 ± 0.9 | 0.9996 |
| N-Acetyl-Galactosamine | 106.2 ± 2.3 | 107.5 ± 3.0 | 112.3 ± 0.4 | 99.6 ± 1.1 | 100.1 ± 1.4 | 94.0 ± 2.2 | 0.9998 |
| N-Acetyl-Leucine | 108.0 ± 2.7 | 101.9 ± 2.2 | 90.0 ± 1.7 | 89.4 ± 1.1 | 96.7 ± 2.3 | 98.1 ± 1.3 | 0.9993 |
| N-Acetyl-Methionine | 112.1 ± 2.1 | 86.8 ± 2.2 | 109.8 ± 3.0 | 112.3 ± 2.3 | 101.6 ± 3.0 | 93.4 ± 2.8 | 0.9998 |
| N-Acetylneuraminic acid | 109.2 ± 0.3 | 114.8 ± 0.7 | 90.4 ± 1.8 | 91.7 ± 0.4 | 101.1 ± 1.6 | 96.5 ± 2.4 | 0.9991 |
| N-Acetyl-Proline | 100.7 ± 3.0 | 106.8 ± 2.7 | 85.8 ± 3.0 | 95.3 ± 2.0 | 86.0 ± 0.8 | 94.7 ± 2.5 | 0.9997 |
| N-Acetyl-Putrescine | 110.5 ± 2.4 | 99.3 ± 0.8 | 91.8 ± 2.0 | 101.9 ± 1.0 | 114.8 ± 0.7 | 108.9 ± 2.0 | 0.9998 |
| N-Acetyl-Serine | 91.2 ± 0.3 | 94.7 ± 1.7 | 106.4 ± 1.3 | 104.7 ± 3.1 | 112.4 ± 2.7 | 93.8 ± 1.1 | 0.9993 |
| N-Acetyl-Serotonin | 91.2 ± 2.4 | 98.9 ± 2.3 | 110.9 ± 3.1 | 107.8 ± 2.2 | 113.7 ± 1.0 | 104.2 ± 1.9 | 0.9991 |
| N-Acyl-Homoserine-lactone | 101.9 ± 0.9 | 104.8 ± 1.6 | 93.6 ± 2.6 | 104.2 ± 1.8 | 102.2 ± 2.6 | 92.6 ± 1.5 | 0.9995 |
| NAD | 88.1 ± 1.6 | 114.9 ± 0.7 | 92.6 ± 2.0 | 105.3 ± 1.8 | 85.8 ± 2.0 | 88.5 ± 1.2 | 0.9998 |
| NADH | 89.6 ± 0.5 | 91.5 ± 1.1 | 115.0 ± 2.5 | 112.9 ± 1.1 | 101.7 ± 1.0 | 89.6 ± 2.0 | 0.9998 |
| NADP | 98.4 ± 1.6 | 98.4 ± 0.4 | 96.6 ± 2.7 | 96.1 ± 1.7 | 107.0 ± 2.7 | 90.0 ± 1.4 | 0.9997 |
| NADPH | 95.1 ± 0.9 | 90.0 ± 0.5 | 110.6 ± 1.9 | 101.6 ± 2.0 | 113.9 ± 2.9 | 108.8 ± 2.8 | 0.9996 |
| N-alpha-Acetyl-lysine | 109.8 ± 2.4 | 101.9 ± 2.0 | 105.6 ± 1.3 | 109.5 ± 0.7 | 110.7 ± 3.1 | 85.8 ± 2.2 | 0.9999 |
| N-alpha-Acetyl-Ornithine | 105.4 ± 2.0 | 86.5 ± 2.7 | 96.8 ± 2.1 | 89.2 ± 2.4 | 100.4 ± 0.8 | 107.7 ± 1.7 | 0.9997 |
| N-Carbamoyl-Proline | 98.5 ± 2.9 | 104.8 ± 2.4 | 111.0 ± 0.3 | 93.2 ± 1.9 | 86.1 ± 0.3 | 105.1 ± 0.5 | 0.9994 |
| N-Carbamoyl-Sarcosine | 90.3 ± 1.2 | 99.6 ± 2.7 | 92.9 ± 3.2 | 86.4 ± 2.1 | 110.2 ± 1.5 | 90.6 ± 2.7 | 0.9996 |
| N-Epsilon-Acetyl-lysine | 94.3 ± 2.3 | 107.4 ± 1.7 | 89.3 ± 2.5 | 99.2 ± 1.3 | 91.8 ± 0.7 | 110.0 ± 0.6 | 0.9993 |
| Nicotinamide | 94.7 ± 1.9 | 94.3 ± 0.4 | 85.1 ± 1.1 | 107.1 ± 1.5 | 105.7 ± 2.4 | 95.9 ± 0.5 | 0.9997 |
| N-Methyl-Aspartic Acid | 108.9 ± 0.8 | 109.8 ± 2.0 | 111.3 ± 3.1 | 104.7 ± 1.7 | 110.3 ± 3.0 | 97.6 ± 1.7 | 0.9998 |
| O-Phosphoethanolamine | 101.4 ± 1.1 | 102.3 ± 1.5 | 106.3 ± 1.7 | 110.6 ± 1.9 | 110.5 ± 2.2 | 90.4 ± 2.8 | 0.9997 |
| Ophthalmic acid | 98.9 ± 0.6 | 103.6 ± 3.1 | 107.5 ± 1.8 | 93.0 ± 1.9 | 108.8 ± 2.4 | 99.6 ± 2.2 | 0.9994 |
| Ornithine | 85.6 ± 0.9 | 93.3 ± 2.0 | 93.7 ± 2.5 | 109.6 ± 0.3 | 100.4 ± 1.7 | 102.3 ± 2.1 | 0.9999 |
| Orotidine 5'-monophosphate | 87.7 ± 0.6 | 92.9 ± 2.1 | 109.2 ± 0.3 | 93.6 ± 1.0 | 108.1 ± 3.1 | 99.3 ± 0.5 | 0.9993 |
| Oxaloacetate | 99.7 ± 0.5 | 103.3 ± 1.3 | 108.9 ± 0.5 | 89.2 ± 1.2 | 109.7 ± 1.1 | 94.6 ± 2.1 | 0.9997 |
| Pantothenic acid | 108.2 ± 1.7 | 87.3 ± 0.8 | 106.5 ± 3.0 | 104.2 ± 1.6 | 110.4 ± 2.6 | 102.6 ± 0.8 | 0.9995 |
| Phenylalanine | 111.8 ± 2.6 | 102.6 ± 1.4 | 105.6 ± 1.1 | 99.3 ± 2.2 | 98.1 ± 1.2 | 90.5 ± 2.4 | 0.9994 |
| Phenylpyruvate | 89.7 ± 1.3 | 90.6 ± 2.5 | 93.4 ± 3.0 | 109.5 ± 1.0 | 106.8 ± 2.5 | 109.1 ± 0.8 | 0.9997 |
| Phosphodimethylethanolamine | 104.6 ± 2.7 | 96.0 ± 2.8 | 107.6 ± 1.4 | 92.5 ± 1.5 | 100.4 ± 0.7 | 87.7 ± 1.9 | 0.9997 |
| Phosphoenolpyruvate | 86.8 ± 2.0 | 94.7 ± 2.9 | 97.2 ± 3.0 | 104.5 ± 2.9 | 104.4 ± 2.7 | 86.5 ± 0.6 | 0.9996 |
| Phosphorylcholine | 96.3 ± 1.8 | 114.3 ± 2.1 | 89.4 ± 0.9 | 114.8 ± 0.5 | 110.1 ± 0.9 | 85.9 ± 2.4 | 0.9997 |
| Pipecolic acid | 103.6 ± 2.3 | 86.6 ± 2.4 | 109.8 ± 2.3 | 105.0 ± 2.8 | 109.8 ± 0.7 | 100.1 ± 0.8 | 0.9996 |
| Proline | 91.0 ± 1.6 | 90.4 ± 1.4 | 109.8 ± 2.2 | 93.8 ± 1.7 | 92.0 ± 0.4 | 105.8 ± 2.4 | 0.9997 |
| Putrescine | 113.9 ± 1.8 | 111.2 ± 2.6 | 89.3 ± 1.7 | 107.5 ± 1.9 | 113.3 ± 2.4 | 90.3 ± 0.9 | 0.9998 |
| Pyridoxal | 92.8 ± 3.0 | 102.9 ± 2.5 | 104.5 ± 2.0 | 86.5 ± 1.1 | 106.7 ± 0.8 | 96.2 ± 2.5 | 0.9998 |
| Pyridoxamine | 89.9 ± 0.9 | 113.3 ± 0.6 | 85.5 ± 0.7 | 91.5 ± 1.1 | 102.3 ± 0.4 | 107.6 ± 2.6 | 0.9994 |
| Pyridoxine | 96.6 ± 3.0 | 98.1 ± 0.7 | 98.1 ± 3.1 | 88.1 ± 1.4 | 94.2 ± 1.9 | 88.8 ± 0.6 | 0.9998 |
| Pyroglutamic acid | 107.4 ± 3.2 | 91.3 ± 3.2 | 97.7 ± 1.2 | 96.8 ± 0.8 | 104.7 ± 1.0 | 102.8 ± 0.4 | 0.9992 |
| Pyrrolysine | 96.7 ± 3.1 | 110.2 ± 1.4 | 88.0 ± 1.1 | 114.3 ± 1.3 | 85.0 ± 2.3 | 95.3 ± 1.2 | 0.9997 |
| Pyruvic Acid | 97.1 ± 2.1 | 112.7 ± 1.4 | 106.0 ± 1.8 | 93.6 ± 3.0 | 93.8 ± 0.9 | 86.9 ± 2.6 | 0.9992 |
| Riboflavin | 98.8 ± 2.6 | 92.3 ± 0.8 | 97.1 ± 2.9 | 86.0 ± 1.5 | 91.0 ± 2.8 | 98.1 ± 1.0 | 0.9997 |
| Ribose-5-Phosphate | 91.4 ± 3.0 | 109.9 ± 0.6 | 98.8 ± 1.1 | 99.6 ± 1.8 | 112.0 ± 3.1 | 93.1 ± 1.6 | 0.9998 |
| Ribulose-1-Phosphate | 109.1 ± 2.8 | 111.1 ± 2.8 | 104.5 ± 1.7 | 95.1 ± 2.3 | 92.3 ± 1.7 | 96.6 ± 3.1 | 0.9992 |
| Ribulose-5-Phosphate | 106.9 ± 1.1 | 86.7 ± 0.5 | 88.5 ± 2.9 | 95.3 ± 1.2 | 104.6 ± 2.7 | 104.5 ± 1.1 | 0.9996 |
| Saccharopine | 88.2 ± 0.5 | 100.7 ± 3.0 | 104.0 ± 1.5 | 88.1 ± 1.5 | 112.6 ± 1.1 | 110.1 ± 1.8 | 0.9992 |
| S-Adenosylhomocysteine | 103.8 ± 1.5 | 96.9 ± 1.7 | 92.5 ± 2.1 | 108.7 ± 0.7 | 113.4 ± 0.3 | 86.6 ± 0.9 | 0.9997 |
| S-Adenosylmethioninamine | 100.5 ± 2.8 | 104.4 ± 2.5 | 103.8 ± 0.9 | 96.3 ± 1.1 | 92.7 ± 1.3 | 106.2 ± 1.5 | 0.9992 |
| S-Adenosylmethionine | 90.1 ± 1.8 | 89.1 ± 2.3 | 91.0 ± 1.5 | 110.7 ± 0.9 | 96.4 ± 0.3 | 90.1 ± 2.9 | 0.9997 |
| Sarcosine | 96.3 ± 0.5 | 94.4 ± 2.5 | 111.9 ± 2.1 | 85.4 ± 2.8 | 107.7 ± 0.9 | 106.4 ± 0.5 | 0.9992 |
| Sedoheptulose-7-phosphate | 111.0 ± 1.5 | 98.9 ± 2.9 | 90.2 ± 2.5 | 92.7 ± 2.2 | 112.7 ± 2.5 | 101.6 ± 0.4 | 0.9999 |
| Selenohomocysteine | 109.3 ± 1.9 | 94.0 ± 1.5 | 112.6 ± 2.7 | 99.6 ± 1.8 | 108.9 ± 2.0 | 101.8 ± 2.1 | 0.9995 |
| Selenomethionine | 88.4 ± 3.0 | 101.5 ± 2.3 | 92.1 ± 2.0 | 89.0 ± 2.1 | 91.2 ± 1.9 | 110.6 ± 1.0 | 0.9995 |
| Serine | 110.3 ± 2.2 | 97.4 ± 2.0 | 107.8 ± 1.2 | 94.0 ± 2.3 | 111.6 ± 2.0 | 93.2 ± 0.5 | 0.9992 |
| S-Methyl-Thioinosine | 86.6 ± 2.5 | 111.8 ± 2.7 | 103.9 ± 2.7 | 94.5 ± 0.6 | 107.3 ± 1.4 | 90.5 ± 2.0 | 0.9996 |
| Spermidine | 102.8 ± 0.4 | 104.0 ± 2.3 | 104.2 ± 1.4 | 86.3 ± 2.5 | 97.3 ± 2.7 | 90.6 ± 3.0 | 0.9998 |
| Spermine | 113.0 ± 2.2 | 110.1 ± 1.2 | 94.9 ± 0.5 | 96.0 ± 2.9 | 100.6 ± 1.5 | 113.8 ± 1.8 | 0.9996 |
| Succinate | 102.5 ± 0.5 | 101.1 ± 1.5 | 89.3 ± 0.5 | 114.7 ± 1.0 | 96.3 ± 0.9 | 109.7 ± 1.8 | 0.9998 |
| Sucrose | 88.5 ± 0.9 | 93.7 ± 3.2 | 95.4 ± 1.0 | 91.8 ± 1.4 | 87.4 ± 2.6 | 98.5 ± 1.1 | 0.9996 |
| Symmetric dimethylarginine | 93.0 ± 2.8 | 99.8 ± 0.6 | 105.6 ± 0.4 | 108.0 ± 1.7 | 93.5 ± 0.3 | 102.8 ± 2.6 | 0.9998 |
| Taurine | 100.1 ± 1.8 | 100.7 ± 1.3 | 91.8 ± 2.8 | 99.5 ± 1.2 | 93.1 ± 2.7 | 92.4 ± 2.4 | 0.9995 |
| Tetrahydroneopterin | 99.6 ± 1.2 | 113.8 ± 1.1 | 102.8 ± 2.1 | 86.0 ± 1.3 | 106.7 ± 2.3 | 97.7 ± 1.5 | 0.9993 |
| Thiamine | 109.4 ± 1.2 | 86.3 ± 0.4 | 111.0 ± 0.4 | 103.5 ± 0.7 | 113.7 ± 2.9 | 108.6 ± 0.4 | 0.9994 |
| Thiamine monophosphate | 86.3 ± 1.4 | 108.2 ± 2.7 | 104.2 ± 1.9 | 89.5 ± 2.0 | 101.9 ± 3.2 | 114.7 ± 1.2 | 0.9994 |
| Threonic acid | 114.8 ± 2.7 | 104.5 ± 0.8 | 101.6 ± 1.2 | 114.8 ± 2.9 | 91.7 ± 1.3 | 114.7 ± 1.6 | 0.9993 |
| Threonine | 103.9 ± 1.7 | 105.4 ± 0.6 | 107.0 ± 2.1 | 109.8 ± 0.7 | 112.5 ± 0.4 | 87.6 ± 2.1 | 0.9992 |
| Thymidine | 100.4 ± 2.9 | 107.6 ± 1.6 | 113.6 ± 1.2 | 99.7 ± 2.0 | 97.8 ± 1.4 | 88.3 ± 3.0 | 0.9997 |
| Thymidine Monophosphate | 108.1 ± 2.1 | 113.7 ± 1.8 | 101.5 ± 1.0 | 93.7 ± 0.6 | 110.9 ± 2.4 | 99.8 ± 2.2 | 0.9995 |
| Thymine | 91.6 ± 2.1 | 87.5 ± 1.8 | 89.3 ± 2.3 | 112.2 ± 1.5 | 98.7 ± 2.2 | 109.3 ± 0.4 | 0.9997 |
| Trigonelleine | 107.7 ± 0.4 | 88.6 ± 0.6 | 91.1 ± 2.7 | 108.9 ± 2.9 | 97.2 ± 3.2 | 105.4 ± 1.4 | 0.9991 |
| Trimethylamine N-oxide | 85.8 ± 2.1 | 94.2 ± 1.8 | 99.4 ± 2.2 | 109.2 ± 0.8 | 111.7 ± 0.7 | 112.1 ± 2.3 | 0.9993 |
| Trimethyllysine | 90.1 ± 1.9 | 113.9 ± 0.8 | 110.1 ± 1.4 | 106.3 ± 2.2 | 90.1 ± 2.9 | 88.7 ± 1.5 | 0.9995 |
| Tryptophan | 109.4 ± 0.9 | 90.1 ± 2.6 | 91.4 ± 2.2 | 107.1 ± 1.3 | 93.6 ± 0.9 | 109.8 ± 1.0 | 0.9991 |
| Tyramine | 103.4 ± 0.9 | 98.2 ± 0.5 | 90.3 ± 2.2 | 93.1 ± 0.4 | 96.0 ± 0.8 | 99.8 ± 1.8 | 0.9996 |
| Tyrosine | 102.5 ± 3.2 | 93.9 ± 0.4 | 111.6 ± 0.5 | 87.0 ± 1.1 | 102.6 ± 1.5 | 89.1 ± 1.9 | 0.9996 |
| Uracil | 110.2 ± 2.3 | 92.3 ± 2.0 | 106.1 ± 0.6 | 113.1 ± 2.8 | 101.6 ± 2.4 | 96.4 ± 2.7 | 0.9992 |
| Uridine 5'-diphosphate | 93.7 ± 1.0 | 93.2 ± 1.2 | 111.9 ± 1.7 | 104.3 ± 2.6 | 105.6 ± 0.8 | 85.7 ± 1.4 | 0.9998 |
| Uridine Monophosphate | 93.8 ± 0.8 | 105.1 ± 1.9 | 107.2 ± 0.6 | 88.3 ± 0.5 | 111.9 ± 1.6 | 109.6 ± 0.6 | 0.9992 |
| Uridine-5-Diphosphogalactose | 86.7 ± 2.1 | 101.3 ± 1.7 | 101.4 ± 1.3 | 114.1 ± 1.3 | 108.1 ± 2.6 | 114.1 ± 1.2 | 0.9993 |
| Urocanic acid | 100.5 ± 2.8 | 93.8 ± 2.0 | 101.4 ± 2.7 | 90.2 ± 1.7 | 92.1 ± 2.1 | 94.2 ± 3.2 | 0.9997 |
| UTP | 112.0 ± 0.9 | 102.2 ± 0.4 | 89.0 ± 2.7 | 93.0 ± 1.4 | 101.7 ± 1.1 | 86.0 ± 0.8 | 0.9997 |
| Valine | 105.6 ± 0.7 | 102.3 ± 3.1 | 98.9 ± 2.3 | 91.0 ± 1.8 | 97.2 ± 1.2 | 107.3 ± 2.8 | 0.9991 |
| XMP | 111.9 ± 1.5 | 91.2 ± 2.9 | 110.2 ± 0.9 | 106.9 ± 2.6 | 104.3 ± 2.6 | 88.0 ± 2.8 | 0.9997 |
| Xylulose-5-Phosphate | 85.4 ± 0.8 | 88.8 ± 2.9 | 94.1 ± 2.5 | 105.4 ± 2.1 | 98.1 ± 3.1 | 92.8 ± 1.4 | 0.9999 |

### **Table S10** | Calibration values for final method conditions (r^2^-value) and percent found of nominal calibrator concentration (Cell extract as matrix).

| Calibrator concentration accuracy  [%] | 10 ng/mL | 25 ng/mL | 50 ng/mL | 100 ng/mL | 500 ng/mL | 1000 ng/mL | r^2^-Value  matrix-matched calibration |
| --- | --- | --- | --- | --- | --- | --- | --- |
| 1-Methylguanine | 91.2 ± 3.2 | 99.3 ± 0.3 | 110.0 ± 0.8 | 107.5 ± 1.9 | 114.9 ± 3.2 | 112.1 ± 2.1 | 0.9995 |
| 1-Methylhistidine | 86.0 ± 1.0 | 114.8 ± 2.8 | 85.5 ± 0.5 | 107.9 ± 0.7 | 112.7 ± 2.7 | 105.5 ± 2.4 | 0.9995 |
| 1-Methylnicotinamide | 96.5 ± 2.2 | 111.6 ± 0.5 | 86.8 ± 2.7 | 88.2 ± 3.2 | 94.3 ± 3.0 | 100.2 ± 0.5 | 0.9997 |
| 2-Aminoadipic Acid | 100.4 ± 2.4 | 96.3 ± 3.2 | 106.3 ± 2.3 | 105.3 ± 1.9 | 95.3 ± 0.8 | 97.1 ± 3.0 | 0.9995 |
| 2-Deoxyadenosine | 99.8 ± 1.5 | 105.1 ± 0.9 | 88.3 ± 0.7 | 100.3 ± 1.6 | 112.0 ± 0.7 | 106.7 ± 3.1 | 0.9997 |
| 2'-Deoxyadenosine 5'-monophosphate | 98.9 ± 2.9 | 105.8 ± 2.6 | 107.0 ± 1.7 | 94.1 ± 2.7 | 95.4 ± 0.4 | 97.3 ± 2.7 | 0.9998 |
| 2'-deoxyadenosine 5'-triphosphate | 111.7 ± 3.0 | 93.1 ± 1.0 | 87.4 ± 2.7 | 112.1 ± 2.2 | 102.8 ± 1.9 | 105.5 ± 2.0 | 0.9998 |
| 2'-Deoxycytidine-5'-monophosphate | 102.6 ± 3.2 | 106.8 ± 1.3 | 105.2 ± 1.5 | 111.6 ± 3.2 | 108.4 ± 3.1 | 105.7 ± 1.8 | 0.9993 |
| 2-Deoxyguanosine | 92.6 ± 3.1 | 108.2 ± 2.8 | 87.8 ± 0.4 | 87.3 ± 2.1 | 108.2 ± 0.5 | 111.7 ± 3.0 | 0.9993 |
| 2-hydroxybutyric acid | 85.3 ± 0.8 | 103.8 ± 0.7 | 110.3 ± 2.1 | 95.5 ± 2.3 | 86.9 ± 1.9 | 102.6 ± 0.4 | 0.9995 |
| 2-hydroxyglutaric acid | 113.0 ± 0.9 | 86.3 ± 1.7 | 113.8 ± 1.4 | 109.2 ± 0.8 | 89.0 ± 1.2 | 113.7 ± 2.3 | 0.9998 |
| 2-Hydroxyisocaproic acid | 103.3 ± 0.6 | 109.6 ± 2.9 | 101.7 ± 1.9 | 109.7 ± 2.6 | 103.5 ± 0.5 | 91.6 ± 1.9 | 0.9992 |
| 2-ketobutyric acid | 105.3 ± 1.4 | 89.9 ± 0.5 | 92.3 ± 2.0 | 91.3 ± 2.3 | 114.2 ± 2.6 | 113.7 ± 0.4 | 0.9998 |
| 2-oxoadipic acid | 110.4 ± 0.3 | 114.4 ± 0.4 | 102.3 ± 0.5 | 111.3 ± 2.3 | 102.1 ± 1.1 | 103.1 ± 2.7 | 0.9992 |
| 2-Phenylglycine | 96.0 ± 0.8 | 88.0 ± 1.8 | 111.1 ± 1.9 | 86.2 ± 1.5 | 101.5 ± 2.6 | 96.1 ± 2.4 | 0.9993 |
| 2-phosphoglycerate | 102.2 ± 2.3 | 92.1 ± 3.0 | 86.4 ± 2.3 | 87.4 ± 1.6 | 104.4 ± 0.7 | 89.0 ± 3.0 | 0.9992 |
| 3-hydroxy-3-methyl-2-oxobutanoate | 94.9 ± 0.7 | 98.6 ± 3.0 | 89.6 ± 2.1 | 97.1 ± 1.6 | 109.0 ± 2.8 | 90.4 ± 2.4 | 0.9992 |
| 3-hydroxybutyric acid | 105.3 ± 0.8 | 108.8 ± 1.0 | 96.2 ± 1.7 | 93.0 ± 1.0 | 111.3 ± 0.3 | 104.9 ± 1.6 | 0.9993 |
| 3-Methylhistidine | 88.0 ± 0.6 | 100.5 ± 1.5 | 106.6 ± 0.8 | 111.5 ± 3.1 | 102.3 ± 2.9 | 96.6 ± 0.5 | 0.9996 |
| 3-Phosphoglyceraldehyde | 95.1 ± 1.8 | 114.3 ± 1.6 | 103.4 ± 2.7 | 96.6 ± 1.0 | 86.1 ± 2.5 | 99.2 ± 2.2 | 0.9994 |
| 3-phosphoglycerate | 102.8 ± 0.5 | 114.3 ± 1.4 | 86.2 ± 2.4 | 100.9 ± 2.4 | 106.9 ± 1.6 | 96.9 ± 0.4 | 0.9996 |
| 4-amino-butyric acid | 96.6 ± 2.5 | 100.7 ± 1.8 | 110.2 ± 1.5 | 109.9 ± 1.5 | 103.7 ± 1.1 | 112.7 ± 2.0 | 0.9993 |
| 4-aspartyl-phosphate | 100.7 ± 0.8 | 93.7 ± 2.5 | 112.0 ± 1.8 | 92.0 ± 2.4 | 95.9 ± 0.3 | 101.0 ± 2.2 | 0.9995 |
| 4-Hydroxybenzoic acid | 96.6 ± 1.4 | 108.2 ± 2.2 | 93.4 ± 2.4 | 100.6 ± 3.2 | 93.6 ± 0.3 | 95.3 ± 2.3 | 0.9993 |
| 4-Hydroxyphenylacetylglycine | 86.1 ± 2.5 | 90.5 ± 2.7 | 92.4 ± 2.9 | 97.5 ± 2.0 | 114.8 ± 2.8 | 104.9 ± 1.7 | 0.9991 |
| 4-methyl-2-oxopentanoate | 105.0 ± 2.2 | 99.6 ± 2.3 | 93.5 ± 0.9 | 109.5 ± 2.8 | 90.0 ± 1.3 | 103.0 ± 2.8 | 0.9996 |
| 4-Oxoglutaramate | 103.4 ± 1.9 | 95.8 ± 2.1 | 97.0 ± 3.1 | 88.4 ± 1.7 | 104.5 ± 1.7 | 95.4 ± 0.7 | 0.9992 |
| 5-Hydroxyindolepyruvate | 100.1 ± 2.8 | 95.5 ± 0.5 | 101.4 ± 1.2 | 89.1 ± 0.9 | 110.6 ± 1.8 | 86.0 ± 0.6 | 0.9994 |
| 5-Hydroxylysine | 100.1 ± 1.1 | 85.3 ± 0.6 | 94.8 ± 1.9 | 103.4 ± 1.3 | 103.3 ± 2.3 | 99.8 ± 2.1 | 0.9997 |
| 5-Hydroxy-Tryptophan | 97.5 ± 0.5 | 109.5 ± 1.9 | 101.7 ± 3.1 | 85.9 ± 2.6 | 114.1 ± 1.6 | 108.4 ± 0.9 | 0.9997 |
| 5-Phosphoribosylamine | 104.9 ± 1.1 | 96.2 ± 0.7 | 101.2 ± 1.3 | 96.5 ± 0.8 | 110.0 ± 2.4 | 89.4 ± 1.8 | 0.9997 |
| 6-phosphogluconate | 92.0 ± 1.7 | 87.5 ± 1.5 | 108.9 ± 2.4 | 111.4 ± 0.9 | 99.9 ± 0.6 | 92.7 ± 2.8 | 0.9996 |
| Acetylcarnitine | 91.8 ± 1.7 | 108.6 ± 2.2 | 114.9 ± 1.9 | 94.8 ± 2.6 | 97.5 ± 2.3 | 94.7 ± 0.9 | 0.9996 |
| Aconitic Acid | 94.9 ± 3.1 | 110.1 ± 2.1 | 109.7 ± 2.1 | 89.1 ± 1.7 | 108.3 ± 0.4 | 93.8 ± 3.1 | 0.9998 |
| Adenine | 96.8 ± 0.6 | 88.2 ± 0.8 | 96.8 ± 2.4 | 104.3 ± 1.1 | 110.2 ± 2.2 | 87.3 ± 2.8 | 0.9993 |
| Adenosine | 89.8 ± 1.3 | 93.2 ± 0.5 | 101.6 ± 1.3 | 88.5 ± 2.4 | 86.5 ± 0.8 | 101.0 ± 1.6 | 0.9991 |
| Adenosine 3',5'-cyclic monophosphate | 92.9 ± 0.8 | 100.8 ± 1.4 | 112.0 ± 2.8 | 86.3 ± 1.9 | 97.9 ± 2.9 | 106.5 ± 2.1 | 0.9994 |
| Adenosine Monophosphate | 106.6 ± 1.4 | 109.0 ± 0.8 | 88.4 ± 2.1 | 85.2 ± 2.2 | 108.1 ± 1.5 | 98.1 ± 2.7 | 0.9997 |
| Adenosine-5'-diphosphate | 109.4 ± 1.8 | 104.7 ± 0.9 | 107.5 ± 1.4 | 85.6 ± 3.1 | 104.8 ± 1.5 | 114.0 ± 3.1 | 0.9997 |
| Adenosine-5'-triphosphate | 95.6 ± 1.0 | 89.3 ± 2.5 | 86.1 ± 1.5 | 94.3 ± 2.2 | 104.4 ± 2.9 | 101.5 ± 1.6 | 0.9995 |
| Adenylosuccinate | 87.4 ± 1.8 | 98.3 ± 0.5 | 87.5 ± 2.6 | 102.1 ± 3.2 | 110.4 ± 1.8 | 88.1 ± 1.7 | 0.9999 |
| Alanine | 110.2 ± 2.7 | 94.1 ± 2.6 | 94.6 ± 1.1 | 93.8 ± 2.3 | 85.8 ± 1.8 | 87.3 ± 0.7 | 0.9995 |
| alpha-ketoglutaric acid | 92.9 ± 2.3 | 103.7 ± 2.0 | 102.2 ± 2.9 | 88.3 ± 2.6 | 108.6 ± 0.6 | 109.4 ± 2.1 | 0.9992 |
| Anserine | 99.8 ± 2.3 | 114.1 ± 2.0 | 107.1 ± 2.9 | 91.8 ± 3.1 | 101.7 ± 2.3 | 89.0 ± 2.4 | 0.9992 |
| Arginine | 87.5 ± 2.9 | 114.1 ± 1.4 | 113.5 ± 1.9 | 86.4 ± 0.6 | 103.9 ± 2.1 | 104.2 ± 3.0 | 0.9992 |
| Argininosuccinic Acid | 87.2 ± 2.5 | 88.3 ± 1.1 | 100.3 ± 0.8 | 85.7 ± 2.5 | 89.4 ± 1.4 | 96.9 ± 3.1 | 0.9993 |
| Asparagine | 113.6 ± 1.9 | 100.4 ± 1.3 | 110.3 ± 2.5 | 106.8 ± 3.0 | 97.8 ± 3.0 | 111.2 ± 3.0 | 0.9991 |
| Aspartic acid | 97.4 ± 3.1 | 86.8 ± 1.5 | 107.5 ± 2.3 | 105.4 ± 0.7 | 95.8 ± 0.6 | 105.5 ± 0.5 | 0.9998 |
| Asymmetric dimethylarginine | 107.7 ± 2.8 | 98.5 ± 2.2 | 103.9 ± 2.1 | 107.7 ± 2.9 | 112.5 ± 2.8 | 101.6 ± 1.8 | 0.9993 |
| Azelaic acid | 112.7 ± 3.2 | 91.0 ± 0.6 | 109.9 ± 1.3 | 89.1 ± 1.2 | 103.4 ± 2.2 | 98.4 ± 2.9 | 0.9995 |
| beta-Alanine | 108.5 ± 0.9 | 106.6 ± 1.1 | 114.3 ± 1.7 | 90.5 ± 1.6 | 112.1 ± 2.6 | 112.8 ± 3.0 | 0.9998 |
| Betaine | 106.5 ± 0.6 | 106.2 ± 2.3 | 102.9 ± 1.0 | 108.2 ± 2.4 | 90.1 ± 2.3 | 96.9 ± 1.8 | 0.9997 |
| Biopterin | 114.7 ± 2.2 | 92.2 ± 2.0 | 95.6 ± 1.3 | 104.5 ± 1.0 | 111.1 ± 0.5 | 108.9 ± 2.3 | 0.9994 |
| Biotin | 88.6 ± 2.2 | 114.8 ± 1.0 | 104.8 ± 0.8 | 88.9 ± 0.9 | 102.0 ± 3.0 | 114.2 ± 2.0 | 0.9991 |
| Carnitine | 111.4 ± 0.3 | 94.0 ± 2.4 | 104.3 ± 2.8 | 109.9 ± 1.0 | 90.5 ± 1.1 | 106.7 ± 1.1 | 0.9997 |
| Carnosine | 92.7 ± 0.4 | 102.2 ± 2.4 | 112.9 ± 1.7 | 107.3 ± 2.3 | 88.1 ± 1.2 | 90.6 ± 2.6 | 0.9999 |
| CDP-choline | 106.6 ± 1.4 | 105.3 ± 2.3 | 88.7 ± 2.3 | 94.9 ± 0.6 | 111.1 ± 2.0 | 104.4 ± 0.4 | 0.9993 |
| cGMP | 109.3 ± 1.2 | 89.1 ± 0.3 | 108.2 ± 2.4 | 92.8 ± 2.2 | 98.5 ± 0.8 | 93.1 ± 1.5 | 0.9994 |
| Choline | 85.6 ± 1.4 | 112.5 ± 0.6 | 97.8 ± 2.4 | 114.3 ± 1.3 | 97.1 ± 1.8 | 98.4 ± 3.2 | 0.9991 |
| Cinnamic acid | 114.2 ± 0.3 | 113.6 ± 2.2 | 99.0 ± 1.1 | 97.5 ± 1.5 | 107.9 ± 0.7 | 93.8 ± 1.1 | 0.9996 |
| Citraconic acid | 93.7 ± 0.3 | 108.5 ± 3.0 | 108.8 ± 0.5 | 89.5 ± 2.6 | 91.6 ± 1.7 | 114.1 ± 1.2 | 0.9998 |
| Citramalic acid | 87.1 ± 2.8 | 88.0 ± 1.5 | 101.1 ± 0.6 | 90.2 ± 1.3 | 105.9 ± 1.3 | 88.1 ± 2.0 | 0.9998 |
| Citric Acid | 104.2 ± 2.1 | 85.6 ± 2.5 | 110.7 ± 1.0 | 111.2 ± 0.7 | 87.3 ± 1.2 | 96.9 ± 2.7 | 0.9998 |
| Citrulline | 102.6 ± 1.5 | 91.0 ± 0.6 | 93.4 ± 1.1 | 98.9 ± 1.1 | 113.4 ± 2.1 | 104.2 ± 1.2 | 0.9997 |
| Creatine | 109.0 ± 0.8 | 87.1 ± 2.5 | 103.7 ± 1.4 | 106.4 ± 0.8 | 86.3 ± 1.4 | 107.3 ± 1.8 | 0.9994 |
| Creatinine | 85.9 ± 2.6 | 113.0 ± 0.6 | 114.3 ± 0.6 | 87.5 ± 1.1 | 94.2 ± 0.9 | 109.8 ± 1.3 | 0.9999 |
| Cystathionine | 100.9 ± 0.9 | 96.9 ± 2.5 | 89.3 ± 3.0 | 98.6 ± 1.9 | 100.4 ± 2.0 | 103.1 ± 1.7 | 0.9994 |
| Cysteamine | 89.1 ± 0.9 | 91.3 ± 1.5 | 91.3 ± 0.7 | 114.4 ± 3.0 | 111.5 ± 2.7 | 108.0 ± 2.5 | 0.9993 |
| Cysteine | 92.5 ± 1.1 | 107.6 ± 0.6 | 85.0 ± 2.2 | 105.8 ± 2.7 | 85.3 ± 2.8 | 89.0 ± 1.3 | 0.9993 |
| Cysteinylglycine | 93.8 ± 2.5 | 110.4 ± 1.8 | 94.4 ± 2.2 | 101.6 ± 2.4 | 89.9 ± 1.0 | 114.9 ± 1.5 | 0.9996 |
| Cystine | 113.9 ± 1.3 | 97.1 ± 2.3 | 86.7 ± 1.9 | 101.2 ± 3.1 | 113.8 ± 1.7 | 88.8 ± 3.0 | 0.9996 |
| Cytidine | 98.9 ± 2.1 | 98.7 ± 2.6 | 89.7 ± 2.7 | 112.9 ± 1.1 | 101.0 ± 0.3 | 114.9 ± 1.8 | 0.9997 |
| Cytidine 3',5'-cyclic monophosphate | 95.9 ± 2.0 | 93.6 ± 2.5 | 108.9 ± 0.6 | 108.4 ± 2.0 | 100.5 ± 1.1 | 113.5 ± 1.1 | 0.9994 |
| Cytidine 5'-diphosphate | 92.9 ± 1.1 | 87.6 ± 0.7 | 94.9 ± 2.5 | 104.7 ± 1.7 | 87.8 ± 1.1 | 106.5 ± 2.8 | 0.9996 |
| Cytidine 5'-triphosphate | 92.1 ± 2.4 | 100.0 ± 0.9 | 101.8 ± 1.2 | 105.6 ± 1.9 | 113.1 ± 1.1 | 91.0 ± 2.1 | 0.9998 |
| Cytidine monophosphate | 106.6 ± 0.3 | 95.7 ± 2.4 | 106.7 ± 0.4 | 114.7 ± 0.5 | 108.8 ± 2.7 | 101.9 ± 1.9 | 0.9992 |
| Cytosine | 86.5 ± 2.2 | 90.5 ± 1.9 | 114.8 ± 2.4 | 114.7 ± 2.1 | 98.4 ± 1.1 | 92.5 ± 1.3 | 0.9998 |
| dADP | 97.3 ± 2.8 | 101.6 ± 1.1 | 104.4 ± 3.0 | 85.4 ± 3.2 | 90.1 ± 1.9 | 91.7 ± 1.8 | 0.9997 |
| Diethanolamine | 100.1 ± 1.8 | 111.2 ± 1.5 | 89.9 ± 0.5 | 92.8 ± 2.1 | 109.3 ± 3.1 | 92.6 ± 0.4 | 0.9993 |
| Dihydrobiopterin | 112.7 ± 0.8 | 97.9 ± 1.9 | 110.9 ± 3.0 | 90.8 ± 2.7 | 104.1 ± 1.9 | 107.8 ± 1.2 | 0.9994 |
| Dihydroneopterin triphosphate | 85.0 ± 0.8 | 97.1 ± 2.8 | 110.2 ± 3.0 | 107.9 ± 1.0 | 105.4 ± 2.7 | 100.4 ± 0.8 | 0.9995 |
| Dihydroxyacetonephosphate | 112.0 ± 3.1 | 88.6 ± 2.3 | 106.9 ± 2.0 | 91.1 ± 0.9 | 96.1 ± 2.4 | 109.5 ± 0.8 | 0.9992 |
| Dimethylglycine | 94.6 ± 3.0 | 91.1 ± 2.4 | 106.3 ± 2.2 | 86.1 ± 1.6 | 112.7 ± 1.4 | 103.6 ± 1.1 | 0.9998 |
| Ectoine | 97.4 ± 2.1 | 92.9 ± 1.1 | 98.4 ± 3.1 | 97.8 ± 2.3 | 93.1 ± 1.2 | 94.6 ± 1.9 | 0.9994 |
| Erythrose-4-phosphate | 101.6 ± 0.5 | 111.0 ± 0.4 | 98.7 ± 2.8 | 92.3 ± 0.8 | 96.3 ± 0.9 | 101.9 ± 2.6 | 0.9998 |
| FAD | 107.8 ± 0.6 | 100.4 ± 2.1 | 85.4 ± 1.4 | 91.8 ± 1.0 | 91.2 ± 2.2 | 99.9 ± 0.7 | 0.9998 |
| Folic acid | 92.8 ± 0.5 | 108.6 ± 3.0 | 102.0 ± 3.0 | 97.0 ± 2.6 | 111.3 ± 1.9 | 100.9 ± 2.5 | 0.9991 |
| Fructose-6-phosphate | 94.5 ± 2.9 | 109.9 ± 1.7 | 94.7 ± 2.4 | 90.7 ± 0.6 | 88.0 ± 1.6 | 85.2 ± 2.6 | 0.9995 |
| Fumaric Acid | 114.5 ± 1.1 | 98.7 ± 1.2 | 105.2 ± 3.1 | 114.7 ± 1.0 | 106.3 ± 2.0 | 108.5 ± 3.0 | 0.9997 |
| Galactosamine | 88.0 ± 2.7 | 100.1 ± 0.8 | 95.1 ± 2.1 | 90.9 ± 2.7 | 88.7 ± 2.4 | 95.7 ± 2.6 | 0.9997 |
| Galactose-1-phosphate | 112.2 ± 1.3 | 97.9 ± 3.0 | 109.8 ± 1.1 | 91.8 ± 1.2 | 107.9 ± 2.3 | 92.6 ± 0.6 | 0.9992 |
| Galactose-6-Phosphate | 105.8 ± 2.7 | 114.5 ± 2.8 | 102.3 ± 1.7 | 112.8 ± 1.6 | 89.6 ± 0.7 | 110.8 ± 3.0 | 0.9999 |
| gamma-Glutamylcysteine | 85.7 ± 1.8 | 94.2 ± 1.8 | 110.6 ± 1.3 | 113.3 ± 3.0 | 111.2 ± 1.8 | 106.9 ± 1.7 | 0.9997 |
| Glucosamine | 96.0 ± 0.9 | 105.4 ± 2.3 | 89.2 ± 2.2 | 102.7 ± 0.3 | 105.9 ± 1.7 | 95.8 ± 1.7 | 0.9997 |
| Glucosamine-6-Phosphate | 89.5 ± 1.4 | 104.3 ± 2.3 | 113.9 ± 0.5 | 87.4 ± 1.9 | 103.1 ± 1.3 | 90.5 ± 0.4 | 0.9992 |
| Glucose | 91.1 ± 0.3 | 105.3 ± 2.6 | 86.1 ± 1.1 | 114.1 ± 3.2 | 106.5 ± 1.7 | 104.8 ± 2.3 | 0.9992 |
| Glucose-1-phosphate | 98.8 ± 0.4 | 109.1 ± 0.4 | 92.0 ± 2.2 | 108.0 ± 2.9 | 101.7 ± 2.2 | 102.4 ± 2.7 | 0.9993 |
| Glucose-6-Phosphate | 105.3 ± 0.7 | 90.1 ± 1.8 | 106.2 ± 2.1 | 94.9 ± 0.7 | 104.6 ± 3.2 | 93.6 ± 0.4 | 0.9996 |
| Glucuronic acid | 91.1 ± 1.7 | 103.7 ± 1.9 | 97.5 ± 1.7 | 106.1 ± 0.4 | 97.3 ± 2.0 | 103.4 ± 2.4 | 0.9996 |
| Glutamic acid | 102.6 ± 3.2 | 95.5 ± 1.5 | 108.0 ± 3.0 | 91.3 ± 1.0 | 97.7 ± 1.0 | 93.6 ± 2.0 | 0.9995 |
| Glutamine | 97.1 ± 0.9 | 97.9 ± 1.9 | 99.4 ± 2.5 | 98.1 ± 2.1 | 104.3 ± 3.2 | 90.4 ± 1.8 | 0.9997 |
| Glutathione | 104.5 ± 3.2 | 87.2 ± 1.5 | 87.7 ± 2.7 | 95.5 ± 2.7 | 113.9 ± 3.0 | 100.5 ± 2.0 | 0.9993 |
| Glutathionylspermine | 87.0 ± 3.0 | 101.3 ± 2.3 | 96.9 ± 1.4 | 89.9 ± 1.9 | 95.5 ± 2.1 | 114.0 ± 2.0 | 0.9997 |
| Glycerophosphocholine | 106.0 ± 3.0 | 97.6 ± 1.9 | 98.7 ± 2.8 | 102.1 ± 2.6 | 86.8 ± 1.8 | 102.8 ± 3.2 | 0.9993 |
| Glycine | 92.4 ± 2.9 | 110.0 ± 0.9 | 95.5 ± 2.3 | 97.9 ± 2.5 | 88.8 ± 1.6 | 103.4 ± 2.3 | 0.9996 |
| Glycolic acid | 94.0 ± 1.6 | 98.2 ± 2.6 | 110.1 ± 1.4 | 86.1 ± 0.3 | 112.4 ± 2.6 | 86.2 ± 0.3 | 0.9996 |
| Guanidinoacetic acid | 113.0 ± 1.9 | 88.7 ± 0.7 | 95.8 ± 1.2 | 90.8 ± 2.6 | 97.0 ± 2.1 | 89.0 ± 0.5 | 0.9996 |
| Guanidinobutanoic acid | 90.5 ± 0.9 | 109.9 ± 2.6 | 113.6 ± 2.1 | 108.2 ± 0.5 | 87.7 ± 1.7 | 105.2 ± 1.3 | 0.9992 |
| Guanine | 113.3 ± 1.6 | 95.6 ± 2.5 | 93.4 ± 2.1 | 107.8 ± 2.0 | 95.2 ± 3.0 | 107.2 ± 2.7 | 0.9993 |
| Guanosine | 89.3 ± 1.2 | 103.6 ± 1.3 | 97.5 ± 0.6 | 113.0 ± 1.8 | 91.6 ± 1.3 | 88.0 ± 1.6 | 0.9996 |
| Guanosine 5'-diphosphate | 86.8 ± 1.2 | 109.9 ± 1.0 | 103.7 ± 3.0 | 106.6 ± 2.4 | 109.9 ± 2.2 | 93.3 ± 1.9 | 0.9998 |
| Guanosine-5'-monophosphate | 112.0 ± 1.0 | 105.0 ± 1.6 | 114.6 ± 1.1 | 109.6 ± 1.5 | 95.6 ± 1.9 | 113.9 ± 1.6 | 0.9998 |
| Hercynine | 97.3 ± 1.8 | 89.3 ± 0.5 | 98.8 ± 0.8 | 93.2 ± 1.1 | 112.8 ± 1.7 | 104.8 ± 1.9 | 0.9992 |
| Hippuric acid | 109.5 ± 2.8 | 109.9 ± 0.3 | 104.6 ± 1.8 | 90.5 ± 0.5 | 114.9 ± 1.3 | 111.0 ± 1.2 | 0.9992 |
| Histamine | 102.6 ± 1.5 | 107.4 ± 0.9 | 104.0 ± 3.0 | 96.8 ± 0.8 | 96.4 ± 2.9 | 96.5 ± 2.1 | 0.9994 |
| Histidine | 99.8 ± 3.2 | 103.5 ± 3.1 | 90.2 ± 0.6 | 93.7 ± 2.7 | 92.8 ± 2.2 | 99.2 ± 2.2 | 0.9996 |
| Homoalanine | 97.1 ± 1.4 | 112.7 ± 2.9 | 102.1 ± 0.3 | 98.9 ± 1.1 | 94.6 ± 1.1 | 111.0 ± 0.7 | 0.9993 |
| Homoarginine | 113.2 ± 0.8 | 110.0 ± 3.0 | 107.3 ± 0.5 | 104.0 ± 2.2 | 105.5 ± 1.4 | 111.7 ± 2.7 | 0.9996 |
| Homocysteine | 102.0 ± 1.9 | 90.8 ± 0.5 | 108.4 ± 3.1 | 103.8 ± 1.2 | 87.3 ± 2.7 | 106.0 ± 1.3 | 0.9999 |
| Homogentisic acid | 90.2 ± 0.9 | 93.4 ± 2.5 | 113.6 ± 3.0 | 114.9 ± 3.1 | 95.8 ± 0.7 | 94.1 ± 3.0 | 0.9991 |
| Homoleucine | 109.6 ± 1.0 | 91.7 ± 2.7 | 92.3 ± 0.4 | 114.3 ± 1.0 | 88.1 ± 2.2 | 90.8 ± 1.0 | 0.9992 |
| Homovanillic acid | 109.7 ± 2.0 | 114.1 ± 1.3 | 105.5 ± 2.9 | 86.8 ± 2.6 | 92.5 ± 0.9 | 92.0 ± 1.0 | 0.9996 |
| Hydantoin-5-propionate | 102.9 ± 1.6 | 107.7 ± 0.9 | 109.3 ± 1.8 | 102.8 ± 0.7 | 102.4 ± 2.2 | 109.5 ± 2.0 | 0.9995 |
| Hypotaurine | 102.0 ± 2.9 | 88.2 ± 3.2 | 110.3 ± 2.0 | 104.2 ± 0.9 | 98.6 ± 2.0 | 105.2 ± 2.0 | 0.9996 |
| Hypoxanthine | 92.2 ± 1.3 | 90.7 ± 1.2 | 90.0 ± 1.2 | 109.2 ± 1.4 | 101.4 ± 2.9 | 99.9 ± 3.1 | 0.9995 |
| Indole-3-acetamide | 100.7 ± 0.5 | 85.7 ± 1.8 | 112.6 ± 0.7 | 88.3 ± 1.5 | 99.0 ± 1.1 | 98.2 ± 2.5 | 0.9997 |
| Inosine | 102.4 ± 2.4 | 90.2 ± 2.9 | 113.0 ± 2.5 | 85.8 ± 2.4 | 86.3 ± 2.5 | 99.1 ± 0.5 | 0.9994 |
| Inosine 5'-monophosphate | 86.1 ± 0.7 | 107.3 ± 2.1 | 96.9 ± 3.0 | 98.9 ± 2.3 | 94.0 ± 2.4 | 107.0 ± 0.5 | 0.9995 |
| Isocitric Acid | 97.8 ± 1.7 | 96.0 ± 1.6 | 92.4 ± 2.4 | 105.4 ± 2.9 | 104.8 ± 2.1 | 102.8 ± 1.4 | 0.9996 |
| Isoleucine | 106.5 ± 2.6 | 88.3 ± 1.1 | 96.6 ± 1.8 | 96.2 ± 2.8 | 112.3 ± 2.9 | 98.7 ± 1.1 | 0.9997 |
| Lactic Acid | 88.2 ± 0.9 | 89.2 ± 3.2 | 87.4 ± 2.9 | 96.0 ± 0.8 | 94.1 ± 2.8 | 102.8 ± 1.3 | 0.9995 |
| Leucine | 85.7 ± 2.0 | 95.6 ± 0.5 | 85.7 ± 0.7 | 87.2 ± 0.7 | 90.1 ± 1.2 | 112.2 ± 3.2 | 0.9995 |
| Levulinic acid | 104.8 ± 3.0 | 101.7 ± 2.5 | 90.4 ± 2.9 | 113.9 ± 2.4 | 96.9 ± 1.7 | 86.8 ± 1.9 | 0.9993 |
| Lysine | 114.6 ± 1.9 | 111.4 ± 2.0 | 96.9 ± 1.0 | 95.7 ± 3.1 | 86.6 ± 1.7 | 104.1 ± 1.8 | 0.9993 |
| Malic Acid | 101.9 ± 2.9 | 93.9 ± 0.5 | 103.0 ± 1.0 | 86.0 ± 2.6 | 110.2 ± 1.2 | 85.8 ± 1.3 | 0.9992 |
| Mannose-1 with Mannose-6-Phosphate | 111.7 ± 2.8 | 103.2 ± 3.1 | 93.6 ± 2.7 | 97.8 ± 3.0 | 100.4 ± 1.0 | 88.1 ± 0.5 | 0.9992 |
| Mesaconic acid | 99.5 ± 0.8 | 86.1 ± 0.4 | 96.5 ± 2.5 | 88.5 ± 1.6 | 106.6 ± 1.1 | 99.2 ± 1.9 | 0.9994 |
| Methionine | 100.5 ± 2.4 | 102.7 ± 2.4 | 110.8 ± 2.9 | 96.6 ± 1.1 | 100.0 ± 0.9 | 94.3 ± 0.6 | 0.9994 |
| Methionine sulfoxide | 91.1 ± 1.0 | 92.7 ± 0.7 | 96.7 ± 2.3 | 94.1 ± 0.9 | 86.2 ± 2.6 | 112.5 ± 1.5 | 0.9994 |
| Methylmalonic Acid | 106.7 ± 1.5 | 89.8 ± 1.2 | 97.2 ± 0.7 | 106.6 ± 2.3 | 97.3 ± 1.9 | 88.1 ± 2.4 | 0.9991 |
| N1-Acetylspermidine | 100.5 ± 2.9 | 112.4 ± 0.6 | 111.5 ± 3.1 | 90.2 ± 2.4 | 98.1 ± 2.4 | 112.0 ± 2.8 | 0.9993 |
| N1-Acetylspermine | 86.1 ± 1.1 | 89.7 ± 1.3 | 108.7 ± 2.4 | 107.9 ± 1.9 | 86.9 ± 0.9 | 94.6 ± 2.9 | 0.9993 |
| N8-Acetylspermidine | 86.1 ± 2.4 | 101.7 ± 2.4 | 97.3 ± 0.3 | 92.0 ± 0.4 | 105.7 ± 1.2 | 91.0 ± 2.7 | 0.9995 |
| N-Acetyl-Aspartyl-Glutamate | 105.3 ± 2.6 | 92.8 ± 2.4 | 101.6 ± 0.3 | 99.1 ± 2.0 | 108.3 ± 1.6 | 94.8 ± 1.4 | 0.9995 |
| N-Acetyl-Aspartyl-Glutamyl-Glutamate | 93.8 ± 1.6 | 87.0 ± 2.9 | 107.8 ± 1.3 | 107.9 ± 2.3 | 104.7 ± 1.5 | 111.5 ± 0.5 | 0.9993 |
| N-Acetyl-Galactosamine | 98.3 ± 1.1 | 102.7 ± 0.5 | 106.5 ± 3.1 | 93.9 ± 1.2 | 92.6 ± 0.4 | 90.7 ± 3.2 | 0.9993 |
| N-Acetyl-Leucine | 109.7 ± 0.9 | 114.0 ± 0.8 | 91.1 ± 1.8 | 97.0 ± 2.9 | 93.8 ± 0.7 | 112.0 ± 2.2 | 0.9994 |
| N-Acetyl-Methionine | 96.8 ± 1.0 | 113.2 ± 1.2 | 106.2 ± 1.5 | 91.9 ± 2.2 | 101.6 ± 3.0 | 86.4 ± 2.9 | 0.9991 |
| N-Acetylneuraminic acid | 106.7 ± 1.4 | 111.7 ± 1.1 | 109.9 ± 1.9 | 94.1 ± 2.0 | 107.6 ± 2.8 | 93.7 ± 2.5 | 0.9991 |
| N-Acetyl-Proline | 102.4 ± 0.9 | 105.5 ± 2.6 | 92.4 ± 1.0 | 105.3 ± 1.8 | 86.1 ± 0.6 | 110.4 ± 1.7 | 0.9994 |
| N-Acetyl-Putrescine | 109.8 ± 1.7 | 98.2 ± 1.3 | 91.9 ± 2.0 | 104.0 ± 2.7 | 112.2 ± 1.2 | 106.8 ± 1.8 | 0.9991 |
| N-Acetyl-Serine | 86.8 ± 2.4 | 105.5 ± 2.5 | 94.1 ± 1.4 | 111.4 ± 3.1 | 98.2 ± 2.1 | 100.0 ± 3.0 | 0.9999 |
| N-Acetyl-Serotonin | 90.4 ± 1.6 | 94.7 ± 0.8 | 94.6 ± 0.8 | 99.2 ± 0.8 | 87.3 ± 0.5 | 92.2 ± 0.6 | 0.9994 |
| N-Acyl-Homoserine-lactone | 113.3 ± 1.0 | 102.0 ± 2.0 | 94.7 ± 3.1 | 103.0 ± 1.6 | 95.2 ± 2.1 | 108.7 ± 3.0 | 0.9999 |
| NAD | 102.5 ± 2.6 | 113.5 ± 0.8 | 91.3 ± 2.5 | 99.7 ± 1.3 | 113.1 ± 1.5 | 109.9 ± 0.6 | 0.9997 |
| NADH | 114.4 ± 0.3 | 94.9 ± 1.4 | 89.1 ± 2.0 | 88.3 ± 0.4 | 91.4 ± 1.4 | 108.0 ± 2.9 | 0.9991 |
| NADP | 99.6 ± 3.2 | 112.2 ± 1.3 | 98.6 ± 1.3 | 90.2 ± 2.1 | 92.0 ± 2.9 | 105.1 ± 0.6 | 0.9996 |
| NADPH | 88.8 ± 2.4 | 113.2 ± 2.8 | 90.5 ± 2.8 | 87.4 ± 3.0 | 110.7 ± 1.0 | 113.4 ± 1.3 | 0.9998 |
| N-alpha-Acetyl-lysine | 94.4 ± 3.1 | 94.8 ± 0.4 | 114.4 ± 1.9 | 114.7 ± 1.3 | 103.9 ± 0.5 | 101.2 ± 2.7 | 0.9995 |
| N-alpha-Acetyl-Ornithine | 107.9 ± 0.8 | 95.2 ± 0.8 | 104.2 ± 1.5 | 102.6 ± 0.8 | 86.5 ± 0.3 | 112.9 ± 2.6 | 0.9993 |
| N-Carbamoyl-Proline | 105.6 ± 0.5 | 98.4 ± 2.9 | 100.3 ± 2.1 | 108.5 ± 0.7 | 92.2 ± 0.5 | 104.6 ± 1.0 | 0.9996 |
| N-Carbamoyl-Sarcosine | 94.1 ± 1.8 | 108.0 ± 1.0 | 110.3 ± 2.7 | 87.7 ± 2.1 | 98.0 ± 0.3 | 110.6 ± 2.2 | 0.9996 |
| N-Epsilon-Acetyl-lysine | 108.0 ± 0.3 | 92.0 ± 0.6 | 112.6 ± 2.3 | 89.4 ± 0.7 | 89.8 ± 0.7 | 98.2 ± 2.9 | 0.9996 |
| Nicotinamide | 101.8 ± 2.9 | 88.6 ± 1.6 | 108.8 ± 2.1 | 85.3 ± 1.4 | 105.9 ± 1.6 | 110.9 ± 1.8 | 0.9993 |
| N-Methyl-Aspartic Acid | 86.1 ± 3.0 | 96.5 ± 1.5 | 105.1 ± 3.1 | 89.7 ± 3.0 | 108.2 ± 1.6 | 108.0 ± 2.2 | 0.9994 |
| O-Phosphoethanolamine | 109.4 ± 2.8 | 91.2 ± 0.5 | 113.9 ± 0.8 | 90.0 ± 2.8 | 111.2 ± 1.3 | 103.7 ± 1.9 | 0.9993 |
| Ophthalmic acid | 94.7 ± 1.9 | 85.4 ± 0.9 | 114.1 ± 0.7 | 104.6 ± 1.8 | 92.3 ± 0.7 | 85.5 ± 1.7 | 0.9998 |
| Ornithine | 90.2 ± 1.4 | 98.4 ± 1.3 | 102.0 ± 2.4 | 107.9 ± 2.6 | 104.8 ± 0.5 | 106.8 ± 2.8 | 0.9998 |
| Orotidine 5'-monophosphate | 88.3 ± 0.8 | 91.7 ± 2.8 | 107.6 ± 3.1 | 107.7 ± 1.7 | 91.3 ± 2.1 | 94.8 ± 2.1 | 0.9997 |
| Oxaloacetate | 104.7 ± 3.0 | 109.6 ± 1.8 | 110.7 ± 2.2 | 88.6 ± 0.5 | 85.6 ± 0.6 | 91.8 ± 3.1 | 0.9992 |
| Pantothenic acid | 87.2 ± 1.3 | 88.7 ± 1.6 | 87.1 ± 1.3 | 103.3 ± 1.0 | 96.0 ± 1.5 | 90.0 ± 0.6 | 0.9992 |
| Phenylalanine | 112.8 ± 0.4 | 102.0 ± 2.7 | 99.4 ± 2.8 | 109.8 ± 1.1 | 109.7 ± 1.0 | 111.9 ± 1.2 | 0.9992 |
| Phenylpyruvate | 96.0 ± 3.0 | 92.5 ± 0.6 | 103.6 ± 2.5 | 106.6 ± 1.0 | 87.0 ± 1.2 | 93.7 ± 2.1 | 0.9992 |
| Phosphodimethylethanolamine | 107.0 ± 1.5 | 85.3 ± 1.6 | 95.3 ± 0.9 | 105.3 ± 1.9 | 88.4 ± 2.7 | 111.5 ± 0.6 | 0.9992 |
| Phosphoenolpyruvate | 113.2 ± 1.2 | 114.8 ± 3.2 | 105.9 ± 2.9 | 93.9 ± 1.4 | 97.7 ± 1.9 | 112.6 ± 1.0 | 0.9994 |
| Phosphorylcholine | 109.9 ± 1.3 | 100.4 ± 0.9 | 89.4 ± 1.1 | 96.7 ± 2.2 | 91.7 ± 2.1 | 87.5 ± 2.9 | 0.9997 |
| Pipecolic acid | 105.3 ± 2.5 | 104.5 ± 3.1 | 93.1 ± 0.6 | 111.5 ± 2.0 | 94.7 ± 0.9 | 102.0 ± 2.6 | 0.9993 |
| Proline | 89.3 ± 1.1 | 87.0 ± 2.0 | 96.5 ± 1.1 | 107.8 ± 3.0 | 110.5 ± 0.9 | 97.5 ± 0.3 | 0.9996 |
| Putrescine | 95.6 ± 2.5 | 100.0 ± 3.2 | 93.1 ± 2.5 | 87.6 ± 1.9 | 91.4 ± 2.9 | 101.7 ± 0.4 | 0.9994 |
| Pyridoxal | 114.1 ± 3.2 | 94.7 ± 2.1 | 85.7 ± 0.7 | 103.7 ± 3.1 | 98.8 ± 1.0 | 103.3 ± 1.4 | 0.9993 |
| Pyridoxamine | 88.0 ± 0.4 | 102.5 ± 2.2 | 89.9 ± 0.3 | 114.9 ± 1.8 | 100.9 ± 2.2 | 113.1 ± 0.9 | 0.9992 |
| Pyridoxine | 106.8 ± 1.0 | 111.7 ± 1.3 | 99.6 ± 1.9 | 100.4 ± 0.9 | 91.3 ± 0.3 | 105.3 ± 1.9 | 0.9999 |
| Pyroglutamic acid | 89.4 ± 1.9 | 102.5 ± 0.9 | 106.1 ± 2.8 | 103.1 ± 3.0 | 104.0 ± 2.0 | 95.7 ± 2.7 | 0.9994 |
| Pyrrolysine | 100.0 ± 1.5 | 103.1 ± 2.9 | 87.8 ± 1.6 | 106.1 ± 1.4 | 94.1 ± 2.5 | 108.6 ± 0.6 | 0.9995 |
| Pyruvic Acid | 101.4 ± 2.9 | 92.3 ± 3.1 | 93.7 ± 2.2 | 109.7 ± 2.1 | 86.2 ± 1.4 | 97.1 ± 1.3 | 0.9992 |
| Riboflavin | 92.7 ± 1.6 | 97.3 ± 2.9 | 85.3 ± 2.6 | 92.6 ± 2.3 | 96.3 ± 3.0 | 111.9 ± 0.9 | 0.9994 |
| Ribose-5-Phosphate | 104.0 ± 2.4 | 113.1 ± 2.0 | 87.2 ± 2.0 | 112.1 ± 2.2 | 94.0 ± 2.7 | 86.3 ± 1.4 | 0.9995 |
| Ribulose-1-Phosphate | 85.7 ± 1.1 | 109.2 ± 1.9 | 108.5 ± 1.0 | 114.3 ± 2.2 | 89.6 ± 2.5 | 88.4 ± 1.2 | 0.9996 |
| Ribulose-5-Phosphate | 86.1 ± 2.2 | 107.1 ± 2.8 | 97.5 ± 2.5 | 88.6 ± 2.9 | 96.9 ± 0.9 | 114.9 ± 0.9 | 0.9997 |
| Saccharopine | 91.9 ± 2.9 | 85.4 ± 2.4 | 100.3 ± 1.9 | 88.2 ± 0.4 | 100.4 ± 2.4 | 91.1 ± 1.2 | 0.9993 |
| S-Adenosylhomocysteine | 103.8 ± 1.1 | 104.8 ± 3.1 | 105.6 ± 1.1 | 99.8 ± 0.9 | 87.9 ± 3.0 | 87.1 ± 2.4 | 0.9997 |
| S-Adenosylmethioninamine | 107.3 ± 3.0 | 113.1 ± 2.9 | 108.9 ± 2.3 | 99.1 ± 0.5 | 114.1 ± 2.7 | 114.7 ± 2.9 | 0.9991 |
| S-Adenosylmethionine | 98.2 ± 1.6 | 112.3 ± 2.9 | 113.8 ± 2.5 | 110.4 ± 0.4 | 85.6 ± 0.3 | 85.5 ± 2.6 | 0.9996 |
| Sarcosine | 112.6 ± 2.1 | 93.1 ± 1.5 | 92.8 ± 0.8 | 114.8 ± 1.4 | 100.7 ± 2.1 | 87.1 ± 1.5 | 0.9995 |
| Sedoheptulose-7-phosphate | 90.3 ± 2.9 | 105.7 ± 2.6 | 114.0 ± 1.4 | 104.5 ± 0.5 | 102.1 ± 0.3 | 103.4 ± 2.2 | 0.9997 |
| Selenohomocysteine | 85.8 ± 1.3 | 111.5 ± 2.0 | 112.9 ± 2.4 | 90.7 ± 0.7 | 96.1 ± 3.2 | 114.4 ± 2.0 | 0.9995 |
| Selenomethionine | 113.1 ± 1.0 | 100.5 ± 2.1 | 113.6 ± 0.8 | 110.9 ± 1.4 | 97.9 ± 3.0 | 101.8 ± 1.0 | 0.9997 |
| Serine | 103.6 ± 1.6 | 89.1 ± 1.5 | 99.6 ± 1.3 | 86.2 ± 1.2 | 103.0 ± 0.9 | 106.5 ± 3.1 | 0.9997 |
| S-Methyl-Thioinosine | 94.1 ± 1.2 | 90.9 ± 0.8 | 105.7 ± 0.9 | 97.4 ± 1.1 | 96.9 ± 2.8 | 87.2 ± 1.7 | 0.9994 |
| Spermidine | 109.5 ± 3.1 | 98.4 ± 3.1 | 110.3 ± 0.4 | 105.0 ± 1.0 | 95.7 ± 0.7 | 93.3 ± 2.8 | 0.9998 |
| Spermine | 91.7 ± 1.1 | 92.2 ± 0.9 | 85.2 ± 1.0 | 93.1 ± 3.0 | 87.9 ± 3.0 | 98.7 ± 1.7 | 0.9995 |
| Succinate | 85.0 ± 2.4 | 103.8 ± 0.4 | 111.1 ± 0.8 | 112.4 ± 0.6 | 108.0 ± 1.1 | 86.6 ± 1.2 | 0.9995 |
| Sucrose | 88.6 ± 2.7 | 93.6 ± 0.9 | 95.0 ± 2.7 | 103.4 ± 2.4 | 95.8 ± 1.9 | 89.8 ± 0.6 | 0.9996 |
| Symmetric dimethylarginine | 113.7 ± 0.8 | 114.9 ± 1.4 | 87.0 ± 0.8 | 113.5 ± 1.2 | 97.8 ± 3.0 | 93.1 ± 2.2 | 0.9997 |
| Taurine | 109.0 ± 0.3 | 90.6 ± 2.4 | 106.3 ± 1.2 | 114.4 ± 1.8 | 99.8 ± 0.4 | 113.7 ± 1.4 | 0.9999 |
| Tetrahydroneopterin | 94.5 ± 0.9 | 89.7 ± 3.0 | 86.3 ± 3.2 | 85.9 ± 1.0 | 88.0 ± 2.6 | 91.6 ± 2.0 | 0.9995 |
| Thiamine | 88.6 ± 0.7 | 101.5 ± 2.7 | 99.9 ± 2.6 | 114.9 ± 0.6 | 93.0 ± 0.5 | 114.4 ± 1.9 | 0.9996 |
| Thiamine monophosphate | 113.0 ± 3.0 | 114.3 ± 1.4 | 98.3 ± 0.4 | 86.7 ± 1.2 | 101.8 ± 1.6 | 114.2 ± 3.0 | 0.9997 |
| Threonic acid | 91.8 ± 2.9 | 91.6 ± 1.0 | 96.8 ± 1.2 | 95.2 ± 3.1 | 98.9 ± 1.1 | 112.9 ± 2.6 | 0.9994 |
| Threonine | 105.6 ± 2.1 | 94.6 ± 0.3 | 101.2 ± 2.2 | 87.5 ± 2.2 | 91.0 ± 3.0 | 88.4 ± 2.8 | 0.9998 |
| Thymidine | 104.5 ± 1.2 | 88.6 ± 0.8 | 91.9 ± 2.5 | 88.7 ± 3.2 | 97.9 ± 1.6 | 113.1 ± 1.0 | 0.9995 |
| Thymidine Monophosphate | 103.1 ± 2.1 | 112.1 ± 2.4 | 102.1 ± 1.0 | 91.4 ± 2.0 | 87.5 ± 1.9 | 98.5 ± 1.5 | 0.9991 |
| Thymine | 86.4 ± 2.5 | 91.8 ± 2.8 | 88.4 ± 2.7 | 110.5 ± 3.1 | 103.9 ± 2.5 | 105.8 ± 0.5 | 0.9998 |
| Trigonelleine | 86.2 ± 2.6 | 89.6 ± 2.2 | 100.1 ± 2.3 | 94.9 ± 0.5 | 109.1 ± 0.3 | 108.2 ± 0.7 | 0.9996 |
| Trimethylamine N-oxide | 92.5 ± 2.3 | 102.2 ± 0.7 | 101.5 ± 1.6 | 87.1 ± 0.4 | 91.3 ± 2.0 | 98.1 ± 0.9 | 0.9991 |
| Trimethyllysine | 105.9 ± 2.1 | 109.3 ± 1.8 | 106.7 ± 3.0 | 89.4 ± 1.5 | 105.2 ± 2.1 | 93.4 ± 1.1 | 0.9995 |
| Tryptophan | 92.0 ± 0.4 | 100.3 ± 2.7 | 88.4 ± 1.4 | 97.6 ± 2.7 | 88.7 ± 2.6 | 86.9 ± 2.9 | 0.9997 |
| Tyramine | 104.2 ± 1.0 | 101.7 ± 1.8 | 89.7 ± 2.4 | 107.7 ± 0.9 | 94.0 ± 2.9 | 90.5 ± 1.2 | 0.9996 |
| Tyrosine | 102.5 ± 1.1 | 107.9 ± 1.0 | 98.1 ± 3.0 | 113.9 ± 2.8 | 107.7 ± 2.6 | 111.3 ± 1.7 | 0.9994 |
| Uracil | 112.3 ± 0.9 | 105.5 ± 0.9 | 85.2 ± 2.8 | 101.7 ± 2.3 | 108.6 ± 0.8 | 89.4 ± 3.2 | 0.9994 |
| Uridine 5'-diphosphate | 107.1 ± 3.0 | 88.3 ± 1.8 | 112.8 ± 2.6 | 105.3 ± 1.3 | 101.6 ± 0.7 | 91.7 ± 2.8 | 0.9997 |
| Uridine Monophosphate | 113.7 ± 2.3 | 86.1 ± 2.1 | 114.9 ± 2.4 | 114.1 ± 1.1 | 93.0 ± 2.9 | 88.2 ± 0.4 | 0.9999 |
| Uridine-5-Diphosphogalactose | 108.0 ± 0.6 | 86.3 ± 1.4 | 98.3 ± 0.6 | 114.6 ± 2.5 | 111.0 ± 1.1 | 104.7 ± 2.3 | 0.9995 |
| Urocanic acid | 94.7 ± 0.7 | 101.8 ± 2.0 | 96.2 ± 0.6 | 112.2 ± 2.3 | 97.8 ± 0.4 | 105.9 ± 0.4 | 0.9996 |
| UTP | 85.8 ± 0.5 | 92.7 ± 1.0 | 89.6 ± 1.9 | 104.4 ± 1.6 | 105.6 ± 0.5 | 102.2 ± 1.6 | 0.9996 |
| Valine | 97.9 ± 2.6 | 112.1 ± 0.7 | 85.3 ± 2.4 | 88.6 ± 3.2 | 114.2 ± 2.0 | 109.1 ± 3.1 | 0.9993 |
| XMP | 110.4 ± 2.5 | 92.7 ± 0.4 | 97.6 ± 0.5 | 98.1 ± 1.1 | 91.7 ± 0.7 | 106.0 ± 1.5 | 0.9991 |
| Xylulose-5-Phosphate | 87.7 ± 2.0 | 97.5 ± 2.0 | 96.8 ± 0.5 | 101.9 ± 2.6 | 89.8 ± 1.0 | 96.7 ± 3.0 | 0.9998 |

### **Table S11** | Calibration values for final method conditions (r^2^-value) and percent found of nominal calibrator concentration (Urine extract as matrix).

| Calibrator concentration accuracy  [%] | 10 ng/mL | 25 ng/mL | 50 ng/mL | 100 ng/mL | 500 ng/mL | 1000 ng/mL | r^2^-Value  matrix-matched calibration |
| --- | --- | --- | --- | --- | --- | --- | --- |
| 1-Methylguanine | 109.6 ± 3.1 | 105.8 ± 2.1 | 94.5 ± 3.0 | 95.6 ± 1.8 | 85.0 ± 2.6 | 97.7 ± 2.2 | 0.9997 |
| 1-Methylhistidine | 96.2 ± 1.5 | 114.5 ± 2.6 | 105.2 ± 2.7 | 96.3 ± 3.1 | 91.5 ± 1.9 | 106.4 ± 1.0 | 0.9995 |
| 1-Methylnicotinamide | 105.1 ± 0.6 | 87.6 ± 1.5 | 102.2 ± 2.0 | 102.6 ± 0.5 | 95.5 ± 1.9 | 105.1 ± 2.7 | 0.9997 |
| 2-Aminoadipic Acid | 86.6 ± 0.4 | 97.3 ± 2.9 | 97.6 ± 2.7 | 113.7 ± 2.8 | 112.1 ± 1.3 | 104.4 ± 1.9 | 0.9998 |
| 2-Deoxyadenosine | 98.1 ± 1.1 | 112.6 ± 2.9 | 98.5 ± 3.2 | 98.6 ± 1.3 | 105.6 ± 0.5 | 100.3 ± 0.4 | 0.9992 |
| 2'-Deoxyadenosine 5'-monophosphate | 85.7 ± 1.7 | 94.7 ± 0.8 | 112.5 ± 2.1 | 102.4 ± 1.7 | 101.1 ± 0.6 | 113.1 ± 2.1 | 0.9998 |
| 2'-deoxyadenosine 5'-triphosphate | 108.7 ± 2.1 | 104.9 ± 0.7 | 99.2 ± 2.7 | 102.8 ± 2.9 | 92.6 ± 1.4 | 105.7 ± 2.0 | 0.9996 |
| 2'-Deoxycytidine-5'-monophosphate | 90.4 ± 1.2 | 87.0 ± 0.6 | 103.3 ± 2.7 | 98.5 ± 2.5 | 93.8 ± 2.4 | 86.6 ± 1.6 | 0.9994 |
| 2-Deoxyguanosine | 94.2 ± 2.3 | 109.7 ± 1.0 | 85.9 ± 2.2 | 109.4 ± 2.4 | 112.8 ± 1.3 | 107.0 ± 1.3 | 0.9997 |
| 2-hydroxybutyric acid | 113.2 ± 1.9 | 98.3 ± 1.1 | 109.7 ± 1.6 | 95.2 ± 0.9 | 114.5 ± 2.3 | 91.9 ± 1.4 | 0.9997 |
| 2-hydroxyglutaric acid | 89.2 ± 1.3 | 86.0 ± 2.2 | 92.3 ± 1.3 | 89.1 ± 0.7 | 91.2 ± 2.8 | 97.2 ± 2.0 | 0.9998 |
| 2-Hydroxyisocaproic acid | 112.1 ± 1.4 | 108.0 ± 1.2 | 89.7 ± 2.3 | 101.1 ± 2.4 | 98.1 ± 1.5 | 101.3 ± 1.5 | 0.9998 |
| 2-ketobutyric acid | 98.3 ± 0.7 | 86.1 ± 1.5 | 96.1 ± 2.8 | 114.6 ± 3.0 | 86.0 ± 1.3 | 101.0 ± 0.8 | 0.9998 |
| 2-oxoadipic acid | 87.5 ± 0.7 | 94.5 ± 2.9 | 90.5 ± 0.7 | 111.1 ± 0.5 | 89.9 ± 0.5 | 106.3 ± 1.3 | 0.9995 |
| 2-Phenylglycine | 101.5 ± 0.5 | 97.5 ± 0.7 | 99.7 ± 0.4 | 101.5 ± 2.6 | 85.9 ± 1.7 | 89.8 ± 1.4 | 0.9999 |
| 2-phosphoglycerate | 103.5 ± 1.6 | 105.3 ± 1.4 | 91.9 ± 1.8 | 93.4 ± 1.7 | 91.6 ± 1.2 | 91.5 ± 2.7 | 0.9997 |
| 3-hydroxy-3-methyl-2-oxobutanoate | 86.3 ± 0.6 | 97.1 ± 1.2 | 109.1 ± 1.2 | 98.2 ± 2.2 | 105.1 ± 1.9 | 91.0 ± 1.4 | 0.9996 |
| 3-hydroxybutyric acid | 91.3 ± 2.2 | 95.3 ± 2.3 | 99.2 ± 2.6 | 97.2 ± 1.0 | 87.7 ± 0.3 | 88.1 ± 1.9 | 0.9997 |
| 3-Methylhistidine | 100.3 ± 0.4 | 105.5 ± 1.7 | 100.4 ± 0.8 | 85.0 ± 2.4 | 101.1 ± 0.5 | 107.0 ± 0.5 | 0.9991 |
| 3-Phosphoglyceraldehyde | 108.7 ± 1.2 | 92.5 ± 1.2 | 91.1 ± 1.8 | 111.6 ± 1.3 | 94.8 ± 0.5 | 107.2 ± 0.8 | 0.9993 |
| 3-phosphoglycerate | 99.1 ± 3.1 | 98.6 ± 2.4 | 112.5 ± 1.2 | 88.0 ± 0.6 | 85.2 ± 1.7 | 114.5 ± 2.9 | 0.9996 |
| 4-amino-butyric acid | 90.6 ± 0.6 | 107.7 ± 0.8 | 112.1 ± 1.3 | 87.7 ± 2.3 | 107.4 ± 1.2 | 112.6 ± 2.0 | 0.9997 |
| 4-aspartyl-phosphate | 85.1 ± 3.1 | 101.7 ± 2.3 | 100.0 ± 1.8 | 96.4 ± 2.7 | 88.0 ± 3.0 | 108.7 ± 1.7 | 0.9993 |
| 4-Hydroxybenzoic acid | 95.3 ± 1.9 | 114.0 ± 2.0 | 93.9 ± 0.7 | 101.3 ± 2.9 | 88.5 ± 1.9 | 96.0 ± 3.1 | 0.9995 |
| 4-Hydroxyphenylacetylglycine | 94.3 ± 2.1 | 110.9 ± 1.0 | 97.0 ± 3.1 | 94.3 ± 1.6 | 90.3 ± 1.2 | 88.3 ± 2.1 | 0.9995 |
| 4-methyl-2-oxopentanoate | 88.8 ± 0.3 | 89.4 ± 1.3 | 108.3 ± 1.5 | 108.6 ± 3.1 | 86.5 ± 2.4 | 113.8 ± 0.4 | 0.9996 |
| 4-Oxoglutaramate | 91.1 ± 2.8 | 105.3 ± 1.1 | 86.8 ± 2.0 | 96.7 ± 2.7 | 94.3 ± 1.7 | 86.3 ± 0.8 | 0.9995 |
| 5-Hydroxyindolepyruvate | 112.3 ± 2.3 | 108.6 ± 0.9 | 111.9 ± 1.2 | 85.8 ± 1.2 | 100.6 ± 3.1 | 112.3 ± 3.0 | 0.9995 |
| 5-Hydroxylysine | 104.4 ± 2.3 | 99.4 ± 2.7 | 100.7 ± 0.5 | 108.2 ± 2.5 | 110.4 ± 1.9 | 104.2 ± 2.7 | 0.9992 |
| 5-Hydroxy-Tryptophan | 96.4 ± 0.5 | 85.5 ± 2.2 | 88.0 ± 3.1 | 106.0 ± 2.4 | 109.3 ± 2.0 | 111.6 ± 2.0 | 0.9995 |
| 5-Phosphoribosylamine | 92.8 ± 1.3 | 86.1 ± 1.3 | 105.3 ± 2.7 | 108.3 ± 0.5 | 109.3 ± 2.3 | 111.2 ± 0.5 | 0.9996 |
| 6-phosphogluconate | 96.9 ± 1.9 | 104.9 ± 1.8 | 113.4 ± 0.8 | 113.5 ± 0.6 | 105.7 ± 2.4 | 114.2 ± 2.3 | 0.9999 |
| Acetylcarnitine | 90.4 ± 3.2 | 114.2 ± 1.2 | 114.1 ± 1.4 | 89.1 ± 1.1 | 97.5 ± 0.7 | 99.9 ± 2.6 | 0.9995 |
| Aconitic Acid | 90.5 ± 2.7 | 86.1 ± 1.2 | 102.0 ± 0.4 | 98.4 ± 2.3 | 102.3 ± 0.9 | 99.2 ± 2.2 | 0.9996 |
| Adenine | 109.3 ± 2.8 | 95.9 ± 1.0 | 107.7 ± 2.7 | 104.2 ± 1.9 | 88.1 ± 1.9 | 101.8 ± 3.1 | 0.9993 |
| Adenosine | 89.8 ± 2.3 | 109.2 ± 2.2 | 103.4 ± 1.0 | 91.0 ± 1.5 | 103.4 ± 2.4 | 87.6 ± 2.4 | 0.9997 |
| Adenosine 3',5'-cyclic monophosphate | 106.2 ± 0.6 | 103.7 ± 1.1 | 110.3 ± 0.8 | 93.0 ± 2.9 | 98.9 ± 2.9 | 106.0 ± 2.1 | 0.9996 |
| Adenosine Monophosphate | 87.8 ± 2.7 | 91.1 ± 1.4 | 89.4 ± 2.0 | 85.5 ± 1.9 | 101.0 ± 0.7 | 94.8 ± 2.8 | 0.9993 |
| Adenosine-5'-diphosphate | 114.2 ± 0.7 | 92.3 ± 3.0 | 90.0 ± 1.9 | 87.8 ± 2.0 | 105.9 ± 1.3 | 89.8 ± 1.6 | 0.9994 |
| Adenosine-5'-triphosphate | 99.5 ± 1.3 | 94.5 ± 2.1 | 107.4 ± 2.3 | 107.3 ± 2.2 | 93.1 ± 0.8 | 113.9 ± 2.6 | 0.9995 |
| Adenylosuccinate | 96.2 ± 1.7 | 93.0 ± 1.9 | 104.3 ± 2.6 | 108.0 ± 3.1 | 112.5 ± 1.8 | 103.7 ± 1.6 | 0.9997 |
| Alanine | 92.2 ± 0.4 | 101.2 ± 0.8 | 93.1 ± 1.6 | 106.1 ± 2.8 | 113.3 ± 1.0 | 91.9 ± 1.1 | 0.9992 |
| alpha-ketoglutaric acid | 111.6 ± 2.8 | 96.5 ± 0.7 | 109.2 ± 2.3 | 106.0 ± 1.4 | 104.4 ± 1.7 | 94.3 ± 1.2 | 0.9993 |
| Anserine | 113.9 ± 1.7 | 114.6 ± 2.1 | 96.9 ± 0.3 | 96.1 ± 1.0 | 92.1 ± 0.3 | 94.4 ± 0.8 | 0.9996 |
| Arginine | 112.6 ± 1.5 | 99.5 ± 2.9 | 115.0 ± 2.9 | 107.9 ± 2.8 | 113.8 ± 2.0 | 109.8 ± 2.0 | 0.9996 |
| Argininosuccinic Acid | 102.3 ± 1.3 | 104.1 ± 2.8 | 113.3 ± 2.2 | 97.8 ± 2.5 | 92.6 ± 1.6 | 85.9 ± 2.0 | 0.9996 |
| Asparagine | 88.8 ± 3.1 | 85.9 ± 1.8 | 113.4 ± 1.4 | 105.9 ± 1.4 | 93.2 ± 0.5 | 100.1 ± 3.1 | 0.9998 |
| Aspartic acid | 113.2 ± 2.7 | 105.6 ± 0.6 | 90.6 ± 2.1 | 113.0 ± 3.0 | 114.7 ± 1.1 | 91.1 ± 0.6 | 0.9996 |
| Asymmetric dimethylarginine | 102.6 ± 1.6 | 107.8 ± 2.9 | 105.9 ± 1.7 | 101.7 ± 1.6 | 100.6 ± 1.3 | 89.4 ± 2.2 | 0.9991 |
| Azelaic acid | 98.2 ± 1.1 | 93.9 ± 1.8 | 114.5 ± 1.8 | 109.7 ± 1.8 | 113.5 ± 2.4 | 96.5 ± 0.4 | 0.9994 |
| beta-Alanine | 96.4 ± 0.6 | 95.2 ± 3.0 | 110.1 ± 2.7 | 109.9 ± 1.4 | 101.5 ± 2.3 | 112.3 ± 3.1 | 0.9993 |
| Betaine | 113.0 ± 2.2 | 114.3 ± 2.5 | 97.2 ± 0.6 | 90.3 ± 1.7 | 100.1 ± 2.7 | 102.6 ± 2.7 | 0.9997 |
| Biopterin | 96.7 ± 2.4 | 107.3 ± 0.7 | 86.9 ± 1.1 | 92.2 ± 1.6 | 90.9 ± 1.8 | 88.5 ± 1.1 | 0.9997 |
| Biotin | 108.2 ± 3.0 | 113.4 ± 2.9 | 107.0 ± 2.5 | 109.5 ± 0.6 | 91.0 ± 2.0 | 104.5 ± 3.0 | 0.9991 |
| Carnitine | 98.1 ± 2.7 | 102.8 ± 2.1 | 92.0 ± 3.1 | 109.2 ± 1.7 | 94.7 ± 1.4 | 108.2 ± 0.6 | 0.9993 |
| Carnosine | 91.4 ± 0.7 | 98.3 ± 2.9 | 103.5 ± 3.0 | 105.7 ± 2.1 | 90.4 ± 0.7 | 114.0 ± 1.0 | 0.9996 |
| CDP-choline | 99.3 ± 2.4 | 102.5 ± 1.9 | 88.7 ± 2.4 | 98.5 ± 0.5 | 94.4 ± 2.7 | 98.0 ± 3.0 | 0.9994 |
| cGMP | 89.6 ± 2.7 | 110.2 ± 1.5 | 106.6 ± 2.5 | 88.5 ± 1.2 | 112.8 ± 1.0 | 105.8 ± 1.2 | 0.9997 |
| Choline | 105.7 ± 3.1 | 114.7 ± 3.1 | 109.6 ± 2.8 | 113.4 ± 2.7 | 101.5 ± 0.9 | 97.8 ± 1.0 | 0.9995 |
| Cinnamic acid | 91.5 ± 2.1 | 113.7 ± 2.8 | 104.6 ± 1.8 | 105.4 ± 1.0 | 114.9 ± 2.5 | 97.0 ± 1.2 | 0.9997 |
| Citraconic acid | 106.2 ± 1.6 | 114.0 ± 0.6 | 97.1 ± 2.9 | 96.1 ± 2.6 | 93.2 ± 0.8 | 85.6 ± 1.2 | 0.9997 |
| Citramalic acid | 110.6 ± 2.7 | 101.5 ± 1.2 | 113.4 ± 2.5 | 96.8 ± 0.4 | 93.5 ± 2.1 | 109.5 ± 1.8 | 0.9991 |
| Citric Acid | 107.7 ± 2.1 | 97.7 ± 0.3 | 88.0 ± 0.4 | 111.1 ± 1.0 | 113.1 ± 1.4 | 114.6 ± 0.3 | 0.9997 |
| Citrulline | 86.0 ± 0.6 | 108.7 ± 1.1 | 107.7 ± 1.5 | 92.5 ± 1.8 | 107.3 ± 2.5 | 114.0 ± 2.8 | 0.9997 |
| Creatine | 108.0 ± 1.1 | 91.3 ± 2.4 | 96.8 ± 2.8 | 97.2 ± 2.7 | 86.7 ± 2.3 | 98.2 ± 2.5 | 0.9998 |
| Creatinine | 88.0 ± 2.4 | 105.0 ± 2.8 | 103.3 ± 0.8 | 112.7 ± 2.7 | 110.9 ± 3.1 | 91.1 ± 1.7 | 0.9998 |
| Cystathionine | 87.8 ± 1.8 | 86.8 ± 2.8 | 94.7 ± 0.7 | 100.4 ± 2.2 | 88.3 ± 1.8 | 100.5 ± 2.9 | 0.9992 |
| Cysteamine | 107.1 ± 1.6 | 100.5 ± 2.2 | 102.6 ± 1.6 | 85.4 ± 1.6 | 94.7 ± 2.4 | 112.1 ± 1.4 | 0.9994 |
| Cysteine | 114.3 ± 2.1 | 109.7 ± 3.0 | 87.8 ± 0.7 | 96.8 ± 1.9 | 110.6 ± 2.8 | 113.1 ± 3.0 | 0.9995 |
| Cysteinylglycine | 104.7 ± 0.9 | 112.9 ± 0.9 | 85.3 ± 0.7 | 90.4 ± 0.8 | 112.3 ± 2.0 | 88.8 ± 2.2 | 0.9997 |
| Cystine | 97.3 ± 1.8 | 91.7 ± 2.3 | 88.7 ± 2.8 | 107.7 ± 1.8 | 111.4 ± 0.9 | 100.3 ± 1.8 | 0.9999 |
| Cytidine | 88.4 ± 0.4 | 113.8 ± 1.2 | 109.8 ± 2.8 | 92.6 ± 0.4 | 112.7 ± 0.8 | 106.0 ± 0.7 | 0.9997 |
| Cytidine 3',5'-cyclic monophosphate | 103.1 ± 1.6 | 98.0 ± 2.1 | 100.8 ± 1.3 | 92.0 ± 2.5 | 113.1 ± 2.2 | 86.7 ± 1.0 | 0.9998 |
| Cytidine 5'-diphosphate | 99.2 ± 1.5 | 89.8 ± 2.6 | 99.0 ± 1.8 | 109.2 ± 1.5 | 101.6 ± 1.6 | 95.4 ± 1.3 | 0.9998 |
| Cytidine 5'-triphosphate | 98.5 ± 3.1 | 96.2 ± 1.0 | 91.3 ± 1.6 | 112.1 ± 1.9 | 98.8 ± 1.3 | 99.7 ± 0.5 | 0.9992 |
| Cytidine monophosphate | 90.2 ± 1.4 | 98.4 ± 1.5 | 89.3 ± 0.8 | 91.8 ± 2.5 | 103.1 ± 1.4 | 111.8 ± 1.5 | 0.9991 |
| Cytosine | 110.7 ± 0.3 | 87.9 ± 2.1 | 86.6 ± 0.4 | 110.1 ± 2.3 | 100.2 ± 2.8 | 110.9 ± 1.5 | 0.9998 |
| dADP | 103.6 ± 1.7 | 107.9 ± 1.8 | 111.8 ± 1.0 | 111.9 ± 1.9 | 107.2 ± 1.8 | 97.4 ± 3.0 | 0.9996 |
| Diethanolamine | 97.1 ± 2.2 | 94.1 ± 0.9 | 87.5 ± 0.3 | 89.5 ± 3.1 | 97.7 ± 2.0 | 99.2 ± 1.1 | 0.9999 |
| Dihydrobiopterin | 104.6 ± 3.0 | 101.0 ± 1.7 | 111.7 ± 2.7 | 93.3 ± 2.1 | 89.9 ± 2.3 | 99.8 ± 2.5 | 0.9992 |
| Dihydroneopterin triphosphate | 103.6 ± 1.5 | 111.2 ± 2.6 | 100.9 ± 3.2 | 96.3 ± 1.5 | 114.2 ± 0.9 | 88.0 ± 2.5 | 0.9993 |
| Dihydroxyacetonephosphate | 85.6 ± 2.9 | 110.4 ± 2.3 | 112.1 ± 0.5 | 93.7 ± 2.6 | 94.7 ± 2.5 | 99.5 ± 2.8 | 0.9994 |
| Dimethylglycine | 96.7 ± 0.9 | 113.9 ± 2.9 | 101.0 ± 2.8 | 101.9 ± 2.3 | 85.2 ± 0.6 | 105.2 ± 2.3 | 0.9991 |
| Ectoine | 111.5 ± 2.6 | 103.0 ± 2.4 | 87.1 ± 1.2 | 87.3 ± 2.7 | 102.7 ± 2.8 | 110.4 ± 2.5 | 0.9999 |
| Erythrose-4-phosphate | 100.6 ± 0.6 | 91.6 ± 3.0 | 103.1 ± 1.9 | 96.3 ± 0.8 | 94.3 ± 1.2 | 94.9 ± 1.2 | 0.9994 |
| FAD | 102.6 ± 1.9 | 101.5 ± 0.7 | 95.7 ± 1.6 | 101.5 ± 3.2 | 104.8 ± 0.5 | 110.4 ± 1.0 | 0.9992 |
| Folic acid | 90.6 ± 0.8 | 93.2 ± 0.8 | 94.8 ± 2.5 | 109.9 ± 0.9 | 85.4 ± 1.8 | 86.7 ± 3.2 | 0.9998 |
| Fructose-6-phosphate | 101.0 ± 0.9 | 88.6 ± 1.3 | 96.2 ± 2.3 | 104.0 ± 3.0 | 107.4 ± 0.9 | 108.1 ± 1.2 | 0.9999 |
| Fumaric Acid | 92.6 ± 0.6 | 106.0 ± 2.7 | 102.0 ± 2.2 | 111.4 ± 0.9 | 103.1 ± 0.9 | 106.0 ± 2.8 | 0.9994 |
| Galactosamine | 113.7 ± 0.6 | 110.3 ± 2.4 | 103.9 ± 1.8 | 108.2 ± 0.6 | 106.2 ± 1.0 | 86.2 ± 0.7 | 0.9993 |
| Galactose-1-phosphate | 114.2 ± 1.1 | 98.4 ± 1.8 | 87.1 ± 0.5 | 98.6 ± 2.4 | 85.9 ± 2.3 | 90.3 ± 2.6 | 0.9994 |
| Galactose-6-Phosphate | 114.8 ± 2.6 | 96.7 ± 2.8 | 87.9 ± 1.5 | 91.2 ± 2.5 | 96.1 ± 2.6 | 98.0 ± 0.3 | 0.9994 |
| gamma-Glutamylcysteine | 113.8 ± 0.6 | 105.4 ± 1.0 | 97.5 ± 2.0 | 99.0 ± 0.6 | 106.7 ± 3.1 | 94.9 ± 2.1 | 0.9998 |
| Glucosamine | 102.6 ± 0.8 | 88.6 ± 2.6 | 112.2 ± 3.0 | 91.4 ± 2.3 | 111.7 ± 1.3 | 95.2 ± 1.0 | 0.9996 |
| Glucosamine-6-Phosphate | 114.3 ± 1.7 | 93.0 ± 1.6 | 110.9 ± 2.9 | 105.8 ± 1.5 | 95.1 ± 2.8 | 114.4 ± 1.5 | 0.9993 |
| Glucose | 112.2 ± 2.9 | 93.8 ± 1.3 | 104.6 ± 2.0 | 95.8 ± 1.4 | 104.2 ± 1.7 | 88.9 ± 2.8 | 0.9993 |
| Glucose-1-phosphate | 107.1 ± 1.6 | 109.2 ± 1.3 | 89.6 ± 0.7 | 85.9 ± 1.1 | 90.8 ± 2.1 | 89.7 ± 1.6 | 0.9991 |
| Glucose-6-Phosphate | 104.8 ± 1.0 | 112.7 ± 1.3 | 110.0 ± 1.4 | 90.3 ± 1.0 | 103.0 ± 0.7 | 103.6 ± 2.0 | 0.9993 |
| Glucuronic acid | 100.3 ± 1.3 | 113.2 ± 0.5 | 101.9 ± 1.6 | 106.8 ± 1.1 | 100.4 ± 2.3 | 106.4 ± 1.3 | 0.9997 |
| Glutamic acid | 112.9 ± 3.2 | 113.0 ± 1.0 | 107.5 ± 1.7 | 113.0 ± 2.8 | 110.9 ± 1.8 | 112.2 ± 2.2 | 0.9996 |
| Glutamine | 86.8 ± 1.4 | 107.9 ± 1.5 | 114.6 ± 3.0 | 89.7 ± 2.0 | 90.2 ± 2.2 | 87.1 ± 2.7 | 0.9995 |
| Glutathione | 92.8 ± 0.6 | 101.0 ± 1.2 | 102.7 ± 1.9 | 90.7 ± 1.4 | 86.0 ± 0.5 | 110.2 ± 0.5 | 0.9997 |
| Glutathionylspermine | 96.6 ± 0.4 | 104.3 ± 1.2 | 105.0 ± 0.6 | 97.5 ± 0.8 | 98.9 ± 2.0 | 107.0 ± 1.2 | 0.9994 |
| Glycerophosphocholine | 85.1 ± 1.6 | 97.6 ± 1.5 | 107.7 ± 1.3 | 94.1 ± 0.4 | 114.1 ± 2.9 | 109.5 ± 3.1 | 0.9993 |
| Glycine | 89.7 ± 1.7 | 112.9 ± 3.0 | 92.3 ± 0.7 | 91.0 ± 1.2 | 87.0 ± 0.3 | 101.3 ± 0.9 | 0.9996 |
| Glycolic acid | 89.4 ± 1.8 | 112.1 ± 3.0 | 102.7 ± 2.2 | 92.0 ± 0.6 | 106.7 ± 2.5 | 99.9 ± 2.1 | 0.9993 |
| Guanidinoacetic acid | 88.7 ± 1.2 | 114.4 ± 0.4 | 88.4 ± 1.1 | 109.6 ± 1.1 | 104.4 ± 1.5 | 92.4 ± 0.5 | 0.9994 |
| Guanidinobutanoic acid | 99.2 ± 2.3 | 101.4 ± 2.4 | 110.9 ± 2.1 | 105.6 ± 1.8 | 95.8 ± 0.8 | 87.6 ± 1.8 | 0.9992 |
| Guanine | 85.1 ± 2.6 | 96.1 ± 1.9 | 108.8 ± 1.9 | 112.6 ± 2.6 | 104.6 ± 3.2 | 104.5 ± 3.0 | 0.9992 |
| Guanosine | 99.4 ± 0.4 | 107.1 ± 1.4 | 94.0 ± 1.5 | 87.7 ± 1.5 | 108.8 ± 1.5 | 90.3 ± 1.6 | 0.9994 |
| Guanosine 5'-diphosphate | 109.2 ± 1.8 | 87.6 ± 0.5 | 111.2 ± 2.9 | 112.9 ± 2.4 | 95.1 ± 1.4 | 90.3 ± 1.6 | 0.9999 |
| Guanosine-5'-monophosphate | 96.7 ± 2.2 | 92.8 ± 0.7 | 87.2 ± 1.4 | 96.1 ± 1.7 | 95.9 ± 0.7 | 108.4 ± 2.5 | 0.9996 |
| Hercynine | 99.5 ± 0.7 | 112.7 ± 2.9 | 87.2 ± 1.3 | 113.5 ± 2.5 | 95.2 ± 0.6 | 109.3 ± 0.5 | 0.9992 |
| Hippuric acid | 110.9 ± 2.1 | 86.5 ± 2.1 | 104.7 ± 2.1 | 103.7 ± 0.6 | 92.0 ± 2.7 | 89.7 ± 3.0 | 0.9997 |
| Histamine | 98.1 ± 2.9 | 86.5 ± 1.5 | 88.7 ± 1.1 | 97.2 ± 2.8 | 105.8 ± 2.6 | 90.5 ± 2.7 | 0.9991 |
| Histidine | 103.8 ± 1.7 | 110.2 ± 1.2 | 98.6 ± 3.1 | 85.4 ± 0.7 | 92.2 ± 1.3 | 94.4 ± 1.9 | 0.9997 |
| Homoalanine | 104.5 ± 2.5 | 108.1 ± 0.6 | 99.8 ± 2.2 | 85.6 ± 2.3 | 87.0 ± 1.5 | 94.8 ± 2.6 | 0.9998 |
| Homoarginine | 110.6 ± 1.4 | 91.9 ± 1.8 | 99.3 ± 0.8 | 111.4 ± 1.9 | 106.3 ± 1.1 | 102.9 ± 1.6 | 0.9993 |
| Homocysteine | 92.1 ± 1.9 | 108.4 ± 1.1 | 96.0 ± 2.4 | 97.5 ± 1.1 | 101.9 ± 1.7 | 96.1 ± 0.8 | 0.9998 |
| Homogentisic acid | 109.9 ± 2.1 | 88.1 ± 2.4 | 108.0 ± 3.0 | 107.5 ± 2.3 | 93.3 ± 2.5 | 86.3 ± 0.9 | 0.9997 |
| Homoleucine | 92.2 ± 3.2 | 88.0 ± 1.2 | 96.7 ± 0.8 | 86.1 ± 1.3 | 90.7 ± 0.5 | 103.1 ± 1.1 | 0.9993 |
| Homovanillic acid | 93.4 ± 2.1 | 107.7 ± 2.7 | 92.6 ± 2.9 | 110.7 ± 2.0 | 86.2 ± 1.5 | 112.0 ± 3.0 | 0.9992 |
| Hydantoin-5-propionate | 111.2 ± 2.0 | 107.0 ± 0.5 | 90.9 ± 2.8 | 94.9 ± 0.4 | 108.8 ± 1.8 | 99.1 ± 2.1 | 0.9996 |
| Hypotaurine | 91.2 ± 1.9 | 96.8 ± 1.6 | 100.7 ± 2.3 | 98.5 ± 0.9 | 89.5 ± 2.5 | 94.7 ± 2.6 | 0.9996 |
| Hypoxanthine | 86.7 ± 2.3 | 99.1 ± 1.0 | 113.0 ± 3.0 | 102.5 ± 2.4 | 111.4 ± 2.9 | 95.3 ± 1.4 | 0.9991 |
| Indole-3-acetamide | 107.8 ± 0.5 | 105.8 ± 0.8 | 92.2 ± 0.8 | 112.9 ± 0.4 | 95.8 ± 0.8 | 108.1 ± 2.5 | 0.9992 |
| Inosine | 94.3 ± 1.5 | 101.1 ± 0.9 | 98.2 ± 2.7 | 96.6 ± 2.1 | 108.7 ± 2.5 | 108.1 ± 3.1 | 0.9995 |
| Inosine 5'-monophosphate | 97.0 ± 0.8 | 86.5 ± 2.6 | 88.4 ± 2.2 | 114.0 ± 0.8 | 95.9 ± 2.0 | 114.0 ± 1.7 | 0.9995 |
| Isocitric Acid | 106.4 ± 1.3 | 103.8 ± 2.3 | 95.9 ± 1.7 | 101.6 ± 0.8 | 114.4 ± 2.2 | 89.2 ± 3.1 | 0.9999 |
| Isoleucine | 102.4 ± 1.5 | 112.5 ± 2.0 | 86.3 ± 1.5 | 102.1 ± 0.6 | 114.5 ± 2.8 | 100.4 ± 1.6 | 0.9994 |
| Lactic Acid | 100.9 ± 1.2 | 107.2 ± 1.4 | 99.6 ± 1.6 | 87.7 ± 2.6 | 88.5 ± 3.1 | 98.3 ± 2.8 | 0.9991 |
| Leucine | 91.4 ± 1.3 | 113.0 ± 0.7 | 113.4 ± 1.9 | 86.9 ± 2.5 | 106.6 ± 2.2 | 107.5 ± 0.4 | 0.9992 |
| Levulinic acid | 93.5 ± 1.3 | 95.2 ± 2.9 | 98.3 ± 2.8 | 99.7 ± 2.5 | 111.3 ± 1.0 | 97.6 ± 3.2 | 0.9994 |
| Lysine | 100.0 ± 2.2 | 104.9 ± 1.3 | 99.9 ± 2.1 | 102.8 ± 1.4 | 114.4 ± 1.8 | 104.9 ± 1.5 | 0.9997 |
| Malic Acid | 107.3 ± 1.2 | 96.2 ± 2.1 | 113.2 ± 2.4 | 90.6 ± 0.8 | 91.9 ± 2.1 | 107.6 ± 1.4 | 0.9992 |
| Mannose-1 with Mannose-6-Phosphate | 114.9 ± 3.1 | 87.1 ± 0.4 | 87.5 ± 0.4 | 92.6 ± 3.0 | 100.9 ± 2.8 | 114.0 ± 1.7 | 0.9991 |
| Mesaconic acid | 91.8 ± 2.7 | 89.7 ± 0.9 | 104.3 ± 0.4 | 112.7 ± 1.6 | 104.1 ± 0.9 | 94.7 ± 2.5 | 0.9995 |
| Methionine | 103.2 ± 2.1 | 98.8 ± 3.0 | 110.9 ± 2.3 | 98.6 ± 0.5 | 103.2 ± 0.6 | 85.3 ± 0.6 | 0.9996 |
| Methionine sulfoxide | 88.4 ± 1.9 | 104.8 ± 0.8 | 114.0 ± 2.6 | 105.7 ± 2.2 | 85.3 ± 2.6 | 92.7 ± 2.2 | 0.9995 |
| Methylmalonic Acid | 110.0 ± 1.9 | 92.0 ± 0.5 | 109.4 ± 2.9 | 99.8 ± 0.4 | 99.8 ± 2.3 | 99.5 ± 1.4 | 0.9997 |
| N1-Acetylspermidine | 104.5 ± 1.7 | 106.1 ± 1.5 | 93.4 ± 1.4 | 88.9 ± 1.3 | 99.2 ± 1.5 | 98.4 ± 2.3 | 0.9998 |
| N1-Acetylspermine | 103.1 ± 1.8 | 87.4 ± 2.6 | 97.2 ± 2.1 | 113.1 ± 1.5 | 114.8 ± 2.8 | 94.9 ± 0.6 | 0.9994 |
| N8-Acetylspermidine | 107.0 ± 1.8 | 96.0 ± 2.6 | 113.7 ± 1.6 | 102.0 ± 3.2 | 95.2 ± 2.3 | 93.6 ± 1.5 | 0.9991 |
| N-Acetyl-Aspartyl-Glutamate | 104.2 ± 1.7 | 107.1 ± 2.4 | 87.7 ± 2.9 | 96.3 ± 0.6 | 90.8 ± 1.4 | 100.8 ± 0.9 | 0.9999 |
| N-Acetyl-Aspartyl-Glutamyl-Glutamate | 95.3 ± 0.8 | 103.9 ± 2.6 | 87.6 ± 0.4 | 97.4 ± 1.0 | 101.1 ± 2.8 | 91.7 ± 1.6 | 0.9995 |
| N-Acetyl-Galactosamine | 91.7 ± 0.3 | 85.8 ± 1.7 | 94.5 ± 1.9 | 103.9 ± 3.1 | 93.9 ± 2.6 | 104.8 ± 2.7 | 0.9991 |
| N-Acetyl-Leucine | 96.8 ± 2.7 | 100.1 ± 0.4 | 111.2 ± 2.8 | 93.4 ± 2.0 | 112.2 ± 1.3 | 109.1 ± 1.8 | 0.9999 |
| N-Acetyl-Methionine | 113.2 ± 0.7 | 102.9 ± 3.2 | 103.8 ± 1.8 | 96.1 ± 3.0 | 105.3 ± 2.3 | 85.8 ± 2.1 | 0.9995 |
| N-Acetylneuraminic acid | 113.1 ± 2.5 | 109.5 ± 2.9 | 97.3 ± 2.9 | 109.4 ± 3.1 | 98.8 ± 0.5 | 94.2 ± 2.5 | 0.9998 |
| N-Acetyl-Proline | 97.9 ± 1.8 | 112.0 ± 2.7 | 103.6 ± 1.5 | 113.2 ± 3.2 | 104.2 ± 2.7 | 94.2 ± 3.0 | 0.9999 |
| N-Acetyl-Putrescine | 96.9 ± 0.5 | 105.5 ± 1.6 | 93.7 ± 1.7 | 94.5 ± 1.9 | 102.5 ± 3.0 | 109.9 ± 2.0 | 0.9997 |
| N-Acetyl-Serine | 85.8 ± 0.5 | 87.6 ± 1.3 | 110.9 ± 1.2 | 91.8 ± 3.1 | 104.1 ± 2.7 | 93.3 ± 2.1 | 0.9997 |
| N-Acetyl-Serotonin | 109.7 ± 2.7 | 106.9 ± 0.3 | 91.4 ± 2.3 | 94.4 ± 3.0 | 96.7 ± 1.8 | 85.9 ± 0.6 | 0.9998 |
| N-Acyl-Homoserine-lactone | 85.0 ± 1.1 | 100.3 ± 1.4 | 101.6 ± 2.9 | 108.4 ± 2.6 | 109.6 ± 2.5 | 96.1 ± 1.9 | 0.9992 |
| NAD | 93.4 ± 2.2 | 89.4 ± 0.7 | 89.4 ± 3.0 | 98.0 ± 1.4 | 91.0 ± 0.4 | 86.1 ± 1.2 | 0.9991 |
| NADH | 88.4 ± 0.9 | 113.2 ± 0.6 | 92.3 ± 2.3 | 102.2 ± 1.4 | 113.4 ± 1.1 | 85.8 ± 1.9 | 0.9997 |
| NADP | 101.6 ± 1.5 | 107.5 ± 0.9 | 101.0 ± 0.8 | 96.1 ± 3.1 | 110.6 ± 1.3 | 101.7 ± 2.0 | 0.9995 |
| NADPH | 89.7 ± 2.4 | 99.5 ± 1.0 | 90.9 ± 1.3 | 105.2 ± 1.2 | 99.8 ± 1.6 | 100.3 ± 2.1 | 0.9998 |
| N-alpha-Acetyl-lysine | 109.4 ± 1.5 | 103.8 ± 1.8 | 100.8 ± 1.7 | 102.1 ± 0.4 | 109.2 ± 1.6 | 101.3 ± 3.0 | 0.9996 |
| N-alpha-Acetyl-Ornithine | 89.8 ± 0.9 | 96.5 ± 2.7 | 110.5 ± 2.8 | 91.1 ± 1.5 | 106.9 ± 0.7 | 109.8 ± 0.5 | 0.9993 |
| N-Carbamoyl-Proline | 110.1 ± 1.4 | 92.5 ± 0.8 | 93.0 ± 1.9 | 89.6 ± 2.1 | 114.2 ± 2.4 | 101.0 ± 3.2 | 0.9998 |
| N-Carbamoyl-Sarcosine | 88.7 ± 2.5 | 113.4 ± 1.2 | 88.2 ± 0.7 | 108.8 ± 2.4 | 106.7 ± 2.2 | 110.4 ± 2.1 | 0.9997 |
| N-Epsilon-Acetyl-lysine | 96.6 ± 1.9 | 86.2 ± 2.8 | 106.7 ± 2.7 | 113.3 ± 2.2 | 112.3 ± 0.9 | 100.4 ± 2.0 | 0.9993 |
| Nicotinamide | 90.4 ± 0.3 | 97.7 ± 2.7 | 99.6 ± 2.4 | 100.1 ± 2.8 | 108.5 ± 0.6 | 89.4 ± 2.1 | 0.9993 |
| N-Methyl-Aspartic Acid | 102.1 ± 2.0 | 92.2 ± 1.7 | 99.2 ± 1.5 | 91.8 ± 0.8 | 91.1 ± 0.3 | 103.9 ± 3.1 | 0.9992 |
| O-Phosphoethanolamine | 110.3 ± 0.6 | 95.5 ± 3.0 | 110.1 ± 0.8 | 88.6 ± 3.0 | 101.5 ± 1.7 | 115.0 ± 2.2 | 0.9995 |
| Ophthalmic acid | 87.1 ± 0.6 | 103.5 ± 2.2 | 114.5 ± 0.8 | 85.4 ± 2.6 | 94.8 ± 2.1 | 108.3 ± 2.0 | 0.9994 |
| Ornithine | 108.9 ± 2.4 | 94.7 ± 1.6 | 110.5 ± 2.3 | 102.4 ± 0.6 | 111.8 ± 0.5 | 105.8 ± 1.0 | 0.9998 |
| Orotidine 5'-monophosphate | 100.2 ± 0.8 | 104.1 ± 2.5 | 89.0 ± 3.0 | 93.9 ± 2.7 | 86.7 ± 1.7 | 112.4 ± 2.9 | 0.9999 |
| Oxaloacetate | 105.1 ± 0.4 | 97.4 ± 1.3 | 110.3 ± 1.5 | 96.4 ± 2.5 | 114.1 ± 2.4 | 111.6 ± 2.0 | 0.9998 |
| Pantothenic acid | 93.4 ± 2.4 | 92.9 ± 1.2 | 103.3 ± 1.1 | 94.3 ± 1.9 | 85.4 ± 2.1 | 110.7 ± 0.4 | 0.9993 |
| Phenylalanine | 106.8 ± 2.9 | 100.8 ± 0.3 | 113.6 ± 3.1 | 109.5 ± 2.2 | 90.3 ± 0.8 | 87.2 ± 1.3 | 0.9996 |
| Phenylpyruvate | 97.9 ± 1.6 | 86.4 ± 3.0 | 105.5 ± 3.1 | 88.5 ± 0.6 | 104.1 ± 1.2 | 107.8 ± 0.3 | 0.9995 |
| Phosphodimethylethanolamine | 86.6 ± 1.2 | 95.9 ± 1.0 | 88.8 ± 2.3 | 102.2 ± 2.9 | 98.3 ± 2.0 | 107.9 ± 0.6 | 0.9994 |
| Phosphoenolpyruvate | 106.5 ± 1.9 | 92.7 ± 0.5 | 91.2 ± 1.0 | 112.1 ± 0.4 | 99.7 ± 1.3 | 111.2 ± 2.5 | 0.9995 |
| Phosphorylcholine | 98.3 ± 1.7 | 113.0 ± 1.1 | 110.1 ± 1.5 | 105.2 ± 1.5 | 96.3 ± 3.2 | 91.8 ± 0.8 | 0.9996 |
| Pipecolic acid | 94.0 ± 1.9 | 110.8 ± 1.5 | 104.8 ± 2.6 | 101.2 ± 1.4 | 85.5 ± 1.4 | 86.7 ± 0.4 | 0.9993 |
| Proline | 89.9 ± 1.5 | 114.8 ± 2.3 | 103.0 ± 2.7 | 102.4 ± 2.0 | 92.1 ± 2.7 | 100.9 ± 2.8 | 0.9998 |
| Putrescine | 106.9 ± 0.4 | 100.3 ± 1.6 | 114.3 ± 3.1 | 107.0 ± 0.8 | 91.6 ± 1.9 | 96.7 ± 2.3 | 0.9998 |
| Pyridoxal | 88.7 ± 0.8 | 110.4 ± 1.9 | 100.4 ± 2.6 | 92.3 ± 0.8 | 96.5 ± 1.3 | 93.5 ± 0.7 | 0.9996 |
| Pyridoxamine | 102.7 ± 2.3 | 103.2 ± 1.8 | 98.0 ± 1.9 | 103.5 ± 1.8 | 111.4 ± 1.0 | 95.8 ± 2.8 | 0.9992 |
| Pyridoxine | 112.4 ± 3.2 | 96.5 ± 2.1 | 92.4 ± 2.0 | 85.7 ± 2.8 | 99.3 ± 1.1 | 114.1 ± 2.9 | 0.9996 |
| Pyroglutamic acid | 112.0 ± 2.6 | 107.6 ± 2.1 | 103.5 ± 0.5 | 87.6 ± 2.9 | 105.1 ± 3.2 | 91.1 ± 2.0 | 0.9998 |
| Pyrrolysine | 97.9 ± 1.0 | 98.7 ± 1.2 | 103.6 ± 3.1 | 89.0 ± 1.6 | 85.4 ± 1.0 | 98.1 ± 2.4 | 0.9996 |
| Pyruvic Acid | 100.9 ± 2.5 | 98.0 ± 0.8 | 93.5 ± 1.2 | 110.0 ± 0.5 | 85.4 ± 1.8 | 92.8 ± 1.7 | 0.9999 |
| Riboflavin | 100.6 ± 2.2 | 114.1 ± 0.8 | 108.8 ± 2.5 | 103.0 ± 1.1 | 108.2 ± 1.3 | 111.3 ± 3.0 | 0.9995 |
| Ribose-5-Phosphate | 88.0 ± 3.0 | 101.6 ± 1.5 | 107.9 ± 1.5 | 90.4 ± 0.7 | 94.0 ± 1.0 | 94.8 ± 0.4 | 0.9992 |
| Ribulose-1-Phosphate | 86.7 ± 0.9 | 85.8 ± 1.1 | 91.1 ± 2.3 | 109.8 ± 1.8 | 101.5 ± 0.6 | 103.7 ± 2.6 | 0.9993 |
| Ribulose-5-Phosphate | 90.1 ± 0.7 | 86.6 ± 3.1 | 85.1 ± 1.8 | 112.1 ± 1.3 | 99.5 ± 2.7 | 87.5 ± 1.0 | 0.9995 |
| Saccharopine | 112.2 ± 2.4 | 99.8 ± 0.7 | 96.5 ± 1.6 | 114.4 ± 1.2 | 106.1 ± 0.7 | 113.1 ± 1.8 | 0.9998 |
| S-Adenosylhomocysteine | 107.5 ± 0.4 | 94.0 ± 2.4 | 96.1 ± 0.7 | 92.1 ± 0.4 | 95.8 ± 1.9 | 109.8 ± 0.7 | 0.9998 |
| S-Adenosylmethioninamine | 101.9 ± 1.2 | 86.5 ± 1.9 | 105.9 ± 3.2 | 102.6 ± 0.8 | 96.5 ± 2.3 | 90.8 ± 2.8 | 0.9993 |
| S-Adenosylmethionine | 111.4 ± 2.4 | 88.8 ± 1.5 | 102.9 ± 2.5 | 91.5 ± 1.8 | 96.3 ± 2.1 | 104.1 ± 0.5 | 0.9996 |
| Sarcosine | 98.7 ± 2.9 | 104.6 ± 1.3 | 114.8 ± 1.9 | 109.3 ± 1.5 | 91.8 ± 2.0 | 105.1 ± 1.9 | 0.9995 |
| Sedoheptulose-7-phosphate | 86.5 ± 2.7 | 98.9 ± 1.4 | 114.9 ± 0.8 | 111.1 ± 0.9 | 91.4 ± 3.0 | 86.2 ± 1.8 | 0.9998 |
| Selenohomocysteine | 102.8 ± 2.8 | 111.3 ± 2.2 | 89.2 ± 3.0 | 90.3 ± 0.3 | 86.5 ± 2.1 | 89.9 ± 0.5 | 0.9998 |
| Selenomethionine | 107.2 ± 0.8 | 87.6 ± 1.1 | 100.0 ± 2.3 | 101.1 ± 1.1 | 96.5 ± 2.7 | 90.3 ± 2.4 | 0.9993 |
| Serine | 103.5 ± 0.5 | 92.3 ± 1.2 | 115.0 ± 2.2 | 86.0 ± 0.4 | 104.7 ± 2.8 | 106.0 ± 2.6 | 0.9997 |
| S-Methyl-Thioinosine | 102.5 ± 2.3 | 94.1 ± 1.3 | 87.1 ± 2.4 | 102.1 ± 1.6 | 107.1 ± 1.4 | 113.0 ± 1.5 | 0.9996 |
| Spermidine | 101.9 ± 2.3 | 91.9 ± 0.9 | 104.0 ± 0.5 | 94.5 ± 3.1 | 114.7 ± 2.6 | 108.5 ± 1.2 | 0.9993 |
| Spermine | 102.6 ± 2.7 | 108.2 ± 1.0 | 89.9 ± 1.1 | 105.6 ± 1.5 | 112.3 ± 0.4 | 95.5 ± 2.2 | 0.9992 |
| Succinate | 88.1 ± 0.6 | 111.3 ± 1.2 | 108.8 ± 2.2 | 87.9 ± 2.4 | 107.8 ± 0.7 | 85.7 ± 1.1 | 0.9995 |
| Sucrose | 112.1 ± 2.7 | 100.9 ± 1.4 | 98.7 ± 2.1 | 102.3 ± 0.6 | 107.2 ± 0.9 | 95.0 ± 2.3 | 0.9998 |
| Symmetric dimethylarginine | 90.4 ± 2.9 | 114.6 ± 1.8 | 96.5 ± 2.0 | 94.3 ± 0.3 | 105.4 ± 2.6 | 88.9 ± 1.6 | 0.9998 |
| Taurine | 104.9 ± 2.2 | 91.0 ± 2.5 | 101.5 ± 1.5 | 108.0 ± 2.6 | 108.7 ± 3.2 | 98.3 ± 2.5 | 0.9992 |
| Tetrahydroneopterin | 96.4 ± 1.3 | 106.9 ± 2.9 | 96.3 ± 0.7 | 93.8 ± 1.4 | 96.6 ± 1.2 | 88.1 ± 1.0 | 0.9996 |
| Thiamine | 98.4 ± 0.8 | 102.3 ± 1.5 | 100.3 ± 2.9 | 85.1 ± 1.2 | 96.8 ± 1.5 | 103.2 ± 1.3 | 0.9991 |
| Thiamine monophosphate | 106.1 ± 2.8 | 86.7 ± 1.5 | 109.5 ± 1.4 | 110.7 ± 0.7 | 87.7 ± 3.0 | 90.4 ± 2.8 | 0.9994 |
| Threonic acid | 95.0 ± 2.5 | 97.6 ± 2.1 | 90.3 ± 2.7 | 105.7 ± 0.5 | 89.3 ± 2.5 | 98.7 ± 2.3 | 0.9997 |
| Threonine | 90.8 ± 1.5 | 89.5 ± 2.3 | 85.6 ± 1.4 | 86.4 ± 1.9 | 112.9 ± 0.5 | 87.1 ± 2.3 | 0.9998 |
| Thymidine | 86.3 ± 2.9 | 97.7 ± 1.7 | 92.6 ± 2.7 | 95.8 ± 2.4 | 97.1 ± 2.7 | 91.9 ± 1.1 | 0.9994 |
| Thymidine Monophosphate | 110.4 ± 0.8 | 102.0 ± 1.6 | 113.1 ± 3.1 | 105.6 ± 0.5 | 110.8 ± 1.7 | 106.2 ± 0.5 | 0.9994 |
| Thymine | 93.7 ± 2.2 | 105.9 ± 0.8 | 92.5 ± 1.3 | 109.2 ± 2.5 | 110.1 ± 1.8 | 100.3 ± 2.9 | 0.9999 |
| Trigonelleine | 109.7 ± 1.9 | 98.9 ± 2.9 | 101.0 ± 1.0 | 105.3 ± 0.9 | 89.1 ± 0.6 | 93.7 ± 1.7 | 0.9998 |
| Trimethylamine N-oxide | 112.6 ± 0.4 | 96.6 ± 2.0 | 111.7 ± 0.7 | 92.2 ± 2.8 | 102.0 ± 2.4 | 85.5 ± 3.1 | 0.9994 |
| Trimethyllysine | 88.4 ± 0.4 | 99.1 ± 1.1 | 90.3 ± 1.8 | 91.2 ± 0.3 | 95.2 ± 1.4 | 100.9 ± 2.1 | 0.9994 |
| Tryptophan | 113.8 ± 0.6 | 91.2 ± 0.9 | 96.4 ± 1.9 | 101.2 ± 0.3 | 98.2 ± 2.4 | 96.9 ± 2.3 | 0.9992 |
| Tyramine | 100.2 ± 1.9 | 106.7 ± 2.5 | 97.3 ± 1.5 | 108.5 ± 0.6 | 90.2 ± 0.9 | 109.1 ± 2.4 | 0.9997 |
| Tyrosine | 99.9 ± 1.0 | 102.1 ± 2.2 | 101.2 ± 3.2 | 90.9 ± 3.1 | 97.4 ± 1.1 | 102.6 ± 1.0 | 0.9993 |
| Uracil | 107.9 ± 1.2 | 90.8 ± 1.5 | 106.1 ± 2.9 | 94.6 ± 1.3 | 99.1 ± 1.7 | 98.0 ± 2.4 | 0.9999 |
| Uridine 5'-diphosphate | 108.2 ± 1.1 | 90.1 ± 3.1 | 97.3 ± 1.6 | 97.7 ± 1.7 | 100.3 ± 2.0 | 102.6 ± 0.4 | 0.9993 |
| Uridine Monophosphate | 114.4 ± 1.1 | 99.0 ± 1.7 | 85.6 ± 2.6 | 110.8 ± 2.2 | 88.3 ± 2.9 | 93.9 ± 2.9 | 0.9994 |
| Uridine-5-Diphosphogalactose | 101.9 ± 2.9 | 109.2 ± 2.8 | 97.4 ± 3.1 | 98.0 ± 1.5 | 105.9 ± 2.2 | 93.6 ± 1.5 | 0.9997 |
| Urocanic acid | 86.6 ± 2.7 | 99.8 ± 2.4 | 113.9 ± 1.1 | 98.1 ± 1.5 | 114.7 ± 0.3 | 88.3 ± 1.9 | 0.9995 |
| UTP | 98.1 ± 1.7 | 108.6 ± 2.6 | 90.0 ± 0.5 | 96.9 ± 1.1 | 107.3 ± 2.9 | 96.3 ± 1.6 | 0.9994 |
| Valine | 94.9 ± 3.0 | 88.7 ± 2.7 | 110.4 ± 1.1 | 99.9 ± 1.2 | 97.7 ± 3.0 | 113.3 ± 1.6 | 0.9994 |
| XMP | 100.7 ± 2.8 | 91.5 ± 0.8 | 87.5 ± 1.6 | 105.3 ± 1.9 | 102.5 ± 1.9 | 114.0 ± 2.6 | 0.9998 |
| Xylulose-5-Phosphate | 85.9 ± 2.5 | 91.9 ± 1.6 | 109.3 ± 1.9 | 112.0 ± 2.8 | 85.9 ± 0.9 | 114.3 ± 1.9 | 0.9995 |

### **Table S12** | Intra – day precision (as % RSD) (Colorectal organoid extract as matrix, n = 5). The concentrations of QC_LLOQ_, QC_Low_, QC_Mid_ and QC_High_ were 10 ng/mL, 25 ng/mL, 100 ng/mL and 500 ng/mL respectively.

| Precision, % | QC_LLOQ_ | QC_low_ | QC_mid_ | QC_high_ |
| --- | --- | --- | --- | --- |
| 1-Methylguanine | 5.3 ± 0.3 | 2.8 ± 0.5 | 7.1 ± 0.4 | 6.8 ± 0.3 |
| 1-Methylhistidine | 10.7 ± 0.8 | 4.5 ± 1.0 | 5.2 ± 0.6 | 2.0 ± 0.4 |
| 1-Methylnicotinamide | 5.0 ± 0.8 | 5.3 ± 0.5 | 1.7 ± 0.3 | 7.0 ± 1.0 |
| 2-Aminoadipic Acid | 6.7 ± 0.9 | 7.6 ± 0.7 | 6.9 ± 1.0 | 1.8 ± 0.2 |
| 2-Deoxyadenosine | 3.0 ± 0.4 | 6.9 ± 0.7 | 4.4 ± 0.5 | 5.5 ± 0.3 |
| 2'-Deoxyadenosine 5'-monophosphate | 4.7 ± 0.9 | 7.8 ± 0.3 | 5.4 ± 0.4 | 3.9 ± 0.9 |
| 2'-deoxyadenosine 5'-triphosphate | 11.8 ± 1.0 | 2.0 ± 0.3 | 1.8 ± 0.2 | 5.9 ± 0.3 |
| 2'-Deoxycytidine-5'-monophosphate | 9.0 ± 0.5 | 3.0 ± 0.6 | 3.7 ± 0.8 | 7.7 ± 0.5 |
| 2-Deoxyguanosine | 8.1 ± 0.3 | 3.5 ± 0.4 | 1.6 ± 0.3 | 5.7 ± 0.8 |
| 2-hydroxybutyric acid | 11.6 ± 0.3 | 1.8 ± 0.4 | 4.4 ± 0.3 | 6.5 ± 0.5 |
| 2-hydroxyglutaric acid | 11.3 ± 1.0 | 2.8 ± 0.2 | 2.8 ± 0.5 | 7.8 ± 0.5 |
| 2-Hydroxyisocaproic acid | 8.1 ± 0.6 | 1.7 ± 0.3 | 6.7 ± 0.6 | 5.8 ± 0.6 |
| 2-ketobutyric acid | 6.0 ± 0.4 | 5.8 ± 0.7 | 5.3 ± 0.6 | 2.1 ± 0.4 |
| 2-oxoadipic acid | 12.8 ± 0.9 | 4.2 ± 0.9 | 7.1 ± 0.8 | 5.9 ± 0.9 |
| 2-Phenylglycine | 5.7 ± 0.5 | 6.5 ± 0.2 | 6.4 ± 0.8 | 1.8 ± 0.4 |
| 2-phosphoglycerate | 5.2 ± 0.6 | 7.2 ± 0.4 | 2.7 ± 0.6 | 1.5 ± 0.4 |
| 3-hydroxy-3-methyl-2-oxobutanoate | 4.6 ± 0.5 | 4.8 ± 0.9 | 4.7 ± 0.4 | 4.6 ± 1.0 |
| 3-hydroxybutyric acid | 12.1 ± 0.3 | 6.8 ± 0.9 | 6.1 ± 1.0 | 6.6 ± 0.4 |
| 3-Methylhistidine | 12.5 ± 0.6 | 2.5 ± 0.5 | 7.7 ± 0.4 | 7.2 ± 0.7 |
| 3-Phosphoglyceraldehyde | 3.5 ± 0.3 | 5.3 ± 0.5 | 2.0 ± 0.3 | 4.0 ± 0.7 |
| 3-phosphoglycerate | 5.8 ± 0.6 | 1.7 ± 0.2 | 2.8 ± 0.4 | 3.0 ± 0.3 |
| 4-amino-butyric acid | 3.4 ± 0.7 | 1.9 ± 0.4 | 7.4 ± 0.2 | 4.6 ± 0.8 |
| 4-aspartyl-phosphate | 3.7 ± 0.8 | 3.6 ± 0.3 | 4.5 ± 0.3 | 7.9 ± 0.4 |
| 4-Hydroxybenzoic acid | 6.3 ± 0.5 | 5.2 ± 0.9 | 5.2 ± 0.7 | 4.4 ± 0.3 |
| 4-Hydroxyphenylacetylglycine | 4.6 ± 0.8 | 2.7 ± 0.3 | 5.5 ± 1.0 | 3.5 ± 0.9 |
| 4-methyl-2-oxopentanoate | 6.5 ± 0.5 | 6.3 ± 0.5 | 6.2 ± 0.3 | 7.5 ± 0.8 |
| 4-Oxoglutaramate | 11.9 ± 0.6 | 5.6 ± 0.2 | 4.9 ± 0.9 | 4.7 ± 0.7 |
| 5-Hydroxyindolepyruvate | 5.9 ± 1.0 | 6.4 ± 0.5 | 2.5 ± 0.4 | 5.2 ± 0.9 |
| 5-Hydroxylysine | 13.0 ± 0.6 | 6.4 ± 0.8 | 6.3 ± 0.9 | 3.9 ± 0.3 |
| 5-Hydroxy-Tryptophan | 11.7 ± 0.3 | 2.4 ± 0.3 | 2.1 ± 0.3 | 5.3 ± 0.8 |
| 5-Phosphoribosylamine | 9.7 ± 0.2 | 7.6 ± 0.6 | 4.9 ± 0.2 | 7.1 ± 0.5 |
| 6-phosphogluconate | 11.9 ± 0.8 | 6.6 ± 0.8 | 4.9 ± 0.4 | 3.1 ± 0.5 |
| Acetylcarnitine | 4.7 ± 0.4 | 5.0 ± 0.5 | 5.4 ± 0.7 | 4.2 ± 0.9 |
| Aconitic Acid | 7.0 ± 0.7 | 3.6 ± 0.3 | 2.7 ± 0.4 | 6.1 ± 0.2 |
| Adenine | 12.9 ± 0.9 | 4.3 ± 1.0 | 7.3 ± 0.7 | 7.0 ± 0.5 |
| Adenosine | 9.2 ± 0.7 | 5.7 ± 0.4 | 1.5 ± 0.3 | 2.3 ± 0.6 |
| Adenosine 3',5'-cyclic monophosphate | 8.3 ± 0.8 | 7.7 ± 1.0 | 6.3 ± 0.4 | 5.8 ± 0.4 |
| Adenosine Monophosphate | 9.4 ± 0.4 | 1.8 ± 0.3 | 5.7 ± 0.7 | 6.8 ± 0.5 |
| Adenosine-5'-diphosphate | 8.0 ± 0.9 | 3.7 ± 0.3 | 3.5 ± 0.7 | 3.4 ± 0.3 |
| Adenosine-5'-triphosphate | 12.9 ± 0.3 | 7.5 ± 0.5 | 2.0 ± 0.3 | 5.8 ± 0.8 |
| Adenylosuccinate | 11.8 ± 0.7 | 5.0 ± 0.8 | 2.8 ± 0.5 | 5.2 ± 0.8 |
| Alanine | 6.0 ± 1.0 | 7.0 ± 0.9 | 2.2 ± 0.5 | 6.5 ± 0.6 |
| alpha-ketoglutaric acid | 9.8 ± 0.8 | 6.7 ± 0.4 | 4.7 ± 1.0 | 2.5 ± 0.6 |
| Anserine | 6.1 ± 0.5 | 3.7 ± 0.2 | 5.0 ± 0.6 | 4.1 ± 0.5 |
| Arginine | 6.9 ± 0.7 | 3.7 ± 0.9 | 1.7 ± 0.4 | 6.5 ± 0.5 |
| Argininosuccinic Acid | 5.8 ± 0.5 | 1.6 ± 0.2 | 7.9 ± 1.0 | 2.1 ± 0.3 |
| Asparagine | 10.5 ± 0.6 | 3.8 ± 0.8 | 5.0 ± 0.8 | 7.9 ± 0.7 |
| Aspartic acid | 10.1 ± 0.2 | 1.5 ± 0.3 | 6.5 ± 0.5 | 2.1 ± 0.3 |
| Asymmetric dimethylarginine | 11.1 ± 0.4 | 5.9 ± 0.7 | 4.6 ± 0.6 | 7.3 ± 0.7 |
| Azelaic acid | 6.9 ± 0.5 | 7.0 ± 1.0 | 4.8 ± 0.3 | 1.8 ± 0.4 |
| beta-Alanine | 6.4 ± 0.8 | 1.5 ± 0.2 | 2.3 ± 0.5 | 6.9 ± 0.3 |
| Betaine | 7.9 ± 0.3 | 4.8 ± 0.4 | 3.6 ± 0.8 | 5.3 ± 0.9 |
| Biopterin | 7.3 ± 0.4 | 4.2 ± 0.5 | 2.2 ± 0.4 | 4.6 ± 0.3 |
| Biotin | 4.0 ± 0.4 | 2.0 ± 0.4 | 7.5 ± 0.7 | 4.7 ± 0.5 |
| Carnitine | 7.9 ± 0.8 | 7.0 ± 0.5 | 3.3 ± 0.5 | 5.9 ± 0.3 |
| Carnosine | 6.9 ± 0.5 | 2.6 ± 0.4 | 5.8 ± 0.8 | 4.0 ± 0.3 |
| CDP-choline | 8.4 ± 0.5 | 4.1 ± 0.6 | 7.6 ± 0.7 | 5.4 ± 0.3 |
| cGMP | 6.8 ± 1.0 | 2.2 ± 0.2 | 6.9 ± 0.5 | 4.2 ± 0.3 |
| Choline | 4.3 ± 0.3 | 7.2 ± 0.6 | 7.5 ± 0.5 | 7.4 ± 0.7 |
| Cinnamic acid | 8.9 ± 0.9 | 2.4 ± 0.5 | 7.1 ± 0.9 | 5.7 ± 0.8 |
| Citraconic acid | 7.0 ± 0.6 | 2.6 ± 0.6 | 5.1 ± 0.8 | 2.9 ± 0.3 |
| Citramalic acid | 12.0 ± 0.8 | 4.3 ± 1.0 | 4.1 ± 0.3 | 6.9 ± 0.7 |
| Citric Acid | 8.5 ± 0.8 | 7.9 ± 1.0 | 2.8 ± 0.5 | 3.1 ± 0.8 |
| Citrulline | 8.7 ± 0.8 | 6.0 ± 0.7 | 4.4 ± 0.7 | 4.6 ± 0.9 |
| Creatine | 5.4 ± 0.5 | 3.8 ± 0.3 | 5.7 ± 0.6 | 3.7 ± 0.7 |
| Creatinine | 6.5 ± 0.6 | 4.0 ± 0.8 | 2.8 ± 0.2 | 6.7 ± 0.3 |
| Cystathionine | 11.2 ± 0.4 | 1.5 ± 0.2 | 4.7 ± 0.4 | 7.8 ± 0.7 |
| Cysteamine | 3.6 ± 0.4 | 7.4 ± 0.5 | 3.4 ± 0.5 | 3.2 ± 0.5 |
| Cysteine | 3.3 ± 0.6 | 7.2 ± 0.5 | 6.0 ± 0.8 | 5.6 ± 0.8 |
| Cysteinylglycine | 11.9 ± 0.5 | 5.5 ± 0.3 | 6.3 ± 0.7 | 8.0 ± 0.4 |
| Cystine | 8.0 ± 0.9 | 5.8 ± 0.7 | 5.2 ± 0.3 | 2.5 ± 0.6 |
| Cytidine | 10.1 ± 0.5 | 5.9 ± 0.3 | 4.6 ± 0.3 | 7.1 ± 0.4 |
| Cytidine 3',5'-cyclic monophosphate | 3.5 ± 0.6 | 6.3 ± 0.3 | 4.4 ± 0.6 | 6.9 ± 0.7 |
| Cytidine 5'-diphosphate | 4.0 ± 1.0 | 7.9 ± 0.2 | 6.9 ± 0.8 | 4.7 ± 0.2 |
| Cytidine 5'-triphosphate | 12.1 ± 0.7 | 1.6 ± 0.2 | 7.1 ± 1.0 | 7.9 ± 0.5 |
| Cytidine monophosphate | 3.1 ± 0.2 | 4.6 ± 0.7 | 4.0 ± 0.5 | 8.0 ± 0.7 |
| Cytosine | 8.9 ± 0.3 | 5.7 ± 0.5 | 5.4 ± 0.6 | 5.6 ± 0.2 |
| dADP | 9.0 ± 0.6 | 5.0 ± 0.6 | 2.3 ± 0.4 | 7.5 ± 1.0 |
| Diethanolamine | 4.7 ± 0.9 | 4.8 ± 0.6 | 1.6 ± 0.4 | 3.9 ± 0.8 |
| Dihydrobiopterin | 7.2 ± 0.3 | 2.7 ± 0.2 | 3.6 ± 0.6 | 4.9 ± 0.9 |
| Dihydroneopterin triphosphate | 12.2 ± 0.3 | 2.0 ± 0.4 | 1.7 ± 0.2 | 5.5 ± 0.8 |
| Dihydroxyacetonephosphate | 6.8 ± 0.7 | 5.8 ± 0.9 | 7.7 ± 1.0 | 1.7 ± 0.2 |
| Dimethylglycine | 6.9 ± 0.5 | 7.6 ± 0.8 | 4.4 ± 1.0 | 2.4 ± 0.5 |
| Ectoine | 7.2 ± 0.4 | 2.2 ± 0.5 | 6.8 ± 0.8 | 2.3 ± 0.4 |
| Erythrose-4-phosphate | 12.7 ± 0.9 | 3.2 ± 0.3 | 6.8 ± 0.4 | 7.8 ± 0.9 |
| FAD | 3.5 ± 0.4 | 6.7 ± 1.0 | 6.2 ± 0.9 | 1.6 ± 0.4 |
| Folic acid | 8.9 ± 0.9 | 3.2 ± 0.6 | 4.3 ± 0.8 | 3.9 ± 0.7 |
| Fructose-6-phosphate | 12.2 ± 0.3 | 1.7 ± 0.3 | 7.3 ± 0.4 | 1.9 ± 0.2 |
| Fumaric Acid | 6.8 ± 0.8 | 2.9 ± 0.3 | 2.4 ± 0.6 | 2.1 ± 0.3 |
| Galactosamine | 9.9 ± 0.2 | 5.4 ± 0.6 | 5.9 ± 0.3 | 4.8 ± 0.6 |
| Galactose-1-phosphate | 8.0 ± 0.7 | 7.4 ± 0.2 | 5.8 ± 0.3 | 7.2 ± 0.2 |
| Galactose-6-Phosphate | 7.9 ± 0.9 | 5.2 ± 0.6 | 5.2 ± 0.3 | 5.3 ± 0.8 |
| gamma-Glutamylcysteine | 5.5 ± 0.9 | 3.6 ± 0.5 | 2.3 ± 0.2 | 7.7 ± 0.7 |
| Glucosamine | 7.1 ± 0.9 | 3.5 ± 0.8 | 3.0 ± 0.2 | 1.9 ± 0.3 |
| Glucosamine-6-Phosphate | 5.9 ± 0.7 | 3.0 ± 0.2 | 7.3 ± 0.5 | 7.5 ± 0.5 |
| Glucose | 9.0 ± 0.5 | 6.4 ± 1.0 | 4.4 ± 0.6 | 2.3 ± 0.3 |
| Glucose-1-phosphate | 4.2 ± 0.8 | 4.1 ± 1.0 | 4.3 ± 0.2 | 2.1 ± 0.3 |
| Glucose-6-Phosphate | 11.5 ± 0.4 | 7.9 ± 0.3 | 3.9 ± 0.5 | 2.2 ± 0.4 |
| Glucuronic acid | 12.9 ± 0.8 | 2.3 ± 0.4 | 3.6 ± 0.5 | 6.8 ± 1.0 |
| Glutamic acid | 9.0 ± 0.3 | 7.9 ± 0.9 | 2.7 ± 0.3 | 6.1 ± 0.4 |
| Glutamine | 3.4 ± 0.8 | 2.1 ± 0.2 | 3.4 ± 0.2 | 5.4 ± 0.9 |
| Glutathione | 6.5 ± 0.4 | 2.0 ± 0.3 | 4.1 ± 0.5 | 4.0 ± 0.5 |
| Glutathionylspermine | 6.6 ± 0.9 | 4.2 ± 0.4 | 3.9 ± 1.0 | 3.8 ± 0.3 |
| Glycerophosphocholine | 11.3 ± 0.7 | 2.7 ± 0.5 | 2.8 ± 0.2 | 6.2 ± 0.4 |
| Glycine | 3.2 ± 0.8 | 3.6 ± 0.4 | 7.2 ± 0.8 | 3.3 ± 0.8 |
| Glycolic acid | 10.1 ± 0.7 | 7.6 ± 0.6 | 4.7 ± 0.8 | 3.6 ± 0.8 |
| Guanidinoacetic acid | 4.1 ± 0.8 | 1.5 ± 0.4 | 6.6 ± 0.6 | 3.4 ± 0.6 |
| Guanidinobutanoic acid | 3.8 ± 0.4 | 2.2 ± 0.3 | 7.4 ± 0.7 | 7.1 ± 0.4 |
| Guanine | 8.6 ± 0.5 | 1.7 ± 0.3 | 3.0 ± 0.7 | 2.0 ± 0.5 |
| Guanosine | 12.7 ± 0.8 | 4.9 ± 0.2 | 3.6 ± 0.9 | 7.6 ± 0.8 |
| Guanosine 5'-diphosphate | 4.3 ± 0.2 | 2.2 ± 0.3 | 6.4 ± 0.6 | 6.8 ± 0.9 |
| Guanosine-5'-monophosphate | 8.9 ± 0.4 | 6.4 ± 0.4 | 6.1 ± 0.2 | 7.6 ± 0.2 |
| Hercynine | 12.4 ± 0.9 | 4.6 ± 0.4 | 7.6 ± 0.2 | 5.1 ± 0.7 |
| Hippuric acid | 5.3 ± 0.5 | 7.5 ± 0.6 | 2.3 ± 0.3 | 3.9 ± 0.6 |
| Histamine | 12.6 ± 0.4 | 7.1 ± 0.5 | 2.9 ± 0.3 | 4.1 ± 0.5 |
| Histidine | 11.9 ± 0.9 | 3.2 ± 0.6 | 7.8 ± 0.6 | 1.5 ± 0.2 |
| Homoalanine | 10.0 ± 0.7 | 4.1 ± 0.4 | 6.3 ± 0.7 | 5.6 ± 0.2 |
| Homoarginine | 3.6 ± 0.9 | 7.0 ± 0.4 | 2.5 ± 0.4 | 5.1 ± 0.9 |
| Homocysteine | 10.8 ± 0.5 | 6.9 ± 1.0 | 4.7 ± 0.7 | 2.8 ± 0.5 |
| Homogentisic acid | 7.7 ± 0.6 | 4.7 ± 0.4 | 7.4 ± 0.3 | 5.5 ± 0.5 |
| Homoleucine | 8.5 ± 0.8 | 5.7 ± 0.5 | 8.0 ± 0.9 | 2.3 ± 0.3 |
| Homovanillic acid | 12.9 ± 0.9 | 6.3 ± 0.5 | 7.0 ± 0.9 | 5.1 ± 0.9 |
| Hydantoin-5-propionate | 10.3 ± 0.4 | 2.1 ± 0.2 | 2.8 ± 0.6 | 4.4 ± 0.7 |
| Hypotaurine | 12.9 ± 0.6 | 2.7 ± 0.3 | 4.8 ± 0.5 | 3.2 ± 0.4 |
| Hypoxanthine | 7.3 ± 1.0 | 1.4 ± 0.3 | 5.1 ± 0.5 | 2.7 ± 0.7 |
| Indole-3-acetamide | 4.4 ± 0.6 | 4.6 ± 0.3 | 4.1 ± 0.3 | 7.4 ± 1.0 |
| Inosine | 4.0 ± 0.4 | 2.6 ± 0.6 | 5.0 ± 0.2 | 7.7 ± 0.4 |
| Inosine 5'-monophosphate | 3.4 ± 0.3 | 4.9 ± 1.0 | 2.3 ± 0.3 | 5.6 ± 0.8 |
| Isocitric Acid | 7.4 ± 0.6 | 3.5 ± 0.5 | 4.2 ± 0.3 | 6.7 ± 0.9 |
| Isoleucine | 5.6 ± 0.3 | 6.3 ± 0.6 | 7.4 ± 0.6 | 7.3 ± 0.7 |
| Lactic Acid | 6.7 ± 0.5 | 2.8 ± 0.2 | 6.6 ± 0.3 | 3.3 ± 0.4 |
| Leucine | 7.4 ± 0.7 | 6.5 ± 0.2 | 4.5 ± 0.9 | 1.8 ± 0.4 |
| Levulinic acid | 3.2 ± 0.5 | 4.3 ± 0.6 | 7.4 ± 0.4 | 7.4 ± 0.5 |
| Lysine | 8.1 ± 0.6 | 6.7 ± 0.3 | 7.3 ± 0.4 | 6.6 ± 1.0 |
| Malic Acid | 6.0 ± 0.6 | 3.6 ± 0.3 | 7.7 ± 0.4 | 4.8 ± 0.6 |
| Mannose-1 with Mannose-6-Phosphate | 8.6 ± 0.3 | 6.9 ± 0.4 | 4.4 ± 0.9 | 7.6 ± 0.4 |
| Mesaconic acid | 9.6 ± 1.0 | 1.9 ± 0.4 | 4.3 ± 0.4 | 1.4 ± 0.3 |
| Methionine | 3.0 ± 0.4 | 4.3 ± 0.8 | 4.6 ± 0.5 | 4.5 ± 0.8 |
| Methionine sulfoxide | 5.8 ± 0.8 | 5.9 ± 0.7 | 6.3 ± 0.3 | 5.8 ± 0.5 |
| Methylmalonic Acid | 5.1 ± 0.9 | 6.5 ± 0.7 | 2.7 ± 0.4 | 6.4 ± 0.6 |
| N1-Acetylspermidine | 12.8 ± 0.3 | 2.3 ± 0.3 | 7.4 ± 0.3 | 6.9 ± 0.8 |
| N1-Acetylspermine | 9.3 ± 1.0 | 7.9 ± 0.4 | 2.5 ± 0.2 | 2.5 ± 0.6 |
| N8-Acetylspermidine | 4.5 ± 0.2 | 1.9 ± 0.3 | 1.5 ± 0.3 | 4.0 ± 1.0 |
| N-Acetyl-Aspartyl-Glutamate | 9.8 ± 0.6 | 5.8 ± 0.2 | 4.6 ± 0.2 | 5.9 ± 0.3 |
| N-Acetyl-Aspartyl-Glutamyl-Glutamate | 10.0 ± 0.4 | 7.9 ± 0.4 | 3.0 ± 0.4 | 4.4 ± 0.8 |
| N-Acetyl-Galactosamine | 10.0 ± 0.6 | 4.9 ± 0.8 | 6.0 ± 0.2 | 6.4 ± 0.6 |
| N-Acetyl-Leucine | 5.3 ± 0.5 | 4.9 ± 0.4 | 5.3 ± 0.6 | 6.2 ± 0.7 |
| N-Acetyl-Methionine | 3.1 ± 0.4 | 5.3 ± 0.7 | 5.1 ± 0.7 | 5.1 ± 0.3 |
| N-Acetylneuraminic acid | 10.4 ± 0.7 | 4.4 ± 1.0 | 6.2 ± 0.9 | 1.5 ± 0.3 |
| N-Acetyl-Proline | 11.8 ± 0.9 | 6.3 ± 0.6 | 6.2 ± 0.4 | 6.5 ± 0.8 |
| N-Acetyl-Putrescine | 7.3 ± 0.3 | 6.6 ± 0.7 | 7.4 ± 0.5 | 5.2 ± 0.9 |
| N-Acetyl-Serine | 6.1 ± 0.7 | 7.8 ± 0.4 | 3.4 ± 0.7 | 3.3 ± 0.4 |
| N-Acetyl-Serotonin | 7.5 ± 0.8 | 6.3 ± 0.3 | 5.8 ± 0.3 | 1.7 ± 0.3 |
| N-Acyl-Homoserine-lactone | 12.4 ± 0.4 | 2.6 ± 0.5 | 3.4 ± 0.8 | 4.1 ± 0.9 |
| NAD | 4.6 ± 0.6 | 6.2 ± 0.9 | 2.3 ± 0.4 | 3.4 ± 0.3 |
| NADH | 9.1 ± 0.6 | 2.6 ± 0.4 | 7.6 ± 1.0 | 5.8 ± 0.7 |
| NADP | 5.7 ± 0.2 | 2.2 ± 0.2 | 7.4 ± 1.0 | 2.8 ± 0.4 |
| NADPH | 7.8 ± 0.3 | 2.6 ± 0.5 | 3.1 ± 0.6 | 3.0 ± 0.7 |
| N-alpha-Acetyl-lysine | 4.7 ± 0.4 | 2.0 ± 0.3 | 6.4 ± 0.9 | 6.1 ± 0.6 |
| N-alpha-Acetyl-Ornithine | 10.5 ± 0.6 | 3.6 ± 0.9 | 3.9 ± 0.8 | 1.4 ± 0.3 |
| N-Carbamoyl-Proline | 11.9 ± 0.3 | 7.1 ± 0.9 | 3.1 ± 0.3 | 2.1 ± 0.5 |
| N-Carbamoyl-Sarcosine | 12.8 ± 0.9 | 1.5 ± 0.2 | 7.0 ± 0.9 | 6.8 ± 0.7 |
| N-Epsilon-Acetyl-lysine | 8.0 ± 0.4 | 6.4 ± 0.7 | 5.5 ± 0.2 | 5.4 ± 1.0 |
| Nicotinamide | 8.4 ± 0.2 | 5.9 ± 0.8 | 7.8 ± 0.3 | 4.0 ± 0.2 |
| N-Methyl-Aspartic Acid | 12.0 ± 0.6 | 5.0 ± 0.5 | 7.3 ± 0.3 | 6.6 ± 0.3 |
| O-Phosphoethanolamine | 4.2 ± 0.6 | 7.2 ± 0.9 | 3.1 ± 0.3 | 5.1 ± 0.2 |
| Ophthalmic acid | 11.9 ± 0.4 | 4.8 ± 0.2 | 1.7 ± 0.2 | 7.6 ± 0.6 |
| Ornithine | 10.4 ± 0.8 | 2.7 ± 0.7 | 5.9 ± 1.0 | 2.7 ± 0.6 |
| Orotidine 5'-monophosphate | 5.4 ± 1.0 | 4.0 ± 0.8 | 2.9 ± 0.6 | 5.8 ± 0.7 |
| Oxaloacetate | 6.2 ± 0.3 | 4.1 ± 0.2 | 2.4 ± 0.3 | 1.7 ± 0.2 |
| Pantothenic acid | 4.0 ± 0.2 | 7.1 ± 0.7 | 3.7 ± 0.9 | 7.2 ± 0.9 |
| Phenylalanine | 13.0 ± 1.0 | 6.6 ± 0.5 | 2.7 ± 0.6 | 7.6 ± 0.6 |
| Phenylpyruvate | 7.5 ± 0.2 | 6.7 ± 0.5 | 3.6 ± 0.2 | 3.6 ± 0.2 |
| Phosphodimethylethanolamine | 12.6 ± 0.3 | 5.9 ± 0.7 | 6.1 ± 0.7 | 2.2 ± 0.3 |
| Phosphoenolpyruvate | 9.2 ± 0.9 | 2.8 ± 0.3 | 2.8 ± 0.4 | 2.2 ± 0.4 |
| Phosphorylcholine | 3.0 ± 0.7 | 2.4 ± 0.4 | 2.8 ± 0.6 | 6.8 ± 0.2 |
| Pipecolic acid | 10.0 ± 0.2 | 5.5 ± 0.5 | 3.0 ± 0.6 | 3.9 ± 0.6 |
| Proline | 11.5 ± 0.5 | 3.5 ± 0.2 | 3.7 ± 0.8 | 3.7 ± 0.6 |
| Putrescine | 8.1 ± 0.8 | 7.3 ± 0.6 | 6.8 ± 0.8 | 1.6 ± 0.3 |
| Pyridoxal | 7.8 ± 0.3 | 1.8 ± 0.3 | 3.8 ± 0.6 | 5.8 ± 0.7 |
| Pyridoxamine | 12.4 ± 0.6 | 6.4 ± 0.8 | 7.6 ± 0.3 | 4.2 ± 0.3 |
| Pyridoxine | 10.0 ± 0.7 | 2.1 ± 0.3 | 5.4 ± 0.6 | 4.6 ± 0.8 |
| Pyroglutamic acid | 9.7 ± 0.3 | 3.9 ± 0.6 | 1.8 ± 0.4 | 2.9 ± 0.4 |
| Pyrrolysine | 3.2 ± 0.6 | 1.9 ± 0.3 | 3.7 ± 0.5 | 4.5 ± 0.9 |
| Pyruvic Acid | 10.7 ± 0.8 | 4.2 ± 0.3 | 2.3 ± 0.3 | 2.2 ± 0.4 |
| Riboflavin | 4.5 ± 0.9 | 1.9 ± 0.3 | 1.5 ± 0.3 | 7.9 ± 0.4 |
| Ribose-5-Phosphate | 5.8 ± 0.2 | 4.1 ± 0.7 | 6.1 ± 0.7 | 1.4 ± 0.2 |
| Ribulose-1-Phosphate | 5.3 ± 0.8 | 4.4 ± 1.0 | 3.7 ± 0.8 | 6.9 ± 0.9 |
| Ribulose-5-Phosphate | 3.0 ± 0.3 | 3.9 ± 0.7 | 4.5 ± 0.3 | 2.3 ± 0.5 |
| Saccharopine | 7.1 ± 0.5 | 6.8 ± 0.6 | 6.7 ± 0.5 | 2.5 ± 0.3 |
| S-Adenosylhomocysteine | 11.5 ± 0.5 | 7.6 ± 1.0 | 6.2 ± 0.3 | 2.1 ± 0.5 |
| S-Adenosylmethioninamine | 11.0 ± 0.5 | 1.9 ± 0.2 | 8.0 ± 0.8 | 4.1 ± 0.3 |
| S-Adenosylmethionine | 9.0 ± 0.6 | 5.1 ± 0.9 | 3.9 ± 0.6 | 3.4 ± 0.5 |
| Sarcosine | 7.3 ± 0.2 | 5.0 ± 0.5 | 2.5 ± 0.2 | 1.7 ± 0.4 |
| Sedoheptulose-7-phosphate | 7.5 ± 0.2 | 1.5 ± 0.2 | 4.1 ± 0.3 | 4.8 ± 0.4 |
| Selenohomocysteine | 4.0 ± 0.3 | 1.6 ± 0.4 | 3.9 ± 0.4 | 7.1 ± 0.8 |
| Selenomethionine | 7.0 ± 0.6 | 4.8 ± 0.9 | 3.2 ± 0.3 | 2.3 ± 0.4 |
| Serine | 9.9 ± 0.7 | 2.9 ± 0.6 | 2.5 ± 0.2 | 3.8 ± 0.3 |
| S-Methyl-Thioinosine | 5.7 ± 0.7 | 1.9 ± 0.5 | 6.3 ± 1.0 | 1.4 ± 0.3 |
| Spermidine | 7.6 ± 0.9 | 5.0 ± 0.7 | 6.7 ± 0.5 | 3.7 ± 0.3 |
| Spermine | 9.0 ± 0.7 | 3.0 ± 0.7 | 6.5 ± 0.7 | 3.9 ± 0.6 |
| Succinate | 9.6 ± 0.6 | 4.1 ± 0.6 | 5.8 ± 0.2 | 3.3 ± 0.3 |
| Sucrose | 6.5 ± 0.6 | 2.3 ± 0.5 | 3.5 ± 0.7 | 8.0 ± 0.7 |
| Symmetric dimethylarginine | 9.7 ± 1.0 | 4.1 ± 0.6 | 8.0 ± 0.9 | 6.6 ± 0.7 |
| Taurine | 11.2 ± 1.0 | 1.7 ± 0.3 | 4.4 ± 0.2 | 4.8 ± 0.8 |
| Tetrahydroneopterin | 6.7 ± 0.7 | 3.8 ± 0.5 | 4.7 ± 0.7 | 3.6 ± 0.4 |
| Thiamine | 4.2 ± 0.2 | 6.6 ± 0.8 | 4.0 ± 0.3 | 2.4 ± 0.5 |
| Thiamine monophosphate | 12.1 ± 1.0 | 3.4 ± 0.2 | 3.1 ± 0.7 | 5.1 ± 0.3 |
| Threonic acid | 8.7 ± 0.8 | 3.3 ± 0.4 | 7.8 ± 0.8 | 4.3 ± 0.8 |
| Threonine | 6.6 ± 0.4 | 5.0 ± 0.7 | 4.5 ± 0.9 | 3.5 ± 0.7 |
| Thymidine | 3.8 ± 0.3 | 6.2 ± 0.9 | 5.4 ± 0.7 | 5.0 ± 0.3 |
| Thymidine Monophosphate | 11.1 ± 0.4 | 5.8 ± 0.5 | 4.8 ± 0.8 | 7.4 ± 0.5 |
| Thymine | 11.1 ± 0.6 | 4.2 ± 1.0 | 3.4 ± 0.4 | 4.5 ± 0.6 |
| Trigonelleine | 8.1 ± 0.7 | 2.6 ± 0.5 | 7.9 ± 0.4 | 6.4 ± 0.9 |
| Trimethylamine N-oxide | 3.9 ± 0.5 | 2.3 ± 0.4 | 2.4 ± 0.5 | 6.4 ± 0.7 |
| Trimethyllysine | 4.3 ± 1.0 | 2.1 ± 0.3 | 3.6 ± 0.6 | 4.2 ± 0.7 |
| Tryptophan | 10.2 ± 0.8 | 1.4 ± 0.2 | 3.6 ± 0.6 | 3.6 ± 0.7 |
| Tyramine | 12.9 ± 0.4 | 3.7 ± 0.3 | 5.3 ± 0.8 | 2.4 ± 0.2 |
| Tyrosine | 9.6 ± 0.3 | 5.6 ± 0.4 | 7.6 ± 0.3 | 4.7 ± 0.5 |
| Uracil | 6.7 ± 0.3 | 7.4 ± 0.3 | 7.3 ± 0.4 | 5.5 ± 0.4 |
| Uridine 5'-diphosphate | 10.5 ± 1.0 | 2.2 ± 0.3 | 7.0 ± 0.7 | 6.7 ± 0.9 |
| Uridine Monophosphate | 9.7 ± 0.3 | 7.1 ± 0.4 | 5.7 ± 0.3 | 4.9 ± 0.5 |
| Uridine-5-Diphosphogalactose | 8.1 ± 0.5 | 8.0 ± 0.4 | 7.0 ± 0.4 | 2.7 ± 0.4 |
| Urocanic acid | 4.2 ± 0.7 | 3.1 ± 0.6 | 1.8 ± 0.4 | 6.4 ± 0.3 |
| UTP | 12.8 ± 0.9 | 2.6 ± 0.4 | 4.4 ± 0.9 | 3.5 ± 0.4 |
| Valine | 3.2 ± 0.7 | 4.4 ± 0.7 | 7.2 ± 0.5 | 7.9 ± 0.9 |
| XMP | 10.9 ± 0.5 | 6.8 ± 0.7 | 4.7 ± 0.3 | 7.3 ± 0.5 |
| Xylulose-5-Phosphate | 3.3 ± 0.3 | 6.1 ± 0.9 | 3.7 ± 0.7 | 2.5 ± 0.4 |

### **Table S13** | Intra – day precision (as % RSD) (Tumor tissue extract as matrix, n = 5). The concentrations of QC_LLOQ_, QC_Low_, QC_Mid_ and QC_High_ were 10 ng/mL, 25 ng/mL, 100 ng/mL and 500 ng/mL respectively.

| Precision, % | QC_LLOQ_ | QC_low_ | QC_mid_ | QC_high_ |
| --- | --- | --- | --- | --- |
| 1-Methylguanine | 5.7 ± 1.1 | 5.6 ± 0.1 | 1.7 ± 0.6 | 2.3 ± 1.2 |
| 1-Methylhistidine | 7.1 ± 0.2 | 8.3 ± 0.8 | 5.7 ± 0.9 | 3.6 ± 0.8 |
| 1-Methylnicotinamide | 6.7 ± 0.1 | 6.2 ± 1.1 | 2.1 ± 0.9 | 2.1 ± 0.3 |
| 2-Aminoadipic Acid | 12.5 ± 0.3 | 12.3 ± 1.0 | 4.9 ± 0.8 | 1.9 ± 0.8 |
| 2-Deoxyadenosine | 5.6 ± 0.3 | 11.7 ± 0.3 | 4.6 ± 1.0 | 2.3 ± 1.0 |
| 2'-Deoxyadenosine 5'-monophosphate | 6.3 ± 0.5 | 9.9 ± 0.2 | 4.4 ± 0.7 | 2.5 ± 0.3 |
| 2'-deoxyadenosine 5'-triphosphate | 7.6 ± 0.5 | 7.3 ± 0.9 | 4.4 ± 0.2 | 4.5 ± 1.0 |
| 2'-Deoxycytidine-5'-monophosphate | 10.9 ± 0.5 | 10.8 ± 0.7 | 2.3 ± 0.8 | 3.0 ± 0.3 |
| 2-Deoxyguanosine | 7.7 ± 0.6 | 8.4 ± 0.2 | 5.7 ± 0.6 | 2.7 ± 0.4 |
| 2-hydroxybutyric acid | 12.4 ± 0.7 | 8.2 ± 1.0 | 4.3 ± 1.0 | 4.9 ± 0.3 |
| 2-hydroxyglutaric acid | 9.8 ± 1.1 | 12.2 ± 1.0 | 5.2 ± 1.0 | 2.7 ± 1.0 |
| 2-Hydroxyisocaproic acid | 10.6 ± 0.8 | 12.6 ± 0.1 | 4.2 ± 1.1 | 2.5 ± 0.7 |
| 2-ketobutyric acid | 10.2 ± 0.9 | 9.4 ± 0.9 | 4.6 ± 0.4 | 2.7 ± 1.0 |
| 2-oxoadipic acid | 9.1 ± 0.9 | 9.6 ± 0.6 | 4.3 ± 0.3 | 3.6 ± 1.1 |
| 2-Phenylglycine | 6.5 ± 0.3 | 11.0 ± 1.1 | 3.0 ± 0.2 | 2.0 ± 0.8 |
| 2-phosphoglycerate | 11.7 ± 0.4 | 12.3 ± 0.3 | 5.2 ± 0.5 | 2.7 ± 0.9 |
| 3-hydroxy-3-methyl-2-oxobutanoate | 12.0 ± 0.3 | 12.2 ± 0.6 | 4.9 ± 0.6 | 4.2 ± 0.7 |
| 3-hydroxybutyric acid | 11.1 ± 1.1 | 10.3 ± 0.8 | 3.5 ± 0.9 | 4.0 ± 0.3 |
| 3-Methylhistidine | 11.0 ± 0.9 | 10.5 ± 1.1 | 2.4 ± 0.7 | 1.9 ± 0.3 |
| 3-Phosphoglyceraldehyde | 8.3 ± 0.6 | 8.3 ± 0.2 | 5.9 ± 0.4 | 1.8 ± 0.9 |
| 3-phosphoglycerate | 12.9 ± 0.3 | 7.8 ± 0.3 | 4.3 ± 0.8 | 4.8 ± 0.3 |
| 4-amino-butyric acid | 7.4 ± 0.9 | 12.0 ± 0.3 | 5.0 ± 0.3 | 2.7 ± 0.4 |
| 4-aspartyl-phosphate | 8.5 ± 0.1 | 11.7 ± 1.1 | 4.6 ± 0.8 | 3.3 ± 0.9 |
| 4-Hydroxybenzoic acid | 8.2 ± 0.7 | 10.9 ± 0.8 | 5.2 ± 0.2 | 2.3 ± 0.7 |
| 4-Hydroxyphenylacetylglycine | 12.4 ± 0.2 | 10.7 ± 0.4 | 3.4 ± 0.5 | 3.6 ± 0.2 |
| 4-methyl-2-oxopentanoate | 5.5 ± 0.4 | 10.1 ± 0.8 | 1.6 ± 0.5 | 2.7 ± 1.0 |
| 4-Oxoglutaramate | 10.8 ± 0.2 | 8.3 ± 1.0 | 4.1 ± 1.0 | 4.1 ± 0.3 |
| 5-Hydroxyindolepyruvate | 11.9 ± 0.4 | 6.9 ± 0.6 | 2.6 ± 0.2 | 2.5 ± 0.7 |
| 5-Hydroxylysine | 9.3 ± 0.6 | 12.6 ± 0.8 | 1.7 ± 1.0 | 4.3 ± 0.4 |
| 5-Hydroxy-Tryptophan | 9.3 ± 0.8 | 10.3 ± 0.6 | 3.3 ± 0.6 | 4.1 ± 0.4 |
| 5-Phosphoribosylamine | 10.1 ± 0.3 | 12.3 ± 0.9 | 2.2 ± 0.8 | 3.6 ± 1.0 |
| 6-phosphogluconate | 6.3 ± 0.2 | 11.7 ± 1.1 | 3.4 ± 0.8 | 2.7 ± 0.3 |
| Acetylcarnitine | 12.4 ± 0.5 | 12.2 ± 0.6 | 3.7 ± 1.1 | 4.5 ± 1.2 |
| Aconitic Acid | 10.3 ± 1.0 | 11.8 ± 0.6 | 2.8 ± 0.4 | 2.6 ± 0.1 |
| Adenine | 10.6 ± 0.2 | 11.5 ± 0.9 | 2.5 ± 1.0 | 4.7 ± 0.3 |
| Adenosine | 6.4 ± 0.9 | 11.4 ± 0.7 | 5.4 ± 0.4 | 4.3 ± 0.7 |
| Adenosine 3',5'-cyclic monophosphate | 6.7 ± 0.2 | 11.2 ± 0.2 | 3.0 ± 0.5 | 3.5 ± 0.3 |
| Adenosine Monophosphate | 8.9 ± 0.5 | 11.7 ± 0.5 | 5.8 ± 0.2 | 2.6 ± 0.2 |
| Adenosine-5'-diphosphate | 6.9 ± 0.3 | 6.2 ± 1.2 | 5.7 ± 0.9 | 2.1 ± 1.1 |
| Adenosine-5'-triphosphate | 5.6 ± 0.9 | 10.7 ± 0.9 | 3.3 ± 0.6 | 1.9 ± 0.3 |
| Adenylosuccinate | 9.7 ± 0.2 | 11.6 ± 0.7 | 5.7 ± 0.7 | 2.0 ± 1.0 |
| Alanine | 11.1 ± 1.0 | 11.6 ± 1.0 | 3.8 ± 0.6 | 3.2 ± 0.4 |
| alpha-ketoglutaric acid | 7.0 ± 0.9 | 7.1 ± 0.6 | 3.6 ± 0.1 | 2.4 ± 0.6 |
| Anserine | 8.8 ± 0.3 | 7.9 ± 0.2 | 4.7 ± 0.6 | 1.8 ± 0.2 |
| Arginine | 5.2 ± 0.2 | 12.3 ± 0.1 | 2.4 ± 1.0 | 4.7 ± 0.8 |
| Argininosuccinic Acid | 8.7 ± 0.8 | 7.9 ± 0.6 | 2.2 ± 0.9 | 2.8 ± 0.8 |
| Asparagine | 12.9 ± 1.0 | 12.2 ± 1.0 | 5.0 ± 0.6 | 3.2 ± 0.2 |
| Aspartic acid | 6.7 ± 1.0 | 10.0 ± 1.0 | 4.4 ± 1.2 | 4.9 ± 0.9 |
| Asymmetric dimethylarginine | 7.6 ± 0.5 | 5.8 ± 0.4 | 3.9 ± 0.6 | 4.6 ± 0.8 |
| Azelaic acid | 8.8 ± 0.3 | 9.8 ± 0.6 | 1.5 ± 1.0 | 3.5 ± 0.2 |
| beta-Alanine | 7.8 ± 0.5 | 12.1 ± 0.7 | 1.9 ± 0.6 | 3.6 ± 0.9 |
| Betaine | 9.5 ± 1.0 | 11.3 ± 0.9 | 3.5 ± 0.9 | 1.8 ± 1.0 |
| Biopterin | 10.2 ± 1.1 | 9.5 ± 0.7 | 1.8 ± 0.8 | 4.6 ± 0.5 |
| Biotin | 5.0 ± 0.7 | 6.2 ± 1.0 | 5.8 ± 0.9 | 3.9 ± 0.5 |
| Carnitine | 12.8 ± 0.5 | 6.4 ± 0.2 | 3.9 ± 0.7 | 3.1 ± 0.5 |
| Carnosine | 8.6 ± 0.7 | 11.5 ± 0.7 | 5.1 ± 0.7 | 2.6 ± 0.9 |
| CDP-choline | 8.4 ± 0.4 | 10.4 ± 0.9 | 2.7 ± 0.4 | 2.1 ± 1.1 |
| cGMP | 10.5 ± 1.0 | 10.2 ± 0.9 | 5.0 ± 1.0 | 3.9 ± 0.8 |
| Choline | 11.1 ± 1.0 | 5.9 ± 1.1 | 2.8 ± 0.4 | 2.2 ± 0.6 |
| Cinnamic acid | 7.4 ± 1.1 | 7.9 ± 0.9 | 2.0 ± 0.7 | 4.7 ± 0.7 |
| Citraconic acid | 8.0 ± 0.6 | 9.3 ± 0.5 | 1.6 ± 0.6 | 4.5 ± 1.2 |
| Citramalic acid | 6.9 ± 1.1 | 7.2 ± 0.4 | 3.6 ± 0.2 | 2.4 ± 0.8 |
| Citric Acid | 10.7 ± 0.8 | 8.3 ± 0.6 | 5.9 ± 0.9 | 3.7 ± 0.4 |
| Citrulline | 9.6 ± 0.5 | 6.8 ± 1.0 | 2.6 ± 1.0 | 1.7 ± 0.7 |
| Creatine | 6.0 ± 0.9 | 9.7 ± 1.1 | 2.6 ± 1.1 | 1.6 ± 0.2 |
| Creatinine | 6.8 ± 0.5 | 11.8 ± 1.0 | 2.8 ± 1.1 | 2.7 ± 0.3 |
| Cystathionine | 9.4 ± 1.0 | 8.2 ± 0.7 | 2.2 ± 0.7 | 4.7 ± 0.2 |
| Cysteamine | 7.0 ± 0.9 | 9.7 ± 0.8 | 3.1 ± 0.4 | 2.3 ± 1.2 |
| Cysteine | 5.2 ± 1.1 | 12.5 ± 1.1 | 2.1 ± 0.2 | 3.4 ± 0.2 |
| Cysteinylglycine | 6.9 ± 0.3 | 6.6 ± 0.3 | 5.4 ± 0.5 | 4.9 ± 0.2 |
| Cystine | 11.0 ± 0.9 | 8.9 ± 1.2 | 2.0 ± 0.6 | 4.8 ± 1.0 |
| Cytidine | 9.0 ± 1.2 | 7.4 ± 0.3 | 5.3 ± 0.1 | 3.7 ± 0.8 |
| Cytidine 3',5'-cyclic monophosphate | 7.1 ± 1.0 | 8.4 ± 0.2 | 3.5 ± 0.4 | 2.0 ± 0.2 |
| Cytidine 5'-diphosphate | 9.3 ± 0.5 | 8.3 ± 0.4 | 2.1 ± 0.2 | 2.1 ± 0.5 |
| Cytidine 5'-triphosphate | 8.1 ± 0.6 | 6.3 ± 1.1 | 2.2 ± 0.3 | 3.8 ± 0.6 |
| Cytidine monophosphate | 11.5 ± 0.4 | 9.3 ± 0.7 | 4.9 ± 1.1 | 2.6 ± 0.8 |
| Cytosine | 11.4 ± 0.1 | 12.1 ± 0.8 | 5.7 ± 0.8 | 4.2 ± 0.5 |
| dADP | 9.2 ± 0.2 | 9.8 ± 0.8 | 3.6 ± 0.7 | 3.2 ± 0.6 |
| Diethanolamine | 10.0 ± 0.7 | 10.2 ± 1.0 | 2.4 ± 0.5 | 2.7 ± 0.7 |
| Dihydrobiopterin | 10.9 ± 0.8 | 7.7 ± 1.0 | 3.2 ± 0.4 | 3.5 ± 1.1 |
| Dihydroneopterin triphosphate | 5.8 ± 0.9 | 5.9 ± 0.2 | 5.2 ± 0.9 | 5.0 ± 1.0 |
| Dihydroxyacetonephosphate | 12.1 ± 1.0 | 10.5 ± 0.9 | 5.0 ± 0.9 | 1.6 ± 0.6 |
| Dimethylglycine | 10.5 ± 0.1 | 6.1 ± 0.1 | 5.4 ± 0.8 | 4.2 ± 1.2 |
| Ectoine | 7.2 ± 0.4 | 5.9 ± 0.4 | 1.8 ± 1.0 | 1.9 ± 0.4 |
| Erythrose-4-phosphate | 10.0 ± 0.7 | 5.8 ± 0.8 | 2.4 ± 0.4 | 3.2 ± 0.8 |
| FAD | 7.3 ± 0.1 | 7.8 ± 0.5 | 5.6 ± 0.6 | 2.0 ± 0.3 |
| Folic acid | 6.5 ± 0.3 | 9.1 ± 0.1 | 3.1 ± 0.4 | 3.3 ± 0.9 |
| Fructose-6-phosphate | 8.2 ± 0.5 | 7.2 ± 1.1 | 6.0 ± 0.9 | 2.7 ± 0.4 |
| Fumaric Acid | 7.9 ± 0.1 | 8.2 ± 1.0 | 1.6 ± 1.0 | 2.8 ± 0.2 |
| Galactosamine | 6.4 ± 1.2 | 12.2 ± 0.2 | 3.8 ± 0.1 | 3.3 ± 0.9 |
| Galactose-1-phosphate | 7.0 ± 0.2 | 7.9 ± 1.1 | 5.5 ± 0.8 | 2.4 ± 1.2 |
| Galactose-6-Phosphate | 5.7 ± 1.1 | 6.4 ± 1.0 | 1.7 ± 0.4 | 3.6 ± 1.0 |
| gamma-Glutamylcysteine | 9.6 ± 0.8 | 11.2 ± 0.4 | 4.2 ± 1.1 | 4.1 ± 1.1 |
| Glucosamine | 8.9 ± 0.5 | 8.8 ± 0.6 | 5.7 ± 0.3 | 3.4 ± 1.0 |
| Glucosamine-6-Phosphate | 6.3 ± 0.5 | 11.2 ± 0.2 | 5.3 ± 0.7 | 1.5 ± 0.8 |
| Glucose | 9.7 ± 0.6 | 5.4 ± 0.9 | 2.7 ± 0.8 | 3.9 ± 0.7 |
| Glucose-1-phosphate | 12.9 ± 0.4 | 5.2 ± 0.9 | 2.1 ± 0.3 | 3.9 ± 0.2 |
| Glucose-6-Phosphate | 11.6 ± 0.9 | 10.5 ± 0.4 | 2.7 ± 0.1 | 2.9 ± 0.9 |
| Glucuronic acid | 5.8 ± 0.4 | 11.7 ± 0.1 | 3.4 ± 0.5 | 4.8 ± 0.9 |
| Glutamic acid | 9.1 ± 0.7 | 7.2 ± 0.7 | 4.8 ± 0.4 | 2.3 ± 1.2 |
| Glutamine | 6.0 ± 0.5 | 6.5 ± 0.6 | 3.0 ± 1.1 | 2.1 ± 0.7 |
| Glutathione | 12.9 ± 0.8 | 12.9 ± 0.7 | 6.0 ± 0.6 | 4.8 ± 0.8 |
| Glutathionylspermine | 7.3 ± 0.6 | 11.4 ± 0.4 | 3.2 ± 1.1 | 2.6 ± 0.7 |
| Glycerophosphocholine | 7.6 ± 1.1 | 7.6 ± 0.4 | 4.6 ± 1.1 | 2.2 ± 0.4 |
| Glycine | 8.0 ± 1.0 | 7.4 ± 0.5 | 5.9 ± 0.3 | 2.2 ± 0.9 |
| Glycolic acid | 9.1 ± 0.3 | 6.7 ± 0.8 | 4.0 ± 0.8 | 2.8 ± 0.2 |
| Guanidinoacetic acid | 7.6 ± 1.1 | 11.7 ± 0.6 | 3.3 ± 1.2 | 1.6 ± 0.3 |
| Guanidinobutanoic acid | 5.2 ± 0.7 | 5.5 ± 1.1 | 3.8 ± 1.0 | 3.0 ± 0.3 |
| Guanine | 11.5 ± 0.6 | 8.1 ± 0.2 | 4.9 ± 0.4 | 2.4 ± 0.5 |
| Guanosine | 6.6 ± 0.8 | 6.5 ± 0.3 | 4.8 ± 1.2 | 3.8 ± 0.7 |
| Guanosine 5'-diphosphate | 6.6 ± 0.7 | 6.6 ± 0.4 | 3.6 ± 0.2 | 1.9 ± 1.2 |
| Guanosine-5'-monophosphate | 8.3 ± 0.2 | 12.1 ± 0.9 | 2.5 ± 0.8 | 1.7 ± 0.2 |
| Hercynine | 5.4 ± 0.6 | 12.8 ± 0.9 | 5.5 ± 1.1 | 4.7 ± 0.2 |
| Hippuric acid | 8.6 ± 0.6 | 9.7 ± 1.1 | 3.2 ± 0.9 | 3.2 ± 0.2 |
| Histamine | 5.8 ± 0.6 | 11.4 ± 0.6 | 5.7 ± 0.1 | 4.2 ± 0.9 |
| Histidine | 11.8 ± 0.8 | 10.0 ± 0.8 | 3.6 ± 0.2 | 2.2 ± 0.1 |
| Homoalanine | 12.4 ± 0.4 | 8.0 ± 1.0 | 4.3 ± 1.1 | 2.0 ± 0.2 |
| Homoarginine | 9.7 ± 0.4 | 7.2 ± 0.3 | 5.0 ± 0.5 | 3.2 ± 0.8 |
| Homocysteine | 5.4 ± 1.2 | 11.1 ± 1.1 | 4.4 ± 0.1 | 4.8 ± 1.0 |
| Homogentisic acid | 6.9 ± 1.0 | 12.5 ± 0.5 | 4.4 ± 0.6 | 3.1 ± 0.8 |
| Homoleucine | 6.5 ± 0.3 | 5.5 ± 0.3 | 4.0 ± 0.8 | 2.5 ± 0.5 |
| Homovanillic acid | 11.2 ± 0.9 | 10.8 ± 0.9 | 3.4 ± 0.1 | 4.1 ± 0.7 |
| Hydantoin-5-propionate | 8.8 ± 0.9 | 5.1 ± 0.8 | 5.2 ± 0.3 | 2.1 ± 0.7 |
| Hypotaurine | 10.9 ± 0.3 | 10.4 ± 1.1 | 1.8 ± 1.1 | 1.8 ± 0.8 |
| Hypoxanthine | 12.4 ± 0.4 | 5.2 ± 0.2 | 3.6 ± 0.4 | 1.8 ± 0.5 |
| Indole-3-acetamide | 5.9 ± 0.3 | 12.5 ± 0.1 | 3.1 ± 0.4 | 4.0 ± 1.2 |
| Inosine | 10.8 ± 1.1 | 9.3 ± 0.2 | 3.6 ± 1.1 | 3.2 ± 1.1 |
| Inosine 5'-monophosphate | 7.5 ± 0.7 | 8.2 ± 1.1 | 3.7 ± 0.8 | 1.6 ± 0.2 |
| Isocitric Acid | 8.1 ± 1.0 | 7.3 ± 0.7 | 4.3 ± 0.1 | 2.4 ± 1.0 |
| Isoleucine | 11.1 ± 0.9 | 5.9 ± 0.4 | 5.3 ± 0.7 | 2.0 ± 1.1 |
| Lactic Acid | 6.1 ± 0.8 | 6.8 ± 0.5 | 6.0 ± 0.8 | 2.6 ± 0.3 |
| Leucine | 8.8 ± 0.3 | 9.9 ± 1.1 | 5.3 ± 0.4 | 4.0 ± 1.0 |
| Levulinic acid | 10.9 ± 1.2 | 9.6 ± 0.3 | 1.8 ± 1.2 | 2.7 ± 1.0 |
| Lysine | 8.1 ± 0.1 | 13.0 ± 0.3 | 4.6 ± 0.8 | 4.7 ± 0.5 |
| Malic Acid | 6.4 ± 0.7 | 5.4 ± 1.1 | 5.1 ± 0.6 | 3.7 ± 0.8 |
| Mannose-1 with Mannose-6-Phosphate | 11.2 ± 0.6 | 6.8 ± 0.6 | 3.6 ± 0.2 | 3.2 ± 1.1 |
| Mesaconic acid | 7.1 ± 1.2 | 12.0 ± 0.2 | 1.9 ± 0.5 | 2.0 ± 0.4 |
| Methionine | 11.0 ± 0.4 | 7.4 ± 0.3 | 5.7 ± 0.5 | 4.1 ± 1.1 |
| Methionine sulfoxide | 11.6 ± 0.8 | 5.6 ± 0.1 | 5.8 ± 1.0 | 4.3 ± 1.0 |
| Methylmalonic Acid | 10.6 ± 0.8 | 12.2 ± 0.9 | 4.1 ± 1.1 | 2.9 ± 0.2 |
| N1-Acetylspermidine | 7.3 ± 1.2 | 8.5 ± 0.6 | 2.3 ± 0.9 | 3.5 ± 0.2 |
| N1-Acetylspermine | 8.7 ± 0.3 | 11.0 ± 0.2 | 5.1 ± 0.7 | 4.7 ± 0.2 |
| N8-Acetylspermidine | 12.6 ± 0.2 | 5.0 ± 0.9 | 5.1 ± 0.4 | 2.1 ± 0.2 |
| N-Acetyl-Aspartyl-Glutamate | 9.9 ± 0.4 | 5.2 ± 0.8 | 4.1 ± 0.9 | 2.8 ± 0.8 |
| N-Acetyl-Aspartyl-Glutamyl-Glutamate | 8.0 ± 0.8 | 11.9 ± 0.5 | 4.2 ± 0.1 | 2.2 ± 0.3 |
| N-Acetyl-Galactosamine | 7.5 ± 0.8 | 9.3 ± 0.8 | 4.0 ± 0.7 | 3.8 ± 0.8 |
| N-Acetyl-Leucine | 12.5 ± 1.2 | 7.8 ± 0.9 | 2.0 ± 1.1 | 2.4 ± 1.2 |
| N-Acetyl-Methionine | 12.7 ± 0.3 | 7.1 ± 1.1 | 5.4 ± 0.9 | 4.5 ± 0.9 |
| N-Acetylneuraminic acid | 9.0 ± 0.5 | 10.4 ± 0.1 | 4.0 ± 0.9 | 2.0 ± 0.9 |
| N-Acetyl-Proline | 6.3 ± 0.8 | 11.0 ± 0.9 | 4.3 ± 0.5 | 1.7 ± 0.6 |
| N-Acetyl-Putrescine | 10.8 ± 0.9 | 9.5 ± 0.3 | 2.6 ± 0.2 | 4.0 ± 0.4 |
| N-Acetyl-Serine | 6.8 ± 0.6 | 10.4 ± 0.7 | 2.9 ± 1.0 | 3.0 ± 0.1 |
| N-Acetyl-Serotonin | 9.3 ± 0.6 | 8.1 ± 0.3 | 2.8 ± 0.4 | 2.4 ± 1.0 |
| N-Acyl-Homoserine-lactone | 5.2 ± 0.7 | 13.0 ± 0.4 | 5.9 ± 0.3 | 4.7 ± 0.7 |
| NAD | 8.2 ± 0.6 | 6.6 ± 1.2 | 2.9 ± 0.5 | 2.8 ± 0.5 |
| NADH | 6.1 ± 0.5 | 12.3 ± 0.7 | 2.5 ± 0.8 | 3.7 ± 0.5 |
| NADP | 8.6 ± 0.7 | 10.0 ± 0.7 | 3.0 ± 0.8 | 3.9 ± 1.0 |
| NADPH | 5.9 ± 0.3 | 6.6 ± 0.1 | 4.6 ± 0.9 | 4.9 ± 1.0 |
| N-alpha-Acetyl-lysine | 10.3 ± 1.1 | 8.2 ± 0.6 | 3.8 ± 0.3 | 2.0 ± 0.1 |
| N-alpha-Acetyl-Ornithine | 9.5 ± 0.5 | 5.8 ± 0.2 | 4.7 ± 1.0 | 4.1 ± 1.1 |
| N-Carbamoyl-Proline | 5.5 ± 1.1 | 9.0 ± 0.7 | 5.6 ± 0.6 | 2.5 ± 0.9 |
| N-Carbamoyl-Sarcosine | 11.7 ± 0.6 | 9.9 ± 0.4 | 3.6 ± 0.9 | 2.0 ± 0.3 |
| N-Epsilon-Acetyl-lysine | 12.0 ± 0.3 | 6.5 ± 1.0 | 2.0 ± 0.7 | 1.5 ± 1.0 |
| Nicotinamide | 6.4 ± 0.5 | 6.1 ± 0.2 | 3.5 ± 0.2 | 3.7 ± 0.2 |
| N-Methyl-Aspartic Acid | 12.0 ± 0.5 | 10.3 ± 0.7 | 5.8 ± 0.7 | 2.5 ± 0.4 |
| O-Phosphoethanolamine | 12.6 ± 0.4 | 5.4 ± 1.0 | 3.6 ± 0.6 | 1.9 ± 0.6 |
| Ophthalmic acid | 6.0 ± 0.4 | 10.0 ± 0.2 | 3.8 ± 0.9 | 1.6 ± 0.9 |
| Ornithine | 8.4 ± 0.4 | 6.8 ± 0.7 | 4.0 ± 0.5 | 2.5 ± 0.5 |
| Orotidine 5'-monophosphate | 7.7 ± 0.3 | 9.5 ± 1.2 | 1.6 ± 0.4 | 3.9 ± 0.2 |
| Oxaloacetate | 9.4 ± 1.1 | 12.7 ± 0.7 | 5.4 ± 0.9 | 4.8 ± 1.2 |
| Pantothenic acid | 13.0 ± 0.2 | 12.0 ± 0.7 | 5.3 ± 1.2 | 1.9 ± 0.2 |
| Phenylalanine | 11.3 ± 0.3 | 5.9 ± 0.2 | 3.3 ± 0.3 | 1.6 ± 0.2 |
| Phenylpyruvate | 12.9 ± 0.2 | 5.7 ± 0.8 | 3.0 ± 1.1 | 2.5 ± 0.7 |
| Phosphodimethylethanolamine | 7.8 ± 0.6 | 5.6 ± 0.9 | 3.1 ± 0.9 | 3.7 ± 0.7 |
| Phosphoenolpyruvate | 7.1 ± 0.6 | 11.3 ± 0.7 | 1.7 ± 0.6 | 4.3 ± 0.9 |
| Phosphorylcholine | 12.6 ± 0.5 | 10.7 ± 0.7 | 4.7 ± 1.0 | 3.9 ± 0.7 |
| Pipecolic acid | 10.9 ± 0.4 | 9.8 ± 1.0 | 1.9 ± 0.4 | 3.1 ± 0.2 |
| Proline | 9.2 ± 0.6 | 6.3 ± 0.7 | 4.1 ± 0.3 | 2.9 ± 0.7 |
| Putrescine | 12.2 ± 0.4 | 8.5 ± 0.9 | 4.1 ± 0.2 | 1.6 ± 0.9 |
| Pyridoxal | 11.7 ± 0.5 | 9.8 ± 1.2 | 1.9 ± 0.7 | 3.3 ± 1.1 |
| Pyridoxamine | 11.6 ± 1.1 | 9.2 ± 1.0 | 4.5 ± 1.2 | 1.7 ± 0.3 |
| Pyridoxine | 7.8 ± 0.9 | 6.9 ± 0.5 | 5.5 ± 0.1 | 3.2 ± 0.9 |
| Pyroglutamic acid | 7.3 ± 0.5 | 12.5 ± 1.1 | 2.0 ± 0.7 | 4.5 ± 0.8 |
| Pyrrolysine | 12.0 ± 0.6 | 6.5 ± 0.7 | 5.5 ± 0.5 | 3.5 ± 1.0 |
| Pyruvic Acid | 10.2 ± 1.0 | 7.6 ± 0.3 | 4.9 ± 0.9 | 2.8 ± 0.5 |
| Riboflavin | 9.3 ± 1.0 | 9.2 ± 1.0 | 4.2 ± 1.1 | 2.3 ± 0.9 |
| Ribose-5-Phosphate | 5.0 ± 0.8 | 6.5 ± 0.3 | 1.6 ± 1.1 | 4.5 ± 0.3 |
| Ribulose-1-Phosphate | 12.2 ± 0.3 | 9.2 ± 0.3 | 5.3 ± 0.8 | 5.0 ± 0.9 |
| Ribulose-5-Phosphate | 12.9 ± 0.9 | 11.8 ± 1.2 | 5.4 ± 0.6 | 3.9 ± 0.6 |
| Saccharopine | 9.5 ± 0.2 | 11.7 ± 0.3 | 5.1 ± 0.9 | 2.7 ± 0.2 |
| S-Adenosylhomocysteine | 7.7 ± 0.8 | 12.4 ± 0.7 | 5.1 ± 0.8 | 3.6 ± 1.1 |
| S-Adenosylmethioninamine | 8.1 ± 0.4 | 12.6 ± 0.8 | 1.9 ± 0.3 | 4.3 ± 0.1 |
| S-Adenosylmethionine | 10.3 ± 0.6 | 11.0 ± 0.3 | 5.5 ± 0.7 | 2.8 ± 0.3 |
| Sarcosine | 11.2 ± 0.5 | 10.7 ± 1.1 | 4.4 ± 0.1 | 2.7 ± 1.0 |
| Sedoheptulose-7-phosphate | 6.4 ± 1.1 | 7.8 ± 0.7 | 4.0 ± 0.3 | 1.8 ± 0.9 |
| Selenohomocysteine | 10.4 ± 1.0 | 7.5 ± 1.0 | 4.1 ± 0.3 | 4.8 ± 0.8 |
| Selenomethionine | 12.7 ± 0.5 | 7.4 ± 1.1 | 1.6 ± 0.5 | 4.5 ± 1.1 |
| Serine | 9.3 ± 1.0 | 7.9 ± 0.5 | 5.4 ± 0.8 | 2.0 ± 0.2 |
| S-Methyl-Thioinosine | 7.7 ± 0.6 | 9.5 ± 1.0 | 2.1 ± 0.9 | 3.1 ± 1.0 |
| Spermidine | 12.7 ± 0.2 | 5.5 ± 0.8 | 5.3 ± 0.6 | 2.5 ± 0.8 |
| Spermine | 8.9 ± 1.1 | 9.6 ± 1.0 | 5.6 ± 0.8 | 4.1 ± 0.3 |
| Succinate | 9.6 ± 1.1 | 6.5 ± 1.1 | 3.7 ± 0.4 | 2.1 ± 0.5 |
| Sucrose | 5.3 ± 1.2 | 11.0 ± 0.7 | 5.0 ± 0.7 | 2.5 ± 1.1 |
| Symmetric dimethylarginine | 10.1 ± 0.5 | 10.1 ± 0.9 | 2.1 ± 0.2 | 2.0 ± 0.3 |
| Taurine | 11.1 ± 0.3 | 11.0 ± 0.6 | 5.2 ± 0.2 | 1.8 ± 0.6 |
| Tetrahydroneopterin | 11.9 ± 0.3 | 6.5 ± 0.6 | 3.5 ± 0.7 | 1.7 ± 0.4 |
| Thiamine | 5.6 ± 1.2 | 9.8 ± 0.8 | 4.3 ± 0.6 | 2.1 ± 0.4 |
| Thiamine monophosphate | 10.0 ± 1.2 | 7.6 ± 0.6 | 3.5 ± 0.7 | 1.9 ± 0.8 |
| Threonic acid | 7.2 ± 0.4 | 6.7 ± 0.8 | 5.8 ± 1.2 | 2.8 ± 0.3 |
| Threonine | 7.4 ± 0.9 | 12.8 ± 1.0 | 2.0 ± 1.0 | 4.1 ± 0.9 |
| Thymidine | 11.6 ± 0.2 | 11.0 ± 0.8 | 3.8 ± 0.5 | 2.3 ± 0.6 |
| Thymidine Monophosphate | 10.6 ± 0.8 | 9.8 ± 0.7 | 1.6 ± 1.1 | 4.7 ± 0.6 |
| Thymine | 11.4 ± 0.1 | 5.9 ± 0.3 | 3.4 ± 0.8 | 1.7 ± 0.5 |
| Trigonelleine | 11.7 ± 0.2 | 12.7 ± 1.0 | 3.3 ± 0.9 | 4.9 ± 0.9 |
| Trimethylamine N-oxide | 11.6 ± 0.1 | 8.6 ± 0.3 | 2.9 ± 0.8 | 2.0 ± 1.0 |
| Trimethyllysine | 7.7 ± 0.9 | 7.7 ± 0.5 | 3.2 ± 0.1 | 2.6 ± 1.2 |
| Tryptophan | 6.0 ± 0.2 | 5.8 ± 1.0 | 2.6 ± 1.1 | 3.8 ± 1.0 |
| Tyramine | 7.3 ± 1.0 | 6.7 ± 0.5 | 5.7 ± 0.5 | 2.0 ± 1.1 |
| Tyrosine | 8.4 ± 0.7 | 7.1 ± 0.8 | 4.8 ± 0.2 | 2.7 ± 0.5 |
| Uracil | 12.5 ± 0.8 | 9.5 ± 0.8 | 3.9 ± 1.0 | 3.5 ± 1.0 |
| Uridine 5'-diphosphate | 10.5 ± 0.6 | 8.3 ± 0.7 | 1.9 ± 0.3 | 3.8 ± 0.4 |
| Uridine Monophosphate | 11.8 ± 0.9 | 8.7 ± 0.8 | 4.6 ± 0.5 | 4.8 ± 0.1 |
| Uridine-5-Diphosphogalactose | 7.9 ± 0.2 | 11.7 ± 0.5 | 2.5 ± 0.6 | 2.4 ± 0.7 |
| Urocanic acid | 11.0 ± 1.1 | 6.4 ± 0.5 | 3.3 ± 0.2 | 3.9 ± 1.1 |
| UTP | 8.8 ± 1.1 | 5.4 ± 0.7 | 5.7 ± 0.2 | 2.3 ± 0.5 |
| Valine | 7.9 ± 0.3 | 7.4 ± 0.2 | 2.2 ± 1.1 | 2.7 ± 0.6 |
| XMP | 6.1 ± 1.1 | 11.0 ± 1.1 | 4.5 ± 0.1 | 3.6 ± 0.3 |
| Xylulose-5-Phosphate | 12.0 ± 0.9 | 6.4 ± 0.7 | 4.8 ± 0.4 | 4.4 ± 1.1 |

### **Table S14** | Intra – day precision (as % RSD) (Plasma extract as matrix, n = 5). The concentrations of QC_LLOQ_, QC_Low_, QC_Mid_ and QC_High_ were 10 ng/mL, 25 ng/mL, 100 ng/mL and 500 ng/mL respectively.

| Precision, % | QC_LLOQ_ | QC_low_ | QC_mid_ | QC_high_ |
| --- | --- | --- | --- | --- |
| 1-Methylguanine | 9.2 ± 0.8 | 2.6 ± 1.1 | 5.7 ± 1.2 | 4.5 ± 0.8 |
| 1-Methylhistidine | 10.1 ± 0.9 | 4.9 ± 0.5 | 2.7 ± 0.6 | 4.5 ± 0.3 |
| 1-Methylnicotinamide | 9.1 ± 1.1 | 2.4 ± 1.2 | 4.4 ± 0.6 | 5.4 ± 0.6 |
| 2-Aminoadipic Acid | 6.0 ± 0.5 | 5.7 ± 0.2 | 5.1 ± 1.1 | 4.2 ± 1.0 |
| 2-Deoxyadenosine | 10.7 ± 0.8 | 5.0 ± 0.6 | 4.3 ± 1.1 | 4.8 ± 1.1 |
| 2'-Deoxyadenosine 5'-monophosphate | 7.7 ± 0.3 | 3.5 ± 1.1 | 4.2 ± 1.0 | 2.9 ± 0.8 |
| 2'-deoxyadenosine 5'-triphosphate | 8.2 ± 1.0 | 3.9 ± 0.7 | 2.7 ± 0.7 | 4.2 ± 0.5 |
| 2'-Deoxycytidine-5'-monophosphate | 5.1 ± 0.9 | 4.5 ± 0.3 | 2.1 ± 0.7 | 2.2 ± 1.0 |
| 2-Deoxyguanosine | 7.9 ± 0.8 | 5.9 ± 0.6 | 2.8 ± 0.6 | 3.0 ± 1.1 |
| 2-hydroxybutyric acid | 5.5 ± 1.2 | 3.7 ± 0.3 | 5.9 ± 1.0 | 4.3 ± 0.3 |
| 2-hydroxyglutaric acid | 8.4 ± 0.7 | 2.9 ± 0.7 | 5.2 ± 0.9 | 4.5 ± 0.7 |
| 2-Hydroxyisocaproic acid | 6.6 ± 0.2 | 5.9 ± 0.2 | 4.2 ± 0.3 | 2.5 ± 0.4 |
| 2-ketobutyric acid | 5.5 ± 1.1 | 4.4 ± 0.9 | 2.2 ± 0.5 | 4.7 ± 0.1 |
| 2-oxoadipic acid | 8.4 ± 0.8 | 3.9 ± 0.5 | 4.6 ± 0.4 | 2.3 ± 0.3 |
| 2-Phenylglycine | 10.3 ± 0.7 | 5.6 ± 0.7 | 4.1 ± 0.7 | 2.1 ± 0.1 |
| 2-phosphoglycerate | 10.6 ± 1.0 | 5.8 ± 0.4 | 3.3 ± 0.4 | 3.7 ± 0.8 |
| 3-hydroxy-3-methyl-2-oxobutanoate | 8.7 ± 0.6 | 3.8 ± 0.2 | 5.6 ± 1.2 | 2.5 ± 0.7 |
| 3-hydroxybutyric acid | 10.9 ± 1.1 | 3.9 ± 1.1 | 4.4 ± 0.3 | 2.5 ± 0.7 |
| 3-Methylhistidine | 5.0 ± 0.6 | 5.6 ± 0.5 | 5.9 ± 1.1 | 2.1 ± 0.4 |
| 3-Phosphoglyceraldehyde | 7.8 ± 0.2 | 5.4 ± 0.7 | 4.2 ± 0.4 | 5.8 ± 0.4 |
| 3-phosphoglycerate | 10.8 ± 0.3 | 2.1 ± 1.2 | 5.9 ± 0.8 | 2.9 ± 0.5 |
| 4-amino-butyric acid | 10.9 ± 1.0 | 3.6 ± 0.7 | 4.4 ± 0.9 | 6.0 ± 1.1 |
| 4-aspartyl-phosphate | 9.1 ± 1.1 | 4.4 ± 0.6 | 2.6 ± 1.1 | 5.1 ± 0.6 |
| 4-Hydroxybenzoic acid | 9.1 ± 0.3 | 3.3 ± 0.8 | 3.2 ± 0.3 | 3.2 ± 0.7 |
| 4-Hydroxyphenylacetylglycine | 5.6 ± 0.6 | 5.7 ± 0.9 | 5.2 ± 0.2 | 5.8 ± 0.4 |
| 4-methyl-2-oxopentanoate | 8.7 ± 0.7 | 3.5 ± 0.4 | 4.3 ± 0.8 | 5.3 ± 1.1 |
| 4-Oxoglutaramate | 9.0 ± 0.2 | 4.9 ± 0.4 | 4.5 ± 0.4 | 4.4 ± 0.9 |
| 5-Hydroxyindolepyruvate | 9.0 ± 0.4 | 2.8 ± 0.6 | 3.1 ± 0.2 | 3.5 ± 0.2 |
| 5-Hydroxylysine | 9.2 ± 0.4 | 4.6 ± 0.1 | 4.5 ± 0.6 | 3.5 ± 0.2 |
| 5-Hydroxy-Tryptophan | 9.1 ± 0.6 | 5.4 ± 0.6 | 3.6 ± 1.2 | 5.0 ± 0.6 |
| 5-Phosphoribosylamine | 8.7 ± 0.4 | 5.0 ± 0.6 | 4.7 ± 1.1 | 4.0 ± 0.9 |
| 6-phosphogluconate | 10.4 ± 0.2 | 4.4 ± 0.5 | 5.3 ± 0.3 | 4.8 ± 0.4 |
| Acetylcarnitine | 8.0 ± 0.8 | 4.7 ± 0.6 | 4.5 ± 0.9 | 4.8 ± 0.9 |
| Aconitic Acid | 7.8 ± 0.7 | 5.5 ± 1.0 | 3.4 ± 0.8 | 4.2 ± 0.5 |
| Adenine | 7.4 ± 0.9 | 3.2 ± 0.3 | 5.3 ± 0.9 | 5.5 ± 1.1 |
| Adenosine | 7.9 ± 0.7 | 4.0 ± 0.1 | 4.5 ± 1.0 | 5.7 ± 0.9 |
| Adenosine 3',5'-cyclic monophosphate | 6.6 ± 0.1 | 6.0 ± 0.7 | 3.7 ± 0.9 | 4.6 ± 0.8 |
| Adenosine Monophosphate | 5.7 ± 0.5 | 2.5 ± 1.0 | 2.6 ± 0.6 | 4.6 ± 0.8 |
| Adenosine-5'-diphosphate | 10.9 ± 0.8 | 3.3 ± 0.9 | 2.5 ± 0.8 | 4.8 ± 0.3 |
| Adenosine-5'-triphosphate | 8.5 ± 0.5 | 5.9 ± 0.1 | 5.1 ± 0.6 | 2.0 ± 0.7 |
| Adenylosuccinate | 8.4 ± 0.8 | 3.9 ± 0.5 | 5.7 ± 0.5 | 3.5 ± 0.6 |
| Alanine | 10.0 ± 1.2 | 4.4 ± 0.3 | 5.9 ± 0.9 | 2.8 ± 0.1 |
| alpha-ketoglutaric acid | 5.2 ± 1.0 | 4.4 ± 0.8 | 2.2 ± 0.3 | 3.6 ± 0.3 |
| Anserine | 10.2 ± 0.6 | 4.6 ± 0.9 | 4.5 ± 0.7 | 4.2 ± 0.5 |
| Arginine | 9.9 ± 0.6 | 4.3 ± 0.4 | 3.7 ± 0.7 | 4.4 ± 0.9 |
| Argininosuccinic Acid | 10.9 ± 0.5 | 6.0 ± 0.6 | 4.9 ± 0.3 | 4.9 ± 0.2 |
| Asparagine | 5.6 ± 0.1 | 2.2 ± 1.2 | 3.3 ± 1.2 | 4.8 ± 1.2 |
| Aspartic acid | 10.2 ± 0.6 | 5.1 ± 0.3 | 3.4 ± 0.2 | 5.4 ± 0.6 |
| Asymmetric dimethylarginine | 6.2 ± 0.4 | 2.3 ± 0.5 | 4.5 ± 1.0 | 5.7 ± 0.9 |
| Azelaic acid | 7.8 ± 0.8 | 3.3 ± 0.6 | 3.1 ± 0.8 | 4.4 ± 0.3 |
| beta-Alanine | 7.1 ± 0.7 | 2.4 ± 0.6 | 2.9 ± 0.4 | 4.4 ± 0.7 |
| Betaine | 11.0 ± 0.6 | 3.6 ± 0.3 | 2.1 ± 0.9 | 5.6 ± 0.6 |
| Biopterin | 9.6 ± 0.2 | 5.3 ± 0.8 | 5.2 ± 1.1 | 2.6 ± 0.2 |
| Biotin | 10.4 ± 0.4 | 2.1 ± 1.2 | 5.8 ± 0.3 | 2.0 ± 0.3 |
| Carnitine | 7.2 ± 0.8 | 2.9 ± 1.0 | 4.5 ± 0.9 | 3.1 ± 0.2 |
| Carnosine | 6.1 ± 0.5 | 2.2 ± 1.1 | 3.4 ± 0.7 | 2.8 ± 0.2 |
| CDP-choline | 7.8 ± 0.3 | 4.3 ± 0.7 | 4.2 ± 1.0 | 2.9 ± 0.6 |
| cGMP | 9.6 ± 0.3 | 4.8 ± 0.2 | 2.3 ± 1.1 | 3.0 ± 0.1 |
| Choline | 5.6 ± 1.1 | 2.4 ± 0.7 | 3.9 ± 0.4 | 4.5 ± 0.9 |
| Cinnamic acid | 6.2 ± 0.3 | 4.9 ± 0.6 | 2.8 ± 0.8 | 3.4 ± 0.5 |
| Citraconic acid | 5.2 ± 0.2 | 4.1 ± 0.4 | 4.2 ± 1.1 | 5.5 ± 1.2 |
| Citramalic acid | 8.0 ± 0.3 | 5.5 ± 0.5 | 4.6 ± 0.7 | 5.9 ± 0.6 |
| Citric Acid | 8.6 ± 0.4 | 5.8 ± 0.2 | 4.9 ± 0.4 | 4.0 ± 0.6 |
| Citrulline | 5.2 ± 0.9 | 3.6 ± 0.7 | 3.7 ± 1.0 | 2.9 ± 0.8 |
| Creatine | 7.5 ± 0.9 | 5.8 ± 0.5 | 3.7 ± 0.6 | 3.4 ± 0.2 |
| Creatinine | 5.8 ± 0.3 | 4.9 ± 0.5 | 6.0 ± 0.3 | 4.9 ± 0.4 |
| Cystathionine | 8.9 ± 0.7 | 5.5 ± 0.5 | 4.7 ± 0.1 | 5.3 ± 0.7 |
| Cysteamine | 5.8 ± 0.2 | 2.6 ± 1.0 | 3.1 ± 0.1 | 5.7 ± 0.8 |
| Cysteine | 7.2 ± 0.2 | 5.7 ± 0.2 | 4.7 ± 1.1 | 4.3 ± 0.1 |
| Cysteinylglycine | 8.2 ± 0.6 | 4.2 ± 1.0 | 5.1 ± 1.1 | 4.8 ± 0.4 |
| Cystine | 5.0 ± 0.8 | 3.5 ± 0.9 | 4.5 ± 1.2 | 5.7 ± 1.2 |
| Cytidine | 8.8 ± 0.1 | 4.6 ± 0.6 | 3.3 ± 0.5 | 3.6 ± 0.3 |
| Cytidine 3',5'-cyclic monophosphate | 10.1 ± 0.7 | 3.5 ± 1.1 | 4.3 ± 0.7 | 4.8 ± 0.7 |
| Cytidine 5'-diphosphate | 8.6 ± 0.9 | 5.3 ± 0.5 | 3.6 ± 0.1 | 4.4 ± 0.2 |
| Cytidine 5'-triphosphate | 9.1 ± 1.0 | 3.1 ± 0.1 | 5.6 ± 0.6 | 2.2 ± 1.0 |
| Cytidine monophosphate | 9.7 ± 0.4 | 3.1 ± 0.9 | 3.6 ± 0.7 | 5.3 ± 0.2 |
| Cytosine | 9.8 ± 0.5 | 2.6 ± 1.0 | 5.7 ± 0.9 | 4.9 ± 1.1 |
| dADP | 5.4 ± 1.0 | 4.3 ± 0.8 | 2.7 ± 0.6 | 4.5 ± 1.1 |
| Diethanolamine | 6.3 ± 0.9 | 4.8 ± 1.1 | 5.6 ± 0.6 | 2.7 ± 0.5 |
| Dihydrobiopterin | 6.6 ± 0.8 | 5.3 ± 0.2 | 3.3 ± 0.4 | 3.0 ± 1.1 |
| Dihydroneopterin triphosphate | 7.6 ± 0.9 | 3.2 ± 0.3 | 4.8 ± 1.1 | 4.4 ± 0.2 |
| Dihydroxyacetonephosphate | 9.2 ± 0.3 | 2.0 ± 0.6 | 6.0 ± 0.5 | 5.2 ± 1.0 |
| Dimethylglycine | 7.3 ± 0.4 | 5.3 ± 1.0 | 2.9 ± 0.6 | 2.8 ± 0.6 |
| Ectoine | 9.3 ± 0.8 | 5.2 ± 0.6 | 4.8 ± 0.9 | 4.3 ± 1.0 |
| Erythrose-4-phosphate | 7.7 ± 0.6 | 5.3 ± 0.2 | 3.4 ± 0.3 | 3.7 ± 1.0 |
| FAD | 7.8 ± 0.3 | 2.4 ± 0.7 | 2.0 ± 0.7 | 2.1 ± 1.2 |
| Folic acid | 10.5 ± 0.9 | 3.9 ± 0.4 | 6.0 ± 0.4 | 5.2 ± 0.8 |
| Fructose-6-phosphate | 10.9 ± 0.9 | 4.8 ± 0.3 | 3.1 ± 1.1 | 4.5 ± 0.5 |
| Fumaric Acid | 8.6 ± 0.6 | 2.7 ± 0.7 | 3.9 ± 0.4 | 5.1 ± 0.4 |
| Galactosamine | 9.6 ± 0.3 | 3.8 ± 0.2 | 3.9 ± 0.1 | 5.8 ± 0.3 |
| Galactose-1-phosphate | 8.8 ± 0.6 | 2.4 ± 0.1 | 2.3 ± 0.5 | 4.9 ± 0.9 |
| Galactose-6-Phosphate | 10.3 ± 0.2 | 2.2 ± 0.8 | 4.5 ± 0.2 | 2.8 ± 1.1 |
| gamma-Glutamylcysteine | 7.3 ± 0.3 | 3.0 ± 0.1 | 5.4 ± 0.9 | 5.6 ± 0.6 |
| Glucosamine | 5.2 ± 0.7 | 3.7 ± 0.8 | 3.5 ± 0.3 | 4.8 ± 0.7 |
| Glucosamine-6-Phosphate | 7.0 ± 0.9 | 2.4 ± 0.6 | 4.3 ± 1.0 | 5.8 ± 0.7 |
| Glucose | 8.2 ± 0.8 | 5.9 ± 1.0 | 4.8 ± 1.0 | 5.8 ± 0.2 |
| Glucose-1-phosphate | 10.2 ± 0.6 | 3.6 ± 0.6 | 4.0 ± 0.9 | 2.7 ± 0.6 |
| Glucose-6-Phosphate | 8.8 ± 0.6 | 4.8 ± 1.1 | 2.1 ± 0.4 | 2.6 ± 1.2 |
| Glucuronic acid | 8.9 ± 0.9 | 6.0 ± 0.9 | 4.2 ± 0.5 | 5.0 ± 0.8 |
| Glutamic acid | 9.8 ± 1.0 | 2.1 ± 0.7 | 4.7 ± 0.6 | 5.0 ± 1.2 |
| Glutamine | 10.6 ± 1.1 | 3.8 ± 1.2 | 5.3 ± 1.0 | 5.5 ± 0.1 |
| Glutathione | 7.6 ± 1.1 | 4.4 ± 0.5 | 4.4 ± 0.7 | 3.8 ± 0.4 |
| Glutathionylspermine | 9.8 ± 0.1 | 3.9 ± 1.2 | 2.5 ± 1.2 | 4.1 ± 0.9 |
| Glycerophosphocholine | 6.2 ± 0.7 | 2.4 ± 0.5 | 4.2 ± 0.5 | 4.6 ± 0.7 |
| Glycine | 8.8 ± 0.3 | 3.2 ± 0.5 | 5.4 ± 0.3 | 4.2 ± 0.7 |
| Glycolic acid | 8.7 ± 0.1 | 3.6 ± 1.1 | 2.5 ± 0.4 | 2.7 ± 0.6 |
| Guanidinoacetic acid | 7.3 ± 1.1 | 5.7 ± 0.3 | 2.9 ± 0.9 | 3.0 ± 0.3 |
| Guanidinobutanoic acid | 7.5 ± 0.9 | 2.9 ± 1.1 | 5.2 ± 1.0 | 3.6 ± 0.9 |
| Guanine | 7.7 ± 0.8 | 6.0 ± 1.1 | 5.8 ± 1.0 | 2.7 ± 1.0 |
| Guanosine | 9.5 ± 0.6 | 2.1 ± 0.5 | 5.8 ± 0.8 | 3.5 ± 1.0 |
| Guanosine 5'-diphosphate | 9.6 ± 0.6 | 5.6 ± 0.5 | 5.6 ± 0.7 | 4.7 ± 0.9 |
| Guanosine-5'-monophosphate | 9.4 ± 0.4 | 5.2 ± 0.8 | 5.1 ± 0.1 | 4.8 ± 0.7 |
| Hercynine | 10.6 ± 0.3 | 4.2 ± 1.1 | 2.0 ± 0.8 | 2.5 ± 0.8 |
| Hippuric acid | 7.8 ± 0.9 | 3.1 ± 0.4 | 3.3 ± 1.2 | 3.0 ± 0.5 |
| Histamine | 9.9 ± 0.6 | 3.2 ± 1.0 | 3.7 ± 0.4 | 2.5 ± 0.4 |
| Histidine | 8.2 ± 0.9 | 5.0 ± 0.9 | 3.5 ± 0.9 | 2.2 ± 0.3 |
| Homoalanine | 10.5 ± 0.8 | 4.9 ± 0.4 | 4.6 ± 0.4 | 3.7 ± 0.9 |
| Homoarginine | 9.1 ± 0.2 | 5.5 ± 0.3 | 4.0 ± 0.2 | 3.5 ± 0.2 |
| Homocysteine | 6.5 ± 1.1 | 2.1 ± 0.4 | 3.9 ± 0.2 | 3.6 ± 0.7 |
| Homogentisic acid | 5.9 ± 0.7 | 3.7 ± 0.7 | 4.1 ± 1.1 | 5.7 ± 0.1 |
| Homoleucine | 9.9 ± 1.1 | 3.1 ± 0.7 | 2.1 ± 1.1 | 4.7 ± 0.5 |
| Homovanillic acid | 9.9 ± 0.4 | 3.5 ± 0.5 | 2.9 ± 0.8 | 3.5 ± 1.0 |
| Hydantoin-5-propionate | 5.1 ± 0.1 | 4.4 ± 1.1 | 3.0 ± 0.8 | 5.1 ± 0.5 |
| Hypotaurine | 10.5 ± 0.8 | 3.9 ± 0.6 | 2.6 ± 0.6 | 5.3 ± 0.9 |
| Hypoxanthine | 8.9 ± 0.2 | 5.1 ± 0.9 | 2.7 ± 0.7 | 4.3 ± 0.3 |
| Indole-3-acetamide | 10.4 ± 0.5 | 4.6 ± 0.9 | 2.0 ± 1.1 | 2.2 ± 0.1 |
| Inosine | 6.0 ± 1.0 | 4.6 ± 0.2 | 3.1 ± 0.2 | 5.6 ± 0.9 |
| Inosine 5'-monophosphate | 6.4 ± 0.6 | 3.5 ± 1.1 | 3.1 ± 0.5 | 4.3 ± 0.6 |
| Isocitric Acid | 7.3 ± 0.7 | 5.5 ± 0.7 | 4.3 ± 0.5 | 4.7 ± 0.3 |
| Isoleucine | 5.5 ± 0.7 | 2.2 ± 0.8 | 5.7 ± 0.5 | 5.6 ± 1.0 |
| Lactic Acid | 6.3 ± 0.4 | 3.1 ± 0.2 | 3.6 ± 1.2 | 5.1 ± 1.2 |
| Leucine | 9.2 ± 0.7 | 3.4 ± 1.2 | 6.0 ± 1.2 | 5.8 ± 0.6 |
| Levulinic acid | 8.2 ± 0.5 | 4.9 ± 1.0 | 3.4 ± 0.8 | 4.2 ± 0.9 |
| Lysine | 6.2 ± 1.1 | 4.7 ± 0.8 | 5.8 ± 1.1 | 2.0 ± 0.8 |
| Malic Acid | 9.7 ± 1.0 | 3.7 ± 0.3 | 2.1 ± 0.9 | 5.4 ± 1.0 |
| Mannose-1 with Mannose-6-Phosphate | 6.6 ± 0.7 | 3.2 ± 0.7 | 5.0 ± 1.0 | 3.1 ± 0.2 |
| Mesaconic acid | 9.0 ± 1.1 | 5.4 ± 0.9 | 5.1 ± 0.4 | 3.5 ± 1.1 |
| Methionine | 10.7 ± 0.3 | 5.6 ± 0.5 | 5.5 ± 0.6 | 3.7 ± 0.5 |
| Methionine sulfoxide | 10.8 ± 0.8 | 2.8 ± 1.0 | 2.9 ± 0.4 | 2.0 ± 0.7 |
| Methylmalonic Acid | 7.5 ± 0.2 | 2.1 ± 0.5 | 5.1 ± 0.5 | 3.5 ± 0.8 |
| N1-Acetylspermidine | 7.0 ± 0.7 | 5.2 ± 0.1 | 2.9 ± 1.1 | 2.5 ± 0.3 |
| N1-Acetylspermine | 8.9 ± 1.1 | 5.9 ± 0.9 | 4.3 ± 0.5 | 3.1 ± 0.6 |
| N8-Acetylspermidine | 8.3 ± 0.8 | 5.3 ± 1.2 | 3.9 ± 0.2 | 4.4 ± 1.1 |
| N-Acetyl-Aspartyl-Glutamate | 6.0 ± 0.9 | 4.6 ± 0.4 | 6.0 ± 0.4 | 3.8 ± 0.4 |
| N-Acetyl-Aspartyl-Glutamyl-Glutamate | 10.0 ± 1.1 | 3.8 ± 0.2 | 3.8 ± 0.3 | 4.9 ± 1.1 |
| N-Acetyl-Galactosamine | 8.7 ± 1.0 | 4.2 ± 0.1 | 4.0 ± 1.0 | 4.7 ± 0.6 |
| N-Acetyl-Leucine | 10.8 ± 0.1 | 4.9 ± 0.6 | 4.6 ± 0.9 | 2.1 ± 0.7 |
| N-Acetyl-Methionine | 8.8 ± 0.9 | 2.1 ± 0.3 | 3.1 ± 0.6 | 5.9 ± 0.9 |
| N-Acetylneuraminic acid | 7.7 ± 0.9 | 4.3 ± 0.1 | 3.5 ± 0.3 | 2.1 ± 0.6 |
| N-Acetyl-Proline | 9.5 ± 0.1 | 3.6 ± 0.9 | 2.4 ± 0.8 | 5.4 ± 0.7 |
| N-Acetyl-Putrescine | 5.8 ± 0.3 | 3.6 ± 0.9 | 4.7 ± 1.0 | 2.5 ± 1.0 |
| N-Acetyl-Serine | 10.5 ± 0.4 | 2.9 ± 1.1 | 4.2 ± 0.5 | 5.7 ± 1.2 |
| N-Acetyl-Serotonin | 7.2 ± 1.2 | 2.7 ± 0.4 | 4.2 ± 1.0 | 5.8 ± 0.7 |
| N-Acyl-Homoserine-lactone | 8.3 ± 1.0 | 4.5 ± 0.3 | 4.9 ± 0.7 | 2.1 ± 0.7 |
| NAD | 6.2 ± 0.9 | 3.2 ± 1.0 | 2.4 ± 0.9 | 5.5 ± 0.3 |
| NADH | 8.2 ± 0.6 | 3.2 ± 0.5 | 4.1 ± 0.9 | 2.7 ± 0.5 |
| NADP | 6.5 ± 0.8 | 3.6 ± 0.8 | 5.1 ± 0.1 | 5.5 ± 0.9 |
| NADPH | 7.5 ± 1.0 | 2.3 ± 0.1 | 3.8 ± 0.9 | 2.2 ± 0.9 |
| N-alpha-Acetyl-lysine | 8.2 ± 0.6 | 5.0 ± 0.6 | 3.3 ± 0.4 | 4.9 ± 1.1 |
| N-alpha-Acetyl-Ornithine | 5.4 ± 0.5 | 3.7 ± 0.7 | 4.0 ± 1.1 | 4.8 ± 0.8 |
| N-Carbamoyl-Proline | 10.4 ± 1.0 | 5.8 ± 0.1 | 5.0 ± 0.7 | 5.6 ± 0.1 |
| N-Carbamoyl-Sarcosine | 9.8 ± 0.9 | 4.4 ± 0.4 | 3.3 ± 0.8 | 5.6 ± 1.0 |
| N-Epsilon-Acetyl-lysine | 10.2 ± 0.9 | 2.9 ± 0.5 | 4.7 ± 0.7 | 2.7 ± 0.6 |
| Nicotinamide | 5.4 ± 0.8 | 2.5 ± 0.4 | 3.8 ± 1.1 | 5.0 ± 1.0 |
| N-Methyl-Aspartic Acid | 8.7 ± 0.4 | 5.4 ± 0.6 | 3.8 ± 0.2 | 3.0 ± 0.5 |
| O-Phosphoethanolamine | 5.8 ± 0.3 | 4.7 ± 0.1 | 4.2 ± 0.5 | 3.9 ± 0.4 |
| Ophthalmic acid | 7.9 ± 0.6 | 2.8 ± 0.8 | 4.4 ± 0.7 | 3.5 ± 0.9 |
| Ornithine | 9.2 ± 1.0 | 5.9 ± 0.5 | 2.8 ± 0.9 | 3.5 ± 1.1 |
| Orotidine 5'-monophosphate | 8.2 ± 0.2 | 2.8 ± 1.0 | 5.3 ± 0.5 | 2.5 ± 0.4 |
| Oxaloacetate | 10.6 ± 0.4 | 4.7 ± 0.2 | 3.8 ± 1.1 | 2.3 ± 0.4 |
| Pantothenic acid | 9.7 ± 0.3 | 5.1 ± 1.1 | 2.0 ± 0.9 | 2.2 ± 0.4 |
| Phenylalanine | 8.7 ± 0.6 | 2.2 ± 0.1 | 2.4 ± 0.2 | 3.5 ± 0.5 |
| Phenylpyruvate | 6.9 ± 0.2 | 3.4 ± 0.2 | 3.7 ± 0.4 | 2.7 ± 1.0 |
| Phosphodimethylethanolamine | 9.5 ± 0.1 | 3.0 ± 0.3 | 3.6 ± 0.3 | 3.4 ± 1.1 |
| Phosphoenolpyruvate | 5.4 ± 0.9 | 5.4 ± 0.7 | 5.1 ± 1.1 | 5.7 ± 1.1 |
| Phosphorylcholine | 6.8 ± 0.4 | 3.3 ± 0.4 | 3.7 ± 0.7 | 2.2 ± 0.7 |
| Pipecolic acid | 9.1 ± 0.2 | 3.0 ± 0.3 | 2.3 ± 0.8 | 2.2 ± 0.7 |
| Proline | 8.0 ± 0.6 | 5.7 ± 0.4 | 5.6 ± 0.2 | 3.5 ± 0.2 |
| Putrescine | 8.4 ± 0.8 | 3.7 ± 0.6 | 4.1 ± 0.6 | 3.8 ± 0.9 |
| Pyridoxal | 5.7 ± 0.6 | 4.4 ± 0.8 | 2.3 ± 0.8 | 5.3 ± 0.8 |
| Pyridoxamine | 7.8 ± 0.5 | 3.9 ± 0.5 | 4.4 ± 0.3 | 3.7 ± 0.7 |
| Pyridoxine | 7.9 ± 0.1 | 5.5 ± 0.6 | 3.3 ± 0.4 | 4.6 ± 0.8 |
| Pyroglutamic acid | 7.6 ± 0.7 | 5.3 ± 0.4 | 2.6 ± 1.0 | 2.7 ± 0.4 |
| Pyrrolysine | 7.4 ± 0.4 | 2.9 ± 0.9 | 2.0 ± 0.2 | 5.8 ± 0.7 |
| Pyruvic Acid | 10.3 ± 1.2 | 2.6 ± 0.3 | 4.1 ± 1.0 | 2.6 ± 0.2 |
| Riboflavin | 10.2 ± 0.5 | 4.1 ± 0.8 | 4.6 ± 0.5 | 3.4 ± 0.7 |
| Ribose-5-Phosphate | 10.6 ± 1.1 | 5.4 ± 0.1 | 4.5 ± 0.3 | 4.4 ± 0.2 |
| Ribulose-1-Phosphate | 9.0 ± 0.6 | 2.3 ± 0.4 | 2.9 ± 0.5 | 2.9 ± 0.2 |
| Ribulose-5-Phosphate | 8.0 ± 0.6 | 2.8 ± 0.2 | 4.4 ± 0.9 | 6.0 ± 0.3 |
| Saccharopine | 10.9 ± 1.2 | 4.2 ± 0.3 | 5.5 ± 0.8 | 2.9 ± 1.1 |
| S-Adenosylhomocysteine | 6.5 ± 0.4 | 4.8 ± 0.8 | 4.0 ± 0.6 | 2.5 ± 0.7 |
| S-Adenosylmethioninamine | 8.6 ± 1.2 | 4.7 ± 0.3 | 2.4 ± 1.0 | 3.7 ± 0.7 |
| S-Adenosylmethionine | 6.7 ± 0.9 | 5.3 ± 0.5 | 2.7 ± 1.0 | 4.1 ± 1.0 |
| Sarcosine | 7.8 ± 1.1 | 5.8 ± 0.9 | 5.1 ± 0.8 | 4.3 ± 0.7 |
| Sedoheptulose-7-phosphate | 8.4 ± 0.8 | 3.9 ± 0.6 | 2.5 ± 0.4 | 5.7 ± 0.6 |
| Selenohomocysteine | 8.1 ± 0.6 | 3.3 ± 0.9 | 2.2 ± 0.3 | 5.4 ± 0.6 |
| Selenomethionine | 5.7 ± 0.4 | 5.3 ± 0.5 | 4.3 ± 0.3 | 5.0 ± 0.8 |
| Serine | 5.2 ± 0.6 | 4.2 ± 0.4 | 2.6 ± 0.6 | 2.6 ± 0.8 |
| S-Methyl-Thioinosine | 5.6 ± 1.0 | 4.0 ± 0.6 | 3.8 ± 0.4 | 4.1 ± 1.2 |
| Spermidine | 9.4 ± 0.3 | 3.2 ± 0.7 | 5.6 ± 0.6 | 2.4 ± 1.0 |
| Spermine | 7.3 ± 0.1 | 3.3 ± 0.6 | 5.1 ± 1.2 | 2.2 ± 0.5 |
| Succinate | 9.2 ± 0.9 | 5.2 ± 0.7 | 2.8 ± 0.8 | 2.4 ± 0.4 |
| Sucrose | 9.0 ± 0.4 | 2.6 ± 1.1 | 2.1 ± 0.9 | 5.3 ± 0.7 |
| Symmetric dimethylarginine | 10.7 ± 0.2 | 5.7 ± 0.5 | 4.4 ± 0.8 | 3.3 ± 0.5 |
| Taurine | 5.3 ± 0.6 | 5.6 ± 1.1 | 5.6 ± 0.3 | 3.8 ± 0.9 |
| Tetrahydroneopterin | 10.5 ± 0.8 | 3.0 ± 1.2 | 5.0 ± 0.6 | 2.3 ± 0.8 |
| Thiamine | 5.6 ± 0.9 | 3.9 ± 0.7 | 2.6 ± 0.7 | 3.9 ± 0.2 |
| Thiamine monophosphate | 5.7 ± 1.1 | 5.9 ± 0.9 | 4.9 ± 0.2 | 5.3 ± 1.0 |
| Threonic acid | 10.3 ± 0.8 | 4.6 ± 0.4 | 5.8 ± 1.0 | 4.4 ± 1.1 |
| Threonine | 6.8 ± 0.8 | 2.5 ± 0.8 | 3.1 ± 1.1 | 3.7 ± 0.3 |
| Thymidine | 10.2 ± 0.6 | 2.6 ± 1.0 | 2.2 ± 0.9 | 2.6 ± 0.4 |
| Thymidine Monophosphate | 10.4 ± 0.6 | 2.9 ± 0.5 | 5.0 ± 1.0 | 5.5 ± 0.3 |
| Thymine | 6.3 ± 0.6 | 5.1 ± 0.2 | 3.4 ± 1.1 | 2.2 ± 1.0 |
| Trigonelleine | 9.0 ± 0.1 | 3.3 ± 0.8 | 2.5 ± 0.3 | 4.4 ± 1.0 |
| Trimethylamine N-oxide | 7.2 ± 0.7 | 2.5 ± 0.2 | 3.8 ± 0.5 | 2.1 ± 0.2 |
| Trimethyllysine | 6.8 ± 0.2 | 6.0 ± 0.9 | 4.1 ± 0.6 | 3.9 ± 0.1 |
| Tryptophan | 10.0 ± 0.6 | 3.5 ± 0.8 | 2.8 ± 0.8 | 5.9 ± 0.6 |
| Tyramine | 7.7 ± 0.7 | 3.8 ± 0.7 | 5.5 ± 0.6 | 5.5 ± 0.3 |
| Tyrosine | 10.4 ± 1.0 | 4.4 ± 0.9 | 2.1 ± 0.7 | 4.3 ± 0.5 |
| Uracil | 10.8 ± 1.1 | 3.2 ± 1.1 | 3.6 ± 0.5 | 2.1 ± 1.1 |
| Uridine 5'-diphosphate | 10.8 ± 0.5 | 5.7 ± 0.7 | 2.3 ± 0.7 | 5.4 ± 0.9 |
| Uridine Monophosphate | 5.6 ± 0.4 | 2.1 ± 0.1 | 4.7 ± 0.9 | 4.3 ± 0.4 |
| Uridine-5-Diphosphogalactose | 7.7 ± 0.7 | 2.3 ± 0.4 | 5.6 ± 0.5 | 2.3 ± 1.0 |
| Urocanic acid | 7.4 ± 0.7 | 3.0 ± 1.0 | 4.4 ± 0.9 | 2.6 ± 1.1 |
| UTP | 6.1 ± 1.2 | 5.2 ± 1.0 | 3.6 ± 1.1 | 4.0 ± 0.5 |
| Valine | 7.4 ± 0.5 | 4.0 ± 0.4 | 4.3 ± 0.2 | 5.6 ± 0.8 |
| XMP | 5.2 ± 0.7 | 4.4 ± 1.2 | 2.2 ± 0.6 | 5.2 ± 0.6 |
| Xylulose-5-Phosphate | 9.7 ± 0.5 | 4.2 ± 1.2 | 3.8 ± 0.5 | 2.5 ± 0.2 |

### **Table S15** | Intra – day precision (as % RSD) (Cell extract as matrix, n = 5). The concentrations of QC_LLOQ_, QC_Low_, QC_Mid_ and QC_High_ were 10 ng/mL, 25 ng/mL, 100 ng/mL and 500 ng/mL respectively.

| Precision, % | QC_LLOQ_ | QC_low_ | QC_mid_ | QC_high_ |
| --- | --- | --- | --- | --- |
| 1-Methylguanine | 8.3 ± 0.5 | 2.6 ± 0.5 | 2.9 ± 0.5 | 2.1 ± 0.3 |
| 1-Methylhistidine | 4.5 ± 0.5 | 5.5 ± 0.9 | 3.8 ± 0.4 | 4.7 ± 0.4 |
| 1-Methylnicotinamide | 8.0 ± 1.2 | 6.0 ± 0.9 | 2.1 ± 0.6 | 5.7 ± 0.1 |
| 2-Aminoadipic Acid | 10.1 ± 0.2 | 4.1 ± 0.6 | 2.3 ± 1.0 | 4.1 ± 0.2 |
| 2-Deoxyadenosine | 8.1 ± 1.2 | 1.3 ± 0.6 | 4.1 ± 0.5 | 3.7 ± 0.7 |
| 2'-Deoxyadenosine 5'-monophosphate | 2.1 ± 0.8 | 5.4 ± 1.0 | 5.9 ± 1.1 | 1.8 ± 0.6 |
| 2'-deoxyadenosine 5'-triphosphate | 5.1 ± 0.8 | 3.9 ± 0.5 | 5.0 ± 0.3 | 2.9 ± 0.3 |
| 2'-Deoxycytidine-5'-monophosphate | 1.3 ± 0.5 | 6.5 ± 0.1 | 6.3 ± 1.2 | 1.4 ± 0.3 |
| 2-Deoxyguanosine | 4.8 ± 0.8 | 6.4 ± 0.9 | 5.5 ± 0.8 | 5.5 ± 0.8 |
| 2-hydroxybutyric acid | 6.1 ± 0.4 | 4.1 ± 0.9 | 2.9 ± 1.2 | 4.5 ± 0.2 |
| 2-hydroxyglutaric acid | 8.1 ± 0.8 | 5.4 ± 0.3 | 3.4 ± 0.2 | 4.7 ± 0.5 |
| 2-Hydroxyisocaproic acid | 3.9 ± 0.4 | 4.5 ± 0.9 | 5.2 ± 0.9 | 4.1 ± 0.9 |
| 2-ketobutyric acid | 2.2 ± 0.4 | 3.3 ± 0.9 | 2.4 ± 0.4 | 5.8 ± 0.5 |
| 2-oxoadipic acid | 10.4 ± 0.8 | 3.4 ± 1.2 | 2.0 ± 0.5 | 1.9 ± 0.7 |
| 2-Phenylglycine | 3.2 ± 1.2 | 6.0 ± 0.6 | 5.0 ± 0.2 | 2.2 ± 0.8 |
| 2-phosphoglycerate | 3.6 ± 0.2 | 4.2 ± 1.0 | 1.4 ± 1.0 | 5.7 ± 0.7 |
| 3-hydroxy-3-methyl-2-oxobutanoate | 9.0 ± 1.2 | 2.6 ± 0.8 | 4.1 ± 0.3 | 3.6 ± 0.7 |
| 3-hydroxybutyric acid | 3.5 ± 0.5 | 5.9 ± 0.2 | 1.3 ± 0.2 | 6.2 ± 0.7 |
| 3-Methylhistidine | 4.0 ± 0.6 | 1.8 ± 0.3 | 4.1 ± 1.2 | 2.8 ± 0.3 |
| 3-Phosphoglyceraldehyde | 3.3 ± 0.1 | 5.7 ± 1.1 | 1.5 ± 1.2 | 6.1 ± 0.4 |
| 3-phosphoglycerate | 9.4 ± 1.2 | 4.2 ± 1.2 | 3.4 ± 0.9 | 5.0 ± 0.4 |
| 4-amino-butyric acid | 1.5 ± 0.6 | 2.4 ± 0.3 | 5.8 ± 0.8 | 2.0 ± 0.4 |
| 4-aspartyl-phosphate | 7.3 ± 0.3 | 3.1 ± 0.1 | 2.3 ± 1.1 | 3.9 ± 1.2 |
| 4-Hydroxybenzoic acid | 8.2 ± 0.7 | 3.2 ± 1.0 | 4.9 ± 0.5 | 4.8 ± 0.8 |
| 4-Hydroxyphenylacetylglycine | 8.4 ± 0.5 | 4.6 ± 0.5 | 5.5 ± 0.9 | 5.4 ± 0.6 |
| 4-methyl-2-oxopentanoate | 7.7 ± 0.4 | 5.9 ± 0.3 | 1.9 ± 0.8 | 3.6 ± 0.3 |
| 4-Oxoglutaramate | 8.1 ± 0.1 | 2.9 ± 0.3 | 5.8 ± 0.5 | 5.4 ± 0.2 |
| 5-Hydroxyindolepyruvate | 2.8 ± 0.8 | 6.0 ± 0.8 | 6.1 ± 1.0 | 4.1 ± 0.2 |
| 5-Hydroxylysine | 3.0 ± 0.2 | 4.0 ± 0.2 | 3.7 ± 0.9 | 5.3 ± 0.7 |
| 5-Hydroxy-Tryptophan | 9.5 ± 0.2 | 4.3 ± 0.1 | 6.0 ± 0.4 | 1.5 ± 0.9 |
| 5-Phosphoribosylamine | 11.8 ± 0.9 | 5.9 ± 0.2 | 5.2 ± 1.0 | 4.0 ± 1.1 |
| 6-phosphogluconate | 8.7 ± 0.9 | 2.5 ± 1.2 | 2.1 ± 1.1 | 2.1 ± 0.4 |
| Acetylcarnitine | 2.1 ± 0.3 | 4.3 ± 0.4 | 2.6 ± 1.0 | 5.0 ± 0.6 |
| Aconitic Acid | 9.5 ± 1.0 | 2.9 ± 0.8 | 1.5 ± 0.1 | 2.1 ± 0.3 |
| Adenine | 10.4 ± 0.6 | 2.1 ± 0.8 | 2.8 ± 0.4 | 1.2 ± 0.5 |
| Adenosine | 6.8 ± 1.2 | 6.4 ± 0.1 | 6.1 ± 0.5 | 4.8 ± 0.9 |
| Adenosine 3',5'-cyclic monophosphate | 9.5 ± 0.2 | 3.5 ± 0.3 | 3.8 ± 0.2 | 1.9 ± 0.5 |
| Adenosine Monophosphate | 7.7 ± 1.2 | 1.3 ± 0.8 | 5.8 ± 0.5 | 3.2 ± 0.9 |
| Adenosine-5'-diphosphate | 3.9 ± 0.1 | 5.7 ± 0.9 | 1.3 ± 1.1 | 1.9 ± 0.7 |
| Adenosine-5'-triphosphate | 8.1 ± 0.8 | 3.7 ± 0.4 | 2.0 ± 0.8 | 5.3 ± 1.0 |
| Adenylosuccinate | 2.4 ± 0.4 | 4.4 ± 0.6 | 1.3 ± 1.1 | 5.6 ± 0.7 |
| Alanine | 3.9 ± 0.4 | 2.3 ± 0.4 | 1.4 ± 1.1 | 5.6 ± 0.9 |
| alpha-ketoglutaric acid | 7.0 ± 0.8 | 1.9 ± 1.0 | 5.3 ± 0.2 | 3.1 ± 1.0 |
| Anserine | 10.5 ± 1.1 | 2.7 ± 0.2 | 5.2 ± 0.5 | 5.0 ± 0.9 |
| Arginine | 2.7 ± 0.7 | 3.4 ± 1.0 | 3.7 ± 1.1 | 2.6 ± 0.9 |
| Argininosuccinic Acid | 11.9 ± 0.5 | 3.3 ± 0.1 | 2.4 ± 0.7 | 4.1 ± 0.4 |
| Asparagine | 10.0 ± 0.3 | 5.5 ± 0.8 | 1.8 ± 0.7 | 1.1 ± 0.8 |
| Aspartic acid | 1.5 ± 0.7 | 1.4 ± 0.7 | 4.9 ± 1.0 | 5.1 ± 1.2 |
| Asymmetric dimethylarginine | 10.7 ± 0.6 | 4.2 ± 1.1 | 4.1 ± 0.4 | 2.5 ± 0.7 |
| Azelaic acid | 7.0 ± 1.0 | 1.2 ± 0.7 | 6.2 ± 0.3 | 5.6 ± 0.3 |
| beta-Alanine | 1.5 ± 0.7 | 4.2 ± 1.1 | 6.4 ± 1.2 | 4.6 ± 0.2 |
| Betaine | 10.1 ± 0.2 | 3.8 ± 0.8 | 5.8 ± 0.9 | 3.0 ± 0.6 |
| Biopterin | 8.8 ± 1.2 | 5.6 ± 0.6 | 3.2 ± 0.9 | 3.4 ± 0.5 |
| Biotin | 4.2 ± 1.1 | 2.6 ± 0.9 | 5.8 ± 0.9 | 4.1 ± 0.2 |
| Carnitine | 6.9 ± 0.5 | 1.5 ± 0.8 | 2.0 ± 1.2 | 4.4 ± 1.1 |
| Carnosine | 10.0 ± 1.1 | 2.1 ± 0.9 | 2.3 ± 0.3 | 2.5 ± 0.5 |
| CDP-choline | 3.8 ± 1.0 | 2.0 ± 0.3 | 6.5 ± 0.7 | 2.8 ± 1.0 |
| cGMP | 2.7 ± 0.9 | 2.3 ± 0.1 | 5.4 ± 0.9 | 4.8 ± 0.5 |
| Choline | 9.5 ± 0.8 | 3.7 ± 0.9 | 3.3 ± 0.2 | 1.2 ± 0.2 |
| Cinnamic acid | 9.1 ± 0.7 | 6.3 ± 0.5 | 5.2 ± 0.6 | 4.7 ± 0.7 |
| Citraconic acid | 3.3 ± 0.9 | 5.1 ± 0.4 | 4.3 ± 0.7 | 2.7 ± 1.1 |
| Citramalic acid | 4.3 ± 1.0 | 1.3 ± 0.9 | 3.1 ± 0.4 | 4.7 ± 0.7 |
| Citric Acid | 3.0 ± 0.4 | 1.5 ± 0.7 | 1.2 ± 0.1 | 2.2 ± 0.2 |
| Citrulline | 9.5 ± 0.8 | 1.1 ± 1.0 | 6.2 ± 1.0 | 5.2 ± 0.4 |
| Creatine | 10.9 ± 0.9 | 4.6 ± 0.7 | 2.1 ± 0.2 | 5.4 ± 0.1 |
| Creatinine | 3.4 ± 0.8 | 2.5 ± 0.1 | 2.7 ± 0.8 | 6.1 ± 0.4 |
| Cystathionine | 7.0 ± 0.2 | 5.4 ± 0.4 | 6.0 ± 1.0 | 3.7 ± 0.9 |
| Cysteamine | 9.5 ± 1.0 | 2.6 ± 0.2 | 2.3 ± 0.9 | 3.7 ± 0.9 |
| Cysteine | 5.4 ± 0.6 | 3.6 ± 0.9 | 5.8 ± 0.3 | 1.7 ± 1.0 |
| Cysteinylglycine | 1.5 ± 0.6 | 4.5 ± 0.5 | 5.7 ± 0.6 | 1.5 ± 0.4 |
| Cystine | 1.3 ± 0.2 | 3.9 ± 0.8 | 1.5 ± 0.5 | 6.2 ± 1.2 |
| Cytidine | 8.9 ± 0.5 | 1.4 ± 0.9 | 5.1 ± 1.0 | 2.0 ± 0.3 |
| Cytidine 3',5'-cyclic monophosphate | 3.6 ± 0.3 | 5.0 ± 0.7 | 6.1 ± 0.5 | 5.2 ± 1.0 |
| Cytidine 5'-diphosphate | 7.1 ± 0.6 | 1.4 ± 0.6 | 2.1 ± 0.4 | 5.7 ± 1.1 |
| Cytidine 5'-triphosphate | 9.5 ± 1.0 | 4.7 ± 1.2 | 1.3 ± 0.7 | 3.3 ± 0.3 |
| Cytidine monophosphate | 8.2 ± 1.0 | 4.3 ± 0.9 | 3.8 ± 0.5 | 3.0 ± 1.1 |
| Cytosine | 4.3 ± 0.2 | 5.0 ± 0.5 | 2.4 ± 1.2 | 1.1 ± 0.5 |
| dADP | 4.3 ± 0.5 | 4.9 ± 0.4 | 6.3 ± 1.0 | 4.2 ± 0.5 |
| Diethanolamine | 9.9 ± 0.7 | 2.2 ± 0.7 | 2.2 ± 0.3 | 2.8 ± 0.8 |
| Dihydrobiopterin | 2.2 ± 0.3 | 2.5 ± 0.2 | 2.2 ± 1.0 | 4.6 ± 0.9 |
| Dihydroneopterin triphosphate | 5.1 ± 0.2 | 5.3 ± 0.9 | 3.1 ± 0.4 | 5.4 ± 0.8 |
| Dihydroxyacetonephosphate | 2.3 ± 0.5 | 5.9 ± 0.4 | 1.1 ± 0.6 | 1.2 ± 0.9 |
| Dimethylglycine | 4.8 ± 0.2 | 2.7 ± 0.8 | 6.0 ± 0.6 | 4.3 ± 1.0 |
| Ectoine | 7.6 ± 0.8 | 2.4 ± 1.1 | 1.0 ± 0.3 | 3.2 ± 0.7 |
| Erythrose-4-phosphate | 9.6 ± 0.9 | 5.5 ± 0.2 | 4.4 ± 0.7 | 4.8 ± 0.8 |
| FAD | 6.1 ± 0.8 | 4.8 ± 0.4 | 3.7 ± 0.9 | 3.9 ± 0.2 |
| Folic acid | 5.2 ± 0.7 | 2.1 ± 0.4 | 4.0 ± 0.9 | 5.4 ± 0.2 |
| Fructose-6-phosphate | 7.8 ± 0.8 | 2.0 ± 0.1 | 6.5 ± 0.5 | 5.7 ± 1.0 |
| Fumaric Acid | 10.9 ± 1.1 | 3.4 ± 0.6 | 5.0 ± 0.7 | 3.9 ± 1.2 |
| Galactosamine | 6.6 ± 0.9 | 2.3 ± 0.8 | 3.4 ± 1.0 | 2.6 ± 0.9 |
| Galactose-1-phosphate | 5.4 ± 0.2 | 3.2 ± 0.6 | 3.5 ± 0.4 | 6.2 ± 0.5 |
| Galactose-6-Phosphate | 6.3 ± 0.8 | 5.0 ± 0.4 | 4.8 ± 0.9 | 6.4 ± 0.8 |
| gamma-Glutamylcysteine | 3.3 ± 0.2 | 6.3 ± 0.5 | 1.4 ± 0.6 | 3.0 ± 0.6 |
| Glucosamine | 7.0 ± 1.2 | 4.5 ± 1.1 | 6.0 ± 0.7 | 3.0 ± 0.4 |
| Glucosamine-6-Phosphate | 3.1 ± 0.3 | 5.2 ± 0.4 | 5.2 ± 0.1 | 1.7 ± 0.1 |
| Glucose | 11.7 ± 0.4 | 6.1 ± 0.7 | 4.1 ± 0.3 | 5.7 ± 0.9 |
| Glucose-1-phosphate | 9.6 ± 0.2 | 2.5 ± 0.1 | 5.4 ± 1.0 | 5.1 ± 0.4 |
| Glucose-6-Phosphate | 7.7 ± 0.2 | 4.1 ± 0.2 | 3.4 ± 0.5 | 2.9 ± 0.6 |
| Glucuronic acid | 3.8 ± 0.7 | 6.4 ± 1.1 | 6.0 ± 0.8 | 3.6 ± 0.7 |
| Glutamic acid | 8.8 ± 0.7 | 4.1 ± 0.8 | 1.2 ± 0.9 | 5.1 ± 0.4 |
| Glutamine | 6.7 ± 0.6 | 1.2 ± 0.2 | 6.0 ± 0.5 | 6.3 ± 0.2 |
| Glutathione | 5.8 ± 0.9 | 1.4 ± 0.5 | 6.5 ± 0.5 | 3.2 ± 0.4 |
| Glutathionylspermine | 2.8 ± 0.6 | 1.6 ± 0.6 | 3.3 ± 0.4 | 1.8 ± 0.4 |
| Glycerophosphocholine | 7.4 ± 0.2 | 3.3 ± 0.2 | 3.3 ± 0.6 | 2.1 ± 1.2 |
| Glycine | 6.9 ± 0.5 | 4.1 ± 1.1 | 1.9 ± 0.3 | 2.0 ± 1.0 |
| Glycolic acid | 6.0 ± 0.7 | 2.3 ± 1.0 | 1.2 ± 0.3 | 1.1 ± 1.0 |
| Guanidinoacetic acid | 6.3 ± 0.1 | 3.0 ± 0.2 | 1.5 ± 1.0 | 2.5 ± 0.2 |
| Guanidinobutanoic acid | 1.7 ± 0.9 | 2.5 ± 1.0 | 5.4 ± 0.2 | 1.9 ± 0.5 |
| Guanine | 1.5 ± 0.5 | 4.8 ± 0.1 | 3.4 ± 0.8 | 2.8 ± 1.1 |
| Guanosine | 4.8 ± 1.0 | 2.6 ± 0.7 | 1.8 ± 1.0 | 3.8 ± 1.1 |
| Guanosine 5'-diphosphate | 7.7 ± 0.4 | 6.2 ± 0.2 | 6.0 ± 0.5 | 3.7 ± 0.9 |
| Guanosine-5'-monophosphate | 11.2 ± 1.0 | 5.5 ± 0.3 | 5.4 ± 1.0 | 1.1 ± 0.2 |
| Hercynine | 11.5 ± 0.3 | 3.2 ± 1.1 | 5.6 ± 0.3 | 1.1 ± 0.2 |
| Hippuric acid | 5.7 ± 0.9 | 2.8 ± 1.1 | 6.5 ± 0.3 | 1.1 ± 0.6 |
| Histamine | 2.8 ± 0.6 | 1.7 ± 0.2 | 1.7 ± 0.6 | 4.9 ± 0.8 |
| Histidine | 1.7 ± 0.2 | 3.0 ± 0.5 | 1.7 ± 1.0 | 3.7 ± 0.3 |
| Homoalanine | 7.1 ± 0.4 | 1.7 ± 0.8 | 3.8 ± 0.3 | 2.5 ± 0.3 |
| Homoarginine | 2.9 ± 0.3 | 1.7 ± 1.1 | 2.6 ± 0.2 | 5.4 ± 0.9 |
| Homocysteine | 10.0 ± 0.8 | 4.1 ± 1.0 | 3.5 ± 0.7 | 1.1 ± 0.6 |
| Homogentisic acid | 7.0 ± 1.1 | 1.8 ± 0.7 | 6.2 ± 0.3 | 5.6 ± 1.1 |
| Homoleucine | 10.4 ± 0.1 | 3.6 ± 0.2 | 5.8 ± 0.2 | 6.4 ± 0.5 |
| Homovanillic acid | 1.6 ± 0.8 | 6.4 ± 1.0 | 6.0 ± 1.1 | 5.4 ± 0.8 |
| Hydantoin-5-propionate | 8.3 ± 1.0 | 5.6 ± 0.5 | 3.8 ± 0.9 | 5.8 ± 0.5 |
| Hypotaurine | 8.7 ± 0.8 | 6.2 ± 0.4 | 4.6 ± 0.2 | 3.7 ± 0.5 |
| Hypoxanthine | 7.2 ± 0.8 | 1.8 ± 1.1 | 5.3 ± 0.5 | 2.4 ± 1.1 |
| Indole-3-acetamide | 3.8 ± 0.4 | 2.8 ± 0.9 | 1.4 ± 0.8 | 2.4 ± 0.8 |
| Inosine | 8.1 ± 0.4 | 4.7 ± 0.6 | 2.1 ± 0.4 | 2.5 ± 0.1 |
| Inosine 5'-monophosphate | 4.7 ± 0.5 | 5.2 ± 0.8 | 3.9 ± 1.0 | 1.3 ± 0.7 |
| Isocitric Acid | 6.4 ± 0.3 | 2.9 ± 0.2 | 2.2 ± 0.9 | 1.8 ± 1.1 |
| Isoleucine | 11.5 ± 0.6 | 5.3 ± 1.1 | 6.4 ± 0.9 | 6.0 ± 0.7 |
| Lactic Acid | 6.5 ± 0.4 | 4.2 ± 0.9 | 1.4 ± 0.5 | 3.9 ± 0.2 |
| Leucine | 3.8 ± 1.2 | 5.4 ± 0.8 | 2.2 ± 1.2 | 2.6 ± 0.9 |
| Levulinic acid | 9.6 ± 0.8 | 3.0 ± 0.9 | 2.5 ± 0.4 | 4.7 ± 0.6 |
| Lysine | 5.5 ± 0.8 | 1.8 ± 0.4 | 6.4 ± 0.8 | 3.2 ± 1.1 |
| Malic Acid | 5.0 ± 0.8 | 1.0 ± 0.7 | 1.9 ± 1.0 | 2.2 ± 0.6 |
| Mannose-1 with Mannose-6-Phosphate | 3.1 ± 0.3 | 3.3 ± 1.2 | 4.7 ± 0.7 | 4.3 ± 0.6 |
| Mesaconic acid | 4.7 ± 1.0 | 6.2 ± 1.1 | 3.3 ± 1.0 | 2.1 ± 0.9 |
| Methionine | 5.7 ± 0.4 | 1.2 ± 1.2 | 5.6 ± 0.6 | 1.1 ± 0.4 |
| Methionine sulfoxide | 11.8 ± 0.6 | 2.2 ± 0.3 | 1.7 ± 0.2 | 1.7 ± 0.5 |
| Methylmalonic Acid | 1.2 ± 0.8 | 3.6 ± 0.9 | 1.1 ± 1.0 | 6.2 ± 0.7 |
| N1-Acetylspermidine | 3.0 ± 0.8 | 1.7 ± 1.1 | 4.1 ± 0.2 | 3.1 ± 0.7 |
| N1-Acetylspermine | 5.9 ± 0.1 | 2.9 ± 0.6 | 4.5 ± 1.1 | 6.0 ± 0.4 |
| N8-Acetylspermidine | 3.5 ± 1.0 | 5.9 ± 0.5 | 4.5 ± 0.4 | 2.1 ± 0.2 |
| N-Acetyl-Aspartyl-Glutamate | 3.8 ± 0.5 | 4.2 ± 1.0 | 6.1 ± 1.1 | 1.1 ± 0.4 |
| N-Acetyl-Aspartyl-Glutamyl-Glutamate | 10.5 ± 0.8 | 6.3 ± 0.2 | 4.2 ± 1.1 | 2.2 ± 0.6 |
| N-Acetyl-Galactosamine | 6.8 ± 0.6 | 3.6 ± 0.6 | 2.4 ± 0.3 | 1.1 ± 1.1 |
| N-Acetyl-Leucine | 9.9 ± 0.8 | 5.4 ± 0.9 | 1.9 ± 1.0 | 3.6 ± 0.8 |
| N-Acetyl-Methionine | 3.7 ± 1.0 | 5.7 ± 1.1 | 1.7 ± 0.2 | 3.6 ± 0.5 |
| N-Acetylneuraminic acid | 9.6 ± 0.5 | 3.5 ± 0.3 | 1.1 ± 0.2 | 1.2 ± 0.7 |
| N-Acetyl-Proline | 2.7 ± 0.6 | 2.0 ± 0.1 | 5.0 ± 0.8 | 2.9 ± 0.7 |
| N-Acetyl-Putrescine | 6.3 ± 0.1 | 4.5 ± 0.3 | 4.2 ± 0.5 | 5.2 ± 0.6 |
| N-Acetyl-Serine | 4.9 ± 0.5 | 3.3 ± 1.0 | 6.4 ± 1.1 | 3.5 ± 0.3 |
| N-Acetyl-Serotonin | 2.6 ± 1.1 | 3.7 ± 0.1 | 5.8 ± 1.0 | 4.0 ± 0.2 |
| N-Acyl-Homoserine-lactone | 10.8 ± 1.2 | 1.8 ± 0.8 | 2.8 ± 0.4 | 4.7 ± 0.8 |
| NAD | 6.8 ± 0.7 | 2.9 ± 0.9 | 2.7 ± 0.4 | 5.2 ± 0.2 |
| NADH | 6.4 ± 1.0 | 3.9 ± 0.1 | 2.4 ± 0.6 | 6.1 ± 0.8 |
| NADP | 6.8 ± 0.3 | 3.9 ± 0.9 | 3.6 ± 0.7 | 3.3 ± 0.8 |
| NADPH | 7.1 ± 0.5 | 6.2 ± 0.5 | 2.4 ± 0.8 | 5.4 ± 0.5 |
| N-alpha-Acetyl-lysine | 11.9 ± 0.7 | 1.7 ± 1.1 | 1.6 ± 1.0 | 4.6 ± 0.6 |
| N-alpha-Acetyl-Ornithine | 9.9 ± 1.2 | 1.6 ± 1.0 | 4.3 ± 0.6 | 2.4 ± 0.9 |
| N-Carbamoyl-Proline | 7.5 ± 0.2 | 6.3 ± 0.8 | 3.7 ± 0.5 | 5.4 ± 1.0 |
| N-Carbamoyl-Sarcosine | 3.9 ± 0.4 | 6.3 ± 1.1 | 1.6 ± 0.3 | 3.9 ± 0.6 |
| N-Epsilon-Acetyl-lysine | 7.0 ± 0.7 | 4.6 ± 0.6 | 1.7 ± 0.5 | 6.2 ± 0.2 |
| Nicotinamide | 1.9 ± 0.3 | 1.7 ± 1.1 | 2.6 ± 0.8 | 4.4 ± 0.4 |
| N-Methyl-Aspartic Acid | 10.4 ± 0.6 | 1.4 ± 0.5 | 1.9 ± 0.1 | 6.1 ± 0.3 |
| O-Phosphoethanolamine | 4.1 ± 0.6 | 3.6 ± 0.1 | 1.8 ± 0.4 | 2.0 ± 0.5 |
| Ophthalmic acid | 7.8 ± 0.1 | 3.0 ± 1.2 | 5.5 ± 0.6 | 3.4 ± 1.0 |
| Ornithine | 11.3 ± 1.1 | 6.2 ± 0.2 | 6.0 ± 0.2 | 6.0 ± 0.9 |
| Orotidine 5'-monophosphate | 11.1 ± 0.8 | 2.1 ± 0.9 | 6.0 ± 0.6 | 4.9 ± 0.8 |
| Oxaloacetate | 4.8 ± 0.3 | 1.4 ± 0.2 | 2.7 ± 1.0 | 1.5 ± 1.1 |
| Pantothenic acid | 6.4 ± 0.6 | 4.3 ± 0.4 | 4.3 ± 0.4 | 5.6 ± 0.8 |
| Phenylalanine | 1.4 ± 0.6 | 1.1 ± 0.7 | 5.3 ± 1.2 | 4.4 ± 0.5 |
| Phenylpyruvate | 10.4 ± 1.1 | 1.3 ± 0.6 | 3.6 ± 1.1 | 5.4 ± 0.4 |
| Phosphodimethylethanolamine | 3.0 ± 0.8 | 5.0 ± 1.1 | 4.0 ± 0.8 | 6.3 ± 1.2 |
| Phosphoenolpyruvate | 10.3 ± 0.1 | 3.7 ± 1.1 | 6.4 ± 0.8 | 2.0 ± 1.2 |
| Phosphorylcholine | 9.4 ± 0.8 | 3.8 ± 1.1 | 1.1 ± 1.0 | 3.9 ± 0.3 |
| Pipecolic acid | 4.9 ± 0.5 | 3.4 ± 0.1 | 5.8 ± 1.2 | 4.5 ± 0.5 |
| Proline | 8.3 ± 0.8 | 5.9 ± 0.2 | 2.5 ± 1.1 | 3.6 ± 0.2 |
| Putrescine | 1.1 ± 0.6 | 1.2 ± 0.4 | 5.3 ± 1.1 | 5.9 ± 0.4 |
| Pyridoxal | 7.9 ± 0.7 | 2.4 ± 0.5 | 4.6 ± 0.6 | 5.8 ± 0.4 |
| Pyridoxamine | 10.5 ± 0.9 | 2.7 ± 0.2 | 4.5 ± 0.7 | 1.8 ± 0.8 |
| Pyridoxine | 10.5 ± 0.3 | 4.0 ± 0.1 | 1.2 ± 1.1 | 5.6 ± 0.7 |
| Pyroglutamic acid | 8.2 ± 0.5 | 3.8 ± 0.7 | 2.8 ± 0.2 | 1.0 ± 0.9 |
| Pyrrolysine | 6.8 ± 0.3 | 6.4 ± 0.8 | 5.4 ± 0.7 | 6.5 ± 1.0 |
| Pyruvic Acid | 9.6 ± 0.2 | 6.3 ± 1.0 | 2.4 ± 1.0 | 2.1 ± 0.3 |
| Riboflavin | 4.7 ± 0.5 | 5.6 ± 0.5 | 4.3 ± 0.3 | 5.1 ± 0.2 |
| Ribose-5-Phosphate | 4.0 ± 1.1 | 1.9 ± 0.4 | 5.3 ± 0.6 | 6.0 ± 0.9 |
| Ribulose-1-Phosphate | 2.1 ± 0.9 | 3.9 ± 0.6 | 3.9 ± 0.4 | 1.3 ± 0.2 |
| Ribulose-5-Phosphate | 1.1 ± 1.0 | 4.9 ± 0.2 | 2.8 ± 0.3 | 2.1 ± 1.2 |
| Saccharopine | 3.3 ± 1.1 | 2.0 ± 1.1 | 5.9 ± 0.1 | 6.2 ± 0.7 |
| S-Adenosylhomocysteine | 8.5 ± 0.5 | 4.5 ± 0.9 | 2.0 ± 0.8 | 1.5 ± 1.2 |
| S-Adenosylmethioninamine | 5.4 ± 0.9 | 6.0 ± 0.9 | 3.9 ± 0.1 | 3.7 ± 0.7 |
| S-Adenosylmethionine | 11.3 ± 1.2 | 2.5 ± 0.9 | 5.7 ± 0.3 | 1.7 ± 0.7 |
| Sarcosine | 12.0 ± 0.2 | 1.2 ± 0.6 | 3.3 ± 1.0 | 5.1 ± 0.4 |
| Sedoheptulose-7-phosphate | 1.1 ± 0.8 | 5.3 ± 1.1 | 1.4 ± 0.8 | 2.2 ± 0.7 |
| Selenohomocysteine | 10.2 ± 0.6 | 4.7 ± 1.0 | 2.3 ± 0.3 | 4.7 ± 0.7 |
| Selenomethionine | 8.9 ± 0.4 | 3.6 ± 0.6 | 6.1 ± 0.7 | 5.1 ± 0.6 |
| Serine | 7.9 ± 0.9 | 2.8 ± 0.7 | 4.0 ± 1.1 | 5.6 ± 0.5 |
| S-Methyl-Thioinosine | 9.4 ± 0.7 | 6.4 ± 0.2 | 3.8 ± 0.7 | 1.3 ± 1.2 |
| Spermidine | 10.9 ± 0.6 | 4.2 ± 0.5 | 2.7 ± 1.1 | 2.3 ± 0.7 |
| Spermine | 7.5 ± 0.2 | 4.4 ± 0.7 | 6.2 ± 0.9 | 6.4 ± 0.4 |
| Succinate | 7.8 ± 0.7 | 1.7 ± 1.0 | 1.6 ± 1.1 | 6.1 ± 0.7 |
| Sucrose | 4.4 ± 0.6 | 2.2 ± 0.3 | 4.4 ± 0.2 | 4.6 ± 0.5 |
| Symmetric dimethylarginine | 8.2 ± 0.8 | 6.0 ± 1.0 | 4.3 ± 1.1 | 5.6 ± 0.5 |
| Taurine | 1.7 ± 0.6 | 3.4 ± 0.9 | 3.8 ± 0.2 | 6.1 ± 1.0 |
| Tetrahydroneopterin | 2.4 ± 0.4 | 1.7 ± 0.3 | 5.7 ± 0.7 | 2.6 ± 0.1 |
| Thiamine | 7.2 ± 0.3 | 2.0 ± 1.0 | 5.8 ± 0.5 | 4.9 ± 0.2 |
| Thiamine monophosphate | 5.3 ± 1.0 | 1.7 ± 1.1 | 1.6 ± 0.7 | 6.5 ± 0.8 |
| Threonic acid | 2.9 ± 0.5 | 1.7 ± 0.2 | 3.8 ± 0.3 | 2.6 ± 1.2 |
| Threonine | 9.9 ± 0.9 | 6.0 ± 1.0 | 5.4 ± 0.5 | 4.3 ± 0.8 |
| Thymidine | 7.9 ± 0.4 | 2.8 ± 0.1 | 4.8 ± 0.2 | 6.1 ± 0.7 |
| Thymidine Monophosphate | 10.4 ± 0.7 | 2.3 ± 0.3 | 3.6 ± 1.2 | 5.5 ± 0.4 |
| Thymine | 8.5 ± 0.4 | 3.3 ± 0.3 | 5.1 ± 0.3 | 6.4 ± 1.2 |
| Trigonelleine | 1.5 ± 0.3 | 3.7 ± 0.2 | 4.3 ± 0.3 | 1.7 ± 1.1 |
| Trimethylamine N-oxide | 10.5 ± 0.5 | 2.2 ± 0.4 | 3.7 ± 0.8 | 2.7 ± 0.8 |
| Trimethyllysine | 11.7 ± 0.3 | 5.1 ± 0.2 | 1.9 ± 1.2 | 2.4 ± 0.8 |
| Tryptophan | 11.9 ± 1.0 | 2.4 ± 0.7 | 1.4 ± 1.2 | 1.2 ± 1.0 |
| Tyramine | 2.8 ± 0.8 | 2.5 ± 0.2 | 1.9 ± 0.2 | 4.5 ± 0.9 |
| Tyrosine | 10.9 ± 1.0 | 2.3 ± 0.3 | 1.2 ± 0.3 | 1.8 ± 0.6 |
| Uracil | 6.0 ± 0.4 | 6.2 ± 0.5 | 5.7 ± 0.6 | 2.7 ± 1.1 |
| Uridine 5'-diphosphate | 10.1 ± 0.7 | 2.9 ± 0.7 | 2.0 ± 0.2 | 6.2 ± 0.5 |
| Uridine Monophosphate | 3.4 ± 0.6 | 1.7 ± 1.1 | 2.6 ± 0.2 | 1.7 ± 1.0 |
| Uridine-5-Diphosphogalactose | 4.3 ± 0.9 | 3.9 ± 0.1 | 1.4 ± 1.1 | 1.4 ± 1.1 |
| Urocanic acid | 9.5 ± 0.7 | 5.3 ± 0.8 | 6.0 ± 0.6 | 4.2 ± 0.3 |
| UTP | 9.4 ± 0.5 | 4.9 ± 0.8 | 4.1 ± 0.8 | 5.9 ± 0.5 |
| Valine | 1.0 ± 1.1 | 5.9 ± 0.9 | 5.3 ± 0.8 | 1.9 ± 0.6 |
| XMP | 10.6 ± 1.2 | 5.3 ± 0.8 | 4.7 ± 0.4 | 3.3 ± 0.3 |
| Xylulose-5-Phosphate | 11.0 ± 1.0 | 6.5 ± 0.5 | 4.7 ± 1.0 | 3.8 ± 1.0 |

### **Table S16** | Intra – day precision (as % RSD) (Urine extract as matrix, n = 5). The concentrations of QC_LLOQ_, QC_Low_, QC_Mid_ and QC_High_ were 10 ng/mL, 25 ng/mL, 100 ng/mL and 500 ng/mL respectively.

| Precision, % | QC_LLOQ_ | QC_low_ | QC_mid_ | QC_high_ |
| --- | --- | --- | --- | --- |
| 1-Methylguanine | 10.7 ± 0.5 | 5.0 ± 0.8 | 1.8 ± 0.7 | 3.2 ± 0.9 |
| 1-Methylhistidine | 3.0 ± 0.4 | 1.7 ± 0.1 | 2.8 ± 0.6 | 2.7 ± 1.0 |
| 1-Methylnicotinamide | 9.4 ± 1.1 | 4.6 ± 0.8 | 4.0 ± 0.6 | 1.8 ± 0.7 |
| 2-Aminoadipic Acid | 10.6 ± 0.3 | 4.8 ± 0.2 | 5.1 ± 1.0 | 3.1 ± 0.8 |
| 2-Deoxyadenosine | 7.7 ± 1.0 | 5.3 ± 1.2 | 5.0 ± 0.8 | 3.6 ± 0.6 |
| 2'-Deoxyadenosine 5'-monophosphate | 4.6 ± 0.3 | 1.8 ± 1.1 | 3.9 ± 0.2 | 5.4 ± 1.1 |
| 2'-deoxyadenosine 5'-triphosphate | 3.9 ± 0.5 | 5.1 ± 0.8 | 3.3 ± 1.2 | 3.4 ± 0.9 |
| 2'-Deoxycytidine-5'-monophosphate | 3.1 ± 0.2 | 1.6 ± 0.7 | 2.8 ± 0.6 | 3.2 ± 0.9 |
| 2-Deoxyguanosine | 11.7 ± 1.0 | 4.4 ± 0.9 | 5.5 ± 0.7 | 5.5 ± 0.2 |
| 2-hydroxybutyric acid | 9.4 ± 0.8 | 4.3 ± 0.8 | 2.3 ± 0.6 | 1.3 ± 0.8 |
| 2-hydroxyglutaric acid | 3.3 ± 0.8 | 5.3 ± 1.1 | 5.1 ± 0.2 | 5.5 ± 1.1 |
| 2-Hydroxyisocaproic acid | 3.5 ± 0.9 | 5.5 ± 0.6 | 3.3 ± 0.4 | 1.6 ± 0.5 |
| 2-ketobutyric acid | 10.0 ± 0.3 | 5.4 ± 0.5 | 4.0 ± 0.6 | 2.0 ± 0.3 |
| 2-oxoadipic acid | 5.4 ± 1.0 | 5.7 ± 1.1 | 2.7 ± 0.5 | 3.9 ± 0.4 |
| 2-Phenylglycine | 6.9 ± 0.8 | 3.3 ± 0.5 | 1.6 ± 0.8 | 2.9 ± 0.8 |
| 2-phosphoglycerate | 3.2 ± 0.8 | 3.3 ± 0.8 | 4.6 ± 0.6 | 4.2 ± 0.7 |
| 3-hydroxy-3-methyl-2-oxobutanoate | 3.2 ± 0.7 | 6.0 ± 0.2 | 2.3 ± 0.6 | 3.7 ± 0.7 |
| 3-hydroxybutyric acid | 9.5 ± 1.1 | 2.5 ± 1.0 | 1.8 ± 1.2 | 3.3 ± 0.3 |
| 3-Methylhistidine | 8.6 ± 0.1 | 3.5 ± 0.2 | 2.6 ± 0.5 | 5.6 ± 1.0 |
| 3-Phosphoglyceraldehyde | 6.7 ± 0.4 | 5.4 ± 0.4 | 3.0 ± 1.1 | 3.4 ± 0.6 |
| 3-phosphoglycerate | 10.3 ± 0.8 | 3.6 ± 1.1 | 1.6 ± 0.1 | 2.4 ± 0.5 |
| 4-amino-butyric acid | 11.1 ± 0.4 | 1.4 ± 0.5 | 2.1 ± 0.6 | 5.6 ± 0.7 |
| 4-aspartyl-phosphate | 7.8 ± 0.9 | 3.3 ± 0.6 | 3.9 ± 0.6 | 2.4 ± 1.1 |
| 4-Hydroxybenzoic acid | 5.7 ± 0.8 | 5.3 ± 0.7 | 4.9 ± 0.2 | 4.4 ± 0.8 |
| 4-Hydroxyphenylacetylglycine | 4.9 ± 1.0 | 2.0 ± 0.9 | 3.0 ± 0.9 | 3.3 ± 0.4 |
| 4-methyl-2-oxopentanoate | 2.2 ± 0.7 | 1.7 ± 1.0 | 4.6 ± 0.4 | 1.9 ± 0.9 |
| 4-Oxoglutaramate | 4.2 ± 0.4 | 6.0 ± 1.0 | 1.8 ± 0.6 | 3.7 ± 0.6 |
| 5-Hydroxyindolepyruvate | 6.6 ± 0.6 | 2.2 ± 0.7 | 2.5 ± 0.2 | 6.0 ± 0.8 |
| 5-Hydroxylysine | 11.2 ± 0.8 | 3.5 ± 1.2 | 5.0 ± 0.4 | 4.9 ± 0.9 |
| 5-Hydroxy-Tryptophan | 2.7 ± 0.6 | 4.9 ± 1.0 | 2.8 ± 0.4 | 5.5 ± 0.3 |
| 5-Phosphoribosylamine | 9.2 ± 0.9 | 4.7 ± 0.2 | 1.3 ± 0.2 | 5.9 ± 0.2 |
| 6-phosphogluconate | 11.9 ± 0.9 | 4.1 ± 0.2 | 2.6 ± 0.1 | 4.9 ± 1.1 |
| Acetylcarnitine | 11.9 ± 1.1 | 2.4 ± 0.3 | 3.4 ± 1.1 | 4.0 ± 0.6 |
| Aconitic Acid | 2.3 ± 0.4 | 5.5 ± 1.0 | 3.0 ± 0.2 | 5.6 ± 0.8 |
| Adenine | 8.2 ± 1.2 | 2.2 ± 0.3 | 3.8 ± 0.3 | 5.6 ± 0.1 |
| Adenosine | 5.2 ± 0.3 | 3.0 ± 0.1 | 5.9 ± 0.6 | 4.6 ± 0.4 |
| Adenosine 3',5'-cyclic monophosphate | 11.9 ± 1.1 | 5.4 ± 0.3 | 2.0 ± 0.4 | 3.6 ± 0.4 |
| Adenosine Monophosphate | 6.2 ± 0.7 | 3.8 ± 0.5 | 3.6 ± 1.0 | 4.9 ± 0.1 |
| Adenosine-5'-diphosphate | 4.7 ± 0.9 | 5.0 ± 0.3 | 5.9 ± 0.9 | 4.9 ± 1.1 |
| Adenosine-5'-triphosphate | 2.0 ± 0.4 | 3.2 ± 0.6 | 1.4 ± 1.1 | 1.3 ± 0.7 |
| Adenylosuccinate | 9.6 ± 0.8 | 4.2 ± 0.9 | 3.7 ± 0.5 | 4.4 ± 0.4 |
| Alanine | 11.9 ± 0.2 | 1.9 ± 0.7 | 4.5 ± 0.6 | 2.9 ± 0.7 |
| alpha-ketoglutaric acid | 5.3 ± 1.1 | 2.6 ± 0.9 | 3.0 ± 0.8 | 4.0 ± 0.6 |
| Anserine | 11.4 ± 0.6 | 4.6 ± 1.1 | 3.5 ± 0.7 | 2.3 ± 0.8 |
| Arginine | 10.7 ± 1.0 | 5.3 ± 0.3 | 3.4 ± 1.0 | 3.9 ± 0.6 |
| Argininosuccinic Acid | 8.9 ± 0.6 | 3.9 ± 0.2 | 2.8 ± 0.4 | 1.5 ± 0.6 |
| Asparagine | 11.9 ± 1.1 | 2.3 ± 0.8 | 3.9 ± 0.6 | 3.9 ± 0.6 |
| Aspartic acid | 8.2 ± 0.9 | 2.7 ± 0.4 | 4.8 ± 0.3 | 3.1 ± 1.1 |
| Asymmetric dimethylarginine | 6.8 ± 1.1 | 3.2 ± 0.7 | 4.4 ± 1.1 | 5.9 ± 0.3 |
| Azelaic acid | 3.5 ± 0.3 | 4.1 ± 0.7 | 5.1 ± 0.7 | 2.4 ± 1.2 |
| beta-Alanine | 8.5 ± 1.1 | 4.2 ± 1.1 | 1.3 ± 0.2 | 3.4 ± 0.7 |
| Betaine | 11.6 ± 0.7 | 3.5 ± 1.0 | 3.4 ± 0.3 | 5.8 ± 1.0 |
| Biopterin | 10.2 ± 0.6 | 1.6 ± 1.0 | 4.4 ± 0.6 | 4.4 ± 0.5 |
| Biotin | 9.4 ± 0.9 | 5.1 ± 0.3 | 3.3 ± 0.1 | 3.5 ± 0.1 |
| Carnitine | 3.1 ± 0.3 | 2.7 ± 0.5 | 2.9 ± 1.2 | 2.6 ± 0.3 |
| Carnosine | 8.9 ± 0.5 | 5.9 ± 0.3 | 5.6 ± 0.6 | 4.9 ± 1.1 |
| CDP-choline | 4.5 ± 0.3 | 5.4 ± 0.8 | 3.4 ± 0.8 | 2.4 ± 0.7 |
| cGMP | 8.6 ± 0.1 | 2.4 ± 0.5 | 1.5 ± 0.5 | 3.1 ± 0.1 |
| Choline | 3.0 ± 1.1 | 5.0 ± 0.4 | 1.9 ± 0.4 | 3.7 ± 0.8 |
| Cinnamic acid | 9.3 ± 0.6 | 1.9 ± 0.6 | 4.6 ± 0.8 | 1.5 ± 0.3 |
| Citraconic acid | 4.0 ± 0.8 | 4.9 ± 0.5 | 6.0 ± 1.1 | 3.4 ± 0.3 |
| Citramalic acid | 6.6 ± 0.5 | 1.9 ± 1.2 | 4.2 ± 0.3 | 3.7 ± 1.1 |
| Citric Acid | 11.0 ± 0.3 | 3.4 ± 0.1 | 1.7 ± 0.7 | 3.1 ± 1.1 |
| Citrulline | 7.8 ± 0.7 | 3.9 ± 1.1 | 5.4 ± 0.5 | 2.1 ± 0.2 |
| Creatine | 8.0 ± 0.1 | 1.9 ± 0.8 | 4.9 ± 0.4 | 2.0 ± 1.1 |
| Creatinine | 8.1 ± 0.8 | 1.5 ± 0.2 | 1.9 ± 0.9 | 2.9 ± 1.0 |
| Cystathionine | 7.8 ± 0.3 | 1.6 ± 0.2 | 4.6 ± 0.1 | 4.5 ± 0.2 |
| Cysteamine | 9.0 ± 0.1 | 3.6 ± 0.4 | 3.5 ± 1.1 | 2.1 ± 0.9 |
| Cysteine | 2.7 ± 0.8 | 5.6 ± 0.9 | 4.3 ± 0.9 | 4.3 ± 1.2 |
| Cysteinylglycine | 6.7 ± 0.3 | 4.7 ± 1.0 | 1.6 ± 0.9 | 2.4 ± 1.2 |
| Cystine | 4.5 ± 0.7 | 1.5 ± 0.2 | 1.8 ± 0.6 | 2.2 ± 0.9 |
| Cytidine | 3.1 ± 0.3 | 4.1 ± 0.1 | 5.8 ± 1.0 | 4.2 ± 1.1 |
| Cytidine 3',5'-cyclic monophosphate | 11.4 ± 1.2 | 1.8 ± 0.7 | 4.1 ± 0.2 | 5.1 ± 0.2 |
| Cytidine 5'-diphosphate | 8.4 ± 0.3 | 1.4 ± 0.1 | 3.6 ± 1.0 | 5.6 ± 1.0 |
| Cytidine 5'-triphosphate | 4.3 ± 0.8 | 2.1 ± 0.8 | 1.7 ± 0.4 | 5.4 ± 0.5 |
| Cytidine monophosphate | 10.8 ± 1.1 | 2.7 ± 0.9 | 1.8 ± 1.1 | 5.5 ± 0.4 |
| Cytosine | 2.3 ± 0.8 | 5.4 ± 0.4 | 4.3 ± 0.3 | 4.2 ± 1.0 |
| dADP | 5.3 ± 1.2 | 5.7 ± 0.9 | 5.6 ± 0.7 | 5.2 ± 0.4 |
| Diethanolamine | 9.4 ± 1.1 | 5.9 ± 0.2 | 3.4 ± 0.7 | 4.2 ± 0.3 |
| Dihydrobiopterin | 8.7 ± 1.0 | 1.6 ± 0.8 | 4.3 ± 0.6 | 2.2 ± 1.0 |
| Dihydroneopterin triphosphate | 9.9 ± 1.1 | 4.1 ± 0.9 | 1.8 ± 0.7 | 4.7 ± 0.5 |
| Dihydroxyacetonephosphate | 6.1 ± 0.4 | 5.5 ± 0.5 | 5.6 ± 1.2 | 4.4 ± 0.5 |
| Dimethylglycine | 6.0 ± 0.9 | 2.9 ± 1.0 | 4.6 ± 0.2 | 3.5 ± 1.1 |
| Ectoine | 7.1 ± 0.2 | 5.4 ± 0.7 | 2.0 ± 1.0 | 4.2 ± 1.2 |
| Erythrose-4-phosphate | 8.6 ± 0.8 | 5.9 ± 0.7 | 5.9 ± 0.8 | 1.5 ± 1.1 |
| FAD | 2.6 ± 0.2 | 2.5 ± 1.2 | 5.1 ± 0.5 | 4.4 ± 0.1 |
| Folic acid | 8.7 ± 1.0 | 2.8 ± 0.9 | 5.7 ± 0.1 | 5.5 ± 0.5 |
| Fructose-6-phosphate | 10.9 ± 0.4 | 1.7 ± 0.2 | 5.0 ± 0.9 | 4.3 ± 0.7 |
| Fumaric Acid | 8.8 ± 0.3 | 2.2 ± 1.0 | 3.7 ± 1.0 | 2.0 ± 1.1 |
| Galactosamine | 6.0 ± 0.9 | 1.8 ± 1.1 | 4.7 ± 0.4 | 4.6 ± 0.2 |
| Galactose-1-phosphate | 11.1 ± 1.2 | 2.7 ± 1.0 | 1.4 ± 0.8 | 3.9 ± 1.0 |
| Galactose-6-Phosphate | 5.1 ± 0.9 | 3.3 ± 0.6 | 3.3 ± 0.9 | 2.1 ± 0.7 |
| gamma-Glutamylcysteine | 3.1 ± 0.5 | 3.8 ± 0.4 | 4.0 ± 1.0 | 2.1 ± 0.6 |
| Glucosamine | 4.9 ± 0.7 | 1.8 ± 0.9 | 5.0 ± 0.6 | 2.1 ± 1.2 |
| Glucosamine-6-Phosphate | 8.6 ± 0.8 | 5.3 ± 0.7 | 4.0 ± 1.0 | 5.4 ± 1.0 |
| Glucose | 6.5 ± 0.6 | 2.7 ± 0.6 | 5.0 ± 0.5 | 1.8 ± 0.2 |
| Glucose-1-phosphate | 8.6 ± 0.5 | 1.5 ± 0.9 | 5.8 ± 0.8 | 3.5 ± 0.5 |
| Glucose-6-Phosphate | 5.6 ± 0.7 | 3.1 ± 1.2 | 4.7 ± 0.5 | 1.8 ± 0.5 |
| Glucuronic acid | 11.4 ± 0.3 | 4.3 ± 0.1 | 1.6 ± 0.8 | 4.0 ± 0.6 |
| Glutamic acid | 10.5 ± 0.2 | 2.8 ± 0.7 | 4.0 ± 1.1 | 3.6 ± 1.1 |
| Glutamine | 5.8 ± 1.1 | 2.3 ± 1.2 | 4.5 ± 0.7 | 1.4 ± 0.4 |
| Glutathione | 6.8 ± 0.1 | 4.4 ± 1.0 | 4.6 ± 0.2 | 5.6 ± 0.2 |
| Glutathionylspermine | 8.2 ± 0.5 | 3.0 ± 1.0 | 5.5 ± 0.2 | 2.2 ± 1.1 |
| Glycerophosphocholine | 10.8 ± 0.4 | 3.3 ± 0.8 | 3.5 ± 0.7 | 5.6 ± 0.6 |
| Glycine | 9.4 ± 0.2 | 2.2 ± 0.3 | 3.3 ± 0.9 | 1.8 ± 0.4 |
| Glycolic acid | 7.0 ± 0.3 | 1.8 ± 0.3 | 4.6 ± 0.8 | 3.7 ± 0.1 |
| Guanidinoacetic acid | 3.8 ± 0.4 | 4.4 ± 0.8 | 5.5 ± 0.9 | 4.5 ± 0.6 |
| Guanidinobutanoic acid | 5.8 ± 0.1 | 4.5 ± 0.9 | 2.0 ± 0.8 | 4.9 ± 0.7 |
| Guanine | 11.2 ± 1.1 | 3.6 ± 0.5 | 2.3 ± 0.9 | 4.4 ± 0.5 |
| Guanosine | 10.5 ± 1.2 | 3.3 ± 0.8 | 5.5 ± 1.0 | 2.0 ± 0.4 |
| Guanosine 5'-diphosphate | 7.7 ± 0.6 | 2.1 ± 1.0 | 3.5 ± 0.4 | 2.3 ± 0.8 |
| Guanosine-5'-monophosphate | 9.4 ± 0.9 | 5.6 ± 1.1 | 2.7 ± 0.4 | 5.9 ± 0.6 |
| Hercynine | 11.6 ± 1.0 | 5.8 ± 0.8 | 2.6 ± 0.9 | 2.2 ± 1.0 |
| Hippuric acid | 8.1 ± 0.8 | 4.8 ± 0.7 | 5.4 ± 0.3 | 5.6 ± 1.1 |
| Histamine | 7.5 ± 0.5 | 2.0 ± 0.5 | 4.2 ± 0.9 | 3.9 ± 0.9 |
| Histidine | 2.8 ± 0.6 | 5.1 ± 0.4 | 2.6 ± 0.3 | 3.5 ± 0.6 |
| Homoalanine | 3.9 ± 1.0 | 5.0 ± 0.3 | 5.1 ± 0.9 | 4.9 ± 0.5 |
| Homoarginine | 6.3 ± 0.3 | 4.2 ± 0.9 | 5.2 ± 1.0 | 4.2 ± 0.8 |
| Homocysteine | 8.7 ± 1.1 | 3.5 ± 0.3 | 5.6 ± 0.3 | 5.0 ± 0.9 |
| Homogentisic acid | 6.5 ± 0.7 | 5.1 ± 0.4 | 3.6 ± 0.2 | 2.8 ± 1.2 |
| Homoleucine | 9.5 ± 0.7 | 2.0 ± 0.2 | 4.0 ± 0.2 | 4.5 ± 0.8 |
| Homovanillic acid | 11.0 ± 0.2 | 5.0 ± 0.5 | 2.6 ± 0.8 | 4.0 ± 0.6 |
| Hydantoin-5-propionate | 8.7 ± 0.6 | 5.4 ± 1.1 | 4.1 ± 1.0 | 1.3 ± 0.2 |
| Hypotaurine | 3.6 ± 0.8 | 5.3 ± 0.8 | 3.2 ± 1.1 | 3.9 ± 0.6 |
| Hypoxanthine | 5.8 ± 0.3 | 2.9 ± 0.2 | 5.7 ± 0.9 | 5.9 ± 0.5 |
| Indole-3-acetamide | 11.5 ± 0.8 | 5.7 ± 0.7 | 5.1 ± 1.1 | 2.5 ± 0.2 |
| Inosine | 11.1 ± 0.4 | 3.8 ± 0.4 | 4.5 ± 0.8 | 4.3 ± 0.7 |
| Inosine 5'-monophosphate | 5.8 ± 1.0 | 1.4 ± 1.0 | 5.5 ± 0.1 | 2.2 ± 0.6 |
| Isocitric Acid | 2.5 ± 0.2 | 3.8 ± 0.5 | 1.4 ± 0.6 | 1.7 ± 1.0 |
| Isoleucine | 5.6 ± 0.6 | 2.1 ± 0.3 | 3.3 ± 0.3 | 5.2 ± 1.1 |
| Lactic Acid | 8.7 ± 0.6 | 2.8 ± 0.9 | 5.0 ± 0.8 | 3.2 ± 1.2 |
| Leucine | 3.3 ± 0.9 | 3.7 ± 1.0 | 5.9 ± 0.9 | 5.9 ± 0.9 |
| Levulinic acid | 11.5 ± 0.4 | 3.1 ± 0.9 | 4.5 ± 0.7 | 2.8 ± 0.2 |
| Lysine | 5.1 ± 0.9 | 1.5 ± 0.7 | 1.7 ± 0.2 | 5.7 ± 0.6 |
| Malic Acid | 3.1 ± 0.6 | 2.7 ± 0.3 | 5.8 ± 1.2 | 2.0 ± 0.8 |
| Mannose-1 with Mannose-6-Phosphate | 2.9 ± 0.2 | 4.9 ± 0.9 | 4.5 ± 0.3 | 4.8 ± 0.1 |
| Mesaconic acid | 3.4 ± 0.1 | 2.5 ± 1.1 | 4.1 ± 0.6 | 3.8 ± 1.0 |
| Methionine | 9.9 ± 1.1 | 3.3 ± 0.1 | 4.4 ± 0.7 | 4.2 ± 1.1 |
| Methionine sulfoxide | 11.8 ± 0.5 | 3.0 ± 1.0 | 5.1 ± 0.1 | 3.5 ± 0.3 |
| Methylmalonic Acid | 4.3 ± 1.0 | 5.8 ± 1.1 | 4.8 ± 0.4 | 1.6 ± 0.3 |
| N1-Acetylspermidine | 5.6 ± 0.6 | 1.5 ± 0.6 | 5.7 ± 0.6 | 2.4 ± 0.9 |
| N1-Acetylspermine | 9.1 ± 1.2 | 2.8 ± 0.6 | 3.1 ± 1.1 | 2.2 ± 1.0 |
| N8-Acetylspermidine | 5.2 ± 0.8 | 4.2 ± 0.7 | 3.7 ± 0.6 | 2.9 ± 0.7 |
| N-Acetyl-Aspartyl-Glutamate | 11.2 ± 1.2 | 3.0 ± 0.6 | 3.0 ± 0.7 | 2.4 ± 0.2 |
| N-Acetyl-Aspartyl-Glutamyl-Glutamate | 6.4 ± 1.2 | 1.5 ± 0.9 | 1.5 ± 0.3 | 4.5 ± 0.4 |
| N-Acetyl-Galactosamine | 10.5 ± 0.3 | 4.5 ± 0.2 | 3.2 ± 1.0 | 4.6 ± 1.1 |
| N-Acetyl-Leucine | 10.4 ± 0.9 | 5.0 ± 1.0 | 2.7 ± 0.8 | 5.1 ± 1.1 |
| N-Acetyl-Methionine | 5.1 ± 0.5 | 5.7 ± 0.4 | 2.3 ± 0.2 | 3.0 ± 0.5 |
| N-Acetylneuraminic acid | 10.1 ± 0.7 | 5.4 ± 0.2 | 3.8 ± 0.9 | 4.5 ± 0.4 |
| N-Acetyl-Proline | 9.7 ± 0.8 | 4.8 ± 0.3 | 2.8 ± 0.9 | 5.3 ± 0.3 |
| N-Acetyl-Putrescine | 6.2 ± 0.8 | 1.5 ± 1.2 | 4.5 ± 0.4 | 2.0 ± 0.7 |
| N-Acetyl-Serine | 11.5 ± 1.1 | 5.1 ± 0.1 | 4.4 ± 0.5 | 2.3 ± 0.4 |
| N-Acetyl-Serotonin | 7.4 ± 0.4 | 5.9 ± 0.7 | 1.5 ± 0.8 | 3.7 ± 0.3 |
| N-Acyl-Homoserine-lactone | 11.7 ± 0.8 | 2.7 ± 0.7 | 4.6 ± 1.2 | 1.7 ± 0.4 |
| NAD | 7.6 ± 0.2 | 2.7 ± 1.1 | 4.7 ± 0.3 | 5.1 ± 0.7 |
| NADH | 7.3 ± 0.8 | 2.3 ± 0.6 | 2.4 ± 0.8 | 3.2 ± 0.4 |
| NADP | 10.1 ± 0.8 | 3.9 ± 0.6 | 3.5 ± 1.0 | 4.2 ± 0.5 |
| NADPH | 3.8 ± 0.3 | 5.0 ± 0.1 | 4.2 ± 1.1 | 4.9 ± 0.4 |
| N-alpha-Acetyl-lysine | 2.4 ± 0.2 | 3.9 ± 0.9 | 5.4 ± 0.2 | 6.0 ± 0.3 |
| N-alpha-Acetyl-Ornithine | 6.7 ± 0.7 | 4.7 ± 0.3 | 4.1 ± 0.8 | 1.3 ± 0.4 |
| N-Carbamoyl-Proline | 4.6 ± 0.5 | 1.6 ± 1.1 | 6.0 ± 1.2 | 3.3 ± 0.4 |
| N-Carbamoyl-Sarcosine | 2.5 ± 0.2 | 4.2 ± 0.6 | 1.7 ± 0.4 | 2.6 ± 0.5 |
| N-Epsilon-Acetyl-lysine | 7.4 ± 0.4 | 3.8 ± 1.1 | 5.5 ± 0.2 | 5.1 ± 0.7 |
| Nicotinamide | 10.3 ± 0.7 | 1.3 ± 1.1 | 2.9 ± 0.9 | 4.8 ± 1.2 |
| N-Methyl-Aspartic Acid | 8.7 ± 0.7 | 3.6 ± 0.8 | 4.5 ± 0.6 | 4.6 ± 0.2 |
| O-Phosphoethanolamine | 4.6 ± 0.9 | 1.6 ± 0.6 | 5.7 ± 0.1 | 2.6 ± 0.6 |
| Ophthalmic acid | 6.3 ± 1.0 | 4.8 ± 0.1 | 5.3 ± 0.8 | 2.2 ± 0.8 |
| Ornithine | 6.9 ± 0.4 | 2.2 ± 0.1 | 3.7 ± 0.7 | 3.3 ± 0.8 |
| Orotidine 5'-monophosphate | 3.4 ± 0.2 | 1.4 ± 0.6 | 1.3 ± 0.9 | 2.8 ± 1.0 |
| Oxaloacetate | 9.0 ± 1.1 | 3.6 ± 0.1 | 2.1 ± 0.5 | 5.6 ± 0.3 |
| Pantothenic acid | 6.2 ± 0.8 | 2.5 ± 0.7 | 1.9 ± 0.9 | 1.7 ± 0.9 |
| Phenylalanine | 2.2 ± 0.1 | 4.7 ± 0.6 | 4.3 ± 0.6 | 2.1 ± 0.4 |
| Phenylpyruvate | 2.2 ± 0.2 | 1.8 ± 0.8 | 3.9 ± 0.7 | 2.9 ± 1.1 |
| Phosphodimethylethanolamine | 7.9 ± 0.2 | 3.7 ± 1.0 | 1.6 ± 0.3 | 2.4 ± 1.2 |
| Phosphoenolpyruvate | 9.5 ± 0.9 | 4.0 ± 1.1 | 3.5 ± 0.7 | 3.8 ± 0.4 |
| Phosphorylcholine | 5.0 ± 0.9 | 2.2 ± 0.5 | 4.9 ± 0.7 | 3.8 ± 1.0 |
| Pipecolic acid | 2.8 ± 1.0 | 2.2 ± 1.2 | 2.4 ± 0.6 | 4.9 ± 0.7 |
| Proline | 7.1 ± 0.9 | 1.9 ± 0.3 | 1.9 ± 0.7 | 5.7 ± 0.4 |
| Putrescine | 8.3 ± 0.1 | 3.1 ± 0.6 | 1.6 ± 1.2 | 2.0 ± 0.9 |
| Pyridoxal | 9.1 ± 1.1 | 4.7 ± 0.5 | 3.1 ± 0.3 | 4.8 ± 1.2 |
| Pyridoxamine | 11.2 ± 0.3 | 2.7 ± 0.2 | 5.9 ± 0.1 | 1.4 ± 0.9 |
| Pyridoxine | 9.3 ± 0.6 | 5.4 ± 0.2 | 1.6 ± 0.9 | 4.6 ± 1.2 |
| Pyroglutamic acid | 10.0 ± 0.4 | 1.3 ± 0.8 | 4.4 ± 1.0 | 2.1 ± 0.7 |
| Pyrrolysine | 9.1 ± 1.0 | 2.9 ± 0.4 | 2.5 ± 0.7 | 2.9 ± 0.1 |
| Pyruvic Acid | 2.6 ± 0.9 | 2.1 ± 0.5 | 5.5 ± 1.1 | 1.5 ± 0.9 |
| Riboflavin | 10.7 ± 0.8 | 2.7 ± 0.8 | 1.9 ± 0.6 | 1.4 ± 0.5 |
| Ribose-5-Phosphate | 4.7 ± 0.2 | 3.6 ± 0.1 | 4.8 ± 0.9 | 6.0 ± 0.2 |
| Ribulose-1-Phosphate | 10.6 ± 0.4 | 3.5 ± 0.2 | 1.5 ± 0.7 | 2.5 ± 0.2 |
| Ribulose-5-Phosphate | 9.0 ± 0.8 | 4.8 ± 0.9 | 2.5 ± 1.1 | 3.5 ± 0.2 |
| Saccharopine | 11.9 ± 1.2 | 1.5 ± 0.6 | 3.3 ± 0.8 | 4.1 ± 0.4 |
| S-Adenosylhomocysteine | 10.1 ± 0.9 | 5.0 ± 0.1 | 4.3 ± 1.0 | 2.8 ± 0.9 |
| S-Adenosylmethioninamine | 8.2 ± 0.6 | 2.8 ± 1.1 | 4.5 ± 1.2 | 3.3 ± 0.6 |
| S-Adenosylmethionine | 11.0 ± 0.4 | 5.3 ± 0.2 | 2.6 ± 0.7 | 3.4 ± 0.2 |
| Sarcosine | 2.4 ± 0.2 | 1.8 ± 1.0 | 2.3 ± 1.1 | 4.6 ± 0.6 |
| Sedoheptulose-7-phosphate | 5.7 ± 1.0 | 1.6 ± 0.3 | 2.1 ± 0.5 | 1.8 ± 1.0 |
| Selenohomocysteine | 8.7 ± 0.2 | 2.2 ± 0.2 | 2.0 ± 0.1 | 2.7 ± 0.5 |
| Selenomethionine | 10.5 ± 0.8 | 3.9 ± 0.4 | 3.1 ± 0.7 | 2.7 ± 0.4 |
| Serine | 9.5 ± 0.7 | 4.0 ± 0.1 | 4.9 ± 0.8 | 3.0 ± 1.1 |
| S-Methyl-Thioinosine | 2.8 ± 0.4 | 4.9 ± 0.3 | 2.0 ± 0.3 | 3.2 ± 0.2 |
| Spermidine | 11.1 ± 1.1 | 2.2 ± 0.8 | 3.3 ± 1.2 | 5.5 ± 0.7 |
| Spermine | 4.6 ± 0.7 | 1.8 ± 0.7 | 3.3 ± 0.2 | 1.9 ± 0.8 |
| Succinate | 7.3 ± 0.9 | 4.3 ± 0.8 | 3.0 ± 0.2 | 2.2 ± 0.4 |
| Sucrose | 7.0 ± 0.5 | 5.6 ± 0.2 | 5.3 ± 0.4 | 3.9 ± 0.4 |
| Symmetric dimethylarginine | 9.9 ± 1.1 | 1.9 ± 0.8 | 4.5 ± 0.9 | 2.6 ± 0.8 |
| Taurine | 8.8 ± 0.1 | 5.2 ± 0.7 | 5.8 ± 0.6 | 5.7 ± 0.3 |
| Tetrahydroneopterin | 11.5 ± 0.4 | 2.3 ± 0.5 | 4.6 ± 0.3 | 2.1 ± 0.3 |
| Thiamine | 6.3 ± 0.2 | 3.3 ± 1.1 | 3.9 ± 0.4 | 3.3 ± 0.7 |
| Thiamine monophosphate | 6.9 ± 0.9 | 5.0 ± 0.6 | 2.0 ± 1.1 | 2.7 ± 0.5 |
| Threonic acid | 4.5 ± 0.9 | 4.5 ± 0.5 | 4.7 ± 0.7 | 4.9 ± 0.7 |
| Threonine | 2.6 ± 0.1 | 6.0 ± 0.8 | 3.0 ± 0.6 | 1.7 ± 0.3 |
| Thymidine | 9.4 ± 0.2 | 1.4 ± 0.9 | 5.8 ± 1.0 | 4.8 ± 1.0 |
| Thymidine Monophosphate | 5.3 ± 1.0 | 2.1 ± 0.8 | 1.4 ± 0.4 | 3.4 ± 0.3 |
| Thymine | 4.9 ± 1.0 | 2.3 ± 1.0 | 4.7 ± 0.8 | 4.6 ± 1.1 |
| Trigonelleine | 10.8 ± 0.2 | 2.6 ± 0.1 | 3.0 ± 0.9 | 5.3 ± 1.1 |
| Trimethylamine N-oxide | 9.8 ± 0.5 | 3.3 ± 1.0 | 1.4 ± 0.2 | 1.5 ± 0.3 |
| Trimethyllysine | 7.3 ± 0.3 | 2.2 ± 0.7 | 2.1 ± 0.1 | 3.0 ± 0.8 |
| Tryptophan | 11.9 ± 0.8 | 2.7 ± 0.8 | 1.9 ± 1.0 | 2.3 ± 0.9 |
| Tyramine | 5.3 ± 0.1 | 2.2 ± 0.9 | 2.2 ± 0.9 | 5.1 ± 0.4 |
| Tyrosine | 5.7 ± 0.8 | 4.8 ± 1.1 | 4.5 ± 1.0 | 5.1 ± 0.2 |
| Uracil | 5.4 ± 0.3 | 3.2 ± 0.9 | 1.5 ± 0.8 | 3.2 ± 1.0 |
| Uridine 5'-diphosphate | 3.9 ± 0.9 | 1.4 ± 1.1 | 3.6 ± 0.8 | 3.3 ± 1.0 |
| Uridine Monophosphate | 8.4 ± 0.9 | 4.6 ± 0.5 | 4.4 ± 0.4 | 4.6 ± 0.7 |
| Uridine-5-Diphosphogalactose | 2.2 ± 1.1 | 3.3 ± 0.3 | 4.0 ± 0.3 | 5.3 ± 1.1 |
| Urocanic acid | 3.2 ± 0.1 | 3.3 ± 1.0 | 3.3 ± 0.3 | 2.0 ± 0.2 |
| UTP | 7.6 ± 0.4 | 1.6 ± 1.0 | 2.6 ± 1.1 | 4.9 ± 0.3 |
| Valine | 2.0 ± 1.1 | 4.8 ± 0.3 | 4.1 ± 0.2 | 4.7 ± 0.8 |
| XMP | 11.3 ± 0.7 | 2.5 ± 0.3 | 5.0 ± 1.0 | 4.6 ± 0.7 |
| Xylulose-5-Phosphate | 9.1 ± 1.0 | 1.4 ± 0.5 | 2.0 ± 0.7 | 3.1 ± 0.7 |

### **Table S17** | Inter – day precision (as % RSD) (Colorectal organoid extract as matrix, n = 5). The concentrations of QC_LLOQ_, QC_Low_, QC_Mid_ and QC_High_ were 10 ng/mL, 25 ng/mL, 100 ng/mL and 500 ng/mL respectively.

| Metabolite | QC_LLOQ_ | QC_Low_ | QC_Mid_ | QC_High_ |
| --- | --- | --- | --- | --- |
| 1-Methylguanine | 8.3 ± 0.6 | 2.6 ± 0.5 | 5.4 ± 0.6 | 3.5 ± 0.3 |
| 1-Methylhistidine | 11.7 ± 0.8 | 5.8 ± 0.8 | 1.7 ± 0.2 | 3.3 ± 0.4 |
| 1-Methylnicotinamide | 4.8 ± 0.8 | 5.4 ± 1.0 | 6.3 ± 0.8 | 4.2 ± 0.8 |
| 2-Aminoadipic Acid | 11.5 ± 0.6 | 2.5 ± 0.5 | 5.0 ± 0.8 | 2.8 ± 0.4 |
| 2-Deoxyadenosine | 8.7 ± 0.9 | 7.7 ± 0.4 | 6.0 ± 0.7 | 5.1 ± 0.5 |
| 2'-Deoxyadenosine 5'-monophosphate | 12.3 ± 0.7 | 7.1 ± 0.8 | 1.8 ± 0.2 | 5.9 ± 0.2 |
| 2'-deoxyadenosine 5'-triphosphate | 4.0 ± 0.6 | 2.7 ± 0.3 | 2.2 ± 0.5 | 2.5 ± 0.4 |
| 2'-Deoxycytidine-5'-monophosphate | 6.8 ± 0.6 | 1.5 ± 0.4 | 2.4 ± 0.5 | 2.2 ± 0.4 |
| 2-Deoxyguanosine | 11.5 ± 0.8 | 3.4 ± 0.2 | 5.8 ± 0.4 | 2.5 ± 0.6 |
| 2-hydroxybutyric acid | 12.7 ± 0.9 | 4.4 ± 0.5 | 5.8 ± 0.3 | 7.0 ± 0.4 |
| 2-hydroxyglutaric acid | 6.1 ± 0.7 | 3.9 ± 0.6 | 6.2 ± 0.4 | 7.7 ± 0.9 |
| 2-Hydroxyisocaproic acid | 10.4 ± 0.9 | 2.6 ± 0.4 | 4.1 ± 0.4 | 1.7 ± 0.3 |
| 2-ketobutyric acid | 6.0 ± 0.9 | 7.6 ± 0.6 | 4.7 ± 0.5 | 1.9 ± 0.4 |
| 2-oxoadipic acid | 4.8 ± 0.8 | 6.9 ± 0.4 | 7.3 ± 0.8 | 1.5 ± 0.3 |
| 2-Phenylglycine | 5.7 ± 0.2 | 1.9 ± 0.3 | 1.7 ± 0.3 | 7.0 ± 0.2 |
| 2-phosphoglycerate | 12.4 ± 0.6 | 7.1 ± 0.7 | 3.0 ± 0.5 | 1.8 ± 0.4 |
| 3-hydroxy-3-methyl-2-oxobutanoate | 8.2 ± 0.6 | 2.1 ± 0.4 | 6.2 ± 0.8 | 3.5 ± 0.7 |
| 3-hydroxybutyric acid | 5.5 ± 0.5 | 5.0 ± 0.8 | 7.7 ± 0.4 | 3.1 ± 0.4 |
| 3-Methylhistidine | 7.6 ± 1.0 | 6.7 ± 0.8 | 3.3 ± 0.6 | 4.4 ± 0.8 |
| 3-Phosphoglyceraldehyde | 11.8 ± 0.3 | 6.6 ± 0.9 | 3.6 ± 0.4 | 7.4 ± 0.7 |
| 3-phosphoglycerate | 11.3 ± 0.4 | 4.0 ± 0.8 | 4.2 ± 0.6 | 7.2 ± 0.8 |
| 4-amino-butyric acid | 3.5 ± 0.5 | 6.7 ± 0.7 | 6.8 ± 0.6 | 6.6 ± 0.3 |
| 4-aspartyl-phosphate | 11.1 ± 0.5 | 7.9 ± 0.3 | 6.1 ± 0.6 | 7.4 ± 0.3 |
| 4-Hydroxybenzoic acid | 3.9 ± 0.4 | 8.0 ± 0.4 | 6.5 ± 0.6 | 7.1 ± 0.5 |
| 4-Hydroxyphenylacetylglycine | 12.3 ± 0.6 | 3.0 ± 0.4 | 4.3 ± 0.5 | 4.5 ± 0.5 |
| 4-methyl-2-oxopentanoate | 12.5 ± 0.3 | 7.3 ± 0.7 | 5.9 ± 0.9 | 2.6 ± 0.6 |
| 4-Oxoglutaramate | 9.8 ± 0.9 | 7.9 ± 0.6 | 4.1 ± 0.4 | 6.3 ± 0.7 |
| 5-Hydroxyindolepyruvate | 3.7 ± 0.7 | 5.5 ± 0.3 | 2.5 ± 0.2 | 4.8 ± 0.7 |
| 5-Hydroxylysine | 11.1 ± 0.8 | 4.0 ± 0.5 | 2.8 ± 0.3 | 7.1 ± 0.5 |
| 5-Hydroxy-Tryptophan | 3.3 ± 0.6 | 3.7 ± 0.5 | 2.2 ± 0.4 | 5.7 ± 0.8 |
| 5-Phosphoribosylamine | 10.4 ± 0.7 | 1.9 ± 0.3 | 7.1 ± 0.7 | 4.6 ± 0.9 |
| 6-phosphogluconate | 8.6 ± 0.6 | 1.9 ± 0.3 | 1.5 ± 0.3 | 3.6 ± 0.5 |
| Acetylcarnitine | 8.1 ± 0.7 | 7.6 ± 0.9 | 4.2 ± 0.7 | 1.6 ± 0.3 |
| Aconitic Acid | 4.1 ± 0.2 | 2.6 ± 0.4 | 5.6 ± 0.9 | 1.9 ± 0.2 |
| Adenine | 11.4 ± 0.4 | 6.0 ± 1.0 | 6.5 ± 0.8 | 2.8 ± 0.4 |
| Adenosine | 10.5 ± 0.6 | 4.6 ± 0.5 | 1.5 ± 0.3 | 3.1 ± 0.3 |
| Adenosine 3',5'-cyclic monophosphate | 10.2 ± 0.9 | 3.4 ± 0.6 | 3.8 ± 0.5 | 3.6 ± 0.2 |
| Adenosine Monophosphate | 12.2 ± 0.4 | 7.0 ± 0.4 | 6.5 ± 0.7 | 7.1 ± 0.6 |
| Adenosine-5'-diphosphate | 5.0 ± 1.0 | 7.2 ± 0.2 | 6.7 ± 0.8 | 7.9 ± 0.7 |
| Adenosine-5'-triphosphate | 6.9 ± 0.4 | 1.6 ± 0.3 | 2.9 ± 0.2 | 6.1 ± 0.4 |
| Adenylosuccinate | 5.3 ± 0.7 | 2.1 ± 0.3 | 5.3 ± 0.3 | 7.7 ± 0.6 |
| Alanine | 8.2 ± 1.0 | 4.6 ± 0.9 | 2.6 ± 0.4 | 4.1 ± 0.4 |
| alpha-ketoglutaric acid | 8.5 ± 0.2 | 3.0 ± 0.7 | 2.3 ± 0.4 | 2.1 ± 0.4 |
| Anserine | 6.4 ± 0.7 | 5.9 ± 0.2 | 6.1 ± 0.9 | 3.8 ± 0.9 |
| Arginine | 12.1 ± 0.8 | 7.9 ± 0.7 | 6.7 ± 0.4 | 5.5 ± 0.4 |
| Argininosuccinic Acid | 5.2 ± 0.6 | 3.2 ± 0.3 | 3.6 ± 0.3 | 4.0 ± 0.2 |
| Asparagine | 8.8 ± 0.6 | 4.6 ± 0.5 | 6.9 ± 0.7 | 7.4 ± 0.9 |
| Aspartic acid | 9.4 ± 0.9 | 4.3 ± 0.3 | 1.5 ± 0.2 | 5.9 ± 0.3 |
| Asymmetric dimethylarginine | 7.7 ± 0.4 | 3.8 ± 0.9 | 3.2 ± 0.6 | 3.6 ± 0.3 |
| Azelaic acid | 6.0 ± 0.5 | 5.6 ± 0.5 | 4.1 ± 0.3 | 3.9 ± 0.5 |
| beta-Alanine | 12.5 ± 0.3 | 1.8 ± 0.4 | 7.4 ± 0.4 | 4.1 ± 0.9 |
| Betaine | 7.9 ± 0.4 | 3.9 ± 0.8 | 6.1 ± 0.3 | 2.6 ± 0.3 |
| Biopterin | 7.8 ± 0.9 | 2.3 ± 0.4 | 1.9 ± 0.4 | 2.1 ± 0.3 |
| Biotin | 5.9 ± 1.0 | 5.7 ± 0.5 | 7.2 ± 0.5 | 7.5 ± 0.7 |
| Carnitine | 12.8 ± 0.5 | 8.0 ± 0.2 | 5.1 ± 0.4 | 3.6 ± 0.3 |
| Carnosine | 11.6 ± 0.8 | 3.5 ± 0.3 | 4.3 ± 0.7 | 4.3 ± 0.5 |
| CDP-choline | 9.3 ± 0.9 | 7.0 ± 0.7 | 5.4 ± 0.9 | 1.8 ± 0.4 |
| cGMP | 3.7 ± 0.7 | 1.8 ± 0.4 | 3.7 ± 0.5 | 1.6 ± 0.2 |
| Choline | 8.0 ± 0.3 | 5.0 ± 0.6 | 1.6 ± 0.4 | 4.8 ± 0.8 |
| Cinnamic acid | 3.3 ± 0.6 | 1.8 ± 0.4 | 3.4 ± 0.8 | 5.7 ± 0.4 |
| Citraconic acid | 11.3 ± 0.8 | 7.2 ± 0.8 | 7.1 ± 0.6 | 1.6 ± 0.2 |
| Citramalic acid | 12.1 ± 0.4 | 3.1 ± 0.5 | 4.5 ± 1.0 | 3.2 ± 0.6 |
| Citric Acid | 8.4 ± 0.6 | 4.2 ± 0.7 | 6.7 ± 1.0 | 3.4 ± 0.7 |
| Citrulline | 12.9 ± 0.8 | 5.2 ± 0.2 | 5.4 ± 1.0 | 6.0 ± 0.7 |
| Creatine | 11.1 ± 1.0 | 1.6 ± 0.4 | 2.7 ± 0.6 | 5.8 ± 0.5 |
| Creatinine | 4.2 ± 0.5 | 5.0 ± 1.0 | 6.3 ± 0.2 | 3.1 ± 0.5 |
| Cystathionine | 4.6 ± 0.4 | 7.0 ± 0.4 | 6.1 ± 0.9 | 6.9 ± 0.5 |
| Cysteamine | 5.4 ± 0.9 | 2.3 ± 0.6 | 2.0 ± 0.3 | 5.0 ± 0.7 |
| Cysteine | 6.2 ± 0.5 | 6.0 ± 0.6 | 4.2 ± 0.7 | 2.2 ± 0.2 |
| Cysteinylglycine | 12.5 ± 0.5 | 7.5 ± 0.7 | 3.9 ± 0.9 | 6.8 ± 0.9 |
| Cystine | 5.2 ± 0.2 | 5.4 ± 0.9 | 3.9 ± 0.6 | 2.9 ± 0.3 |
| Cytidine | 5.3 ± 0.9 | 6.4 ± 0.5 | 4.8 ± 0.3 | 1.5 ± 0.4 |
| Cytidine 3',5'-cyclic monophosphate | 9.0 ± 0.4 | 5.1 ± 0.7 | 4.5 ± 0.7 | 4.1 ± 0.5 |
| Cytidine 5'-diphosphate | 8.5 ± 0.4 | 7.1 ± 0.5 | 6.1 ± 0.3 | 4.2 ± 0.7 |
| Cytidine 5'-triphosphate | 11.3 ± 0.3 | 2.8 ± 0.4 | 6.0 ± 0.4 | 7.6 ± 0.7 |
| Cytidine monophosphate | 5.2 ± 0.3 | 5.2 ± 0.9 | 4.5 ± 0.3 | 4.6 ± 0.9 |
| Cytosine | 3.3 ± 0.8 | 7.3 ± 0.5 | 6.1 ± 0.6 | 2.8 ± 0.2 |
| dADP | 5.0 ± 0.7 | 5.1 ± 0.4 | 2.0 ± 0.3 | 2.8 ± 0.5 |
| Diethanolamine | 10.7 ± 0.3 | 6.5 ± 0.3 | 3.8 ± 0.8 | 2.6 ± 0.5 |
| Dihydrobiopterin | 6.6 ± 0.3 | 4.7 ± 0.7 | 7.6 ± 0.9 | 6.3 ± 0.5 |
| Dihydroneopterin triphosphate | 12.7 ± 0.7 | 2.1 ± 0.3 | 4.6 ± 0.3 | 7.3 ± 0.4 |
| Dihydroxyacetonephosphate | 5.6 ± 0.7 | 6.7 ± 0.7 | 5.2 ± 0.8 | 7.8 ± 1.0 |
| Dimethylglycine | 4.5 ± 0.5 | 4.0 ± 0.7 | 4.0 ± 0.2 | 6.3 ± 0.6 |
| Ectoine | 10.3 ± 0.6 | 2.4 ± 0.5 | 2.8 ± 0.6 | 7.2 ± 0.7 |
| Erythrose-4-phosphate | 6.6 ± 0.6 | 7.1 ± 0.3 | 6.5 ± 0.5 | 6.6 ± 0.8 |
| FAD | 9.9 ± 0.4 | 2.9 ± 0.3 | 5.3 ± 0.5 | 1.9 ± 0.3 |
| Folic acid | 8.2 ± 0.3 | 7.3 ± 0.5 | 5.9 ± 0.4 | 7.7 ± 0.4 |
| Fructose-6-phosphate | 4.5 ± 0.5 | 5.1 ± 0.2 | 2.0 ± 0.4 | 2.1 ± 0.5 |
| Fumaric Acid | 3.6 ± 0.3 | 5.0 ± 0.2 | 3.0 ± 0.2 | 7.0 ± 0.3 |
| Galactosamine | 11.8 ± 0.8 | 7.7 ± 0.3 | 3.4 ± 0.6 | 2.7 ± 0.6 |
| Galactose-1-phosphate | 3.2 ± 0.5 | 7.9 ± 0.3 | 6.5 ± 0.4 | 2.7 ± 0.6 |
| Galactose-6-Phosphate | 4.0 ± 0.8 | 1.8 ± 0.4 | 5.3 ± 0.7 | 3.7 ± 0.6 |
| gamma-Glutamylcysteine | 5.6 ± 0.6 | 1.5 ± 0.3 | 5.7 ± 1.0 | 7.3 ± 0.5 |
| Glucosamine | 11.7 ± 0.8 | 3.9 ± 0.3 | 3.3 ± 0.6 | 7.6 ± 0.7 |
| Glucosamine-6-Phosphate | 11.0 ± 0.7 | 5.5 ± 0.5 | 4.2 ± 0.2 | 4.5 ± 1.0 |
| Glucose | 7.1 ± 0.6 | 2.2 ± 0.3 | 5.1 ± 1.0 | 3.7 ± 0.8 |
| Glucose-1-phosphate | 12.8 ± 0.4 | 2.4 ± 0.4 | 7.4 ± 1.0 | 3.0 ± 0.4 |
| Glucose-6-Phosphate | 3.4 ± 0.8 | 2.7 ± 0.2 | 1.7 ± 0.3 | 3.3 ± 0.3 |
| Glucuronic acid | 12.5 ± 0.7 | 3.0 ± 0.4 | 4.6 ± 0.3 | 2.2 ± 0.5 |
| Glutamic acid | 6.9 ± 0.6 | 2.7 ± 0.5 | 5.7 ± 0.5 | 4.9 ± 0.2 |
| Glutamine | 12.1 ± 0.6 | 5.2 ± 0.7 | 7.2 ± 0.5 | 6.3 ± 0.5 |
| Glutathione | 8.5 ± 0.7 | 3.5 ± 0.4 | 7.3 ± 0.7 | 3.6 ± 0.5 |
| Glutathionylspermine | 9.0 ± 0.5 | 2.3 ± 0.4 | 2.3 ± 0.2 | 3.9 ± 0.3 |
| Glycerophosphocholine | 8.6 ± 1.0 | 5.5 ± 0.4 | 2.1 ± 0.4 | 2.3 ± 0.5 |
| Glycine | 5.3 ± 0.5 | 7.4 ± 0.7 | 5.8 ± 0.6 | 1.9 ± 0.3 |
| Glycolic acid | 10.8 ± 0.4 | 4.3 ± 0.7 | 3.3 ± 0.7 | 6.3 ± 0.9 |
| Guanidinoacetic acid | 12.2 ± 0.8 | 3.1 ± 0.4 | 4.9 ± 0.6 | 1.7 ± 0.3 |
| Guanidinobutanoic acid | 4.0 ± 0.3 | 4.6 ± 0.9 | 7.6 ± 0.6 | 6.3 ± 0.3 |
| Guanine | 6.5 ± 0.9 | 6.7 ± 0.6 | 4.1 ± 0.3 | 3.0 ± 0.6 |
| Guanosine | 8.6 ± 1.0 | 5.1 ± 0.5 | 5.0 ± 0.3 | 2.9 ± 0.2 |
| Guanosine 5'-diphosphate | 9.1 ± 0.3 | 1.9 ± 0.5 | 2.9 ± 0.5 | 3.8 ± 0.6 |
| Guanosine-5'-monophosphate | 9.2 ± 0.4 | 4.4 ± 1.0 | 3.9 ± 0.8 | 5.2 ± 0.9 |
| Hercynine | 10.8 ± 0.4 | 2.7 ± 0.4 | 6.0 ± 1.0 | 5.0 ± 0.7 |
| Hippuric acid | 9.3 ± 0.4 | 8.0 ± 0.9 | 5.7 ± 0.5 | 6.7 ± 0.5 |
| Histamine | 3.3 ± 0.7 | 7.2 ± 0.4 | 6.8 ± 0.5 | 1.8 ± 0.4 |
| Histidine | 3.2 ± 0.4 | 6.5 ± 0.5 | 5.8 ± 0.2 | 3.9 ± 0.5 |
| Homoalanine | 4.9 ± 0.7 | 3.5 ± 0.4 | 5.4 ± 0.9 | 6.6 ± 0.8 |
| Homoarginine | 6.6 ± 0.8 | 1.9 ± 0.4 | 5.1 ± 0.6 | 1.5 ± 0.3 |
| Homocysteine | 6.4 ± 0.9 | 3.4 ± 0.3 | 3.1 ± 0.2 | 4.8 ± 0.5 |
| Homogentisic acid | 3.6 ± 0.5 | 2.8 ± 0.2 | 3.5 ± 0.5 | 7.6 ± 0.3 |
| Homoleucine | 7.8 ± 0.9 | 7.1 ± 0.4 | 5.4 ± 0.7 | 2.7 ± 0.6 |
| Homovanillic acid | 3.8 ± 0.2 | 4.7 ± 0.8 | 7.2 ± 0.6 | 6.4 ± 0.7 |
| Hydantoin-5-propionate | 8.9 ± 0.4 | 3.4 ± 0.6 | 7.2 ± 0.4 | 7.5 ± 0.7 |
| Hypotaurine | 7.7 ± 0.2 | 3.6 ± 0.8 | 2.5 ± 0.3 | 8.0 ± 0.9 |
| Hypoxanthine | 10.2 ± 0.7 | 3.8 ± 0.2 | 3.4 ± 0.6 | 1.7 ± 0.4 |
| Indole-3-acetamide | 9.1 ± 0.4 | 1.8 ± 0.3 | 4.4 ± 1.0 | 5.0 ± 0.2 |
| Inosine | 3.2 ± 0.3 | 6.3 ± 0.9 | 7.5 ± 0.2 | 6.6 ± 0.2 |
| Inosine 5'-monophosphate | 7.3 ± 0.9 | 6.9 ± 0.3 | 7.7 ± 0.5 | 4.7 ± 1.0 |
| Isocitric Acid | 11.8 ± 0.4 | 3.1 ± 0.6 | 3.6 ± 0.7 | 3.2 ± 0.5 |
| Isoleucine | 11.6 ± 0.3 | 2.4 ± 0.3 | 4.8 ± 1.0 | 3.7 ± 0.6 |
| Lactic Acid | 12.0 ± 1.0 | 2.2 ± 0.4 | 7.9 ± 1.0 | 5.9 ± 0.9 |
| Leucine | 6.0 ± 0.9 | 7.3 ± 0.3 | 4.6 ± 0.4 | 3.5 ± 0.4 |
| Levulinic acid | 5.6 ± 0.9 | 2.9 ± 0.5 | 3.2 ± 0.6 | 4.1 ± 0.8 |
| Lysine | 12.2 ± 0.8 | 5.1 ± 0.4 | 8.0 ± 0.8 | 6.1 ± 0.7 |
| Malic Acid | 7.0 ± 0.7 | 4.9 ± 0.9 | 2.2 ± 0.3 | 7.2 ± 1.0 |
| Mannose-1 with Mannose-6-Phosphate | 3.8 ± 0.5 | 6.4 ± 0.5 | 5.4 ± 0.9 | 6.7 ± 0.6 |
| Mesaconic acid | 4.4 ± 0.5 | 5.5 ± 0.7 | 6.8 ± 0.8 | 2.2 ± 0.3 |
| Methionine | 8.1 ± 0.7 | 6.6 ± 0.7 | 1.7 ± 0.2 | 5.0 ± 0.8 |
| Methionine sulfoxide | 10.5 ± 0.2 | 1.9 ± 0.5 | 6.1 ± 0.9 | 1.5 ± 0.3 |
| Methylmalonic Acid | 12.4 ± 0.3 | 5.2 ± 0.5 | 1.9 ± 0.3 | 4.7 ± 0.4 |
| N1-Acetylspermidine | 11.1 ± 0.3 | 5.9 ± 0.9 | 5.6 ± 0.4 | 4.9 ± 0.3 |
| N1-Acetylspermine | 6.7 ± 0.4 | 6.1 ± 0.4 | 4.0 ± 0.3 | 1.7 ± 0.3 |
| N8-Acetylspermidine | 12.7 ± 0.8 | 4.1 ± 0.9 | 6.8 ± 0.6 | 5.5 ± 0.3 |
| N-Acetyl-Aspartyl-Glutamate | 3.3 ± 0.5 | 2.5 ± 0.5 | 5.8 ± 0.6 | 6.3 ± 0.5 |
| N-Acetyl-Aspartyl-Glutamyl-Glutamate | 5.0 ± 0.4 | 6.6 ± 0.8 | 7.3 ± 0.2 | 5.4 ± 0.8 |
| N-Acetyl-Galactosamine | 3.0 ± 0.7 | 3.6 ± 0.6 | 7.3 ± 1.0 | 3.3 ± 0.4 |
| N-Acetyl-Leucine | 8.6 ± 0.9 | 1.9 ± 0.3 | 3.9 ± 0.2 | 6.0 ± 0.3 |
| N-Acetyl-Methionine | 3.7 ± 0.3 | 2.5 ± 0.4 | 7.1 ± 0.8 | 2.1 ± 0.3 |
| N-Acetylneuraminic acid | 6.2 ± 0.5 | 4.4 ± 1.0 | 3.6 ± 0.2 | 4.2 ± 0.3 |
| N-Acetyl-Proline | 11.1 ± 0.6 | 7.2 ± 0.7 | 5.1 ± 0.9 | 5.0 ± 0.8 |
| N-Acetyl-Putrescine | 6.3 ± 0.5 | 4.9 ± 0.9 | 6.1 ± 0.2 | 5.2 ± 1.0 |
| N-Acetyl-Serine | 9.0 ± 0.2 | 5.7 ± 0.9 | 2.1 ± 0.5 | 2.5 ± 0.5 |
| N-Acetyl-Serotonin | 5.9 ± 0.5 | 5.2 ± 0.9 | 2.1 ± 0.4 | 1.5 ± 0.2 |
| N-Acyl-Homoserine-lactone | 6.2 ± 0.4 | 5.1 ± 0.9 | 3.5 ± 0.7 | 3.5 ± 0.6 |
| NAD | 8.4 ± 0.9 | 5.9 ± 0.4 | 7.7 ± 0.6 | 2.0 ± 0.4 |
| NADH | 4.4 ± 0.4 | 1.6 ± 0.3 | 5.5 ± 0.4 | 5.3 ± 0.3 |
| NADP | 6.9 ± 0.9 | 5.0 ± 0.3 | 5.3 ± 0.3 | 1.9 ± 0.4 |
| NADPH | 10.6 ± 0.3 | 7.0 ± 0.6 | 7.4 ± 0.4 | 2.6 ± 0.6 |
| N-alpha-Acetyl-lysine | 5.2 ± 0.3 | 6.2 ± 0.8 | 2.3 ± 0.4 | 5.9 ± 0.9 |
| N-alpha-Acetyl-Ornithine | 11.1 ± 0.9 | 5.4 ± 0.3 | 4.6 ± 0.3 | 6.4 ± 0.7 |
| N-Carbamoyl-Proline | 9.8 ± 0.6 | 2.9 ± 0.6 | 6.6 ± 0.5 | 2.1 ± 0.3 |
| N-Carbamoyl-Sarcosine | 8.3 ± 0.8 | 3.7 ± 0.5 | 7.2 ± 0.3 | 3.4 ± 0.7 |
| N-Epsilon-Acetyl-lysine | 12.6 ± 0.9 | 4.8 ± 0.3 | 4.6 ± 0.5 | 5.9 ± 0.2 |
| Nicotinamide | 9.1 ± 0.8 | 4.4 ± 0.9 | 2.0 ± 0.4 | 3.3 ± 0.8 |
| N-Methyl-Aspartic Acid | 7.6 ± 0.9 | 7.5 ± 0.5 | 2.2 ± 0.5 | 3.2 ± 0.6 |
| O-Phosphoethanolamine | 9.4 ± 0.2 | 7.8 ± 0.3 | 4.7 ± 0.9 | 1.5 ± 0.4 |
| Ophthalmic acid | 12.6 ± 0.4 | 6.3 ± 0.5 | 7.7 ± 0.8 | 5.2 ± 1.0 |
| Ornithine | 7.0 ± 1.0 | 1.4 ± 0.2 | 5.9 ± 0.5 | 1.7 ± 0.3 |
| Orotidine 5'-monophosphate | 6.4 ± 0.7 | 4.9 ± 0.3 | 2.7 ± 0.3 | 5.4 ± 0.2 |
| Oxaloacetate | 6.4 ± 0.6 | 4.5 ± 0.5 | 3.9 ± 0.6 | 4.3 ± 0.8 |
| Pantothenic acid | 8.1 ± 0.7 | 3.8 ± 0.3 | 2.9 ± 0.3 | 7.2 ± 0.4 |
| Phenylalanine | 9.8 ± 0.6 | 8.0 ± 0.3 | 4.7 ± 0.3 | 6.9 ± 0.6 |
| Phenylpyruvate | 7.0 ± 0.4 | 2.8 ± 0.6 | 4.2 ± 0.5 | 1.6 ± 0.4 |
| Phosphodimethylethanolamine | 11.2 ± 0.7 | 7.3 ± 0.8 | 3.7 ± 0.5 | 5.8 ± 0.7 |
| Phosphoenolpyruvate | 6.8 ± 1.0 | 3.8 ± 0.4 | 7.8 ± 0.7 | 3.6 ± 0.8 |
| Phosphorylcholine | 12.0 ± 0.4 | 3.5 ± 0.7 | 4.6 ± 1.0 | 4.7 ± 0.3 |
| Pipecolic acid | 11.3 ± 0.2 | 2.1 ± 0.4 | 3.0 ± 0.3 | 4.8 ± 0.7 |
| Proline | 6.9 ± 0.7 | 3.5 ± 0.6 | 5.9 ± 1.0 | 3.1 ± 0.5 |
| Putrescine | 11.9 ± 0.5 | 5.9 ± 0.5 | 2.4 ± 0.2 | 7.0 ± 0.7 |
| Pyridoxal | 3.2 ± 0.3 | 6.9 ± 0.5 | 3.9 ± 0.3 | 1.7 ± 0.4 |
| Pyridoxamine | 6.4 ± 0.9 | 3.7 ± 0.5 | 2.3 ± 0.3 | 3.1 ± 0.3 |
| Pyridoxine | 9.3 ± 0.7 | 6.7 ± 0.4 | 2.2 ± 0.3 | 2.3 ± 0.5 |
| Pyroglutamic acid | 10.1 ± 0.7 | 2.6 ± 0.4 | 7.2 ± 0.5 | 2.4 ± 0.5 |
| Pyrrolysine | 6.3 ± 0.9 | 5.3 ± 0.6 | 3.4 ± 0.8 | 6.9 ± 0.9 |
| Pyruvic Acid | 12.0 ± 0.2 | 7.6 ± 0.9 | 7.1 ± 0.7 | 7.2 ± 0.6 |
| Riboflavin | 12.3 ± 0.5 | 4.0 ± 0.7 | 3.5 ± 0.7 | 7.6 ± 0.6 |
| Ribose-5-Phosphate | 11.4 ± 0.8 | 3.8 ± 0.3 | 1.8 ± 0.4 | 4.0 ± 0.6 |
| Ribulose-1-Phosphate | 11.2 ± 0.7 | 2.4 ± 0.3 | 4.4 ± 0.8 | 2.4 ± 0.4 |
| Ribulose-5-Phosphate | 4.7 ± 0.9 | 7.3 ± 0.4 | 4.7 ± 0.9 | 7.2 ± 0.6 |
| Saccharopine | 4.8 ± 0.3 | 6.7 ± 0.2 | 7.4 ± 0.4 | 4.0 ± 1.0 |
| S-Adenosylhomocysteine | 6.9 ± 0.9 | 6.5 ± 0.3 | 1.7 ± 0.4 | 7.4 ± 0.5 |
| S-Adenosylmethioninamine | 10.5 ± 0.6 | 5.3 ± 0.9 | 4.9 ± 0.8 | 5.2 ± 0.8 |
| S-Adenosylmethionine | 9.3 ± 0.3 | 6.2 ± 0.3 | 1.7 ± 0.3 | 5.6 ± 0.6 |
| Sarcosine | 12.4 ± 0.4 | 5.2 ± 0.7 | 7.5 ± 0.3 | 1.5 ± 0.3 |
| Sedoheptulose-7-phosphate | 5.1 ± 0.8 | 3.4 ± 0.3 | 4.1 ± 0.4 | 7.7 ± 0.8 |
| Selenohomocysteine | 3.5 ± 0.8 | 6.5 ± 0.9 | 6.0 ± 0.7 | 7.0 ± 0.8 |
| Selenomethionine | 6.1 ± 0.3 | 4.0 ± 0.9 | 2.2 ± 0.5 | 4.4 ± 1.0 |
| Serine | 8.7 ± 0.4 | 2.0 ± 0.3 | 4.6 ± 0.8 | 6.9 ± 0.3 |
| S-Methyl-Thioinosine | 6.6 ± 0.2 | 5.1 ± 0.9 | 3.0 ± 0.4 | 3.7 ± 0.6 |
| Spermidine | 12.8 ± 0.9 | 6.8 ± 0.5 | 3.5 ± 0.8 | 5.5 ± 0.8 |
| Spermine | 10.1 ± 0.3 | 7.1 ± 0.4 | 6.6 ± 0.2 | 5.0 ± 0.9 |
| Succinate | 5.0 ± 0.2 | 3.7 ± 0.7 | 6.2 ± 0.9 | 7.3 ± 0.7 |
| Sucrose | 6.8 ± 0.5 | 7.4 ± 0.9 | 7.1 ± 0.4 | 3.0 ± 0.4 |
| Symmetric dimethylarginine | 8.6 ± 0.3 | 6.1 ± 0.4 | 7.1 ± 0.2 | 3.8 ± 0.7 |
| Taurine | 5.2 ± 0.7 | 2.2 ± 0.3 | 7.6 ± 0.7 | 6.2 ± 0.2 |
| Tetrahydroneopterin | 6.9 ± 0.9 | 5.0 ± 0.6 | 1.5 ± 0.3 | 5.6 ± 0.5 |
| Thiamine | 7.6 ± 0.6 | 2.1 ± 0.4 | 3.6 ± 0.3 | 1.6 ± 0.3 |
| Thiamine monophosphate | 6.2 ± 0.9 | 4.1 ± 0.5 | 2.9 ± 0.7 | 4.9 ± 0.8 |
| Threonic acid | 9.9 ± 1.0 | 2.8 ± 0.4 | 1.8 ± 0.3 | 4.8 ± 0.7 |
| Threonine | 4.7 ± 0.7 | 1.5 ± 0.3 | 6.5 ± 0.8 | 5.0 ± 0.7 |
| Thymidine | 4.1 ± 0.7 | 5.8 ± 0.5 | 3.4 ± 0.5 | 3.3 ± 0.4 |
| Thymidine Monophosphate | 12.6 ± 0.5 | 4.2 ± 0.4 | 3.2 ± 0.7 | 5.3 ± 0.6 |
| Thymine | 12.5 ± 0.6 | 3.4 ± 0.7 | 4.9 ± 0.7 | 7.3 ± 0.3 |
| Trigonelleine | 12.0 ± 0.8 | 6.5 ± 0.9 | 3.9 ± 0.3 | 8.0 ± 0.5 |
| Trimethylamine N-oxide | 10.9 ± 0.5 | 1.4 ± 0.2 | 3.9 ± 0.6 | 6.8 ± 0.5 |
| Trimethyllysine | 3.5 ± 0.7 | 1.6 ± 0.4 | 4.9 ± 0.5 | 5.3 ± 0.7 |
| Tryptophan | 10.4 ± 0.4 | 1.5 ± 0.3 | 3.4 ± 0.2 | 7.7 ± 0.4 |
| Tyramine | 11.2 ± 0.7 | 3.8 ± 0.8 | 2.5 ± 0.4 | 2.0 ± 0.4 |
| Tyrosine | 7.9 ± 0.5 | 2.6 ± 0.3 | 3.4 ± 0.7 | 4.1 ± 1.0 |
| Uracil | 11.0 ± 0.8 | 6.2 ± 0.5 | 6.6 ± 0.4 | 3.1 ± 0.5 |
| Uridine 5'-diphosphate | 6.4 ± 0.6 | 3.7 ± 0.7 | 8.0 ± 0.9 | 7.4 ± 0.4 |
| Uridine Monophosphate | 10.6 ± 0.9 | 3.1 ± 0.4 | 3.3 ± 0.3 | 3.2 ± 0.7 |
| Uridine-5-Diphosphogalactose | 12.7 ± 0.4 | 4.6 ± 0.4 | 6.1 ± 0.5 | 3.8 ± 0.6 |
| Urocanic acid | 8.5 ± 0.8 | 5.6 ± 0.4 | 2.4 ± 0.2 | 6.1 ± 0.4 |
| UTP | 11.2 ± 0.7 | 6.2 ± 0.7 | 2.4 ± 0.3 | 2.8 ± 0.2 |
| Valine | 4.9 ± 0.9 | 6.3 ± 0.3 | 4.5 ± 0.2 | 4.8 ± 0.7 |
| XMP | 8.8 ± 0.4 | 2.1 ± 0.3 | 7.4 ± 1.0 | 4.5 ± 0.3 |
| Xylulose-5-Phosphate | 12.1 ± 0.7 | 6.0 ± 0.6 | 7.8 ± 0.9 | 2.5 ± 0.3 |

### **Table S18** | Inter – day precision (as % RSD) (Tumor tissue extract as matrix, n = 5). The concentrations of QC_LLOQ_, QC_Low_, QC_Mid_ and QC_High_ were 10 ng/mL, 25 ng/mL, 100 ng/mL and 500 ng/mL respectively.

| Metabolite | QC_LLOQ_ | QC_Low_ | QC_Mid_ | QC_High_ |
| --- | --- | --- | --- | --- |
| 1-Methylguanine | 6.6 ± 0.7 | 3.1 ± 1.1 | 3.1 ± 0.7 | 2.1 ± 0.8 |
| 1-Methylhistidine | 10.0 ± 0.5 | 2.0 ± 0.8 | 5.3 ± 0.5 | 3.6 ± 0.4 |
| 1-Methylnicotinamide | 8.8 ± 0.7 | 6.3 ± 1.1 | 2.9 ± 0.8 | 6.9 ± 0.7 |
| 2-Aminoadipic Acid | 8.0 ± 0.7 | 4.2 ± 0.6 | 7.4 ± 0.7 | 2.8 ± 0.5 |
| 2-Deoxyadenosine | 9.9 ± 0.2 | 7.0 ± 1.1 | 3.1 ± 0.4 | 5.3 ± 0.6 |
| 2'-Deoxyadenosine 5'-monophosphate | 6.6 ± 0.6 | 3.9 ± 1.2 | 2.2 ± 0.5 | 4.0 ± 0.3 |
| 2'-deoxyadenosine 5'-triphosphate | 6.7 ± 0.8 | 6.2 ± 0.8 | 7.2 ± 1.1 | 4.1 ± 1.2 |
| 2'-Deoxycytidine-5'-monophosphate | 8.2 ± 0.3 | 4.2 ± 1.0 | 4.8 ± 0.9 | 2.3 ± 0.6 |
| 2-Deoxyguanosine | 10.5 ± 0.9 | 6.0 ± 0.7 | 4.6 ± 0.1 | 6.7 ± 0.9 |
| 2-hydroxybutyric acid | 9.0 ± 1.1 | 2.1 ± 1.1 | 6.5 ± 0.7 | 4.5 ± 0.7 |
| 2-hydroxyglutaric acid | 7.3 ± 0.4 | 6.9 ± 0.5 | 3.9 ± 0.5 | 5.6 ± 0.3 |
| 2-Hydroxyisocaproic acid | 8.8 ± 0.6 | 6.9 ± 0.3 | 7.9 ± 0.7 | 3.5 ± 0.8 |
| 2-ketobutyric acid | 8.0 ± 0.9 | 4.4 ± 1.0 | 3.0 ± 0.9 | 3.2 ± 0.6 |
| 2-oxoadipic acid | 7.0 ± 1.1 | 4.3 ± 0.9 | 8.0 ± 0.2 | 7.9 ± 0.9 |
| 2-Phenylglycine | 8.9 ± 0.5 | 3.3 ± 0.3 | 7.8 ± 0.2 | 3.5 ± 0.7 |
| 2-phosphoglycerate | 8.0 ± 1.0 | 5.3 ± 0.2 | 5.7 ± 0.7 | 6.4 ± 0.6 |
| 3-hydroxy-3-methyl-2-oxobutanoate | 7.6 ± 1.0 | 5.0 ± 0.4 | 5.6 ± 0.1 | 7.6 ± 0.8 |
| 3-hydroxybutyric acid | 6.1 ± 1.1 | 6.6 ± 0.9 | 5.9 ± 0.1 | 7.8 ± 0.4 |
| 3-Methylhistidine | 8.2 ± 0.5 | 2.9 ± 1.0 | 3.0 ± 1.2 | 3.8 ± 1.1 |
| 3-Phosphoglyceraldehyde | 9.1 ± 0.1 | 4.7 ± 0.4 | 6.5 ± 0.4 | 5.3 ± 0.8 |
| 3-phosphoglycerate | 10.3 ± 0.4 | 3.1 ± 1.0 | 4.1 ± 1.2 | 4.0 ± 1.1 |
| 4-amino-butyric acid | 6.7 ± 0.4 | 8.0 ± 0.9 | 6.0 ± 0.4 | 7.9 ± 0.8 |
| 4-aspartyl-phosphate | 6.1 ± 0.5 | 4.5 ± 0.8 | 2.2 ± 0.3 | 6.4 ± 0.2 |
| 4-Hydroxybenzoic acid | 10.4 ± 0.3 | 4.4 ± 0.5 | 3.0 ± 1.1 | 3.0 ± 0.7 |
| 4-Hydroxyphenylacetylglycine | 8.4 ± 0.5 | 4.5 ± 1.0 | 7.5 ± 0.4 | 5.2 ± 0.2 |
| 4-methyl-2-oxopentanoate | 7.6 ± 0.8 | 6.9 ± 1.1 | 3.4 ± 0.3 | 6.1 ± 1.0 |
| 4-Oxoglutaramate | 8.7 ± 0.7 | 6.8 ± 0.5 | 2.9 ± 0.5 | 7.4 ± 0.3 |
| 5-Hydroxyindolepyruvate | 9.7 ± 1.2 | 2.5 ± 1.1 | 4.4 ± 0.7 | 5.1 ± 0.7 |
| 5-Hydroxylysine | 10.6 ± 0.9 | 6.9 ± 0.6 | 3.1 ± 0.2 | 2.8 ± 1.2 |
| 5-Hydroxy-Tryptophan | 8.3 ± 1.1 | 2.5 ± 1.1 | 6.1 ± 0.6 | 4.3 ± 0.6 |
| 5-Phosphoribosylamine | 6.8 ± 0.8 | 6.5 ± 0.9 | 5.6 ± 0.8 | 7.0 ± 0.5 |
| 6-phosphogluconate | 6.9 ± 1.1 | 6.3 ± 0.9 | 3.2 ± 0.8 | 2.3 ± 1.1 |
| Acetylcarnitine | 11.0 ± 0.8 | 3.2 ± 0.9 | 5.6 ± 0.8 | 7.8 ± 0.9 |
| Aconitic Acid | 6.1 ± 0.6 | 7.0 ± 1.1 | 3.2 ± 0.1 | 5.3 ± 0.9 |
| Adenine | 9.7 ± 0.2 | 7.7 ± 0.2 | 5.3 ± 1.1 | 2.5 ± 1.0 |
| Adenosine | 8.4 ± 0.2 | 7.6 ± 0.7 | 4.1 ± 0.4 | 7.0 ± 0.3 |
| Adenosine 3',5'-cyclic monophosphate | 10.4 ± 0.4 | 2.1 ± 0.8 | 5.5 ± 0.5 | 6.6 ± 0.6 |
| Adenosine Monophosphate | 6.7 ± 0.9 | 2.9 ± 0.7 | 4.9 ± 0.4 | 4.9 ± 0.8 |
| Adenosine-5'-diphosphate | 7.6 ± 0.4 | 2.2 ± 0.5 | 5.3 ± 1.0 | 7.9 ± 0.6 |
| Adenosine-5'-triphosphate | 6.5 ± 1.0 | 2.6 ± 1.1 | 7.2 ± 0.7 | 7.2 ± 1.1 |
| Adenylosuccinate | 10.4 ± 1.1 | 5.9 ± 0.3 | 7.4 ± 0.3 | 2.1 ± 0.7 |
| Alanine | 9.8 ± 0.5 | 4.7 ± 0.4 | 5.0 ± 0.2 | 5.8 ± 0.7 |
| alpha-ketoglutaric acid | 6.5 ± 1.0 | 6.0 ± 1.0 | 6.5 ± 1.1 | 2.4 ± 0.7 |
| Anserine | 9.8 ± 0.4 | 3.1 ± 0.1 | 5.2 ± 0.7 | 3.0 ± 0.3 |
| Arginine | 10.9 ± 0.1 | 5.5 ± 0.1 | 5.8 ± 0.8 | 3.1 ± 1.1 |
| Argininosuccinic Acid | 6.2 ± 1.0 | 4.3 ± 0.6 | 3.4 ± 0.6 | 6.9 ± 0.4 |
| Asparagine | 9.4 ± 0.3 | 7.1 ± 0.9 | 3.2 ± 0.1 | 7.1 ± 0.7 |
| Aspartic acid | 10.0 ± 0.5 | 4.7 ± 1.1 | 5.0 ± 0.4 | 7.2 ± 1.2 |
| Asymmetric dimethylarginine | 7.0 ± 1.1 | 2.4 ± 1.0 | 6.1 ± 1.2 | 4.3 ± 0.5 |
| Azelaic acid | 7.5 ± 0.4 | 7.4 ± 0.8 | 5.5 ± 0.5 | 7.4 ± 0.8 |
| beta-Alanine | 7.6 ± 1.0 | 6.5 ± 1.0 | 3.9 ± 0.6 | 2.2 ± 0.7 |
| Betaine | 9.5 ± 0.1 | 6.5 ± 0.4 | 4.4 ± 0.9 | 7.1 ± 1.1 |
| Biopterin | 10.3 ± 1.0 | 2.8 ± 0.7 | 3.8 ± 1.1 | 2.5 ± 0.1 |
| Biotin | 6.9 ± 0.7 | 5.3 ± 0.9 | 3.5 ± 0.4 | 4.2 ± 0.1 |
| Carnitine | 8.5 ± 0.2 | 2.5 ± 1.1 | 7.6 ± 0.2 | 6.1 ± 0.8 |
| Carnosine | 9.3 ± 0.2 | 3.5 ± 0.3 | 2.2 ± 0.7 | 5.0 ± 1.0 |
| CDP-choline | 7.7 ± 1.0 | 3.2 ± 1.1 | 5.4 ± 0.6 | 2.9 ± 0.5 |
| cGMP | 7.8 ± 1.1 | 5.6 ± 0.7 | 7.5 ± 1.0 | 7.3 ± 0.8 |
| Choline | 7.7 ± 0.5 | 3.5 ± 0.4 | 2.4 ± 1.0 | 7.3 ± 0.8 |
| Cinnamic acid | 8.2 ± 0.8 | 4.6 ± 0.5 | 7.4 ± 0.6 | 4.3 ± 0.3 |
| Citraconic acid | 6.6 ± 0.1 | 5.2 ± 0.9 | 3.0 ± 0.9 | 7.1 ± 0.9 |
| Citramalic acid | 8.2 ± 1.0 | 3.2 ± 0.3 | 5.0 ± 0.2 | 2.3 ± 0.3 |
| Citric Acid | 6.9 ± 0.6 | 4.8 ± 0.6 | 4.3 ± 0.6 | 5.8 ± 0.6 |
| Citrulline | 10.6 ± 0.5 | 4.3 ± 0.5 | 7.4 ± 0.7 | 4.0 ± 0.2 |
| Creatine | 9.9 ± 1.0 | 6.5 ± 1.0 | 6.3 ± 0.2 | 7.3 ± 0.4 |
| Creatinine | 9.5 ± 0.6 | 6.5 ± 0.3 | 6.1 ± 0.4 | 5.8 ± 0.3 |
| Cystathionine | 8.8 ± 0.2 | 2.5 ± 0.7 | 6.4 ± 0.4 | 6.4 ± 1.0 |
| Cysteamine | 8.4 ± 0.4 | 7.8 ± 1.1 | 3.0 ± 0.1 | 4.2 ± 1.0 |
| Cysteine | 8.2 ± 0.2 | 7.2 ± 0.7 | 5.6 ± 0.4 | 5.5 ± 0.2 |
| Cysteinylglycine | 6.0 ± 0.6 | 5.4 ± 0.9 | 7.8 ± 0.7 | 5.7 ± 0.4 |
| Cystine | 6.1 ± 1.2 | 7.2 ± 1.2 | 6.9 ± 0.3 | 4.5 ± 0.6 |
| Cytidine | 6.7 ± 0.9 | 6.6 ± 0.9 | 2.2 ± 0.4 | 7.4 ± 0.5 |
| Cytidine 3',5'-cyclic monophosphate | 10.5 ± 1.0 | 8.0 ± 0.3 | 2.2 ± 1.2 | 5.7 ± 0.9 |
| Cytidine 5'-diphosphate | 11.0 ± 1.1 | 4.4 ± 0.7 | 6.3 ± 0.2 | 6.0 ± 0.3 |
| Cytidine 5'-triphosphate | 7.9 ± 1.0 | 2.7 ± 0.3 | 4.2 ± 0.7 | 5.3 ± 0.9 |
| Cytidine monophosphate | 6.0 ± 1.2 | 7.0 ± 0.6 | 4.7 ± 0.6 | 5.3 ± 0.3 |
| Cytosine | 7.1 ± 1.2 | 3.7 ± 1.1 | 7.7 ± 0.2 | 2.5 ± 0.3 |
| dADP | 9.4 ± 0.6 | 4.9 ± 0.6 | 5.7 ± 0.8 | 7.1 ± 0.4 |
| Diethanolamine | 6.5 ± 0.2 | 7.8 ± 1.0 | 3.4 ± 1.1 | 4.8 ± 0.6 |
| Dihydrobiopterin | 6.6 ± 0.8 | 5.8 ± 0.5 | 3.4 ± 0.2 | 3.3 ± 0.4 |
| Dihydroneopterin triphosphate | 11.0 ± 0.2 | 4.0 ± 1.1 | 6.5 ± 0.5 | 2.1 ± 0.8 |
| Dihydroxyacetonephosphate | 8.8 ± 1.0 | 7.1 ± 1.1 | 4.4 ± 1.0 | 2.0 ± 0.7 |
| Dimethylglycine | 6.8 ± 0.4 | 5.7 ± 1.2 | 6.9 ± 0.6 | 7.5 ± 0.7 |
| Ectoine | 6.9 ± 1.1 | 5.7 ± 1.0 | 2.5 ± 0.9 | 6.4 ± 0.5 |
| Erythrose-4-phosphate | 10.0 ± 0.4 | 7.9 ± 0.4 | 3.9 ± 1.0 | 2.2 ± 0.4 |
| FAD | 8.7 ± 0.4 | 4.5 ± 0.9 | 7.3 ± 0.8 | 4.5 ± 0.9 |
| Folic acid | 9.4 ± 0.3 | 3.6 ± 0.4 | 4.9 ± 0.8 | 2.8 ± 0.9 |
| Fructose-6-phosphate | 8.3 ± 1.2 | 5.5 ± 0.6 | 7.7 ± 0.8 | 6.1 ± 0.7 |
| Fumaric Acid | 10.2 ± 0.6 | 4.8 ± 1.0 | 2.0 ± 1.1 | 5.0 ± 0.9 |
| Galactosamine | 10.7 ± 1.0 | 3.3 ± 1.0 | 3.2 ± 0.2 | 5.3 ± 0.3 |
| Galactose-1-phosphate | 10.6 ± 0.3 | 2.6 ± 1.0 | 2.4 ± 0.4 | 6.6 ± 0.6 |
| Galactose-6-Phosphate | 10.4 ± 0.3 | 4.8 ± 0.7 | 6.3 ± 0.4 | 6.8 ± 1.1 |
| gamma-Glutamylcysteine | 10.5 ± 0.9 | 6.2 ± 0.5 | 2.2 ± 0.2 | 3.6 ± 0.1 |
| Glucosamine | 9.3 ± 0.6 | 4.0 ± 0.8 | 7.4 ± 1.2 | 5.2 ± 0.3 |
| Glucosamine-6-Phosphate | 6.2 ± 0.4 | 4.6 ± 0.9 | 5.7 ± 0.9 | 2.4 ± 0.3 |
| Glucose | 10.7 ± 1.0 | 3.3 ± 0.4 | 2.1 ± 0.9 | 3.9 ± 1.1 |
| Glucose-1-phosphate | 8.3 ± 0.7 | 7.8 ± 0.2 | 4.7 ± 1.0 | 7.5 ± 0.8 |
| Glucose-6-Phosphate | 6.9 ± 0.7 | 4.9 ± 0.3 | 3.0 ± 0.6 | 2.7 ± 0.4 |
| Glucuronic acid | 9.5 ± 0.4 | 7.8 ± 0.6 | 2.0 ± 1.2 | 4.4 ± 0.7 |
| Glutamic acid | 8.3 ± 0.3 | 4.8 ± 0.3 | 5.9 ± 0.5 | 6.7 ± 0.1 |
| Glutamine | 6.0 ± 0.9 | 7.8 ± 0.2 | 3.2 ± 0.1 | 5.8 ± 0.8 |
| Glutathione | 6.0 ± 0.3 | 4.7 ± 1.0 | 6.7 ± 0.3 | 2.1 ± 0.5 |
| Glutathionylspermine | 6.4 ± 0.7 | 3.2 ± 0.8 | 2.9 ± 1.0 | 5.3 ± 0.7 |
| Glycerophosphocholine | 7.1 ± 0.7 | 7.4 ± 0.1 | 3.2 ± 0.4 | 4.4 ± 0.5 |
| Glycine | 6.6 ± 0.8 | 5.0 ± 0.2 | 6.2 ± 0.1 | 5.9 ± 0.9 |
| Glycolic acid | 6.6 ± 0.8 | 3.9 ± 0.9 | 7.1 ± 0.7 | 4.7 ± 0.7 |
| Guanidinoacetic acid | 6.3 ± 0.9 | 7.8 ± 0.6 | 5.6 ± 0.4 | 2.8 ± 0.5 |
| Guanidinobutanoic acid | 6.8 ± 1.1 | 3.6 ± 0.2 | 3.4 ± 0.9 | 6.8 ± 0.1 |
| Guanine | 11.0 ± 1.2 | 3.5 ± 1.0 | 7.6 ± 0.8 | 4.7 ± 0.9 |
| Guanosine | 10.0 ± 0.9 | 6.2 ± 1.1 | 3.2 ± 0.8 | 6.2 ± 0.4 |
| Guanosine 5'-diphosphate | 9.4 ± 0.8 | 3.4 ± 1.0 | 3.8 ± 0.1 | 6.5 ± 1.2 |
| Guanosine-5'-monophosphate | 8.8 ± 0.7 | 7.4 ± 0.4 | 7.5 ± 1.2 | 2.2 ± 1.0 |
| Hercynine | 10.2 ± 1.2 | 4.3 ± 1.1 | 7.8 ± 0.5 | 4.9 ± 1.2 |
| Hippuric acid | 10.8 ± 1.1 | 7.4 ± 0.9 | 4.3 ± 0.5 | 3.6 ± 0.1 |
| Histamine | 8.5 ± 0.8 | 5.2 ± 1.1 | 5.3 ± 0.8 | 3.2 ± 0.8 |
| Histidine | 6.8 ± 0.9 | 5.9 ± 0.3 | 6.8 ± 1.0 | 2.1 ± 0.9 |
| Homoalanine | 6.8 ± 0.6 | 2.2 ± 0.5 | 6.5 ± 0.2 | 4.3 ± 0.2 |
| Homoarginine | 6.7 ± 0.1 | 4.6 ± 0.4 | 2.7 ± 0.4 | 3.5 ± 0.5 |
| Homocysteine | 7.8 ± 0.4 | 2.8 ± 0.3 | 7.6 ± 0.9 | 4.2 ± 0.7 |
| Homogentisic acid | 10.5 ± 0.6 | 6.0 ± 0.4 | 4.9 ± 0.3 | 6.3 ± 0.4 |
| Homoleucine | 7.8 ± 0.2 | 3.0 ± 0.9 | 2.3 ± 0.7 | 7.7 ± 0.7 |
| Homovanillic acid | 8.3 ± 1.0 | 4.8 ± 0.9 | 4.5 ± 0.9 | 3.2 ± 0.9 |
| Hydantoin-5-propionate | 8.4 ± 0.7 | 2.3 ± 0.5 | 4.7 ± 0.4 | 2.4 ± 0.6 |
| Hypotaurine | 10.9 ± 0.5 | 5.4 ± 0.2 | 2.1 ± 0.7 | 7.7 ± 0.7 |
| Hypoxanthine | 8.4 ± 1.1 | 2.4 ± 1.1 | 5.7 ± 0.6 | 2.4 ± 1.0 |
| Indole-3-acetamide | 8.0 ± 0.6 | 6.0 ± 0.9 | 3.9 ± 0.6 | 5.6 ± 0.7 |
| Inosine | 10.8 ± 0.1 | 2.3 ± 0.2 | 6.7 ± 1.0 | 5.2 ± 1.0 |
| Inosine 5'-monophosphate | 7.5 ± 0.7 | 3.6 ± 0.4 | 6.4 ± 0.2 | 5.7 ± 0.5 |
| Isocitric Acid | 6.6 ± 0.5 | 6.8 ± 0.3 | 7.0 ± 0.3 | 4.5 ± 0.4 |
| Isoleucine | 8.1 ± 0.6 | 3.4 ± 1.1 | 6.2 ± 0.8 | 6.3 ± 0.3 |
| Lactic Acid | 10.8 ± 0.3 | 3.8 ± 0.4 | 7.7 ± 1.1 | 7.5 ± 0.6 |
| Leucine | 10.7 ± 0.3 | 4.3 ± 0.2 | 6.3 ± 0.5 | 5.4 ± 1.2 |
| Levulinic acid | 9.9 ± 0.3 | 4.7 ± 1.1 | 3.5 ± 0.2 | 5.9 ± 0.9 |
| Lysine | 7.5 ± 0.4 | 3.9 ± 1.1 | 7.1 ± 0.4 | 2.1 ± 0.6 |
| Malic Acid | 6.5 ± 0.9 | 6.8 ± 0.9 | 2.1 ± 0.8 | 7.2 ± 0.4 |
| Mannose-1 with Mannose-6-Phosphate | 7.4 ± 0.4 | 6.6 ± 1.1 | 5.9 ± 0.3 | 3.1 ± 0.3 |
| Mesaconic acid | 10.8 ± 1.1 | 7.2 ± 0.5 | 5.9 ± 0.5 | 7.9 ± 0.4 |
| Methionine | 9.6 ± 0.4 | 5.5 ± 1.0 | 3.1 ± 1.1 | 3.7 ± 0.7 |
| Methionine sulfoxide | 7.4 ± 0.3 | 3.0 ± 0.5 | 3.9 ± 1.1 | 7.0 ± 0.2 |
| Methylmalonic Acid | 8.0 ± 0.1 | 6.8 ± 0.8 | 2.1 ± 0.7 | 3.7 ± 1.1 |
| N1-Acetylspermidine | 6.3 ± 1.0 | 7.0 ± 0.6 | 7.5 ± 0.9 | 6.8 ± 0.2 |
| N1-Acetylspermine | 10.5 ± 0.8 | 5.2 ± 0.4 | 2.1 ± 0.6 | 2.6 ± 0.7 |
| N8-Acetylspermidine | 7.9 ± 0.9 | 5.9 ± 0.8 | 5.9 ± 0.6 | 4.5 ± 0.8 |
| N-Acetyl-Aspartyl-Glutamate | 7.3 ± 0.6 | 6.6 ± 0.2 | 5.4 ± 0.8 | 3.2 ± 0.6 |
| N-Acetyl-Aspartyl-Glutamyl-Glutamate | 6.2 ± 1.0 | 2.4 ± 0.7 | 6.7 ± 1.0 | 4.9 ± 0.6 |
| N-Acetyl-Galactosamine | 6.4 ± 0.4 | 4.1 ± 0.5 | 5.1 ± 0.2 | 7.2 ± 0.3 |
| N-Acetyl-Leucine | 7.5 ± 0.8 | 7.5 ± 0.3 | 2.8 ± 0.8 | 5.6 ± 0.6 |
| N-Acetyl-Methionine | 8.3 ± 0.4 | 6.8 ± 0.3 | 5.5 ± 1.2 | 2.5 ± 0.5 |
| N-Acetylneuraminic acid | 8.1 ± 1.0 | 4.6 ± 1.1 | 3.6 ± 0.8 | 2.1 ± 1.0 |
| N-Acetyl-Proline | 8.7 ± 0.4 | 5.7 ± 0.3 | 3.0 ± 0.4 | 5.2 ± 0.5 |
| N-Acetyl-Putrescine | 6.6 ± 1.1 | 6.9 ± 1.0 | 7.2 ± 1.0 | 3.1 ± 1.1 |
| N-Acetyl-Serine | 10.6 ± 0.4 | 7.0 ± 1.0 | 3.1 ± 0.9 | 7.9 ± 0.5 |
| N-Acetyl-Serotonin | 9.3 ± 0.5 | 2.1 ± 0.1 | 2.6 ± 1.0 | 4.2 ± 0.8 |
| N-Acyl-Homoserine-lactone | 9.3 ± 0.6 | 7.3 ± 0.8 | 7.5 ± 1.0 | 5.5 ± 0.4 |
| NAD | 6.5 ± 0.6 | 3.7 ± 1.0 | 3.7 ± 0.4 | 6.1 ± 0.4 |
| NADH | 6.5 ± 0.3 | 3.8 ± 1.0 | 2.8 ± 0.2 | 6.3 ± 0.5 |
| NADP | 10.1 ± 0.2 | 7.0 ± 0.2 | 4.8 ± 0.7 | 5.3 ± 1.0 |
| NADPH | 10.7 ± 0.7 | 6.0 ± 0.8 | 7.1 ± 0.7 | 4.3 ± 0.8 |
| N-alpha-Acetyl-lysine | 9.0 ± 0.2 | 3.7 ± 1.2 | 7.1 ± 1.1 | 3.9 ± 0.7 |
| N-alpha-Acetyl-Ornithine | 7.8 ± 0.2 | 5.4 ± 0.2 | 7.3 ± 1.1 | 4.4 ± 0.7 |
| N-Carbamoyl-Proline | 8.8 ± 0.6 | 3.6 ± 0.6 | 7.4 ± 0.8 | 5.5 ± 0.4 |
| N-Carbamoyl-Sarcosine | 6.6 ± 0.4 | 6.1 ± 0.5 | 7.1 ± 0.9 | 3.9 ± 1.0 |
| N-Epsilon-Acetyl-lysine | 9.1 ± 1.1 | 3.4 ± 0.2 | 6.5 ± 0.9 | 7.6 ± 0.3 |
| Nicotinamide | 9.9 ± 0.8 | 5.7 ± 0.4 | 4.0 ± 0.8 | 6.9 ± 0.9 |
| N-Methyl-Aspartic Acid | 6.5 ± 1.2 | 7.5 ± 0.9 | 2.9 ± 0.8 | 3.4 ± 0.6 |
| O-Phosphoethanolamine | 6.7 ± 0.8 | 6.8 ± 1.2 | 5.4 ± 0.6 | 2.2 ± 1.0 |
| Ophthalmic acid | 9.3 ± 0.9 | 2.1 ± 0.8 | 3.2 ± 0.3 | 5.6 ± 0.6 |
| Ornithine | 10.3 ± 0.3 | 7.9 ± 0.9 | 6.1 ± 0.8 | 2.0 ± 1.1 |
| Orotidine 5'-monophosphate | 6.9 ± 0.3 | 5.4 ± 0.1 | 5.3 ± 0.7 | 3.5 ± 0.2 |
| Oxaloacetate | 9.7 ± 0.1 | 5.9 ± 0.8 | 4.0 ± 0.5 | 3.5 ± 1.2 |
| Pantothenic acid | 10.5 ± 0.6 | 4.0 ± 0.6 | 5.6 ± 0.8 | 7.4 ± 0.7 |
| Phenylalanine | 9.8 ± 1.2 | 7.2 ± 1.0 | 3.7 ± 0.5 | 4.0 ± 1.2 |
| Phenylpyruvate | 7.7 ± 0.6 | 7.7 ± 0.5 | 6.4 ± 0.3 | 7.9 ± 1.0 |
| Phosphodimethylethanolamine | 9.6 ± 0.7 | 5.5 ± 0.2 | 5.5 ± 0.2 | 4.2 ± 0.2 |
| Phosphoenolpyruvate | 9.7 ± 1.1 | 2.6 ± 1.0 | 5.6 ± 1.0 | 3.7 ± 0.9 |
| Phosphorylcholine | 10.8 ± 0.5 | 7.9 ± 0.7 | 7.6 ± 0.2 | 2.5 ± 0.6 |
| Pipecolic acid | 9.4 ± 0.2 | 7.6 ± 1.1 | 7.4 ± 0.3 | 7.4 ± 0.2 |
| Proline | 10.2 ± 0.2 | 6.2 ± 0.4 | 4.1 ± 0.8 | 5.2 ± 0.5 |
| Putrescine | 9.7 ± 0.5 | 6.3 ± 0.7 | 3.0 ± 0.6 | 8.0 ± 0.6 |
| Pyridoxal | 10.6 ± 0.2 | 4.9 ± 1.0 | 4.8 ± 1.0 | 3.2 ± 0.7 |
| Pyridoxamine | 10.7 ± 0.4 | 2.8 ± 1.0 | 7.2 ± 0.4 | 2.8 ± 0.7 |
| Pyridoxine | 8.4 ± 0.2 | 7.1 ± 0.5 | 2.9 ± 1.1 | 4.5 ± 0.7 |
| Pyroglutamic acid | 7.6 ± 1.0 | 3.4 ± 0.7 | 7.1 ± 1.0 | 8.0 ± 0.9 |
| Pyrrolysine | 8.7 ± 0.4 | 7.6 ± 0.9 | 5.7 ± 1.2 | 4.4 ± 0.8 |
| Pyruvic Acid | 8.7 ± 0.4 | 3.7 ± 0.2 | 7.4 ± 1.1 | 6.0 ± 1.0 |
| Riboflavin | 8.5 ± 1.0 | 7.6 ± 1.1 | 2.1 ± 0.3 | 4.6 ± 1.1 |
| Ribose-5-Phosphate | 9.3 ± 0.6 | 6.6 ± 0.4 | 7.3 ± 1.0 | 7.2 ± 0.8 |
| Ribulose-1-Phosphate | 10.6 ± 1.0 | 3.7 ± 0.1 | 7.3 ± 0.8 | 3.7 ± 1.1 |
| Ribulose-5-Phosphate | 6.5 ± 1.0 | 2.1 ± 1.1 | 6.1 ± 0.4 | 2.0 ± 1.0 |
| Saccharopine | 6.9 ± 0.4 | 5.6 ± 0.8 | 7.3 ± 0.4 | 3.0 ± 0.6 |
| S-Adenosylhomocysteine | 8.3 ± 0.4 | 3.4 ± 0.2 | 4.7 ± 0.5 | 4.7 ± 0.6 |
| S-Adenosylmethioninamine | 9.8 ± 0.9 | 5.5 ± 0.4 | 5.5 ± 0.2 | 3.0 ± 0.6 |
| S-Adenosylmethionine | 6.5 ± 0.4 | 5.4 ± 0.4 | 4.5 ± 0.4 | 3.7 ± 0.1 |
| Sarcosine | 7.3 ± 1.0 | 8.0 ± 0.8 | 2.2 ± 0.2 | 5.2 ± 0.7 |
| Sedoheptulose-7-phosphate | 8.2 ± 0.9 | 6.8 ± 1.1 | 4.9 ± 1.1 | 2.6 ± 1.1 |
| Selenohomocysteine | 8.3 ± 0.8 | 3.9 ± 0.2 | 7.4 ± 0.6 | 2.6 ± 0.8 |
| Selenomethionine | 8.9 ± 0.9 | 7.3 ± 1.0 | 3.0 ± 1.1 | 2.8 ± 0.4 |
| Serine | 6.5 ± 0.5 | 5.0 ± 0.4 | 6.4 ± 1.0 | 2.0 ± 0.2 |
| S-Methyl-Thioinosine | 7.6 ± 0.2 | 6.4 ± 1.2 | 4.4 ± 1.1 | 7.0 ± 0.7 |
| Spermidine | 6.4 ± 0.7 | 3.0 ± 1.1 | 2.1 ± 0.5 | 2.9 ± 0.4 |
| Spermine | 9.6 ± 0.5 | 6.3 ± 1.0 | 3.2 ± 1.1 | 2.3 ± 0.5 |
| Succinate | 8.7 ± 0.4 | 2.7 ± 0.4 | 4.7 ± 0.5 | 4.5 ± 0.2 |
| Sucrose | 9.8 ± 0.3 | 2.3 ± 0.7 | 3.7 ± 0.4 | 6.8 ± 1.0 |
| Symmetric dimethylarginine | 6.1 ± 1.1 | 5.7 ± 0.2 | 4.5 ± 0.1 | 2.3 ± 0.4 |
| Taurine | 9.5 ± 0.8 | 6.5 ± 1.0 | 6.6 ± 0.8 | 7.7 ± 1.1 |
| Tetrahydroneopterin | 10.7 ± 0.3 | 2.2 ± 0.3 | 2.7 ± 0.8 | 4.4 ± 0.5 |
| Thiamine | 10.1 ± 0.6 | 7.9 ± 1.1 | 5.5 ± 0.9 | 7.1 ± 0.1 |
| Thiamine monophosphate | 6.6 ± 0.6 | 3.5 ± 1.0 | 6.1 ± 1.0 | 6.4 ± 1.0 |
| Threonic acid | 7.3 ± 0.3 | 6.8 ± 0.7 | 7.6 ± 0.7 | 2.6 ± 0.3 |
| Threonine | 6.5 ± 0.7 | 5.7 ± 1.2 | 3.2 ± 0.6 | 2.7 ± 0.7 |
| Thymidine | 8.7 ± 0.4 | 3.3 ± 0.6 | 4.6 ± 0.2 | 4.0 ± 0.3 |
| Thymidine Monophosphate | 10.0 ± 0.8 | 2.8 ± 1.0 | 3.9 ± 0.2 | 3.7 ± 0.7 |
| Thymine | 8.1 ± 0.2 | 5.8 ± 1.0 | 6.9 ± 0.3 | 4.9 ± 0.3 |
| Trigonelleine | 7.0 ± 1.2 | 4.7 ± 0.7 | 2.4 ± 0.3 | 7.4 ± 1.1 |
| Trimethylamine N-oxide | 10.4 ± 1.0 | 5.6 ± 0.5 | 5.6 ± 0.4 | 6.1 ± 0.9 |
| Trimethyllysine | 9.7 ± 0.2 | 2.5 ± 0.8 | 7.9 ± 0.7 | 5.7 ± 0.9 |
| Tryptophan | 6.1 ± 0.3 | 3.2 ± 1.1 | 2.6 ± 0.9 | 3.3 ± 0.2 |
| Tyramine | 9.7 ± 1.0 | 3.0 ± 0.6 | 2.2 ± 1.0 | 6.0 ± 0.6 |
| Tyrosine | 8.0 ± 0.8 | 5.2 ± 0.6 | 4.3 ± 0.7 | 7.7 ± 0.9 |
| Uracil | 8.7 ± 0.8 | 7.6 ± 0.2 | 2.3 ± 0.4 | 3.3 ± 1.1 |
| Uridine 5'-diphosphate | 6.5 ± 0.8 | 4.5 ± 0.8 | 3.5 ± 1.0 | 4.6 ± 0.8 |
| Uridine Monophosphate | 7.5 ± 0.9 | 4.7 ± 0.6 | 7.1 ± 0.8 | 6.9 ± 0.3 |
| Uridine-5-Diphosphogalactose | 7.5 ± 0.9 | 3.1 ± 0.1 | 6.8 ± 0.7 | 3.3 ± 0.4 |
| Urocanic acid | 10.5 ± 0.4 | 4.7 ± 0.8 | 3.2 ± 0.3 | 4.2 ± 1.0 |
| UTP | 7.9 ± 0.4 | 3.6 ± 0.3 | 5.7 ± 0.8 | 2.9 ± 0.6 |
| Valine | 6.4 ± 0.6 | 2.2 ± 0.9 | 3.4 ± 0.2 | 7.9 ± 0.9 |
| XMP | 6.1 ± 0.6 | 4.3 ± 0.4 | 6.4 ± 0.2 | 6.6 ± 0.7 |
| Xylulose-5-Phosphate | 6.7 ± 0.3 | 2.7 ± 0.8 | 2.8 ± 0.4 | 4.2 ± 0.7 |

### **Table S19** | Inter – day precision (as % RSD) (Plasma extract as matrix, n = 5). The concentrations of QC_LLOQ_, QC_Low_, QC_Mid_ and QC_High_ were 10 ng/mL, 25 ng/mL, 100 ng/mL and 500 ng/mL respectively.

| Metabolite | QC_LLOQ_ | QC_Low_ | QC_Mid_ | QC_High_ |
| --- | --- | --- | --- | --- |
| 1-Methylguanine | 6.0 ± 0.7 | 4.2 ± 1.0 | 2.2 ± 0.3 | 3.3 ± 0.7 |
| 1-Methylhistidine | 9.2 ± 0.7 | 4.8 ± 0.6 | 5.6 ± 1.0 | 3.2 ± 0.5 |
| 1-Methylnicotinamide | 8.6 ± 0.5 | 5.3 ± 0.8 | 4.4 ± 0.6 | 4.9 ± 0.1 |
| 2-Aminoadipic Acid | 9.4 ± 0.4 | 5.3 ± 0.7 | 4.3 ± 0.3 | 2.3 ± 0.5 |
| 2-Deoxyadenosine | 9.4 ± 0.7 | 3.5 ± 0.8 | 4.3 ± 0.2 | 2.5 ± 1.1 |
| 2'-Deoxyadenosine 5'-monophosphate | 7.3 ± 0.5 | 4.8 ± 1.0 | 4.2 ± 0.5 | 4.9 ± 1.2 |
| 2'-deoxyadenosine 5'-triphosphate | 6.9 ± 1.1 | 4.9 ± 0.3 | 2.9 ± 0.8 | 3.3 ± 0.4 |
| 2'-Deoxycytidine-5'-monophosphate | 5.6 ± 1.1 | 4.9 ± 0.4 | 4.7 ± 0.5 | 5.7 ± 0.3 |
| 2-Deoxyguanosine | 8.9 ± 0.6 | 2.7 ± 0.5 | 2.6 ± 0.5 | 3.4 ± 0.5 |
| 2-hydroxybutyric acid | 6.2 ± 0.9 | 2.6 ± 1.2 | 5.9 ± 0.9 | 3.1 ± 0.2 |
| 2-hydroxyglutaric acid | 7.3 ± 0.8 | 3.0 ± 1.1 | 4.5 ± 0.7 | 3.5 ± 1.2 |
| 2-Hydroxyisocaproic acid | 8.2 ± 0.5 | 2.3 ± 0.2 | 3.8 ± 0.9 | 4.8 ± 0.9 |
| 2-ketobutyric acid | 7.2 ± 0.9 | 5.2 ± 0.2 | 3.5 ± 0.7 | 3.0 ± 1.1 |
| 2-oxoadipic acid | 5.9 ± 0.6 | 4.8 ± 0.6 | 5.1 ± 0.8 | 3.1 ± 0.7 |
| 2-Phenylglycine | 8.5 ± 0.6 | 4.3 ± 0.9 | 3.3 ± 0.4 | 4.8 ± 0.1 |
| 2-phosphoglycerate | 8.1 ± 0.9 | 3.8 ± 0.3 | 2.8 ± 0.3 | 2.6 ± 1.1 |
| 3-hydroxy-3-methyl-2-oxobutanoate | 9.2 ± 0.4 | 2.8 ± 0.7 | 2.7 ± 0.6 | 3.0 ± 0.5 |
| 3-hydroxybutyric acid | 5.1 ± 0.8 | 2.0 ± 0.8 | 2.8 ± 0.9 | 5.8 ± 0.6 |
| 3-Methylhistidine | 8.6 ± 0.6 | 4.5 ± 1.1 | 3.1 ± 0.2 | 2.7 ± 0.9 |
| 3-Phosphoglyceraldehyde | 9.5 ± 0.6 | 4.2 ± 1.0 | 3.3 ± 0.9 | 3.6 ± 0.7 |
| 3-phosphoglycerate | 8.5 ± 1.0 | 4.9 ± 0.6 | 4.7 ± 1.0 | 5.3 ± 0.8 |
| 4-amino-butyric acid | 7.9 ± 0.5 | 2.5 ± 0.6 | 3.5 ± 1.1 | 3.2 ± 0.8 |
| 4-aspartyl-phosphate | 8.3 ± 1.1 | 3.3 ± 0.1 | 4.2 ± 0.9 | 3.0 ± 0.9 |
| 4-Hydroxybenzoic acid | 8.8 ± 0.3 | 4.9 ± 0.4 | 5.8 ± 0.7 | 3.9 ± 0.3 |
| 4-Hydroxyphenylacetylglycine | 8.5 ± 0.9 | 3.4 ± 0.6 | 2.8 ± 0.7 | 3.2 ± 0.9 |
| 4-methyl-2-oxopentanoate | 6.4 ± 0.6 | 2.3 ± 1.0 | 5.6 ± 1.0 | 4.5 ± 1.2 |
| 4-Oxoglutaramate | 7.8 ± 0.3 | 3.4 ± 0.8 | 3.1 ± 0.1 | 2.8 ± 0.4 |
| 5-Hydroxyindolepyruvate | 5.3 ± 1.0 | 2.2 ± 0.6 | 4.3 ± 1.0 | 4.0 ± 0.4 |
| 5-Hydroxylysine | 6.0 ± 0.7 | 2.5 ± 0.2 | 3.2 ± 1.0 | 5.2 ± 0.3 |
| 5-Hydroxy-Tryptophan | 5.3 ± 0.7 | 2.5 ± 0.1 | 3.6 ± 0.2 | 4.9 ± 0.3 |
| 5-Phosphoribosylamine | 9.8 ± 1.0 | 4.3 ± 0.9 | 4.1 ± 1.1 | 5.6 ± 0.5 |
| 6-phosphogluconate | 6.6 ± 0.7 | 5.3 ± 0.3 | 4.0 ± 0.2 | 3.6 ± 0.8 |
| Acetylcarnitine | 10.2 ± 0.3 | 5.0 ± 0.8 | 5.7 ± 0.2 | 5.7 ± 0.2 |
| Aconitic Acid | 9.1 ± 0.3 | 5.2 ± 0.2 | 5.0 ± 0.6 | 4.0 ± 0.5 |
| Adenine | 5.3 ± 0.2 | 4.6 ± 0.4 | 4.8 ± 0.6 | 5.1 ± 0.7 |
| Adenosine | 10.9 ± 0.7 | 2.3 ± 0.9 | 5.4 ± 1.0 | 5.1 ± 0.8 |
| Adenosine 3',5'-cyclic monophosphate | 6.0 ± 0.6 | 2.1 ± 0.2 | 4.1 ± 0.4 | 5.9 ± 0.7 |
| Adenosine Monophosphate | 9.6 ± 1.1 | 2.2 ± 0.6 | 5.2 ± 0.9 | 5.0 ± 0.5 |
| Adenosine-5'-diphosphate | 5.6 ± 0.2 | 4.1 ± 0.6 | 5.9 ± 1.1 | 2.5 ± 0.5 |
| Adenosine-5'-triphosphate | 7.2 ± 0.6 | 5.2 ± 0.9 | 4.6 ± 0.8 | 3.7 ± 0.6 |
| Adenylosuccinate | 5.8 ± 1.0 | 5.2 ± 0.3 | 4.5 ± 0.2 | 5.6 ± 0.2 |
| Alanine | 10.2 ± 0.7 | 3.6 ± 1.1 | 4.5 ± 0.2 | 2.2 ± 1.0 |
| alpha-ketoglutaric acid | 11.0 ± 0.1 | 4.7 ± 0.7 | 2.9 ± 0.8 | 4.3 ± 0.1 |
| Anserine | 10.8 ± 0.5 | 5.5 ± 1.1 | 3.4 ± 0.8 | 4.4 ± 0.3 |
| Arginine | 10.8 ± 1.1 | 3.1 ± 0.1 | 4.8 ± 1.0 | 4.8 ± 0.2 |
| Argininosuccinic Acid | 9.7 ± 0.5 | 2.5 ± 0.2 | 5.7 ± 0.6 | 5.3 ± 1.1 |
| Asparagine | 10.9 ± 0.6 | 4.5 ± 0.5 | 3.7 ± 0.6 | 5.8 ± 0.5 |
| Aspartic acid | 5.3 ± 0.2 | 5.9 ± 0.6 | 2.6 ± 0.7 | 5.5 ± 0.8 |
| Asymmetric dimethylarginine | 10.6 ± 0.2 | 5.7 ± 1.0 | 2.9 ± 0.9 | 2.9 ± 1.2 |
| Azelaic acid | 6.1 ± 0.9 | 2.8 ± 0.8 | 2.7 ± 0.6 | 2.5 ± 0.2 |
| beta-Alanine | 5.1 ± 0.9 | 3.2 ± 0.6 | 4.3 ± 0.3 | 5.4 ± 1.0 |
| Betaine | 8.7 ± 0.4 | 5.3 ± 1.0 | 3.7 ± 0.7 | 4.7 ± 0.4 |
| Biopterin | 8.0 ± 0.5 | 3.5 ± 0.3 | 4.5 ± 0.9 | 4.7 ± 0.6 |
| Biotin | 7.5 ± 0.5 | 5.3 ± 1.2 | 5.1 ± 0.9 | 3.5 ± 1.1 |
| Carnitine | 7.5 ± 0.8 | 2.6 ± 0.1 | 3.4 ± 0.3 | 4.3 ± 0.2 |
| Carnosine | 6.5 ± 0.1 | 5.0 ± 0.5 | 5.0 ± 0.3 | 3.2 ± 0.9 |
| CDP-choline | 6.1 ± 0.6 | 3.5 ± 0.4 | 3.9 ± 0.6 | 2.8 ± 0.3 |
| cGMP | 7.7 ± 1.0 | 4.8 ± 0.6 | 3.5 ± 0.7 | 4.0 ± 0.2 |
| Choline | 10.2 ± 0.9 | 2.4 ± 0.5 | 3.8 ± 0.5 | 2.5 ± 1.2 |
| Cinnamic acid | 8.4 ± 0.3 | 4.2 ± 0.9 | 2.8 ± 1.0 | 5.2 ± 1.1 |
| Citraconic acid | 6.9 ± 1.0 | 4.0 ± 0.9 | 3.3 ± 1.0 | 3.2 ± 0.4 |
| Citramalic acid | 7.4 ± 0.9 | 4.0 ± 0.1 | 4.2 ± 0.8 | 4.9 ± 0.3 |
| Citric Acid | 5.7 ± 0.1 | 5.6 ± 0.8 | 2.5 ± 1.2 | 6.0 ± 0.8 |
| Citrulline | 10.8 ± 1.1 | 3.7 ± 0.8 | 4.3 ± 0.4 | 4.2 ± 0.8 |
| Creatine | 9.2 ± 0.5 | 2.4 ± 0.7 | 4.5 ± 1.2 | 4.0 ± 0.4 |
| Creatinine | 5.4 ± 0.5 | 5.3 ± 0.8 | 4.8 ± 0.1 | 4.2 ± 0.2 |
| Cystathionine | 9.9 ± 0.8 | 3.7 ± 0.6 | 4.1 ± 1.0 | 4.7 ± 1.0 |
| Cysteamine | 9.3 ± 0.5 | 5.5 ± 0.3 | 3.0 ± 0.9 | 5.6 ± 0.5 |
| Cysteine | 10.6 ± 0.7 | 5.5 ± 0.9 | 2.6 ± 0.9 | 2.7 ± 1.1 |
| Cysteinylglycine | 8.5 ± 0.6 | 5.0 ± 0.2 | 2.7 ± 0.8 | 3.3 ± 0.2 |
| Cystine | 8.4 ± 0.3 | 3.4 ± 0.6 | 5.9 ± 0.4 | 4.3 ± 0.1 |
| Cytidine | 10.2 ± 0.7 | 3.4 ± 0.6 | 2.3 ± 1.2 | 5.8 ± 0.7 |
| Cytidine 3',5'-cyclic monophosphate | 8.8 ± 1.1 | 3.0 ± 1.0 | 4.9 ± 0.4 | 5.5 ± 0.4 |
| Cytidine 5'-diphosphate | 8.2 ± 0.4 | 5.7 ± 0.7 | 3.5 ± 0.9 | 2.8 ± 0.7 |
| Cytidine 5'-triphosphate | 6.3 ± 1.2 | 3.8 ± 0.8 | 2.4 ± 1.0 | 5.4 ± 0.2 |
| Cytidine monophosphate | 7.5 ± 0.5 | 3.9 ± 0.5 | 3.5 ± 0.5 | 5.4 ± 0.4 |
| Cytosine | 9.6 ± 0.9 | 3.0 ± 1.0 | 4.6 ± 0.5 | 5.9 ± 0.7 |
| dADP | 8.1 ± 0.6 | 3.6 ± 0.4 | 5.6 ± 0.8 | 4.6 ± 0.2 |
| Diethanolamine | 10.9 ± 0.7 | 2.3 ± 0.2 | 6.0 ± 0.6 | 4.0 ± 0.8 |
| Dihydrobiopterin | 9.3 ± 0.6 | 4.6 ± 0.8 | 5.9 ± 0.9 | 3.0 ± 0.1 |
| Dihydroneopterin triphosphate | 9.9 ± 0.3 | 3.1 ± 0.3 | 3.9 ± 1.0 | 2.2 ± 0.4 |
| Dihydroxyacetonephosphate | 5.1 ± 0.7 | 5.4 ± 0.6 | 3.5 ± 1.0 | 4.8 ± 0.6 |
| Dimethylglycine | 5.8 ± 0.8 | 3.8 ± 0.3 | 5.1 ± 0.4 | 2.9 ± 0.5 |
| Ectoine | 7.5 ± 0.4 | 3.5 ± 0.2 | 2.1 ± 0.3 | 4.6 ± 0.8 |
| Erythrose-4-phosphate | 5.7 ± 0.5 | 3.3 ± 0.6 | 5.8 ± 0.3 | 2.0 ± 0.4 |
| FAD | 8.2 ± 0.3 | 5.3 ± 1.2 | 4.5 ± 1.1 | 2.6 ± 1.0 |
| Folic acid | 9.0 ± 0.7 | 3.9 ± 0.4 | 5.8 ± 1.0 | 3.3 ± 0.6 |
| Fructose-6-phosphate | 10.9 ± 0.3 | 3.9 ± 0.4 | 3.7 ± 0.2 | 5.2 ± 0.2 |
| Fumaric Acid | 9.6 ± 0.8 | 4.8 ± 1.1 | 3.2 ± 1.0 | 2.1 ± 0.8 |
| Galactosamine | 9.1 ± 0.6 | 3.0 ± 0.9 | 2.4 ± 0.6 | 3.2 ± 0.7 |
| Galactose-1-phosphate | 9.6 ± 1.2 | 3.9 ± 0.1 | 2.1 ± 0.6 | 2.4 ± 0.8 |
| Galactose-6-Phosphate | 6.9 ± 1.1 | 4.7 ± 0.7 | 3.1 ± 0.7 | 2.4 ± 0.7 |
| gamma-Glutamylcysteine | 5.3 ± 0.9 | 4.3 ± 0.2 | 3.1 ± 1.2 | 4.6 ± 0.6 |
| Glucosamine | 6.2 ± 0.7 | 3.1 ± 0.8 | 5.4 ± 1.0 | 5.2 ± 0.8 |
| Glucosamine-6-Phosphate | 7.5 ± 0.1 | 4.4 ± 0.2 | 3.4 ± 0.4 | 6.0 ± 0.5 |
| Glucose | 7.1 ± 0.9 | 3.2 ± 1.2 | 2.4 ± 0.5 | 5.1 ± 0.4 |
| Glucose-1-phosphate | 10.0 ± 0.8 | 3.4 ± 0.8 | 3.5 ± 1.2 | 4.3 ± 1.1 |
| Glucose-6-Phosphate | 10.8 ± 0.6 | 5.0 ± 0.5 | 5.5 ± 0.8 | 5.5 ± 0.5 |
| Glucuronic acid | 7.8 ± 0.2 | 2.7 ± 0.8 | 3.0 ± 0.8 | 4.2 ± 0.6 |
| Glutamic acid | 9.1 ± 0.1 | 2.2 ± 1.0 | 2.6 ± 0.7 | 2.6 ± 0.5 |
| Glutamine | 5.6 ± 0.8 | 5.3 ± 1.2 | 4.8 ± 0.6 | 2.2 ± 0.7 |
| Glutathione | 9.5 ± 0.7 | 3.9 ± 0.5 | 3.1 ± 1.1 | 5.9 ± 0.2 |
| Glutathionylspermine | 7.7 ± 0.6 | 2.1 ± 0.7 | 4.0 ± 0.5 | 3.2 ± 1.1 |
| Glycerophosphocholine | 10.2 ± 0.6 | 5.7 ± 0.1 | 4.5 ± 0.8 | 3.3 ± 0.5 |
| Glycine | 5.9 ± 1.0 | 4.7 ± 1.1 | 3.8 ± 0.5 | 4.5 ± 0.6 |
| Glycolic acid | 8.8 ± 0.6 | 5.6 ± 0.3 | 3.6 ± 0.7 | 2.1 ± 0.9 |
| Guanidinoacetic acid | 6.2 ± 0.6 | 2.4 ± 0.5 | 2.2 ± 0.4 | 4.2 ± 1.0 |
| Guanidinobutanoic acid | 6.3 ± 0.3 | 4.7 ± 0.5 | 2.1 ± 1.1 | 4.6 ± 0.7 |
| Guanine | 10.4 ± 0.3 | 2.0 ± 0.5 | 2.1 ± 0.3 | 3.9 ± 1.0 |
| Guanosine | 8.4 ± 0.7 | 2.2 ± 0.6 | 2.0 ± 0.5 | 4.5 ± 0.5 |
| Guanosine 5'-diphosphate | 7.2 ± 1.0 | 6.0 ± 0.3 | 2.1 ± 0.4 | 4.3 ± 1.0 |
| Guanosine-5'-monophosphate | 5.8 ± 1.1 | 2.3 ± 1.1 | 2.5 ± 0.7 | 3.8 ± 1.1 |
| Hercynine | 8.5 ± 1.1 | 3.3 ± 1.0 | 4.2 ± 0.6 | 5.4 ± 0.8 |
| Hippuric acid | 7.7 ± 0.8 | 4.1 ± 0.6 | 2.5 ± 1.0 | 4.2 ± 0.2 |
| Histamine | 10.3 ± 0.9 | 3.4 ± 0.6 | 3.1 ± 1.1 | 3.0 ± 1.0 |
| Histidine | 8.9 ± 0.7 | 3.6 ± 1.1 | 5.5 ± 1.0 | 2.2 ± 0.7 |
| Homoalanine | 10.2 ± 0.6 | 2.1 ± 0.8 | 3.6 ± 0.5 | 3.6 ± 0.6 |
| Homoarginine | 7.9 ± 1.1 | 3.6 ± 0.5 | 3.1 ± 1.1 | 2.4 ± 1.1 |
| Homocysteine | 8.9 ± 0.5 | 5.7 ± 0.5 | 2.8 ± 0.4 | 5.4 ± 1.2 |
| Homogentisic acid | 7.5 ± 0.9 | 2.6 ± 0.2 | 5.1 ± 0.8 | 3.9 ± 1.0 |
| Homoleucine | 7.7 ± 0.6 | 5.0 ± 0.5 | 4.0 ± 0.1 | 2.4 ± 1.0 |
| Homovanillic acid | 5.8 ± 0.7 | 3.7 ± 0.2 | 4.7 ± 0.8 | 2.2 ± 1.2 |
| Hydantoin-5-propionate | 10.7 ± 1.1 | 3.5 ± 1.1 | 5.1 ± 0.3 | 2.2 ± 1.1 |
| Hypotaurine | 9.8 ± 1.1 | 2.7 ± 0.7 | 2.8 ± 0.4 | 3.1 ± 0.9 |
| Hypoxanthine | 11.0 ± 1.0 | 2.0 ± 0.6 | 4.1 ± 0.4 | 2.9 ± 0.8 |
| Indole-3-acetamide | 8.8 ± 1.0 | 4.0 ± 0.5 | 5.2 ± 0.9 | 5.7 ± 0.6 |
| Inosine | 5.9 ± 0.7 | 3.6 ± 0.2 | 2.3 ± 0.4 | 2.1 ± 0.2 |
| Inosine 5'-monophosphate | 6.3 ± 0.1 | 3.7 ± 1.0 | 2.5 ± 1.2 | 5.3 ± 0.5 |
| Isocitric Acid | 6.4 ± 0.6 | 2.7 ± 0.5 | 4.0 ± 1.0 | 2.4 ± 0.4 |
| Isoleucine | 9.5 ± 0.6 | 5.5 ± 0.9 | 4.2 ± 0.2 | 3.6 ± 0.9 |
| Lactic Acid | 9.5 ± 1.2 | 4.7 ± 0.8 | 5.3 ± 0.7 | 3.3 ± 0.3 |
| Leucine | 5.6 ± 0.5 | 2.5 ± 0.5 | 5.6 ± 1.1 | 3.1 ± 0.1 |
| Levulinic acid | 8.9 ± 0.7 | 4.7 ± 0.3 | 4.1 ± 1.2 | 4.2 ± 0.7 |
| Lysine | 6.4 ± 0.4 | 4.2 ± 1.2 | 4.4 ± 0.6 | 5.7 ± 0.4 |
| Malic Acid | 10.0 ± 1.2 | 3.3 ± 0.4 | 4.5 ± 1.0 | 5.5 ± 1.0 |
| Mannose-1 with Mannose-6-Phosphate | 6.7 ± 1.2 | 4.4 ± 0.3 | 3.7 ± 0.3 | 3.9 ± 1.2 |
| Mesaconic acid | 8.0 ± 0.4 | 3.0 ± 1.2 | 2.9 ± 0.3 | 3.5 ± 0.2 |
| Methionine | 7.1 ± 0.4 | 2.3 ± 0.2 | 3.4 ± 0.2 | 5.4 ± 0.9 |
| Methionine sulfoxide | 10.7 ± 0.6 | 3.7 ± 0.4 | 5.3 ± 0.2 | 4.3 ± 0.5 |
| Methylmalonic Acid | 6.1 ± 0.2 | 2.0 ± 0.7 | 2.3 ± 0.7 | 4.0 ± 0.7 |
| N1-Acetylspermidine | 5.6 ± 0.7 | 2.7 ± 1.0 | 4.7 ± 0.8 | 4.1 ± 1.1 |
| N1-Acetylspermine | 5.1 ± 0.3 | 4.2 ± 0.9 | 3.9 ± 0.4 | 3.0 ± 0.7 |
| N8-Acetylspermidine | 5.3 ± 0.1 | 4.6 ± 0.8 | 5.0 ± 0.2 | 4.4 ± 0.3 |
| N-Acetyl-Aspartyl-Glutamate | 6.0 ± 0.2 | 5.6 ± 1.0 | 3.9 ± 0.6 | 2.8 ± 1.1 |
| N-Acetyl-Aspartyl-Glutamyl-Glutamate | 10.0 ± 0.6 | 2.1 ± 0.9 | 4.2 ± 0.6 | 4.3 ± 0.9 |
| N-Acetyl-Galactosamine | 8.7 ± 0.1 | 4.9 ± 0.6 | 3.8 ± 1.1 | 2.3 ± 0.2 |
| N-Acetyl-Leucine | 5.9 ± 0.7 | 4.0 ± 0.3 | 4.9 ± 0.6 | 2.5 ± 0.5 |
| N-Acetyl-Methionine | 8.8 ± 0.3 | 5.2 ± 1.2 | 3.1 ± 0.2 | 4.0 ± 1.0 |
| N-Acetylneuraminic acid | 9.5 ± 0.9 | 2.1 ± 0.3 | 4.5 ± 0.6 | 4.6 ± 0.4 |
| N-Acetyl-Proline | 10.2 ± 0.7 | 3.9 ± 1.2 | 4.4 ± 0.1 | 4.6 ± 0.8 |
| N-Acetyl-Putrescine | 7.0 ± 0.2 | 5.5 ± 0.5 | 4.1 ± 0.8 | 4.7 ± 0.5 |
| N-Acetyl-Serine | 5.5 ± 0.6 | 5.3 ± 0.4 | 4.4 ± 0.3 | 4.4 ± 0.1 |
| N-Acetyl-Serotonin | 6.6 ± 1.0 | 4.9 ± 0.1 | 5.4 ± 0.3 | 2.4 ± 0.7 |
| N-Acyl-Homoserine-lactone | 5.8 ± 0.9 | 3.8 ± 0.7 | 4.4 ± 0.2 | 4.9 ± 0.5 |
| NAD | 6.4 ± 0.8 | 5.2 ± 0.5 | 3.5 ± 1.2 | 3.0 ± 0.7 |
| NADH | 9.2 ± 0.3 | 4.7 ± 0.6 | 4.7 ± 1.0 | 3.8 ± 1.1 |
| NADP | 10.0 ± 1.0 | 2.9 ± 0.3 | 5.6 ± 1.1 | 5.8 ± 0.1 |
| NADPH | 9.0 ± 1.2 | 2.0 ± 0.1 | 3.9 ± 0.8 | 2.3 ± 0.7 |
| N-alpha-Acetyl-lysine | 8.3 ± 1.1 | 2.8 ± 0.7 | 5.4 ± 0.5 | 5.6 ± 0.5 |
| N-alpha-Acetyl-Ornithine | 5.5 ± 0.3 | 2.9 ± 0.1 | 5.4 ± 0.8 | 2.1 ± 0.4 |
| N-Carbamoyl-Proline | 5.5 ± 0.1 | 2.8 ± 1.2 | 3.4 ± 0.8 | 3.8 ± 0.7 |
| N-Carbamoyl-Sarcosine | 6.4 ± 0.4 | 2.9 ± 1.0 | 5.2 ± 0.8 | 4.1 ± 0.4 |
| N-Epsilon-Acetyl-lysine | 9.1 ± 0.4 | 4.7 ± 0.5 | 2.3 ± 0.7 | 4.8 ± 0.7 |
| Nicotinamide | 6.1 ± 0.2 | 3.0 ± 1.1 | 3.7 ± 0.2 | 2.0 ± 0.4 |
| N-Methyl-Aspartic Acid | 7.4 ± 0.3 | 4.4 ± 0.3 | 4.4 ± 0.6 | 2.0 ± 0.3 |
| O-Phosphoethanolamine | 8.4 ± 0.5 | 5.6 ± 0.7 | 4.4 ± 1.0 | 5.3 ± 0.5 |
| Ophthalmic acid | 8.1 ± 0.3 | 2.4 ± 1.0 | 4.5 ± 0.7 | 4.4 ± 0.5 |
| Ornithine | 5.9 ± 0.8 | 3.6 ± 0.8 | 5.3 ± 0.2 | 2.9 ± 0.5 |
| Orotidine 5'-monophosphate | 10.4 ± 0.7 | 4.6 ± 0.9 | 4.1 ± 1.2 | 4.2 ± 1.0 |
| Oxaloacetate | 7.2 ± 0.5 | 3.1 ± 0.9 | 4.2 ± 0.3 | 3.0 ± 1.0 |
| Pantothenic acid | 6.5 ± 0.8 | 2.5 ± 0.4 | 5.3 ± 0.7 | 5.9 ± 1.0 |
| Phenylalanine | 5.8 ± 0.2 | 2.9 ± 0.6 | 5.6 ± 0.8 | 5.9 ± 1.1 |
| Phenylpyruvate | 10.5 ± 0.5 | 5.4 ± 0.4 | 3.6 ± 1.1 | 3.8 ± 0.2 |
| Phosphodimethylethanolamine | 10.5 ± 0.1 | 2.9 ± 0.8 | 4.5 ± 0.3 | 2.5 ± 0.5 |
| Phosphoenolpyruvate | 8.7 ± 0.9 | 3.5 ± 0.3 | 5.5 ± 0.7 | 4.9 ± 0.6 |
| Phosphorylcholine | 6.2 ± 0.7 | 3.6 ± 0.1 | 5.5 ± 1.1 | 4.5 ± 1.0 |
| Pipecolic acid | 10.2 ± 0.2 | 5.2 ± 0.7 | 3.2 ± 0.2 | 5.6 ± 0.3 |
| Proline | 8.9 ± 0.7 | 4.9 ± 0.6 | 2.2 ± 0.2 | 5.5 ± 0.5 |
| Putrescine | 8.3 ± 0.5 | 4.0 ± 0.6 | 5.9 ± 1.1 | 3.0 ± 0.4 |
| Pyridoxal | 6.5 ± 0.5 | 4.2 ± 0.9 | 3.3 ± 0.9 | 3.7 ± 0.6 |
| Pyridoxamine | 10.4 ± 0.5 | 5.4 ± 1.1 | 4.0 ± 1.2 | 2.8 ± 1.1 |
| Pyridoxine | 8.0 ± 0.1 | 5.2 ± 1.1 | 4.7 ± 0.2 | 3.3 ± 1.0 |
| Pyroglutamic acid | 8.1 ± 0.6 | 3.6 ± 1.0 | 5.2 ± 1.2 | 5.4 ± 0.8 |
| Pyrrolysine | 10.1 ± 0.3 | 4.9 ± 1.2 | 5.7 ± 0.8 | 3.6 ± 1.1 |
| Pyruvic Acid | 8.8 ± 0.5 | 2.6 ± 0.7 | 5.7 ± 0.3 | 3.7 ± 0.9 |
| Riboflavin | 10.7 ± 0.8 | 2.7 ± 0.9 | 2.5 ± 0.1 | 4.8 ± 1.2 |
| Ribose-5-Phosphate | 8.7 ± 1.0 | 5.0 ± 0.4 | 4.9 ± 0.7 | 5.9 ± 0.8 |
| Ribulose-1-Phosphate | 5.1 ± 0.5 | 5.3 ± 0.2 | 5.8 ± 0.8 | 2.4 ± 0.2 |
| Ribulose-5-Phosphate | 6.2 ± 1.0 | 5.7 ± 1.0 | 5.3 ± 0.2 | 4.8 ± 0.8 |
| Saccharopine | 6.4 ± 0.2 | 3.5 ± 0.7 | 4.5 ± 0.2 | 5.0 ± 0.5 |
| S-Adenosylhomocysteine | 5.2 ± 0.1 | 5.8 ± 0.8 | 3.1 ± 0.4 | 4.6 ± 0.9 |
| S-Adenosylmethioninamine | 9.3 ± 0.7 | 2.7 ± 0.1 | 4.8 ± 0.5 | 4.5 ± 0.9 |
| S-Adenosylmethionine | 7.8 ± 0.6 | 4.1 ± 0.7 | 3.3 ± 0.8 | 4.0 ± 0.3 |
| Sarcosine | 10.8 ± 0.7 | 3.8 ± 0.6 | 5.8 ± 0.5 | 2.4 ± 0.7 |
| Sedoheptulose-7-phosphate | 9.0 ± 1.0 | 4.6 ± 0.9 | 4.2 ± 0.8 | 3.4 ± 1.1 |
| Selenohomocysteine | 7.1 ± 0.3 | 2.7 ± 0.3 | 4.0 ± 0.8 | 4.1 ± 0.4 |
| Selenomethionine | 8.0 ± 0.3 | 3.7 ± 0.8 | 4.0 ± 0.5 | 5.6 ± 0.4 |
| Serine | 9.1 ± 0.5 | 2.7 ± 0.4 | 4.4 ± 0.7 | 5.7 ± 0.9 |
| S-Methyl-Thioinosine | 7.2 ± 0.6 | 3.4 ± 1.0 | 2.1 ± 1.0 | 2.8 ± 0.4 |
| Spermidine | 9.6 ± 0.4 | 4.3 ± 0.7 | 3.7 ± 0.2 | 3.8 ± 0.7 |
| Spermine | 7.2 ± 1.0 | 2.0 ± 1.1 | 4.4 ± 0.2 | 2.4 ± 0.2 |
| Succinate | 10.4 ± 0.8 | 2.3 ± 0.1 | 3.8 ± 0.6 | 4.0 ± 0.9 |
| Sucrose | 9.2 ± 1.0 | 3.3 ± 1.2 | 3.5 ± 0.5 | 5.3 ± 0.2 |
| Symmetric dimethylarginine | 6.8 ± 0.3 | 4.2 ± 0.5 | 3.6 ± 0.4 | 4.3 ± 0.1 |
| Taurine | 8.6 ± 1.2 | 5.0 ± 0.9 | 2.4 ± 0.1 | 5.6 ± 0.4 |
| Tetrahydroneopterin | 7.8 ± 0.8 | 2.2 ± 0.3 | 3.7 ± 0.8 | 5.8 ± 0.1 |
| Thiamine | 7.3 ± 0.3 | 5.6 ± 0.1 | 4.2 ± 0.8 | 5.5 ± 0.3 |
| Thiamine monophosphate | 8.3 ± 0.2 | 3.5 ± 0.3 | 5.9 ± 0.4 | 4.8 ± 0.6 |
| Threonic acid | 10.4 ± 0.5 | 2.3 ± 0.4 | 2.5 ± 0.6 | 2.9 ± 0.9 |
| Threonine | 8.4 ± 0.4 | 5.9 ± 1.0 | 2.5 ± 0.3 | 6.0 ± 1.1 |
| Thymidine | 8.6 ± 1.1 | 3.1 ± 0.4 | 4.4 ± 1.0 | 4.4 ± 1.2 |
| Thymidine Monophosphate | 10.5 ± 0.9 | 3.4 ± 0.2 | 2.2 ± 0.4 | 5.0 ± 0.4 |
| Thymine | 6.0 ± 1.1 | 6.0 ± 1.0 | 3.9 ± 0.7 | 2.0 ± 0.3 |
| Trigonelleine | 6.8 ± 1.0 | 5.7 ± 0.5 | 2.3 ± 0.8 | 3.2 ± 0.7 |
| Trimethylamine N-oxide | 9.3 ± 0.1 | 2.2 ± 1.0 | 4.3 ± 0.6 | 4.0 ± 0.3 |
| Trimethyllysine | 10.6 ± 0.3 | 3.4 ± 0.4 | 4.1 ± 0.9 | 5.8 ± 0.5 |
| Tryptophan | 9.3 ± 0.6 | 3.5 ± 0.5 | 2.1 ± 0.6 | 5.5 ± 0.6 |
| Tyramine | 10.3 ± 0.4 | 4.9 ± 0.8 | 2.7 ± 0.6 | 2.2 ± 0.5 |
| Tyrosine | 10.0 ± 0.7 | 2.9 ± 0.7 | 3.6 ± 0.6 | 5.6 ± 0.4 |
| Uracil | 6.2 ± 1.1 | 2.3 ± 0.6 | 5.8 ± 0.1 | 5.0 ± 0.7 |
| Uridine 5'-diphosphate | 10.3 ± 0.4 | 4.9 ± 0.2 | 4.6 ± 0.9 | 6.0 ± 1.1 |
| Uridine Monophosphate | 6.2 ± 1.1 | 3.7 ± 0.2 | 3.8 ± 1.2 | 2.2 ± 1.2 |
| Uridine-5-Diphosphogalactose | 9.8 ± 1.0 | 4.3 ± 0.4 | 4.1 ± 0.9 | 2.8 ± 1.0 |
| Urocanic acid | 7.6 ± 0.1 | 4.3 ± 0.6 | 2.5 ± 1.0 | 4.5 ± 0.4 |
| UTP | 9.9 ± 1.2 | 3.9 ± 0.7 | 3.3 ± 1.0 | 3.8 ± 0.4 |
| Valine | 9.4 ± 1.0 | 3.9 ± 0.8 | 3.2 ± 0.6 | 3.3 ± 0.3 |
| XMP | 8.1 ± 0.2 | 4.3 ± 1.2 | 5.1 ± 0.1 | 4.1 ± 1.0 |
| Xylulose-5-Phosphate | 7.1 ± 0.6 | 2.3 ± 0.6 | 2.1 ± 0.6 | 3.0 ± 0.2 |

### **Table S20** | Inter – day precision (as % RSD) (Cell extract as matrix, n = 5). The concentrations of QC_LLOQ_, QC_Low_, QC_Mid_ and QC_High_ were 10 ng/mL, 25 ng/mL, 100 ng/mL and 500 ng/mL respectively.

| Metabolite | QC_LLOQ_ | QC_Low_ | QC_Mid_ | QC_High_ |
| --- | --- | --- | --- | --- |
| 1-Methylguanine | 9.2 ± 0.7 | 5.0 ± 0.8 | 1.1 ± 0.2 | 1.0 ± 1.1 |
| 1-Methylhistidine | 11.9 ± 0.1 | 4.9 ± 1.1 | 6.1 ± 1.0 | 2.8 ± 1.0 |
| 1-Methylnicotinamide | 4.2 ± 0.9 | 1.8 ± 0.6 | 5.1 ± 1.2 | 3.0 ± 0.9 |
| 2-Aminoadipic Acid | 5.1 ± 0.8 | 5.7 ± 0.9 | 2.9 ± 1.2 | 3.1 ± 0.2 |
| 2-Deoxyadenosine | 11.3 ± 0.6 | 3.7 ± 0.8 | 3.6 ± 0.9 | 3.2 ± 0.3 |
| 2'-Deoxyadenosine 5'-monophosphate | 9.6 ± 1.1 | 5.8 ± 1.1 | 1.9 ± 0.5 | 6.4 ± 0.6 |
| 2'-deoxyadenosine 5'-triphosphate | 8.9 ± 0.6 | 5.8 ± 1.2 | 5.4 ± 0.6 | 2.3 ± 0.1 |
| 2'-Deoxycytidine-5'-monophosphate | 9.9 ± 0.7 | 2.7 ± 1.1 | 2.1 ± 1.2 | 1.5 ± 0.2 |
| 2-Deoxyguanosine | 7.0 ± 1.1 | 5.0 ± 1.2 | 6.3 ± 0.5 | 2.0 ± 1.2 |
| 2-hydroxybutyric acid | 3.4 ± 0.2 | 1.8 ± 1.1 | 1.0 ± 0.4 | 4.0 ± 1.1 |
| 2-hydroxyglutaric acid | 5.1 ± 1.0 | 2.6 ± 0.9 | 4.7 ± 1.0 | 4.2 ± 0.5 |
| 2-Hydroxyisocaproic acid | 3.8 ± 1.1 | 5.5 ± 0.8 | 3.0 ± 0.7 | 3.4 ± 0.4 |
| 2-ketobutyric acid | 7.9 ± 1.1 | 5.3 ± 1.1 | 6.4 ± 0.5 | 2.3 ± 1.0 |
| 2-oxoadipic acid | 10.9 ± 0.5 | 3.8 ± 1.1 | 2.3 ± 0.5 | 5.4 ± 0.6 |
| 2-Phenylglycine | 9.1 ± 0.5 | 5.6 ± 0.6 | 3.1 ± 0.3 | 4.0 ± 0.5 |
| 2-phosphoglycerate | 2.2 ± 0.6 | 3.9 ± 0.2 | 3.8 ± 0.6 | 5.8 ± 0.2 |
| 3-hydroxy-3-methyl-2-oxobutanoate | 5.1 ± 0.3 | 3.8 ± 0.4 | 4.4 ± 0.5 | 4.5 ± 0.3 |
| 3-hydroxybutyric acid | 3.6 ± 0.6 | 3.4 ± 0.5 | 5.4 ± 0.6 | 3.2 ± 0.2 |
| 3-Methylhistidine | 10.8 ± 0.9 | 3.8 ± 0.6 | 1.2 ± 0.5 | 1.8 ± 0.3 |
| 3-Phosphoglyceraldehyde | 5.5 ± 0.9 | 6.1 ± 0.9 | 5.1 ± 0.7 | 3.8 ± 0.4 |
| 3-phosphoglycerate | 2.6 ± 1.1 | 1.1 ± 0.5 | 4.7 ± 0.7 | 4.3 ± 0.3 |
| 4-amino-butyric acid | 1.4 ± 0.3 | 3.0 ± 0.7 | 1.6 ± 0.8 | 2.0 ± 1.1 |
| 4-aspartyl-phosphate | 10.4 ± 0.6 | 5.4 ± 0.1 | 4.2 ± 0.5 | 3.3 ± 0.9 |
| 4-Hydroxybenzoic acid | 2.8 ± 1.2 | 6.1 ± 0.8 | 4.1 ± 1.0 | 3.0 ± 1.0 |
| 4-Hydroxyphenylacetylglycine | 8.2 ± 0.2 | 6.2 ± 1.1 | 3.2 ± 0.6 | 2.4 ± 0.1 |
| 4-methyl-2-oxopentanoate | 1.1 ± 0.9 | 3.4 ± 0.3 | 6.2 ± 0.5 | 1.4 ± 0.9 |
| 4-Oxoglutaramate | 6.8 ± 0.6 | 3.9 ± 0.8 | 1.5 ± 1.0 | 4.8 ± 0.7 |
| 5-Hydroxyindolepyruvate | 5.6 ± 1.1 | 4.3 ± 0.3 | 6.1 ± 0.6 | 5.0 ± 0.7 |
| 5-Hydroxylysine | 5.6 ± 1.1 | 1.1 ± 1.2 | 4.7 ± 0.8 | 1.6 ± 0.7 |
| 5-Hydroxy-Tryptophan | 10.6 ± 1.1 | 3.6 ± 0.2 | 5.3 ± 0.4 | 3.3 ± 0.4 |
| 5-Phosphoribosylamine | 4.9 ± 0.4 | 1.4 ± 0.3 | 4.4 ± 1.0 | 5.1 ± 0.5 |
| 6-phosphogluconate | 10.4 ± 0.9 | 6.5 ± 0.9 | 4.9 ± 0.8 | 1.5 ± 1.0 |
| Acetylcarnitine | 4.5 ± 1.2 | 4.7 ± 0.4 | 5.9 ± 1.0 | 4.1 ± 0.4 |
| Aconitic Acid | 2.4 ± 0.6 | 1.2 ± 0.4 | 2.6 ± 0.8 | 1.8 ± 0.8 |
| Adenine | 4.0 ± 0.6 | 5.1 ± 0.3 | 3.3 ± 1.2 | 1.6 ± 1.1 |
| Adenosine | 6.5 ± 0.1 | 5.7 ± 0.3 | 6.2 ± 1.0 | 2.6 ± 0.9 |
| Adenosine 3',5'-cyclic monophosphate | 3.4 ± 0.8 | 6.5 ± 0.8 | 4.5 ± 0.7 | 2.8 ± 1.0 |
| Adenosine Monophosphate | 3.4 ± 1.1 | 3.1 ± 0.7 | 3.8 ± 0.4 | 2.7 ± 0.8 |
| Adenosine-5'-diphosphate | 11.4 ± 0.7 | 2.3 ± 0.2 | 2.6 ± 0.5 | 4.5 ± 1.1 |
| Adenosine-5'-triphosphate | 8.1 ± 0.5 | 5.7 ± 1.0 | 5.8 ± 0.6 | 1.1 ± 0.7 |
| Adenylosuccinate | 11.9 ± 0.7 | 2.1 ± 0.6 | 1.3 ± 1.2 | 2.1 ± 1.0 |
| Alanine | 8.6 ± 0.6 | 2.6 ± 0.7 | 2.9 ± 1.1 | 5.8 ± 0.9 |
| alpha-ketoglutaric acid | 9.6 ± 0.7 | 2.8 ± 0.8 | 4.1 ± 0.5 | 1.7 ± 1.0 |
| Anserine | 10.5 ± 0.4 | 5.9 ± 1.1 | 2.9 ± 1.1 | 3.4 ± 0.6 |
| Arginine | 7.7 ± 0.9 | 2.8 ± 0.7 | 5.0 ± 0.3 | 5.1 ± 1.0 |
| Argininosuccinic Acid | 7.1 ± 0.1 | 5.5 ± 0.7 | 2.9 ± 0.3 | 4.3 ± 0.8 |
| Asparagine | 9.8 ± 0.3 | 5.6 ± 1.0 | 5.8 ± 1.0 | 2.2 ± 0.1 |
| Aspartic acid | 5.7 ± 0.8 | 3.5 ± 0.2 | 1.5 ± 0.6 | 3.9 ± 0.9 |
| Asymmetric dimethylarginine | 9.0 ± 0.9 | 5.3 ± 0.8 | 5.4 ± 0.2 | 1.1 ± 0.6 |
| Azelaic acid | 3.8 ± 1.0 | 5.0 ± 0.7 | 4.1 ± 0.7 | 3.5 ± 0.9 |
| beta-Alanine | 5.7 ± 0.8 | 2.2 ± 0.5 | 5.8 ± 0.5 | 4.4 ± 0.3 |
| Betaine | 2.2 ± 0.6 | 3.8 ± 1.1 | 4.0 ± 1.1 | 6.2 ± 0.4 |
| Biopterin | 11.0 ± 1.2 | 6.1 ± 0.9 | 3.3 ± 0.2 | 5.1 ± 1.1 |
| Biotin | 5.8 ± 0.5 | 6.3 ± 0.3 | 5.2 ± 1.1 | 4.3 ± 0.8 |
| Carnitine | 3.6 ± 0.5 | 4.9 ± 1.2 | 6.5 ± 0.9 | 2.6 ± 0.6 |
| Carnosine | 8.4 ± 0.8 | 5.0 ± 0.8 | 2.1 ± 0.1 | 1.7 ± 0.4 |
| CDP-choline | 2.5 ± 0.3 | 5.4 ± 0.8 | 1.9 ± 0.9 | 4.2 ± 1.1 |
| cGMP | 11.5 ± 0.8 | 4.1 ± 0.5 | 5.9 ± 0.8 | 4.9 ± 0.7 |
| Choline | 6.6 ± 1.0 | 1.3 ± 1.0 | 3.4 ± 0.6 | 5.6 ± 0.4 |
| Cinnamic acid | 3.1 ± 0.5 | 4.9 ± 0.4 | 5.4 ± 1.1 | 2.1 ± 0.3 |
| Citraconic acid | 7.6 ± 1.1 | 5.8 ± 0.7 | 2.9 ± 0.5 | 4.3 ± 0.5 |
| Citramalic acid | 2.6 ± 0.3 | 4.6 ± 0.8 | 2.5 ± 1.1 | 4.6 ± 1.1 |
| Citric Acid | 8.4 ± 0.7 | 3.5 ± 0.4 | 2.0 ± 0.6 | 3.5 ± 0.4 |
| Citrulline | 8.9 ± 0.4 | 6.4 ± 0.4 | 3.0 ± 0.4 | 3.8 ± 0.9 |
| Creatine | 3.7 ± 1.0 | 3.1 ± 0.1 | 2.6 ± 0.8 | 1.6 ± 0.3 |
| Creatinine | 9.2 ± 1.1 | 4.6 ± 0.1 | 1.7 ± 0.4 | 5.5 ± 0.3 |
| Cystathionine | 1.3 ± 0.6 | 4.8 ± 0.4 | 2.7 ± 0.4 | 2.8 ± 0.4 |
| Cysteamine | 2.8 ± 0.6 | 5.3 ± 0.6 | 5.2 ± 0.7 | 5.3 ± 0.8 |
| Cysteine | 10.2 ± 0.7 | 6.4 ± 0.1 | 4.3 ± 1.1 | 2.4 ± 0.4 |
| Cysteinylglycine | 1.8 ± 0.4 | 2.3 ± 1.1 | 4.4 ± 0.7 | 1.3 ± 0.1 |
| Cystine | 4.5 ± 0.6 | 5.1 ± 0.2 | 4.0 ± 1.2 | 5.8 ± 0.7 |
| Cytidine | 3.7 ± 1.1 | 2.0 ± 0.8 | 4.4 ± 0.4 | 4.2 ± 0.5 |
| Cytidine 3',5'-cyclic monophosphate | 10.1 ± 1.1 | 2.1 ± 0.7 | 4.7 ± 0.6 | 6.0 ± 0.4 |
| Cytidine 5'-diphosphate | 9.4 ± 0.1 | 3.2 ± 0.6 | 1.4 ± 0.6 | 1.6 ± 0.8 |
| Cytidine 5'-triphosphate | 10.8 ± 0.8 | 6.3 ± 0.6 | 2.3 ± 1.1 | 2.6 ± 0.4 |
| Cytidine monophosphate | 10.1 ± 0.9 | 6.1 ± 0.5 | 5.1 ± 1.2 | 3.1 ± 0.3 |
| Cytosine | 5.1 ± 0.5 | 6.4 ± 0.3 | 2.6 ± 1.1 | 4.5 ± 0.6 |
| dADP | 6.0 ± 1.2 | 3.1 ± 1.0 | 4.7 ± 1.0 | 4.0 ± 0.2 |
| Diethanolamine | 11.8 ± 1.0 | 1.3 ± 0.8 | 6.4 ± 1.1 | 2.8 ± 0.9 |
| Dihydrobiopterin | 9.1 ± 0.5 | 3.4 ± 1.0 | 4.0 ± 0.2 | 1.0 ± 0.5 |
| Dihydroneopterin triphosphate | 11.1 ± 1.1 | 2.2 ± 1.1 | 6.4 ± 0.8 | 3.0 ± 0.5 |
| Dihydroxyacetonephosphate | 6.6 ± 0.8 | 4.5 ± 0.9 | 1.7 ± 1.1 | 3.7 ± 0.7 |
| Dimethylglycine | 1.8 ± 1.0 | 4.8 ± 0.4 | 3.2 ± 1.0 | 5.5 ± 0.3 |
| Ectoine | 5.6 ± 1.0 | 3.8 ± 1.1 | 2.7 ± 0.3 | 3.4 ± 0.1 |
| Erythrose-4-phosphate | 5.0 ± 0.6 | 2.1 ± 0.3 | 1.5 ± 0.6 | 2.0 ± 0.7 |
| FAD | 4.9 ± 0.5 | 3.0 ± 0.2 | 2.0 ± 1.1 | 1.6 ± 0.8 |
| Folic acid | 11.7 ± 0.7 | 6.4 ± 0.5 | 2.2 ± 0.8 | 1.7 ± 1.1 |
| Fructose-6-phosphate | 5.2 ± 0.8 | 1.0 ± 1.2 | 3.9 ± 0.9 | 1.3 ± 0.4 |
| Fumaric Acid | 6.3 ± 0.3 | 1.5 ± 0.5 | 2.6 ± 0.9 | 4.1 ± 0.2 |
| Galactosamine | 8.2 ± 0.3 | 3.7 ± 0.6 | 5.4 ± 1.0 | 5.0 ± 0.6 |
| Galactose-1-phosphate | 7.3 ± 0.6 | 1.0 ± 0.6 | 2.0 ± 0.8 | 5.7 ± 0.5 |
| Galactose-6-Phosphate | 8.3 ± 0.8 | 5.5 ± 1.2 | 2.5 ± 0.7 | 6.3 ± 0.8 |
| gamma-Glutamylcysteine | 2.2 ± 0.7 | 5.5 ± 0.9 | 3.3 ± 0.7 | 1.9 ± 0.9 |
| Glucosamine | 2.1 ± 0.6 | 2.1 ± 0.1 | 5.6 ± 0.4 | 2.9 ± 0.2 |
| Glucosamine-6-Phosphate | 8.9 ± 0.2 | 1.6 ± 1.0 | 1.5 ± 0.4 | 2.6 ± 1.0 |
| Glucose | 11.6 ± 0.6 | 4.7 ± 0.1 | 4.0 ± 0.9 | 1.8 ± 0.5 |
| Glucose-1-phosphate | 1.3 ± 0.9 | 3.8 ± 0.8 | 4.0 ± 0.3 | 5.3 ± 0.4 |
| Glucose-6-Phosphate | 10.5 ± 0.6 | 2.0 ± 0.5 | 2.3 ± 0.1 | 4.7 ± 0.9 |
| Glucuronic acid | 7.1 ± 0.6 | 5.0 ± 0.4 | 5.7 ± 0.4 | 4.6 ± 0.7 |
| Glutamic acid | 10.2 ± 1.0 | 2.7 ± 0.7 | 3.6 ± 0.9 | 4.3 ± 0.8 |
| Glutamine | 2.3 ± 0.5 | 1.2 ± 0.9 | 4.0 ± 0.2 | 5.6 ± 0.7 |
| Glutathione | 11.7 ± 0.6 | 5.6 ± 0.6 | 5.4 ± 0.1 | 1.4 ± 0.8 |
| Glutathionylspermine | 10.2 ± 0.2 | 5.3 ± 1.0 | 3.9 ± 0.2 | 5.0 ± 1.0 |
| Glycerophosphocholine | 4.4 ± 1.0 | 4.4 ± 1.1 | 3.3 ± 0.7 | 4.9 ± 1.1 |
| Glycine | 2.6 ± 0.4 | 1.8 ± 0.6 | 3.6 ± 0.5 | 6.3 ± 0.5 |
| Glycolic acid | 8.7 ± 0.5 | 4.6 ± 0.5 | 1.5 ± 0.1 | 5.3 ± 1.0 |
| Guanidinoacetic acid | 2.8 ± 0.9 | 5.5 ± 1.1 | 2.7 ± 1.0 | 5.3 ± 0.5 |
| Guanidinobutanoic acid | 2.0 ± 0.3 | 3.9 ± 0.3 | 4.7 ± 0.6 | 1.1 ± 1.1 |
| Guanine | 7.5 ± 0.6 | 6.3 ± 1.0 | 2.4 ± 0.7 | 3.8 ± 0.4 |
| Guanosine | 11.1 ± 0.7 | 6.3 ± 0.4 | 4.4 ± 0.7 | 4.9 ± 0.9 |
| Guanosine 5'-diphosphate | 4.3 ± 0.3 | 2.0 ± 0.9 | 3.9 ± 1.1 | 1.0 ± 0.6 |
| Guanosine-5'-monophosphate | 4.8 ± 0.4 | 2.1 ± 0.5 | 3.6 ± 1.1 | 2.2 ± 1.1 |
| Hercynine | 2.3 ± 0.5 | 2.9 ± 0.9 | 1.2 ± 0.2 | 4.6 ± 0.2 |
| Hippuric acid | 3.7 ± 0.1 | 5.3 ± 1.1 | 4.1 ± 0.2 | 2.1 ± 0.8 |
| Histamine | 4.3 ± 1.1 | 4.9 ± 0.3 | 3.3 ± 0.8 | 1.5 ± 0.5 |
| Histidine | 9.1 ± 0.3 | 1.1 ± 0.5 | 3.1 ± 0.1 | 1.2 ± 0.8 |
| Homoalanine | 3.9 ± 0.8 | 6.2 ± 0.8 | 1.5 ± 1.0 | 1.1 ± 0.4 |
| Homoarginine | 4.0 ± 0.6 | 3.6 ± 0.2 | 4.5 ± 0.9 | 2.4 ± 0.9 |
| Homocysteine | 4.7 ± 0.9 | 3.5 ± 0.6 | 3.7 ± 0.4 | 5.0 ± 0.5 |
| Homogentisic acid | 2.9 ± 0.6 | 5.9 ± 0.5 | 4.0 ± 0.1 | 3.9 ± 0.9 |
| Homoleucine | 2.7 ± 1.1 | 5.9 ± 0.5 | 2.0 ± 0.4 | 1.2 ± 0.6 |
| Homovanillic acid | 7.6 ± 1.0 | 4.1 ± 0.7 | 4.0 ± 0.5 | 4.9 ± 0.8 |
| Hydantoin-5-propionate | 6.1 ± 0.5 | 3.9 ± 1.0 | 1.7 ± 0.3 | 3.6 ± 0.2 |
| Hypotaurine | 5.4 ± 0.3 | 1.2 ± 0.9 | 3.3 ± 0.5 | 4.0 ± 1.0 |
| Hypoxanthine | 1.1 ± 0.4 | 1.3 ± 0.4 | 5.8 ± 1.0 | 1.4 ± 0.4 |
| Indole-3-acetamide | 8.9 ± 0.8 | 4.8 ± 0.5 | 5.1 ± 0.2 | 2.8 ± 0.9 |
| Inosine | 8.7 ± 0.6 | 3.9 ± 0.4 | 3.0 ± 1.0 | 2.1 ± 0.5 |
| Inosine 5'-monophosphate | 11.8 ± 0.5 | 4.7 ± 0.5 | 4.3 ± 0.7 | 3.8 ± 0.8 |
| Isocitric Acid | 3.4 ± 0.7 | 3.7 ± 0.6 | 4.2 ± 0.1 | 4.9 ± 1.1 |
| Isoleucine | 5.6 ± 0.8 | 4.7 ± 0.7 | 5.7 ± 1.2 | 2.3 ± 0.4 |
| Lactic Acid | 5.9 ± 0.5 | 2.0 ± 0.3 | 2.9 ± 0.7 | 2.5 ± 0.5 |
| Leucine | 3.1 ± 0.7 | 4.0 ± 0.9 | 2.8 ± 0.8 | 5.7 ± 1.1 |
| Levulinic acid | 1.3 ± 0.3 | 2.0 ± 1.2 | 1.9 ± 0.3 | 2.0 ± 0.4 |
| Lysine | 6.0 ± 0.2 | 4.0 ± 1.2 | 4.3 ± 0.9 | 1.8 ± 0.1 |
| Malic Acid | 6.0 ± 0.4 | 4.1 ± 0.2 | 6.1 ± 0.3 | 4.3 ± 0.8 |
| Mannose-1 with Mannose-6-Phosphate | 10.8 ± 1.1 | 4.1 ± 0.7 | 1.5 ± 1.2 | 5.9 ± 1.0 |
| Mesaconic acid | 5.5 ± 0.3 | 2.2 ± 0.4 | 4.1 ± 0.5 | 5.1 ± 0.3 |
| Methionine | 7.0 ± 0.9 | 5.8 ± 0.2 | 4.1 ± 0.2 | 3.3 ± 0.7 |
| Methionine sulfoxide | 3.0 ± 1.0 | 3.1 ± 1.0 | 6.0 ± 0.6 | 4.3 ± 0.9 |
| Methylmalonic Acid | 9.7 ± 0.6 | 5.9 ± 0.4 | 2.5 ± 1.0 | 5.7 ± 0.1 |
| N1-Acetylspermidine | 7.1 ± 0.7 | 3.8 ± 0.4 | 1.5 ± 0.2 | 3.2 ± 0.9 |
| N1-Acetylspermine | 2.6 ± 0.2 | 3.6 ± 0.3 | 1.1 ± 0.1 | 2.0 ± 0.1 |
| N8-Acetylspermidine | 4.4 ± 0.2 | 2.4 ± 0.3 | 2.1 ± 0.4 | 2.7 ± 0.2 |
| N-Acetyl-Aspartyl-Glutamate | 2.1 ± 0.4 | 6.4 ± 0.4 | 5.6 ± 0.8 | 2.1 ± 1.1 |
| N-Acetyl-Aspartyl-Glutamyl-Glutamate | 6.4 ± 0.2 | 5.3 ± 0.5 | 4.1 ± 0.4 | 4.9 ± 0.7 |
| N-Acetyl-Galactosamine | 10.4 ± 0.9 | 5.2 ± 0.3 | 3.6 ± 0.3 | 2.5 ± 0.9 |
| N-Acetyl-Leucine | 4.4 ± 1.0 | 3.8 ± 0.5 | 6.5 ± 0.6 | 2.9 ± 0.2 |
| N-Acetyl-Methionine | 2.3 ± 0.8 | 2.1 ± 0.8 | 2.4 ± 0.9 | 3.3 ± 0.5 |
| N-Acetylneuraminic acid | 1.9 ± 0.8 | 4.3 ± 0.5 | 5.9 ± 0.9 | 1.6 ± 0.5 |
| N-Acetyl-Proline | 4.1 ± 0.6 | 2.4 ± 0.8 | 2.7 ± 0.8 | 1.2 ± 0.3 |
| N-Acetyl-Putrescine | 2.0 ± 0.4 | 3.7 ± 0.7 | 2.2 ± 0.3 | 5.6 ± 1.1 |
| N-Acetyl-Serine | 11.7 ± 0.3 | 4.8 ± 0.1 | 2.2 ± 0.5 | 3.5 ± 0.9 |
| N-Acetyl-Serotonin | 9.8 ± 0.9 | 6.1 ± 0.3 | 1.4 ± 1.0 | 2.0 ± 1.0 |
| N-Acyl-Homoserine-lactone | 1.5 ± 0.3 | 1.4 ± 1.2 | 4.6 ± 0.6 | 2.4 ± 1.1 |
| NAD | 11.3 ± 0.4 | 5.3 ± 0.7 | 1.7 ± 0.6 | 6.2 ± 0.3 |
| NADH | 7.4 ± 0.3 | 1.4 ± 0.4 | 3.0 ± 1.0 | 1.9 ± 0.3 |
| NADP | 5.6 ± 0.5 | 3.9 ± 0.5 | 3.6 ± 1.1 | 4.2 ± 0.2 |
| NADPH | 2.7 ± 0.3 | 3.7 ± 0.8 | 5.3 ± 0.7 | 1.8 ± 0.5 |
| N-alpha-Acetyl-lysine | 6.9 ± 0.6 | 4.9 ± 0.1 | 1.3 ± 0.3 | 5.1 ± 0.2 |
| N-alpha-Acetyl-Ornithine | 4.1 ± 1.0 | 1.3 ± 0.6 | 6.2 ± 0.1 | 5.0 ± 0.3 |
| N-Carbamoyl-Proline | 9.5 ± 0.3 | 5.1 ± 1.1 | 5.2 ± 0.3 | 3.4 ± 0.2 |
| N-Carbamoyl-Sarcosine | 6.4 ± 1.1 | 5.2 ± 0.6 | 4.1 ± 0.6 | 1.6 ± 0.7 |
| N-Epsilon-Acetyl-lysine | 9.2 ± 1.1 | 1.7 ± 0.5 | 4.4 ± 0.4 | 2.0 ± 0.2 |
| Nicotinamide | 5.7 ± 0.8 | 3.5 ± 1.1 | 4.4 ± 1.0 | 3.4 ± 1.1 |
| N-Methyl-Aspartic Acid | 1.2 ± 0.8 | 5.0 ± 0.6 | 2.2 ± 0.6 | 3.8 ± 0.3 |
| O-Phosphoethanolamine | 8.9 ± 0.7 | 1.6 ± 0.1 | 1.6 ± 0.3 | 2.1 ± 1.1 |
| Ophthalmic acid | 6.8 ± 0.2 | 6.3 ± 1.2 | 2.4 ± 0.2 | 2.7 ± 0.7 |
| Ornithine | 9.2 ± 0.6 | 4.2 ± 1.0 | 4.4 ± 1.2 | 4.1 ± 0.5 |
| Orotidine 5'-monophosphate | 7.5 ± 0.6 | 2.6 ± 0.7 | 1.8 ± 0.9 | 1.8 ± 0.3 |
| Oxaloacetate | 2.6 ± 0.5 | 4.8 ± 0.6 | 1.3 ± 0.1 | 2.3 ± 0.8 |
| Pantothenic acid | 2.7 ± 0.6 | 5.8 ± 1.1 | 4.8 ± 0.3 | 3.5 ± 0.6 |
| Phenylalanine | 5.6 ± 1.0 | 3.5 ± 1.0 | 2.5 ± 0.3 | 4.5 ± 0.6 |
| Phenylpyruvate | 1.8 ± 1.1 | 3.0 ± 0.3 | 3.9 ± 0.2 | 5.6 ± 1.1 |
| Phosphodimethylethanolamine | 11.5 ± 0.2 | 1.9 ± 1.0 | 4.4 ± 1.0 | 4.8 ± 0.3 |
| Phosphoenolpyruvate | 9.4 ± 0.9 | 1.9 ± 0.4 | 2.2 ± 0.3 | 5.4 ± 0.3 |
| Phosphorylcholine | 8.7 ± 1.0 | 6.3 ± 0.1 | 2.6 ± 0.7 | 6.2 ± 0.6 |
| Pipecolic acid | 11.3 ± 1.1 | 4.4 ± 0.3 | 6.1 ± 0.8 | 6.4 ± 0.1 |
| Proline | 10.6 ± 0.7 | 1.7 ± 0.2 | 2.6 ± 0.6 | 2.8 ± 0.8 |
| Putrescine | 5.6 ± 0.7 | 4.6 ± 0.7 | 2.1 ± 0.4 | 4.1 ± 0.2 |
| Pyridoxal | 10.0 ± 0.4 | 5.2 ± 0.7 | 3.8 ± 0.2 | 5.9 ± 0.6 |
| Pyridoxamine | 3.2 ± 0.7 | 3.5 ± 0.9 | 2.2 ± 0.6 | 4.5 ± 0.9 |
| Pyridoxine | 1.3 ± 1.1 | 3.7 ± 0.9 | 2.7 ± 1.0 | 6.3 ± 0.9 |
| Pyroglutamic acid | 5.0 ± 0.7 | 6.2 ± 0.5 | 6.0 ± 0.3 | 2.6 ± 0.7 |
| Pyrrolysine | 4.5 ± 1.1 | 4.6 ± 0.2 | 1.8 ± 0.3 | 5.8 ± 0.5 |
| Pyruvic Acid | 8.2 ± 0.2 | 1.2 ± 0.9 | 4.2 ± 1.2 | 5.4 ± 0.8 |
| Riboflavin | 5.3 ± 1.0 | 1.2 ± 1.0 | 3.1 ± 0.6 | 1.5 ± 0.7 |
| Ribose-5-Phosphate | 10.2 ± 0.5 | 4.5 ± 0.8 | 1.0 ± 1.1 | 4.9 ± 1.1 |
| Ribulose-1-Phosphate | 8.0 ± 1.1 | 3.6 ± 0.2 | 1.3 ± 0.3 | 3.4 ± 1.0 |
| Ribulose-5-Phosphate | 9.6 ± 0.9 | 6.2 ± 0.8 | 1.3 ± 0.9 | 4.1 ± 1.0 |
| Saccharopine | 5.4 ± 0.5 | 4.9 ± 0.6 | 5.4 ± 0.4 | 4.9 ± 1.1 |
| S-Adenosylhomocysteine | 6.6 ± 0.9 | 4.7 ± 0.6 | 3.3 ± 0.2 | 1.3 ± 0.3 |
| S-Adenosylmethioninamine | 5.1 ± 1.1 | 2.3 ± 1.0 | 5.3 ± 0.2 | 2.5 ± 0.6 |
| S-Adenosylmethionine | 3.0 ± 1.0 | 5.0 ± 0.7 | 2.9 ± 0.9 | 2.8 ± 0.5 |
| Sarcosine | 4.0 ± 0.5 | 5.6 ± 0.8 | 5.1 ± 0.6 | 3.0 ± 0.6 |
| Sedoheptulose-7-phosphate | 2.3 ± 0.1 | 1.6 ± 1.1 | 4.1 ± 0.5 | 2.1 ± 0.4 |
| Selenohomocysteine | 6.3 ± 0.2 | 1.0 ± 1.2 | 4.8 ± 0.5 | 4.2 ± 0.6 |
| Selenomethionine | 6.9 ± 0.1 | 1.8 ± 1.1 | 5.0 ± 1.0 | 3.9 ± 0.6 |
| Serine | 8.0 ± 1.2 | 1.3 ± 0.2 | 1.1 ± 1.0 | 5.2 ± 0.4 |
| S-Methyl-Thioinosine | 2.0 ± 0.7 | 3.5 ± 1.1 | 4.5 ± 1.1 | 5.1 ± 0.6 |
| Spermidine | 5.1 ± 1.1 | 2.5 ± 0.5 | 3.3 ± 1.0 | 2.1 ± 1.1 |
| Spermine | 3.5 ± 0.1 | 1.1 ± 0.8 | 4.5 ± 0.7 | 2.0 ± 0.7 |
| Succinate | 9.8 ± 0.7 | 5.9 ± 0.5 | 4.0 ± 1.2 | 4.9 ± 0.7 |
| Sucrose | 5.5 ± 0.4 | 3.7 ± 0.2 | 1.2 ± 1.2 | 6.2 ± 0.2 |
| Symmetric dimethylarginine | 9.6 ± 1.2 | 4.5 ± 0.2 | 1.9 ± 0.3 | 3.3 ± 0.5 |
| Taurine | 3.9 ± 0.9 | 5.7 ± 0.3 | 5.2 ± 0.8 | 5.1 ± 0.4 |
| Tetrahydroneopterin | 8.4 ± 0.6 | 2.7 ± 0.9 | 4.0 ± 0.5 | 5.4 ± 0.8 |
| Thiamine | 6.2 ± 0.6 | 4.8 ± 0.3 | 4.0 ± 0.6 | 1.1 ± 1.0 |
| Thiamine monophosphate | 7.3 ± 0.3 | 3.6 ± 1.2 | 6.3 ± 1.2 | 4.5 ± 1.1 |
| Threonic acid | 10.1 ± 0.5 | 1.6 ± 0.5 | 3.9 ± 0.4 | 6.0 ± 1.0 |
| Threonine | 8.5 ± 0.6 | 6.3 ± 0.9 | 2.0 ± 0.5 | 2.3 ± 0.8 |
| Thymidine | 11.9 ± 1.0 | 2.3 ± 1.0 | 1.4 ± 1.1 | 2.4 ± 1.2 |
| Thymidine Monophosphate | 4.7 ± 0.7 | 1.7 ± 0.8 | 5.2 ± 0.7 | 3.3 ± 0.8 |
| Thymine | 7.2 ± 0.2 | 1.5 ± 1.1 | 3.8 ± 0.2 | 3.8 ± 0.6 |
| Trigonelleine | 8.6 ± 1.0 | 2.6 ± 1.1 | 4.0 ± 0.5 | 1.8 ± 0.3 |
| Trimethylamine N-oxide | 11.4 ± 0.1 | 2.3 ± 0.1 | 5.8 ± 1.1 | 3.3 ± 1.0 |
| Trimethyllysine | 7.2 ± 0.3 | 4.7 ± 0.7 | 2.6 ± 0.8 | 2.3 ± 0.8 |
| Tryptophan | 11.4 ± 0.4 | 3.2 ± 0.1 | 2.8 ± 0.2 | 6.2 ± 0.2 |
| Tyramine | 10.5 ± 0.3 | 6.3 ± 0.3 | 4.5 ± 0.9 | 6.3 ± 0.9 |
| Tyrosine | 3.6 ± 0.2 | 1.6 ± 0.7 | 4.6 ± 0.6 | 4.0 ± 0.8 |
| Uracil | 3.4 ± 0.5 | 6.1 ± 0.1 | 5.4 ± 0.2 | 5.3 ± 0.5 |
| Uridine 5'-diphosphate | 7.5 ± 0.9 | 4.8 ± 0.3 | 2.6 ± 1.1 | 1.3 ± 0.9 |
| Uridine Monophosphate | 1.4 ± 0.4 | 1.0 ± 0.2 | 4.7 ± 0.6 | 4.4 ± 0.5 |
| Uridine-5-Diphosphogalactose | 7.3 ± 0.5 | 5.7 ± 0.3 | 4.5 ± 0.3 | 5.2 ± 0.9 |
| Urocanic acid | 11.9 ± 0.3 | 5.7 ± 1.2 | 2.2 ± 1.0 | 5.6 ± 0.4 |
| UTP | 2.6 ± 0.9 | 3.4 ± 0.4 | 4.4 ± 0.3 | 5.8 ± 0.2 |
| Valine | 9.7 ± 1.0 | 4.7 ± 0.8 | 5.1 ± 1.1 | 3.5 ± 0.6 |
| XMP | 6.1 ± 0.5 | 4.0 ± 1.2 | 4.0 ± 0.1 | 6.4 ± 1.1 |
| Xylulose-5-Phosphate | 8.8 ± 1.2 | 5.3 ± 0.2 | 1.3 ± 1.2 | 1.3 ± 0.4 |

### **Table S21** | Inter – day precision (as % RSD) (Urine extract as matrix, n = 5). The concentrations of QC_LLOQ_, QC_Low_, QC_Mid_ and QC_High_ were 10 ng/mL, 25 ng/mL, 100 ng/mL and 500 ng/mL respectively.

| Metabolite | QC_LLOQ_ | QC_Low_ | QC_Mid_ | QC_High_ |
| --- | --- | --- | --- | --- |
| 1-Methylguanine | 5.7 ± 0.5 | 5.9 ± 1.0 | 4.9 ± 0.3 | 5.2 ± 0.6 |
| 1-Methylhistidine | 9.8 ± 1.1 | 4.2 ± 1.0 | 3.0 ± 0.9 | 1.5 ± 0.7 |
| 1-Methylnicotinamide | 5.4 ± 0.1 | 2.2 ± 0.4 | 3.3 ± 0.2 | 4.2 ± 1.0 |
| 2-Aminoadipic Acid | 7.7 ± 0.8 | 5.3 ± 1.0 | 6.0 ± 1.1 | 4.5 ± 0.7 |
| 2-Deoxyadenosine | 4.1 ± 0.7 | 2.3 ± 0.9 | 5.2 ± 0.1 | 3.4 ± 1.1 |
| 2'-Deoxyadenosine 5'-monophosphate | 11.7 ± 0.9 | 3.4 ± 1.0 | 3.4 ± 0.5 | 2.3 ± 0.6 |
| 2'-deoxyadenosine 5'-triphosphate | 2.1 ± 0.4 | 4.9 ± 0.5 | 4.2 ± 0.6 | 4.3 ± 0.7 |
| 2'-Deoxycytidine-5'-monophosphate | 7.7 ± 0.6 | 1.8 ± 0.2 | 5.8 ± 1.1 | 5.4 ± 1.1 |
| 2-Deoxyguanosine | 7.3 ± 0.6 | 3.3 ± 1.1 | 1.8 ± 0.2 | 2.8 ± 0.4 |
| 2-hydroxybutyric acid | 7.7 ± 0.2 | 4.4 ± 0.8 | 3.7 ± 0.4 | 2.5 ± 0.3 |
| 2-hydroxyglutaric acid | 4.8 ± 0.8 | 2.5 ± 0.1 | 5.3 ± 0.7 | 5.8 ± 0.2 |
| 2-Hydroxyisocaproic acid | 3.8 ± 0.4 | 1.3 ± 0.2 | 4.1 ± 0.6 | 2.0 ± 0.4 |
| 2-ketobutyric acid | 7.1 ± 1.0 | 3.4 ± 0.8 | 4.5 ± 0.5 | 3.1 ± 1.1 |
| 2-oxoadipic acid | 10.3 ± 1.1 | 4.6 ± 0.8 | 4.4 ± 0.2 | 3.3 ± 0.7 |
| 2-Phenylglycine | 10.1 ± 1.1 | 5.1 ± 0.7 | 5.1 ± 1.2 | 5.7 ± 0.6 |
| 2-phosphoglycerate | 2.2 ± 1.1 | 3.1 ± 0.3 | 4.2 ± 0.5 | 5.4 ± 0.9 |
| 3-hydroxy-3-methyl-2-oxobutanoate | 11.0 ± 0.8 | 1.5 ± 0.6 | 2.8 ± 0.3 | 5.4 ± 1.1 |
| 3-hydroxybutyric acid | 5.5 ± 0.3 | 1.3 ± 0.3 | 5.4 ± 1.0 | 5.8 ± 0.1 |
| 3-Methylhistidine | 4.9 ± 0.3 | 5.0 ± 0.6 | 4.5 ± 1.2 | 1.9 ± 0.1 |
| 3-Phosphoglyceraldehyde | 8.7 ± 0.8 | 3.1 ± 0.2 | 2.7 ± 1.1 | 3.1 ± 0.3 |
| 3-phosphoglycerate | 8.6 ± 1.0 | 3.2 ± 0.4 | 1.5 ± 0.3 | 3.7 ± 0.5 |
| 4-amino-butyric acid | 6.2 ± 1.0 | 3.9 ± 0.2 | 1.9 ± 0.9 | 2.7 ± 0.1 |
| 4-aspartyl-phosphate | 8.8 ± 0.5 | 4.0 ± 1.1 | 2.7 ± 0.9 | 5.2 ± 0.3 |
| 4-Hydroxybenzoic acid | 3.5 ± 0.4 | 2.3 ± 0.7 | 3.3 ± 1.0 | 3.4 ± 0.4 |
| 4-Hydroxyphenylacetylglycine | 3.2 ± 1.1 | 3.9 ± 1.0 | 1.7 ± 0.2 | 3.4 ± 0.8 |
| 4-methyl-2-oxopentanoate | 2.6 ± 0.2 | 2.1 ± 0.3 | 3.7 ± 0.8 | 4.5 ± 1.2 |
| 4-Oxoglutaramate | 5.7 ± 0.5 | 2.1 ± 0.7 | 2.5 ± 0.4 | 2.5 ± 0.2 |
| 5-Hydroxyindolepyruvate | 9.4 ± 0.8 | 5.9 ± 1.2 | 5.0 ± 0.3 | 4.4 ± 0.7 |
| 5-Hydroxylysine | 10.8 ± 0.8 | 2.6 ± 0.3 | 1.5 ± 0.5 | 1.8 ± 1.1 |
| 5-Hydroxy-Tryptophan | 6.7 ± 0.3 | 1.5 ± 0.3 | 2.9 ± 0.3 | 2.1 ± 0.2 |
| 5-Phosphoribosylamine | 6.7 ± 0.5 | 3.4 ± 1.2 | 1.6 ± 0.9 | 2.5 ± 1.0 |
| 6-phosphogluconate | 11.8 ± 1.0 | 5.6 ± 0.2 | 4.5 ± 1.1 | 1.8 ± 0.9 |
| Acetylcarnitine | 11.6 ± 0.1 | 2.7 ± 0.5 | 5.2 ± 0.8 | 4.0 ± 0.3 |
| Aconitic Acid | 7.4 ± 1.0 | 1.8 ± 0.4 | 3.6 ± 0.3 | 3.9 ± 1.0 |
| Adenine | 10.2 ± 0.1 | 5.7 ± 0.9 | 4.8 ± 0.1 | 3.9 ± 0.8 |
| Adenosine | 5.8 ± 0.2 | 4.7 ± 0.4 | 1.9 ± 0.8 | 4.2 ± 0.5 |
| Adenosine 3',5'-cyclic monophosphate | 5.9 ± 1.1 | 2.8 ± 0.9 | 5.8 ± 0.3 | 5.8 ± 0.5 |
| Adenosine Monophosphate | 4.0 ± 1.0 | 2.8 ± 0.9 | 5.4 ± 1.0 | 5.1 ± 0.9 |
| Adenosine-5'-diphosphate | 9.3 ± 0.1 | 3.0 ± 0.4 | 3.2 ± 0.7 | 2.5 ± 0.9 |
| Adenosine-5'-triphosphate | 5.0 ± 0.5 | 4.2 ± 0.8 | 1.5 ± 1.1 | 4.0 ± 0.4 |
| Adenylosuccinate | 6.0 ± 0.4 | 5.6 ± 0.1 | 4.4 ± 1.0 | 4.6 ± 0.7 |
| Alanine | 10.4 ± 0.5 | 3.9 ± 0.9 | 3.5 ± 0.8 | 2.4 ± 0.5 |
| alpha-ketoglutaric acid | 8.6 ± 0.6 | 4.0 ± 0.6 | 1.9 ± 1.2 | 3.6 ± 0.8 |
| Anserine | 10.0 ± 0.2 | 4.4 ± 0.4 | 2.0 ± 0.8 | 6.0 ± 0.7 |
| Arginine | 2.7 ± 0.3 | 5.1 ± 1.0 | 1.9 ± 0.9 | 5.5 ± 0.9 |
| Argininosuccinic Acid | 10.3 ± 0.4 | 5.1 ± 0.9 | 3.4 ± 0.7 | 3.3 ± 0.4 |
| Asparagine | 7.0 ± 0.9 | 5.0 ± 1.0 | 5.2 ± 0.7 | 3.5 ± 1.0 |
| Aspartic acid | 6.3 ± 0.4 | 1.5 ± 0.7 | 3.2 ± 1.2 | 4.2 ± 0.5 |
| Asymmetric dimethylarginine | 10.4 ± 0.3 | 4.2 ± 1.0 | 5.4 ± 0.6 | 4.1 ± 0.9 |
| Azelaic acid | 10.4 ± 0.3 | 3.1 ± 0.8 | 2.0 ± 0.6 | 2.2 ± 0.7 |
| beta-Alanine | 5.5 ± 0.2 | 5.0 ± 0.7 | 2.3 ± 1.2 | 1.5 ± 0.4 |
| Betaine | 5.2 ± 0.3 | 4.3 ± 1.1 | 5.8 ± 0.4 | 5.6 ± 0.7 |
| Biopterin | 2.8 ± 0.9 | 3.4 ± 0.3 | 1.7 ± 0.6 | 1.7 ± 0.7 |
| Biotin | 7.7 ± 0.2 | 4.3 ± 0.5 | 3.7 ± 0.6 | 3.0 ± 1.1 |
| Carnitine | 8.7 ± 0.2 | 5.6 ± 0.8 | 5.9 ± 0.7 | 1.4 ± 0.5 |
| Carnosine | 7.8 ± 0.9 | 1.3 ± 0.3 | 2.0 ± 0.1 | 4.0 ± 1.0 |
| CDP-choline | 5.8 ± 0.5 | 4.9 ± 1.0 | 5.3 ± 0.1 | 4.1 ± 1.2 |
| cGMP | 6.6 ± 0.7 | 3.4 ± 0.7 | 5.6 ± 0.7 | 5.4 ± 0.6 |
| Choline | 7.3 ± 0.2 | 1.9 ± 0.2 | 4.9 ± 0.7 | 5.9 ± 0.9 |
| Cinnamic acid | 7.9 ± 1.1 | 5.1 ± 1.1 | 5.0 ± 1.1 | 4.5 ± 0.1 |
| Citraconic acid | 5.2 ± 1.2 | 3.1 ± 0.9 | 2.9 ± 0.9 | 4.8 ± 1.0 |
| Citramalic acid | 10.0 ± 0.4 | 4.3 ± 0.9 | 3.0 ± 0.5 | 5.9 ± 1.2 |
| Citric Acid | 10.2 ± 0.2 | 4.2 ± 1.1 | 1.3 ± 0.8 | 2.8 ± 0.1 |
| Citrulline | 7.3 ± 0.5 | 5.2 ± 0.9 | 3.7 ± 0.8 | 4.0 ± 0.2 |
| Creatine | 10.5 ± 0.7 | 4.3 ± 0.5 | 3.6 ± 1.2 | 1.7 ± 1.0 |
| Creatinine | 9.6 ± 1.1 | 4.8 ± 0.1 | 5.0 ± 0.7 | 3.7 ± 0.5 |
| Cystathionine | 6.5 ± 0.6 | 5.8 ± 1.0 | 4.6 ± 0.4 | 3.6 ± 0.6 |
| Cysteamine | 4.7 ± 1.0 | 5.4 ± 0.8 | 3.6 ± 0.3 | 2.2 ± 1.0 |
| Cysteine | 6.5 ± 0.5 | 5.7 ± 1.2 | 5.2 ± 1.0 | 6.0 ± 0.7 |
| Cysteinylglycine | 3.6 ± 0.9 | 3.2 ± 0.6 | 1.8 ± 1.1 | 5.6 ± 0.9 |
| Cystine | 10.3 ± 0.7 | 4.2 ± 0.8 | 2.1 ± 0.2 | 4.4 ± 0.8 |
| Cytidine | 3.2 ± 0.4 | 4.1 ± 1.0 | 3.9 ± 0.9 | 2.6 ± 0.6 |
| Cytidine 3',5'-cyclic monophosphate | 8.0 ± 0.3 | 4.9 ± 0.2 | 3.8 ± 0.2 | 6.0 ± 0.2 |
| Cytidine 5'-diphosphate | 5.2 ± 0.5 | 2.6 ± 0.9 | 4.4 ± 0.5 | 3.8 ± 0.9 |
| Cytidine 5'-triphosphate | 8.7 ± 0.8 | 4.1 ± 0.7 | 3.3 ± 0.3 | 3.3 ± 1.0 |
| Cytidine monophosphate | 8.4 ± 0.3 | 4.9 ± 1.0 | 1.4 ± 1.1 | 5.4 ± 0.5 |
| Cytosine | 12.0 ± 0.7 | 4.1 ± 0.7 | 2.2 ± 1.0 | 2.8 ± 0.4 |
| dADP | 3.2 ± 0.9 | 4.2 ± 0.7 | 4.3 ± 0.9 | 2.2 ± 1.1 |
| Diethanolamine | 5.5 ± 0.2 | 5.5 ± 1.1 | 2.0 ± 0.5 | 5.4 ± 0.2 |
| Dihydrobiopterin | 12.0 ± 0.8 | 2.0 ± 1.0 | 2.2 ± 0.6 | 5.7 ± 0.9 |
| Dihydroneopterin triphosphate | 8.3 ± 0.2 | 5.2 ± 1.0 | 5.9 ± 0.6 | 2.1 ± 0.4 |
| Dihydroxyacetonephosphate | 7.1 ± 1.0 | 5.9 ± 0.5 | 1.9 ± 0.4 | 4.3 ± 0.4 |
| Dimethylglycine | 3.5 ± 0.9 | 5.2 ± 0.9 | 3.6 ± 1.1 | 1.8 ± 0.8 |
| Ectoine | 10.8 ± 0.2 | 4.2 ± 0.5 | 2.8 ± 0.2 | 2.1 ± 0.3 |
| Erythrose-4-phosphate | 8.9 ± 0.3 | 1.6 ± 0.9 | 2.7 ± 1.2 | 5.3 ± 0.9 |
| FAD | 4.4 ± 0.8 | 2.8 ± 1.2 | 4.5 ± 0.6 | 3.4 ± 1.1 |
| Folic acid | 6.5 ± 1.0 | 1.7 ± 0.8 | 3.4 ± 0.6 | 4.7 ± 0.7 |
| Fructose-6-phosphate | 5.4 ± 0.2 | 1.4 ± 0.2 | 4.5 ± 0.3 | 1.5 ± 1.1 |
| Fumaric Acid | 3.8 ± 0.2 | 2.3 ± 1.2 | 1.9 ± 0.9 | 4.2 ± 0.1 |
| Galactosamine | 2.1 ± 0.3 | 1.7 ± 0.8 | 3.3 ± 0.5 | 2.5 ± 0.8 |
| Galactose-1-phosphate | 6.2 ± 0.4 | 1.7 ± 1.0 | 2.3 ± 0.1 | 5.5 ± 0.7 |
| Galactose-6-Phosphate | 11.2 ± 0.9 | 4.8 ± 0.2 | 4.5 ± 0.2 | 3.7 ± 0.5 |
| gamma-Glutamylcysteine | 2.3 ± 0.2 | 3.3 ± 0.2 | 3.3 ± 1.1 | 4.3 ± 1.0 |
| Glucosamine | 9.9 ± 0.6 | 1.6 ± 0.4 | 5.3 ± 0.4 | 3.1 ± 1.2 |
| Glucosamine-6-Phosphate | 6.7 ± 0.4 | 2.8 ± 1.1 | 2.9 ± 0.9 | 5.4 ± 0.8 |
| Glucose | 3.2 ± 0.4 | 4.1 ± 1.1 | 4.4 ± 0.2 | 4.6 ± 0.4 |
| Glucose-1-phosphate | 4.3 ± 0.4 | 3.6 ± 0.5 | 2.9 ± 0.3 | 5.2 ± 1.1 |
| Glucose-6-Phosphate | 9.7 ± 1.0 | 4.4 ± 0.2 | 1.6 ± 0.9 | 3.8 ± 1.0 |
| Glucuronic acid | 10.3 ± 1.0 | 5.4 ± 0.4 | 2.6 ± 1.0 | 3.2 ± 0.7 |
| Glutamic acid | 7.6 ± 0.2 | 3.4 ± 0.9 | 2.9 ± 0.8 | 4.1 ± 0.2 |
| Glutamine | 2.1 ± 0.6 | 1.7 ± 0.4 | 2.1 ± 0.5 | 4.2 ± 0.4 |
| Glutathione | 9.6 ± 0.4 | 2.9 ± 1.0 | 2.6 ± 0.8 | 4.4 ± 0.9 |
| Glutathionylspermine | 6.9 ± 1.1 | 4.2 ± 0.4 | 5.4 ± 1.0 | 2.3 ± 0.9 |
| Glycerophosphocholine | 6.0 ± 0.2 | 3.7 ± 1.2 | 3.3 ± 0.6 | 1.4 ± 0.3 |
| Glycine | 8.3 ± 1.0 | 5.4 ± 0.7 | 4.1 ± 1.2 | 6.0 ± 0.3 |
| Glycolic acid | 9.5 ± 0.2 | 3.5 ± 0.6 | 4.5 ± 0.6 | 6.0 ± 1.2 |
| Guanidinoacetic acid | 3.2 ± 0.9 | 2.1 ± 0.2 | 4.1 ± 0.2 | 4.5 ± 0.6 |
| Guanidinobutanoic acid | 4.1 ± 0.4 | 5.8 ± 0.4 | 1.5 ± 0.6 | 3.5 ± 0.2 |
| Guanine | 5.1 ± 0.7 | 1.8 ± 1.0 | 4.3 ± 1.1 | 3.3 ± 1.0 |
| Guanosine | 11.0 ± 0.7 | 3.2 ± 0.1 | 4.3 ± 0.6 | 3.3 ± 0.4 |
| Guanosine 5'-diphosphate | 4.1 ± 0.2 | 2.2 ± 0.3 | 2.2 ± 1.1 | 2.1 ± 0.7 |
| Guanosine-5'-monophosphate | 5.9 ± 0.4 | 5.0 ± 0.9 | 4.5 ± 0.7 | 1.8 ± 0.2 |
| Hercynine | 10.8 ± 0.7 | 1.4 ± 0.5 | 1.3 ± 0.1 | 2.4 ± 0.5 |
| Hippuric acid | 10.7 ± 1.1 | 5.1 ± 0.6 | 1.8 ± 1.0 | 1.6 ± 0.9 |
| Histamine | 8.4 ± 0.3 | 4.0 ± 1.2 | 3.6 ± 0.4 | 1.9 ± 1.0 |
| Histidine | 10.0 ± 0.4 | 1.3 ± 0.6 | 2.8 ± 0.9 | 1.9 ± 0.6 |
| Homoalanine | 8.3 ± 0.2 | 5.9 ± 0.5 | 1.4 ± 0.3 | 3.2 ± 0.9 |
| Homoarginine | 8.1 ± 1.1 | 1.3 ± 1.2 | 4.0 ± 0.4 | 2.7 ± 0.5 |
| Homocysteine | 7.8 ± 0.9 | 4.7 ± 0.2 | 5.5 ± 0.7 | 5.0 ± 0.4 |
| Homogentisic acid | 8.7 ± 0.7 | 1.6 ± 0.2 | 4.8 ± 0.4 | 5.7 ± 0.9 |
| Homoleucine | 11.4 ± 0.9 | 3.8 ± 0.9 | 4.9 ± 0.9 | 3.7 ± 1.2 |
| Homovanillic acid | 2.5 ± 0.7 | 2.9 ± 0.9 | 1.5 ± 0.5 | 2.0 ± 0.5 |
| Hydantoin-5-propionate | 6.3 ± 0.8 | 2.8 ± 1.2 | 2.9 ± 0.9 | 1.5 ± 0.2 |
| Hypotaurine | 3.4 ± 1.2 | 1.9 ± 0.9 | 2.3 ± 0.8 | 2.8 ± 1.0 |
| Hypoxanthine | 9.1 ± 0.9 | 4.9 ± 1.0 | 3.3 ± 0.7 | 3.4 ± 1.1 |
| Indole-3-acetamide | 10.8 ± 0.7 | 2.6 ± 0.4 | 3.5 ± 0.9 | 2.7 ± 0.9 |
| Inosine | 7.3 ± 0.9 | 4.9 ± 1.2 | 2.0 ± 1.1 | 5.4 ± 0.5 |
| Inosine 5'-monophosphate | 4.0 ± 1.0 | 5.9 ± 1.1 | 5.1 ± 0.7 | 2.3 ± 0.1 |
| Isocitric Acid | 11.0 ± 0.2 | 2.2 ± 0.4 | 2.2 ± 1.2 | 2.2 ± 0.8 |
| Isoleucine | 5.4 ± 0.4 | 5.3 ± 0.2 | 1.9 ± 0.8 | 3.1 ± 1.1 |
| Lactic Acid | 3.1 ± 1.1 | 4.3 ± 0.8 | 6.0 ± 0.5 | 4.8 ± 0.1 |
| Leucine | 4.1 ± 1.0 | 5.8 ± 0.7 | 1.7 ± 0.2 | 4.3 ± 0.2 |
| Levulinic acid | 2.5 ± 0.3 | 2.3 ± 0.5 | 5.8 ± 1.1 | 3.6 ± 0.9 |
| Lysine | 6.8 ± 0.9 | 2.4 ± 0.1 | 2.7 ± 0.5 | 5.0 ± 1.2 |
| Malic Acid | 11.4 ± 0.2 | 1.7 ± 0.9 | 4.2 ± 0.6 | 1.4 ± 0.5 |
| Mannose-1 with Mannose-6-Phosphate | 9.3 ± 0.3 | 1.3 ± 1.1 | 2.0 ± 1.0 | 2.4 ± 0.9 |
| Mesaconic acid | 2.6 ± 0.8 | 2.4 ± 0.9 | 3.0 ± 0.3 | 3.8 ± 1.2 |
| Methionine | 11.3 ± 1.1 | 5.1 ± 0.7 | 5.8 ± 0.9 | 2.5 ± 0.9 |
| Methionine sulfoxide | 8.9 ± 0.6 | 4.2 ± 0.3 | 5.0 ± 0.4 | 4.8 ± 1.2 |
| Methylmalonic Acid | 6.7 ± 1.2 | 5.5 ± 1.0 | 5.8 ± 0.3 | 2.3 ± 0.6 |
| N1-Acetylspermidine | 8.0 ± 0.8 | 2.5 ± 0.9 | 3.6 ± 0.6 | 3.2 ± 0.8 |
| N1-Acetylspermine | 6.2 ± 0.9 | 3.2 ± 0.6 | 3.3 ± 0.8 | 4.4 ± 0.6 |
| N8-Acetylspermidine | 9.2 ± 1.1 | 1.5 ± 0.6 | 2.3 ± 0.1 | 2.6 ± 0.2 |
| N-Acetyl-Aspartyl-Glutamate | 11.3 ± 0.2 | 3.1 ± 0.2 | 5.9 ± 1.0 | 5.2 ± 0.8 |
| N-Acetyl-Aspartyl-Glutamyl-Glutamate | 11.5 ± 0.8 | 3.3 ± 0.7 | 3.8 ± 0.7 | 1.8 ± 0.7 |
| N-Acetyl-Galactosamine | 8.9 ± 0.9 | 6.0 ± 0.8 | 2.0 ± 1.2 | 2.2 ± 0.2 |
| N-Acetyl-Leucine | 3.0 ± 0.3 | 2.0 ± 0.1 | 3.3 ± 0.3 | 4.0 ± 1.0 |
| N-Acetyl-Methionine | 11.7 ± 0.4 | 5.8 ± 0.3 | 2.8 ± 0.4 | 2.9 ± 0.5 |
| N-Acetylneuraminic acid | 11.9 ± 0.4 | 1.6 ± 0.1 | 3.8 ± 0.7 | 4.9 ± 0.5 |
| N-Acetyl-Proline | 7.5 ± 0.9 | 3.6 ± 0.3 | 5.3 ± 0.3 | 2.4 ± 0.6 |
| N-Acetyl-Putrescine | 5.3 ± 0.8 | 5.5 ± 0.9 | 4.2 ± 0.2 | 2.2 ± 0.2 |
| N-Acetyl-Serine | 3.3 ± 0.8 | 3.9 ± 1.1 | 4.9 ± 0.1 | 4.2 ± 0.2 |
| N-Acetyl-Serotonin | 3.5 ± 0.1 | 1.4 ± 0.1 | 6.0 ± 0.3 | 4.2 ± 1.0 |
| N-Acyl-Homoserine-lactone | 5.7 ± 0.9 | 4.1 ± 1.0 | 4.0 ± 1.1 | 5.7 ± 0.1 |
| NAD | 7.4 ± 1.1 | 5.0 ± 0.6 | 5.2 ± 0.4 | 5.9 ± 0.6 |
| NADH | 10.0 ± 0.8 | 1.4 ± 0.5 | 1.8 ± 0.8 | 3.8 ± 0.6 |
| NADP | 6.3 ± 0.7 | 3.4 ± 0.5 | 1.6 ± 0.2 | 1.5 ± 0.2 |
| NADPH | 7.1 ± 0.9 | 4.3 ± 0.8 | 3.2 ± 1.0 | 3.3 ± 1.1 |
| N-alpha-Acetyl-lysine | 6.1 ± 0.8 | 4.9 ± 1.1 | 5.4 ± 0.7 | 1.7 ± 1.2 |
| N-alpha-Acetyl-Ornithine | 7.0 ± 0.4 | 1.5 ± 0.9 | 5.4 ± 0.9 | 1.5 ± 1.2 |
| N-Carbamoyl-Proline | 7.6 ± 1.1 | 1.9 ± 0.9 | 4.9 ± 0.3 | 3.4 ± 1.0 |
| N-Carbamoyl-Sarcosine | 6.8 ± 1.1 | 3.1 ± 0.2 | 1.9 ± 0.5 | 5.8 ± 0.8 |
| N-Epsilon-Acetyl-lysine | 10.8 ± 0.4 | 2.5 ± 0.2 | 4.0 ± 1.1 | 4.7 ± 0.9 |
| Nicotinamide | 3.0 ± 0.4 | 3.3 ± 0.5 | 4.5 ± 0.3 | 3.7 ± 0.8 |
| N-Methyl-Aspartic Acid | 7.9 ± 0.2 | 5.8 ± 1.1 | 2.4 ± 1.1 | 1.5 ± 0.7 |
| O-Phosphoethanolamine | 12.0 ± 1.2 | 5.6 ± 0.2 | 5.4 ± 0.9 | 2.2 ± 0.9 |
| Ophthalmic acid | 8.1 ± 0.8 | 3.3 ± 0.8 | 2.2 ± 0.2 | 1.9 ± 0.6 |
| Ornithine | 7.9 ± 0.3 | 1.6 ± 0.2 | 5.3 ± 0.2 | 2.1 ± 0.6 |
| Orotidine 5'-monophosphate | 7.8 ± 0.5 | 2.3 ± 0.4 | 4.9 ± 0.2 | 1.7 ± 0.9 |
| Oxaloacetate | 5.1 ± 0.3 | 2.8 ± 1.2 | 4.4 ± 1.2 | 4.5 ± 0.8 |
| Pantothenic acid | 10.3 ± 0.4 | 5.3 ± 0.1 | 3.1 ± 0.5 | 5.8 ± 1.1 |
| Phenylalanine | 10.8 ± 0.5 | 1.8 ± 0.9 | 1.8 ± 0.4 | 4.2 ± 1.0 |
| Phenylpyruvate | 4.7 ± 0.4 | 2.5 ± 0.3 | 3.1 ± 0.8 | 5.8 ± 0.3 |
| Phosphodimethylethanolamine | 5.0 ± 0.5 | 4.9 ± 0.1 | 5.6 ± 1.0 | 2.9 ± 0.4 |
| Phosphoenolpyruvate | 7.0 ± 0.3 | 4.1 ± 0.9 | 2.1 ± 0.7 | 5.3 ± 0.5 |
| Phosphorylcholine | 6.4 ± 1.0 | 3.1 ± 0.5 | 5.7 ± 0.9 | 3.2 ± 0.8 |
| Pipecolic acid | 6.0 ± 0.3 | 1.5 ± 0.5 | 1.9 ± 1.1 | 1.6 ± 0.2 |
| Proline | 6.5 ± 0.3 | 4.8 ± 0.8 | 1.4 ± 0.2 | 5.3 ± 0.3 |
| Putrescine | 8.7 ± 1.1 | 5.8 ± 0.6 | 3.4 ± 1.1 | 2.9 ± 0.2 |
| Pyridoxal | 3.7 ± 0.1 | 5.6 ± 0.5 | 5.8 ± 0.3 | 3.9 ± 0.8 |
| Pyridoxamine | 7.5 ± 0.6 | 4.6 ± 0.3 | 3.1 ± 0.6 | 5.6 ± 0.8 |
| Pyridoxine | 9.9 ± 1.0 | 3.6 ± 0.8 | 4.1 ± 0.7 | 3.1 ± 0.8 |
| Pyroglutamic acid | 10.0 ± 1.1 | 3.9 ± 0.4 | 5.6 ± 0.8 | 4.7 ± 0.2 |
| Pyrrolysine | 3.8 ± 0.4 | 4.2 ± 0.4 | 5.0 ± 0.8 | 5.4 ± 1.0 |
| Pyruvic Acid | 7.8 ± 0.3 | 2.2 ± 0.6 | 1.7 ± 1.1 | 3.7 ± 0.7 |
| Riboflavin | 3.1 ± 0.8 | 2.7 ± 0.2 | 4.9 ± 0.3 | 5.6 ± 0.6 |
| Ribose-5-Phosphate | 9.2 ± 0.2 | 4.6 ± 0.3 | 5.2 ± 0.5 | 5.5 ± 0.8 |
| Ribulose-1-Phosphate | 2.1 ± 1.0 | 3.7 ± 0.4 | 2.0 ± 0.3 | 4.6 ± 1.1 |
| Ribulose-5-Phosphate | 6.5 ± 1.1 | 3.0 ± 0.3 | 5.3 ± 0.9 | 4.6 ± 0.9 |
| Saccharopine | 11.7 ± 1.2 | 5.3 ± 1.1 | 2.3 ± 0.8 | 4.0 ± 0.6 |
| S-Adenosylhomocysteine | 10.4 ± 0.2 | 2.6 ± 0.5 | 2.0 ± 1.1 | 6.0 ± 1.0 |
| S-Adenosylmethioninamine | 3.3 ± 0.8 | 2.7 ± 1.1 | 4.6 ± 0.8 | 3.0 ± 1.0 |
| S-Adenosylmethionine | 11.2 ± 0.1 | 2.8 ± 1.2 | 2.7 ± 0.6 | 5.8 ± 0.1 |
| Sarcosine | 2.5 ± 1.1 | 2.0 ± 0.8 | 1.6 ± 0.5 | 1.7 ± 0.6 |
| Sedoheptulose-7-phosphate | 9.6 ± 0.8 | 3.5 ± 0.4 | 1.8 ± 0.7 | 5.9 ± 0.7 |
| Selenohomocysteine | 3.3 ± 0.8 | 1.5 ± 0.7 | 5.4 ± 1.0 | 2.3 ± 0.6 |
| Selenomethionine | 5.3 ± 0.3 | 2.7 ± 1.0 | 5.1 ± 1.0 | 3.6 ± 0.4 |
| Serine | 4.0 ± 1.2 | 1.6 ± 0.5 | 4.1 ± 0.3 | 2.9 ± 0.6 |
| S-Methyl-Thioinosine | 4.2 ± 1.1 | 5.3 ± 0.6 | 4.0 ± 0.6 | 2.2 ± 0.4 |
| Spermidine | 11.6 ± 1.1 | 3.7 ± 0.5 | 4.4 ± 1.1 | 1.5 ± 1.1 |
| Spermine | 9.3 ± 0.7 | 5.7 ± 0.3 | 1.5 ± 1.1 | 4.7 ± 0.8 |
| Succinate | 7.1 ± 1.1 | 5.8 ± 0.8 | 5.7 ± 0.7 | 3.7 ± 1.0 |
| Sucrose | 10.2 ± 0.9 | 4.9 ± 0.6 | 2.8 ± 1.1 | 1.4 ± 0.7 |
| Symmetric dimethylarginine | 9.5 ± 0.7 | 1.7 ± 0.2 | 4.9 ± 0.4 | 5.2 ± 0.8 |
| Taurine | 7.2 ± 0.8 | 5.4 ± 0.2 | 6.0 ± 1.1 | 2.5 ± 1.0 |
| Tetrahydroneopterin | 6.3 ± 0.2 | 1.5 ± 0.7 | 3.5 ± 0.5 | 5.5 ± 0.8 |
| Thiamine | 7.6 ± 0.5 | 4.6 ± 1.0 | 2.0 ± 0.7 | 5.3 ± 1.2 |
| Thiamine monophosphate | 6.7 ± 1.1 | 3.7 ± 0.5 | 6.0 ± 0.2 | 3.9 ± 0.6 |
| Threonic acid | 6.9 ± 0.1 | 2.8 ± 0.1 | 4.4 ± 1.2 | 5.2 ± 0.4 |
| Threonine | 10.6 ± 0.7 | 4.3 ± 0.2 | 4.4 ± 0.4 | 4.2 ± 0.4 |
| Thymidine | 5.7 ± 0.6 | 5.3 ± 0.5 | 2.4 ± 1.0 | 4.2 ± 1.0 |
| Thymidine Monophosphate | 4.9 ± 0.3 | 2.4 ± 0.6 | 4.1 ± 0.3 | 1.4 ± 0.5 |
| Thymine | 6.6 ± 0.7 | 2.0 ± 0.5 | 5.4 ± 0.9 | 2.5 ± 0.4 |
| Trigonelleine | 2.9 ± 1.2 | 2.1 ± 0.6 | 5.0 ± 0.6 | 3.6 ± 0.4 |
| Trimethylamine N-oxide | 10.3 ± 0.6 | 5.8 ± 0.3 | 2.5 ± 1.0 | 5.2 ± 0.6 |
| Trimethyllysine | 6.7 ± 1.2 | 3.7 ± 0.5 | 3.4 ± 1.2 | 5.3 ± 1.2 |
| Tryptophan | 5.1 ± 0.3 | 3.0 ± 0.4 | 1.4 ± 0.8 | 1.4 ± 0.9 |
| Tyramine | 5.1 ± 0.9 | 3.2 ± 0.3 | 4.5 ± 0.7 | 4.6 ± 1.0 |
| Tyrosine | 9.4 ± 1.0 | 4.7 ± 0.4 | 4.7 ± 0.5 | 5.6 ± 0.4 |
| Uracil | 4.5 ± 1.0 | 2.6 ± 1.0 | 2.2 ± 0.1 | 1.6 ± 0.9 |
| Uridine 5'-diphosphate | 4.9 ± 0.9 | 5.9 ± 0.4 | 2.7 ± 0.3 | 1.9 ± 0.1 |
| Uridine Monophosphate | 5.5 ± 1.2 | 5.6 ± 1.2 | 3.2 ± 0.4 | 4.5 ± 0.3 |
| Uridine-5-Diphosphogalactose | 4.5 ± 0.5 | 1.7 ± 0.1 | 5.2 ± 0.2 | 3.0 ± 0.8 |
| Urocanic acid | 11.0 ± 0.8 | 1.6 ± 0.5 | 3.6 ± 0.2 | 5.2 ± 0.1 |
| UTP | 4.8 ± 1.0 | 2.9 ± 0.4 | 5.3 ± 0.2 | 2.5 ± 1.1 |
| Valine | 3.2 ± 1.0 | 3.2 ± 0.9 | 5.6 ± 0.7 | 5.4 ± 0.6 |
| XMP | 10.0 ± 0.9 | 3.1 ± 0.5 | 4.7 ± 0.1 | 2.2 ± 0.4 |
| Xylulose-5-Phosphate | 4.4 ± 0.4 | 5.2 ± 1.2 | 5.4 ± 0.9 | 5.8 ± 1.0 |

### **Table S22** | Intra – day accuracy (as determined by % recovery) (Colorectal organoid extract as matrix, n = 5). The concentrations of QC_LLOQ_, QC_Low_, QC_Mid_ and QC_High_ were 10 ng/mL, 25 ng/mL, 100 ng/mL and 500 ng/mL respectively.

| Accuracy, % | QC_LLOQ_ | QC_Low_ | QC_Mid_ | QC_High_ |
| --- | --- | --- | --- | --- |
| 1-Methylguanine | 113.0 ± 1.1 | 101.3 ± 1.7 | 98.4 ± 0.3 | 88.2 ± 1.4 |
| 1-Methylhistidine | 106.4 ± 1.8 | 106.1 ± 1.1 | 114.3 ± 1.5 | 100.8 ± 1.4 |
| 1-Methylnicotinamide | 106.0 ± 1.4 | 97.4 ± 0.7 | 101.3 ± 1.0 | 102.1 ± 1.0 |
| 2-Aminoadipic Acid | 101.1 ± 1.7 | 103.9 ± 0.7 | 98.2 ± 0.5 | 97.8 ± 0.8 |
| 2-Deoxyadenosine | 98.4 ± 1.6 | 92.9 ± 1.2 | 105.9 ± 0.4 | 102.2 ± 0.3 |
| 2'-Deoxyadenosine 5'-monophosphate | 91.2 ± 1.1 | 87.9 ± 0.5 | 93.0 ± 0.3 | 91.5 ± 1.1 |
| 2'-deoxyadenosine 5'-triphosphate | 100.3 ± 1.6 | 113.9 ± 1.8 | 98.1 ± 0.2 | 88.8 ± 1.2 |
| 2'-Deoxycytidine-5'-monophosphate | 93.4 ± 1.3 | 98.0 ± 0.6 | 98.5 ± 0.6 | 101.7 ± 0.4 |
| 2-Deoxyguanosine | 109.6 ± 0.9 | 103.6 ± 0.6 | 100.2 ± 0.5 | 105.0 ± 1.9 |
| 2-hydroxybutyric acid | 105.6 ± 1.0 | 97.3 ± 1.3 | 98.2 ± 1.2 | 105.2 ± 0.9 |
| 2-hydroxyglutaric acid | 107.7 ± 1.6 | 85.6 ± 1.2 | 90.9 ± 0.8 | 101.5 ± 1.1 |
| 2-Hydroxyisocaproic acid | 114.7 ± 0.9 | 95.2 ± 1.4 | 101.2 ± 2.0 | 95.6 ± 1.4 |
| 2-ketobutyric acid | 88.4 ± 1.1 | 109.0 ± 0.4 | 89.9 ± 1.5 | 89.0 ± 0.3 |
| 2-oxoadipic acid | 92.6 ± 1.6 | 99.0 ± 1.1 | 96.6 ± 0.6 | 85.8 ± 0.8 |
| 2-Phenylglycine | 93.2 ± 1.0 | 103.7 ± 1.5 | 104.5 ± 1.2 | 103.8 ± 0.8 |
| 2-phosphoglycerate | 94.6 ± 0.7 | 101.8 ± 2.0 | 104.9 ± 1.8 | 104.1 ± 1.6 |
| 3-hydroxy-3-methyl-2-oxobutanoate | 110.9 ± 1.8 | 91.9 ± 1.1 | 94.5 ± 1.7 | 86.8 ± 0.8 |
| 3-hydroxybutyric acid | 93.0 ± 1.6 | 104.8 ± 1.5 | 96.4 ± 0.7 | 104.3 ± 0.4 |
| 3-Methylhistidine | 94.2 ± 0.4 | 100.2 ± 0.6 | 88.2 ± 0.8 | 94.5 ± 0.8 |
| 3-Phosphoglyceraldehyde | 97.6 ± 1.0 | 100.9 ± 0.3 | 104.8 ± 1.8 | 97.4 ± 0.3 |
| 3-phosphoglycerate | 87.2 ± 0.7 | 92.8 ± 0.3 | 90.2 ± 0.4 | 86.0 ± 1.4 |
| 4-amino-butyric acid | 89.9 ± 1.6 | 95.8 ± 1.1 | 104.4 ± 1.0 | 101.3 ± 0.5 |
| 4-aspartyl-phosphate | 93.7 ± 0.4 | 104.3 ± 1.0 | 100.4 ± 0.6 | 86.9 ± 1.4 |
| 4-Hydroxybenzoic acid | 100.6 ± 1.5 | 99.7 ± 1.0 | 98.4 ± 1.4 | 103.7 ± 1.0 |
| 4-Hydroxyphenylacetylglycine | 89.0 ± 0.7 | 85.1 ± 0.6 | 93.4 ± 0.5 | 109.5 ± 1.6 |
| 4-methyl-2-oxopentanoate | 92.2 ± 1.1 | 106.5 ± 0.5 | 101.1 ± 1.6 | 87.5 ± 1.4 |
| 4-Oxoglutaramate | 105.8 ± 0.4 | 110.0 ± 1.9 | 91.7 ± 0.4 | 92.5 ± 1.5 |
| 5-Hydroxyindolepyruvate | 109.6 ± 1.4 | 90.7 ± 1.8 | 96.8 ± 0.3 | 114.1 ± 0.4 |
| 5-Hydroxylysine | 111.1 ± 1.9 | 103.0 ± 1.2 | 99.1 ± 1.2 | 93.3 ± 0.5 |
| 5-Hydroxy-Tryptophan | 107.0 ± 0.3 | 95.2 ± 1.3 | 104.1 ± 0.3 | 104.4 ± 0.6 |
| 5-Phosphoribosylamine | 114.7 ± 1.8 | 112.5 ± 1.1 | 105.1 ± 0.2 | 105.1 ± 1.0 |
| 6-phosphogluconate | 100.2 ± 1.6 | 102.2 ± 0.4 | 98.1 ± 0.7 | 98.1 ± 0.2 |
| Acetylcarnitine | 103.2 ± 0.8 | 94.9 ± 1.9 | 102.3 ± 1.2 | 89.4 ± 1.6 |
| Aconitic Acid | 110.6 ± 1.2 | 99.2 ± 1.3 | 95.1 ± 1.0 | 113.3 ± 1.8 |
| Adenine | 96.9 ± 0.8 | 101.6 ± 0.8 | 97.3 ± 1.1 | 97.3 ± 0.2 |
| Adenosine | 94.6 ± 1.3 | 100.8 ± 1.1 | 97.4 ± 0.8 | 99.1 ± 1.8 |
| Adenosine 3',5'-cyclic monophosphate | 111.1 ± 0.4 | 104.3 ± 0.3 | 114.6 ± 0.4 | 97.6 ± 1.4 |
| Adenosine Monophosphate | 85.8 ± 0.4 | 92.2 ± 0.8 | 94.5 ± 1.4 | 85.3 ± 1.0 |
| Adenosine-5'-diphosphate | 110.2 ± 0.3 | 86.1 ± 0.4 | 94.4 ± 1.7 | 100.3 ± 1.4 |
| Adenosine-5'-triphosphate | 102.7 ± 0.2 | 91.5 ± 1.3 | 96.6 ± 0.2 | 90.5 ± 1.6 |
| Adenylosuccinate | 108.0 ± 0.6 | 88.4 ± 0.6 | 90.4 ± 1.0 | 86.3 ± 0.4 |
| Alanine | 92.1 ± 1.5 | 93.6 ± 1.4 | 114.4 ± 1.4 | 111.8 ± 1.4 |
| alpha-ketoglutaric acid | 104.2 ± 1.1 | 99.2 ± 0.3 | 86.4 ± 0.6 | 93.5 ± 0.8 |
| Anserine | 98.2 ± 0.5 | 101.3 ± 1.3 | 108.9 ± 0.3 | 85.1 ± 0.8 |
| Arginine | 99.5 ± 1.0 | 96.1 ± 1.1 | 103.2 ± 1.8 | 98.0 ± 0.9 |
| Argininosuccinic Acid | 105.0 ± 1.3 | 102.1 ± 1.3 | 94.2 ± 0.4 | 99.3 ± 0.5 |
| Asparagine | 98.2 ± 0.9 | 104.0 ± 2.0 | 93.2 ± 0.3 | 114.4 ± 1.3 |
| Aspartic acid | 89.0 ± 1.2 | 102.7 ± 1.0 | 105.9 ± 0.3 | 101.9 ± 1.9 |
| Asymmetric dimethylarginine | 91.5 ± 1.4 | 96.3 ± 1.1 | 88.3 ± 0.4 | 97.8 ± 1.8 |
| Azelaic acid | 108.7 ± 0.9 | 101.5 ± 0.4 | 103.2 ± 1.9 | 104.3 ± 0.4 |
| beta-Alanine | 95.8 ± 1.7 | 102.6 ± 0.8 | 101.6 ± 0.4 | 98.9 ± 0.6 |
| Betaine | 112.4 ± 0.5 | 97.7 ± 1.7 | 100.3 ± 1.6 | 99.7 ± 0.4 |
| Biopterin | 97.4 ± 1.0 | 104.6 ± 0.2 | 97.5 ± 0.7 | 98.0 ± 0.8 |
| Biotin | 111.7 ± 0.9 | 104.9 ± 1.2 | 100.0 ± 1.1 | 100.3 ± 2.0 |
| Carnitine | 91.4 ± 0.7 | 100.2 ± 0.5 | 100.1 ± 0.3 | 103.3 ± 0.4 |
| Carnosine | 104.7 ± 0.3 | 98.3 ± 1.6 | 97.0 ± 1.4 | 101.0 ± 1.7 |
| CDP-choline | 90.2 ± 1.6 | 100.6 ± 0.6 | 98.9 ± 1.3 | 100.8 ± 1.1 |
| cGMP | 96.5 ± 1.3 | 99.0 ± 1.8 | 105.8 ± 0.9 | 101.6 ± 0.9 |
| Choline | 112.8 ± 0.9 | 85.9 ± 0.2 | 91.2 ± 1.4 | 95.8 ± 0.7 |
| Cinnamic acid | 100.2 ± 1.6 | 104.4 ± 1.9 | 105.8 ± 0.3 | 105.6 ± 1.6 |
| Citraconic acid | 106.0 ± 1.9 | 112.8 ± 0.2 | 113.3 ± 1.8 | 110.1 ± 0.4 |
| Citramalic acid | 96.5 ± 1.0 | 87.7 ± 1.4 | 106.8 ± 1.0 | 99.2 ± 1.1 |
| Citric Acid | 110.6 ± 0.8 | 105.4 ± 1.3 | 108.2 ± 1.7 | 106.5 ± 1.7 |
| Citrulline | 112.4 ± 0.6 | 98.0 ± 0.4 | 105.9 ± 0.6 | 101.2 ± 1.0 |
| Creatine | 96.0 ± 1.4 | 95.7 ± 1.8 | 96.6 ± 1.0 | 97.9 ± 1.9 |
| Creatinine | 87.2 ± 0.4 | 88.5 ± 1.4 | 114.8 ± 0.5 | 112.2 ± 1.5 |
| Cystathionine | 106.8 ± 0.8 | 102.0 ± 0.5 | 102.5 ± 1.1 | 104.6 ± 0.9 |
| Cysteamine | 94.5 ± 0.5 | 99.0 ± 0.6 | 107.1 ± 1.8 | 98.7 ± 0.4 |
| Cysteine | 100.5 ± 1.3 | 103.6 ± 1.8 | 101.4 ± 0.6 | 102.1 ± 2.0 |
| Cysteinylglycine | 107.0 ± 1.4 | 99.1 ± 1.0 | 104.1 ± 1.3 | 97.4 ± 1.8 |
| Cystine | 93.9 ± 0.9 | 107.4 ± 1.0 | 95.6 ± 1.8 | 98.0 ± 1.0 |
| Cytidine | 86.1 ± 1.5 | 96.3 ± 0.6 | 110.5 ± 1.1 | 113.0 ± 0.8 |
| Cytidine 3',5'-cyclic monophosphate | 110.9 ± 0.8 | 96.9 ± 0.7 | 103.6 ± 0.3 | 106.7 ± 1.6 |
| Cytidine 5'-diphosphate | 101.5 ± 0.4 | 103.6 ± 1.6 | 104.1 ± 1.9 | 103.0 ± 0.4 |
| Cytidine 5'-triphosphate | 100.2 ± 0.6 | 103.3 ± 1.6 | 96.6 ± 1.3 | 99.4 ± 0.3 |
| Cytidine monophosphate | 101.9 ± 1.8 | 102.0 ± 0.9 | 101.5 ± 0.6 | 100.6 ± 1.8 |
| Cytosine | 102.9 ± 1.7 | 97.4 ± 1.9 | 96.1 ± 0.4 | 100.0 ± 1.3 |
| dADP | 114.8 ± 0.4 | 99.8 ± 1.5 | 99.0 ± 0.8 | 88.8 ± 0.4 |
| Diethanolamine | 95.6 ± 1.2 | 98.4 ± 1.1 | 103.1 ± 0.9 | 99.1 ± 0.9 |
| Dihydrobiopterin | 104.7 ± 1.0 | 100.4 ± 1.0 | 104.6 ± 1.9 | 114.7 ± 0.5 |
| Dihydroneopterin triphosphate | 106.6 ± 1.5 | 103.1 ± 1.3 | 99.5 ± 1.3 | 98.7 ± 1.2 |
| Dihydroxyacetonephosphate | 104.0 ± 0.7 | 102.2 ± 0.2 | 106.0 ± 0.6 | 103.6 ± 0.9 |
| Dimethylglycine | 94.8 ± 0.9 | 104.7 ± 0.9 | 87.0 ± 1.5 | 90.7 ± 1.4 |
| Ectoine | 111.4 ± 1.0 | 113.5 ± 1.3 | 104.4 ± 1.4 | 90.6 ± 1.7 |
| Erythrose-4-phosphate | 98.5 ± 0.8 | 98.7 ± 1.3 | 104.3 ± 1.6 | 103.7 ± 1.4 |
| FAD | 90.5 ± 1.0 | 106.7 ± 1.8 | 109.9 ± 1.9 | 89.7 ± 0.9 |
| Folic acid | 96.5 ± 0.3 | 102.2 ± 1.8 | 96.3 ± 1.4 | 102.2 ± 0.5 |
| Fructose-6-phosphate | 111.4 ± 1.3 | 104.7 ± 1.5 | 100.9 ± 1.8 | 102.1 ± 0.9 |
| Fumaric Acid | 109.9 ± 1.9 | 102.3 ± 0.7 | 101.1 ± 0.7 | 103.2 ± 1.6 |
| Galactosamine | 102.9 ± 0.9 | 86.5 ± 0.8 | 112.1 ± 2.0 | 88.0 ± 1.3 |
| Galactose-1-phosphate | 94.5 ± 0.3 | 103.6 ± 1.3 | 102.2 ± 1.7 | 99.5 ± 0.7 |
| Galactose-6-Phosphate | 90.2 ± 1.6 | 110.5 ± 2.0 | 90.3 ± 0.8 | 96.0 ± 0.6 |
| gamma-Glutamylcysteine | 86.1 ± 0.5 | 99.4 ± 1.7 | 96.7 ± 0.9 | 104.9 ± 0.8 |
| Glucosamine | 92.5 ± 1.8 | 95.9 ± 1.5 | 100.7 ± 2.0 | 106.1 ± 0.9 |
| Glucosamine-6-Phosphate | 112.9 ± 0.9 | 94.8 ± 1.3 | 87.1 ± 1.7 | 89.0 ± 1.0 |
| Glucose | 111.7 ± 0.4 | 98.3 ± 0.3 | 103.6 ± 0.2 | 100.1 ± 1.4 |
| Glucose-1-phosphate | 114.3 ± 0.2 | 99.5 ± 0.8 | 92.0 ± 0.5 | 102.1 ± 1.9 |
| Glucose-6-Phosphate | 88.8 ± 0.8 | 101.6 ± 1.3 | 101.0 ± 1.1 | 105.6 ± 1.7 |
| Glucuronic acid | 95.0 ± 1.8 | 99.2 ± 0.7 | 97.1 ± 0.5 | 100.7 ± 1.1 |
| Glutamic acid | 95.1 ± 0.9 | 98.4 ± 0.8 | 104.2 ± 0.9 | 104.9 ± 0.3 |
| Glutamine | 114.0 ± 0.7 | 100.5 ± 2.0 | 99.1 ± 0.2 | 105.6 ± 0.5 |
| Glutathione | 90.1 ± 1.7 | 103.9 ± 1.4 | 98.5 ± 1.3 | 106.2 ± 1.4 |
| Glutathionylspermine | 105.4 ± 1.2 | 96.4 ± 1.8 | 100.9 ± 0.7 | 101.7 ± 0.9 |
| Glycerophosphocholine | 93.7 ± 0.9 | 99.5 ± 0.7 | 102.3 ± 1.3 | 102.7 ± 0.9 |
| Glycine | 103.0 ± 1.4 | 98.7 ± 1.8 | 109.8 ± 1.8 | 107.9 ± 2.0 |
| Glycolic acid | 102.1 ± 0.7 | 101.1 ± 0.7 | 100.2 ± 1.5 | 102.9 ± 2.0 |
| Guanidinoacetic acid | 111.7 ± 0.6 | 95.6 ± 0.8 | 106.4 ± 0.6 | 95.5 ± 1.6 |
| Guanidinobutanoic acid | 98.8 ± 0.3 | 91.6 ± 1.5 | 97.2 ± 0.2 | 86.3 ± 1.2 |
| Guanine | 90.9 ± 0.2 | 104.0 ± 1.8 | 97.5 ± 1.7 | 101.9 ± 2.0 |
| Guanosine | 91.8 ± 0.9 | 104.0 ± 1.7 | 103.2 ± 2.0 | 104.5 ± 1.0 |
| Guanosine 5'-diphosphate | 96.1 ± 1.1 | 113.1 ± 0.8 | 98.2 ± 0.8 | 107.0 ± 0.3 |
| Guanosine-5'-monophosphate | 102.2 ± 1.6 | 95.6 ± 1.1 | 99.6 ± 0.2 | 102.2 ± 1.7 |
| Hercynine | 97.6 ± 1.6 | 95.7 ± 0.5 | 98.5 ± 1.2 | 104.4 ± 1.5 |
| Hippuric acid | 111.6 ± 0.9 | 100.1 ± 1.7 | 105.0 ± 1.6 | 105.3 ± 1.1 |
| Histamine | 111.1 ± 1.8 | 100.8 ± 1.1 | 104.7 ± 1.5 | 100.2 ± 1.4 |
| Histidine | 95.7 ± 1.4 | 100.8 ± 1.8 | 102.6 ± 0.4 | 99.8 ± 1.7 |
| Homoalanine | 97.0 ± 1.0 | 104.0 ± 1.8 | 98.8 ± 1.4 | 103.2 ± 0.5 |
| Homoarginine | 105.9 ± 1.0 | 104.2 ± 1.4 | 102.4 ± 1.1 | 97.0 ± 1.8 |
| Homocysteine | 100.1 ± 0.8 | 94.1 ± 1.2 | 100.5 ± 0.3 | 96.0 ± 0.9 |
| Homogentisic acid | 107.1 ± 0.8 | 91.5 ± 1.3 | 94.2 ± 0.7 | 95.5 ± 1.6 |
| Homoleucine | 91.4 ± 0.9 | 96.4 ± 0.2 | 97.1 ± 1.2 | 106.3 ± 1.3 |
| Homovanillic acid | 90.3 ± 1.6 | 101.6 ± 1.5 | 102.4 ± 1.9 | 104.5 ± 0.5 |
| Hydantoin-5-propionate | 87.2 ± 0.4 | 112.6 ± 1.8 | 93.4 ± 1.3 | 86.9 ± 0.8 |
| Hypotaurine | 113.2 ± 0.6 | 92.9 ± 1.6 | 99.9 ± 1.3 | 113.1 ± 0.6 |
| Hypoxanthine | 89.5 ± 0.8 | 104.2 ± 1.4 | 104.2 ± 1.3 | 100.9 ± 1.3 |
| Indole-3-acetamide | 111.4 ± 0.9 | 104.0 ± 1.9 | 113.3 ± 1.2 | 101.0 ± 1.5 |
| Inosine | 102.1 ± 0.3 | 92.4 ± 1.3 | 92.7 ± 1.3 | 109.1 ± 1.6 |
| Inosine 5'-monophosphate | 98.4 ± 0.5 | 85.6 ± 1.0 | 111.1 ± 1.0 | 88.3 ± 0.9 |
| Isocitric Acid | 99.0 ± 0.8 | 104.6 ± 0.7 | 101.1 ± 0.3 | 105.4 ± 1.8 |
| Isoleucine | 95.8 ± 1.9 | 102.0 ± 0.9 | 90.9 ± 0.6 | 92.2 ± 0.3 |
| Lactic Acid | 99.5 ± 1.9 | 97.9 ± 1.4 | 105.3 ± 0.7 | 98.1 ± 1.2 |
| Leucine | 90.1 ± 1.7 | 97.0 ± 0.9 | 105.0 ± 1.4 | 101.3 ± 1.1 |
| Levulinic acid | 104.6 ± 0.9 | 100.9 ± 0.9 | 96.7 ± 0.5 | 97.6 ± 1.8 |
| Lysine | 103.8 ± 0.5 | 112.0 ± 0.4 | 85.3 ± 0.6 | 93.2 ± 1.7 |
| Malic Acid | 110.7 ± 0.8 | 105.0 ± 0.4 | 94.2 ± 0.4 | 88.4 ± 0.3 |
| Mannose-1 with Mannose-6-Phosphate | 100.1 ± 1.2 | 108.4 ± 1.2 | 92.7 ± 1.4 | 104.4 ± 1.2 |
| Mesaconic acid | 100.7 ± 0.9 | 96.1 ± 1.6 | 96.4 ± 1.6 | 103.6 ± 1.9 |
| Methionine | 103.5 ± 0.7 | 104.9 ± 0.8 | 102.9 ± 0.8 | 97.5 ± 1.9 |
| Methionine sulfoxide | 100.4 ± 0.7 | 102.9 ± 1.4 | 103.0 ± 0.6 | 104.2 ± 1.1 |
| Methylmalonic Acid | 113.0 ± 1.1 | 104.0 ± 0.5 | 98.5 ± 1.4 | 103.2 ± 0.8 |
| N1-Acetylspermidine | 93.9 ± 0.7 | 114.7 ± 1.8 | 98.3 ± 0.8 | 110.8 ± 1.4 |
| N1-Acetylspermine | 91.6 ± 0.9 | 102.0 ± 1.6 | 104.0 ± 1.6 | 105.2 ± 1.2 |
| N8-Acetylspermidine | 103.4 ± 0.7 | 104.5 ± 0.5 | 105.1 ± 1.1 | 94.7 ± 1.7 |
| N-Acetyl-Aspartyl-Glutamate | 110.4 ± 1.7 | 105.0 ± 0.2 | 86.9 ± 0.8 | 92.6 ± 1.6 |
| N-Acetyl-Aspartyl-Glutamyl-Glutamate | 106.6 ± 0.3 | 99.9 ± 0.8 | 111.6 ± 0.6 | 90.6 ± 1.4 |
| N-Acetyl-Galactosamine | 109.7 ± 1.1 | 103.5 ± 1.4 | 97.0 ± 0.3 | 100.2 ± 0.5 |
| N-Acetyl-Leucine | 108.8 ± 1.6 | 103.2 ± 0.3 | 101.0 ± 0.5 | 100.0 ± 1.2 |
| N-Acetyl-Methionine | 86.5 ± 0.8 | 104.3 ± 0.5 | 101.5 ± 1.7 | 106.4 ± 1.6 |
| N-Acetylneuraminic acid | 86.4 ± 1.1 | 96.2 ± 1.3 | 111.5 ± 1.7 | 109.5 ± 1.3 |
| N-Acetyl-Proline | 101.7 ± 1.7 | 100.8 ± 0.3 | 97.2 ± 0.3 | 102.1 ± 1.5 |
| N-Acetyl-Putrescine | 113.7 ± 0.5 | 101.5 ± 0.8 | 99.4 ± 1.3 | 103.9 ± 0.3 |
| N-Acetyl-Serine | 103.0 ± 0.9 | 99.4 ± 0.4 | 97.3 ± 0.6 | 103.8 ± 1.5 |
| N-Acetyl-Serotonin | 104.6 ± 0.9 | 99.1 ± 1.5 | 105.5 ± 0.4 | 98.4 ± 1.8 |
| N-Acyl-Homoserine-lactone | 94.8 ± 1.6 | 103.3 ± 0.3 | 104.6 ± 1.4 | 99.9 ± 0.6 |
| NAD | 88.1 ± 0.4 | 100.7 ± 1.5 | 103.9 ± 0.4 | 97.9 ± 1.9 |
| NADH | 102.4 ± 1.8 | 96.1 ± 1.8 | 104.1 ± 1.6 | 103.1 ± 1.4 |
| NADP | 105.4 ± 0.9 | 90.2 ± 1.1 | 107.4 ± 0.9 | 101.6 ± 0.5 |
| NADPH | 103.3 ± 1.8 | 96.8 ± 1.0 | 98.9 ± 1.6 | 105.5 ± 0.7 |
| N-alpha-Acetyl-lysine | 92.6 ± 0.4 | 87.3 ± 1.3 | 98.3 ± 0.5 | 91.9 ± 1.2 |
| N-alpha-Acetyl-Ornithine | 102.3 ± 0.6 | 88.6 ± 1.6 | 86.6 ± 0.3 | 113.9 ± 1.8 |
| N-Carbamoyl-Proline | 109.0 ± 1.3 | 98.6 ± 1.5 | 105.3 ± 1.7 | 101.9 ± 1.6 |
| N-Carbamoyl-Sarcosine | 104.9 ± 1.2 | 100.7 ± 0.3 | 96.7 ± 0.8 | 106.5 ± 1.9 |
| N-Epsilon-Acetyl-lysine | 98.8 ± 1.7 | 103.5 ± 1.2 | 105.2 ± 1.9 | 103.8 ± 0.7 |
| Nicotinamide | 89.3 ± 0.6 | 103.6 ± 1.6 | 96.5 ± 0.5 | 98.4 ± 0.4 |
| N-Methyl-Aspartic Acid | 103.2 ± 2.0 | 99.3 ± 1.5 | 104.0 ± 1.6 | 106.4 ± 1.7 |
| O-Phosphoethanolamine | 104.2 ± 1.2 | 102.8 ± 0.3 | 102.5 ± 1.3 | 102.1 ± 0.9 |
| Ophthalmic acid | 101.4 ± 1.4 | 96.9 ± 1.1 | 98.1 ± 0.4 | 98.4 ± 2.0 |
| Ornithine | 89.1 ± 1.7 | 103.7 ± 1.9 | 103.4 ± 0.7 | 97.1 ± 1.7 |
| Orotidine 5'-monophosphate | 96.2 ± 1.6 | 103.1 ± 0.6 | 105.4 ± 1.9 | 99.6 ± 1.8 |
| Oxaloacetate | 97.7 ± 1.0 | 103.9 ± 0.9 | 104.0 ± 0.9 | 97.8 ± 1.4 |
| Pantothenic acid | 94.7 ± 1.5 | 102.5 ± 0.4 | 104.5 ± 1.3 | 98.2 ± 1.5 |
| Phenylalanine | 106.2 ± 1.5 | 98.4 ± 1.7 | 104.0 ± 1.5 | 105.8 ± 1.7 |
| Phenylpyruvate | 100.1 ± 1.1 | 113.5 ± 1.4 | 103.5 ± 0.8 | 99.7 ± 1.9 |
| Phosphodimethylethanolamine | 108.1 ± 0.4 | 100.8 ± 0.7 | 100.8 ± 0.2 | 92.5 ± 0.4 |
| Phosphoenolpyruvate | 96.2 ± 0.5 | 98.1 ± 0.6 | 105.6 ± 1.9 | 105.2 ± 0.4 |
| Phosphorylcholine | 106.2 ± 1.9 | 103.0 ± 1.2 | 103.0 ± 1.8 | 85.4 ± 0.5 |
| Pipecolic acid | 102.8 ± 1.3 | 104.3 ± 1.3 | 105.5 ± 0.6 | 100.4 ± 0.3 |
| Proline | 111.1 ± 1.8 | 104.9 ± 1.5 | 97.4 ± 0.2 | 100.2 ± 0.9 |
| Putrescine | 113.5 ± 0.4 | 91.0 ± 1.2 | 94.0 ± 0.6 | 101.2 ± 1.8 |
| Pyridoxal | 95.8 ± 1.6 | 103.7 ± 0.6 | 100.6 ± 0.7 | 97.4 ± 0.8 |
| Pyridoxamine | 108.4 ± 1.7 | 105.0 ± 1.3 | 88.6 ± 0.7 | 107.6 ± 1.2 |
| Pyridoxine | 111.6 ± 1.4 | 104.8 ± 0.3 | 102.5 ± 1.0 | 99.9 ± 0.4 |
| Pyroglutamic acid | 88.6 ± 1.7 | 101.2 ± 1.0 | 102.4 ± 0.3 | 103.3 ± 1.9 |
| Pyrrolysine | 107.5 ± 0.8 | 104.5 ± 0.3 | 103.9 ± 0.8 | 98.4 ± 1.6 |
| Pyruvic Acid | 99.2 ± 1.8 | 102.8 ± 0.5 | 98.6 ± 0.3 | 101.6 ± 1.6 |
| Riboflavin | 101.4 ± 1.1 | 102.1 ± 1.4 | 100.1 ± 1.1 | 105.0 ± 1.7 |
| Ribose-5-Phosphate | 109.0 ± 1.4 | 98.8 ± 0.8 | 99.1 ± 1.9 | 105.2 ± 0.8 |
| Ribulose-1-Phosphate | 101.3 ± 0.6 | 85.2 ± 0.7 | 106.2 ± 0.6 | 96.9 ± 1.6 |
| Ribulose-5-Phosphate | 103.0 ± 0.7 | 98.6 ± 0.7 | 94.7 ± 0.5 | 113.2 ± 0.2 |
| Saccharopine | 93.2 ± 1.0 | 96.4 ± 1.4 | 97.9 ± 1.0 | 101.6 ± 0.4 |
| S-Adenosylhomocysteine | 94.3 ± 1.0 | 103.8 ± 1.1 | 102.8 ± 0.2 | 101.8 ± 1.2 |
| S-Adenosylmethioninamine | 92.8 ± 1.1 | 100.4 ± 1.2 | 104.3 ± 1.2 | 101.9 ± 0.4 |
| S-Adenosylmethionine | 94.4 ± 0.5 | 99.6 ± 0.3 | 106.0 ± 0.6 | 104.4 ± 1.7 |
| Sarcosine | 96.2 ± 1.8 | 100.4 ± 0.8 | 105.6 ± 1.8 | 100.1 ± 0.3 |
| Sedoheptulose-7-phosphate | 88.9 ± 1.7 | 87.4 ± 0.3 | 94.6 ± 0.6 | 112.4 ± 1.7 |
| Selenohomocysteine | 108.9 ± 0.6 | 101.4 ± 1.2 | 94.5 ± 0.7 | 108.6 ± 1.3 |
| Selenomethionine | 88.4 ± 1.1 | 87.5 ± 1.1 | 100.5 ± 1.9 | 112.4 ± 1.6 |
| Serine | 100.7 ± 1.3 | 87.7 ± 1.5 | 100.1 ± 1.5 | 96.8 ± 0.5 |
| S-Methyl-Thioinosine | 110.0 ± 0.6 | 98.9 ± 1.3 | 85.9 ± 1.6 | 101.0 ± 0.4 |
| Spermidine | 95.7 ± 1.1 | 113.2 ± 1.9 | 110.8 ± 0.3 | 100.5 ± 0.3 |
| Spermine | 97.7 ± 1.9 | 85.6 ± 0.2 | 97.9 ± 1.5 | 95.1 ± 1.3 |
| Succinate | 91.6 ± 1.7 | 100.7 ± 1.4 | 88.1 ± 1.8 | 106.7 ± 2.0 |
| Sucrose | 89.4 ± 0.3 | 98.0 ± 1.3 | 96.2 ± 0.8 | 100.3 ± 1.2 |
| Symmetric dimethylarginine | 110.7 ± 0.7 | 104.1 ± 0.9 | 97.8 ± 0.7 | 97.9 ± 1.1 |
| Taurine | 103.2 ± 1.3 | 99.6 ± 1.8 | 102.0 ± 1.3 | 105.8 ± 1.9 |
| Tetrahydroneopterin | 99.9 ± 0.9 | 90.2 ± 0.9 | 93.4 ± 0.6 | 95.2 ± 1.1 |
| Thiamine | 102.4 ± 1.2 | 109.2 ± 1.5 | 104.9 ± 0.7 | 88.2 ± 1.4 |
| Thiamine monophosphate | 90.4 ± 0.7 | 89.8 ± 0.5 | 93.5 ± 0.3 | 94.2 ± 1.9 |
| Threonic acid | 108.1 ± 1.4 | 113.5 ± 1.7 | 85.9 ± 0.9 | 106.2 ± 1.6 |
| Threonine | 111.1 ± 1.7 | 105.6 ± 1.4 | 93.1 ± 0.9 | 86.1 ± 0.6 |
| Thymidine | 114.5 ± 2.0 | 88.3 ± 1.0 | 94.1 ± 1.4 | 94.1 ± 0.6 |
| Thymidine Monophosphate | 106.2 ± 1.0 | 105.7 ± 1.7 | 97.1 ± 0.8 | 93.4 ± 0.8 |
| Thymine | 108.2 ± 1.4 | 98.4 ± 1.9 | 92.5 ± 1.6 | 91.7 ± 1.7 |
| Trigonelleine | 109.0 ± 0.4 | 103.4 ± 1.0 | 99.4 ± 0.7 | 102.5 ± 0.4 |
| Trimethylamine N-oxide | 106.9 ± 0.6 | 96.5 ± 1.0 | 104.5 ± 0.7 | 103.2 ± 2.0 |
| Trimethyllysine | 111.4 ± 0.2 | 100.0 ± 1.0 | 99.6 ± 1.9 | 104.6 ± 2.0 |
| Tryptophan | 99.6 ± 0.3 | 97.5 ± 0.8 | 98.6 ± 1.0 | 105.8 ± 1.8 |
| Tyramine | 98.8 ± 1.2 | 97.1 ± 0.6 | 102.1 ± 1.4 | 106.3 ± 0.8 |
| Tyrosine | 99.2 ± 1.6 | 96.7 ± 0.9 | 97.9 ± 1.9 | 100.0 ± 0.5 |
| Uracil | 105.4 ± 2.0 | 103.9 ± 0.7 | 100.1 ± 0.5 | 99.3 ± 0.9 |
| Uridine 5'-diphosphate | 113.3 ± 1.1 | 98.8 ± 2.0 | 104.9 ± 1.9 | 102.3 ± 0.6 |
| Uridine Monophosphate | 106.6 ± 1.5 | 107.0 ± 1.0 | 86.7 ± 1.4 | 111.8 ± 1.7 |
| Uridine-5-Diphosphogalactose | 91.8 ± 1.8 | 96.3 ± 0.5 | 110.2 ± 1.5 | 111.6 ± 0.7 |
| Urocanic acid | 111.3 ± 1.5 | 112.5 ± 0.6 | 108.5 ± 1.0 | 94.4 ± 1.3 |
| UTP | 98.7 ± 0.6 | 104.7 ± 0.4 | 89.2 ± 0.9 | 96.4 ± 0.6 |
| Valine | 101.3 ± 0.9 | 99.3 ± 1.6 | 105.8 ± 1.4 | 105.5 ± 1.5 |
| XMP | 96.8 ± 1.5 | 103.5 ± 0.3 | 101.1 ± 0.9 | 103.0 ± 1.8 |
| Xylulose-5-Phosphate | 111.8 ± 1.8 | 103.9 ± 0.6 | 100.9 ± 2.0 | 105.9 ± 0.6 |

### **Table S23**| Intra – day accuracy (as determined by % recovery) (Tumor tissue extract as matrix, n = 5). The concentrations of QC_LLOQ_, QC_Low_, QC_Mid_ and QC_High_ were 10 ng/mL, 25 ng/mL, 100 ng/mL and 500 ng/mL respectively.

| Accuracy, % | QC_LLOQ_ | QC_Low_ | QC_Mid_ | QC_High_ |
| --- | --- | --- | --- | --- |
| 1-Methylguanine | 96.4 ± 2.6 | 102.5 ± 1.9 | 94.5 ± 4.5 | 112.0 ± 2.2 |
| 1-Methylhistidine | 93.3 ± 4.2 | 97.8 ± 4.1 | 92.0 ± 5.4 | 102.6 ± 0.7 |
| 1-Methylnicotinamide | 85.5 ± 4.9 | 102.2 ± 5.8 | 90.1 ± 1.4 | 94.3 ± 4.3 |
| 2-Aminoadipic Acid | 91.9 ± 2.9 | 99.8 ± 0.9 | 101.8 ± 0.8 | 88.4 ± 5.9 |
| 2-Deoxyadenosine | 92.5 ± 0.8 | 99.2 ± 3.3 | 97.8 ± 6.1 | 112.5 ± 4.6 |
| 2'-Deoxyadenosine 5'-monophosphate | 94.1 ± 2.0 | 98.3 ± 4.7 | 104.0 ± 4.5 | 98.7 ± 6.3 |
| 2'-deoxyadenosine 5'-triphosphate | 88.7 ± 3.1 | 95.4 ± 4.1 | 105.1 ± 2.0 | 88.1 ± 1.3 |
| 2'-Deoxycytidine-5'-monophosphate | 89.4 ± 4.6 | 94.5 ± 1.8 | 97.0 ± 3.5 | 90.4 ± 4.1 |
| 2-Deoxyguanosine | 89.5 ± 1.5 | 99.2 ± 3.9 | 91.3 ± 3.5 | 98.4 ± 2.1 |
| 2-hydroxybutyric acid | 86.3 ± 2.8 | 90.1 ± 4.7 | 96.7 ± 1.5 | 89.7 ± 4.1 |
| 2-hydroxyglutaric acid | 95.9 ± 2.8 | 97.8 ± 4.3 | 111.1 ± 3.7 | 113.8 ± 2.0 |
| 2-Hydroxyisocaproic acid | 89.6 ± 5.9 | 101.3 ± 0.2 | 104.2 ± 1.2 | 108.8 ± 4.8 |
| 2-ketobutyric acid | 87.3 ± 2.6 | 94.2 ± 2.9 | 111.1 ± 5.2 | 99.2 ± 3.6 |
| 2-oxoadipic acid | 89.2 ± 2.0 | 96.3 ± 0.5 | 99.8 ± 4.9 | 105.8 ± 1.4 |
| 2-Phenylglycine | 94.3 ± 1.4 | 99.7 ± 1.2 | 100.9 ± 3.2 | 95.4 ± 2.4 |
| 2-phosphoglycerate | 91.4 ± 5.6 | 97.0 ± 1.9 | 94.4 ± 0.4 | 101.3 ± 4.7 |
| 3-hydroxy-3-methyl-2-oxobutanoate | 94.3 ± 0.2 | 99.8 ± 0.2 | 100.9 ± 4.9 | 111.4 ± 5.9 |
| 3-hydroxybutyric acid | 91.1 ± 1.3 | 91.5 ± 5.9 | 100.1 ± 5.1 | 97.0 ± 2.2 |
| 3-Methylhistidine | 95.1 ± 2.5 | 98.9 ± 1.5 | 109.6 ± 0.7 | 99.2 ± 2.6 |
| 3-Phosphoglyceraldehyde | 97.5 ± 5.7 | 94.0 ± 5.2 | 107.6 ± 0.5 | 90.6 ± 4.0 |
| 3-phosphoglycerate | 87.8 ± 5.0 | 95.8 ± 5.6 | 90.6 ± 3.0 | 99.8 ± 2.0 |
| 4-amino-butyric acid | 89.4 ± 1.2 | 100.8 ± 2.3 | 95.1 ± 4.0 | 102.1 ± 6.2 |
| 4-aspartyl-phosphate | 87.4 ± 4.4 | 97.1 ± 0.4 | 93.4 ± 4.7 | 111.8 ± 1.0 |
| 4-Hydroxybenzoic acid | 86.8 ± 3.3 | 97.5 ± 1.2 | 99.5 ± 3.0 | 107.8 ± 0.4 |
| 4-Hydroxyphenylacetylglycine | 90.4 ± 4.7 | 93.4 ± 5.0 | 103.0 ± 0.4 | 109.2 ± 0.4 |
| 4-methyl-2-oxopentanoate | 95.5 ± 4.9 | 101.4 ± 6.1 | 109.8 ± 3.6 | 94.1 ± 2.4 |
| 4-Oxoglutaramate | 95.0 ± 2.6 | 100.0 ± 4.2 | 110.5 ± 4.3 | 88.2 ± 5.5 |
| 5-Hydroxyindolepyruvate | 89.1 ± 5.0 | 93.1 ± 5.9 | 108.1 ± 5.2 | 89.2 ± 2.2 |
| 5-Hydroxylysine | 93.0 ± 4.2 | 91.9 ± 4.4 | 90.6 ± 4.6 | 108.9 ± 1.6 |
| 5-Hydroxy-Tryptophan | 85.0 ± 4.5 | 89.5 ± 2.2 | 109.6 ± 0.5 | 93.4 ± 3.9 |
| 5-Phosphoribosylamine | 91.6 ± 2.3 | 101.1 ± 2.2 | 92.7 ± 6.1 | 95.4 ± 4.3 |
| 6-phosphogluconate | 96.3 ± 3.1 | 94.8 ± 4.5 | 103.9 ± 3.1 | 103.7 ± 4.8 |
| Acetylcarnitine | 90.7 ± 2.9 | 88.6 ± 3.4 | 91.8 ± 2.7 | 91.7 ± 6.2 |
| Aconitic Acid | 91.5 ± 0.9 | 99.1 ± 2.9 | 91.0 ± 3.5 | 106.6 ± 1.9 |
| Adenine | 93.2 ± 0.6 | 99.3 ± 0.8 | 98.4 ± 5.0 | 103.5 ± 5.4 |
| Adenosine | 93.0 ± 4.3 | 90.5 ± 0.7 | 90.7 ± 1.9 | 94.3 ± 3.9 |
| Adenosine 3',5'-cyclic monophosphate | 89.1 ± 2.5 | 99.4 ± 2.0 | 92.4 ± 2.2 | 113.3 ± 5.9 |
| Adenosine Monophosphate | 89.0 ± 1.0 | 88.7 ± 0.7 | 100.8 ± 0.4 | 109.4 ± 3.2 |
| Adenosine-5'-diphosphate | 92.4 ± 4.0 | 102.7 ± 4.9 | 95.3 ± 2.7 | 95.3 ± 5.1 |
| Adenosine-5'-triphosphate | 92.4 ± 2.6 | 90.9 ± 6.0 | 99.2 ± 3.9 | 98.4 ± 0.9 |
| Adenylosuccinate | 86.4 ± 6.1 | 92.8 ± 5.9 | 111.4 ± 1.8 | 88.5 ± 3.0 |
| Alanine | 85.6 ± 2.6 | 90.9 ± 1.0 | 111.7 ± 3.9 | 104.0 ± 2.1 |
| alpha-ketoglutaric acid | 87.8 ± 5.8 | 95.7 ± 3.2 | 100.1 ± 3.6 | 101.4 ± 1.7 |
| Anserine | 90.0 ± 1.1 | 99.1 ± 3.7 | 104.2 ± 1.1 | 95.1 ± 1.8 |
| Arginine | 97.7 ± 0.8 | 89.0 ± 2.8 | 95.4 ± 0.6 | 104.0 ± 1.4 |
| Argininosuccinic Acid | 91.2 ± 3.2 | 100.2 ± 2.2 | 109.4 ± 3.5 | 104.7 ± 3.7 |
| Asparagine | 88.7 ± 5.5 | 92.0 ± 2.5 | 105.6 ± 2.3 | 109.0 ± 2.4 |
| Aspartic acid | 90.6 ± 4.3 | 93.5 ± 2.7 | 102.7 ± 4.4 | 99.7 ± 1.6 |
| Asymmetric dimethylarginine | 85.8 ± 6.1 | 101.5 ± 6.2 | 99.5 ± 4.0 | 102.3 ± 1.4 |
| Azelaic acid | 85.6 ± 5.7 | 89.2 ± 1.6 | 91.9 ± 3.5 | 110.8 ± 1.3 |
| beta-Alanine | 89.9 ± 2.6 | 98.2 ± 5.8 | 106.8 ± 3.7 | 112.1 ± 2.8 |
| Betaine | 86.4 ± 1.9 | 98.5 ± 1.0 | 106.7 ± 0.9 | 104.8 ± 4.5 |
| Biopterin | 92.8 ± 3.7 | 98.4 ± 0.2 | 91.5 ± 3.1 | 108.6 ± 5.7 |
| Biotin | 96.1 ± 0.9 | 99.4 ± 4.6 | 93.2 ± 4.5 | 95.5 ± 2.3 |
| Carnitine | 97.1 ± 1.6 | 100.8 ± 1.5 | 99.9 ± 6.0 | 111.7 ± 3.2 |
| Carnosine | 91.2 ± 5.4 | 100.1 ± 2.7 | 110.8 ± 1.6 | 99.3 ± 4.5 |
| CDP-choline | 90.4 ± 4.3 | 96.7 ± 3.9 | 105.6 ± 4.6 | 89.4 ± 5.3 |
| cGMP | 89.7 ± 0.1 | 98.5 ± 1.6 | 90.2 ± 3.9 | 90.1 ± 5.0 |
| Choline | 91.6 ± 6.0 | 91.9 ± 4.1 | 91.3 ± 0.4 | 90.6 ± 0.4 |
| Cinnamic acid | 97.3 ± 3.9 | 93.3 ± 0.9 | 104.6 ± 0.5 | 88.7 ± 4.9 |
| Citraconic acid | 90.6 ± 5.8 | 101.7 ± 0.9 | 112.0 ± 3.5 | 89.9 ± 0.9 |
| Citramalic acid | 94.2 ± 2.7 | 95.5 ± 5.3 | 103.3 ± 0.3 | 103.2 ± 0.2 |
| Citric Acid | 92.0 ± 0.1 | 88.3 ± 4.5 | 102.8 ± 3.5 | 113.3 ± 5.5 |
| Citrulline | 88.6 ± 4.7 | 99.3 ± 4.4 | 109.6 ± 3.3 | 112.1 ± 4.4 |
| Creatine | 97.4 ± 2.7 | 90.5 ± 0.3 | 93.6 ± 2.8 | 100.4 ± 1.7 |
| Creatinine | 96.6 ± 2.9 | 90.5 ± 3.0 | 101.3 ± 5.8 | 94.9 ± 5.8 |
| Cystathionine | 97.4 ± 4.6 | 102.5 ± 1.3 | 101.7 ± 6.1 | 113.3 ± 3.2 |
| Cysteamine | 97.0 ± 0.3 | 97.9 ± 5.2 | 102.4 ± 1.0 | 94.3 ± 1.3 |
| Cysteine | 95.2 ± 4.1 | 95.8 ± 3.2 | 109.0 ± 6.0 | 97.5 ± 6.3 |
| Cysteinylglycine | 89.7 ± 1.7 | 99.5 ± 1.1 | 107.1 ± 0.2 | 100.8 ± 2.8 |
| Cystine | 90.2 ± 1.2 | 90.7 ± 2.9 | 97.9 ± 3.8 | 108.3 ± 1.7 |
| Cytidine | 95.0 ± 3.3 | 89.4 ± 3.9 | 101.4 ± 4.0 | 105.7 ± 4.8 |
| Cytidine 3',5'-cyclic monophosphate | 85.4 ± 0.7 | 97.1 ± 3.5 | 99.3 ± 5.8 | 97.0 ± 2.6 |
| Cytidine 5'-diphosphate | 92.3 ± 2.5 | 101.1 ± 4.8 | 96.5 ± 5.6 | 104.2 ± 2.4 |
| Cytidine 5'-triphosphate | 93.6 ± 0.2 | 97.4 ± 1.2 | 100.2 ± 4.7 | 102.5 ± 4.3 |
| Cytidine monophosphate | 95.1 ± 1.1 | 97.5 ± 1.5 | 107.2 ± 0.4 | 87.1 ± 4.2 |
| Cytosine | 87.8 ± 2.9 | 101.7 ± 5.2 | 91.0 ± 6.2 | 102.9 ± 1.6 |
| dADP | 89.3 ± 2.0 | 90.0 ± 0.2 | 91.3 ± 3.7 | 87.2 ± 3.6 |
| Diethanolamine | 92.6 ± 5.9 | 100.2 ± 0.8 | 100.5 ± 1.7 | 113.3 ± 5.1 |
| Dihydrobiopterin | 86.7 ± 3.6 | 100.5 ± 2.9 | 100.7 ± 1.6 | 101.1 ± 1.7 |
| Dihydroneopterin triphosphate | 86.1 ± 3.6 | 90.7 ± 1.8 | 95.9 ± 6.2 | 106.4 ± 0.6 |
| Dihydroxyacetonephosphate | 85.0 ± 1.1 | 88.7 ± 3.8 | 103.5 ± 2.6 | 100.7 ± 2.1 |
| Dimethylglycine | 85.2 ± 2.1 | 91.1 ± 6.2 | 105.8 ± 1.9 | 95.0 ± 4.8 |
| Ectoine | 93.3 ± 6.2 | 94.9 ± 1.8 | 92.2 ± 5.0 | 88.0 ± 3.9 |
| Erythrose-4-phosphate | 92.8 ± 1.5 | 91.6 ± 6.2 | 94.9 ± 2.9 | 104.8 ± 1.2 |
| FAD | 94.6 ± 0.9 | 92.2 ± 3.3 | 106.5 ± 4.3 | 100.3 ± 1.8 |
| Folic acid | 95.9 ± 5.3 | 99.6 ± 6.0 | 99.8 ± 0.9 | 91.5 ± 5.2 |
| Fructose-6-phosphate | 88.3 ± 5.6 | 101.0 ± 6.1 | 94.8 ± 1.6 | 96.4 ± 2.3 |
| Fumaric Acid | 87.6 ± 4.2 | 91.7 ± 4.6 | 102.9 ± 5.9 | 106.7 ± 0.6 |
| Galactosamine | 88.3 ± 3.9 | 99.6 ± 5.6 | 95.5 ± 3.3 | 109.6 ± 4.3 |
| Galactose-1-phosphate | 93.1 ± 5.0 | 88.3 ± 2.2 | 90.2 ± 5.8 | 105.9 ± 5.4 |
| Galactose-6-Phosphate | 85.2 ± 0.4 | 99.2 ± 3.4 | 95.7 ± 1.7 | 102.6 ± 3.5 |
| gamma-Glutamylcysteine | 85.2 ± 4.8 | 96.8 ± 1.0 | 111.5 ± 2.5 | 95.9 ± 4.8 |
| Glucosamine | 93.9 ± 1.7 | 102.6 ± 5.3 | 103.4 ± 1.5 | 97.8 ± 0.7 |
| Glucosamine-6-Phosphate | 92.3 ± 5.8 | 102.7 ± 3.9 | 96.9 ± 5.1 | 112.4 ± 4.9 |
| Glucose | 93.0 ± 0.2 | 102.9 ± 5.5 | 99.7 ± 2.1 | 92.1 ± 1.9 |
| Glucose-1-phosphate | 96.6 ± 5.6 | 99.0 ± 3.1 | 98.1 ± 3.9 | 93.2 ± 3.0 |
| Glucose-6-Phosphate | 96.0 ± 0.8 | 91.7 ± 0.6 | 111.7 ± 0.4 | 94.8 ± 4.7 |
| Glucuronic acid | 88.4 ± 1.7 | 90.9 ± 5.9 | 110.5 ± 5.8 | 110.0 ± 5.3 |
| Glutamic acid | 86.8 ± 1.5 | 88.4 ± 0.5 | 103.8 ± 0.5 | 96.3 ± 2.0 |
| Glutamine | 90.3 ± 1.8 | 100.1 ± 2.0 | 102.4 ± 4.2 | 94.1 ± 0.1 |
| Glutathione | 93.6 ± 2.2 | 93.3 ± 1.4 | 109.1 ± 5.6 | 113.7 ± 5.0 |
| Glutathionylspermine | 94.4 ± 2.4 | 102.4 ± 5.5 | 104.8 ± 5.5 | 96.8 ± 3.4 |
| Glycerophosphocholine | 97.0 ± 3.4 | 96.0 ± 2.0 | 102.3 ± 0.4 | 96.0 ± 3.6 |
| Glycine | 88.9 ± 5.7 | 102.3 ± 5.5 | 105.4 ± 4.9 | 108.1 ± 2.6 |
| Glycolic acid | 86.4 ± 1.1 | 97.4 ± 1.3 | 91.6 ± 4.8 | 109.8 ± 3.5 |
| Guanidinoacetic acid | 95.2 ± 6.0 | 99.5 ± 3.5 | 110.8 ± 3.5 | 99.5 ± 0.4 |
| Guanidinobutanoic acid | 87.3 ± 4.9 | 101.0 ± 2.1 | 99.0 ± 4.1 | 109.3 ± 4.5 |
| Guanine | 90.8 ± 0.5 | 98.7 ± 0.9 | 111.2 ± 1.5 | 108.1 ± 3.0 |
| Guanosine | 88.1 ± 1.3 | 93.7 ± 1.0 | 97.6 ± 2.2 | 107.0 ± 1.2 |
| Guanosine 5'-diphosphate | 88.5 ± 1.6 | 90.2 ± 5.6 | 94.0 ± 4.2 | 100.9 ± 1.5 |
| Guanosine-5'-monophosphate | 89.8 ± 0.5 | 96.3 ± 3.7 | 105.6 ± 1.3 | 104.9 ± 3.1 |
| Hercynine | 97.3 ± 1.7 | 96.7 ± 4.8 | 93.6 ± 5.4 | 103.9 ± 6.2 |
| Hippuric acid | 91.8 ± 1.9 | 97.0 ± 5.8 | 101.1 ± 1.7 | 111.6 ± 2.8 |
| Histamine | 94.6 ± 4.5 | 96.5 ± 5.3 | 102.5 ± 5.0 | 92.7 ± 1.5 |
| Histidine | 85.5 ± 0.9 | 91.0 ± 0.3 | 92.3 ± 0.5 | 109.9 ± 4.5 |
| Homoalanine | 96.0 ± 6.1 | 100.4 ± 1.8 | 99.2 ± 3.1 | 108.9 ± 1.6 |
| Homoarginine | 89.6 ± 1.9 | 102.3 ± 6.0 | 105.0 ± 3.2 | 103.0 ± 4.3 |
| Homocysteine | 95.3 ± 5.8 | 91.4 ± 2.1 | 108.9 ± 6.1 | 95.5 ± 1.1 |
| Homogentisic acid | 87.4 ± 1.6 | 102.3 ± 3.3 | 103.1 ± 0.7 | 113.3 ± 2.4 |
| Homoleucine | 97.4 ± 0.8 | 100.9 ± 1.9 | 110.1 ± 6.1 | 102.1 ± 2.4 |
| Homovanillic acid | 92.0 ± 1.5 | 102.1 ± 4.4 | 108.5 ± 1.2 | 88.4 ± 2.0 |
| Hydantoin-5-propionate | 91.3 ± 2.6 | 90.4 ± 2.6 | 102.4 ± 5.0 | 113.0 ± 0.3 |
| Hypotaurine | 90.3 ± 5.5 | 89.7 ± 1.7 | 98.3 ± 0.4 | 98.2 ± 4.9 |
| Hypoxanthine | 94.5 ± 5.1 | 96.1 ± 5.4 | 107.9 ± 4.3 | 104.7 ± 3.8 |
| Indole-3-acetamide | 89.1 ± 4.5 | 89.3 ± 1.3 | 110.3 ± 5.5 | 113.5 ± 5.3 |
| Inosine | 92.8 ± 1.0 | 89.1 ± 3.0 | 106.6 ± 0.1 | 95.7 ± 1.6 |
| Inosine 5'-monophosphate | 86.2 ± 5.5 | 93.2 ± 4.4 | 102.4 ± 3.7 | 110.1 ± 0.6 |
| Isocitric Acid | 95.7 ± 3.5 | 99.3 ± 0.7 | 101.5 ± 4.9 | 103.2 ± 1.5 |
| Isoleucine | 97.0 ± 5.9 | 90.1 ± 4.7 | 106.2 ± 0.7 | 113.8 ± 5.3 |
| Lactic Acid | 94.3 ± 4.9 | 98.7 ± 6.1 | 101.9 ± 1.6 | 113.6 ± 1.0 |
| Leucine | 94.3 ± 6.1 | 98.5 ± 2.5 | 105.3 ± 0.7 | 103.8 ± 1.2 |
| Levulinic acid | 89.7 ± 5.0 | 96.4 ± 5.8 | 94.6 ± 0.5 | 104.2 ± 1.5 |
| Lysine | 88.0 ± 4.4 | 99.5 ± 1.2 | 97.6 ± 5.9 | 90.8 ± 0.9 |
| Malic Acid | 97.7 ± 5.8 | 96.8 ± 4.8 | 109.0 ± 4.5 | 106.3 ± 4.3 |
| Mannose-1 with Mannose-6-Phosphate | 96.3 ± 1.3 | 89.6 ± 2.4 | 92.6 ± 5.7 | 91.8 ± 1.9 |
| Mesaconic acid | 85.3 ± 5.0 | 94.8 ± 0.7 | 103.5 ± 5.4 | 89.9 ± 3.6 |
| Methionine | 90.0 ± 2.1 | 94.8 ± 6.1 | 107.5 ± 0.9 | 108.4 ± 5.4 |
| Methionine sulfoxide | 88.1 ± 3.3 | 96.2 ± 5.2 | 108.8 ± 0.8 | 107.1 ± 5.6 |
| Methylmalonic Acid | 90.8 ± 3.9 | 92.1 ± 0.3 | 95.1 ± 1.1 | 93.8 ± 1.2 |
| N1-Acetylspermidine | 97.5 ± 1.3 | 102.2 ± 2.3 | 100.8 ± 3.2 | 110.0 ± 4.0 |
| N1-Acetylspermine | 85.3 ± 3.5 | 91.3 ± 5.6 | 108.9 ± 1.4 | 104.8 ± 5.5 |
| N8-Acetylspermidine | 93.9 ± 5.9 | 102.5 ± 2.8 | 97.6 ± 4.3 | 95.3 ± 5.5 |
| N-Acetyl-Aspartyl-Glutamate | 86.3 ± 5.6 | 89.3 ± 2.2 | 91.6 ± 4.5 | 105.5 ± 2.7 |
| N-Acetyl-Aspartyl-Glutamyl-Glutamate | 85.1 ± 3.4 | 90.5 ± 2.5 | 110.1 ± 2.5 | 89.3 ± 6.2 |
| N-Acetyl-Galactosamine | 90.7 ± 6.0 | 102.1 ± 3.3 | 97.9 ± 1.4 | 97.7 ± 2.8 |
| N-Acetyl-Leucine | 87.1 ± 2.8 | 99.8 ± 1.7 | 91.9 ± 0.9 | 106.5 ± 5.1 |
| N-Acetyl-Methionine | 94.9 ± 1.1 | 101.7 ± 2.3 | 108.4 ± 2.5 | 105.8 ± 4.3 |
| N-Acetylneuraminic acid | 94.1 ± 5.3 | 92.4 ± 5.3 | 111.5 ± 3.7 | 100.2 ± 2.1 |
| N-Acetyl-Proline | 89.5 ± 0.5 | 98.9 ± 4.9 | 111.1 ± 4.5 | 108.2 ± 2.3 |
| N-Acetyl-Putrescine | 96.0 ± 3.8 | 90.3 ± 3.7 | 108.6 ± 2.8 | 87.1 ± 4.5 |
| N-Acetyl-Serine | 92.4 ± 3.5 | 96.1 ± 3.1 | 93.7 ± 3.5 | 113.4 ± 2.8 |
| N-Acetyl-Serotonin | 92.0 ± 6.0 | 93.6 ± 5.5 | 98.3 ± 4.2 | 87.2 ± 1.8 |
| N-Acyl-Homoserine-lactone | 97.8 ± 1.2 | 93.1 ± 4.3 | 100.6 ± 1.2 | 112.2 ± 2.9 |
| NAD | 91.7 ± 1.9 | 91.9 ± 0.7 | 101.7 ± 4.6 | 107.7 ± 0.8 |
| NADH | 88.8 ± 0.4 | 94.3 ± 1.2 | 100.5 ± 0.8 | 101.2 ± 4.6 |
| NADP | 94.0 ± 5.8 | 102.7 ± 5.7 | 92.0 ± 5.1 | 112.8 ± 5.4 |
| NADPH | 85.9 ± 4.6 | 101.2 ± 4.1 | 106.2 ± 1.3 | 98.4 ± 2.9 |
| N-alpha-Acetyl-lysine | 92.2 ± 4.1 | 88.4 ± 1.3 | 104.4 ± 3.5 | 106.8 ± 5.3 |
| N-alpha-Acetyl-Ornithine | 89.9 ± 3.5 | 89.7 ± 2.3 | 105.0 ± 3.2 | 106.4 ± 3.9 |
| N-Carbamoyl-Proline | 88.9 ± 5.9 | 90.9 ± 3.4 | 91.9 ± 6.1 | 114.0 ± 3.8 |
| N-Carbamoyl-Sarcosine | 96.8 ± 1.5 | 101.2 ± 3.3 | 107.2 ± 4.1 | 107.5 ± 5.5 |
| N-Epsilon-Acetyl-lysine | 90.0 ± 5.8 | 88.5 ± 2.5 | 105.5 ± 4.2 | 92.5 ± 0.9 |
| Nicotinamide | 93.2 ± 5.3 | 91.9 ± 2.1 | 96.8 ± 3.7 | 102.6 ± 0.1 |
| N-Methyl-Aspartic Acid | 85.2 ± 2.0 | 94.2 ± 5.5 | 108.0 ± 0.5 | 110.5 ± 1.5 |
| O-Phosphoethanolamine | 92.0 ± 0.7 | 88.5 ± 3.8 | 108.9 ± 3.5 | 89.2 ± 6.3 |
| Ophthalmic acid | 109.8 ± 3.4 | 111.6 ± 2.3 | 113.7 ± 2.8 | 111.7 ± 1.6 |
| Ornithine | 90.8 ± 2.3 | 103.8 ± 0.9 | 113.3 ± 3.6 | 105.2 ± 0.5 |
| Orotidine 5'-monophosphate | 92.6 ± 2.9 | 94.7 ± 0.8 | 86.9 ± 0.6 | 109.8 ± 2.7 |
| Oxaloacetate | 92.1 ± 1.9 | 87.7 ± 1.0 | 101.1 ± 3.5 | 99.5 ± 2.1 |
| Pantothenic acid | 102.8 ± 3.0 | 93.6 ± 2.3 | 87.9 ± 2.1 | 85.9 ± 3.1 |
| Phenylalanine | 93.4 ± 2.5 | 112.0 ± 3.5 | 109.1 ± 1.1 | 102.8 ± 1.8 |
| Phenylpyruvate | 109.0 ± 2.9 | 96.5 ± 1.8 | 107.4 ± 2.1 | 106.5 ± 1.5 |
| Phosphodimethylethanolamine | 108.1 ± 1.2 | 113.3 ± 2.8 | 92.6 ± 0.7 | 103.7 ± 1.7 |
| Phosphoenolpyruvate | 107.4 ± 1.7 | 89.9 ± 2.1 | 100.8 ± 2.0 | 93.4 ± 2.6 |
| Phosphorylcholine | 99.5 ± 2.7 | 101.3 ± 3.9 | 104.7 ± 1.6 | 105.4 ± 2.7 |
| Pipecolic acid | 107.6 ± 1.5 | 112.4 ± 0.6 | 91.8 ± 2.1 | 95.7 ± 1.7 |
| Proline | 107.2 ± 2.6 | 107.0 ± 1.5 | 103.9 ± 1.9 | 96.7 ± 1.1 |
| Putrescine | 87.1 ± 2.2 | 95.1 ± 1.5 | 103.4 ± 2.9 | 111.9 ± 2.9 |
| Pyridoxal | 109.4 ± 0.9 | 102.0 ± 1.1 | 99.2 ± 1.7 | 103.4 ± 2.4 |
| Pyridoxamine | 86.9 ± 3.9 | 86.1 ± 1.8 | 105.0 ± 3.5 | 105.2 ± 1.0 |
| Pyridoxine | 107.7 ± 3.3 | 107.3 ± 2.8 | 104.1 ± 0.5 | 107.4 ± 2.6 |
| Pyroglutamic acid | 99.7 ± 0.5 | 87.4 ± 2.0 | 99.1 ± 3.4 | 112.0 ± 1.8 |
| Pyrrolysine | 114.4 ± 2.5 | 96.0 ± 2.4 | 99.5 ± 1.5 | 103.5 ± 2.7 |
| Pyruvic Acid | 112.6 ± 3.2 | 110.8 ± 0.3 | 114.6 ± 2.7 | 94.9 ± 0.8 |
| Riboflavin | 108.4 ± 3.3 | 95.4 ± 1.3 | 109.0 ± 0.6 | 108.6 ± 3.3 |
| Ribose-5-Phosphate | 102.9 ± 3.4 | 85.3 ± 0.4 | 90.9 ± 1.2 | 96.7 ± 3.9 |
| Ribulose-1-Phosphate | 107.4 ± 0.9 | 86.2 ± 1.1 | 97.9 ± 1.2 | 92.0 ± 2.5 |
| Ribulose-5-Phosphate | 97.1 ± 1.5 | 96.3 ± 1.0 | 97.0 ± 2.2 | 104.0 ± 2.9 |
| Saccharopine | 110.5 ± 1.2 | 93.0 ± 2.3 | 87.1 ± 2.4 | 85.3 ± 1.5 |
| S-Adenosylhomocysteine | 97.6 ± 1.7 | 105.0 ± 0.5 | 93.9 ± 1.5 | 105.8 ± 3.2 |
| S-Adenosylmethioninamine | 90.6 ± 3.6 | 109.4 ± 0.7 | 101.2 ± 0.4 | 114.2 ± 2.2 |
| S-Adenosylmethionine | 108.5 ± 1.3 | 114.7 ± 2.0 | 91.0 ± 3.5 | 107.9 ± 1.6 |
| Sarcosine | 114.0 ± 0.5 | 88.4 ± 2.6 | 105.5 ± 2.2 | 101.5 ± 0.3 |
| Sedoheptulose-7-phosphate | 103.2 ± 3.8 | 98.6 ± 3.5 | 99.0 ± 1.1 | 94.3 ± 3.3 |
| Selenohomocysteine | 103.7 ± 2.4 | 100.8 ± 3.3 | 112.1 ± 1.1 | 95.2 ± 2.0 |
| Selenomethionine | 109.0 ± 2.6 | 100.4 ± 1.9 | 111.8 ± 3.0 | 105.3 ± 2.8 |
| Serine | 98.4 ± 3.7 | 96.3 ± 2.7 | 101.5 ± 3.4 | 105.1 ± 2.5 |
| S-Methyl-Thioinosine | 105.5 ± 3.7 | 102.7 ± 2.5 | 90.1 ± 3.1 | 90.7 ± 0.3 |
| Spermidine | 114.5 ± 2.2 | 114.1 ± 1.2 | 97.0 ± 1.2 | 111.1 ± 0.9 |
| Spermine | 92.2 ± 4.0 | 113.3 ± 0.4 | 90.1 ± 0.9 | 104.1 ± 1.0 |
| Succinate | 112.9 ± 0.7 | 108.3 ± 2.6 | 88.1 ± 3.1 | 97.2 ± 2.3 |
| Sucrose | 99.2 ± 2.2 | 99.4 ± 1.4 | 105.1 ± 2.9 | 88.9 ± 2.8 |
| Symmetric dimethylarginine | 90.3 ± 1.7 | 100.4 ± 0.7 | 93.5 ± 3.1 | 114.7 ± 1.9 |
| Taurine | 113.8 ± 1.3 | 99.0 ± 2.0 | 87.6 ± 2.1 | 114.1 ± 2.7 |
| Tetrahydroneopterin | 99.9 ± 3.8 | 98.9 ± 3.4 | 102.4 ± 0.9 | 103.0 ± 2.4 |
| Thiamine | 98.7 ± 2.3 | 104.8 ± 1.4 | 111.3 ± 2.8 | 111.0 ± 1.4 |
| Thiamine monophosphate | 108.9 ± 1.1 | 92.9 ± 3.2 | 109.7 ± 1.1 | 111.1 ± 1.3 |
| Threonic acid | 105.6 ± 1.5 | 109.2 ± 3.3 | 88.1 ± 0.5 | 104.5 ± 1.0 |
| Threonine | 109.1 ± 2.2 | 111.1 ± 0.6 | 107.4 ± 3.6 | 92.5 ± 0.7 |
| Thymidine | 85.6 ± 0.3 | 114.2 ± 1.6 | 112.5 ± 4.0 | 86.1 ± 3.5 |
| Thymidine Monophosphate | 105.4 ± 0.4 | 106.7 ± 1.8 | 100.9 ± 3.8 | 87.0 ± 0.6 |
| Thymine | 91.4 ± 3.3 | 95.5 ± 2.6 | 111.3 ± 0.6 | 104.0 ± 1.3 |
| Trigonelleine | 97.5 ± 1.7 | 85.3 ± 0.4 | 91.1 ± 2.8 | 108.1 ± 3.0 |
| Trimethylamine N-oxide | 95.7 ± 2.3 | 109.1 ± 0.4 | 103.2 ± 2.0 | 91.4 ± 3.6 |
| Trimethyllysine | 101.3 ± 0.4 | 94.3 ± 3.6 | 89.8 ± 0.5 | 114.4 ± 3.7 |
| Tryptophan | 111.3 ± 1.7 | 108.9 ± 3.8 | 104.9 ± 3.3 | 93.4 ± 0.3 |
| Tyramine | 85.3 ± 3.5 | 98.7 ± 2.9 | 97.4 ± 2.9 | 101.6 ± 3.3 |
| Tyrosine | 104.8 ± 3.2 | 106.1 ± 0.4 | 113.9 ± 1.6 | 88.2 ± 3.5 |
| Uracil | 106.5 ± 1.7 | 110.3 ± 0.6 | 108.7 ± 2.7 | 104.2 ± 1.2 |
| Uridine 5'-diphosphate | 96.3 ± 2.9 | 91.8 ± 0.5 | 93.6 ± 1.6 | 94.4 ± 1.5 |
| Uridine Monophosphate | 103.4 ± 0.3 | 112.7 ± 2.1 | 106.0 ± 2.1 | 100.5 ± 1.4 |
| Uridine-5-Diphosphogalactose | 98.5 ± 1.5 | 95.3 ± 3.7 | 107.4 ± 0.8 | 101.5 ± 2.8 |
| Urocanic acid | 94.1 ± 2.9 | 105.0 ± 2.0 | 99.1 ± 1.4 | 98.6 ± 3.5 |
| UTP | 109.6 ± 3.9 | 110.1 ± 2.5 | 99.7 ± 2.1 | 95.1 ± 3.3 |
| Valine | 102.3 ± 2.2 | 88.0 ± 2.2 | 100.1 ± 3.3 | 85.8 ± 2.2 |
| XMP | 114.5 ± 1.6 | 94.0 ± 3.9 | 112.7 ± 2.5 | 91.5 ± 0.4 |
| Xylulose-5-Phosphate | 111.2 ± 0.4 | 105.7 ± 0.6 | 112.0 ± 1.1 | 105.4 ± 1.3 |

### **Table S24** | Intra – day accuracy (as determined by % recovery) (Plasma extract as matrix, n = 5). The concentrations of QC_LLOQ_, QC_Low_, QC_Mid_ and QC_High_ were 10 ng/mL, 25 ng/mL, 100 ng/mL and 500 ng/mL respectively.

| Accuracy, % | QC_LLOQ_ | QC_Low_ | QC_Mid_ | QC_High_ |
| --- | --- | --- | --- | --- |
| 1-Methylguanine | 96.0 ± 5.5 | 103.9 ± 3.4 | 93.3 ± 0.5 | 85.5 ± 2.9 |
| 1-Methylhistidine | 106.5 ± 3.5 | 92.7 ± 2.2 | 86.4 ± 5.9 | 92.7 ± 0.2 |
| 1-Methylnicotinamide | 99.0 ± 2.4 | 113.9 ± 2.6 | 106.6 ± 4.0 | 107.9 ± 3.9 |
| 2-Aminoadipic Acid | 91.2 ± 1.2 | 108.7 ± 3.3 | 114.6 ± 2.8 | 113.7 ± 2.7 |
| 2-Deoxyadenosine | 103.2 ± 1.5 | 101.0 ± 1.1 | 104.0 ± 2.3 | 110.5 ± 2.3 |
| 2'-Deoxyadenosine 5'-monophosphate | 113.0 ± 4.7 | 94.8 ± 6.1 | 103.0 ± 0.8 | 103.8 ± 6.2 |
| 2'-deoxyadenosine 5'-triphosphate | 88.7 ± 4.8 | 106.0 ± 1.9 | 89.2 ± 2.4 | 89.3 ± 3.3 |
| 2'-Deoxycytidine-5'-monophosphate | 99.3 ± 0.7 | 113.3 ± 5.5 | 113.9 ± 2.4 | 98.3 ± 2.8 |
| 2-Deoxyguanosine | 111.5 ± 2.1 | 107.0 ± 3.9 | 98.7 ± 2.8 | 113.5 ± 0.5 |
| 2-hydroxybutyric acid | 103.9 ± 0.7 | 113.7 ± 4.1 | 89.2 ± 2.7 | 95.0 ± 1.9 |
| 2-hydroxyglutaric acid | 98.0 ± 4.7 | 111.9 ± 5.7 | 94.4 ± 4.6 | 104.8 ± 4.6 |
| 2-Hydroxyisocaproic acid | 89.8 ± 0.3 | 96.8 ± 5.5 | 99.8 ± 0.6 | 114.3 ± 4.6 |
| 2-ketobutyric acid | 87.9 ± 5.9 | 97.9 ± 4.4 | 104.9 ± 2.0 | 109.7 ± 1.5 |
| 2-oxoadipic acid | 114.6 ± 1.4 | 99.1 ± 0.8 | 102.8 ± 0.8 | 108.2 ± 3.0 |
| 2-Phenylglycine | 99.8 ± 2.0 | 93.3 ± 0.3 | 91.7 ± 4.1 | 85.7 ± 5.7 |
| 2-phosphoglycerate | 87.5 ± 1.2 | 91.0 ± 3.5 | 93.7 ± 3.0 | 99.5 ± 4.5 |
| 3-hydroxy-3-methyl-2-oxobutanoate | 91.9 ± 5.7 | 98.2 ± 2.4 | 98.6 ± 5.3 | 91.3 ± 2.4 |
| 3-hydroxybutyric acid | 91.1 ± 2.1 | 107.0 ± 0.8 | 87.7 ± 2.4 | 96.0 ± 1.4 |
| 3-Methylhistidine | 103.8 ± 2.1 | 114.9 ± 2.4 | 91.3 ± 0.4 | 88.8 ± 6.3 |
| 3-Phosphoglyceraldehyde | 98.7 ± 6.1 | 98.3 ± 6.3 | 89.4 ± 1.2 | 112.2 ± 0.7 |
| 3-phosphoglycerate | 100.7 ± 4.1 | 104.7 ± 4.4 | 102.4 ± 1.8 | 99.1 ± 5.7 |
| 4-amino-butyric acid | 85.2 ± 0.3 | 108.4 ± 2.9 | 108.4 ± 2.0 | 106.4 ± 0.9 |
| 4-aspartyl-phosphate | 101.3 ± 0.5 | 87.8 ± 3.5 | 97.4 ± 4.4 | 95.2 ± 2.1 |
| 4-Hydroxybenzoic acid | 85.9 ± 5.3 | 88.3 ± 3.4 | 96.5 ± 1.5 | 92.4 ± 2.5 |
| 4-Hydroxyphenylacetylglycine | 98.7 ± 1.5 | 114.6 ± 1.6 | 93.0 ± 2.7 | 109.2 ± 1.3 |
| 4-methyl-2-oxopentanoate | 87.1 ± 1.5 | 109.5 ± 0.8 | 111.9 ± 5.7 | 99.7 ± 5.7 |
| 4-Oxoglutaramate | 103.9 ± 0.3 | 96.0 ± 3.8 | 104.6 ± 4.6 | 106.1 ± 5.9 |
| 5-Hydroxyindolepyruvate | 108.5 ± 3.5 | 89.8 ± 2.3 | 87.1 ± 1.3 | 88.5 ± 4.3 |
| 5-Hydroxylysine | 95.1 ± 0.6 | 114.4 ± 4.7 | 100.4 ± 3.9 | 97.5 ± 4.7 |
| 5-Hydroxy-Tryptophan | 90.4 ± 1.5 | 108.0 ± 4.2 | 90.4 ± 6.0 | 113.7 ± 2.8 |
| 5-Phosphoribosylamine | 99.5 ± 5.7 | 113.7 ± 1.0 | 103.5 ± 5.4 | 112.9 ± 1.9 |
| 6-phosphogluconate | 97.2 ± 0.4 | 92.1 ± 4.4 | 85.2 ± 0.1 | 100.3 ± 3.4 |
| Acetylcarnitine | 98.8 ± 4.9 | 112.2 ± 3.6 | 105.9 ± 2.1 | 105.9 ± 1.9 |
| Aconitic Acid | 105.6 ± 1.6 | 108.9 ± 3.0 | 96.5 ± 1.9 | 105.4 ± 5.1 |
| Adenine | 103.1 ± 1.4 | 95.2 ± 1.3 | 98.0 ± 4.2 | 87.7 ± 5.4 |
| Adenosine | 99.3 ± 0.7 | 87.9 ± 4.2 | 93.0 ± 4.7 | 85.3 ± 4.2 |
| Adenosine 3',5'-cyclic monophosphate | 110.5 ± 4.3 | 108.4 ± 4.3 | 105.9 ± 1.2 | 100.8 ± 2.7 |
| Adenosine Monophosphate | 102.3 ± 3.4 | 95.7 ± 6.2 | 97.4 ± 6.1 | 106.2 ± 0.2 |
| Adenosine-5'-diphosphate | 99.0 ± 4.4 | 95.8 ± 5.3 | 112.0 ± 0.6 | 86.5 ± 0.1 |
| Adenosine-5'-triphosphate | 89.4 ± 1.7 | 106.2 ± 3.5 | 103.9 ± 3.6 | 108.4 ± 4.9 |
| Adenylosuccinate | 109.5 ± 1.8 | 87.6 ± 4.1 | 114.8 ± 3.9 | 105.1 ± 0.5 |
| Alanine | 90.0 ± 3.2 | 90.4 ± 5.2 | 93.9 ± 3.8 | 94.4 ± 5.9 |
| alpha-ketoglutaric acid | 94.9 ± 3.8 | 107.1 ± 1.0 | 114.0 ± 0.8 | 92.4 ± 5.9 |
| Anserine | 108.2 ± 1.4 | 97.5 ± 2.8 | 107.7 ± 3.3 | 98.9 ± 3.9 |
| Arginine | 99.0 ± 6.1 | 111.7 ± 5.8 | 88.3 ± 2.5 | 112.8 ± 4.6 |
| Argininosuccinic Acid | 87.7 ± 4.8 | 89.9 ± 0.5 | 87.9 ± 1.8 | 99.5 ± 1.3 |
| Asparagine | 89.5 ± 1.2 | 89.3 ± 5.3 | 98.9 ± 3.8 | 97.2 ± 4.2 |
| Aspartic acid | 107.0 ± 4.8 | 100.7 ± 2.4 | 112.1 ± 1.8 | 111.9 ± 2.9 |
| Asymmetric dimethylarginine | 87.9 ± 0.6 | 92.1 ± 4.1 | 87.8 ± 5.3 | 104.2 ± 4.6 |
| Azelaic acid | 111.9 ± 3.1 | 91.7 ± 3.1 | 85.2 ± 1.6 | 106.1 ± 1.7 |
| beta-Alanine | 104.8 ± 2.9 | 104.4 ± 2.2 | 110.5 ± 6.1 | 88.0 ± 2.2 |
| Betaine | 113.2 ± 5.3 | 96.1 ± 3.4 | 99.5 ± 3.1 | 114.1 ± 0.9 |
| Biopterin | 97.0 ± 2.9 | 90.5 ± 4.5 | 100.1 ± 1.1 | 102.8 ± 0.3 |
| Biotin | 99.4 ± 3.3 | 111.1 ± 3.5 | 108.1 ± 3.2 | 91.0 ± 5.9 |
| Carnitine | 102.6 ± 4.8 | 99.0 ± 0.5 | 103.7 ± 2.0 | 113.3 ± 1.6 |
| Carnosine | 90.0 ± 1.5 | 86.7 ± 1.2 | 102.4 ± 5.7 | 108.8 ± 6.2 |
| CDP-choline | 91.4 ± 4.1 | 100.6 ± 5.1 | 99.5 ± 2.7 | 92.8 ± 1.5 |
| cGMP | 104.9 ± 1.9 | 107.9 ± 0.1 | 114.0 ± 3.1 | 88.2 ± 3.3 |
| Choline | 97.5 ± 2.5 | 99.2 ± 5.8 | 113.4 ± 5.7 | 99.5 ± 5.5 |
| Cinnamic acid | 113.0 ± 1.5 | 103.1 ± 1.4 | 99.3 ± 6.3 | 95.3 ± 5.9 |
| Citraconic acid | 90.4 ± 2.6 | 105.7 ± 0.5 | 86.1 ± 4.5 | 97.9 ± 3.0 |
| Citramalic acid | 98.5 ± 1.9 | 99.1 ± 0.2 | 88.7 ± 4.8 | 106.8 ± 6.0 |
| Citric Acid | 108.3 ± 0.2 | 92.2 ± 2.7 | 85.4 ± 2.4 | 97.5 ± 3.3 |
| Citrulline | 114.2 ± 2.6 | 108.9 ± 6.1 | 90.5 ± 3.1 | 105.0 ± 4.8 |
| Creatine | 93.9 ± 2.7 | 87.1 ± 5.7 | 108.5 ± 2.1 | 89.2 ± 2.8 |
| Creatinine | 111.2 ± 1.3 | 105.4 ± 0.5 | 101.7 ± 0.4 | 114.0 ± 4.7 |
| Cystathionine | 114.1 ± 1.0 | 93.9 ± 1.1 | 112.3 ± 2.3 | 102.4 ± 2.9 |
| Cysteamine | 112.8 ± 0.9 | 96.7 ± 0.1 | 92.5 ± 3.9 | 91.4 ± 2.9 |
| Cysteine | 91.6 ± 0.7 | 93.1 ± 1.5 | 96.2 ± 5.8 | 100.5 ± 5.7 |
| Cysteinylglycine | 109.9 ± 4.3 | 93.5 ± 3.8 | 89.1 ± 2.4 | 93.3 ± 4.4 |
| Cystine | 107.7 ± 5.7 | 103.5 ± 1.6 | 108.3 ± 6.3 | 102.4 ± 3.5 |
| Cytidine | 108.0 ± 1.2 | 91.8 ± 0.3 | 86.4 ± 4.8 | 85.6 ± 4.7 |
| Cytidine 3',5'-cyclic monophosphate | 110.3 ± 6.2 | 107.7 ± 4.2 | 96.8 ± 1.1 | 102.0 ± 2.7 |
| Cytidine 5'-diphosphate | 99.9 ± 2.9 | 101.7 ± 5.7 | 109.4 ± 2.2 | 109.2 ± 6.2 |
| Cytidine 5'-triphosphate | 95.2 ± 5.7 | 85.4 ± 2.5 | 107.9 ± 5.5 | 92.4 ± 4.9 |
| Cytidine monophosphate | 89.3 ± 3.7 | 97.8 ± 1.9 | 96.9 ± 5.9 | 99.4 ± 3.1 |
| Cytosine | 89.7 ± 2.1 | 105.5 ± 4.0 | 104.0 ± 5.0 | 100.0 ± 6.1 |
| dADP | 86.2 ± 0.3 | 112.6 ± 4.0 | 86.5 ± 5.1 | 85.9 ± 0.3 |
| Diethanolamine | 89.4 ± 4.4 | 101.1 ± 3.5 | 96.2 ± 0.7 | 108.5 ± 4.2 |
| Dihydrobiopterin | 92.2 ± 1.4 | 104.2 ± 5.7 | 89.9 ± 1.2 | 103.0 ± 5.1 |
| Dihydroneopterin triphosphate | 107.4 ± 3.1 | 103.8 ± 0.4 | 113.3 ± 1.2 | 88.9 ± 2.7 |
| Dihydroxyacetonephosphate | 108.8 ± 6.1 | 95.8 ± 2.8 | 88.0 ± 6.2 | 108.3 ± 1.3 |
| Dimethylglycine | 105.0 ± 5.1 | 95.8 ± 6.1 | 93.1 ± 5.5 | 105.4 ± 3.2 |
| Ectoine | 93.1 ± 1.1 | 102.8 ± 3.4 | 110.0 ± 2.1 | 86.1 ± 6.0 |
| Erythrose-4-phosphate | 97.9 ± 5.6 | 90.3 ± 4.1 | 87.4 ± 4.0 | 96.7 ± 2.2 |
| FAD | 91.4 ± 2.2 | 102.7 ± 6.3 | 109.5 ± 4.4 | 88.2 ± 3.3 |
| Folic acid | 109.2 ± 0.8 | 89.9 ± 5.2 | 103.4 ± 5.3 | 109.9 ± 5.7 |
| Fructose-6-phosphate | 85.8 ± 5.6 | 107.2 ± 2.0 | 102.7 ± 1.2 | 99.1 ± 3.5 |
| Fumaric Acid | 101.1 ± 3.0 | 109.1 ± 3.8 | 93.9 ± 0.1 | 109.5 ± 3.1 |
| Galactosamine | 107.8 ± 1.9 | 101.6 ± 4.8 | 95.9 ± 3.2 | 111.9 ± 5.0 |
| Galactose-1-phosphate | 97.8 ± 2.7 | 113.9 ± 4.4 | 93.8 ± 2.6 | 88.4 ± 6.1 |
| Galactose-6-Phosphate | 85.2 ± 1.3 | 103.6 ± 2.2 | 104.8 ± 5.2 | 89.1 ± 1.6 |
| gamma-Glutamylcysteine | 104.5 ± 0.6 | 92.8 ± 0.2 | 103.2 ± 0.2 | 101.5 ± 1.1 |
| Glucosamine | 98.8 ± 4.1 | 95.6 ± 3.9 | 90.9 ± 5.5 | 105.4 ± 3.0 |
| Glucosamine-6-Phosphate | 105.0 ± 1.8 | 111.1 ± 2.3 | 106.4 ± 3.0 | 113.8 ± 1.3 |
| Glucose | 88.8 ± 0.4 | 102.6 ± 2.5 | 109.7 ± 4.3 | 105.5 ± 3.4 |
| Glucose-1-phosphate | 93.6 ± 3.9 | 92.3 ± 5.9 | 94.1 ± 4.3 | 85.0 ± 3.5 |
| Glucose-6-Phosphate | 87.7 ± 0.7 | 98.4 ± 6.1 | 113.3 ± 3.8 | 100.3 ± 1.0 |
| Glucuronic acid | 93.6 ± 1.5 | 107.1 ± 2.6 | 105.9 ± 5.2 | 87.7 ± 1.9 |
| Glutamic acid | 111.1 ± 3.3 | 100.7 ± 2.5 | 110.5 ± 0.8 | 95.3 ± 3.4 |
| Glutamine | 96.1 ± 5.3 | 90.1 ± 5.8 | 88.7 ± 1.5 | 101.3 ± 3.9 |
| Glutathione | 90.4 ± 1.1 | 88.3 ± 2.1 | 107.2 ± 3.8 | 97.7 ± 1.0 |
| Glutathionylspermine | 91.3 ± 4.1 | 97.1 ± 4.0 | 100.9 ± 3.1 | 111.1 ± 2.5 |
| Glycerophosphocholine | 105.9 ± 1.8 | 113.9 ± 6.0 | 111.1 ± 5.3 | 92.1 ± 6.0 |
| Glycine | 114.4 ± 2.1 | 103.0 ± 6.3 | 86.8 ± 5.8 | 96.3 ± 5.9 |
| Glycolic acid | 94.2 ± 3.0 | 100.6 ± 6.1 | 101.9 ± 3.4 | 111.5 ± 2.4 |
| Guanidinoacetic acid | 92.4 ± 0.2 | 89.8 ± 6.0 | 106.4 ± 2.9 | 113.6 ± 3.0 |
| Guanidinobutanoic acid | 88.4 ± 1.9 | 96.6 ± 2.8 | 91.4 ± 3.0 | 87.2 ± 0.6 |
| Guanine | 92.8 ± 4.1 | 110.4 ± 4.9 | 101.0 ± 3.3 | 101.8 ± 3.1 |
| Guanosine | 106.6 ± 2.7 | 94.9 ± 3.1 | 96.8 ± 5.2 | 112.6 ± 1.8 |
| Guanosine 5'-diphosphate | 97.1 ± 1.0 | 114.2 ± 0.4 | 114.6 ± 4.0 | 91.7 ± 4.4 |
| Guanosine-5'-monophosphate | 97.8 ± 5.0 | 95.3 ± 0.6 | 111.7 ± 2.7 | 97.2 ± 4.7 |
| Hercynine | 94.4 ± 4.3 | 90.8 ± 3.9 | 89.4 ± 2.6 | 89.3 ± 6.3 |
| Hippuric acid | 106.1 ± 5.3 | 113.8 ± 2.8 | 102.6 ± 4.2 | 86.6 ± 2.4 |
| Histamine | 98.5 ± 5.1 | 106.2 ± 0.9 | 106.4 ± 5.7 | 101.7 ± 2.9 |
| Histidine | 107.7 ± 0.5 | 111.3 ± 3.3 | 101.9 ± 1.6 | 104.2 ± 2.9 |
| Homoalanine | 98.8 ± 2.5 | 102.1 ± 3.6 | 90.5 ± 0.5 | 94.9 ± 0.5 |
| Homoarginine | 91.7 ± 1.1 | 95.1 ± 3.6 | 114.6 ± 4.7 | 102.7 ± 4.2 |
| Homocysteine | 95.4 ± 3.1 | 102.1 ± 3.8 | 97.4 ± 2.7 | 88.6 ± 3.1 |
| Homogentisic acid | 114.1 ± 5.3 | 85.9 ± 3.3 | 103.9 ± 4.6 | 104.3 ± 0.6 |
| Homoleucine | 100.8 ± 0.5 | 88.6 ± 2.8 | 103.3 ± 0.3 | 103.0 ± 4.5 |
| Homovanillic acid | 93.2 ± 3.5 | 93.2 ± 2.7 | 99.6 ± 6.1 | 101.3 ± 4.3 |
| Hydantoin-5-propionate | 99.7 ± 4.3 | 110.4 ± 1.3 | 85.1 ± 2.0 | 98.9 ± 5.7 |
| Hypotaurine | 93.2 ± 3.8 | 96.7 ± 1.7 | 87.4 ± 6.1 | 85.2 ± 2.3 |
| Hypoxanthine | 102.4 ± 4.0 | 101.8 ± 3.2 | 91.7 ± 3.4 | 113.9 ± 2.6 |
| Indole-3-acetamide | 103.5 ± 0.2 | 114.2 ± 2.6 | 93.2 ± 2.2 | 94.5 ± 1.2 |
| Inosine | 87.2 ± 6.2 | 89.3 ± 1.3 | 99.2 ± 3.6 | 99.2 ± 3.5 |
| Inosine 5'-monophosphate | 95.5 ± 4.0 | 114.1 ± 0.8 | 100.7 ± 1.2 | 99.4 ± 2.9 |
| Isocitric Acid | 99.5 ± 4.4 | 98.6 ± 2.3 | 101.7 ± 1.8 | 101.0 ± 5.1 |
| Isoleucine | 109.7 ± 2.2 | 96.9 ± 5.8 | 114.3 ± 3.6 | 86.3 ± 2.5 |
| Lactic Acid | 90.7 ± 0.9 | 90.7 ± 5.7 | 90.6 ± 3.4 | 97.6 ± 3.8 |
| Leucine | 85.2 ± 2.6 | 111.8 ± 4.5 | 99.4 ± 5.8 | 94.6 ± 5.3 |
| Levulinic acid | 93.7 ± 3.3 | 91.3 ± 5.6 | 87.4 ± 4.1 | 86.3 ± 5.4 |
| Lysine | 111.8 ± 2.4 | 105.1 ± 5.3 | 101.4 ± 3.7 | 100.1 ± 1.8 |
| Malic Acid | 106.5 ± 3.5 | 104.2 ± 4.4 | 86.7 ± 6.1 | 111.8 ± 5.7 |
| Mannose-1 with Mannose-6-Phosphate | 95.7 ± 3.7 | 110.0 ± 1.6 | 112.4 ± 3.9 | 114.0 ± 4.0 |
| Mesaconic acid | 112.6 ± 5.6 | 91.3 ± 1.5 | 86.9 ± 1.6 | 87.7 ± 5.4 |
| Methionine | 109.6 ± 2.1 | 87.1 ± 3.6 | 108.7 ± 1.8 | 102.8 ± 5.2 |
| Methionine sulfoxide | 106.3 ± 5.7 | 96.5 ± 3.9 | 85.1 ± 5.1 | 91.6 ± 2.6 |
| Methylmalonic Acid | 105.0 ± 2.2 | 103.4 ± 4.3 | 109.9 ± 3.0 | 87.8 ± 0.6 |
| N1-Acetylspermidine | 85.5 ± 0.8 | 96.5 ± 1.2 | 88.6 ± 3.1 | 107.3 ± 3.6 |
| N1-Acetylspermine | 88.6 ± 3.9 | 90.2 ± 3.0 | 102.6 ± 2.3 | 89.0 ± 3.7 |
| N8-Acetylspermidine | 90.8 ± 5.2 | 93.9 ± 4.5 | 93.2 ± 3.1 | 111.2 ± 4.0 |
| N-Acetyl-Aspartyl-Glutamate | 93.0 ± 5.8 | 110.7 ± 4.8 | 110.6 ± 2.1 | 99.4 ± 4.0 |
| N-Acetyl-Aspartyl-Glutamyl-Glutamate | 109.6 ± 6.2 | 94.1 ± 3.8 | 110.2 ± 4.4 | 96.8 ± 4.1 |
| N-Acetyl-Galactosamine | 106.8 ± 4.2 | 95.8 ± 4.0 | 88.8 ± 2.5 | 85.2 ± 3.3 |
| N-Acetyl-Leucine | 94.0 ± 2.4 | 105.5 ± 0.7 | 94.7 ± 4.4 | 88.2 ± 0.5 |
| N-Acetyl-Methionine | 90.5 ± 3.8 | 104.5 ± 3.4 | 98.6 ± 5.4 | 114.7 ± 3.0 |
| N-Acetylneuraminic acid | 93.7 ± 5.9 | 114.1 ± 2.9 | 109.8 ± 5.0 | 90.0 ± 1.1 |
| N-Acetyl-Proline | 85.1 ± 2.7 | 111.5 ± 5.8 | 105.1 ± 4.2 | 114.1 ± 6.2 |
| N-Acetyl-Putrescine | 111.7 ± 3.6 | 106.0 ± 3.8 | 92.4 ± 0.5 | 94.3 ± 1.4 |
| N-Acetyl-Serine | 101.6 ± 1.7 | 114.9 ± 1.7 | 102.3 ± 3.5 | 104.0 ± 3.6 |
| N-Acetyl-Serotonin | 89.9 ± 6.1 | 97.1 ± 6.1 | 96.2 ± 3.9 | 88.1 ± 1.7 |
| N-Acyl-Homoserine-lactone | 107.5 ± 3.2 | 110.5 ± 2.4 | 97.7 ± 3.3 | 102.9 ± 1.9 |
| NAD | 102.1 ± 2.9 | 101.8 ± 5.9 | 95.3 ± 1.5 | 109.3 ± 3.8 |
| NADH | 101.7 ± 6.2 | 106.3 ± 3.8 | 106.1 ± 2.1 | 97.0 ± 6.1 |
| NADP | 106.6 ± 3.6 | 114.0 ± 2.8 | 114.4 ± 3.6 | 98.8 ± 6.1 |
| NADPH | 113.2 ± 6.2 | 113.9 ± 6.0 | 103.0 ± 0.8 | 87.1 ± 1.5 |
| N-alpha-Acetyl-lysine | 114.8 ± 3.2 | 110.2 ± 6.2 | 101.8 ± 4.9 | 85.8 ± 0.9 |
| N-alpha-Acetyl-Ornithine | 104.1 ± 5.6 | 112.4 ± 4.1 | 110.8 ± 1.1 | 107.1 ± 2.0 |
| N-Carbamoyl-Proline | 110.2 ± 5.7 | 93.3 ± 5.2 | 98.0 ± 1.7 | 103.3 ± 4.5 |
| N-Carbamoyl-Sarcosine | 111.7 ± 6.2 | 85.6 ± 0.5 | 114.2 ± 1.4 | 99.4 ± 3.5 |
| N-Epsilon-Acetyl-lysine | 93.4 ± 4.2 | 109.8 ± 1.5 | 112.3 ± 2.4 | 103.3 ± 5.0 |
| Nicotinamide | 103.0 ± 5.2 | 89.3 ± 0.5 | 108.6 ± 0.2 | 102.6 ± 2.9 |
| N-Methyl-Aspartic Acid | 85.8 ± 2.9 | 85.8 ± 4.6 | 93.7 ± 1.3 | 106.9 ± 4.9 |
| O-Phosphoethanolamine | 98.1 ± 3.5 | 100.6 ± 1.3 | 108.8 ± 3.5 | 100.2 ± 2.8 |
| Ophthalmic acid | 96.3 ± 3.3 | 108.9 ± 0.8 | 99.5 ± 2.3 | 95.9 ± 1.3 |
| Ornithine | 110.0 ± 1.1 | 108.7 ± 3.4 | 90.7 ± 0.9 | 107.1 ± 2.3 |
| Orotidine 5'-monophosphate | 97.3 ± 1.2 | 111.5 ± 2.9 | 106.2 ± 2.2 | 97.1 ± 1.8 |
| Oxaloacetate | 90.9 ± 2.2 | 91.6 ± 3.6 | 109.3 ± 3.0 | 91.9 ± 1.1 |
| Pantothenic acid | 113.4 ± 1.3 | 87.1 ± 1.5 | 113.6 ± 3.1 | 98.3 ± 3.8 |
| Phenylalanine | 107.1 ± 2.4 | 100.1 ± 1.5 | 103.2 ± 2.1 | 92.9 ± 2.1 |
| Phenylpyruvate | 107.6 ± 4.0 | 97.7 ± 0.4 | 110.4 ± 2.3 | 109.6 ± 1.5 |
| Phosphodimethylethanolamine | 106.1 ± 0.4 | 106.7 ± 3.8 | 101.9 ± 3.1 | 103.7 ± 3.8 |
| Phosphoenolpyruvate | 95.0 ± 3.1 | 107.6 ± 1.2 | 113.6 ± 1.9 | 104.9 ± 3.9 |
| Phosphorylcholine | 98.5 ± 1.2 | 85.8 ± 2.7 | 98.1 ± 0.9 | 110.4 ± 2.5 |
| Pipecolic acid | 85.5 ± 2.6 | 98.3 ± 3.3 | 90.2 ± 0.7 | 99.4 ± 2.8 |
| Proline | 110.3 ± 0.4 | 91.1 ± 3.9 | 98.4 ± 2.8 | 102.1 ± 1.5 |
| Putrescine | 113.3 ± 1.4 | 90.0 ± 3.0 | 98.6 ± 3.5 | 93.3 ± 3.8 |
| Pyridoxal | 90.1 ± 1.8 | 113.0 ± 3.8 | 94.7 ± 3.6 | 89.5 ± 1.4 |
| Pyridoxamine | 102.4 ± 3.3 | 112.8 ± 3.4 | 107.2 ± 1.1 | 93.7 ± 0.6 |
| Pyridoxine | 92.0 ± 3.8 | 91.5 ± 3.2 | 91.5 ± 1.1 | 109.1 ± 3.5 |
| Pyroglutamic acid | 103.4 ± 0.6 | 96.9 ± 3.9 | 114.9 ± 1.3 | 96.6 ± 0.5 |
| Pyrrolysine | 99.4 ± 0.8 | 94.8 ± 1.1 | 86.9 ± 3.0 | 107.4 ± 1.8 |
| Pyruvic Acid | 104.2 ± 2.2 | 92.2 ± 2.0 | 106.8 ± 2.2 | 91.3 ± 1.0 |
| Riboflavin | 93.9 ± 4.0 | 100.0 ± 3.1 | 106.6 ± 3.9 | 103.3 ± 1.4 |
| Ribose-5-Phosphate | 114.3 ± 0.8 | 88.5 ± 1.3 | 95.5 ± 0.7 | 103.1 ± 1.8 |
| Ribulose-1-Phosphate | 93.4 ± 3.5 | 97.6 ± 3.0 | 113.7 ± 2.8 | 107.3 ± 1.6 |
| Ribulose-5-Phosphate | 96.9 ± 0.7 | 106.0 ± 1.8 | 108.5 ± 1.7 | 99.7 ± 2.1 |
| Saccharopine | 102.8 ± 1.4 | 92.9 ± 2.0 | 96.6 ± 0.8 | 110.8 ± 3.3 |
| S-Adenosylhomocysteine | 86.1 ± 2.1 | 105.8 ± 3.5 | 108.7 ± 2.9 | 92.6 ± 0.4 |
| S-Adenosylmethioninamine | 95.9 ± 1.9 | 95.6 ± 1.0 | 90.9 ± 4.0 | 91.1 ± 3.5 |
| S-Adenosylmethionine | 97.1 ± 0.6 | 114.0 ± 1.5 | 91.8 ± 2.1 | 110.9 ± 0.7 |
| Sarcosine | 91.9 ± 2.4 | 92.0 ± 1.8 | 88.1 ± 1.5 | 114.9 ± 2.7 |
| Sedoheptulose-7-phosphate | 108.7 ± 1.1 | 92.3 ± 0.7 | 94.8 ± 1.1 | 103.8 ± 0.5 |
| Selenohomocysteine | 85.8 ± 0.4 | 95.4 ± 1.3 | 100.9 ± 0.5 | 104.1 ± 3.3 |
| Selenomethionine | 89.4 ± 1.2 | 87.0 ± 1.0 | 88.4 ± 0.8 | 108.5 ± 1.4 |
| Serine | 105.0 ± 1.9 | 91.1 ± 3.4 | 95.8 ± 2.2 | 114.6 ± 1.2 |
| S-Methyl-Thioinosine | 95.6 ± 0.3 | 109.7 ± 2.3 | 88.3 ± 1.7 | 95.5 ± 1.3 |
| Spermidine | 112.9 ± 2.4 | 90.4 ± 2.5 | 111.1 ± 3.9 | 111.0 ± 0.8 |
| Spermine | 90.0 ± 2.3 | 87.1 ± 3.7 | 102.1 ± 3.3 | 104.8 ± 1.4 |
| Succinate | 113.2 ± 3.0 | 114.9 ± 1.0 | 106.0 ± 4.0 | 95.4 ± 1.2 |
| Sucrose | 111.7 ± 2.3 | 109.6 ± 2.2 | 106.1 ± 0.4 | 96.5 ± 2.1 |
| Symmetric dimethylarginine | 111.3 ± 0.5 | 104.2 ± 1.7 | 90.5 ± 1.5 | 93.9 ± 1.6 |
| Taurine | 109.8 ± 2.3 | 97.2 ± 2.4 | 88.6 ± 1.4 | 98.0 ± 2.3 |
| Tetrahydroneopterin | 107.4 ± 0.5 | 101.5 ± 1.9 | 109.3 ± 3.3 | 94.2 ± 2.4 |
| Thiamine | 102.9 ± 3.6 | 95.4 ± 1.9 | 88.6 ± 3.8 | 110.4 ± 1.8 |
| Thiamine monophosphate | 110.2 ± 1.5 | 86.7 ± 3.3 | 98.0 ± 2.0 | 112.6 ± 0.6 |
| Threonic acid | 90.6 ± 2.2 | 113.1 ± 0.8 | 108.2 ± 3.0 | 109.5 ± 3.0 |
| Threonine | 93.5 ± 0.4 | 110.2 ± 0.6 | 100.7 ± 3.8 | 94.8 ± 0.4 |
| Thymidine | 98.6 ± 2.7 | 86.8 ± 0.5 | 90.8 ± 2.4 | 85.8 ± 2.4 |
| Thymidine Monophosphate | 114.8 ± 0.4 | 94.8 ± 0.6 | 102.2 ± 3.9 | 89.3 ± 3.5 |
| Thymine | 89.9 ± 3.8 | 92.6 ± 3.6 | 90.7 ± 1.2 | 92.4 ± 2.3 |
| Trigonelleine | 93.3 ± 2.9 | 101.8 ± 3.6 | 98.4 ± 1.4 | 86.6 ± 3.6 |
| Trimethylamine N-oxide | 103.1 ± 1.3 | 104.3 ± 1.1 | 105.2 ± 0.4 | 113.0 ± 2.7 |
| Trimethyllysine | 87.7 ± 3.9 | 95.5 ± 0.9 | 92.1 ± 2.2 | 89.5 ± 0.3 |
| Tryptophan | 107.3 ± 1.6 | 92.8 ± 3.0 | 95.1 ± 2.9 | 109.2 ± 1.5 |
| Tyramine | 102.7 ± 1.5 | 92.3 ± 0.5 | 97.9 ± 2.0 | 105.4 ± 3.8 |
| Tyrosine | 87.2 ± 3.9 | 111.8 ± 0.7 | 97.0 ± 3.3 | 105.9 ± 3.8 |
| Uracil | 87.2 ± 0.4 | 99.2 ± 1.2 | 106.5 ± 2.4 | 95.4 ± 2.7 |
| Uridine 5'-diphosphate | 112.1 ± 3.3 | 91.3 ± 1.6 | 87.5 ± 3.1 | 106.7 ± 2.1 |
| Uridine Monophosphate | 106.8 ± 3.1 | 89.8 ± 3.2 | 111.4 ± 3.8 | 105.1 ± 1.1 |
| Uridine-5-Diphosphogalactose | 112.9 ± 1.3 | 111.6 ± 2.5 | 111.1 ± 3.4 | 109.1 ± 1.5 |
| Urocanic acid | 111.0 ± 1.8 | 87.6 ± 2.2 | 91.8 ± 3.3 | 97.6 ± 0.6 |
| UTP | 107.7 ± 3.6 | 106.7 ± 1.3 | 108.2 ± 0.3 | 87.3 ± 3.9 |
| Valine | 113.4 ± 2.1 | 104.2 ± 0.3 | 107.7 ± 1.7 | 109.1 ± 2.5 |
| XMP | 100.2 ± 0.8 | 94.3 ± 1.4 | 97.3 ± 1.1 | 85.8 ± 0.3 |
| Xylulose-5-Phosphate | 109.2 ± 2.7 | 89.4 ± 3.8 | 101.6 ± 1.3 | 114.1 ± 3.3 |

### **Table S25** | Intra – day accuracy (as determined by % recovery) (Cell extract as matrix, n = 5). The concentrations of QC_LLOQ_, QC_Low_, QC_Mid_ and QC_High_ were 10 ng/mL, 25 ng/mL, 100 ng/mL and 500 ng/mL respectively.

| Accuracy, % | QC_LLOQ_ | QC_Low_ | QC_Mid_ | QC_High_ |
| --- | --- | --- | --- | --- |
| 1-Methylguanine | 105.6 ± 5.3 | 103.2 ± 0.3 | 101.3 ± 1.6 | 106.6 ± 4.8 |
| 1-Methylhistidine | 103.4 ± 2.6 | 106.7 ± 2.0 | 105.4 ± 6.1 | 108.7 ± 2.9 |
| 1-Methylnicotinamide | 106.7 ± 5.1 | 103.2 ± 2.5 | 100.6 ± 4.2 | 104.5 ± 2.8 |
| 2-Aminoadipic Acid | 104.5 ± 2.7 | 103.4 ± 0.9 | 101.4 ± 5.6 | 98.6 ± 1.2 |
| 2-Deoxyadenosine | 98.7 ± 2.1 | 103.2 ± 0.9 | 101.4 ± 1.0 | 105.6 ± 1.9 |
| 2'-Deoxyadenosine 5'-monophosphate | 97.7 ± 0.4 | 98.8 ± 1.7 | 103.4 ± 0.9 | 102.1 ± 2.4 |
| 2'-deoxyadenosine 5'-triphosphate | 99.6 ± 0.3 | 104.5 ± 1.9 | 102.3 ± 4.0 | 102.1 ± 4.8 |
| 2'-Deoxycytidine-5'-monophosphate | 103.4 ± 0.6 | 98.9 ± 3.9 | 101.2 ± 4.7 | 100.8 ± 0.4 |
| 2-Deoxyguanosine | 97.7 ± 1.3 | 105.6 ± 2.3 | 103.2 ± 4.1 | 108.8 ± 4.4 |
| 2-hydroxybutyric acid | 105.6 ± 2.2 | 98.5 ± 2.1 | 106.7 ± 4.4 | 100.3 ± 1.3 |
| 2-hydroxyglutaric acid | 104.5 ± 4.3 | 106.7 ± 1.5 | 96.6 ± 4.1 | 98.8 ± 3.1 |
| 2-Hydroxyisocaproic acid | 100.2 ± 5.3 | 98.8 ± 0.9 | 97.7 ± 1.8 | 103.4 ± 1.2 |
| 2-ketobutyric acid | 105.6 ± 5.3 | 107.7 ± 5.0 | 103.2 ± 5.7 | 97.7 ± 3.6 |
| 2-oxoadipic acid | 98.8 ± 3.3 | 103.2 ± 1.0 | 100.6 ± 3.3 | 105.6 ± 4.4 |
| 2-Phenylglycine | 96.7 ± 1.4 | 103.2 ± 4.5 | 100.7 ± 1.7 | 108.8 ± 3.1 |
| 2-phosphoglycerate | 98.8 ± 3.6 | 97.7 ± 3.9 | 103.2 ± 4.2 | 101.4 ± 0.5 |
| 3-hydroxy-3-methyl-2-oxobutanoate | 100.8 ± 5.0 | 103.6 ± 1.6 | 105.6 ± 0.6 | 105.2 ± 3.7 |
| 3-hydroxybutyric acid | 103.2 ± 0.7 | 106.7 ± 5.0 | 98.8 ± 3.3 | 103.2 ± 1.0 |
| 3-Methylhistidine | 105.6 ± 1.1 | 98.8 ± 1.5 | 100.9 ± 5.6 | 103.4 ± 2.6 |
| 3-Phosphoglyceraldehyde | 105.6 ± 3.4 | 103.4 ± 3.5 | 108.8 ± 2.0 | 98.8 ± 1.3 |
| 3-phosphoglycerate | 96.6 ± 3.0 | 98.8 ± 1.0 | 100.4 ± 1.9 | 103.2 ± 2.2 |
| 4-amino-butyric acid | 98.6 ± 3.9 | 99.5 ± 5.8 | 103.2 ± 3.0 | 100.1 ± 3.4 |
| 4-aspartyl-phosphate | 100.5 ± 5.7 | 98.6 ± 1.7 | 99.3 ± 5.2 | 106.6 ± 5.6 |
| 4-Hydroxybenzoic acid | 97.8 ± 1.7 | 99.5 ± 3.9 | 103.2 ± 3.7 | 101.3 ± 5.4 |
| 4-Hydroxyphenylacetylglycine | 104.5 ± 3.6 | 98.8 ± 5.1 | 99.3 ± 1.4 | 96.7 ± 4.3 |
| 4-methyl-2-oxopentanoate | 106.7 ± 2.3 | 105.6 ± 1.3 | 108.9 ± 0.7 | 98.8 ± 0.9 |
| 4-Oxoglutaramate | 104.5 ± 5.5 | 98.7 ± 1.9 | 95.6 ± 3.4 | 97.7 ± 4.1 |
| 5-Hydroxyindolepyruvate | 98.9 ± 3.1 | 103.4 ± 0.9 | 101.2 ± 0.1 | 98.8 ± 5.5 |
| 5-Hydroxylysine | 98.8 ± 1.2 | 103.2 ± 1.4 | 101.2 ± 1.2 | 100.9 ± 4.4 |
| 5-Hydroxy-Tryptophan | 95.4 ± 4.9 | 98.7 ± 0.5 | 101.2 ± 2.0 | 105.6 ± 2.5 |
| 5-Phosphoribosylamine | 106.7 ± 4.7 | 103.2 ± 4.3 | 98.8 ± 2.7 | 97.6 ± 4.2 |
| 6-phosphogluconate | 94.6 ± 5.0 | 103.2 ± 5.5 | 101.5 ± 1.3 | 100.6 ± 4.7 |
| Acetylcarnitine | 108.4 ± 2.9 | 97.7 ± 0.4 | 96.6 ± 5.4 | 95.4 ± 2.4 |
| Aconitic Acid | 100.6 ± 4.7 | 104.5 ± 1.8 | 98.8 ± 3.2 | 96.7 ± 6.2 |
| Adenine | 108.9 ± 0.6 | 103.2 ± 0.7 | 97.6 ± 3.4 | 95.4 ± 4.5 |
| Adenosine | 103.2 ± 5.6 | 100.8 ± 3.5 | 98.7 ± 3.5 | 95.6 ± 3.1 |
| Adenosine 3',5'-cyclic monophosphate | 97.8 ± 4.1 | 103.2 ± 5.0 | 101.3 ± 4.3 | 98.5 ± 5.0 |
| Adenosine Monophosphate | 103.4 ± 0.4 | 102.5 ± 2.2 | 99.7 ± 2.0 | 97.8 ± 2.3 |
| Adenosine-5'-diphosphate | 98.8 ± 1.6 | 103.2 ± 6.1 | 104.5 ± 2.9 | 107.7 ± 0.5 |
| Adenosine-5'-triphosphate | 105.6 ± 2.6 | 103.2 ± 4.0 | 104.5 ± 3.3 | 106.7 ± 1.3 |
| Adenylosuccinate | 98.8 ± 0.2 | 96.7 ± 4.1 | 103.2 ± 5.4 | 104.5 ± 3.9 |
| Alanine | 99.5 ± 0.3 | 103.2 ± 2.6 | 100.8 ± 3.7 | 98.6 ± 3.0 |
| alpha-ketoglutaric acid | 99.4 ± 2.6 | 103.2 ± 2.7 | 101.6 ± 3.1 | 100.9 ± 5.7 |
| Anserine | 97.7 ± 4.1 | 103.2 ± 2.2 | 104.5 ± 0.6 | 100.9 ± 4.5 |
| Arginine | 96.5 ± 5.3 | 98.2 ± 1.5 | 95.0 ± 5.8 | 103.1 ± 1.1 |
| Argininosuccinic Acid | 104.0 ± 4.7 | 98.3 ± 1.6 | 97.7 ± 4.8 | 96.5 ± 1.3 |
| Asparagine | 98.6 ± 0.4 | 96.5 ± 6.1 | 103.2 ± 1.6 | 101.5 ± 2.5 |
| Aspartic acid | 104.3 ± 2.1 | 102.1 ± 3.2 | 98.4 ± 5.1 | 97.6 ± 1.2 |
| Asymmetric dimethylarginine | 99.5 ± 4.2 | 103.2 ± 0.3 | 101.2 ± 5.3 | 100.7 ± 3.8 |
| Azelaic acid | 100.8 ± 2.9 | 104.5 ± 2.9 | 108.6 ± 2.5 | 106.5 ± 1.6 |
| beta-Alanine | 98.7 ± 5.3 | 99.5 ± 3.4 | 103.4 ± 0.7 | 101.2 ± 1.4 |
| Betaine | 97.6 ± 4.0 | 103.2 ± 3.6 | 101.3 ± 3.4 | 96.6 ± 3.5 |
| Biopterin | 98.6 ± 4.5 | 103.2 ± 3.0 | 105.4 ± 1.6 | 102.0 ± 6.0 |
| Biotin | 106.7 ± 0.7 | 103.2 ± 3.2 | 104.1 ± 1.2 | 98.8 ± 4.1 |
| Carnitine | 98.3 ± 3.0 | 95.6 ± 2.3 | 103.4 ± 4.1 | 98.8 ± 5.1 |
| Carnosine | 99.4 ± 2.2 | 103.2 ± 0.8 | 101.5 ± 4.0 | 108.9 ± 2.2 |
| CDP-choline | 105.6 ± 1.2 | 103.2 ± 4.8 | 97.9 ± 4.6 | 96.8 ± 1.3 |
| cGMP | 99.8 ± 1.7 | 103.4 ± 0.9 | 98.7 ± 2.0 | 105.6 ± 2.2 |
| Choline | 103.4 ± 4.7 | 105.6 ± 4.5 | 106.7 ± 2.4 | 102.1 ± 3.4 |
| Cinnamic acid | 98.8 ± 4.7 | 96.0 ± 1.9 | 99.6 ± 4.0 | 103.4 ± 5.3 |
| Citraconic acid | 102.1 ± 1.6 | 106.4 ± 3.0 | 98.8 ± 5.9 | 104.5 ± 2.5 |
| Citramalic acid | 97.5 ± 0.6 | 99.7 ± 2.8 | 103.2 ± 2.6 | 104.6 ± 2.7 |
| Citric Acid | 104.2 ± 0.9 | 98.8 ± 0.9 | 99.5 ± 3.4 | 102.1 ± 3.2 |
| Citrulline | 97.7 ± 5.4 | 96.6 ± 0.2 | 102.3 ± 0.9 | 100.8 ± 4.8 |
| Creatine | 98.8 ± 0.7 | 99.4 ± 4.7 | 102.3 ± 4.7 | 100.5 ± 3.0 |
| Creatinine | 100.7 ± 4.3 | 103.4 ± 4.6 | 97.7 ± 4.6 | 96.5 ± 6.0 |
| Cystathionine | 98.8 ± 6.3 | 102.1 ± 2.3 | 106.6 ± 4.5 | 107.2 ± 5.2 |
| Cysteamine | 103.4 ± 1.5 | 100.8 ± 0.5 | 98.8 ± 5.1 | 95.6 ± 2.8 |
| Cysteine | 97.7 ± 1.3 | 95.1 ± 0.9 | 103.4 ± 3.9 | 102.1 ± 2.8 |
| Cysteinylglycine | 100.9 ± 4.5 | 106.5 ± 3.8 | 104.3 ± 0.4 | 109.6 ± 6.2 |
| Cystine | 106.6 ± 2.5 | 104.3 ± 2.6 | 105.6 ± 6.0 | 100.8 ± 5.6 |
| Cytidine | 98.7 ± 0.3 | 99.4 ± 5.3 | 103.2 ± 4.8 | 97.7 ± 4.4 |
| Cytidine 3',5'-cyclic monophosphate | 97.7 ± 5.0 | 95.6 ± 3.0 | 98.1 ± 6.1 | 100.3 ± 2.5 |
| Cytidine 5'-diphosphate | 98.8 ± 2.2 | 103.2 ± 5.3 | 101.3 ± 2.1 | 97.7 ± 5.6 |
| Cytidine 5'-triphosphate | 96.6 ± 4.6 | 103.2 ± 1.1 | 101.4 ± 3.7 | 95.8 ± 3.7 |
| Cytidine monophosphate | 99.3 ± 5.4 | 103.2 ± 3.8 | 101.4 ± 3.9 | 100.8 ± 3.2 |
| Cytosine | 100.6 ± 6.2 | 98.8 ± 0.3 | 103.2 ± 4.0 | 105.6 ± 2.5 |
| dADP | 108.8 ± 4.5 | 99.8 ± 5.4 | 103.2 ± 2.3 | 100.4 ± 6.1 |
| Diethanolamine | 105.6 ± 5.5 | 103.2 ± 3.7 | 101.2 ± 0.3 | 98.8 ± 2.9 |
| Dihydrobiopterin | 104.5 ± 4.7 | 102.1 ± 4.1 | 98.9 ± 3.2 | 106.6 ± 0.5 |
| Dihydroneopterin triphosphate | 105.6 ± 2.7 | 100.8 ± 2.3 | 98.9 ± 4.9 | 99.6 ± 4.9 |
| Dihydroxyacetonephosphate | 100.4 ± 5.7 | 98.8 ± 0.9 | 103.2 ± 2.0 | 96.7 ± 3.2 |
| Dimethylglycine | 98.7 ± 2.3 | 103.2 ± 3.2 | 101.4 ± 5.7 | 100.8 ± 6.0 |
| Ectoine | 98.8 ± 6.0 | 104.3 ± 4.7 | 105.3 ± 0.7 | 100.9 ± 5.2 |
| Erythrose-4-phosphate | 97.7 ± 1.6 | 99.0 ± 6.2 | 95.6 ± 0.9 | 101.3 ± 5.4 |
| FAD | 102.4 ± 0.4 | 101.5 ± 4.7 | 98.0 ± 4.5 | 97.6 ± 1.0 |
| Folic acid | 98.8 ± 1.6 | 99.5 ± 6.3 | 101.3 ± 4.9 | 100.7 ± 5.5 |
| Fructose-6-phosphate | 99.6 ± 3.0 | 103.4 ± 0.6 | 102.1 ± 4.1 | 105.6 ± 3.4 |
| Fumaric Acid | 98.8 ± 2.5 | 103.2 ± 3.5 | 102.0 ± 1.0 | 105.6 ± 1.3 |
| Galactosamine | 104.5 ± 4.8 | 98.8 ± 4.1 | 99.4 ± 0.4 | 103.2 ± 3.3 |
| Galactose-1-phosphate | 105.6 ± 3.3 | 103.4 ± 2.0 | 102.1 ± 0.8 | 98.8 ± 1.6 |
| Galactose-6-Phosphate | 103.2 ± 1.9 | 97.8 ± 1.0 | 96.7 ± 1.0 | 99.5 ± 6.1 |
| gamma-Glutamylcysteine | 99.0 ± 5.3 | 103.2 ± 1.6 | 102.1 ± 5.9 | 101.6 ± 4.2 |
| Glucosamine | 105.6 ± 1.6 | 103.2 ± 3.6 | 98.8 ± 1.0 | 103.2 ± 0.9 |
| Glucosamine-6-Phosphate | 97.7 ± 4.8 | 98.0 ± 3.8 | 99.4 ± 1.6 | 103.2 ± 0.3 |
| Glucose | 99.8 ± 2.9 | 103.2 ± 3.1 | 101.4 ± 4.1 | 105.6 ± 5.0 |
| Glucose-1-phosphate | 106.7 ± 4.2 | 105.4 ± 3.3 | 103.2 ± 2.5 | 98.8 ± 5.1 |
| Glucose-6-Phosphate | 97.6 ± 2.4 | 103.2 ± 6.2 | 101.2 ± 1.4 | 105.6 ± 0.2 |
| Glucuronic acid | 106.1 ± 0.4 | 105.4 ± 2.2 | 98.8 ± 5.8 | 97.7 ± 4.3 |
| Glutamic acid | 96.6 ± 4.8 | 98.8 ± 1.6 | 103.2 ± 2.6 | 104.5 ± 0.2 |
| Glutamine | 102.1 ± 1.8 | 103.5 ± 2.1 | 100.8 ± 1.2 | 99.5 ± 4.7 |
| Glutathione | 97.7 ± 3.9 | 95.6 ± 0.1 | 103.2 ± 5.8 | 106.7 ± 2.2 |
| Glutathionylspermine | 105.6 ± 4.1 | 107.7 ± 4.7 | 104.3 ± 3.8 | 99.6 ± 4.9 |
| Glycerophosphocholine | 99.8 ± 1.8 | 103.2 ± 5.3 | 101.4 ± 5.3 | 105.6 ± 5.3 |
| Glycine | 100.4 ± 1.0 | 98.8 ± 4.4 | 96.5 ± 1.3 | 95.4 ± 5.2 |
| Glycolic acid | 95.7 ± 5.9 | 103.2 ± 2.5 | 101.4 ± 3.6 | 100.8 ± 5.1 |
| Guanidinoacetic acid | 98.9 ± 5.0 | 103.2 ± 5.6 | 101.4 ± 4.8 | 97.6 ± 1.0 |
| Guanidinobutanoic acid | 100.9 ± 1.3 | 97.7 ± 5.9 | 103.2 ± 5.1 | 97.8 ± 1.6 |
| Guanine | 98.8 ± 3.6 | 103.2 ± 4.7 | 100.4 ± 3.2 | 99.5 ± 0.5 |
| Guanosine | 105.6 ± 2.8 | 103.2 ± 5.2 | 98.8 ± 0.4 | 97.7 ± 6.0 |
| Guanosine 5'-diphosphate | 102.4 ± 4.5 | 100.6 ± 0.8 | 99.6 ± 2.2 | 96.7 ± 4.3 |
| Guanosine-5'-monophosphate | 104.5 ± 3.0 | 102.1 ± 6.1 | 100.9 ± 2.8 | 96.7 ± 1.2 |
| Hercynine | 108.9 ± 4.5 | 103.2 ± 2.1 | 97.6 ± 3.9 | 96.7 ± 3.4 |
| Hippuric acid | 103.2 ± 0.4 | 98.8 ± 2.0 | 96.7 ± 0.9 | 95.0 ± 0.8 |
| Histamine | 104.1 ± 4.2 | 102.0 ± 4.7 | 96.8 ± 2.1 | 95.9 ± 3.7 |
| Histidine | 104.3 ± 5.5 | 102.1 ± 4.1 | 99.4 ± 1.0 | 102.3 ± 3.8 |
| Homoalanine | 104.1 ± 6.2 | 105.6 ± 3.8 | 108.9 ± 1.0 | 99.4 ± 4.1 |
| Homoarginine | 99.2 ± 3.7 | 105.6 ± 5.7 | 106.9 ± 4.7 | 103.2 ± 0.8 |
| Homocysteine | 106.7 ± 5.6 | 98.7 ± 1.1 | 95.6 ± 3.2 | 97.9 ± 5.3 |
| Homogentisic acid | 98.0 ± 2.8 | 103.2 ± 5.0 | 102.4 ± 2.5 | 101.6 ± 4.5 |
| Homoleucine | 109.6 ± 5.2 | 106.5 ± 5.7 | 107.8 ± 6.0 | 105.4 ± 2.6 |
| Homovanillic acid | 108.2 ± 4.6 | 105.6 ± 1.8 | 103.2 ± 2.5 | 98.8 ± 0.5 |
| Hydantoin-5-propionate | 97.5 ± 0.8 | 99.6 ± 4.4 | 105.5 ± 4.6 | 103.2 ± 0.6 |
| Hypotaurine | 101.7 ± 0.4 | 107.2 ± 2.9 | 98.2 ± 3.8 | 113.0 ± 1.0 |
| Hypoxanthine | 112.7 ± 3.2 | 113.5 ± 3.3 | 100.0 ± 2.3 | 89.6 ± 2.9 |
| Indole-3-acetamide | 96.0 ± 1.2 | 90.0 ± 1.4 | 114.4 ± 3.3 | 114.6 ± 3.2 |
| Inosine | 110.8 ± 0.4 | 85.2 ± 2.0 | 103.3 ± 2.4 | 95.1 ± 3.7 |
| Inosine 5'-monophosphate | 85.2 ± 0.4 | 87.6 ± 3.3 | 107.5 ± 1.5 | 88.1 ± 3.7 |
| Isocitric Acid | 86.6 ± 3.2 | 100.5 ± 0.3 | 85.9 ± 0.4 | 100.2 ± 1.7 |
| Isoleucine | 109.5 ± 3.4 | 114.4 ± 3.4 | 95.8 ± 1.9 | 100.8 ± 3.7 |
| Lactic Acid | 97.7 ± 1.9 | 91.7 ± 1.4 | 90.7 ± 2.4 | 91.0 ± 1.3 |
| Leucine | 106.8 ± 0.4 | 111.3 ± 2.8 | 103.9 ± 3.5 | 111.1 ± 2.4 |
| Levulinic acid | 88.7 ± 2.6 | 113.7 ± 1.2 | 113.4 ± 2.6 | 112.8 ± 0.8 |
| Lysine | 109.0 ± 1.4 | 95.1 ± 1.5 | 89.9 ± 1.3 | 104.3 ± 3.8 |
| Malic Acid | 107.4 ± 1.6 | 94.7 ± 3.6 | 89.4 ± 2.0 | 95.2 ± 3.3 |
| Mannose-1 with Mannose-6-Phosphate | 86.2 ± 3.4 | 99.0 ± 2.8 | 101.3 ± 2.0 | 104.3 ± 1.4 |
| Mesaconic acid | 98.5 ± 0.7 | 111.5 ± 1.2 | 104.8 ± 1.2 | 111.3 ± 0.5 |
| Methionine | 98.6 ± 1.4 | 95.8 ± 3.0 | 89.6 ± 2.0 | 95.7 ± 3.6 |
| Methionine sulfoxide | 94.0 ± 0.7 | 88.4 ± 3.1 | 112.9 ± 3.6 | 92.5 ± 2.9 |
| Methylmalonic Acid | 86.5 ± 2.9 | 91.6 ± 2.7 | 103.0 ± 3.1 | 96.7 ± 2.0 |
| N1-Acetylspermidine | 85.6 ± 1.0 | 98.7 ± 2.2 | 99.8 ± 0.8 | 93.8 ± 0.8 |
| N1-Acetylspermine | 88.8 ± 1.4 | 86.3 ± 0.6 | 88.2 ± 0.6 | 107.5 ± 1.4 |
| N8-Acetylspermidine | 105.3 ± 2.6 | 92.9 ± 0.6 | 97.2 ± 2.7 | 110.3 ± 0.3 |
| N-Acetyl-Aspartyl-Glutamate | 94.2 ± 2.4 | 88.1 ± 1.6 | 95.9 ± 2.4 | 109.1 ± 2.3 |
| N-Acetyl-Aspartyl-Glutamyl-Glutamate | 98.5 ± 0.5 | 95.1 ± 2.0 | 91.0 ± 4.0 | 109.6 ± 3.2 |
| N-Acetyl-Galactosamine | 91.6 ± 0.8 | 99.9 ± 0.4 | 113.2 ± 2.7 | 101.5 ± 3.1 |
| N-Acetyl-Leucine | 109.4 ± 3.8 | 90.8 ± 0.4 | 105.8 ± 3.4 | 92.4 ± 3.6 |
| N-Acetyl-Methionine | 101.6 ± 3.8 | 86.8 ± 3.4 | 93.6 ± 0.6 | 88.8 ± 1.5 |
| N-Acetylneuraminic acid | 110.4 ± 0.9 | 96.0 ± 1.0 | 110.6 ± 1.9 | 86.5 ± 0.7 |
| N-Acetyl-Proline | 88.0 ± 3.5 | 106.7 ± 0.4 | 101.2 ± 3.5 | 101.2 ± 3.5 |
| N-Acetyl-Putrescine | 111.4 ± 1.5 | 102.0 ± 0.8 | 92.3 ± 0.5 | 95.8 ± 0.7 |
| N-Acetyl-Serine | 112.3 ± 0.5 | 93.4 ± 0.8 | 85.9 ± 0.4 | 88.3 ± 3.4 |
| N-Acetyl-Serotonin | 87.5 ± 2.1 | 101.8 ± 0.9 | 101.6 ± 1.1 | 99.4 ± 3.7 |
| N-Acyl-Homoserine-lactone | 102.6 ± 3.1 | 92.2 ± 3.2 | 98.4 ± 0.5 | 96.1 ± 3.5 |
| NAD | 94.8 ± 2.5 | 90.7 ± 3.2 | 109.8 ± 3.0 | 95.8 ± 1.8 |
| NADH | 92.6 ± 0.4 | 95.4 ± 2.6 | 102.9 ± 3.2 | 105.0 ± 3.6 |
| NADP | 101.2 ± 2.1 | 96.5 ± 2.8 | 85.9 ± 3.8 | 93.7 ± 0.7 |
| NADPH | 86.9 ± 3.1 | 108.7 ± 3.2 | 96.1 ± 1.3 | 93.1 ± 1.1 |
| N-alpha-Acetyl-lysine | 90.9 ± 2.7 | 109.8 ± 2.8 | 86.7 ± 3.9 | 112.3 ± 3.7 |
| N-alpha-Acetyl-Ornithine | 109.1 ± 2.6 | 104.5 ± 0.9 | 104.7 ± 0.7 | 105.3 ± 2.7 |
| N-Carbamoyl-Proline | 95.4 ± 0.4 | 114.1 ± 2.3 | 88.6 ± 1.1 | 97.1 ± 3.9 |
| N-Carbamoyl-Sarcosine | 106.1 ± 0.7 | 86.0 ± 1.2 | 92.4 ± 3.3 | 96.3 ± 2.4 |
| N-Epsilon-Acetyl-lysine | 92.5 ± 3.1 | 107.9 ± 1.3 | 88.3 ± 3.2 | 91.9 ± 3.8 |
| Nicotinamide | 85.9 ± 0.4 | 86.6 ± 1.4 | 101.3 ± 2.5 | 109.0 ± 3.5 |
| N-Methyl-Aspartic Acid | 105.6 ± 1.6 | 102.3 ± 1.8 | 90.7 ± 2.1 | 98.9 ± 2.7 |
| O-Phosphoethanolamine | 106.6 ± 1.0 | 100.5 ± 1.2 | 85.0 ± 2.2 | 112.9 ± 3.8 |
| Ophthalmic acid | 106.8 ± 0.4 | 107.4 ± 0.6 | 108.2 ± 0.4 | 97.4 ± 2.3 |
| Ornithine | 103.7 ± 0.9 | 91.2 ± 0.9 | 94.2 ± 2.1 | 87.9 ± 3.6 |
| Orotidine 5'-monophosphate | 107.7 ± 2.3 | 87.0 ± 1.0 | 103.7 ± 0.7 | 111.2 ± 3.0 |
| Oxaloacetate | 106.2 ± 1.2 | 99.1 ± 3.5 | 89.0 ± 3.9 | 113.4 ± 3.7 |
| Pantothenic acid | 94.1 ± 3.8 | 85.4 ± 1.7 | 88.3 ± 0.3 | 112.9 ± 2.3 |
| Phenylalanine | 87.8 ± 1.7 | 87.5 ± 0.4 | 103.6 ± 1.2 | 113.7 ± 1.4 |
| Phenylpyruvate | 88.4 ± 0.6 | 106.0 ± 3.4 | 113.0 ± 2.2 | 86.7 ± 2.7 |
| Phosphodimethylethanolamine | 105.3 ± 1.4 | 87.3 ± 2.3 | 96.5 ± 3.1 | 101.4 ± 0.6 |
| Phosphoenolpyruvate | 112.0 ± 3.9 | 94.8 ± 2.1 | 93.7 ± 3.0 | 89.9 ± 2.8 |
| Phosphorylcholine | 101.7 ± 2.3 | 113.6 ± 2.1 | 87.6 ± 3.1 | 106.7 ± 3.7 |
| Pipecolic acid | 106.9 ± 1.2 | 108.9 ± 1.6 | 102.4 ± 2.4 | 91.5 ± 0.7 |
| Proline | 101.5 ± 0.7 | 104.8 ± 3.9 | 86.7 ± 3.3 | 109.8 ± 3.4 |
| Putrescine | 111.5 ± 0.7 | 114.5 ± 3.5 | 109.8 ± 4.0 | 87.0 ± 2.6 |
| Pyridoxal | 112.5 ± 3.9 | 102.0 ± 0.6 | 92.8 ± 1.1 | 114.4 ± 1.3 |
| Pyridoxamine | 107.3 ± 0.6 | 114.4 ± 2.3 | 85.1 ± 3.4 | 93.4 ± 1.7 |
| Pyridoxine | 87.9 ± 0.5 | 90.6 ± 1.1 | 113.1 ± 1.6 | 104.8 ± 3.4 |
| Pyroglutamic acid | 91.7 ± 1.0 | 106.5 ± 0.8 | 100.1 ± 2.4 | 86.4 ± 2.9 |
| Pyrrolysine | 90.1 ± 1.3 | 90.5 ± 3.0 | 86.9 ± 2.1 | 106.4 ± 3.6 |
| Pyruvic Acid | 95.0 ± 1.4 | 102.8 ± 1.8 | 114.0 ± 4.0 | 106.2 ± 1.5 |
| Riboflavin | 89.9 ± 3.8 | 100.5 ± 3.0 | 90.8 ± 3.0 | 112.9 ± 2.4 |
| Ribose-5-Phosphate | 96.7 ± 2.2 | 89.9 ± 2.8 | 99.7 ± 0.6 | 87.7 ± 1.2 |
| Ribulose-1-Phosphate | 93.5 ± 2.7 | 92.8 ± 2.9 | 99.5 ± 3.3 | 94.7 ± 2.1 |
| Ribulose-5-Phosphate | 104.9 ± 3.3 | 107.5 ± 3.7 | 94.3 ± 2.9 | 111.3 ± 1.3 |
| Saccharopine | 93.0 ± 2.9 | 102.0 ± 0.5 | 99.5 ± 3.4 | 87.6 ± 2.3 |
| S-Adenosylhomocysteine | 108.8 ± 0.6 | 86.0 ± 3.4 | 86.6 ± 3.0 | 98.7 ± 1.0 |
| S-Adenosylmethioninamine | 112.2 ± 0.9 | 89.6 ± 1.9 | 105.8 ± 3.9 | 103.5 ± 1.9 |
| S-Adenosylmethionine | 91.8 ± 0.8 | 94.0 ± 3.8 | 94.3 ± 3.8 | 101.5 ± 0.7 |
| Sarcosine | 113.3 ± 0.4 | 97.9 ± 3.9 | 91.7 ± 0.9 | 99.5 ± 2.4 |
| Sedoheptulose-7-phosphate | 101.1 ± 2.9 | 99.0 ± 0.4 | 87.4 ± 2.5 | 94.8 ± 2.1 |
| Selenohomocysteine | 95.0 ± 2.0 | 92.8 ± 3.2 | 112.3 ± 1.6 | 114.1 ± 2.9 |
| Selenomethionine | 109.1 ± 0.8 | 113.5 ± 0.8 | 87.1 ± 4.0 | 92.5 ± 4.0 |
| Serine | 111.0 ± 3.7 | 107.5 ± 2.5 | 87.7 ± 2.3 | 94.8 ± 2.9 |
| S-Methyl-Thioinosine | 93.8 ± 1.5 | 107.1 ± 2.7 | 101.5 ± 3.3 | 85.9 ± 0.7 |
| Spermidine | 107.9 ± 1.5 | 89.2 ± 0.5 | 105.4 ± 0.8 | 96.5 ± 1.5 |
| Spermine | 85.5 ± 0.6 | 107.6 ± 0.4 | 103.2 ± 0.3 | 100.9 ± 1.9 |
| Succinate | 112.0 ± 0.7 | 87.3 ± 3.3 | 85.4 ± 2.0 | 102.8 ± 3.2 |
| Sucrose | 110.0 ± 2.7 | 90.2 ± 0.9 | 91.8 ± 2.4 | 100.6 ± 2.2 |
| Symmetric dimethylarginine | 111.8 ± 3.5 | 99.1 ± 2.8 | 112.0 ± 1.5 | 105.9 ± 2.1 |
| Taurine | 92.2 ± 3.7 | 86.2 ± 2.0 | 103.5 ± 1.5 | 110.8 ± 0.9 |
| Tetrahydroneopterin | 102.6 ± 3.7 | 94.3 ± 2.7 | 90.4 ± 0.9 | 99.0 ± 2.1 |
| Thiamine | 86.6 ± 1.5 | 111.7 ± 1.1 | 95.7 ± 1.8 | 91.3 ± 0.7 |
| Thiamine monophosphate | 92.3 ± 1.2 | 109.5 ± 1.7 | 113.6 ± 4.0 | 95.6 ± 1.2 |
| Threonic acid | 109.5 ± 3.4 | 105.6 ± 1.8 | 95.9 ± 1.6 | 92.8 ± 3.4 |
| Threonine | 96.4 ± 1.8 | 102.9 ± 0.8 | 103.3 ± 1.5 | 100.5 ± 2.7 |
| Thymidine | 89.3 ± 3.8 | 92.0 ± 2.4 | 110.2 ± 1.4 | 106.9 ± 3.5 |
| Thymidine Monophosphate | 97.7 ± 2.0 | 90.7 ± 0.5 | 97.5 ± 1.7 | 87.7 ± 2.5 |
| Thymine | 92.2 ± 3.7 | 112.7 ± 2.8 | 97.2 ± 2.6 | 85.9 ± 3.7 |
| Trigonelleine | 106.9 ± 1.9 | 104.6 ± 2.7 | 104.0 ± 2.0 | 100.7 ± 0.7 |
| Trimethylamine N-oxide | 90.3 ± 3.2 | 98.0 ± 0.4 | 91.9 ± 0.5 | 95.0 ± 0.7 |
| Trimethyllysine | 102.7 ± 2.7 | 106.3 ± 0.7 | 105.5 ± 3.1 | 107.5 ± 4.0 |
| Tryptophan | 97.8 ± 1.6 | 94.8 ± 1.5 | 100.0 ± 1.6 | 113.7 ± 0.9 |
| Tyramine | 103.6 ± 3.3 | 96.6 ± 1.3 | 94.7 ± 3.9 | 90.7 ± 3.0 |
| Tyrosine | 106.7 ± 3.5 | 113.5 ± 3.1 | 97.4 ± 2.3 | 111.1 ± 0.7 |
| Uracil | 91.7 ± 1.3 | 109.5 ± 1.2 | 110.2 ± 2.4 | 94.7 ± 2.9 |
| Uridine 5'-diphosphate | 103.7 ± 1.5 | 114.6 ± 2.8 | 99.0 ± 1.4 | 95.7 ± 2.5 |
| Uridine Monophosphate | 91.5 ± 3.0 | 102.3 ± 2.8 | 104.9 ± 1.4 | 86.1 ± 2.0 |
| Uridine-5-Diphosphogalactose | 89.7 ± 1.4 | 104.0 ± 1.9 | 106.5 ± 1.9 | 113.5 ± 3.5 |
| Urocanic acid | 110.9 ± 0.4 | 93.6 ± 3.2 | 94.4 ± 2.5 | 105.1 ± 2.6 |
| UTP | 103.6 ± 0.8 | 113.7 ± 3.2 | 113.8 ± 3.4 | 100.8 ± 2.5 |
| Valine | 92.7 ± 2.5 | 92.9 ± 2.4 | 110.9 ± 2.0 | 114.7 ± 1.8 |
| XMP | 93.9 ± 1.2 | 109.2 ± 2.9 | 107.5 ± 0.4 | 98.5 ± 0.7 |
| Xylulose-5-Phosphate | 97.7 ± 2.3 | 99.5 ± 1.2 | 90.1 ± 3.9 | 111.5 ± 2.2 |

### **Table S26** | Intra – day accuracy (as determined by % recovery) (Urine extract as matrix, n = 5). The concentrations of QC_LLOQ_, QC_Low_, QC_Mid_ and QC_High_ were 10 ng/mL, 25 ng/mL, 100 ng/mL and 500 ng/mL respectively.

| Accuracy, % | QC_LLOQ_ | QC_Low_ | QC_Mid_ | QC_High_ |
| --- | --- | --- | --- | --- |
| 1-Methylguanine | 106.3 ± 1.9 | 107.8 ± 0.2 | 112.3 ± 1.3 | 98.0 ± 5.4 |
| 1-Methylhistidine | 97.3 ± 4.2 | 93.2 ± 3.0 | 111.2 ± 0.2 | 92.8 ± 3.0 |
| 1-Methylnicotinamide | 86.4 ± 5.8 | 105.8 ± 5.5 | 112.3 ± 2.6 | 103.5 ± 2.1 |
| 2-Aminoadipic Acid | 107.3 ± 2.1 | 111.0 ± 2.8 | 98.0 ± 0.9 | 100.1 ± 0.9 |
| 2-Deoxyadenosine | 102.6 ± 1.3 | 106.8 ± 5.6 | 100.6 ± 2.5 | 93.8 ± 4.4 |
| 2'-Deoxyadenosine 5'-monophosphate | 95.4 ± 3.7 | 107.1 ± 4.2 | 93.3 ± 0.5 | 85.6 ± 1.3 |
| 2'-deoxyadenosine 5'-triphosphate | 98.9 ± 3.8 | 102.2 ± 2.8 | 87.0 ± 1.1 | 106.2 ± 3.3 |
| 2'-Deoxycytidine-5'-monophosphate | 92.1 ± 4.2 | 107.3 ± 3.3 | 110.1 ± 5.5 | 108.4 ± 4.1 |
| 2-Deoxyguanosine | 97.0 ± 3.0 | 95.0 ± 4.8 | 109.5 ± 3.4 | 94.6 ± 3.7 |
| 2-hydroxybutyric acid | 103.2 ± 0.9 | 92.1 ± 2.8 | 101.2 ± 3.7 | 93.0 ± 5.4 |
| 2-hydroxyglutaric acid | 91.4 ± 3.4 | 91.2 ± 5.5 | 99.7 ± 3.5 | 91.5 ± 6.2 |
| 2-Hydroxyisocaproic acid | 92.3 ± 2.4 | 109.4 ± 1.2 | 100.5 ± 1.3 | 107.0 ± 1.2 |
| 2-ketobutyric acid | 88.7 ± 4.4 | 95.8 ± 0.4 | 88.8 ± 0.5 | 92.9 ± 0.4 |
| 2-oxoadipic acid | 93.4 ± 1.3 | 89.9 ± 2.5 | 104.9 ± 5.4 | 104.3 ± 6.1 |
| 2-Phenylglycine | 98.2 ± 5.0 | 100.9 ± 4.8 | 100.8 ± 0.5 | 101.3 ± 3.9 |
| 2-phosphoglycerate | 110.4 ± 5.4 | 100.7 ± 2.8 | 104.0 ± 2.9 | 86.6 ± 4.7 |
| 3-hydroxy-3-methyl-2-oxobutanoate | 102.4 ± 0.4 | 93.2 ± 3.0 | 112.9 ± 0.5 | 91.4 ± 3.9 |
| 3-hydroxybutyric acid | 89.5 ± 4.1 | 87.1 ± 0.7 | 89.3 ± 5.8 | 111.0 ± 2.4 |
| 3-Methylhistidine | 110.8 ± 4.6 | 109.9 ± 3.7 | 106.9 ± 4.6 | 95.3 ± 0.8 |
| 3-Phosphoglyceraldehyde | 114.5 ± 1.0 | 94.7 ± 3.6 | 113.7 ± 2.1 | 113.1 ± 1.8 |
| 3-phosphoglycerate | 109.3 ± 2.2 | 111.2 ± 1.4 | 100.3 ± 2.7 | 95.8 ± 3.6 |
| 4-amino-butyric acid | 94.1 ± 0.4 | 99.6 ± 0.4 | 103.7 ± 3.2 | 89.8 ± 1.7 |
| 4-aspartyl-phosphate | 97.0 ± 3.1 | 87.8 ± 3.1 | 85.6 ± 3.5 | 96.2 ± 4.4 |
| 4-Hydroxybenzoic acid | 108.2 ± 3.3 | 99.0 ± 1.8 | 90.2 ± 4.6 | 96.4 ± 5.1 |
| 4-Hydroxyphenylacetylglycine | 114.3 ± 5.8 | 113.8 ± 0.9 | 90.7 ± 0.8 | 93.9 ± 2.9 |
| 4-methyl-2-oxopentanoate | 106.8 ± 3.5 | 112.9 ± 0.4 | 95.6 ± 3.5 | 96.7 ± 5.3 |
| 4-Oxoglutaramate | 114.4 ± 3.7 | 109.0 ± 3.5 | 91.3 ± 0.4 | 111.7 ± 0.8 |
| 5-Hydroxyindolepyruvate | 87.1 ± 1.6 | 112.1 ± 0.7 | 86.0 ± 1.6 | 85.6 ± 1.3 |
| 5-Hydroxylysine | 102.9 ± 4.9 | 88.9 ± 5.0 | 90.5 ± 1.9 | 114.5 ± 0.2 |
| 5-Hydroxy-Tryptophan | 90.1 ± 6.0 | 100.1 ± 2.0 | 101.6 ± 2.7 | 85.6 ± 6.3 |
| 5-Phosphoribosylamine | 107.3 ± 3.5 | 85.9 ± 6.0 | 109.8 ± 2.8 | 104.2 ± 4.7 |
| 6-phosphogluconate | 108.1 ± 5.0 | 92.6 ± 5.9 | 102.1 ± 5.8 | 101.3 ± 0.4 |
| Acetylcarnitine | 97.9 ± 2.8 | 94.4 ± 5.9 | 99.6 ± 5.7 | 100.0 ± 5.5 |
| Aconitic Acid | 111.3 ± 4.5 | 93.2 ± 3.9 | 94.6 ± 3.1 | 101.9 ± 0.7 |
| Adenine | 95.8 ± 1.9 | 104.6 ± 5.0 | 104.4 ± 1.8 | 85.5 ± 1.1 |
| Adenosine | 87.1 ± 4.2 | 106.4 ± 2.7 | 86.4 ± 4.2 | 90.6 ± 4.9 |
| Adenosine 3',5'-cyclic monophosphate | 104.5 ± 2.6 | 104.6 ± 4.5 | 91.0 ± 0.8 | 86.1 ± 4.5 |
| Adenosine Monophosphate | 90.1 ± 1.0 | 102.5 ± 1.1 | 113.0 ± 3.9 | 104.0 ± 3.3 |
| Adenosine-5'-diphosphate | 104.8 ± 6.1 | 86.5 ± 6.0 | 103.5 ± 0.4 | 113.7 ± 2.9 |
| Adenosine-5'-triphosphate | 90.5 ± 5.3 | 112.6 ± 0.3 | 102.8 ± 5.0 | 94.5 ± 2.7 |
| Adenylosuccinate | 113.8 ± 5.9 | 106.8 ± 3.6 | 108.5 ± 3.4 | 106.5 ± 3.3 |
| Alanine | 106.8 ± 6.0 | 91.3 ± 3.3 | 97.4 ± 5.6 | 104.2 ± 3.2 |
| alpha-ketoglutaric acid | 93.4 ± 5.7 | 108.2 ± 6.1 | 100.4 ± 4.1 | 108.9 ± 4.1 |
| Anserine | 111.1 ± 4.9 | 97.8 ± 4.9 | 114.3 ± 2.4 | 110.6 ± 0.9 |
| Arginine | 86.5 ± 0.7 | 86.2 ± 2.9 | 94.8 ± 2.3 | 106.3 ± 2.3 |
| Argininosuccinic Acid | 96.4 ± 2.4 | 105.8 ± 5.6 | 113.4 ± 3.8 | 113.5 ± 0.6 |
| Asparagine | 108.8 ± 2.5 | 100.5 ± 4.4 | 110.2 ± 3.0 | 114.2 ± 4.9 |
| Aspartic acid | 92.1 ± 5.4 | 85.5 ± 2.6 | 93.9 ± 2.0 | 103.1 ± 2.2 |
| Asymmetric dimethylarginine | 88.7 ± 3.0 | 96.1 ± 6.3 | 108.5 ± 4.2 | 94.1 ± 0.8 |
| Azelaic acid | 95.7 ± 3.7 | 95.3 ± 1.6 | 113.3 ± 2.9 | 111.0 ± 1.1 |
| beta-Alanine | 109.4 ± 5.5 | 90.9 ± 5.7 | 96.7 ± 4.2 | 107.7 ± 2.4 |
| Betaine | 105.1 ± 3.8 | 96.5 ± 1.2 | 87.1 ± 5.1 | 107.6 ± 2.1 |
| Biopterin | 108.1 ± 2.5 | 93.3 ± 2.6 | 108.1 ± 5.2 | 109.6 ± 0.3 |
| Biotin | 101.0 ± 4.6 | 94.8 ± 4.2 | 106.0 ± 5.3 | 114.2 ± 4.2 |
| Carnitine | 113.5 ± 4.3 | 111.0 ± 3.5 | 85.5 ± 6.0 | 107.4 ± 1.2 |
| Carnosine | 106.4 ± 5.1 | 114.8 ± 1.9 | 98.2 ± 1.3 | 107.3 ± 4.6 |
| CDP-choline | 103.4 ± 3.1 | 95.6 ± 2.2 | 94.3 ± 1.0 | 97.2 ± 3.1 |
| cGMP | 86.8 ± 3.3 | 106.7 ± 4.2 | 106.4 ± 0.2 | 92.9 ± 5.8 |
| Choline | 103.1 ± 3.9 | 96.1 ± 5.6 | 89.9 ± 5.0 | 94.8 ± 1.4 |
| Cinnamic acid | 105.7 ± 2.2 | 95.1 ± 1.7 | 109.5 ± 2.8 | 109.7 ± 4.5 |
| Citraconic acid | 92.0 ± 5.1 | 114.7 ± 5.2 | 91.0 ± 1.4 | 104.8 ± 0.2 |
| Citramalic acid | 90.9 ± 4.1 | 87.8 ± 2.6 | 107.4 ± 0.5 | 98.1 ± 1.0 |
| Citric Acid | 109.8 ± 0.7 | 99.5 ± 5.9 | 97.7 ± 1.0 | 106.1 ± 2.7 |
| Citrulline | 111.6 ± 5.9 | 108.1 ± 4.9 | 109.4 ± 3.2 | 90.2 ± 2.8 |
| Creatine | 87.5 ± 3.9 | 95.0 ± 0.9 | 90.0 ± 2.2 | 87.8 ± 5.7 |
| Creatinine | 95.9 ± 3.1 | 100.9 ± 0.6 | 97.7 ± 0.2 | 108.1 ± 3.4 |
| Cystathionine | 112.8 ± 6.1 | 96.4 ± 3.6 | 110.9 ± 4.3 | 92.7 ± 0.4 |
| Cysteamine | 102.3 ± 2.6 | 94.1 ± 0.9 | 107.2 ± 4.4 | 91.3 ± 3.1 |
| Cysteine | 112.3 ± 1.8 | 91.9 ± 5.0 | 98.2 ± 1.8 | 110.0 ± 0.1 |
| Cysteinylglycine | 91.5 ± 3.1 | 107.7 ± 1.3 | 93.9 ± 4.8 | 107.7 ± 1.6 |
| Cystine | 103.1 ± 6.2 | 88.4 ± 4.5 | 99.3 ± 6.1 | 103.2 ± 5.7 |
| Cytidine | 85.3 ± 3.0 | 108.1 ± 1.8 | 90.1 ± 3.2 | 111.2 ± 2.1 |
| Cytidine 3',5'-cyclic monophosphate | 88.1 ± 0.3 | 95.6 ± 5.9 | 93.4 ± 5.5 | 90.4 ± 3.7 |
| Cytidine 5'-diphosphate | 112.8 ± 4.2 | 103.0 ± 2.9 | 112.7 ± 0.8 | 94.1 ± 1.5 |
| Cytidine 5'-triphosphate | 105.9 ± 3.6 | 111.7 ± 0.9 | 99.7 ± 2.5 | 89.9 ± 2.8 |
| Cytidine monophosphate | 87.7 ± 1.4 | 93.2 ± 3.2 | 90.7 ± 2.4 | 106.4 ± 3.2 |
| Cytosine | 106.4 ± 3.5 | 112.3 ± 1.5 | 109.2 ± 0.3 | 91.7 ± 1.7 |
| dADP | 104.6 ± 2.7 | 101.5 ± 2.4 | 106.2 ± 3.3 | 110.3 ± 3.5 |
| Diethanolamine | 97.2 ± 2.6 | 88.0 ± 1.9 | 101.7 ± 3.5 | 87.6 ± 1.1 |
| Dihydrobiopterin | 89.9 ± 2.8 | 106.5 ± 2.4 | 109.7 ± 3.3 | 91.4 ± 1.7 |
| Dihydroneopterin triphosphate | 87.2 ± 3.1 | 85.4 ± 4.0 | 108.0 ± 1.6 | 106.0 ± 0.9 |
| Dihydroxyacetonephosphate | 89.9 ± 0.5 | 90.2 ± 1.6 | 93.0 ± 3.2 | 104.0 ± 1.2 |
| Dimethylglycine | 107.5 ± 1.4 | 85.2 ± 0.6 | 97.5 ± 2.9 | 112.8 ± 2.7 |
| Ectoine | 105.8 ± 1.6 | 107.0 ± 0.7 | 104.0 ± 1.5 | 103.1 ± 2.0 |
| Erythrose-4-phosphate | 96.7 ± 3.2 | 97.0 ± 2.6 | 100.2 ± 3.5 | 92.1 ± 2.9 |
| FAD | 99.5 ± 3.7 | 89.8 ± 2.5 | 103.3 ± 1.6 | 104.7 ± 0.5 |
| Folic acid | 107.7 ± 0.8 | 108.2 ± 1.1 | 106.4 ± 0.5 | 114.2 ± 0.9 |
| Fructose-6-phosphate | 114.4 ± 1.2 | 87.7 ± 0.9 | 113.7 ± 3.5 | 103.9 ± 3.0 |
| Fumaric Acid | 99.0 ± 1.3 | 109.8 ± 3.9 | 109.6 ± 1.5 | 104.6 ± 3.8 |
| Galactosamine | 96.1 ± 3.3 | 94.6 ± 1.5 | 100.3 ± 3.9 | 110.4 ± 2.0 |
| Galactose-1-phosphate | 108.9 ± 0.4 | 112.5 ± 0.7 | 87.8 ± 3.7 | 107.9 ± 1.3 |
| Galactose-6-Phosphate | 102.2 ± 3.6 | 104.1 ± 1.9 | 111.7 ± 3.4 | 94.8 ± 3.0 |
| gamma-Glutamylcysteine | 97.2 ± 1.2 | 100.4 ± 2.9 | 108.1 ± 3.2 | 107.2 ± 1.3 |
| Glucosamine | 110.3 ± 1.4 | 114.1 ± 1.6 | 114.1 ± 3.5 | 114.0 ± 2.4 |
| Glucosamine-6-Phosphate | 93.8 ± 0.7 | 106.3 ± 0.3 | 89.9 ± 3.6 | 110.3 ± 3.7 |
| Glucose | 102.3 ± 3.0 | 111.5 ± 3.9 | 105.1 ± 1.9 | 98.7 ± 1.7 |
| Glucose-1-phosphate | 99.7 ± 1.8 | 107.3 ± 3.0 | 104.8 ± 3.9 | 102.1 ± 2.0 |
| Glucose-6-Phosphate | 108.8 ± 4.0 | 113.8 ± 2.3 | 94.6 ± 2.2 | 92.4 ± 3.6 |
| Glucuronic acid | 114.3 ± 0.5 | 86.8 ± 0.8 | 86.0 ± 3.8 | 88.0 ± 2.6 |
| Glutamic acid | 102.2 ± 3.2 | 93.8 ± 1.0 | 108.7 ± 3.8 | 99.7 ± 3.6 |
| Glutamine | 91.1 ± 3.8 | 100.4 ± 2.1 | 93.2 ± 1.5 | 105.7 ± 1.0 |
| Glutathione | 98.8 ± 1.3 | 86.1 ± 2.4 | 113.9 ± 1.7 | 85.1 ± 2.1 |
| Glutathionylspermine | 102.0 ± 3.3 | 96.0 ± 2.9 | 93.2 ± 1.4 | 96.7 ± 0.9 |
| Glycerophosphocholine | 104.8 ± 0.4 | 105.6 ± 0.4 | 97.2 ± 0.4 | 109.8 ± 1.8 |
| Glycine | 88.3 ± 2.0 | 87.9 ± 1.7 | 103.2 ± 0.9 | 104.7 ± 1.5 |
| Glycolic acid | 95.6 ± 3.6 | 87.7 ± 1.0 | 99.1 ± 1.1 | 107.6 ± 2.2 |
| Guanidinoacetic acid | 94.0 ± 1.6 | 86.3 ± 2.2 | 96.5 ± 2.9 | 100.2 ± 2.0 |
| Guanidinobutanoic acid | 104.1 ± 2.3 | 109.3 ± 1.5 | 88.9 ± 1.2 | 111.8 ± 0.6 |
| Guanine | 104.0 ± 1.6 | 108.7 ± 3.7 | 111.1 ± 1.2 | 102.4 ± 1.1 |
| Guanosine | 95.9 ± 0.7 | 93.9 ± 1.6 | 103.6 ± 0.9 | 111.0 ± 2.3 |
| Guanosine 5'-diphosphate | 88.0 ± 3.0 | 85.2 ± 3.4 | 105.3 ± 3.1 | 98.6 ± 3.5 |
| Guanosine-5'-monophosphate | 91.9 ± 0.9 | 102.8 ± 3.8 | 97.0 ± 2.4 | 96.2 ± 2.6 |
| Hercynine | 103.8 ± 3.1 | 98.8 ± 3.7 | 114.5 ± 2.8 | 89.7 ± 1.1 |
| Hippuric acid | 102.7 ± 2.7 | 92.0 ± 2.1 | 86.9 ± 2.7 | 107.7 ± 1.3 |
| Histamine | 104.2 ± 0.7 | 112.0 ± 3.7 | 112.9 ± 3.2 | 92.6 ± 1.6 |
| Histidine | 115.0 ± 1.3 | 93.2 ± 1.9 | 103.5 ± 0.8 | 114.2 ± 0.9 |
| Homoalanine | 98.0 ± 0.6 | 90.3 ± 3.8 | 110.3 ± 0.6 | 88.1 ± 1.1 |
| Homoarginine | 94.2 ± 2.9 | 98.3 ± 1.0 | 91.9 ± 1.4 | 86.1 ± 3.1 |
| Homocysteine | 104.4 ± 2.3 | 104.2 ± 2.2 | 109.7 ± 3.4 | 106.7 ± 0.4 |
| Homogentisic acid | 90.8 ± 3.3 | 87.0 ± 3.6 | 87.8 ± 2.5 | 111.7 ± 3.2 |
| Homoleucine | 93.6 ± 3.6 | 104.2 ± 3.1 | 100.2 ± 1.1 | 98.3 ± 2.5 |
| Homovanillic acid | 113.0 ± 3.8 | 87.1 ± 1.3 | 108.2 ± 1.4 | 87.4 ± 1.5 |
| Hydantoin-5-propionate | 93.8 ± 1.8 | 94.0 ± 2.6 | 101.6 ± 2.4 | 100.7 ± 2.7 |
| Hypotaurine | 94.2 ± 0.7 | 93.6 ± 2.7 | 111.9 ± 3.3 | 107.7 ± 0.6 |
| Hypoxanthine | 110.7 ± 2.2 | 105.5 ± 1.6 | 88.3 ± 2.9 | 108.6 ± 1.3 |
| Indole-3-acetamide | 110.6 ± 1.1 | 85.8 ± 1.2 | 115.0 ± 2.3 | 86.2 ± 2.3 |
| Inosine | 87.4 ± 3.8 | 112.1 ± 2.6 | 111.6 ± 1.9 | 99.6 ± 2.4 |
| Inosine 5'-monophosphate | 113.9 ± 1.6 | 103.0 ± 1.6 | 97.8 ± 2.6 | 94.6 ± 3.4 |
| Isocitric Acid | 85.4 ± 3.5 | 90.0 ± 2.7 | 101.4 ± 3.8 | 85.2 ± 3.6 |
| Isoleucine | 103.9 ± 0.7 | 99.9 ± 3.8 | 101.0 ± 2.8 | 102.2 ± 2.1 |
| Lactic Acid | 99.8 ± 3.4 | 100.8 ± 1.8 | 104.3 ± 3.4 | 101.7 ± 1.3 |
| Leucine | 88.3 ± 2.1 | 108.1 ± 1.2 | 111.9 ± 0.3 | 106.3 ± 3.8 |
| Levulinic acid | 111.5 ± 2.7 | 98.6 ± 2.2 | 93.7 ± 3.8 | 90.1 ± 1.8 |
| Lysine | 91.1 ± 0.6 | 93.6 ± 2.3 | 91.2 ± 2.4 | 111.2 ± 0.9 |
| Malic Acid | 106.3 ± 0.8 | 95.4 ± 1.9 | 93.2 ± 1.0 | 101.3 ± 3.1 |
| Mannose-1 with Mannose-6-Phosphate | 91.5 ± 3.1 | 85.0 ± 2.0 | 113.3 ± 3.1 | 108.0 ± 2.7 |
| Mesaconic acid | 93.1 ± 1.9 | 95.4 ± 2.1 | 94.5 ± 2.8 | 96.5 ± 1.7 |
| Methionine | 109.8 ± 1.1 | 98.6 ± 2.8 | 106.2 ± 1.1 | 109.1 ± 3.9 |
| Methionine sulfoxide | 111.3 ± 3.7 | 89.3 ± 3.6 | 85.9 ± 0.8 | 102.5 ± 3.5 |
| Methylmalonic Acid | 92.6 ± 0.8 | 109.9 ± 1.6 | 91.9 ± 3.8 | 111.2 ± 3.8 |
| N1-Acetylspermidine | 89.6 ± 0.7 | 106.0 ± 1.3 | 93.3 ± 2.4 | 96.0 ± 1.5 |
| N1-Acetylspermine | 109.0 ± 3.2 | 85.7 ± 3.0 | 109.8 ± 2.1 | 114.2 ± 0.7 |
| N8-Acetylspermidine | 91.4 ± 3.6 | 93.4 ± 2.9 | 113.6 ± 1.9 | 105.8 ± 1.8 |
| N-Acetyl-Aspartyl-Glutamate | 107.5 ± 0.8 | 111.3 ± 1.7 | 105.3 ± 2.5 | 106.4 ± 3.2 |
| N-Acetyl-Aspartyl-Glutamyl-Glutamate | 97.6 ± 2.3 | 113.1 ± 1.4 | 110.1 ± 2.8 | 107.5 ± 0.8 |
| N-Acetyl-Galactosamine | 98.5 ± 2.0 | 88.4 ± 0.3 | 110.1 ± 3.3 | 100.9 ± 2.1 |
| N-Acetyl-Leucine | 112.7 ± 2.6 | 90.3 ± 3.3 | 108.3 ± 1.0 | 93.1 ± 0.7 |
| N-Acetyl-Methionine | 92.8 ± 3.6 | 110.6 ± 1.0 | 87.5 ± 2.3 | 90.3 ± 3.2 |
| N-Acetylneuraminic acid | 107.2 ± 0.6 | 101.9 ± 3.4 | 94.4 ± 3.6 | 102.3 ± 1.8 |
| N-Acetyl-Proline | 87.4 ± 0.6 | 99.6 ± 0.9 | 105.4 ± 3.2 | 113.0 ± 2.9 |
| N-Acetyl-Putrescine | 95.0 ± 3.2 | 98.7 ± 1.6 | 102.2 ± 0.7 | 110.8 ± 2.9 |
| N-Acetyl-Serine | 104.1 ± 2.2 | 104.2 ± 3.3 | 95.6 ± 2.5 | 97.5 ± 0.3 |
| N-Acetyl-Serotonin | 107.2 ± 0.4 | 90.5 ± 2.5 | 91.6 ± 2.4 | 91.4 ± 1.4 |
| N-Acyl-Homoserine-lactone | 87.1 ± 0.9 | 88.6 ± 2.3 | 100.9 ± 0.8 | 98.0 ± 3.1 |
| NAD | 96.3 ± 0.9 | 86.9 ± 2.8 | 91.5 ± 0.8 | 113.9 ± 3.7 |
| NADH | 110.9 ± 0.9 | 97.2 ± 3.4 | 106.7 ± 1.5 | 92.0 ± 0.3 |
| NADP | 110.8 ± 0.4 | 102.7 ± 2.5 | 102.9 ± 0.8 | 98.8 ± 3.8 |
| NADPH | 105.7 ± 3.3 | 89.0 ± 3.2 | 114.9 ± 1.2 | 112.5 ± 0.6 |
| N-alpha-Acetyl-lysine | 91.0 ± 0.9 | 91.6 ± 1.9 | 92.5 ± 1.7 | 114.7 ± 0.6 |
| N-alpha-Acetyl-Ornithine | 86.4 ± 2.6 | 88.7 ± 1.9 | 103.5 ± 1.8 | 92.7 ± 0.8 |
| N-Carbamoyl-Proline | 113.4 ± 2.0 | 99.4 ± 3.2 | 97.4 ± 1.6 | 98.8 ± 2.5 |
| N-Carbamoyl-Sarcosine | 109.8 ± 3.3 | 103.6 ± 2.0 | 102.6 ± 1.6 | 104.6 ± 1.7 |
| N-Epsilon-Acetyl-lysine | 91.0 ± 2.1 | 107.6 ± 2.4 | 107.9 ± 2.0 | 114.2 ± 2.5 |
| Nicotinamide | 87.6 ± 0.4 | 87.2 ± 2.2 | 108.8 ± 0.5 | 112.5 ± 2.7 |
| N-Methyl-Aspartic Acid | 104.3 ± 2.7 | 97.8 ± 1.0 | 90.8 ± 3.7 | 106.4 ± 0.8 |
| O-Phosphoethanolamine | 99.9 ± 3.8 | 109.5 ± 2.9 | 109.9 ± 2.5 | 96.1 ± 2.8 |
| Ophthalmic acid | 96.4 ± 0.7 | 107.9 ± 2.8 | 106.5 ± 0.9 | 88.3 ± 3.6 |
| Ornithine | 110.8 ± 3.0 | 112.7 ± 0.8 | 101.8 ± 0.6 | 86.9 ± 3.7 |
| Orotidine 5'-monophosphate | 109.9 ± 2.5 | 86.8 ± 2.1 | 106.3 ± 0.5 | 98.5 ± 2.0 |
| Oxaloacetate | 105.7 ± 3.7 | 102.9 ± 1.6 | 97.0 ± 2.6 | 93.7 ± 2.1 |
| Pantothenic acid | 86.0 ± 3.7 | 103.2 ± 1.0 | 106.9 ± 1.0 | 107.8 ± 1.9 |
| Phenylalanine | 108.2 ± 3.7 | 87.4 ± 1.6 | 96.3 ± 1.2 | 85.0 ± 3.9 |
| Phenylpyruvate | 114.2 ± 3.2 | 107.2 ± 1.5 | 109.3 ± 1.6 | 110.5 ± 1.7 |
| Phosphodimethylethanolamine | 113.3 ± 2.0 | 110.8 ± 3.5 | 87.8 ± 3.9 | 91.7 ± 3.6 |
| Phosphoenolpyruvate | 90.7 ± 3.0 | 86.0 ± 0.7 | 100.3 ± 2.4 | 93.1 ± 3.2 |
| Phosphorylcholine | 103.5 ± 3.1 | 101.7 ± 1.7 | 87.2 ± 0.7 | 88.4 ± 2.4 |
| Pipecolic acid | 104.3 ± 1.8 | 113.7 ± 1.1 | 97.4 ± 0.9 | 112.6 ± 3.9 |
| Proline | 88.4 ± 3.0 | 94.9 ± 3.0 | 100.9 ± 0.7 | 110.6 ± 3.3 |
| Putrescine | 100.2 ± 0.3 | 85.3 ± 0.8 | 108.0 ± 0.5 | 96.1 ± 0.7 |
| Pyridoxal | 106.8 ± 3.4 | 105.8 ± 3.3 | 101.0 ± 1.5 | 90.7 ± 0.6 |
| Pyridoxamine | 87.3 ± 0.9 | 106.4 ± 0.4 | 106.5 ± 3.3 | 100.0 ± 1.2 |
| Pyridoxine | 97.7 ± 2.8 | 97.5 ± 1.3 | 114.2 ± 3.3 | 111.6 ± 3.2 |
| Pyroglutamic acid | 101.4 ± 0.6 | 86.4 ± 1.2 | 102.9 ± 1.7 | 98.6 ± 3.8 |
| Pyrrolysine | 113.5 ± 1.0 | 90.3 ± 3.8 | 112.0 ± 1.1 | 106.2 ± 3.2 |
| Pyruvic Acid | 86.9 ± 1.7 | 110.3 ± 0.7 | 100.2 ± 2.0 | 90.4 ± 3.9 |
| Riboflavin | 89.0 ± 3.3 | 87.8 ± 1.3 | 109.3 ± 2.6 | 92.5 ± 0.6 |
| Ribose-5-Phosphate | 99.3 ± 2.0 | 88.7 ± 1.3 | 102.4 ± 3.8 | 89.0 ± 0.9 |
| Ribulose-1-Phosphate | 95.9 ± 0.6 | 107.9 ± 3.4 | 94.1 ± 0.7 | 92.5 ± 1.1 |
| Ribulose-5-Phosphate | 98.7 ± 3.6 | 114.3 ± 3.4 | 98.1 ± 0.8 | 88.1 ± 0.3 |
| Saccharopine | 113.7 ± 2.6 | 114.5 ± 2.6 | 96.6 ± 3.3 | 108.0 ± 2.6 |
| S-Adenosylhomocysteine | 111.0 ± 1.8 | 107.1 ± 0.5 | 111.9 ± 3.6 | 104.1 ± 3.8 |
| S-Adenosylmethioninamine | 101.6 ± 1.7 | 90.0 ± 1.3 | 103.4 ± 1.3 | 90.6 ± 3.9 |
| S-Adenosylmethionine | 90.8 ± 2.6 | 109.9 ± 1.8 | 113.6 ± 1.6 | 102.8 ± 3.1 |
| Sarcosine | 100.9 ± 3.2 | 109.9 ± 1.4 | 92.1 ± 1.7 | 103.4 ± 2.3 |
| Sedoheptulose-7-phosphate | 95.1 ± 1.2 | 107.2 ± 2.5 | 111.3 ± 1.4 | 94.5 ± 1.3 |
| Selenohomocysteine | 102.6 ± 0.6 | 87.0 ± 3.3 | 90.7 ± 0.6 | 110.9 ± 1.3 |
| Selenomethionine | 91.4 ± 0.9 | 114.5 ± 1.0 | 101.2 ± 1.1 | 107.7 ± 1.2 |
| Serine | 108.1 ± 2.3 | 98.5 ± 2.5 | 109.8 ± 3.9 | 88.9 ± 3.2 |
| S-Methyl-Thioinosine | 108.8 ± 0.6 | 110.5 ± 1.5 | 102.4 ± 3.4 | 111.0 ± 4.0 |
| Spermidine | 86.3 ± 3.3 | 99.6 ± 0.5 | 108.0 ± 2.1 | 85.7 ± 1.4 |
| Spermine | 96.6 ± 1.5 | 108.6 ± 0.6 | 108.2 ± 3.5 | 104.2 ± 0.5 |
| Succinate | 95.6 ± 3.4 | 113.4 ± 0.8 | 107.5 ± 3.8 | 94.0 ± 1.5 |
| Sucrose | 93.6 ± 3.6 | 97.3 ± 2.5 | 88.5 ± 3.2 | 95.9 ± 3.5 |
| Symmetric dimethylarginine | 89.0 ± 2.4 | 111.6 ± 3.9 | 94.7 ± 2.4 | 104.9 ± 0.9 |
| Taurine | 100.6 ± 2.6 | 112.5 ± 1.8 | 102.4 ± 1.2 | 101.8 ± 2.2 |
| Tetrahydroneopterin | 106.1 ± 3.5 | 88.7 ± 2.1 | 86.9 ± 0.4 | 95.3 ± 2.5 |
| Thiamine | 98.8 ± 2.9 | 100.9 ± 2.6 | 109.9 ± 2.5 | 91.5 ± 1.6 |
| Thiamine monophosphate | 100.1 ± 1.0 | 92.1 ± 0.8 | 100.6 ± 1.3 | 115.0 ± 0.8 |
| Threonic acid | 89.1 ± 2.5 | 86.4 ± 3.9 | 104.1 ± 1.1 | 101.7 ± 0.5 |
| Threonine | 102.8 ± 2.8 | 97.4 ± 2.1 | 107.4 ± 0.8 | 109.8 ± 2.4 |
| Thymidine | 108.5 ± 3.7 | 113.0 ± 2.5 | 99.4 ± 3.1 | 93.9 ± 0.5 |
| Thymidine Monophosphate | 100.0 ± 2.6 | 114.7 ± 2.6 | 112.9 ± 1.2 | 101.1 ± 3.8 |
| Thymine | 109.7 ± 2.9 | 103.5 ± 1.4 | 85.7 ± 3.2 | 94.1 ± 1.7 |
| Trigonelleine | 109.0 ± 1.2 | 103.3 ± 2.6 | 111.4 ± 1.1 | 90.4 ± 0.5 |
| Trimethylamine N-oxide | 109.6 ± 4.0 | 109.4 ± 0.7 | 96.0 ± 3.8 | 93.9 ± 3.9 |
| Trimethyllysine | 101.3 ± 3.8 | 105.7 ± 3.1 | 103.6 ± 3.2 | 109.7 ± 2.4 |
| Tryptophan | 108.5 ± 3.2 | 95.9 ± 1.3 | 98.5 ± 0.9 | 113.5 ± 0.7 |
| Tyramine | 108.8 ± 1.1 | 90.3 ± 1.0 | 86.9 ± 0.7 | 110.5 ± 1.4 |
| Tyrosine | 108.6 ± 3.7 | 89.3 ± 1.8 | 102.3 ± 2.7 | 88.7 ± 0.7 |
| Uracil | 90.8 ± 1.7 | 95.1 ± 2.4 | 107.8 ± 2.8 | 113.9 ± 1.7 |
| Uridine 5'-diphosphate | 97.1 ± 3.2 | 96.6 ± 3.5 | 91.1 ± 1.1 | 92.6 ± 3.9 |
| Uridine Monophosphate | 107.3 ± 2.5 | 106.4 ± 2.9 | 114.3 ± 2.3 | 109.3 ± 1.0 |
| Uridine-5-Diphosphogalactose | 89.6 ± 0.6 | 113.9 ± 0.3 | 86.3 ± 0.8 | 113.1 ± 3.1 |
| Urocanic acid | 105.5 ± 3.7 | 95.6 ± 2.4 | 110.3 ± 3.1 | 92.0 ± 3.9 |
| UTP | 101.3 ± 3.6 | 96.8 ± 2.6 | 85.0 ± 3.9 | 93.5 ± 0.9 |
| Valine | 99.2 ± 1.4 | 95.7 ± 2.8 | 93.1 ± 3.0 | 99.4 ± 2.2 |
| XMP | 112.0 ± 2.8 | 111.2 ± 2.7 | 89.0 ± 0.5 | 111.7 ± 2.4 |
| Xylulose-5-Phosphate | 110.3 ± 3.9 | 100.2 ± 2.6 | 91.6 ± 0.5 | 104.1 ± 1.4 |

### **Table S27** | Inter – day accuracy (as determined by % recovery) (Colorectal organoid extract as matrix, n = 5). The concentrations of QC_LLOQ_, QC_Low_, QC_Mid_ and QC_High_ were 10 ng/mL, 25 ng/mL, 100 ng/mL and 500 ng/mL respectively.

| Accuracy, % | QC_LLOQ_ | QC_Low_ | QC_Mid_ | QC_High_ |
| --- | --- | --- | --- | --- |
| 1-Methylguanine | 106.4 ± 1.8 | 106.1 ± 1.1 | 114.3 ± 1.5 | 100.8 ± 1.4 |
| 1-Methylhistidine | 106.0 ± 1.4 | 97.4 ± 0.7 | 101.3 ± 1.0 | 102.1 ± 1.0 |
| 1-Methylnicotinamide | 101.1 ± 1.7 | 103.9 ± 0.7 | 98.2 ± 0.5 | 97.8 ± 0.8 |
| 2-Aminoadipic Acid | 98.4 ± 1.6 | 92.9 ± 1.2 | 105.9 ± 0.4 | 102.2 ± 0.3 |
| 2-Deoxyadenosine | 91.2 ± 1.1 | 87.9 ± 0.5 | 93.0 ± 0.3 | 91.5 ± 1.1 |
| 2'-Deoxyadenosine 5'-monophosphate | 100.3 ± 1.6 | 113.9 ± 1.8 | 98.1 ± 0.2 | 88.8 ± 1.2 |
| 2'-deoxyadenosine 5'-triphosphate | 93.4 ± 1.3 | 98.0 ± 0.6 | 98.5 ± 0.6 | 101.7 ± 0.4 |
| 2'-Deoxycytidine-5'-monophosphate | 109.6 ± 0.9 | 103.6 ± 0.6 | 100.2 ± 0.5 | 105.0 ± 1.9 |
| 2-Deoxyguanosine | 105.6 ± 1.0 | 97.3 ± 1.3 | 98.2 ± 1.2 | 105.2 ± 0.9 |
| 2-hydroxybutyric acid | 107.7 ± 1.6 | 85.6 ± 1.2 | 90.9 ± 0.8 | 101.5 ± 1.1 |
| 2-hydroxyglutaric acid | 114.7 ± 0.9 | 95.2 ± 1.4 | 101.2 ± 2.0 | 95.6 ± 1.4 |
| 2-Hydroxyisocaproic acid | 88.4 ± 1.1 | 109.0 ± 0.4 | 89.9 ± 1.5 | 89.0 ± 0.3 |
| 2-ketobutyric acid | 92.6 ± 1.6 | 99.0 ± 1.1 | 96.6 ± 0.6 | 85.8 ± 0.8 |
| 2-oxoadipic acid | 93.2 ± 1.0 | 103.7 ± 1.5 | 104.5 ± 1.2 | 103.8 ± 0.8 |
| 2-Phenylglycine | 94.6 ± 0.7 | 101.8 ± 2.0 | 104.9 ± 1.8 | 104.1 ± 1.6 |
| 2-phosphoglycerate | 110.9 ± 1.8 | 91.9 ± 1.1 | 94.5 ± 1.7 | 86.8 ± 0.8 |
| 3-hydroxy-3-methyl-2-oxobutanoate | 93.0 ± 1.6 | 104.8 ± 1.5 | 96.4 ± 0.7 | 104.3 ± 0.4 |
| 3-hydroxybutyric acid | 94.2 ± 0.4 | 100.2 ± 0.6 | 88.2 ± 0.8 | 94.5 ± 0.8 |
| 3-Methylhistidine | 97.6 ± 1.0 | 100.9 ± 0.3 | 104.8 ± 1.8 | 97.4 ± 0.3 |
| 3-Phosphoglyceraldehyde | 87.2 ± 0.7 | 92.8 ± 0.3 | 90.2 ± 0.4 | 86.0 ± 1.4 |
| 3-phosphoglycerate | 101.7 ± 0.3 | 100.8 ± 1.0 | 105.0 ± 1.5 | 106.5 ± 0.4 |
| 4-amino-butyric acid | 93.7 ± 0.4 | 104.3 ± 1.0 | 100.4 ± 0.6 | 86.9 ± 1.4 |
| 4-aspartyl-phosphate | 100.6 ± 1.5 | 99.7 ± 1.0 | 98.4 ± 1.4 | 103.7 ± 1.0 |
| 4-Hydroxybenzoic acid | 89.0 ± 0.7 | 85.1 ± 0.6 | 93.4 ± 0.5 | 109.5 ± 1.6 |
| 4-Hydroxyphenylacetylglycine | 92.2 ± 1.1 | 106.5 ± 0.5 | 101.1 ± 1.6 | 87.5 ± 1.4 |
| 4-methyl-2-oxopentanoate | 105.8 ± 0.4 | 110.0 ± 1.9 | 91.7 ± 0.4 | 92.5 ± 1.5 |
| 4-Oxoglutaramate | 109.6 ± 1.4 | 90.7 ± 1.8 | 96.8 ± 0.3 | 114.1 ± 0.4 |
| 5-Hydroxyindolepyruvate | 111.1 ± 1.9 | 103.0 ± 1.2 | 99.1 ± 1.2 | 93.3 ± 0.5 |
| 5-Hydroxylysine | 107.0 ± 0.3 | 95.2 ± 1.3 | 104.1 ± 0.3 | 104.4 ± 0.6 |
| 5-Hydroxy-Tryptophan | 114.7 ± 1.8 | 112.5 ± 1.1 | 105.1 ± 0.2 | 105.1 ± 1.0 |
| 5-Phosphoribosylamine | 100.2 ± 1.6 | 102.2 ± 0.4 | 98.1 ± 0.7 | 98.1 ± 0.2 |
| 6-phosphogluconate | 103.2 ± 0.8 | 94.9 ± 1.9 | 102.3 ± 1.2 | 89.4 ± 1.6 |
| Acetylcarnitine | 110.6 ± 1.2 | 99.2 ± 1.3 | 95.1 ± 1.0 | 113.3 ± 1.8 |
| Aconitic Acid | 96.9 ± 0.8 | 101.6 ± 0.8 | 97.3 ± 1.1 | 97.3 ± 0.2 |
| Adenine | 94.6 ± 1.3 | 100.8 ± 1.1 | 97.4 ± 0.8 | 99.1 ± 1.8 |
| Adenosine | 111.1 ± 0.4 | 104.3 ± 0.3 | 114.6 ± 0.4 | 97.6 ± 1.4 |
| Adenosine 3',5'-cyclic monophosphate | 85.8 ± 0.4 | 92.2 ± 0.8 | 94.5 ± 1.4 | 85.3 ± 1.0 |
| Adenosine Monophosphate | 110.2 ± 0.3 | 86.1 ± 0.4 | 94.4 ± 1.7 | 100.3 ± 1.4 |
| Adenosine-5'-diphosphate | 102.7 ± 0.2 | 91.5 ± 1.3 | 96.6 ± 0.2 | 90.5 ± 1.6 |
| Adenosine-5'-triphosphate | 108.0 ± 0.6 | 88.4 ± 0.6 | 90.4 ± 1.0 | 86.3 ± 0.4 |
| Adenylosuccinate | 92.1 ± 1.5 | 93.6 ± 1.4 | 114.4 ± 1.4 | 111.8 ± 1.4 |
| Alanine | 104.2 ± 1.1 | 99.2 ± 0.3 | 86.4 ± 0.6 | 93.5 ± 0.8 |
| alpha-ketoglutaric acid | 98.2 ± 0.5 | 101.3 ± 1.3 | 108.9 ± 0.3 | 85.1 ± 0.8 |
| Anserine | 99.5 ± 1.0 | 96.1 ± 1.1 | 103.2 ± 1.8 | 98.0 ± 0.9 |
| Arginine | 105.0 ± 1.3 | 102.1 ± 1.3 | 94.2 ± 0.4 | 99.3 ± 0.5 |
| Argininosuccinic Acid | 98.2 ± 0.9 | 104.0 ± 2.0 | 93.2 ± 0.3 | 114.4 ± 1.3 |
| Asparagine | 89.0 ± 1.2 | 102.7 ± 1.0 | 105.9 ± 0.3 | 101.9 ± 1.9 |
| Aspartic acid | 91.5 ± 1.4 | 96.3 ± 1.1 | 88.3 ± 0.4 | 97.8 ± 1.8 |
| Asymmetric dimethylarginine | 108.7 ± 0.9 | 101.5 ± 0.4 | 103.2 ± 1.9 | 104.3 ± 0.4 |
| Azelaic acid | 95.8 ± 1.7 | 102.6 ± 0.8 | 101.6 ± 0.4 | 98.9 ± 0.6 |
| beta-Alanine | 112.4 ± 0.5 | 97.7 ± 1.7 | 100.3 ± 1.6 | 99.7 ± 0.4 |
| Betaine | 97.4 ± 1.0 | 104.6 ± 0.2 | 97.5 ± 0.7 | 98.0 ± 0.8 |
| Biopterin | 111.7 ± 0.9 | 104.9 ± 1.2 | 100.0 ± 1.1 | 100.3 ± 2.0 |
| Biotin | 91.4 ± 0.7 | 100.2 ± 0.5 | 100.1 ± 0.3 | 103.3 ± 0.4 |
| Carnitine | 104.7 ± 0.3 | 98.3 ± 1.6 | 97.0 ± 1.4 | 101.0 ± 1.7 |
| Carnosine | 90.2 ± 1.6 | 100.6 ± 0.6 | 98.9 ± 1.3 | 100.8 ± 1.1 |
| CDP-choline | 96.5 ± 1.3 | 99.0 ± 1.8 | 105.8 ± 0.9 | 101.6 ± 0.9 |
| cGMP | 112.8 ± 0.9 | 85.9 ± 0.2 | 91.2 ± 1.4 | 95.8 ± 0.7 |
| Choline | 100.2 ± 1.6 | 104.4 ± 1.9 | 105.8 ± 0.3 | 105.6 ± 1.6 |
| Cinnamic acid | 106.0 ± 1.9 | 112.8 ± 0.2 | 113.3 ± 1.8 | 110.1 ± 0.4 |
| Citraconic acid | 96.5 ± 1.0 | 87.7 ± 1.4 | 106.8 ± 1.0 | 99.2 ± 1.1 |
| Citramalic acid | 110.6 ± 0.8 | 105.4 ± 1.3 | 108.2 ± 1.7 | 106.5 ± 1.7 |
| Citric Acid | 112.4 ± 0.6 | 98.0 ± 0.4 | 105.9 ± 0.6 | 101.2 ± 1.0 |
| Citrulline | 96.0 ± 1.4 | 95.7 ± 1.8 | 96.6 ± 1.0 | 97.9 ± 1.9 |
| Creatine | 87.2 ± 0.4 | 88.5 ± 1.4 | 114.8 ± 0.5 | 112.2 ± 1.5 |
| Creatinine | 106.8 ± 0.8 | 102.0 ± 0.5 | 102.5 ± 1.1 | 104.6 ± 0.9 |
| Cystathionine | 94.5 ± 0.5 | 99.0 ± 0.6 | 107.1 ± 1.8 | 98.7 ± 0.4 |
| Cysteamine | 100.5 ± 1.3 | 103.6 ± 1.8 | 101.4 ± 0.6 | 102.1 ± 2.0 |
| Cysteine | 107.0 ± 1.4 | 99.1 ± 1.0 | 104.1 ± 1.3 | 97.4 ± 1.8 |
| Cysteinylglycine | 93.9 ± 0.9 | 107.4 ± 1.0 | 95.6 ± 1.8 | 98.0 ± 1.0 |
| Cystine | 86.1 ± 1.5 | 96.3 ± 0.6 | 110.5 ± 1.1 | 113.0 ± 0.8 |
| Cytidine | 110.9 ± 0.8 | 96.9 ± 0.7 | 103.6 ± 0.3 | 106.7 ± 1.6 |
| Cytidine 3',5'-cyclic monophosphate | 101.5 ± 0.4 | 103.6 ± 1.6 | 104.1 ± 1.9 | 103.0 ± 0.4 |
| Cytidine 5'-diphosphate | 100.2 ± 0.6 | 103.3 ± 1.6 | 96.6 ± 1.3 | 99.4 ± 0.3 |
| Cytidine 5'-triphosphate | 101.9 ± 1.8 | 102.0 ± 0.9 | 101.5 ± 0.6 | 100.6 ± 1.8 |
| Cytidine monophosphate | 102.9 ± 1.7 | 97.4 ± 1.9 | 96.1 ± 0.4 | 100.0 ± 1.3 |
| Cytosine | 114.8 ± 0.4 | 99.8 ± 1.5 | 99.0 ± 0.8 | 88.8 ± 0.4 |
| dADP | 95.6 ± 1.2 | 98.4 ± 1.1 | 103.1 ± 0.9 | 99.1 ± 0.9 |
| Diethanolamine | 104.7 ± 1.0 | 100.4 ± 1.0 | 104.6 ± 1.9 | 114.7 ± 0.5 |
| Dihydrobiopterin | 106.6 ± 1.5 | 103.1 ± 1.3 | 99.5 ± 1.3 | 98.7 ± 1.2 |
| Dihydroneopterin triphosphate | 104.0 ± 0.7 | 102.2 ± 0.2 | 106.0 ± 0.6 | 103.6 ± 0.9 |
| Dihydroxyacetonephosphate | 94.8 ± 0.9 | 104.7 ± 0.9 | 87.0 ± 1.5 | 90.7 ± 1.4 |
| Dimethylglycine | 111.4 ± 1.0 | 113.5 ± 1.3 | 104.4 ± 1.4 | 90.6 ± 1.7 |
| Ectoine | 98.5 ± 0.8 | 98.7 ± 1.3 | 104.3 ± 1.6 | 103.7 ± 1.4 |
| Erythrose-4-phosphate | 90.5 ± 1.0 | 106.7 ± 1.8 | 109.9 ± 1.9 | 89.7 ± 0.9 |
| FAD | 96.5 ± 0.3 | 102.2 ± 1.8 | 96.3 ± 1.4 | 102.2 ± 0.5 |
| Folic acid | 111.4 ± 1.3 | 104.7 ± 1.5 | 100.9 ± 1.8 | 102.1 ± 0.9 |
| Fructose-6-phosphate | 109.9 ± 1.9 | 102.3 ± 0.7 | 101.1 ± 0.7 | 103.2 ± 1.6 |
| Fumaric Acid | 102.9 ± 0.9 | 86.5 ± 0.8 | 112.1 ± 2.0 | 88.0 ± 1.3 |
| Galactosamine | 94.5 ± 0.3 | 103.6 ± 1.3 | 102.2 ± 1.7 | 99.5 ± 0.7 |
| Galactose-1-phosphate | 90.2 ± 1.6 | 110.5 ± 2.0 | 90.3 ± 0.8 | 96.0 ± 0.6 |
| Galactose-6-Phosphate | 86.1 ± 0.5 | 99.4 ± 1.7 | 96.7 ± 0.9 | 104.9 ± 0.8 |
| gamma-Glutamylcysteine | 92.5 ± 1.8 | 95.9 ± 1.5 | 100.7 ± 2.0 | 106.1 ± 0.9 |
| Glucosamine | 112.9 ± 0.9 | 94.8 ± 1.3 | 87.1 ± 1.7 | 89.0 ± 1.0 |
| Glucosamine-6-Phosphate | 111.7 ± 0.4 | 98.3 ± 0.3 | 103.6 ± 0.2 | 100.1 ± 1.4 |
| Glucose | 114.3 ± 0.2 | 99.5 ± 0.8 | 92.0 ± 0.5 | 102.1 ± 1.9 |
| Glucose-1-phosphate | 88.8 ± 0.8 | 101.6 ± 1.3 | 101.0 ± 1.1 | 105.6 ± 1.7 |
| Glucose-6-Phosphate | 95.0 ± 1.8 | 99.2 ± 0.7 | 97.1 ± 0.5 | 100.7 ± 1.1 |
| Glucuronic acid | 114.0 ± 0.7 | 100.5 ± 2.0 | 99.1 ± 0.2 | 105.6 ± 0.5 |
| Glutamic acid | 90.1 ± 1.7 | 103.9 ± 1.4 | 98.5 ± 1.3 | 106.2 ± 1.4 |
| Glutamine | 105.4 ± 1.2 | 96.4 ± 1.8 | 100.9 ± 0.7 | 101.7 ± 0.9 |
| Glutathione | 93.7 ± 0.9 | 99.5 ± 0.7 | 102.3 ± 1.3 | 102.7 ± 0.9 |
| Glutathionylspermine | 103.0 ± 1.4 | 98.7 ± 1.8 | 109.8 ± 1.8 | 107.9 ± 2.0 |
| Glycerophosphocholine | 102.1 ± 0.7 | 101.1 ± 0.7 | 100.2 ± 1.5 | 102.9 ± 2.0 |
| Glycine | 111.7 ± 0.6 | 95.6 ± 0.8 | 106.4 ± 0.6 | 95.5 ± 1.6 |
| Glycolic acid | 98.8 ± 0.3 | 91.6 ± 1.5 | 97.2 ± 0.2 | 86.3 ± 1.2 |
| Guanidinoacetic acid | 90.9 ± 0.2 | 104.0 ± 1.8 | 97.5 ± 1.7 | 101.9 ± 2.0 |
| Guanidinobutanoic acid | 91.8 ± 0.9 | 104.0 ± 1.7 | 103.2 ± 2.0 | 104.5 ± 1.0 |
| Guanine | 96.1 ± 1.1 | 113.1 ± 0.8 | 98.2 ± 0.8 | 107.0 ± 0.3 |
| Guanosine | 102.2 ± 1.6 | 95.6 ± 1.1 | 99.6 ± 0.2 | 102.2 ± 1.7 |
| Guanosine 5'-diphosphate | 97.6 ± 1.6 | 95.7 ± 0.5 | 98.5 ± 1.2 | 104.4 ± 1.5 |
| Guanosine-5'-monophosphate | 111.6 ± 0.9 | 100.1 ± 1.7 | 105.0 ± 1.6 | 105.3 ± 1.1 |
| Hercynine | 111.1 ± 1.8 | 100.8 ± 1.1 | 104.7 ± 1.5 | 100.2 ± 1.4 |
| Hippuric acid | 95.7 ± 1.4 | 100.8 ± 1.8 | 102.6 ± 0.4 | 99.8 ± 1.7 |
| Histamine | 97.0 ± 1.0 | 104.0 ± 1.8 | 98.8 ± 1.4 | 103.2 ± 0.5 |
| Histidine | 105.9 ± 1.0 | 104.2 ± 1.4 | 102.4 ± 1.1 | 97.0 ± 1.8 |
| Homoalanine | 100.1 ± 0.8 | 94.1 ± 1.2 | 100.5 ± 0.3 | 96.0 ± 0.9 |
| Homoarginine | 107.1 ± 0.8 | 91.5 ± 1.3 | 94.2 ± 0.7 | 95.5 ± 1.6 |
| Homocysteine | 91.4 ± 0.9 | 96.4 ± 0.2 | 97.1 ± 1.2 | 106.3 ± 1.3 |
| Homogentisic acid | 90.3 ± 1.6 | 101.6 ± 1.5 | 102.4 ± 1.9 | 104.5 ± 0.5 |
| Homoleucine | 87.2 ± 0.4 | 112.6 ± 1.8 | 93.4 ± 1.3 | 86.9 ± 0.8 |
| Homovanillic acid | 113.2 ± 0.6 | 92.9 ± 1.6 | 99.9 ± 1.3 | 113.1 ± 0.6 |
| Hydantoin-5-propionate | 89.5 ± 0.8 | 104.2 ± 1.4 | 104.2 ± 1.3 | 100.9 ± 1.3 |
| Hypotaurine | 111.4 ± 0.9 | 104.0 ± 1.9 | 113.3 ± 1.2 | 101.0 ± 1.5 |
| Hypoxanthine | 102.1 ± 0.3 | 92.4 ± 1.3 | 92.7 ± 1.3 | 109.1 ± 1.6 |
| Indole-3-acetamide | 98.4 ± 0.5 | 85.6 ± 1.0 | 111.1 ± 1.0 | 88.3 ± 0.9 |
| Inosine | 99.0 ± 0.8 | 104.6 ± 0.7 | 101.1 ± 0.3 | 105.4 ± 1.8 |
| Inosine 5'-monophosphate | 95.8 ± 1.9 | 102.0 ± 0.9 | 90.9 ± 0.6 | 92.2 ± 0.3 |
| Isocitric Acid | 99.5 ± 1.9 | 97.9 ± 1.4 | 105.3 ± 0.7 | 98.1 ± 1.2 |
| Isoleucine | 90.1 ± 1.7 | 97.0 ± 0.9 | 105.0 ± 1.4 | 101.3 ± 1.1 |
| Lactic Acid | 104.6 ± 0.9 | 100.9 ± 0.9 | 96.7 ± 0.5 | 97.6 ± 1.8 |
| Leucine | 103.8 ± 0.5 | 112.0 ± 0.4 | 85.3 ± 0.6 | 93.2 ± 1.7 |
| Levulinic acid | 110.7 ± 0.8 | 105.0 ± 0.4 | 94.2 ± 0.4 | 88.4 ± 0.3 |
| Lysine | 100.1 ± 1.2 | 108.4 ± 1.2 | 92.7 ± 1.4 | 104.4 ± 1.2 |
| Malic Acid | 100.7 ± 0.9 | 96.1 ± 1.6 | 96.4 ± 1.6 | 103.6 ± 1.9 |
| Mannose-1 with Mannose-6-Phosphate | 103.5 ± 0.7 | 104.9 ± 0.8 | 102.9 ± 0.8 | 97.5 ± 1.9 |
| Mesaconic acid | 100.4 ± 0.7 | 102.9 ± 1.4 | 103.0 ± 0.6 | 104.2 ± 1.1 |
| Methionine | 113.0 ± 1.1 | 104.0 ± 0.5 | 98.5 ± 1.4 | 103.2 ± 0.8 |
| Methionine sulfoxide | 93.9 ± 0.7 | 114.7 ± 1.8 | 98.3 ± 0.8 | 110.8 ± 1.4 |
| Methylmalonic Acid | 91.6 ± 0.9 | 102.0 ± 1.6 | 104.0 ± 1.6 | 105.2 ± 1.2 |
| N1-Acetylspermidine | 103.4 ± 0.7 | 104.5 ± 0.5 | 105.1 ± 1.1 | 94.7 ± 1.7 |
| N1-Acetylspermine | 110.4 ± 1.7 | 105.0 ± 0.2 | 86.9 ± 0.8 | 92.6 ± 1.6 |
| N8-Acetylspermidine | 106.6 ± 0.3 | 99.9 ± 0.8 | 111.6 ± 0.6 | 90.6 ± 1.4 |
| N-Acetyl-Aspartyl-Glutamate | 109.7 ± 1.1 | 103.5 ± 1.4 | 97.0 ± 0.3 | 100.2 ± 0.5 |
| N-Acetyl-Aspartyl-Glutamyl-Glutamate | 108.8 ± 1.6 | 103.2 ± 0.3 | 101.0 ± 0.5 | 100.0 ± 1.2 |
| N-Acetyl-Galactosamine | 86.5 ± 0.8 | 104.3 ± 0.5 | 101.5 ± 1.7 | 106.4 ± 1.6 |
| N-Acetyl-Leucine | 86.4 ± 1.1 | 96.2 ± 1.3 | 111.5 ± 1.7 | 109.5 ± 1.3 |
| N-Acetyl-Methionine | 101.7 ± 1.7 | 100.8 ± 0.3 | 97.2 ± 0.3 | 102.1 ± 1.5 |
| N-Acetylneuraminic acid | 113.7 ± 0.5 | 101.5 ± 0.8 | 99.4 ± 1.3 | 103.9 ± 0.3 |
| N-Acetyl-Proline | 103.0 ± 0.9 | 99.4 ± 0.4 | 97.3 ± 0.6 | 103.8 ± 1.5 |
| N-Acetyl-Putrescine | 104.6 ± 0.9 | 99.1 ± 1.5 | 105.5 ± 0.4 | 98.4 ± 1.8 |
| N-Acetyl-Serine | 94.8 ± 1.6 | 103.3 ± 0.3 | 104.6 ± 1.4 | 99.9 ± 0.6 |
| N-Acetyl-Serotonin | 88.1 ± 0.4 | 100.7 ± 1.5 | 103.9 ± 0.4 | 97.9 ± 1.9 |
| N-Acyl-Homoserine-lactone | 102.4 ± 1.8 | 96.1 ± 1.8 | 104.1 ± 1.6 | 103.1 ± 1.4 |
| NAD | 105.4 ± 0.9 | 90.2 ± 1.1 | 107.4 ± 0.9 | 101.6 ± 0.5 |
| NADH | 103.3 ± 1.8 | 96.8 ± 1.0 | 98.9 ± 1.6 | 105.5 ± 0.7 |
| NADP | 92.6 ± 0.4 | 87.3 ± 1.3 | 98.3 ± 0.5 | 91.9 ± 1.2 |
| NADPH | 102.3 ± 0.6 | 88.6 ± 1.6 | 86.6 ± 0.3 | 113.9 ± 1.8 |
| N-alpha-Acetyl-lysine | 109.0 ± 1.3 | 98.6 ± 1.5 | 105.3 ± 1.7 | 101.9 ± 1.6 |
| N-alpha-Acetyl-Ornithine | 104.9 ± 1.2 | 100.7 ± 0.3 | 96.7 ± 0.8 | 106.5 ± 1.9 |
| N-Carbamoyl-Proline | 98.8 ± 1.7 | 103.5 ± 1.2 | 105.2 ± 1.9 | 103.8 ± 0.7 |
| N-Carbamoyl-Sarcosine | 89.3 ± 0.6 | 103.6 ± 1.6 | 96.5 ± 0.5 | 98.4 ± 0.4 |
| N-Epsilon-Acetyl-lysine | 103.2 ± 2.0 | 99.3 ± 1.5 | 104.0 ± 1.6 | 106.4 ± 1.7 |
| Nicotinamide | 104.2 ± 1.2 | 102.8 ± 0.3 | 102.5 ± 1.3 | 102.1 ± 0.9 |
| N-Methyl-Aspartic Acid | 101.4 ± 1.4 | 96.9 ± 1.1 | 98.1 ± 0.4 | 98.4 ± 2.0 |
| O-Phosphoethanolamine | 89.1 ± 1.7 | 103.7 ± 1.9 | 103.4 ± 0.7 | 97.1 ± 1.7 |
| Ophthalmic acid | 96.2 ± 1.6 | 103.1 ± 0.6 | 105.4 ± 1.9 | 99.6 ± 1.8 |
| Ornithine | 97.7 ± 1.0 | 103.9 ± 0.9 | 104.0 ± 0.9 | 97.8 ± 1.4 |
| Orotidine 5'-monophosphate | 94.7 ± 1.5 | 102.5 ± 0.4 | 104.5 ± 1.3 | 98.2 ± 1.5 |
| Oxaloacetate | 106.2 ± 1.5 | 98.4 ± 1.7 | 104.0 ± 1.5 | 105.8 ± 1.7 |
| Pantothenic acid | 100.1 ± 1.1 | 113.5 ± 1.4 | 103.5 ± 0.8 | 99.7 ± 1.9 |
| Phenylalanine | 108.1 ± 0.4 | 100.8 ± 0.7 | 100.8 ± 0.2 | 92.5 ± 0.4 |
| Phenylpyruvate | 96.2 ± 0.5 | 98.1 ± 0.6 | 105.6 ± 1.9 | 105.2 ± 0.4 |
| Phosphodimethylethanolamine | 106.2 ± 1.9 | 103.0 ± 1.2 | 103.0 ± 1.8 | 85.4 ± 0.5 |
| Phosphoenolpyruvate | 102.8 ± 1.3 | 104.3 ± 1.3 | 105.5 ± 0.6 | 100.4 ± 0.3 |
| Phosphorylcholine | 111.1 ± 1.8 | 104.9 ± 1.5 | 97.4 ± 0.2 | 100.2 ± 0.9 |
| Pipecolic acid | 113.5 ± 0.4 | 91.0 ± 1.2 | 94.0 ± 0.6 | 101.2 ± 1.8 |
| Proline | 95.8 ± 1.6 | 103.7 ± 0.6 | 100.6 ± 0.7 | 97.4 ± 0.8 |
| Putrescine | 108.4 ± 1.7 | 105.0 ± 1.3 | 88.6 ± 0.7 | 107.6 ± 1.2 |
| Pyridoxal | 111.6 ± 1.4 | 104.8 ± 0.3 | 102.5 ± 1.0 | 99.9 ± 0.4 |
| Pyridoxamine | 88.6 ± 1.7 | 101.2 ± 1.0 | 102.4 ± 0.3 | 103.3 ± 1.9 |
| Pyridoxine | 107.5 ± 0.8 | 104.5 ± 0.3 | 103.9 ± 0.8 | 98.4 ± 1.6 |
| Pyroglutamic acid | 99.2 ± 1.8 | 102.8 ± 0.5 | 98.6 ± 0.3 | 101.6 ± 1.6 |
| Pyrrolysine | 101.4 ± 1.1 | 102.1 ± 1.4 | 100.1 ± 1.1 | 105.0 ± 1.7 |
| Pyruvic Acid | 109.0 ± 1.4 | 98.8 ± 0.8 | 99.1 ± 1.9 | 105.2 ± 0.8 |
| Riboflavin | 101.3 ± 0.6 | 85.2 ± 0.7 | 106.2 ± 0.6 | 96.9 ± 1.6 |
| Ribose-5-Phosphate | 103.0 ± 0.7 | 98.6 ± 0.7 | 94.7 ± 0.5 | 113.2 ± 0.2 |
| Ribulose-1-Phosphate | 93.2 ± 1.0 | 96.4 ± 1.4 | 97.9 ± 1.0 | 101.6 ± 0.4 |
| Ribulose-5-Phosphate | 94.3 ± 1.0 | 103.8 ± 1.1 | 102.8 ± 0.2 | 101.8 ± 1.2 |
| Saccharopine | 92.8 ± 1.1 | 100.4 ± 1.2 | 104.3 ± 1.2 | 101.9 ± 0.4 |
| S-Adenosylhomocysteine | 94.4 ± 0.5 | 99.6 ± 0.3 | 106.0 ± 0.6 | 104.4 ± 1.7 |
| S-Adenosylmethioninamine | 96.2 ± 1.8 | 100.4 ± 0.8 | 105.6 ± 1.8 | 100.1 ± 0.3 |
| S-Adenosylmethionine | 88.9 ± 1.7 | 87.4 ± 0.3 | 94.6 ± 0.6 | 112.4 ± 1.7 |
| Sarcosine | 108.9 ± 0.6 | 101.4 ± 1.2 | 94.5 ± 0.7 | 108.6 ± 1.3 |
| Sedoheptulose-7-phosphate | 88.4 ± 1.1 | 87.5 ± 1.1 | 100.5 ± 1.9 | 112.4 ± 1.6 |
| Selenohomocysteine | 100.7 ± 1.3 | 87.7 ± 1.5 | 100.1 ± 1.5 | 96.8 ± 0.5 |
| Selenomethionine | 110.0 ± 0.6 | 98.9 ± 1.3 | 85.9 ± 1.6 | 101.0 ± 0.4 |
| Serine | 95.7 ± 1.1 | 113.2 ± 1.9 | 110.8 ± 0.3 | 100.5 ± 0.3 |
| S-Methyl-Thioinosine | 97.7 ± 1.9 | 85.6 ± 0.2 | 97.9 ± 1.5 | 95.1 ± 1.3 |
| Spermidine | 91.6 ± 1.7 | 100.7 ± 1.4 | 88.1 ± 1.8 | 106.7 ± 2.0 |
| Spermine | 89.4 ± 0.3 | 98.0 ± 1.3 | 96.2 ± 0.8 | 100.3 ± 1.2 |
| Succinate | 110.7 ± 0.7 | 104.1 ± 0.9 | 97.8 ± 0.7 | 97.9 ± 1.1 |
| Sucrose | 103.2 ± 1.3 | 99.6 ± 1.8 | 102.0 ± 1.3 | 105.8 ± 1.9 |
| Symmetric dimethylarginine | 99.9 ± 0.9 | 90.2 ± 0.9 | 93.4 ± 0.6 | 95.2 ± 1.1 |
| Taurine | 102.4 ± 1.2 | 109.2 ± 1.5 | 104.9 ± 0.7 | 88.2 ± 1.4 |
| Tetrahydroneopterin | 90.4 ± 0.7 | 89.8 ± 0.5 | 93.5 ± 0.3 | 94.2 ± 1.9 |
| Thiamine | 108.1 ± 1.4 | 113.5 ± 1.7 | 85.9 ± 0.9 | 106.2 ± 1.6 |
| Thiamine monophosphate | 111.1 ± 1.7 | 105.6 ± 1.4 | 93.1 ± 0.9 | 86.1 ± 0.6 |
| Threonic acid | 114.5 ± 2.0 | 88.3 ± 1.0 | 94.1 ± 1.4 | 94.1 ± 0.6 |
| Threonine | 106.2 ± 1.0 | 105.7 ± 1.7 | 97.1 ± 0.8 | 93.4 ± 0.8 |
| Thymidine | 108.2 ± 1.4 | 98.4 ± 1.9 | 92.5 ± 1.6 | 91.7 ± 1.7 |
| Thymidine Monophosphate | 109.0 ± 0.4 | 103.4 ± 1.0 | 99.4 ± 0.7 | 102.5 ± 0.4 |
| Thymine | 106.9 ± 0.6 | 96.5 ± 1.0 | 104.5 ± 0.7 | 103.2 ± 2.0 |
| Trigonelleine | 111.4 ± 0.2 | 100.0 ± 1.0 | 99.6 ± 1.9 | 104.6 ± 2.0 |
| Trimethylamine N-oxide | 99.6 ± 0.3 | 97.5 ± 0.8 | 98.6 ± 1.0 | 105.8 ± 1.8 |
| Trimethyllysine | 98.8 ± 1.2 | 97.1 ± 0.6 | 102.1 ± 1.4 | 106.3 ± 0.8 |
| Tryptophan | 99.2 ± 1.6 | 96.7 ± 0.9 | 97.9 ± 1.9 | 100.0 ± 0.5 |
| Tyramine | 105.4 ± 2.0 | 103.9 ± 0.7 | 100.1 ± 0.5 | 99.3 ± 0.9 |
| Tyrosine | 113.3 ± 1.1 | 98.8 ± 2.0 | 104.9 ± 1.9 | 102.3 ± 0.6 |
| Uracil | 106.6 ± 1.5 | 107.0 ± 1.0 | 86.7 ± 1.4 | 111.8 ± 1.7 |
| Uridine 5'-diphosphate | 91.8 ± 1.8 | 96.3 ± 0.5 | 110.2 ± 1.5 | 111.6 ± 0.7 |
| Uridine Monophosphate | 111.3 ± 1.5 | 112.5 ± 0.6 | 108.5 ± 1.0 | 94.4 ± 1.3 |
| Uridine-5-Diphosphogalactose | 98.7 ± 0.6 | 104.7 ± 0.4 | 89.2 ± 0.9 | 96.4 ± 0.6 |
| Urocanic acid | 101.3 ± 0.9 | 99.3 ± 1.6 | 105.8 ± 1.4 | 105.5 ± 1.5 |
| UTP | 96.8 ± 1.5 | 103.5 ± 0.3 | 101.1 ± 0.9 | 103.0 ± 1.8 |
| Valine | 111.8 ± 1.8 | 103.9 ± 0.6 | 100.9 ± 2.0 | 105.9 ± 0.6 |
| XMP | 96.5 ± 0.4 | 100.8 ± 1.5 | 99.8 ± 1.9 | 102.5 ± 0.7 |
| Xylulose-5-Phosphate | 97.8 ± 1.4 | 103.3 ± 1.7 | 105.4 ± 0.7 | 106.8 ± 1.1 |

### **Table S28** | Inter – day accuracy (as determined by % recovery) (Tumor tissue extract as matrix, n = 5). The concentrations of QC_LLOQ_, QC_Low_, QC_Mid_ and QC_High_ were 10 ng/mL, 25 ng/mL, 100 ng/mL and 500 ng/mL respectively.

| Accuracy, % | QC_LLOQ_ | QC_Low_ | QC_Mid_ | QC_High_ |
| --- | --- | --- | --- | --- |
| 1-Methylguanine | 97.8 ± 1.3 | 109.2 ± 1.9 | 98.6 ± 4.8 | 106.3 ± 1.8 |
| 1-Methylhistidine | 99.2 ± 5.4 | 98.3 ± 5.0 | 106.3 ± 0.1 | 101.3 ± 1.6 |
| 1-Methylnicotinamide | 104.2 ± 1.5 | 91.6 ± 4.0 | 103.2 ± 2.1 | 91.0 ± 3.9 |
| 2-Aminoadipic Acid | 103.5 ± 5.1 | 98.1 ± 1.4 | 104.2 ± 4.4 | 107.2 ± 4.2 |
| 2-Deoxyadenosine | 89.3 ± 0.8 | 99.5 ± 3.3 | 108.2 ± 4.4 | 105.3 ± 3.5 |
| 2'-Deoxyadenosine 5'-monophosphate | 90.3 ± 4.7 | 95.5 ± 3.8 | 103.7 ± 4.4 | 109.8 ± 4.8 |
| 2'-deoxyadenosine 5'-triphosphate | 86.4 ± 1.6 | 93.7 ± 3.1 | 107.6 ± 3.6 | 100.0 ± 5.9 |
| 2'-Deoxycytidine-5'-monophosphate | 85.7 ± 6.1 | 109.7 ± 5.6 | 99.8 ± 6.1 | 91.4 ± 5.5 |
| 2-Deoxyguanosine | 95.5 ± 3.9 | 101.6 ± 4.8 | 94.2 ± 2.5 | 99.2 ± 1.1 |
| 2-hydroxybutyric acid | 97.0 ± 4.5 | 94.1 ± 2.0 | 103.1 ± 2.7 | 99.7 ± 4.1 |
| 2-hydroxyglutaric acid | 89.2 ± 0.5 | 90.4 ± 3.9 | 92.3 ± 2.9 | 99.4 ± 3.1 |
| 2-Hydroxyisocaproic acid | 87.9 ± 2.5 | 101.6 ± 1.7 | 95.5 ± 0.8 | 92.2 ± 0.9 |
| 2-ketobutyric acid | 98.8 ± 2.0 | 108.9 ± 1.7 | 107.4 ± 6.0 | 100.7 ± 0.2 |
| 2-oxoadipic acid | 86.0 ± 2.2 | 109.6 ± 5.9 | 94.9 ± 2.3 | 98.6 ± 5.4 |
| 2-Phenylglycine | 100.2 ± 1.8 | 90.9 ± 0.8 | 102.8 ± 3.4 | 105.7 ± 4.8 |
| 2-phosphoglycerate | 86.2 ± 1.3 | 91.7 ± 0.4 | 91.6 ± 3.6 | 108.7 ± 2.8 |
| 3-hydroxy-3-methyl-2-oxobutanoate | 98.1 ± 3.0 | 103.0 ± 6.2 | 96.2 ± 2.1 | 109.1 ± 4.3 |
| 3-hydroxybutyric acid | 86.0 ± 2.0 | 95.7 ± 4.8 | 102.3 ± 6.2 | 102.9 ± 2.3 |
| 3-Methylhistidine | 102.7 ± 4.1 | 102.6 ± 0.4 | 107.9 ± 6.1 | 107.9 ± 0.5 |
| 3-Phosphoglyceraldehyde | 91.4 ± 3.0 | 97.2 ± 2.8 | 92.6 ± 6.2 | 92.3 ± 2.3 |
| 3-phosphoglycerate | 89.6 ± 6.1 | 103.2 ± 5.3 | 97.1 ± 1.1 | 95.4 ± 4.9 |
| 4-amino-butyric acid | 98.1 ± 4.7 | 106.8 ± 2.6 | 97.7 ± 1.1 | 94.9 ± 6.1 |
| 4-aspartyl-phosphate | 90.8 ± 2.8 | 90.6 ± 0.1 | 90.4 ± 2.2 | 95.9 ± 1.3 |
| 4-Hydroxybenzoic acid | 92.7 ± 3.0 | 103.4 ± 5.8 | 94.8 ± 5.7 | 96.1 ± 5.4 |
| 4-Hydroxyphenylacetylglycine | 87.3 ± 1.6 | 91.4 ± 3.4 | 102.0 ± 2.2 | 108.9 ± 4.3 |
| 4-methyl-2-oxopentanoate | 101.8 ± 4.0 | 108.6 ± 0.6 | 100.1 ± 1.1 | 99.7 ± 5.5 |
| 4-Oxoglutaramate | 100.2 ± 2.2 | 92.4 ± 1.8 | 109.6 ± 1.0 | 98.7 ± 4.7 |
| 5-Hydroxyindolepyruvate | 101.4 ± 1.2 | 93.9 ± 4.2 | 95.7 ± 4.3 | 104.6 ± 2.7 |
| 5-Hydroxylysine | 91.7 ± 0.5 | 103.7 ± 1.4 | 101.8 ± 5.5 | 94.7 ± 4.5 |
| 5-Hydroxy-Tryptophan | 90.8 ± 5.7 | 107.1 ± 0.9 | 106.1 ± 2.1 | 95.1 ± 4.0 |
| 5-Phosphoribosylamine | 92.8 ± 6.2 | 100.2 ± 2.6 | 103.2 ± 1.3 | 91.0 ± 5.1 |
| 6-phosphogluconate | 95.7 ± 1.3 | 90.0 ± 2.4 | 97.6 ± 1.9 | 92.1 ± 2.4 |
| Acetylcarnitine | 94.8 ± 5.5 | 91.7 ± 3.9 | 93.5 ± 0.6 | 91.2 ± 5.7 |
| Aconitic Acid | 104.7 ± 0.6 | 98.3 ± 1.8 | 95.6 ± 0.6 | 95.3 ± 0.5 |
| Adenine | 86.1 ± 4.3 | 103.6 ± 6.1 | 90.9 ± 4.6 | 95.8 ± 3.1 |
| Adenosine | 102.5 ± 1.4 | 94.3 ± 3.6 | 100.2 ± 1.4 | 94.9 ± 3.4 |
| Adenosine 3',5'-cyclic monophosphate | 91.3 ± 3.1 | 101.6 ± 4.1 | 90.8 ± 4.9 | 108.4 ± 1.8 |
| Adenosine Monophosphate | 90.4 ± 2.6 | 98.6 ± 1.9 | 99.2 ± 4.3 | 102.2 ± 4.3 |
| Adenosine-5'-diphosphate | 98.8 ± 1.7 | 103.0 ± 0.7 | 99.2 ± 2.2 | 90.9 ± 3.8 |
| Adenosine-5'-triphosphate | 102.2 ± 1.2 | 92.2 ± 3.4 | 99.7 ± 0.2 | 109.5 ± 5.6 |
| Adenylosuccinate | 95.0 ± 4.5 | 92.0 ± 0.6 | 98.8 ± 2.0 | 106.6 ± 5.1 |
| Alanine | 94.5 ± 0.4 | 109.0 ± 3.8 | 106.9 ± 4.2 | 98.1 ± 4.7 |
| alpha-ketoglutaric acid | 102.8 ± 3.4 | 103.9 ± 0.7 | 105.0 ± 5.4 | 105.9 ± 2.8 |
| Anserine | 91.7 ± 0.4 | 98.5 ± 2.1 | 108.3 ± 0.5 | 93.4 ± 6.1 |
| Arginine | 104.0 ± 0.9 | 98.6 ± 0.8 | 109.5 ± 2.1 | 94.1 ± 3.8 |
| Argininosuccinic Acid | 95.8 ± 1.9 | 93.9 ± 4.2 | 98.3 ± 0.6 | 96.1 ± 4.6 |
| Asparagine | 91.1 ± 2.3 | 99.4 ± 3.2 | 95.7 ± 0.9 | 101.0 ± 4.6 |
| Aspartic acid | 101.2 ± 3.8 | 97.1 ± 6.0 | 94.0 ± 0.9 | 94.4 ± 2.2 |
| Asymmetric dimethylarginine | 87.5 ± 5.5 | 90.8 ± 3.0 | 94.2 ± 1.4 | 99.9 ± 5.3 |
| Azelaic acid | 86.5 ± 0.8 | 109.8 ± 6.1 | 107.2 ± 1.3 | 90.3 ± 3.3 |
| beta-Alanine | 96.2 ± 3.4 | 97.9 ± 2.6 | 99.1 ± 5.7 | 99.0 ± 2.9 |
| Betaine | 100.4 ± 4.1 | 107.7 ± 3.2 | 92.7 ± 3.2 | 109.6 ± 0.3 |
| Biopterin | 93.5 ± 0.6 | 103.3 ± 3.2 | 107.2 ± 4.8 | 91.0 ± 2.3 |
| Biotin | 94.2 ± 0.6 | 90.0 ± 2.8 | 104.7 ± 4.5 | 102.8 ± 3.8 |
| Carnitine | 104.2 ± 1.4 | 103.9 ± 5.5 | 91.3 ± 4.1 | 107.9 ± 3.0 |
| Carnosine | 96.3 ± 0.3 | 92.1 ± 3.8 | 100.9 ± 0.3 | 98.4 ± 0.3 |
| CDP-choline | 90.3 ± 4.9 | 107.3 ± 3.5 | 104.8 ± 2.1 | 109.3 ± 2.1 |
| cGMP | 87.1 ± 3.5 | 106.8 ± 3.7 | 98.9 ± 2.6 | 103.1 ± 1.8 |
| Choline | 91.4 ± 1.5 | 101.3 ± 1.4 | 108.9 ± 3.2 | 95.4 ± 2.3 |
| Cinnamic acid | 88.3 ± 4.6 | 92.2 ± 3.2 | 95.0 ± 1.7 | 109.2 ± 4.7 |
| Citraconic acid | 104.2 ± 6.2 | 101.3 ± 1.0 | 94.9 ± 0.3 | 107.5 ± 1.6 |
| Citramalic acid | 93.7 ± 2.4 | 100.0 ± 4.6 | 92.8 ± 5.5 | 101.6 ± 4.0 |
| Citric Acid | 101.8 ± 5.9 | 106.4 ± 1.7 | 102.5 ± 1.4 | 109.0 ± 5.1 |
| Citrulline | 96.6 ± 2.7 | 101.4 ± 2.9 | 98.2 ± 6.2 | 102.3 ± 4.5 |
| Creatine | 87.7 ± 0.4 | 92.2 ± 5.5 | 103.9 ± 3.3 | 106.8 ± 0.6 |
| Creatinine | 101.7 ± 3.3 | 103.4 ± 1.0 | 99.3 ± 0.3 | 105.1 ± 2.2 |
| Cystathionine | 100.3 ± 5.7 | 107.1 ± 3.5 | 91.3 ± 3.7 | 97.3 ± 3.7 |
| Cysteamine | 95.3 ± 4.3 | 92.7 ± 2.8 | 110.0 ± 5.5 | 106.7 ± 1.3 |
| Cysteine | 102.2 ± 3.2 | 94.7 ± 1.1 | 94.9 ± 4.8 | 92.5 ± 2.0 |
| Cysteinylglycine | 88.0 ± 0.9 | 109.4 ± 5.7 | 106.9 ± 4.5 | 101.7 ± 1.7 |
| Cystine | 99.5 ± 2.6 | 96.8 ± 5.8 | 108.2 ± 4.3 | 90.7 ± 4.1 |
| Cytidine | 98.4 ± 3.8 | 100.4 ± 1.3 | 95.2 ± 2.8 | 90.7 ± 1.5 |
| Cytidine 3',5'-cyclic monophosphate | 104.8 ± 2.3 | 99.6 ± 0.9 | 101.5 ± 3.6 | 92.8 ± 3.0 |
| Cytidine 5'-diphosphate | 93.9 ± 6.0 | 100.6 ± 4.7 | 100.0 ± 5.6 | 103.4 ± 4.2 |
| Cytidine 5'-triphosphate | 96.0 ± 0.5 | 101.9 ± 2.2 | 100.7 ± 1.5 | 107.5 ± 4.2 |
| Cytidine monophosphate | 88.7 ± 4.9 | 101.6 ± 2.0 | 100.5 ± 4.9 | 94.4 ± 0.7 |
| Cytosine | 91.3 ± 1.6 | 92.0 ± 4.2 | 97.6 ± 0.6 | 93.0 ± 2.5 |
| dADP | 99.7 ± 0.6 | 104.2 ± 3.7 | 104.0 ± 4.8 | 110.0 ± 1.5 |
| Diethanolamine | 86.6 ± 0.9 | 98.8 ± 6.1 | 96.1 ± 1.0 | 99.5 ± 5.3 |
| Dihydrobiopterin | 96.4 ± 4.2 | 99.0 ± 4.6 | 99.6 ± 0.6 | 109.7 ± 4.1 |
| Dihydroneopterin triphosphate | 91.5 ± 1.4 | 106.6 ± 1.4 | 99.8 ± 6.1 | 102.8 ± 2.7 |
| Dihydroxyacetonephosphate | 104.5 ± 2.4 | 93.9 ± 2.9 | 109.3 ± 2.8 | 104.7 ± 0.2 |
| Dimethylglycine | 101.8 ± 1.2 | 99.4 ± 4.0 | 101.2 ± 1.8 | 94.3 ± 3.5 |
| Ectoine | 85.1 ± 5.1 | 95.4 ± 1.9 | 97.3 ± 3.8 | 91.4 ± 1.9 |
| Erythrose-4-phosphate | 100.6 ± 0.3 | 96.3 ± 3.4 | 102.2 ± 3.8 | 107.8 ± 5.9 |
| FAD | 91.1 ± 1.5 | 106.0 ± 1.3 | 107.0 ± 4.7 | 103.8 ± 3.5 |
| Folic acid | 101.1 ± 1.5 | 90.2 ± 3.8 | 90.5 ± 1.2 | 104.3 ± 2.9 |
| Fructose-6-phosphate | 86.8 ± 3.4 | 103.7 ± 6.2 | 109.3 ± 0.3 | 109.5 ± 3.2 |
| Fumaric Acid | 89.1 ± 3.7 | 102.5 ± 1.7 | 99.9 ± 4.9 | 91.1 ± 1.4 |
| Galactosamine | 103.9 ± 1.0 | 96.6 ± 3.8 | 94.4 ± 5.5 | 103.5 ± 5.7 |
| Galactose-1-phosphate | 89.4 ± 4.1 | 94.4 ± 4.5 | 93.9 ± 0.4 | 96.5 ± 5.5 |
| Galactose-6-Phosphate | 88.0 ± 2.7 | 96.5 ± 5.7 | 105.1 ± 5.2 | 98.1 ± 1.6 |
| gamma-Glutamylcysteine | 104.2 ± 2.3 | 90.2 ± 3.3 | 92.0 ± 1.2 | 98.3 ± 5.0 |
| Glucosamine | 85.5 ± 0.2 | 100.6 ± 3.0 | 95.1 ± 2.2 | 101.2 ± 4.0 |
| Glucosamine-6-Phosphate | 96.6 ± 2.9 | 108.9 ± 5.2 | 95.0 ± 1.8 | 90.9 ± 5.0 |
| Glucose | 87.2 ± 4.8 | 95.2 ± 2.5 | 102.7 ± 5.5 | 102.8 ± 2.1 |
| Glucose-1-phosphate | 99.7 ± 4.9 | 105.1 ± 0.2 | 104.2 ± 2.8 | 108.2 ± 4.7 |
| Glucose-6-Phosphate | 95.8 ± 5.0 | 102.4 ± 4.5 | 98.1 ± 2.7 | 92.9 ± 0.8 |
| Glucuronic acid | 102.4 ± 6.1 | 90.3 ± 4.5 | 101.8 ± 4.7 | 107.0 ± 3.9 |
| Glutamic acid | 88.5 ± 1.1 | 101.6 ± 5.5 | 92.0 ± 1.0 | 107.6 ± 2.9 |
| Glutamine | 87.4 ± 2.3 | 96.7 ± 1.4 | 109.7 ± 4.6 | 108.8 ± 0.7 |
| Glutathione | 99.0 ± 5.1 | 107.4 ± 2.6 | 97.1 ± 2.3 | 95.9 ± 4.6 |
| Glutathionylspermine | 98.0 ± 2.0 | 99.1 ± 2.0 | 90.8 ± 0.1 | 109.3 ± 2.5 |
| Glycerophosphocholine | 85.5 ± 4.1 | 106.6 ± 5.8 | 103.3 ± 5.8 | 99.9 ± 3.6 |
| Glycine | 85.9 ± 0.8 | 108.4 ± 4.7 | 100.9 ± 0.4 | 95.9 ± 3.0 |
| Glycolic acid | 91.9 ± 0.4 | 104.8 ± 4.8 | 106.2 ± 1.9 | 104.9 ± 0.9 |
| Guanidinoacetic acid | 89.3 ± 0.4 | 104.5 ± 4.3 | 109.8 ± 5.2 | 92.4 ± 4.9 |
| Guanidinobutanoic acid | 90.1 ± 5.9 | 90.8 ± 3.1 | 90.5 ± 3.5 | 94.0 ± 5.2 |
| Guanine | 99.5 ± 4.8 | 106.5 ± 4.5 | 93.7 ± 3.0 | 101.8 ± 1.2 |
| Guanosine | 98.0 ± 1.2 | 109.9 ± 5.6 | 95.5 ± 6.2 | 104.9 ± 4.3 |
| Guanosine 5'-diphosphate | 91.1 ± 1.9 | 101.6 ± 1.4 | 94.9 ± 2.6 | 104.6 ± 2.8 |
| Guanosine-5'-monophosphate | 88.6 ± 3.1 | 91.0 ± 5.0 | 104.5 ± 1.2 | 109.2 ± 1.3 |
| Hercynine | 101.4 ± 2.9 | 109.8 ± 1.3 | 91.1 ± 5.3 | 92.6 ± 4.3 |
| Hippuric acid | 95.0 ± 4.2 | 95.5 ± 3.2 | 103.3 ± 1.0 | 107.8 ± 5.3 |
| Histamine | 100.7 ± 3.4 | 104.3 ± 4.4 | 92.9 ± 5.9 | 104.0 ± 5.2 |
| Histidine | 91.7 ± 4.3 | 97.4 ± 5.3 | 91.0 ± 0.1 | 95.5 ± 0.5 |
| Homoalanine | 91.1 ± 1.7 | 92.5 ± 2.6 | 100.0 ± 4.9 | 95.7 ± 5.6 |
| Homoarginine | 104.8 ± 1.2 | 97.9 ± 1.5 | 101.3 ± 1.9 | 102.1 ± 0.6 |
| Homocysteine | 93.9 ± 4.3 | 98.6 ± 4.1 | 106.1 ± 4.3 | 98.2 ± 3.3 |
| Homogentisic acid | 99.5 ± 2.6 | 101.6 ± 4.6 | 107.8 ± 0.6 | 93.3 ± 2.1 |
| Homoleucine | 99.2 ± 1.5 | 100.7 ± 1.8 | 108.6 ± 2.9 | 95.1 ± 2.3 |
| Homovanillic acid | 95.4 ± 0.8 | 109.4 ± 5.3 | 92.0 ± 3.5 | 104.7 ± 1.1 |
| Hydantoin-5-propionate | 99.5 ± 0.7 | 98.7 ± 3.4 | 94.1 ± 4.3 | 100.2 ± 1.5 |
| Hypotaurine | 96.2 ± 5.3 | 108.5 ± 4.8 | 106.3 ± 0.3 | 97.3 ± 3.8 |
| Hypoxanthine | 85.8 ± 5.8 | 91.6 ± 4.1 | 108.1 ± 4.1 | 106.6 ± 1.6 |
| Indole-3-acetamide | 104.7 ± 4.8 | 95.4 ± 4.7 | 98.0 ± 2.1 | 93.2 ± 6.1 |
| Inosine | 86.4 ± 3.3 | 90.8 ± 1.4 | 97.6 ± 4.3 | 110.0 ± 4.4 |
| Inosine 5'-monophosphate | 103.1 ± 2.5 | 107.4 ± 3.5 | 101.6 ± 4.8 | 104.8 ± 6.1 |
| Isocitric Acid | 96.1 ± 1.7 | 91.7 ± 0.2 | 106.4 ± 3.1 | 93.3 ± 1.6 |
| Isoleucine | 86.1 ± 1.1 | 103.7 ± 4.0 | 98.4 ± 5.3 | 90.8 ± 4.9 |
| Lactic Acid | 88.8 ± 6.1 | 109.7 ± 1.7 | 94.2 ± 0.9 | 106.3 ± 4.9 |
| Leucine | 101.2 ± 5.3 | 104.8 ± 1.1 | 96.9 ± 3.0 | 103.0 ± 5.5 |
| Levulinic acid | 103.1 ± 5.0 | 94.1 ± 0.1 | 90.7 ± 1.5 | 100.6 ± 4.5 |
| Lysine | 87.7 ± 4.5 | 103.2 ± 0.2 | 90.7 ± 5.3 | 92.4 ± 2.5 |
| Malic Acid | 97.7 ± 4.9 | 90.0 ± 3.8 | 103.8 ± 5.5 | 96.0 ± 4.9 |
| Mannose-1 with Mannose-6-Phosphate | 85.6 ± 5.2 | 91.9 ± 2.4 | 94.2 ± 5.0 | 98.9 ± 3.8 |
| Mesaconic acid | 93.3 ± 1.8 | 107.7 ± 3.4 | 94.3 ± 2.5 | 92.8 ± 0.5 |
| Methionine | 103.5 ± 3.2 | 107.8 ± 6.1 | 99.1 ± 4.4 | 97.4 ± 3.1 |
| Methionine sulfoxide | 88.3 ± 4.9 | 107.0 ± 1.3 | 103.2 ± 1.6 | 91.5 ± 2.4 |
| Methylmalonic Acid | 94.6 ± 1.4 | 92.1 ± 3.9 | 107.7 ± 0.7 | 92.2 ± 4.0 |
| N1-Acetylspermidine | 85.8 ± 3.1 | 96.4 ± 0.9 | 91.9 ± 0.3 | 93.2 ± 2.3 |
| N1-Acetylspermine | 89.4 ± 6.2 | 103.1 ± 4.1 | 96.4 ± 6.1 | 100.2 ± 4.7 |
| N8-Acetylspermidine | 103.8 ± 5.6 | 102.2 ± 4.5 | 90.8 ± 1.8 | 104.9 ± 4.8 |
| N-Acetyl-Aspartyl-Glutamate | 97.8 ± 6.1 | 102.2 ± 2.0 | 108.7 ± 3.1 | 99.3 ± 3.1 |
| N-Acetyl-Aspartyl-Glutamyl-Glutamate | 104.6 ± 1.8 | 105.9 ± 4.1 | 98.5 ± 2.9 | 94.0 ± 1.3 |
| N-Acetyl-Galactosamine | 96.5 ± 3.1 | 91.2 ± 3.8 | 92.0 ± 1.7 | 104.7 ± 3.9 |
| N-Acetyl-Leucine | 93.7 ± 5.6 | 94.4 ± 2.1 | 106.2 ± 4.3 | 97.3 ± 1.2 |
| N-Acetyl-Methionine | 98.8 ± 0.8 | 95.1 ± 3.7 | 105.7 ± 2.1 | 107.4 ± 2.3 |
| N-Acetylneuraminic acid | 103.2 ± 1.7 | 108.7 ± 3.0 | 93.8 ± 0.7 | 91.3 ± 4.7 |
| N-Acetyl-Proline | 100.4 ± 4.5 | 107.8 ± 5.1 | 99.0 ± 5.7 | 107.0 ± 2.0 |
| N-Acetyl-Putrescine | 97.2 ± 4.8 | 92.6 ± 2.8 | 102.4 ± 5.6 | 104.4 ± 4.4 |
| N-Acetyl-Serine | 94.3 ± 3.1 | 90.1 ± 0.3 | 104.8 ± 1.2 | 104.1 ± 0.8 |
| N-Acetyl-Serotonin | 91.9 ± 1.8 | 106.5 ± 1.2 | 99.8 ± 5.7 | 99.1 ± 5.8 |
| N-Acyl-Homoserine-lactone | 99.7 ± 5.6 | 107.5 ± 3.0 | 103.4 ± 6.2 | 93.8 ± 0.4 |
| NAD | 91.9 ± 3.5 | 91.4 ± 5.8 | 106.8 ± 3.5 | 103.0 ± 4.5 |
| NADH | 90.6 ± 6.1 | 106.8 ± 5.2 | 109.8 ± 2.5 | 100.8 ± 4.3 |
| NADP | 86.2 ± 4.6 | 107.4 ± 3.5 | 93.5 ± 1.2 | 104.0 ± 3.5 |
| NADPH | 104.0 ± 0.6 | 106.1 ± 4.8 | 109.3 ± 4.1 | 97.8 ± 5.0 |
| N-alpha-Acetyl-lysine | 85.6 ± 4.3 | 108.6 ± 3.6 | 108.7 ± 5.5 | 104.5 ± 6.3 |
| N-alpha-Acetyl-Ornithine | 92.0 ± 1.7 | 92.4 ± 3.5 | 93.8 ± 1.9 | 110.0 ± 3.9 |
| N-Carbamoyl-Proline | 101.5 ± 4.9 | 98.1 ± 3.3 | 108.5 ± 5.5 | 93.3 ± 5.7 |
| N-Carbamoyl-Sarcosine | 91.6 ± 4.7 | 94.3 ± 2.7 | 95.6 ± 5.9 | 102.4 ± 0.7 |
| N-Epsilon-Acetyl-lysine | 86.5 ± 4.8 | 107.7 ± 6.3 | 104.9 ± 1.3 | 105.1 ± 1.7 |
| Nicotinamide | 100.4 ± 6.3 | 100.1 ± 3.6 | 105.0 ± 2.9 | 99.3 ± 5.6 |
| N-Methyl-Aspartic Acid | 98.4 ± 0.9 | 96.1 ± 0.9 | 101.3 ± 1.3 | 106.9 ± 4.8 |
| O-Phosphoethanolamine | 104.5 ± 1.6 | 109.1 ± 5.9 | 105.9 ± 6.0 | 101.0 ± 6.0 |
| Ophthalmic acid | 85.4 ± 0.6 | 101.5 ± 3.9 | 93.1 ± 1.3 | 93.3 ± 3.9 |
| Ornithine | 95.2 ± 2.2 | 110.0 ± 0.8 | 109.5 ± 0.7 | 99.5 ± 1.1 |
| Orotidine 5'-monophosphate | 113.2 ± 2.0 | 87.8 ± 0.5 | 105.9 ± 1.9 | 92.7 ± 3.7 |
| Oxaloacetate | 92.3 ± 0.5 | 106.7 ± 0.6 | 106.9 ± 0.4 | 99.1 ± 0.9 |
| Pantothenic acid | 113.9 ± 2.9 | 89.2 ± 2.2 | 87.7 ± 1.1 | 94.8 ± 0.3 |
| Phenylalanine | 115.0 ± 2.7 | 112.1 ± 3.5 | 108.0 ± 1.9 | 94.6 ± 3.5 |
| Phenylpyruvate | 98.6 ± 2.5 | 101.2 ± 2.6 | 105.0 ± 1.1 | 97.8 ± 2.9 |
| Phosphodimethylethanolamine | 107.3 ± 1.1 | 91.6 ± 3.0 | 93.5 ± 3.9 | 113.4 ± 1.0 |
| Phosphoenolpyruvate | 113.6 ± 2.9 | 91.7 ± 1.9 | 112.4 ± 2.1 | 94.7 ± 3.8 |
| Phosphorylcholine | 91.4 ± 2.7 | 88.2 ± 2.3 | 101.2 ± 1.3 | 111.9 ± 3.1 |
| Pipecolic acid | 86.2 ± 0.4 | 111.5 ± 0.9 | 101.4 ± 1.1 | 112.2 ± 1.4 |
| Proline | 102.8 ± 2.9 | 106.1 ± 1.2 | 109.0 ± 2.2 | 102.9 ± 2.6 |
| Putrescine | 104.6 ± 0.5 | 90.0 ± 1.6 | 105.3 ± 0.4 | 88.3 ± 1.1 |
| Pyridoxal | 108.4 ± 0.9 | 89.6 ± 3.1 | 87.8 ± 1.3 | 105.7 ± 3.7 |
| Pyridoxamine | 92.9 ± 1.2 | 91.6 ± 1.5 | 108.1 ± 1.0 | 112.9 ± 0.6 |
| Pyridoxine | 96.9 ± 2.9 | 112.4 ± 3.5 | 107.5 ± 3.4 | 106.0 ± 2.0 |
| Pyroglutamic acid | 97.4 ± 0.4 | 108.2 ± 2.7 | 112.9 ± 1.3 | 104.3 ± 2.6 |
| Pyrrolysine | 99.2 ± 0.5 | 107.0 ± 1.9 | 86.2 ± 1.2 | 101.1 ± 3.5 |
| Pyruvic Acid | 107.8 ± 2.7 | 114.8 ± 2.5 | 111.3 ± 3.8 | 86.9 ± 1.2 |
| Riboflavin | 101.8 ± 2.6 | 94.5 ± 3.7 | 104.8 ± 1.6 | 109.7 ± 2.4 |
| Ribose-5-Phosphate | 102.8 ± 1.9 | 108.4 ± 2.1 | 109.3 ± 1.5 | 102.5 ± 3.8 |
| Ribulose-1-Phosphate | 108.2 ± 1.6 | 96.6 ± 0.7 | 106.2 ± 2.6 | 112.0 ± 2.8 |
| Ribulose-5-Phosphate | 98.1 ± 1.9 | 113.6 ± 2.9 | 85.3 ± 3.7 | 94.3 ± 0.4 |
| Saccharopine | 103.2 ± 2.4 | 108.1 ± 0.4 | 106.8 ± 0.7 | 93.5 ± 1.1 |
| S-Adenosylhomocysteine | 105.9 ± 2.4 | 107.0 ± 2.8 | 110.2 ± 3.2 | 95.8 ± 1.6 |
| S-Adenosylmethioninamine | 111.3 ± 2.5 | 86.7 ± 2.2 | 113.5 ± 2.5 | 88.2 ± 2.1 |
| S-Adenosylmethionine | 97.1 ± 1.5 | 113.8 ± 2.9 | 104.0 ± 2.0 | 94.8 ± 1.9 |
| Sarcosine | 96.6 ± 1.0 | 91.1 ± 3.9 | 92.7 ± 0.7 | 104.8 ± 1.7 |
| Sedoheptulose-7-phosphate | 99.5 ± 0.6 | 93.3 ± 2.0 | 90.9 ± 1.7 | 88.0 ± 3.9 |
| Selenohomocysteine | 108.9 ± 2.3 | 110.6 ± 3.8 | 105.5 ± 2.0 | 94.5 ± 2.5 |
| Selenomethionine | 113.8 ± 2.9 | 103.1 ± 3.7 | 103.5 ± 0.6 | 110.3 ± 2.5 |
| Serine | 104.4 ± 2.8 | 90.2 ± 0.9 | 89.6 ± 0.4 | 86.2 ± 2.4 |
| S-Methyl-Thioinosine | 102.0 ± 1.5 | 92.6 ± 0.3 | 95.1 ± 3.9 | 95.5 ± 2.0 |
| Spermidine | 90.1 ± 2.4 | 107.1 ± 0.9 | 98.7 ± 2.4 | 95.7 ± 2.0 |
| Spermine | 87.6 ± 2.2 | 102.8 ± 0.6 | 112.9 ± 3.5 | 100.5 ± 3.9 |
| Succinate | 108.4 ± 0.6 | 91.1 ± 3.1 | 86.2 ± 2.8 | 89.0 ± 1.1 |
| Sucrose | 106.0 ± 3.7 | 92.2 ± 2.7 | 107.4 ± 3.2 | 92.3 ± 2.8 |
| Symmetric dimethylarginine | 102.5 ± 1.6 | 112.0 ± 2.3 | 102.6 ± 2.6 | 87.9 ± 2.9 |
| Taurine | 95.9 ± 0.5 | 85.1 ± 2.3 | 96.4 ± 3.6 | 106.9 ± 3.2 |
| Tetrahydroneopterin | 102.1 ± 0.7 | 110.9 ± 1.4 | 90.2 ± 0.5 | 107.8 ± 2.0 |
| Thiamine | 97.2 ± 3.3 | 109.7 ± 3.5 | 101.5 ± 2.6 | 106.2 ± 3.4 |
| Thiamine monophosphate | 104.9 ± 2.3 | 85.1 ± 2.3 | 113.8 ± 0.9 | 93.7 ± 2.7 |
| Threonic acid | 107.2 ± 1.6 | 105.4 ± 2.2 | 101.5 ± 1.7 | 90.0 ± 1.7 |
| Threonine | 97.5 ± 2.5 | 98.0 ± 1.3 | 106.7 ± 0.7 | 98.9 ± 3.5 |
| Thymidine | 90.5 ± 1.6 | 107.4 ± 0.7 | 96.8 ± 0.8 | 111.6 ± 0.4 |
| Thymidine Monophosphate | 99.4 ± 3.3 | 98.9 ± 2.5 | 86.6 ± 3.4 | 102.3 ± 3.6 |
| Thymine | 93.1 ± 2.2 | 91.9 ± 2.8 | 90.6 ± 3.5 | 108.8 ± 2.3 |
| Trigonelleine | 94.4 ± 3.8 | 103.1 ± 3.7 | 105.2 ± 1.3 | 93.8 ± 2.6 |
| Trimethylamine N-oxide | 111.0 ± 3.2 | 88.3 ± 1.0 | 112.5 ± 0.4 | 101.0 ± 2.5 |
| Trimethyllysine | 89.7 ± 2.5 | 90.0 ± 2.9 | 85.1 ± 0.5 | 109.3 ± 1.2 |
| Tryptophan | 98.7 ± 1.7 | 113.4 ± 0.5 | 89.0 ± 2.4 | 108.9 ± 3.0 |
| Tyramine | 90.3 ± 0.3 | 94.9 ± 1.2 | 97.2 ± 1.2 | 88.2 ± 1.3 |
| Tyrosine | 107.9 ± 1.5 | 102.4 ± 2.0 | 97.1 ± 3.0 | 106.8 ± 1.6 |
| Uracil | 97.5 ± 2.9 | 108.2 ± 2.1 | 86.4 ± 0.4 | 88.4 ± 3.0 |
| Uridine 5'-diphosphate | 95.5 ± 3.3 | 96.6 ± 2.3 | 114.0 ± 1.9 | 96.7 ± 2.0 |
| Uridine Monophosphate | 98.7 ± 1.6 | 102.4 ± 1.7 | 112.7 ± 1.5 | 100.3 ± 3.0 |
| Uridine-5-Diphosphogalactose | 108.4 ± 1.4 | 113.1 ± 1.6 | 108.0 ± 2.9 | 90.5 ± 2.2 |
| Urocanic acid | 92.7 ± 0.8 | 89.2 ± 3.0 | 105.0 ± 1.7 | 89.9 ± 3.4 |
| UTP | 109.9 ± 1.6 | 98.3 ± 2.9 | 88.9 ± 1.6 | 85.3 ± 3.2 |
| Valine | 88.9 ± 2.0 | 97.4 ± 2.5 | 102.7 ± 3.9 | 89.8 ± 1.6 |
| XMP | 101.1 ± 1.8 | 91.2 ± 1.7 | 95.5 ± 3.8 | 91.1 ± 2.7 |
| Xylulose-5-Phosphate | 99.4 ± 2.2 | 98.7 ± 3.3 | 106.6 ± 2.1 | 110.5 ± 3.0 |

### **Table S29** | Inter – day accuracy (as determined by % recovery) (Plasma extract as matrix, n = 5). The concentrations of QC_LLOQ_, QC_Low_, QC_Mid_ and QC_High_ were 10 ng/mL, 25 ng/mL, 100 ng/mL and 500 ng/mL respectively.

| Accuracy, % | QC_LLOQ_ | QC_Low_ | QC_Mid_ | QC_High_ |
| --- | --- | --- | --- | --- |
| 1-Methylguanine | 95.7 ± 5.7 | 113.0 ± 2.5 | 92.0 ± 0.2 | 99.8 ± 2.8 |
| 1-Methylhistidine | 100.9 ± 4.3 | 109.4 ± 0.5 | 96.3 ± 2.4 | 102.9 ± 1.8 |
| 1-Methylnicotinamide | 110.8 ± 5.9 | 100.5 ± 5.1 | 103.2 ± 3.2 | 101.2 ± 4.2 |
| 2-Aminoadipic Acid | 114.0 ± 6.0 | 101.5 ± 0.9 | 93.3 ± 3.7 | 99.4 ± 6.3 |
| 2-Deoxyadenosine | 110.8 ± 3.1 | 93.9 ± 3.1 | 101.4 ± 1.0 | 106.2 ± 3.8 |
| 2'-Deoxyadenosine 5'-monophosphate | 101.9 ± 2.0 | 113.6 ± 4.8 | 95.3 ± 3.5 | 105.0 ± 5.3 |
| 2'-deoxyadenosine 5'-triphosphate | 96.3 ± 2.7 | 88.2 ± 4.9 | 87.4 ± 2.8 | 86.2 ± 0.8 |
| 2'-Deoxycytidine-5'-monophosphate | 97.3 ± 4.8 | 95.1 ± 6.1 | 98.9 ± 3.3 | 85.7 ± 0.6 |
| 2-Deoxyguanosine | 104.6 ± 0.2 | 92.6 ± 4.7 | 114.5 ± 3.0 | 102.2 ± 0.3 |
| 2-hydroxybutyric acid | 103.0 ± 0.4 | 88.3 ± 0.2 | 105.4 ± 1.6 | 92.6 ± 3.1 |
| 2-hydroxyglutaric acid | 87.5 ± 4.0 | 87.3 ± 1.0 | 93.8 ± 2.8 | 88.6 ± 0.6 |
| 2-Hydroxyisocaproic acid | 88.1 ± 1.1 | 101.4 ± 3.5 | 111.2 ± 2.5 | 114.7 ± 2.2 |
| 2-ketobutyric acid | 105.4 ± 0.4 | 95.3 ± 0.6 | 114.9 ± 6.2 | 101.1 ± 3.2 |
| 2-oxoadipic acid | 98.0 ± 1.5 | 108.2 ± 4.6 | 111.4 ± 3.1 | 101.6 ± 2.1 |
| 2-Phenylglycine | 110.5 ± 6.2 | 95.9 ± 1.0 | 104.5 ± 0.6 | 112.6 ± 6.2 |
| 2-phosphoglycerate | 93.2 ± 2.5 | 108.5 ± 2.9 | 114.6 ± 1.7 | 99.6 ± 2.7 |
| 3-hydroxy-3-methyl-2-oxobutanoate | 99.7 ± 0.7 | 105.6 ± 1.0 | 90.2 ± 2.1 | 99.5 ± 0.6 |
| 3-hydroxybutyric acid | 91.5 ± 3.7 | 113.1 ± 3.4 | 96.0 ± 5.1 | 113.5 ± 5.6 |
| 3-Methylhistidine | 94.3 ± 1.5 | 100.5 ± 3.9 | 104.7 ± 0.6 | 112.7 ± 5.0 |
| 3-Phosphoglyceraldehyde | 88.8 ± 1.7 | 91.2 ± 2.9 | 111.9 ± 2.9 | 104.6 ± 1.4 |
| 3-phosphoglycerate | 95.1 ± 0.2 | 102.9 ± 0.8 | 108.1 ± 4.5 | 93.9 ± 3.2 |
| 4-amino-butyric acid | 105.3 ± 2.0 | 110.7 ± 5.5 | 112.9 ± 5.2 | 102.2 ± 1.0 |
| 4-aspartyl-phosphate | 112.1 ± 2.7 | 95.1 ± 1.8 | 100.3 ± 3.4 | 111.6 ± 5.4 |
| 4-Hydroxybenzoic acid | 93.9 ± 1.9 | 112.3 ± 1.9 | 89.8 ± 5.8 | 86.7 ± 0.6 |
| 4-Hydroxyphenylacetylglycine | 109.8 ± 2.8 | 93.9 ± 3.8 | 107.9 ± 4.7 | 98.6 ± 4.0 |
| 4-methyl-2-oxopentanoate | 101.2 ± 6.1 | 109.5 ± 1.0 | 108.2 ± 1.1 | 85.4 ± 2.2 |
| 4-Oxoglutaramate | 112.0 ± 0.6 | 108.2 ± 1.6 | 91.1 ± 3.5 | 101.7 ± 3.2 |
| 5-Hydroxyindolepyruvate | 111.1 ± 0.6 | 92.9 ± 5.3 | 98.0 ± 1.4 | 106.6 ± 0.2 |
| 5-Hydroxylysine | 104.0 ± 2.6 | 104.4 ± 1.6 | 109.4 ± 4.8 | 91.8 ± 3.6 |
| 5-Hydroxy-Tryptophan | 111.9 ± 0.8 | 94.6 ± 4.9 | 86.0 ± 1.2 | 86.0 ± 3.6 |
| 5-Phosphoribosylamine | 111.0 ± 5.4 | 104.0 ± 6.1 | 114.6 ± 0.7 | 89.8 ± 4.4 |
| 6-phosphogluconate | 90.8 ± 5.6 | 102.6 ± 2.1 | 94.0 ± 5.6 | 110.7 ± 2.6 |
| Acetylcarnitine | 100.7 ± 2.0 | 97.1 ± 2.0 | 87.8 ± 2.0 | 95.7 ± 1.0 |
| Aconitic Acid | 101.7 ± 3.9 | 107.6 ± 5.4 | 113.4 ± 3.3 | 99.2 ± 3.5 |
| Adenine | 93.6 ± 3.5 | 101.4 ± 3.2 | 97.1 ± 2.2 | 100.1 ± 4.4 |
| Adenosine | 104.7 ± 0.5 | 85.3 ± 4.8 | 100.1 ± 4.4 | 106.5 ± 0.6 |
| Adenosine 3',5'-cyclic monophosphate | 91.9 ± 2.0 | 103.3 ± 5.1 | 109.0 ± 0.7 | 112.4 ± 5.8 |
| Adenosine Monophosphate | 92.4 ± 2.4 | 108.6 ± 3.3 | 97.0 ± 4.8 | 87.8 ± 1.3 |
| Adenosine-5'-diphosphate | 95.0 ± 3.0 | 113.8 ± 2.0 | 112.9 ± 3.2 | 105.1 ± 1.6 |
| Adenosine-5'-triphosphate | 105.8 ± 6.1 | 94.0 ± 3.6 | 94.4 ± 1.9 | 100.4 ± 6.0 |
| Adenylosuccinate | 94.1 ± 4.2 | 85.3 ± 0.8 | 85.7 ± 1.9 | 108.6 ± 5.9 |
| Alanine | 113.8 ± 1.8 | 104.2 ± 3.7 | 99.5 ± 5.1 | 85.5 ± 5.6 |
| alpha-ketoglutaric acid | 86.9 ± 4.5 | 113.4 ± 1.4 | 111.7 ± 0.8 | 88.3 ± 3.9 |
| Anserine | 95.7 ± 4.5 | 100.2 ± 2.0 | 96.6 ± 6.2 | 104.6 ± 1.9 |
| Arginine | 108.9 ± 4.2 | 98.6 ± 6.0 | 114.1 ± 6.3 | 111.3 ± 0.3 |
| Argininosuccinic Acid | 102.3 ± 4.3 | 90.8 ± 1.9 | 91.6 ± 1.7 | 95.5 ± 3.7 |
| Asparagine | 87.8 ± 4.2 | 110.2 ± 1.9 | 91.6 ± 0.7 | 101.2 ± 3.7 |
| Aspartic acid | 94.7 ± 2.2 | 110.1 ± 0.7 | 104.9 ± 1.9 | 97.9 ± 1.9 |
| Asymmetric dimethylarginine | 111.6 ± 4.8 | 101.4 ± 5.4 | 104.7 ± 2.2 | 101.0 ± 2.4 |
| Azelaic acid | 94.2 ± 3.3 | 110.5 ± 1.7 | 99.9 ± 6.0 | 108.9 ± 3.8 |
| beta-Alanine | 98.9 ± 5.6 | 85.5 ± 6.1 | 97.9 ± 2.4 | 90.7 ± 0.2 |
| Betaine | 112.6 ± 4.5 | 109.0 ± 4.2 | 106.0 ± 3.0 | 106.2 ± 0.6 |
| Biopterin | 89.8 ± 4.9 | 101.0 ± 3.4 | 110.6 ± 5.4 | 91.7 ± 3.1 |
| Biotin | 93.2 ± 0.9 | 94.3 ± 0.8 | 111.7 ± 4.1 | 108.6 ± 2.6 |
| Carnitine | 107.4 ± 5.8 | 111.4 ± 3.8 | 113.6 ± 4.9 | 101.3 ± 6.0 |
| Carnosine | 95.6 ± 4.2 | 104.3 ± 4.4 | 88.1 ± 0.5 | 89.4 ± 6.1 |
| CDP-choline | 114.3 ± 5.0 | 89.9 ± 5.5 | 93.8 ± 4.4 | 108.5 ± 5.0 |
| cGMP | 100.8 ± 5.8 | 102.2 ± 6.1 | 86.5 ± 4.1 | 86.6 ± 2.5 |
| Choline | 91.1 ± 4.6 | 111.0 ± 2.6 | 94.7 ± 5.2 | 86.8 ± 5.9 |
| Cinnamic acid | 109.2 ± 4.8 | 113.4 ± 5.2 | 101.1 ± 6.2 | 97.8 ± 6.1 |
| Citraconic acid | 108.3 ± 0.8 | 101.1 ± 4.6 | 87.0 ± 4.6 | 96.4 ± 3.5 |
| Citramalic acid | 100.5 ± 2.3 | 109.4 ± 2.8 | 89.7 ± 3.9 | 100.2 ± 1.3 |
| Citric Acid | 103.2 ± 1.4 | 106.7 ± 2.2 | 96.8 ± 1.3 | 87.2 ± 4.3 |
| Citrulline | 101.1 ± 1.5 | 100.1 ± 2.8 | 101.9 ± 2.2 | 108.6 ± 2.1 |
| Creatine | 94.7 ± 0.2 | 94.1 ± 4.0 | 98.3 ± 0.7 | 105.4 ± 4.9 |
| Creatinine | 109.5 ± 5.4 | 106.3 ± 4.1 | 106.8 ± 3.5 | 85.9 ± 1.2 |
| Cystathionine | 95.0 ± 4.6 | 100.5 ± 5.0 | 86.2 ± 5.2 | 106.2 ± 0.9 |
| Cysteamine | 99.0 ± 2.3 | 89.9 ± 2.8 | 114.3 ± 0.8 | 95.3 ± 3.2 |
| Cysteine | 85.3 ± 4.0 | 110.7 ± 2.9 | 103.3 ± 1.2 | 85.6 ± 2.7 |
| Cysteinylglycine | 91.4 ± 4.2 | 90.3 ± 0.5 | 108.8 ± 4.3 | 96.1 ± 1.9 |
| Cystine | 109.6 ± 0.8 | 91.7 ± 0.4 | 96.6 ± 1.9 | 92.9 ± 1.4 |
| Cytidine | 114.3 ± 1.5 | 103.9 ± 1.8 | 92.3 ± 0.5 | 95.0 ± 1.8 |
| Cytidine 3',5'-cyclic monophosphate | 107.9 ± 0.9 | 112.4 ± 0.3 | 89.8 ± 4.3 | 107.8 ± 0.1 |
| Cytidine 5'-diphosphate | 99.8 ± 4.6 | 114.2 ± 1.7 | 90.3 ± 3.0 | 109.6 ± 2.6 |
| Cytidine 5'-triphosphate | 108.7 ± 4.2 | 97.0 ± 4.2 | 92.5 ± 2.6 | 98.7 ± 1.8 |
| Cytidine monophosphate | 103.2 ± 4.6 | 91.7 ± 5.1 | 106.7 ± 2.2 | 92.0 ± 1.6 |
| Cytosine | 112.9 ± 5.4 | 96.7 ± 2.4 | 101.8 ± 5.4 | 102.1 ± 5.1 |
| dADP | 108.9 ± 0.7 | 96.5 ± 3.1 | 99.9 ± 4.0 | 107.3 ± 0.9 |
| Diethanolamine | 93.7 ± 3.3 | 107.7 ± 4.4 | 98.6 ± 1.2 | 108.4 ± 4.3 |
| Dihydrobiopterin | 105.6 ± 1.7 | 106.7 ± 6.2 | 112.0 ± 2.7 | 102.7 ± 3.2 |
| Dihydroneopterin triphosphate | 108.6 ± 5.0 | 105.5 ± 3.9 | 105.0 ± 0.5 | 103.6 ± 3.1 |
| Dihydroxyacetonephosphate | 94.9 ± 5.2 | 86.6 ± 1.9 | 89.1 ± 5.1 | 101.9 ± 6.3 |
| Dimethylglycine | 96.0 ± 1.4 | 91.0 ± 4.7 | 108.2 ± 1.3 | 104.4 ± 6.0 |
| Ectoine | 104.9 ± 1.3 | 108.3 ± 0.2 | 92.1 ± 1.9 | 90.5 ± 3.5 |
| Erythrose-4-phosphate | 110.5 ± 1.6 | 85.7 ± 2.5 | 97.3 ± 6.0 | 102.6 ± 0.7 |
| FAD | 109.1 ± 2.5 | 101.1 ± 4.4 | 114.8 ± 0.2 | 112.7 ± 2.6 |
| Folic acid | 85.6 ± 3.9 | 112.1 ± 2.4 | 98.6 ± 4.2 | 86.2 ± 0.4 |
| Fructose-6-phosphate | 91.3 ± 4.5 | 93.0 ± 1.4 | 102.8 ± 4.6 | 87.9 ± 4.4 |
| Fumaric Acid | 96.9 ± 5.5 | 95.3 ± 0.2 | 96.9 ± 2.6 | 95.4 ± 6.1 |
| Galactosamine | 86.8 ± 2.7 | 89.1 ± 5.8 | 107.3 ± 1.7 | 90.4 ± 5.4 |
| Galactose-1-phosphate | 109.0 ± 0.8 | 111.7 ± 5.5 | 97.0 ± 2.0 | 97.6 ± 4.2 |
| Galactose-6-Phosphate | 98.4 ± 2.5 | 100.5 ± 2.9 | 88.1 ± 6.1 | 96.5 ± 5.4 |
| gamma-Glutamylcysteine | 108.3 ± 1.0 | 97.4 ± 2.2 | 111.0 ± 0.7 | 98.9 ± 0.4 |
| Glucosamine | 93.0 ± 6.1 | 91.1 ± 1.3 | 90.5 ± 1.9 | 95.3 ± 2.8 |
| Glucosamine-6-Phosphate | 91.2 ± 6.1 | 99.7 ± 3.0 | 110.9 ± 1.2 | 109.3 ± 1.3 |
| Glucose | 87.7 ± 2.3 | 86.5 ± 5.5 | 103.7 ± 3.4 | 90.8 ± 1.9 |
| Glucose-1-phosphate | 111.5 ± 1.2 | 109.5 ± 3.0 | 105.9 ± 1.7 | 110.7 ± 3.0 |
| Glucose-6-Phosphate | 88.0 ± 4.1 | 111.6 ± 1.1 | 87.9 ± 0.4 | 93.0 ± 2.2 |
| Glucuronic acid | 99.7 ± 4.4 | 109.4 ± 3.7 | 101.5 ± 0.2 | 87.5 ± 3.9 |
| Glutamic acid | 109.5 ± 1.6 | 89.7 ± 3.5 | 86.8 ± 2.6 | 86.4 ± 3.3 |
| Glutamine | 95.2 ± 1.5 | 111.3 ± 2.9 | 90.5 ± 3.3 | 104.0 ± 6.1 |
| Glutathione | 98.5 ± 0.5 | 107.1 ± 1.7 | 113.9 ± 0.3 | 91.6 ± 5.7 |
| Glutathionylspermine | 108.2 ± 3.1 | 111.6 ± 5.1 | 85.1 ± 4.4 | 91.1 ± 3.6 |
| Glycerophosphocholine | 113.4 ± 0.9 | 106.7 ± 2.3 | 90.8 ± 1.3 | 88.9 ± 5.1 |
| Glycine | 87.8 ± 6.2 | 109.7 ± 2.4 | 102.4 ± 1.3 | 96.5 ± 5.4 |
| Glycolic acid | 100.2 ± 1.9 | 96.4 ± 4.4 | 111.1 ± 0.1 | 90.3 ± 3.3 |
| Guanidinoacetic acid | 94.4 ± 4.7 | 105.3 ± 4.8 | 104.5 ± 5.0 | 95.2 ± 4.1 |
| Guanidinobutanoic acid | 103.5 ± 0.2 | 99.0 ± 0.3 | 99.0 ± 4.7 | 85.2 ± 6.1 |
| Guanine | 87.0 ± 2.2 | 87.3 ± 5.5 | 86.4 ± 2.9 | 109.0 ± 5.0 |
| Guanosine | 89.4 ± 4.4 | 95.1 ± 6.0 | 98.7 ± 4.6 | 112.4 ± 0.5 |
| Guanosine 5'-diphosphate | 102.3 ± 2.7 | 98.7 ± 1.9 | 101.4 ± 2.3 | 88.3 ± 4.1 |
| Guanosine-5'-monophosphate | 101.8 ± 2.7 | 86.8 ± 3.1 | 107.3 ± 1.9 | 94.4 ± 5.7 |
| Hercynine | 114.8 ± 3.7 | 91.7 ± 4.0 | 111.7 ± 3.7 | 103.3 ± 1.2 |
| Hippuric acid | 112.0 ± 5.0 | 91.0 ± 1.1 | 100.4 ± 3.3 | 101.3 ± 6.0 |
| Histamine | 85.5 ± 3.1 | 88.0 ± 3.6 | 114.4 ± 1.7 | 106.2 ± 5.6 |
| Histidine | 93.5 ± 6.1 | 111.6 ± 6.1 | 103.9 ± 0.6 | 108.0 ± 5.2 |
| Homoalanine | 105.8 ± 2.4 | 86.6 ± 4.7 | 98.2 ± 2.7 | 114.0 ± 0.7 |
| Homoarginine | 110.5 ± 4.6 | 97.8 ± 1.7 | 87.1 ± 5.8 | 98.1 ± 2.5 |
| Homocysteine | 113.6 ± 1.4 | 99.4 ± 1.5 | 107.0 ± 5.4 | 93.9 ± 4.6 |
| Homogentisic acid | 96.0 ± 3.1 | 89.9 ± 5.1 | 112.4 ± 4.8 | 99.2 ± 6.2 |
| Homoleucine | 88.5 ± 2.5 | 95.7 ± 4.5 | 95.9 ± 2.2 | 98.7 ± 5.0 |
| Homovanillic acid | 90.8 ± 4.3 | 86.5 ± 3.3 | 94.6 ± 6.1 | 100.7 ± 1.5 |
| Hydantoin-5-propionate | 112.1 ± 6.0 | 101.2 ± 2.0 | 95.7 ± 2.0 | 113.6 ± 0.7 |
| Hypotaurine | 100.4 ± 5.7 | 87.1 ± 0.1 | 105.3 ± 1.5 | 111.4 ± 3.6 |
| Hypoxanthine | 103.3 ± 1.0 | 106.5 ± 6.0 | 101.9 ± 0.4 | 109.6 ± 2.0 |
| Indole-3-acetamide | 103.6 ± 5.6 | 90.6 ± 6.1 | 91.2 ± 4.6 | 106.8 ± 1.8 |
| Inosine | 97.3 ± 5.7 | 113.8 ± 0.9 | 104.2 ± 0.6 | 111.3 ± 2.8 |
| Inosine 5'-monophosphate | 86.8 ± 5.4 | 113.3 ± 3.8 | 97.1 ± 5.9 | 108.6 ± 5.7 |
| Isocitric Acid | 109.7 ± 5.0 | 114.6 ± 4.1 | 92.5 ± 4.1 | 95.7 ± 6.0 |
| Isoleucine | 88.1 ± 3.9 | 92.9 ± 5.5 | 104.8 ± 2.7 | 88.5 ± 0.8 |
| Lactic Acid | 100.4 ± 1.1 | 97.5 ± 3.1 | 109.4 ± 0.9 | 105.4 ± 5.0 |
| Leucine | 88.9 ± 4.7 | 95.0 ± 2.0 | 106.4 ± 6.3 | 99.4 ± 2.5 |
| Levulinic acid | 104.9 ± 1.7 | 97.7 ± 1.6 | 110.6 ± 4.7 | 100.5 ± 3.5 |
| Lysine | 99.7 ± 4.3 | 106.2 ± 2.1 | 91.0 ± 4.9 | 113.7 ± 5.2 |
| Malic Acid | 107.7 ± 5.9 | 95.6 ± 3.5 | 90.3 ± 4.5 | 94.6 ± 3.1 |
| Mannose-1 with Mannose-6-Phosphate | 85.5 ± 0.9 | 102.5 ± 5.5 | 97.0 ± 0.2 | 104.8 ± 0.5 |
| Mesaconic acid | 97.8 ± 3.2 | 113.4 ± 1.1 | 101.1 ± 1.2 | 91.6 ± 5.6 |
| Methionine | 99.2 ± 5.6 | 98.9 ± 0.5 | 105.6 ± 6.0 | 95.5 ± 2.1 |
| Methionine sulfoxide | 87.1 ± 3.2 | 96.3 ± 2.8 | 102.6 ± 4.7 | 87.7 ± 5.9 |
| Methylmalonic Acid | 94.4 ± 6.0 | 91.9 ± 5.1 | 93.8 ± 5.6 | 98.5 ± 4.4 |
| N1-Acetylspermidine | 102.5 ± 5.4 | 100.5 ± 4.7 | 109.8 ± 0.4 | 87.1 ± 2.6 |
| N1-Acetylspermine | 90.5 ± 1.8 | 103.0 ± 1.8 | 109.0 ± 2.9 | 85.2 ± 3.6 |
| N8-Acetylspermidine | 109.4 ± 3.1 | 86.4 ± 2.0 | 88.5 ± 2.0 | 109.3 ± 3.7 |
| N-Acetyl-Aspartyl-Glutamate | 94.5 ± 5.0 | 114.4 ± 0.8 | 88.6 ± 4.5 | 109.3 ± 4.3 |
| N-Acetyl-Aspartyl-Glutamyl-Glutamate | 94.4 ± 1.4 | 109.9 ± 0.3 | 113.9 ± 1.8 | 102.6 ± 1.8 |
| N-Acetyl-Galactosamine | 94.0 ± 1.9 | 105.9 ± 3.2 | 113.7 ± 6.0 | 106.1 ± 3.0 |
| N-Acetyl-Leucine | 110.1 ± 0.4 | 106.1 ± 6.0 | 112.9 ± 5.0 | 113.9 ± 4.9 |
| N-Acetyl-Methionine | 89.1 ± 5.4 | 105.6 ± 3.4 | 94.9 ± 0.7 | 110.9 ± 5.5 |
| N-Acetylneuraminic acid | 105.6 ± 4.5 | 88.7 ± 0.9 | 86.7 ± 1.2 | 88.3 ± 6.3 |
| N-Acetyl-Proline | 88.1 ± 1.5 | 110.5 ± 5.5 | 114.9 ± 1.5 | 109.0 ± 2.3 |
| N-Acetyl-Putrescine | 104.4 ± 0.3 | 94.6 ± 5.2 | 98.9 ± 1.0 | 100.9 ± 2.0 |
| N-Acetyl-Serine | 90.3 ± 5.1 | 114.6 ± 2.6 | 86.8 ± 3.7 | 97.9 ± 2.3 |
| N-Acetyl-Serotonin | 100.4 ± 2.4 | 113.3 ± 5.8 | 110.2 ± 4.2 | 112.8 ± 1.5 |
| N-Acyl-Homoserine-lactone | 108.8 ± 3.3 | 98.7 ± 2.6 | 114.4 ± 2.5 | 95.4 ± 5.9 |
| NAD | 89.4 ± 4.5 | 85.4 ± 0.6 | 108.8 ± 2.5 | 103.9 ± 3.9 |
| NADH | 106.8 ± 3.5 | 94.2 ± 1.3 | 89.9 ± 4.9 | 86.4 ± 3.2 |
| NADP | 90.9 ± 4.6 | 108.4 ± 1.0 | 97.8 ± 3.3 | 109.1 ± 2.8 |
| NADPH | 110.4 ± 0.3 | 96.9 ± 3.7 | 100.7 ± 4.8 | 85.4 ± 5.0 |
| N-alpha-Acetyl-lysine | 94.1 ± 0.5 | 97.6 ± 3.8 | 101.0 ± 6.2 | 88.7 ± 2.2 |
| N-alpha-Acetyl-Ornithine | 106.1 ± 6.0 | 107.7 ± 5.2 | 111.2 ± 3.3 | 101.1 ± 3.2 |
| N-Carbamoyl-Proline | 86.5 ± 0.9 | 87.3 ± 5.5 | 109.2 ± 4.8 | 112.0 ± 0.1 |
| N-Carbamoyl-Sarcosine | 86.4 ± 0.5 | 112.4 ± 5.8 | 105.1 ± 3.5 | 86.9 ± 2.4 |
| N-Epsilon-Acetyl-lysine | 102.7 ± 6.0 | 86.3 ± 4.4 | 88.5 ± 5.1 | 97.8 ± 0.3 |
| Nicotinamide | 86.7 ± 5.9 | 90.2 ± 0.9 | 99.0 ± 1.7 | 108.7 ± 5.7 |
| N-Methyl-Aspartic Acid | 102.3 ± 4.9 | 112.6 ± 6.1 | 103.0 ± 6.0 | 110.7 ± 3.4 |
| O-Phosphoethanolamine | 108.4 ± 0.4 | 112.8 ± 1.2 | 109.6 ± 0.4 | 114.3 ± 0.7 |
| Ophthalmic acid | 103.6 ± 1.6 | 88.4 ± 3.6 | 108.1 ± 3.2 | 105.0 ± 1.7 |
| Ornithine | 111.0 ± 0.7 | 87.6 ± 0.8 | 114.6 ± 0.7 | 107.9 ± 2.5 |
| Orotidine 5'-monophosphate | 90.0 ± 1.6 | 109.6 ± 1.6 | 90.7 ± 3.9 | 87.2 ± 2.0 |
| Oxaloacetate | 99.4 ± 3.6 | 100.2 ± 2.4 | 86.7 ± 3.4 | 111.8 ± 2.2 |
| Pantothenic acid | 93.9 ± 2.5 | 112.2 ± 1.4 | 103.9 ± 0.6 | 105.2 ± 1.8 |
| Phenylalanine | 97.0 ± 3.9 | 94.0 ± 0.8 | 92.5 ± 3.7 | 91.2 ± 1.5 |
| Phenylpyruvate | 87.9 ± 3.4 | 113.2 ± 2.4 | 108.1 ± 3.9 | 104.1 ± 3.5 |
| Phosphodimethylethanolamine | 93.1 ± 2.7 | 114.7 ± 1.2 | 88.2 ± 2.2 | 86.2 ± 1.2 |
| Phosphoenolpyruvate | 96.1 ± 1.1 | 92.4 ± 1.1 | 97.3 ± 3.1 | 114.9 ± 3.7 |
| Phosphorylcholine | 90.9 ± 3.4 | 91.6 ± 2.7 | 99.6 ± 2.4 | 92.4 ± 2.4 |
| Pipecolic acid | 109.9 ± 1.9 | 90.5 ± 3.5 | 98.2 ± 0.4 | 103.8 ± 3.4 |
| Proline | 95.9 ± 3.1 | 85.1 ± 1.4 | 113.2 ± 3.6 | 98.4 ± 3.6 |
| Putrescine | 90.4 ± 2.9 | 105.0 ± 3.2 | 92.9 ± 2.2 | 105.2 ± 1.7 |
| Pyridoxal | 105.8 ± 0.7 | 87.3 ± 0.7 | 113.3 ± 3.0 | 86.8 ± 3.0 |
| Pyridoxamine | 87.6 ± 1.5 | 108.3 ± 3.0 | 114.2 ± 2.1 | 105.3 ± 0.6 |
| Pyridoxine | 109.4 ± 1.3 | 96.5 ± 1.4 | 90.5 ± 3.8 | 97.0 ± 0.7 |
| Pyroglutamic acid | 104.1 ± 2.4 | 110.7 ± 1.6 | 104.9 ± 1.7 | 88.5 ± 3.5 |
| Pyrrolysine | 88.4 ± 1.8 | 93.8 ± 3.1 | 102.3 ± 0.3 | 112.5 ± 0.4 |
| Pyruvic Acid | 112.1 ± 2.6 | 94.7 ± 2.7 | 100.8 ± 1.5 | 88.8 ± 1.2 |
| Riboflavin | 85.4 ± 1.5 | 99.6 ± 0.9 | 112.7 ± 1.1 | 104.1 ± 1.6 |
| Ribose-5-Phosphate | 112.9 ± 1.0 | 94.6 ± 3.9 | 97.3 ± 0.6 | 111.8 ± 3.6 |
| Ribulose-1-Phosphate | 113.8 ± 2.6 | 107.3 ± 2.1 | 89.9 ± 3.6 | 112.6 ± 2.8 |
| Ribulose-5-Phosphate | 97.2 ± 3.2 | 104.6 ± 1.6 | 96.5 ± 3.8 | 93.7 ± 2.4 |
| Saccharopine | 104.0 ± 0.9 | 101.8 ± 2.3 | 94.7 ± 2.1 | 86.2 ± 1.2 |
| S-Adenosylhomocysteine | 99.2 ± 1.1 | 114.2 ± 3.9 | 94.0 ± 1.4 | 113.7 ± 2.1 |
| S-Adenosylmethioninamine | 104.7 ± 3.7 | 97.9 ± 2.7 | 90.8 ± 2.5 | 92.5 ± 3.5 |
| S-Adenosylmethionine | 108.0 ± 1.2 | 94.7 ± 1.4 | 110.9 ± 3.5 | 87.2 ± 2.4 |
| Sarcosine | 89.7 ± 0.4 | 110.8 ± 2.6 | 94.3 ± 1.1 | 93.4 ± 2.8 |
| Sedoheptulose-7-phosphate | 97.5 ± 2.0 | 110.0 ± 2.7 | 110.7 ± 1.4 | 91.6 ± 1.7 |
| Selenohomocysteine | 108.5 ± 3.4 | 94.4 ± 1.6 | 90.7 ± 3.6 | 88.1 ± 2.4 |
| Selenomethionine | 105.8 ± 2.1 | 89.0 ± 1.4 | 114.0 ± 3.1 | 87.5 ± 2.3 |
| Serine | 93.4 ± 1.7 | 105.5 ± 1.7 | 107.0 ± 3.6 | 92.6 ± 3.7 |
| S-Methyl-Thioinosine | 108.1 ± 3.0 | 85.6 ± 1.6 | 105.2 ± 0.7 | 91.2 ± 1.3 |
| Spermidine | 90.8 ± 2.6 | 109.8 ± 1.6 | 89.4 ± 0.6 | 105.3 ± 1.8 |
| Spermine | 103.0 ± 1.0 | 85.3 ± 1.4 | 89.9 ± 0.3 | 97.7 ± 2.5 |
| Succinate | 93.3 ± 1.5 | 106.1 ± 1.3 | 93.2 ± 2.1 | 90.8 ± 2.0 |
| Sucrose | 105.9 ± 2.0 | 99.4 ± 3.6 | 114.8 ± 1.9 | 114.3 ± 1.3 |
| Symmetric dimethylarginine | 111.9 ± 0.6 | 92.3 ± 3.7 | 113.5 ± 4.0 | 98.3 ± 1.4 |
| Taurine | 91.7 ± 2.1 | 97.6 ± 2.7 | 103.5 ± 0.5 | 86.5 ± 3.2 |
| Tetrahydroneopterin | 106.2 ± 2.4 | 111.2 ± 2.9 | 112.9 ± 0.5 | 94.1 ± 3.2 |
| Thiamine | 100.2 ± 3.5 | 99.7 ± 0.8 | 114.4 ± 1.8 | 87.2 ± 1.4 |
| Thiamine monophosphate | 107.0 ± 3.9 | 106.0 ± 1.7 | 98.5 ± 2.5 | 110.9 ± 2.2 |
| Threonic acid | 106.2 ± 2.7 | 90.4 ± 2.5 | 107.8 ± 3.6 | 105.0 ± 3.0 |
| Threonine | 105.4 ± 1.9 | 93.4 ± 3.0 | 97.5 ± 2.0 | 87.7 ± 3.9 |
| Thymidine | 112.0 ± 1.4 | 86.2 ± 1.5 | 102.1 ± 3.7 | 89.9 ± 4.0 |
| Thymidine Monophosphate | 99.3 ± 2.7 | 101.3 ± 2.6 | 104.4 ± 0.5 | 92.9 ± 2.8 |
| Thymine | 94.2 ± 2.1 | 91.8 ± 3.7 | 93.6 ± 3.1 | 111.3 ± 0.6 |
| Trigonelleine | 101.2 ± 3.6 | 85.9 ± 2.2 | 105.4 ± 0.3 | 107.9 ± 3.7 |
| Trimethylamine N-oxide | 91.4 ± 1.7 | 109.4 ± 2.0 | 90.0 ± 0.7 | 91.2 ± 2.8 |
| Trimethyllysine | 107.0 ± 3.9 | 109.4 ± 2.0 | 112.9 ± 1.5 | 92.2 ± 0.8 |
| Tryptophan | 108.2 ± 3.1 | 106.5 ± 3.6 | 103.9 ± 1.3 | 86.5 ± 3.4 |
| Tyramine | 94.5 ± 3.8 | 108.6 ± 2.7 | 100.7 ± 3.2 | 99.0 ± 0.9 |
| Tyrosine | 96.9 ± 1.7 | 88.7 ± 1.7 | 106.3 ± 0.5 | 104.9 ± 3.1 |
| Uracil | 107.3 ± 3.8 | 95.1 ± 3.2 | 86.0 ± 0.5 | 87.2 ± 2.2 |
| Uridine 5'-diphosphate | 98.0 ± 1.3 | 89.3 ± 1.0 | 102.1 ± 3.7 | 101.2 ± 2.4 |
| Uridine Monophosphate | 94.3 ± 0.6 | 92.6 ± 3.3 | 106.5 ± 1.1 | 105.4 ± 3.0 |
| Uridine-5-Diphosphogalactose | 89.1 ± 1.3 | 97.7 ± 2.5 | 108.2 ± 3.3 | 94.1 ± 2.3 |
| Urocanic acid | 100.6 ± 2.3 | 104.6 ± 1.2 | 102.9 ± 3.9 | 88.3 ± 0.8 |
| UTP | 100.5 ± 0.6 | 112.6 ± 3.7 | 114.3 ± 3.7 | 108.8 ± 0.4 |
| Valine | 98.9 ± 1.6 | 95.8 ± 0.9 | 95.8 ± 2.3 | 100.0 ± 0.9 |
| XMP | 94.8 ± 1.2 | 88.7 ± 2.7 | 114.5 ± 1.5 | 101.0 ± 3.2 |
| Xylulose-5-Phosphate | 97.5 ± 1.6 | 110.9 ± 2.7 | 89.8 ± 3.9 | 110.0 ± 3.3 |

### **Table S30**| Inter – day accuracy (as determined by % recovery) (Cell extract as matrix, n = 5). The concentrations of QC_LLOQ_, QC_Low_, QC_Mid_ and QC_High_ were 10 ng/mL, 25 ng/mL, 100 ng/mL and 500 ng/mL respectively.

| Accuracy, % | QC_LLOQ_ | QC_Low_ | QC_Mid_ | QC_High_ |
| --- | --- | --- | --- | --- |
| 1-Methylguanine | 98.4 ± 5.5 | 96.3 ± 1.2 | 104.0 ± 5.6 | 99.0 ± 0.7 |
| 1-Methylhistidine | 103.2 ± 5.4 | 97.8 ± 6.3 | 106.7 ± 6.3 | 96.6 ± 1.1 |
| 1-Methylnicotinamide | 107.0 ± 2.5 | 95.9 ± 1.9 | 106.2 ± 3.0 | 95.4 ± 0.4 |
| 2-Aminoadipic Acid | 106.5 ± 2.7 | 98.7 ± 2.0 | 106.2 ± 5.0 | 99.1 ± 6.0 |
| 2-Deoxyadenosine | 109.0 ± 1.7 | 96.5 ± 4.4 | 104.6 ± 3.1 | 97.0 ± 2.6 |
| 2'-Deoxyadenosine 5'-monophosphate | 103.7 ± 1.2 | 98.1 ± 4.1 | 107.2 ± 3.0 | 97.9 ± 5.1 |
| 2'-deoxyadenosine 5'-triphosphate | 106.1 ± 4.0 | 98.9 ± 1.3 | 101.9 ± 5.9 | 97.3 ± 5.3 |
| 2'-Deoxycytidine-5'-monophosphate | 97.5 ± 3.2 | 97.9 ± 2.3 | 101.9 ± 5.1 | 97.8 ± 2.0 |
| 2-Deoxyguanosine | 102.0 ± 1.6 | 95.9 ± 3.1 | 105.6 ± 1.9 | 95.5 ± 2.5 |
| 2-hydroxybutyric acid | 98.2 ± 5.0 | 97.2 ± 5.9 | 104.2 ± 4.0 | 98.8 ± 0.8 |
| 2-hydroxyglutaric acid | 106.3 ± 3.8 | 96.8 ± 0.8 | 106.1 ± 0.2 | 96.0 ± 2.1 |
| 2-Hydroxyisocaproic acid | 105.6 ± 6.1 | 98.4 ± 5.5 | 106.1 ± 4.2 | 99.2 ± 0.7 |
| 2-ketobutyric acid | 101.5 ± 3.5 | 95.7 ± 4.3 | 106.8 ± 2.0 | 98.6 ± 4.8 |
| 2-oxoadipic acid | 106.7 ± 0.9 | 99.1 ± 5.5 | 106.3 ± 4.5 | 97.2 ± 0.3 |
| 2-Phenylglycine | 107.6 ± 2.3 | 97.6 ± 4.7 | 102.0 ± 4.9 | 99.4 ± 4.9 |
| 2-phosphoglycerate | 107.4 ± 6.3 | 98.3 ± 3.1 | 104.5 ± 5.6 | 96.4 ± 5.4 |
| 3-hydroxy-3-methyl-2-oxobutanoate | 104.4 ± 1.5 | 98.9 ± 4.8 | 106.3 ± 4.8 | 99.3 ± 6.3 |
| 3-hydroxybutyric acid | 104.2 ± 4.0 | 98.6 ± 4.0 | 100.4 ± 2.6 | 96.8 ± 1.9 |
| 3-Methylhistidine | 102.2 ± 3.7 | 98.5 ± 1.9 | 107.0 ± 4.2 | 96.9 ± 3.9 |
| 3-Phosphoglyceraldehyde | 106.4 ± 1.1 | 98.4 ± 5.2 | 101.5 ± 1.5 | 99.3 ± 1.6 |
| 3-phosphoglycerate | 99.5 ± 1.5 | 99.6 ± 6.0 | 100.7 ± 3.2 | 97.2 ± 0.5 |
| 4-amino-butyric acid | 103.0 ± 0.3 | 97.4 ± 5.5 | 106.9 ± 4.4 | 95.8 ± 5.3 |
| 4-aspartyl-phosphate | 108.8 ± 0.4 | 98.6 ± 2.9 | 102.1 ± 2.2 | 95.9 ± 2.6 |
| 4-Hydroxybenzoic acid | 106.5 ± 5.7 | 96.5 ± 1.7 | 105.5 ± 6.1 | 97.8 ± 1.9 |
| 4-Hydroxyphenylacetylglycine | 105.4 ± 0.2 | 97.8 ± 6.1 | 103.9 ± 3.1 | 96.1 ± 4.2 |
| 4-methyl-2-oxopentanoate | 100.2 ± 2.3 | 96.8 ± 3.0 | 103.5 ± 1.9 | 98.5 ± 6.2 |
| 4-Oxoglutaramate | 105.5 ± 5.0 | 96.3 ± 1.2 | 104.6 ± 0.6 | 96.5 ± 3.3 |
| 5-Hydroxyindolepyruvate | 107.1 ± 4.0 | 98.5 ± 4.2 | 103.1 ± 4.6 | 96.9 ± 3.7 |
| 5-Hydroxylysine | 104.6 ± 5.5 | 99.0 ± 2.7 | 104.0 ± 5.6 | 97.1 ± 1.3 |
| 5-Hydroxy-Tryptophan | 105.7 ± 3.0 | 95.8 ± 0.4 | 100.3 ± 0.9 | 97.7 ± 0.3 |
| 5-Phosphoribosylamine | 107.9 ± 2.4 | 97.7 ± 1.7 | 106.7 ± 1.6 | 95.5 ± 0.3 |
| 6-phosphogluconate | 103.2 ± 6.0 | 96.2 ± 2.0 | 105.6 ± 1.2 | 99.0 ± 5.1 |
| Acetylcarnitine | 108.2 ± 1.4 | 98.5 ± 5.1 | 103.8 ± 0.6 | 96.5 ± 4.4 |
| Aconitic Acid | 100.7 ± 0.7 | 98.4 ± 3.6 | 108.3 ± 4.4 | 99.4 ± 5.6 |
| Adenine | 100.8 ± 3.9 | 98.4 ± 1.9 | 106.4 ± 0.8 | 99.2 ± 5.5 |
| Adenosine | 108.7 ± 1.9 | 97.2 ± 2.0 | 108.6 ± 5.8 | 95.6 ± 4.6 |
| Adenosine 3',5'-cyclic monophosphate | 105.2 ± 4.4 | 98.1 ± 2.8 | 104.2 ± 1.5 | 99.1 ± 5.6 |
| Adenosine Monophosphate | 107.4 ± 2.0 | 97.2 ± 2.1 | 100.8 ± 5.5 | 98.1 ± 5.1 |
| Adenosine-5'-diphosphate | 104.3 ± 5.6 | 97.1 ± 1.6 | 105.2 ± 4.8 | 99.1 ± 5.7 |
| Adenosine-5'-triphosphate | 105.1 ± 6.1 | 97.7 ± 1.9 | 102.6 ± 4.5 | 97.2 ± 6.1 |
| Adenylosuccinate | 101.0 ± 2.5 | 96.8 ± 1.7 | 103.3 ± 4.1 | 97.0 ± 2.2 |
| Alanine | 105.4 ± 6.2 | 99.2 ± 0.4 | 100.2 ± 4.4 | 97.4 ± 3.2 |
| alpha-ketoglutaric acid | 105.8 ± 5.5 | 95.3 ± 3.2 | 105.5 ± 2.6 | 96.9 ± 3.8 |
| Anserine | 105.9 ± 2.5 | 98.1 ± 3.3 | 105.3 ± 5.0 | 97.1 ± 2.8 |
| Arginine | 106.8 ± 2.3 | 95.3 ± 3.0 | 106.5 ± 4.7 | 96.8 ± 2.8 |
| Argininosuccinic Acid | 107.0 ± 3.6 | 96.4 ± 5.9 | 108.5 ± 5.3 | 98.5 ± 4.0 |
| Asparagine | 107.2 ± 4.0 | 97.7 ± 3.9 | 107.3 ± 1.4 | 97.1 ± 3.4 |
| Aspartic acid | 106.9 ± 5.8 | 99.6 ± 1.2 | 108.3 ± 0.5 | 95.7 ± 0.2 |
| Asymmetric dimethylarginine | 104.6 ± 4.6 | 99.4 ± 0.2 | 105.6 ± 2.9 | 96.0 ± 5.6 |
| Azelaic acid | 106.3 ± 2.5 | 96.0 ± 2.2 | 105.8 ± 3.3 | 96.5 ± 0.2 |
| beta-Alanine | 105.8 ± 5.5 | 97.3 ± 0.4 | 107.4 ± 4.6 | 98.4 ± 6.3 |
| Betaine | 102.9 ± 2.0 | 96.5 ± 2.3 | 100.7 ± 0.1 | 96.7 ± 6.0 |
| Biopterin | 104.0 ± 1.2 | 95.8 ± 2.6 | 103.9 ± 0.6 | 96.5 ± 5.6 |
| Biotin | 108.1 ± 1.8 | 98.3 ± 5.8 | 104.8 ± 5.5 | 97.1 ± 3.2 |
| Carnitine | 107.6 ± 3.8 | 96.5 ± 1.1 | 103.6 ± 4.8 | 97.3 ± 3.0 |
| Carnosine | 107.7 ± 5.9 | 96.2 ± 6.2 | 108.2 ± 0.6 | 95.8 ± 2.4 |
| CDP-choline | 105.2 ± 4.4 | 96.1 ± 6.1 | 100.0 ± 4.3 | 98.6 ± 0.3 |
| cGMP | 105.9 ± 5.1 | 98.7 ± 3.0 | 105.0 ± 6.1 | 96.8 ± 3.4 |
| Choline | 104.9 ± 4.5 | 98.8 ± 5.3 | 106.0 ± 2.0 | 97.6 ± 0.1 |
| Cinnamic acid | 103.7 ± 2.3 | 98.0 ± 5.7 | 105.8 ± 1.3 | 97.4 ± 6.0 |
| Citraconic acid | 103.0 ± 4.3 | 97.7 ± 2.4 | 107.1 ± 2.1 | 96.0 ± 4.5 |
| Citramalic acid | 105.6 ± 4.9 | 95.7 ± 2.6 | 106.4 ± 4.0 | 98.1 ± 4.6 |
| Citric Acid | 103.1 ± 0.4 | 98.3 ± 0.6 | 106.7 ± 4.4 | 98.3 ± 0.4 |
| Citrulline | 101.0 ± 5.9 | 97.6 ± 5.4 | 106.1 ± 4.7 | 95.7 ± 6.3 |
| Creatine | 107.7 ± 2.9 | 95.5 ± 1.8 | 107.6 ± 1.7 | 95.8 ± 3.0 |
| Creatinine | 106.1 ± 1.0 | 97.7 ± 1.8 | 105.1 ± 0.5 | 98.6 ± 6.1 |
| Cystathionine | 107.8 ± 4.9 | 97.1 ± 6.0 | 104.7 ± 1.0 | 95.8 ± 0.4 |
| Cysteamine | 105.9 ± 5.1 | 99.4 ± 2.1 | 106.4 ± 3.3 | 98.7 ± 2.0 |
| Cysteine | 105.2 ± 5.7 | 96.5 ± 1.2 | 105.0 ± 1.8 | 97.8 ± 1.0 |
| Cysteinylglycine | 108.1 ± 2.1 | 97.2 ± 2.4 | 104.7 ± 1.6 | 95.4 ± 1.7 |
| Cystine | 107.4 ± 4.9 | 99.3 ± 2.9 | 104.2 ± 5.3 | 98.6 ± 3.7 |
| Cytidine | 100.1 ± 5.2 | 96.0 ± 1.0 | 108.2 ± 3.3 | 97.6 ± 3.8 |
| Cytidine 3',5'-cyclic monophosphate | 107.3 ± 3.0 | 96.0 ± 2.9 | 107.2 ± 3.6 | 95.3 ± 1.1 |
| Cytidine 5'-diphosphate | 104.7 ± 2.2 | 98.0 ± 1.7 | 106.4 ± 1.7 | 96.3 ± 1.4 |
| Cytidine 5'-triphosphate | 108.6 ± 1.4 | 98.6 ± 1.3 | 105.9 ± 3.4 | 96.2 ± 1.6 |
| Cytidine monophosphate | 107.8 ± 5.3 | 96.0 ± 1.4 | 104.9 ± 0.7 | 97.4 ± 1.0 |
| Cytosine | 104.6 ± 4.4 | 97.1 ± 4.3 | 107.0 ± 5.9 | 96.7 ± 5.1 |
| dADP | 102.6 ± 3.4 | 97.8 ± 3.1 | 106.6 ± 6.1 | 97.2 ± 2.8 |
| Diethanolamine | 108.2 ± 4.6 | 99.5 ± 0.2 | 104.7 ± 5.9 | 97.6 ± 6.1 |
| Dihydrobiopterin | 106.6 ± 1.5 | 97.9 ± 0.3 | 108.7 ± 0.6 | 98.1 ± 5.3 |
| Dihydroneopterin triphosphate | 106.5 ± 2.1 | 97.7 ± 5.7 | 103.5 ± 2.5 | 97.1 ± 4.6 |
| Dihydroxyacetonephosphate | 108.0 ± 3.4 | 96.4 ± 3.9 | 102.1 ± 5.4 | 97.6 ± 5.0 |
| Dimethylglycine | 108.8 ± 1.4 | 98.8 ± 2.6 | 107.0 ± 2.1 | 98.5 ± 6.1 |
| Ectoine | 111.6 ± 2.9 | 97.2 ± 3.5 | 107.7 ± 2.4 | 96.4 ± 2.3 |
| Erythrose-4-phosphate | 104.8 ± 0.6 | 97.3 ± 1.0 | 107.3 ± 1.8 | 95.5 ± 2.9 |
| FAD | 103.1 ± 2.2 | 96.0 ± 2.5 | 105.4 ± 2.7 | 96.1 ± 2.3 |
| Folic acid | 106.8 ± 3.7 | 99.6 ± 5.0 | 103.6 ± 4.9 | 97.1 ± 3.1 |
| Fructose-6-phosphate | 100.5 ± 2.9 | 95.7 ± 5.9 | 103.1 ± 4.7 | 96.5 ± 3.8 |
| Fumaric Acid | 103.7 ± 0.9 | 99.1 ± 5.5 | 103.2 ± 5.3 | 96.0 ± 3.4 |
| Galactosamine | 94.3 ± 1.7 | 97.4 ± 1.5 | 106.5 ± 2.9 | 97.2 ± 1.1 |
| Galactose-1-phosphate | 97.3 ± 1.7 | 99.3 ± 5.3 | 103.2 ± 3.6 | 97.1 ± 2.4 |
| Galactose-6-Phosphate | 97.6 ± 1.4 | 95.8 ± 6.0 | 106.6 ± 2.4 | 97.4 ± 3.1 |
| gamma-Glutamylcysteine | 108.1 ± 1.0 | 97.0 ± 4.4 | 103.9 ± 3.6 | 97.3 ± 4.3 |
| Glucosamine | 102.8 ± 2.9 | 95.1 ± 2.3 | 108.5 ± 5.5 | 97.8 ± 3.1 |
| Glucosamine-6-Phosphate | 104.0 ± 3.3 | 95.1 ± 2.3 | 103.3 ± 4.9 | 96.7 ± 4.2 |
| Glucose | 100.4 ± 2.9 | 95.2 ± 4.4 | 107.8 ± 0.7 | 97.3 ± 5.1 |
| Glucose-1-phosphate | 98.6 ± 1.3 | 98.1 ± 1.8 | 103.5 ± 0.6 | 97.6 ± 1.8 |
| Glucose-6-Phosphate | 95.0 ± 2.4 | 99.2 ± 5.1 | 103.6 ± 2.8 | 97.9 ± 1.4 |
| Glucuronic acid | 106.9 ± 2.0 | 95.6 ± 1.0 | 108.6 ± 1.4 | 98.3 ± 0.6 |
| Glutamic acid | 91.8 ± 0.3 | 98.2 ± 0.5 | 105.2 ± 6.2 | 96.4 ± 4.1 |
| Glutamine | 86.4 ± 0.6 | 96.6 ± 6.3 | 104.7 ± 0.4 | 96.0 ± 5.6 |
| Glutathione | 93.3 ± 0.9 | 85.3 ± 1.2 | 106.4 ± 0.8 | 95.3 ± 4.2 |
| Glutathionylspermine | 109.3 ± 1.2 | 104.7 ± 3.7 | 106.3 ± 5.1 | 98.2 ± 0.4 |
| Glycerophosphocholine | 103.3 ± 1.6 | 104.7 ± 3.3 | 104.8 ± 3.1 | 97.5 ± 2.3 |
| Glycine | 109.2 ± 3.2 | 91.7 ± 0.5 | 104.4 ± 0.9 | 98.5 ± 1.0 |
| Glycolic acid | 107.8 ± 2.1 | 106.3 ± 1.7 | 106.5 ± 4.7 | 97.2 ± 0.4 |
| Guanidinoacetic acid | 97.6 ± 3.4 | 96.6 ± 2.4 | 105.2 ± 5.1 | 96.1 ± 2.3 |
| Guanidinobutanoic acid | 94.7 ± 0.5 | 108.7 ± 1.6 | 106.2 ± 2.8 | 98.5 ± 4.3 |
| Guanine | 94.5 ± 2.5 | 101.3 ± 2.0 | 108.1 ± 5.6 | 96.6 ± 5.9 |
| Guanosine | 106.2 ± 3.3 | 103.6 ± 1.2 | 108.6 ± 6.2 | 98.3 ± 2.2 |
| Guanosine 5'-diphosphate | 110.7 ± 1.0 | 108.7 ± 1.6 | 108.5 ± 4.8 | 97.6 ± 4.7 |
| Guanosine-5'-monophosphate | 86.6 ± 2.1 | 90.7 ± 2.8 | 106.9 ± 5.7 | 97.0 ± 0.5 |
| Hercynine | 93.7 ± 0.8 | 96.0 ± 0.8 | 107.4 ± 3.3 | 97.7 ± 5.9 |
| Hippuric acid | 112.1 ± 3.1 | 86.5 ± 2.9 | 106.9 ± 2.1 | 97.6 ± 2.4 |
| Histamine | 88.4 ± 3.9 | 111.9 ± 3.6 | 100.7 ± 4.8 | 96.0 ± 5.8 |
| Histidine | 112.5 ± 0.6 | 93.4 ± 2.5 | 105.8 ± 5.0 | 97.7 ± 2.8 |
| Homoalanine | 108.2 ± 1.2 | 112.9 ± 2.8 | 108.2 ± 5.9 | 96.5 ± 1.3 |
| Homoarginine | 87.8 ± 1.3 | 100.9 ± 3.5 | 107.1 ± 4.9 | 86.3 ± 2.4 |
| Homocysteine | 93.5 ± 0.5 | 88.0 ± 3.1 | 112.4 ± 1.5 | 92.4 ± 1.5 |
| Homogentisic acid | 108.4 ± 3.7 | 109.7 ± 3.0 | 85.9 ± 3.5 | 94.3 ± 0.3 |
| Homoleucine | 88.2 ± 3.3 | 104.7 ± 3.7 | 91.5 ± 1.5 | 104.0 ± 1.3 |
| Homovanillic acid | 94.5 ± 3.9 | 113.7 ± 0.8 | 100.8 ± 1.5 | 97.3 ± 2.1 |
| Hydantoin-5-propionate | 98.6 ± 1.1 | 85.8 ± 3.4 | 113.6 ± 1.5 | 98.6 ± 2.7 |
| Hypotaurine | 105.3 ± 0.7 | 88.4 ± 2.4 | 94.6 ± 2.4 | 97.7 ± 0.7 |
| Hypoxanthine | 96.5 ± 2.6 | 102.4 ± 0.7 | 90.6 ± 1.4 | 114.2 ± 1.8 |
| Indole-3-acetamide | 106.6 ± 0.9 | 101.3 ± 3.6 | 97.1 ± 1.5 | 109.0 ± 3.7 |
| Inosine | 92.2 ± 3.7 | 101.2 ± 2.3 | 105.6 ± 0.9 | 109.2 ± 2.2 |
| Inosine 5'-monophosphate | 89.0 ± 1.7 | 97.9 ± 2.5 | 95.0 ± 0.6 | 97.5 ± 1.7 |
| Isocitric Acid | 104.9 ± 0.6 | 104.9 ± 2.5 | 96.9 ± 2.8 | 103.6 ± 0.5 |
| Isoleucine | 107.9 ± 3.7 | 98.2 ± 3.0 | 88.5 ± 0.5 | 98.4 ± 2.8 |
| Lactic Acid | 112.8 ± 1.1 | 99.9 ± 1.1 | 86.5 ± 2.1 | 110.7 ± 0.3 |
| Leucine | 87.0 ± 3.5 | 111.8 ± 0.9 | 89.6 ± 3.5 | 100.9 ± 1.0 |
| Levulinic acid | 101.7 ± 3.3 | 97.3 ± 1.0 | 85.8 ± 0.8 | 98.6 ± 2.3 |
| Lysine | 104.8 ± 1.9 | 99.3 ± 3.8 | 103.2 ± 2.6 | 90.7 ± 1.4 |
| Malic Acid | 107.4 ± 3.3 | 87.8 ± 2.3 | 87.5 ± 1.9 | 94.8 ± 3.3 |
| Mannose-1 with Mannose-6-Phosphate | 97.9 ± 0.6 | 106.6 ± 0.4 | 90.6 ± 2.9 | 88.9 ± 0.7 |
| Mesaconic acid | 96.5 ± 3.9 | 95.6 ± 1.3 | 88.1 ± 2.9 | 95.5 ± 3.4 |
| Methionine | 104.7 ± 3.8 | 101.2 ± 1.7 | 103.8 ± 1.5 | 106.8 ± 2.0 |
| Methionine sulfoxide | 99.5 ± 3.3 | 96.7 ± 1.0 | 86.6 ± 2.6 | 102.1 ± 2.0 |
| Methylmalonic Acid | 102.0 ± 2.4 | 108.7 ± 2.8 | 93.3 ± 1.9 | 85.4 ± 0.9 |
| N1-Acetylspermidine | 95.0 ± 1.5 | 96.1 ± 1.6 | 90.4 ± 0.4 | 87.8 ± 2.3 |
| N1-Acetylspermine | 85.3 ± 1.8 | 96.3 ± 2.4 | 108.3 ± 3.4 | 100.1 ± 2.5 |
| N8-Acetylspermidine | 98.2 ± 1.0 | 100.0 ± 1.5 | 85.5 ± 3.5 | 91.6 ± 3.5 |
| N-Acetyl-Aspartyl-Glutamate | 97.3 ± 1.7 | 113.0 ± 2.8 | 104.9 ± 3.6 | 99.7 ± 2.5 |
| N-Acetyl-Aspartyl-Glutamyl-Glutamate | 85.3 ± 1.8 | 110.1 ± 2.0 | 112.5 ± 3.0 | 109.9 ± 1.4 |
| N-Acetyl-Galactosamine | 104.4 ± 0.4 | 88.6 ± 3.2 | 112.8 ± 3.2 | 88.4 ± 3.4 |
| N-Acetyl-Leucine | 105.4 ± 2.3 | 93.3 ± 1.3 | 86.7 ± 0.7 | 110.0 ± 3.1 |
| N-Acetyl-Methionine | 89.3 ± 1.2 | 89.4 ± 1.2 | 105.9 ± 3.7 | 109.4 ± 2.3 |
| N-Acetylneuraminic acid | 96.5 ± 2.9 | 94.3 ± 1.5 | 97.9 ± 1.2 | 104.9 ± 2.2 |
| N-Acetyl-Proline | 97.1 ± 2.5 | 86.9 ± 2.3 | 100.7 ± 1.4 | 93.2 ± 2.7 |
| N-Acetyl-Putrescine | 107.8 ± 2.9 | 94.5 ± 3.1 | 109.1 ± 3.0 | 98.1 ± 0.6 |
| N-Acetyl-Serine | 99.6 ± 3.0 | 91.8 ± 1.8 | 102.4 ± 2.5 | 114.2 ± 3.3 |
| N-Acetyl-Serotonin | 114.1 ± 3.2 | 111.7 ± 3.3 | 106.7 ± 1.4 | 93.3 ± 2.6 |
| N-Acyl-Homoserine-lactone | 113.6 ± 3.9 | 85.8 ± 1.1 | 101.5 ± 2.8 | 111.9 ± 0.9 |
| NAD | 106.1 ± 4.0 | 107.1 ± 1.1 | 89.8 ± 0.7 | 99.0 ± 3.2 |
| NADH | 99.6 ± 1.1 | 89.7 ± 3.5 | 109.8 ± 1.8 | 104.2 ± 2.5 |
| NADP | 106.0 ± 3.9 | 114.6 ± 0.7 | 100.8 ± 3.6 | 97.6 ± 1.8 |
| NADPH | 92.4 ± 2.4 | 94.7 ± 1.6 | 105.0 ± 3.5 | 101.1 ± 1.1 |
| N-alpha-Acetyl-lysine | 107.7 ± 1.9 | 85.4 ± 3.9 | 96.9 ± 3.9 | 85.9 ± 0.3 |
| N-alpha-Acetyl-Ornithine | 105.5 ± 3.4 | 107.1 ± 0.5 | 97.8 ± 3.2 | 87.7 ± 1.2 |
| N-Carbamoyl-Proline | 89.2 ± 1.0 | 101.3 ± 2.2 | 90.2 ± 0.7 | 85.6 ± 3.1 |
| N-Carbamoyl-Sarcosine | 103.3 ± 3.5 | 112.3 ± 2.1 | 99.8 ± 1.0 | 97.4 ± 2.5 |
| N-Epsilon-Acetyl-lysine | 113.6 ± 3.4 | 100.1 ± 1.8 | 96.2 ± 3.6 | 103.3 ± 0.6 |
| Nicotinamide | 90.3 ± 1.8 | 85.5 ± 3.7 | 91.2 ± 3.9 | 87.0 ± 3.3 |
| N-Methyl-Aspartic Acid | 87.7 ± 3.3 | 107.8 ± 1.0 | 94.0 ± 1.2 | 104.2 ± 1.7 |
| O-Phosphoethanolamine | 114.8 ± 2.7 | 90.1 ± 0.3 | 114.5 ± 2.6 | 110.4 ± 2.7 |
| Ophthalmic acid | 114.1 ± 2.2 | 112.4 ± 2.3 | 111.4 ± 1.0 | 85.8 ± 1.5 |
| Ornithine | 95.8 ± 3.5 | 111.4 ± 3.0 | 86.8 ± 2.9 | 96.8 ± 0.8 |
| Orotidine 5'-monophosphate | 110.7 ± 3.2 | 91.4 ± 2.8 | 88.4 ± 3.0 | 107.5 ± 3.0 |
| Oxaloacetate | 107.0 ± 3.2 | 108.5 ± 1.5 | 102.7 ± 0.6 | 86.3 ± 0.7 |
| Pantothenic acid | 106.0 ± 0.4 | 101.6 ± 1.6 | 88.7 ± 3.1 | 93.6 ± 1.4 |
| Phenylalanine | 112.4 ± 3.7 | 112.4 ± 2.8 | 110.3 ± 2.5 | 109.7 ± 3.7 |
| Phenylpyruvate | 103.7 ± 0.3 | 90.0 ± 3.5 | 102.1 ± 1.6 | 92.7 ± 0.6 |
| Phosphodimethylethanolamine | 90.1 ± 2.4 | 89.5 ± 1.2 | 90.3 ± 2.6 | 95.5 ± 2.4 |
| Phosphoenolpyruvate | 92.2 ± 2.1 | 98.9 ± 0.7 | 85.5 ± 3.0 | 86.3 ± 3.2 |
| Phosphorylcholine | 109.5 ± 3.8 | 101.8 ± 3.9 | 109.4 ± 1.7 | 102.7 ± 1.4 |
| Pipecolic acid | 103.5 ± 3.7 | 103.4 ± 1.7 | 90.2 ± 2.0 | 87.1 ± 1.7 |
| Proline | 101.7 ± 3.1 | 105.3 ± 2.2 | 114.8 ± 3.1 | 96.1 ± 1.6 |
| Putrescine | 100.4 ± 3.4 | 104.3 ± 3.4 | 94.3 ± 3.6 | 112.9 ± 1.4 |
| Pyridoxal | 101.6 ± 0.5 | 106.4 ± 1.7 | 86.6 ± 2.8 | 89.5 ± 3.5 |
| Pyridoxamine | 85.3 ± 3.3 | 104.7 ± 1.1 | 102.2 ± 2.0 | 105.1 ± 3.3 |
| Pyridoxine | 96.0 ± 0.5 | 97.4 ± 4.0 | 103.0 ± 2.4 | 112.5 ± 1.9 |
| Pyroglutamic acid | 97.2 ± 2.2 | 106.5 ± 0.7 | 109.3 ± 3.8 | 112.5 ± 2.6 |
| Pyrrolysine | 102.0 ± 3.1 | 100.0 ± 1.9 | 110.7 ± 2.3 | 99.1 ± 0.6 |
| Pyruvic Acid | 87.6 ± 2.4 | 109.6 ± 3.5 | 93.3 ± 3.5 | 86.5 ± 0.9 |
| Riboflavin | 106.3 ± 3.9 | 109.8 ± 1.7 | 102.8 ± 2.4 | 87.4 ± 2.4 |
| Ribose-5-Phosphate | 95.6 ± 3.5 | 114.7 ± 0.7 | 93.1 ± 3.9 | 86.6 ± 3.1 |
| Ribulose-1-Phosphate | 102.1 ± 2.3 | 89.5 ± 2.2 | 86.0 ± 2.3 | 95.7 ± 1.4 |
| Ribulose-5-Phosphate | 94.5 ± 0.8 | 113.6 ± 3.2 | 93.5 ± 1.0 | 88.7 ± 1.3 |
| Saccharopine | 87.1 ± 1.4 | 113.7 ± 2.2 | 94.2 ± 2.7 | 112.8 ± 3.5 |
| S-Adenosylhomocysteine | 114.3 ± 0.6 | 94.1 ± 0.6 | 107.9 ± 1.2 | 105.6 ± 2.7 |
| S-Adenosylmethioninamine | 100.8 ± 1.1 | 107.6 ± 2.6 | 112.8 ± 1.3 | 101.3 ± 1.0 |
| S-Adenosylmethionine | 100.8 ± 3.5 | 108.0 ± 2.3 | 95.7 ± 3.0 | 91.3 ± 3.3 |
| Sarcosine | 111.2 ± 0.5 | 96.7 ± 2.9 | 86.2 ± 3.4 | 94.5 ± 3.9 |
| Sedoheptulose-7-phosphate | 91.4 ± 1.7 | 90.7 ± 0.7 | 93.6 ± 0.6 | 113.0 ± 4.0 |
| Selenohomocysteine | 91.4 ± 3.3 | 113.0 ± 4.0 | 91.5 ± 3.2 | 106.7 ± 1.1 |
| Selenomethionine | 112.7 ± 3.9 | 86.2 ± 2.4 | 89.5 ± 0.7 | 101.3 ± 0.8 |
| Serine | 86.5 ± 2.6 | 112.7 ± 1.3 | 99.8 ± 3.0 | 110.5 ± 3.9 |
| S-Methyl-Thioinosine | 95.2 ± 0.4 | 90.1 ± 3.6 | 85.3 ± 0.4 | 87.8 ± 1.9 |
| Spermidine | 110.6 ± 2.5 | 106.0 ± 2.9 | 96.2 ± 2.1 | 103.4 ± 2.1 |
| Spermine | 101.5 ± 1.4 | 88.8 ± 3.3 | 100.4 ± 2.6 | 102.9 ± 2.7 |
| Succinate | 98.3 ± 2.3 | 93.3 ± 1.2 | 94.5 ± 1.8 | 97.7 ± 1.7 |
| Sucrose | 100.1 ± 0.5 | 86.4 ± 1.3 | 114.3 ± 3.5 | 111.0 ± 3.1 |
| Symmetric dimethylarginine | 102.0 ± 0.8 | 86.6 ± 1.3 | 88.1 ± 1.1 | 89.9 ± 3.1 |
| Taurine | 104.9 ± 0.9 | 87.9 ± 1.7 | 106.1 ± 0.5 | 85.5 ± 2.0 |
| Tetrahydroneopterin | 112.4 ± 1.9 | 87.3 ± 3.9 | 98.4 ± 3.5 | 105.5 ± 3.4 |
| Thiamine | 96.6 ± 3.4 | 99.1 ± 2.9 | 114.7 ± 1.8 | 106.0 ± 2.6 |
| Thiamine monophosphate | 110.3 ± 1.3 | 98.6 ± 2.2 | 95.1 ± 3.9 | 92.3 ± 0.9 |
| Threonic acid | 99.6 ± 1.1 | 109.4 ± 3.0 | 105.2 ± 1.2 | 103.7 ± 1.9 |
| Threonine | 93.6 ± 2.9 | 105.5 ± 1.3 | 103.0 ± 0.8 | 108.0 ± 1.6 |
| Thymidine | 95.7 ± 1.0 | 98.8 ± 1.8 | 110.9 ± 1.6 | 90.6 ± 2.4 |
| Thymidine Monophosphate | 86.7 ± 2.9 | 114.3 ± 1.5 | 108.2 ± 3.5 | 96.1 ± 3.5 |
| Thymine | 90.3 ± 3.9 | 87.9 ± 1.6 | 92.3 ± 3.9 | 109.3 ± 3.5 |
| Trigonelleine | 109.5 ± 0.4 | 102.8 ± 0.3 | 101.8 ± 1.4 | 94.3 ± 2.4 |
| Trimethylamine N-oxide | 89.1 ± 2.2 | 111.5 ± 1.4 | 110.2 ± 1.8 | 106.0 ± 2.9 |
| Trimethyllysine | 85.8 ± 1.7 | 85.1 ± 2.2 | 98.6 ± 2.4 | 98.2 ± 3.6 |
| Tryptophan | 88.7 ± 3.9 | 107.4 ± 2.9 | 91.7 ± 1.9 | 91.2 ± 3.7 |
| Tyramine | 106.2 ± 0.4 | 107.5 ± 1.7 | 93.5 ± 3.4 | 105.9 ± 3.1 |
| Tyrosine | 96.0 ± 1.1 | 92.9 ± 2.1 | 105.0 ± 2.6 | 101.0 ± 3.0 |
| Uracil | 97.7 ± 3.8 | 100.9 ± 3.3 | 104.7 ± 3.9 | 95.0 ± 3.4 |
| Uridine 5'-diphosphate | 106.6 ± 3.3 | 102.3 ± 1.1 | 86.8 ± 3.7 | 102.9 ± 0.5 |
| Uridine Monophosphate | 95.7 ± 0.9 | 99.3 ± 0.6 | 91.8 ± 3.0 | 99.7 ± 1.7 |
| Uridine-5-Diphosphogalactose | 100.0 ± 2.4 | 105.6 ± 1.2 | 95.1 ± 3.6 | 112.5 ± 2.2 |
| Urocanic acid | 88.1 ± 2.1 | 95.1 ± 0.3 | 112.0 ± 0.6 | 105.3 ± 2.3 |
| UTP | 92.5 ± 3.4 | 109.7 ± 3.3 | 89.0 ± 1.4 | 113.5 ± 3.5 |
| Valine | 92.8 ± 4.0 | 90.8 ± 4.0 | 95.6 ± 2.8 | 88.0 ± 3.1 |
| XMP | 100.4 ± 1.9 | 98.8 ± 0.4 | 105.6 ± 0.8 | 89.4 ± 3.8 |
| Xylulose-5-Phosphate | 93.8 ± 2.6 | 91.0 ± 2.7 | 114.9 ± 3.0 | 107.1 ± 3.3 |

### **Table S31** | Inter – day accuracy (as determined by % recovery) (Urine extract as matrix, n = 5). The concentrations of QC_LLOQ_, QC_Low_, QC_Mid_ and QC_High_ were 10 ng/mL, 25 ng/mL, 100 ng/mL and 500 ng/mL respectively.

| Accuracy, % | QC_LLOQ_ | QC_Low_ | QC_Mid_ | QC_High_ |
| --- | --- | --- | --- | --- |
| 1-Methylguanine | 95.8 ± 3.8 | 109.1 ± 3.5 | 91.7 ± 5.6 | 97.6 ± 4.0 |
| 1-Methylhistidine | 85.3 ± 0.6 | 104.5 ± 1.1 | 113.1 ± 1.2 | 108.7 ± 2.4 |
| 1-Methylnicotinamide | 85.6 ± 3.8 | 95.5 ± 5.1 | 102.9 ± 5.4 | 93.0 ± 2.9 |
| 2-Aminoadipic Acid | 104.6 ± 2.7 | 96.7 ± 5.6 | 112.2 ± 5.6 | 111.9 ± 2.3 |
| 2-Deoxyadenosine | 108.3 ± 2.3 | 100.3 ± 2.6 | 106.3 ± 1.4 | 96.1 ± 3.2 |
| 2'-Deoxyadenosine 5'-monophosphate | 108.4 ± 1.5 | 92.5 ± 4.0 | 92.6 ± 3.2 | 110.9 ± 3.6 |
| 2'-deoxyadenosine 5'-triphosphate | 105.6 ± 2.1 | 85.1 ± 2.4 | 88.0 ± 3.4 | 111.5 ± 4.7 |
| 2'-Deoxycytidine-5'-monophosphate | 97.6 ± 1.2 | 93.9 ± 4.3 | 104.3 ± 4.2 | 94.0 ± 5.9 |
| 2-Deoxyguanosine | 87.8 ± 2.7 | 110.1 ± 0.6 | 94.3 ± 3.6 | 94.9 ± 2.3 |
| 2-hydroxybutyric acid | 93.7 ± 3.3 | 96.8 ± 2.8 | 114.8 ± 3.3 | 94.2 ± 2.4 |
| 2-hydroxyglutaric acid | 87.6 ± 0.6 | 87.1 ± 4.5 | 105.8 ± 5.3 | 91.6 ± 2.8 |
| 2-Hydroxyisocaproic acid | 92.5 ± 0.5 | 101.7 ± 5.2 | 89.1 ± 4.1 | 92.2 ± 2.7 |
| 2-ketobutyric acid | 101.1 ± 2.0 | 105.3 ± 3.5 | 91.3 ± 1.6 | 101.9 ± 3.8 |
| 2-oxoadipic acid | 93.2 ± 1.0 | 102.0 ± 2.2 | 102.0 ± 0.3 | 103.6 ± 2.8 |
| 2-Phenylglycine | 87.8 ± 0.5 | 97.4 ± 5.8 | 89.8 ± 0.7 | 93.0 ± 6.1 |
| 2-phosphoglycerate | 100.0 ± 5.2 | 86.7 ± 2.2 | 107.2 ± 2.6 | 88.1 ± 3.5 |
| 3-hydroxy-3-methyl-2-oxobutanoate | 90.1 ± 3.1 | 104.0 ± 2.2 | 96.6 ± 2.7 | 98.0 ± 2.0 |
| 3-hydroxybutyric acid | 92.9 ± 0.7 | 94.6 ± 2.7 | 89.2 ± 0.4 | 85.5 ± 1.5 |
| 3-Methylhistidine | 110.4 ± 2.5 | 96.2 ± 2.4 | 102.5 ± 6.2 | 96.3 ± 5.2 |
| 3-Phosphoglyceraldehyde | 105.7 ± 5.4 | 100.3 ± 5.2 | 96.0 ± 1.2 | 106.3 ± 5.1 |
| 3-phosphoglycerate | 94.4 ± 0.8 | 104.1 ± 3.5 | 112.5 ± 2.1 | 98.7 ± 0.5 |
| 4-amino-butyric acid | 103.9 ± 1.2 | 88.8 ± 4.3 | 92.3 ± 5.8 | 101.3 ± 6.1 |
| 4-aspartyl-phosphate | 91.8 ± 0.4 | 95.0 ± 4.3 | 111.2 ± 0.8 | 97.6 ± 1.0 |
| 4-Hydroxybenzoic acid | 99.6 ± 2.2 | 90.0 ± 3.5 | 93.7 ± 3.1 | 104.2 ± 3.5 |
| 4-Hydroxyphenylacetylglycine | 101.0 ± 6.2 | 89.2 ± 5.3 | 95.7 ± 2.7 | 101.3 ± 5.4 |
| 4-methyl-2-oxopentanoate | 105.4 ± 3.7 | 95.1 ± 3.8 | 111.5 ± 4.8 | 90.0 ± 1.6 |
| 4-Oxoglutaramate | 109.9 ± 3.0 | 94.8 ± 3.6 | 102.2 ± 3.3 | 100.2 ± 0.3 |
| 5-Hydroxyindolepyruvate | 93.3 ± 0.5 | 110.0 ± 6.2 | 107.5 ± 0.1 | 91.3 ± 5.0 |
| 5-Hydroxylysine | 98.0 ± 5.6 | 97.9 ± 0.8 | 92.5 ± 0.7 | 111.8 ± 3.5 |
| 5-Hydroxy-Tryptophan | 89.2 ± 4.3 | 111.1 ± 2.5 | 102.1 ± 3.1 | 92.7 ± 5.0 |
| 5-Phosphoribosylamine | 97.8 ± 4.2 | 95.8 ± 1.1 | 101.8 ± 2.0 | 87.8 ± 0.7 |
| 6-phosphogluconate | 114.7 ± 3.1 | 92.8 ± 1.7 | 111.9 ± 4.5 | 87.8 ± 4.6 |
| Acetylcarnitine | 102.1 ± 4.0 | 87.9 ± 5.0 | 107.9 ± 5.6 | 92.2 ± 4.2 |
| Aconitic Acid | 111.8 ± 5.5 | 105.2 ± 3.6 | 109.9 ± 4.4 | 86.0 ± 3.0 |
| Adenine | 97.6 ± 4.9 | 113.9 ± 5.2 | 97.4 ± 4.5 | 90.2 ± 3.0 |
| Adenosine | 112.4 ± 1.9 | 91.8 ± 5.4 | 91.8 ± 1.5 | 96.3 ± 3.8 |
| Adenosine 3',5'-cyclic monophosphate | 86.7 ± 2.0 | 94.7 ± 0.5 | 89.9 ± 1.1 | 100.2 ± 2.1 |
| Adenosine Monophosphate | 112.0 ± 0.9 | 96.6 ± 5.9 | 102.0 ± 4.6 | 90.6 ± 3.4 |
| Adenosine-5'-diphosphate | 95.3 ± 2.7 | 88.2 ± 5.6 | 102.1 ± 4.0 | 87.3 ± 5.4 |
| Adenosine-5'-triphosphate | 95.7 ± 3.7 | 97.3 ± 0.2 | 86.0 ± 1.1 | 99.1 ± 4.6 |
| Adenylosuccinate | 93.4 ± 3.1 | 97.1 ± 3.8 | 88.8 ± 2.1 | 103.0 ± 0.2 |
| Alanine | 92.8 ± 0.7 | 97.3 ± 3.0 | 100.7 ± 2.6 | 104.9 ± 2.9 |
| alpha-ketoglutaric acid | 99.6 ± 3.0 | 100.9 ± 2.2 | 105.1 ± 0.8 | 103.9 ± 1.6 |
| Anserine | 108.4 ± 3.8 | 108.8 ± 5.4 | 91.7 ± 3.5 | 85.9 ± 1.9 |
| Arginine | 101.0 ± 0.5 | 98.5 ± 1.2 | 90.2 ± 4.7 | 101.7 ± 0.7 |
| Argininosuccinic Acid | 94.0 ± 5.4 | 107.2 ± 2.6 | 112.1 ± 3.2 | 98.6 ± 0.2 |
| Asparagine | 104.8 ± 3.5 | 97.9 ± 2.9 | 102.0 ± 3.3 | 112.3 ± 2.3 |
| Aspartic acid | 89.1 ± 3.2 | 109.3 ± 0.6 | 104.4 ± 4.7 | 89.0 ± 1.7 |
| Asymmetric dimethylarginine | 95.5 ± 3.1 | 90.4 ± 6.3 | 107.6 ± 2.5 | 111.2 ± 0.2 |
| Azelaic acid | 93.3 ± 3.5 | 110.2 ± 4.1 | 114.9 ± 4.2 | 104.7 ± 1.4 |
| beta-Alanine | 96.8 ± 3.4 | 104.5 ± 5.1 | 113.6 ± 0.4 | 106.0 ± 2.1 |
| Betaine | 104.8 ± 1.2 | 98.0 ± 3.8 | 92.6 ± 1.7 | 93.7 ± 3.2 |
| Biopterin | 89.3 ± 1.7 | 86.2 ± 4.6 | 103.4 ± 5.3 | 88.5 ± 0.4 |
| Biotin | 99.4 ± 3.9 | 87.4 ± 5.5 | 102.4 ± 1.7 | 104.9 ± 4.8 |
| Carnitine | 94.1 ± 5.0 | 95.4 ± 2.0 | 89.5 ± 1.7 | 92.8 ± 3.6 |
| Carnosine | 113.6 ± 5.7 | 97.0 ± 4.7 | 90.8 ± 2.9 | 113.0 ± 0.7 |
| CDP-choline | 93.1 ± 4.2 | 105.4 ± 2.8 | 112.9 ± 5.6 | 113.3 ± 5.8 |
| cGMP | 85.2 ± 6.3 | 105.3 ± 1.5 | 101.9 ± 2.8 | 91.6 ± 4.5 |
| Choline | 112.7 ± 5.0 | 99.2 ± 1.2 | 89.2 ± 1.4 | 103.3 ± 3.7 |
| Cinnamic acid | 103.0 ± 1.1 | 98.6 ± 1.0 | 105.4 ± 2.2 | 107.5 ± 0.6 |
| Citraconic acid | 104.5 ± 3.6 | 106.5 ± 5.1 | 94.4 ± 6.0 | 91.1 ± 0.5 |
| Citramalic acid | 111.5 ± 6.0 | 105.8 ± 6.1 | 86.6 ± 5.1 | 86.8 ± 0.2 |
| Citric Acid | 98.2 ± 2.3 | 113.9 ± 1.6 | 93.1 ± 4.8 | 102.9 ± 5.3 |
| Citrulline | 97.9 ± 4.7 | 111.7 ± 1.4 | 88.2 ± 3.8 | 112.5 ± 5.0 |
| Creatine | 88.8 ± 5.2 | 110.2 ± 4.8 | 106.4 ± 4.9 | 86.9 ± 4.5 |
| Creatinine | 101.9 ± 0.6 | 98.5 ± 4.8 | 99.6 ± 1.4 | 109.0 ± 0.7 |
| Cystathionine | 85.9 ± 0.4 | 91.7 ± 0.6 | 104.2 ± 3.6 | 98.4 ± 2.5 |
| Cysteamine | 86.0 ± 1.6 | 106.2 ± 0.9 | 104.7 ± 5.7 | 89.4 ± 2.9 |
| Cysteine | 111.2 ± 5.2 | 102.5 ± 0.8 | 99.3 ± 2.6 | 85.8 ± 4.6 |
| Cysteinylglycine | 113.5 ± 1.6 | 107.4 ± 2.6 | 98.0 ± 1.0 | 114.7 ± 1.6 |
| Cystine | 96.9 ± 0.5 | 97.2 ± 6.1 | 91.2 ± 5.8 | 100.3 ± 2.4 |
| Cytidine | 100.2 ± 3.7 | 88.4 ± 5.0 | 112.7 ± 5.1 | 92.6 ± 6.2 |
| Cytidine 3',5'-cyclic monophosphate | 90.3 ± 3.9 | 90.6 ± 2.6 | 108.7 ± 3.0 | 94.4 ± 0.9 |
| Cytidine 5'-diphosphate | 107.7 ± 5.5 | 85.6 ± 5.1 | 102.3 ± 3.6 | 88.0 ± 4.5 |
| Cytidine 5'-triphosphate | 109.2 ± 0.8 | 103.1 ± 5.2 | 101.2 ± 3.5 | 108.3 ± 2.4 |
| Cytidine monophosphate | 98.6 ± 3.4 | 90.5 ± 1.0 | 99.7 ± 3.1 | 91.2 ± 0.8 |
| Cytosine | 93.4 ± 5.8 | 114.0 ± 0.9 | 86.0 ± 0.7 | 98.2 ± 2.6 |
| dADP | 91.4 ± 4.0 | 87.7 ± 6.0 | 110.9 ± 1.1 | 112.4 ± 3.2 |
| Diethanolamine | 109.8 ± 0.2 | 93.2 ± 1.1 | 98.6 ± 2.6 | 99.7 ± 6.0 |
| Dihydrobiopterin | 113.6 ± 3.9 | 85.3 ± 4.9 | 87.6 ± 4.4 | 113.5 ± 4.0 |
| Dihydroneopterin triphosphate | 88.8 ± 3.6 | 108.2 ± 5.0 | 98.1 ± 3.8 | 97.0 ± 5.4 |
| Dihydroxyacetonephosphate | 86.3 ± 3.6 | 94.8 ± 4.8 | 104.1 ± 3.8 | 114.1 ± 0.5 |
| Dimethylglycine | 85.4 ± 5.3 | 89.5 ± 3.2 | 102.0 ± 1.8 | 89.5 ± 0.6 |
| Ectoine | 111.5 ± 4.5 | 110.8 ± 4.4 | 113.8 ± 0.1 | 113.6 ± 5.8 |
| Erythrose-4-phosphate | 114.8 ± 0.4 | 104.9 ± 3.5 | 99.4 ± 5.3 | 111.7 ± 5.2 |
| FAD | 108.0 ± 2.3 | 106.8 ± 3.2 | 106.5 ± 3.8 | 102.5 ± 3.8 |
| Folic acid | 96.4 ± 5.9 | 94.3 ± 4.5 | 86.0 ± 6.2 | 85.7 ± 1.5 |
| Fructose-6-phosphate | 87.4 ± 0.7 | 92.3 ± 1.4 | 106.9 ± 0.4 | 102.2 ± 4.5 |
| Fumaric Acid | 93.8 ± 1.4 | 97.8 ± 4.8 | 98.4 ± 1.2 | 102.6 ± 5.0 |
| Galactosamine | 94.1 ± 3.1 | 96.9 ± 1.9 | 114.6 ± 5.7 | 92.8 ± 4.5 |
| Galactose-1-phosphate | 107.8 ± 2.3 | 110.2 ± 0.9 | 86.4 ± 3.2 | 85.8 ± 3.3 |
| Galactose-6-Phosphate | 92.8 ± 4.0 | 110.8 ± 1.4 | 95.6 ± 3.3 | 86.3 ± 2.4 |
| gamma-Glutamylcysteine | 90.9 ± 6.2 | 97.3 ± 6.3 | 114.5 ± 5.5 | 96.1 ± 0.6 |
| Glucosamine | 108.7 ± 5.4 | 106.5 ± 3.9 | 105.7 ± 0.1 | 96.9 ± 4.9 |
| Glucosamine-6-Phosphate | 98.3 ± 4.2 | 92.2 ± 5.5 | 97.7 ± 4.3 | 100.5 ± 2.0 |
| Glucose | 85.9 ± 6.1 | 90.3 ± 6.2 | 99.9 ± 1.7 | 97.9 ± 6.1 |
| Glucose-1-phosphate | 99.3 ± 5.4 | 96.6 ± 4.5 | 102.3 ± 1.7 | 98.8 ± 3.9 |
| Glucose-6-Phosphate | 106.1 ± 5.6 | 112.4 ± 5.6 | 104.0 ± 0.6 | 111.6 ± 5.0 |
| Glucuronic acid | 103.5 ± 5.7 | 96.7 ± 2.9 | 106.4 ± 4.9 | 85.1 ± 5.8 |
| Glutamic acid | 89.2 ± 3.0 | 111.5 ± 4.7 | 99.8 ± 5.0 | 93.5 ± 5.7 |
| Glutamine | 110.2 ± 5.1 | 86.9 ± 4.5 | 92.3 ± 0.5 | 97.7 ± 3.0 |
| Glutathione | 92.5 ± 6.1 | 113.0 ± 2.3 | 114.8 ± 5.2 | 113.3 ± 3.0 |
| Glutathionylspermine | 91.3 ± 0.1 | 104.3 ± 4.3 | 106.5 ± 3.8 | 113.5 ± 5.0 |
| Glycerophosphocholine | 88.0 ± 4.3 | 90.5 ± 1.2 | 92.4 ± 1.7 | 96.0 ± 4.9 |
| Glycine | 86.7 ± 4.4 | 89.3 ± 2.0 | 86.7 ± 3.3 | 103.1 ± 6.3 |
| Glycolic acid | 95.7 ± 1.3 | 95.0 ± 2.8 | 109.2 ± 5.2 | 95.1 ± 3.1 |
| Guanidinoacetic acid | 89.4 ± 6.3 | 105.7 ± 4.4 | 86.8 ± 1.1 | 102.5 ± 2.3 |
| Guanidinobutanoic acid | 94.3 ± 2.7 | 110.7 ± 4.9 | 101.9 ± 1.0 | 91.7 ± 0.7 |
| Guanine | 105.2 ± 6.2 | 105.0 ± 3.3 | 107.5 ± 5.8 | 106.3 ± 4.0 |
| Guanosine | 112.3 ± 5.0 | 106.2 ± 4.1 | 104.3 ± 4.6 | 89.4 ± 5.7 |
| Guanosine 5'-diphosphate | 99.1 ± 3.6 | 98.0 ± 3.6 | 102.6 ± 1.5 | 89.1 ± 2.4 |
| Guanosine-5'-monophosphate | 103.9 ± 4.5 | 92.1 ± 0.6 | 101.5 ± 5.0 | 96.1 ± 5.1 |
| Hercynine | 88.5 ± 4.4 | 86.0 ± 4.2 | 106.6 ± 4.0 | 98.2 ± 1.5 |
| Hippuric acid | 109.7 ± 2.9 | 97.1 ± 4.3 | 91.1 ± 0.7 | 85.1 ± 6.2 |
| Histamine | 103.0 ± 3.5 | 94.1 ± 5.8 | 107.9 ± 5.4 | 106.0 ± 5.0 |
| Histidine | 90.4 ± 1.3 | 106.6 ± 5.2 | 100.4 ± 0.8 | 104.4 ± 3.4 |
| Homoalanine | 91.0 ± 0.3 | 97.5 ± 5.6 | 110.9 ± 0.8 | 107.8 ± 4.2 |
| Homoarginine | 103.6 ± 4.3 | 104.6 ± 2.4 | 113.8 ± 3.2 | 99.5 ± 2.0 |
| Homocysteine | 112.1 ± 1.4 | 92.7 ± 4.5 | 101.5 ± 2.3 | 85.1 ± 4.4 |
| Homogentisic acid | 85.1 ± 0.2 | 86.9 ± 0.7 | 98.9 ± 1.6 | 99.0 ± 3.1 |
| Homoleucine | 88.3 ± 5.4 | 91.4 ± 3.9 | 88.0 ± 4.3 | 114.8 ± 1.0 |
| Homovanillic acid | 110.8 ± 1.5 | 106.2 ± 5.0 | 101.4 ± 5.7 | 108.5 ± 5.5 |
| Hydantoin-5-propionate | 113.5 ± 4.2 | 89.0 ± 4.2 | 107.9 ± 0.1 | 96.3 ± 3.3 |
| Hypotaurine | 102.9 ± 4.9 | 85.5 ± 1.9 | 87.2 ± 5.6 | 103.7 ± 0.8 |
| Hypoxanthine | 89.8 ± 1.0 | 86.5 ± 0.7 | 89.0 ± 1.3 | 89.3 ± 5.4 |
| Indole-3-acetamide | 105.5 ± 4.0 | 100.4 ± 6.1 | 90.9 ± 5.3 | 95.3 ± 0.3 |
| Inosine | 85.1 ± 4.3 | 93.8 ± 1.8 | 97.9 ± 0.8 | 113.1 ± 3.3 |
| Inosine 5'-monophosphate | 105.3 ± 4.2 | 99.7 ± 3.6 | 107.9 ± 0.9 | 110.6 ± 2.8 |
| Isocitric Acid | 112.3 ± 3.6 | 104.8 ± 0.6 | 98.3 ± 4.4 | 105.4 ± 2.0 |
| Isoleucine | 87.3 ± 1.7 | 97.1 ± 0.4 | 97.6 ± 0.6 | 90.8 ± 6.0 |
| Lactic Acid | 109.8 ± 6.2 | 97.7 ± 3.0 | 105.0 ± 3.9 | 99.0 ± 5.9 |
| Leucine | 89.8 ± 5.3 | 114.1 ± 1.5 | 104.4 ± 3.4 | 112.7 ± 4.4 |
| Levulinic acid | 114.6 ± 4.0 | 99.3 ± 5.9 | 110.3 ± 4.3 | 112.5 ± 0.7 |
| Lysine | 111.1 ± 5.5 | 106.8 ± 1.2 | 87.3 ± 4.4 | 111.9 ± 5.1 |
| Malic Acid | 99.0 ± 3.7 | 90.2 ± 5.2 | 86.7 ± 2.5 | 114.8 ± 1.6 |
| Mannose-1 with Mannose-6-Phosphate | 104.7 ± 0.6 | 110.2 ± 3.3 | 105.4 ± 1.2 | 113.8 ± 1.9 |
| Mesaconic acid | 109.2 ± 3.0 | 102.8 ± 0.8 | 88.1 ± 5.4 | 91.9 ± 1.2 |
| Methionine | 105.9 ± 1.6 | 106.5 ± 4.6 | 101.0 ± 5.6 | 102.7 ± 1.2 |
| Methionine sulfoxide | 91.7 ± 6.0 | 87.1 ± 5.1 | 104.8 ± 6.2 | 107.0 ± 1.6 |
| Methylmalonic Acid | 106.0 ± 3.9 | 109.6 ± 3.1 | 108.4 ± 3.5 | 106.1 ± 2.2 |
| N1-Acetylspermidine | 85.4 ± 0.6 | 104.7 ± 3.9 | 107.6 ± 4.9 | 87.0 ± 5.5 |
| N1-Acetylspermine | 89.6 ± 1.9 | 97.7 ± 2.1 | 87.5 ± 1.7 | 107.6 ± 4.2 |
| N8-Acetylspermidine | 104.1 ± 6.0 | 97.9 ± 3.0 | 94.4 ± 2.3 | 102.2 ± 1.7 |
| N-Acetyl-Aspartyl-Glutamate | 111.7 ± 3.6 | 99.7 ± 5.3 | 94.0 ± 4.8 | 102.7 ± 3.8 |
| N-Acetyl-Aspartyl-Glutamyl-Glutamate | 100.5 ± 0.2 | 89.0 ± 3.0 | 88.4 ± 1.3 | 93.0 ± 4.0 |
| N-Acetyl-Galactosamine | 89.4 ± 3.8 | 97.9 ± 1.7 | 104.3 ± 0.4 | 103.5 ± 3.2 |
| N-Acetyl-Leucine | 90.1 ± 5.4 | 114.2 ± 4.5 | 112.9 ± 5.3 | 103.6 ± 1.2 |
| N-Acetyl-Methionine | 92.4 ± 4.5 | 99.0 ± 2.4 | 88.4 ± 0.5 | 90.9 ± 5.6 |
| N-Acetylneuraminic acid | 107.9 ± 3.0 | 98.9 ± 5.8 | 92.0 ± 2.0 | 101.6 ± 2.7 |
| N-Acetyl-Proline | 98.9 ± 1.2 | 94.8 ± 4.9 | 96.8 ± 0.9 | 91.3 ± 2.7 |
| N-Acetyl-Putrescine | 93.7 ± 2.8 | 85.1 ± 4.1 | 92.0 ± 1.8 | 88.0 ± 0.2 |
| N-Acetyl-Serine | 95.5 ± 5.3 | 95.9 ± 5.9 | 104.0 ± 2.6 | 112.5 ± 2.0 |
| N-Acetyl-Serotonin | 88.5 ± 1.9 | 114.1 ± 2.4 | 106.9 ± 3.1 | 99.8 ± 0.1 |
| N-Acyl-Homoserine-lactone | 111.8 ± 0.2 | 89.5 ± 5.5 | 85.0 ± 6.0 | 96.5 ± 1.3 |
| NAD | 104.0 ± 3.6 | 108.9 ± 1.5 | 105.2 ± 0.1 | 100.2 ± 3.7 |
| NADH | 85.2 ± 4.5 | 102.1 ± 4.2 | 89.5 ± 4.6 | 108.2 ± 2.1 |
| NADP | 107.5 ± 2.9 | 88.6 ± 3.5 | 93.6 ± 1.9 | 113.3 ± 5.0 |
| NADPH | 95.8 ± 1.8 | 114.8 ± 5.4 | 85.6 ± 5.1 | 102.9 ± 5.2 |
| N-alpha-Acetyl-lysine | 91.2 ± 2.0 | 96.5 ± 6.1 | 110.2 ± 1.7 | 85.1 ± 3.2 |
| N-alpha-Acetyl-Ornithine | 100.1 ± 3.5 | 93.6 ± 2.8 | 103.1 ± 3.0 | 93.6 ± 2.4 |
| N-Carbamoyl-Proline | 95.6 ± 1.8 | 104.4 ± 3.9 | 114.8 ± 5.1 | 108.3 ± 5.5 |
| N-Carbamoyl-Sarcosine | 104.9 ± 2.7 | 95.5 ± 2.4 | 88.6 ± 0.6 | 109.3 ± 5.7 |
| N-Epsilon-Acetyl-lysine | 105.1 ± 0.3 | 107.3 ± 2.1 | 114.7 ± 4.5 | 110.7 ± 1.0 |
| Nicotinamide | 99.4 ± 1.6 | 99.0 ± 0.5 | 88.2 ± 4.5 | 89.9 ± 1.9 |
| N-Methyl-Aspartic Acid | 110.4 ± 1.7 | 88.0 ± 1.4 | 96.4 ± 3.2 | 92.3 ± 6.2 |
| O-Phosphoethanolamine | 96.3 ± 1.9 | 106.2 ± 1.6 | 99.6 ± 2.0 | 94.1 ± 2.9 |
| Ophthalmic acid | 114.4 ± 1.5 | 87.6 ± 3.4 | 85.5 ± 0.6 | 109.8 ± 2.4 |
| Ornithine | 91.2 ± 2.3 | 88.7 ± 2.7 | 112.7 ± 3.2 | 110.3 ± 1.8 |
| Orotidine 5'-monophosphate | 85.8 ± 1.7 | 96.9 ± 2.9 | 103.4 ± 3.4 | 104.2 ± 2.5 |
| Oxaloacetate | 100.2 ± 3.0 | 106.4 ± 3.8 | 97.6 ± 0.8 | 98.8 ± 3.0 |
| Pantothenic acid | 89.1 ± 2.6 | 101.7 ± 3.1 | 86.6 ± 3.8 | 95.7 ± 3.1 |
| Phenylalanine | 109.3 ± 2.5 | 109.7 ± 1.8 | 96.6 ± 1.7 | 98.1 ± 0.6 |
| Phenylpyruvate | 89.9 ± 1.5 | 107.0 ± 3.6 | 91.1 ± 2.3 | 110.2 ± 0.7 |
| Phosphodimethylethanolamine | 109.8 ± 0.4 | 102.2 ± 0.7 | 87.8 ± 1.7 | 104.6 ± 1.5 |
| Phosphoenolpyruvate | 89.4 ± 3.3 | 95.5 ± 2.2 | 106.0 ± 1.3 | 111.6 ± 3.7 |
| Phosphorylcholine | 90.9 ± 0.5 | 105.6 ± 1.1 | 107.6 ± 3.0 | 104.7 ± 3.5 |
| Pipecolic acid | 111.4 ± 0.7 | 113.0 ± 1.3 | 103.5 ± 1.6 | 105.6 ± 3.7 |
| Proline | 86.1 ± 2.0 | 85.8 ± 2.3 | 100.4 ± 2.2 | 114.7 ± 1.3 |
| Putrescine | 93.1 ± 0.9 | 103.0 ± 3.8 | 107.4 ± 0.7 | 100.2 ± 1.6 |
| Pyridoxal | 92.2 ± 4.0 | 106.0 ± 1.7 | 94.1 ± 0.8 | 97.1 ± 3.4 |
| Pyridoxamine | 97.0 ± 2.4 | 115.0 ± 3.7 | 89.6 ± 3.3 | 103.3 ± 2.8 |
| Pyridoxine | 88.7 ± 3.6 | 103.5 ± 1.6 | 113.6 ± 0.9 | 89.2 ± 1.9 |
| Pyroglutamic acid | 100.8 ± 1.3 | 106.5 ± 2.4 | 97.7 ± 4.0 | 111.9 ± 3.5 |
| Pyrrolysine | 106.3 ± 1.4 | 101.1 ± 1.8 | 88.5 ± 1.1 | 105.0 ± 2.0 |
| Pyruvic Acid | 85.5 ± 2.4 | 94.5 ± 2.5 | 86.3 ± 1.9 | 101.6 ± 0.7 |
| Riboflavin | 89.4 ± 3.8 | 109.9 ± 1.4 | 95.9 ± 0.4 | 98.3 ± 0.4 |
| Ribose-5-Phosphate | 90.4 ± 2.6 | 94.2 ± 0.7 | 89.3 ± 2.6 | 109.2 ± 1.6 |
| Ribulose-1-Phosphate | 110.3 ± 1.6 | 113.8 ± 3.4 | 114.8 ± 0.5 | 113.3 ± 3.2 |
| Ribulose-5-Phosphate | 110.4 ± 0.6 | 99.6 ± 2.5 | 109.7 ± 2.7 | 108.0 ± 3.1 |
| Saccharopine | 91.1 ± 3.0 | 110.1 ± 0.7 | 112.0 ± 1.6 | 100.8 ± 0.6 |
| S-Adenosylhomocysteine | 109.1 ± 1.1 | 92.5 ± 1.1 | 104.3 ± 1.7 | 96.5 ± 1.9 |
| S-Adenosylmethioninamine | 95.1 ± 1.9 | 114.8 ± 1.2 | 94.0 ± 3.5 | 107.5 ± 3.5 |
| S-Adenosylmethionine | 94.6 ± 2.6 | 100.2 ± 2.2 | 94.0 ± 0.8 | 113.0 ± 0.6 |
| Sarcosine | 88.7 ± 2.0 | 89.6 ± 0.9 | 111.4 ± 2.3 | 112.9 ± 1.8 |
| Sedoheptulose-7-phosphate | 100.8 ± 1.4 | 87.3 ± 1.4 | 109.5 ± 3.8 | 112.1 ± 1.6 |
| Selenohomocysteine | 108.6 ± 3.6 | 93.8 ± 0.7 | 99.9 ± 3.2 | 108.9 ± 1.3 |
| Selenomethionine | 96.1 ± 1.7 | 86.9 ± 2.8 | 103.7 ± 1.4 | 98.7 ± 4.0 |
| Serine | 107.1 ± 1.4 | 85.7 ± 3.8 | 99.8 ± 0.4 | 113.2 ± 1.7 |
| S-Methyl-Thioinosine | 93.0 ± 1.7 | 96.5 ± 3.1 | 113.3 ± 2.7 | 109.2 ± 3.1 |
| Spermidine | 109.2 ± 1.0 | 87.4 ± 0.6 | 89.6 ± 1.9 | 106.5 ± 2.8 |
| Spermine | 109.8 ± 2.9 | 96.9 ± 0.6 | 87.9 ± 0.7 | 92.1 ± 3.2 |
| Succinate | 113.4 ± 3.6 | 88.4 ± 3.4 | 113.3 ± 3.5 | 106.3 ± 2.9 |
| Sucrose | 110.7 ± 2.2 | 93.5 ± 2.2 | 111.3 ± 0.9 | 106.4 ± 1.5 |
| Symmetric dimethylarginine | 98.1 ± 1.9 | 85.1 ± 3.9 | 103.0 ± 1.9 | 98.1 ± 0.9 |
| Taurine | 93.8 ± 1.8 | 95.2 ± 3.6 | 101.4 ± 3.9 | 94.5 ± 1.6 |
| Tetrahydroneopterin | 86.7 ± 1.1 | 98.7 ± 1.2 | 102.8 ± 2.8 | 93.7 ± 2.0 |
| Thiamine | 94.9 ± 2.3 | 102.2 ± 2.1 | 99.7 ± 0.9 | 87.1 ± 3.5 |
| Thiamine monophosphate | 105.6 ± 2.1 | 88.8 ± 2.1 | 91.7 ± 2.0 | 114.9 ± 2.9 |
| Threonic acid | 105.6 ± 1.7 | 110.3 ± 2.9 | 100.7 ± 2.6 | 105.5 ± 2.5 |
| Threonine | 106.9 ± 0.7 | 109.9 ± 3.7 | 114.7 ± 1.9 | 97.3 ± 0.6 |
| Thymidine | 112.6 ± 1.7 | 98.7 ± 1.3 | 109.4 ± 2.5 | 106.2 ± 0.5 |
| Thymidine Monophosphate | 88.6 ± 0.9 | 90.0 ± 3.6 | 88.6 ± 2.3 | 113.9 ± 3.4 |
| Thymine | 86.0 ± 3.7 | 96.9 ± 1.5 | 111.6 ± 3.9 | 108.3 ± 3.8 |
| Trigonelleine | 91.6 ± 2.8 | 112.5 ± 3.4 | 85.1 ± 0.4 | 107.6 ± 2.1 |
| Trimethylamine N-oxide | 92.5 ± 3.0 | 87.8 ± 2.2 | 98.5 ± 2.4 | 96.2 ± 2.0 |
| Trimethyllysine | 109.1 ± 3.0 | 94.6 ± 3.1 | 95.3 ± 3.9 | 91.1 ± 0.7 |
| Tryptophan | 114.8 ± 3.4 | 100.6 ± 0.9 | 94.4 ± 3.7 | 104.3 ± 2.7 |
| Tyramine | 101.5 ± 2.6 | 88.2 ± 3.0 | 90.2 ± 1.8 | 87.8 ± 4.0 |
| Tyrosine | 107.7 ± 0.4 | 105.1 ± 1.2 | 112.7 ± 1.9 | 108.8 ± 2.0 |
| Uracil | 112.1 ± 1.9 | 95.5 ± 1.8 | 88.2 ± 2.4 | 88.1 ± 3.7 |
| Uridine 5'-diphosphate | 107.7 ± 1.3 | 114.0 ± 1.2 | 101.5 ± 1.5 | 97.1 ± 1.8 |
| Uridine Monophosphate | 107.5 ± 1.9 | 101.0 ± 2.7 | 96.2 ± 2.3 | 96.1 ± 3.0 |
| Uridine-5-Diphosphogalactose | 114.2 ± 1.5 | 103.7 ± 1.7 | 95.4 ± 0.6 | 85.1 ± 3.0 |
| Urocanic acid | 97.2 ± 2.8 | 103.3 ± 0.8 | 95.0 ± 0.6 | 99.3 ± 1.6 |
| UTP | 99.3 ± 3.9 | 91.1 ± 2.8 | 85.7 ± 0.5 | 85.7 ± 2.2 |
| Valine | 91.1 ± 2.9 | 100.5 ± 2.2 | 89.0 ± 2.7 | 88.6 ± 0.5 |
| XMP | 97.3 ± 1.0 | 106.1 ± 1.7 | 87.1 ± 0.4 | 95.1 ± 3.8 |
| Xylulose-5-Phosphate | 102.3 ± 0.6 | 86.9 ± 1.7 | 93.6 ± 2.3 | 112.6 ± 0.7 |

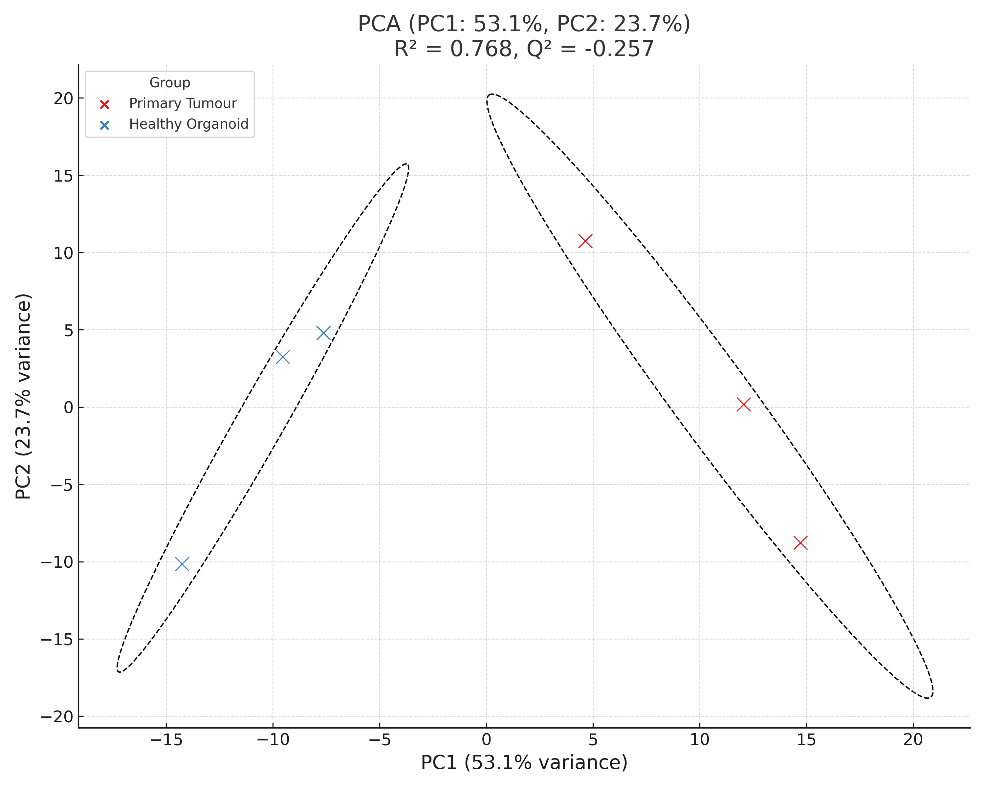

### **Figure S1** | Principal Component Analysis (PCA) of colorectal organoid metabolomic profiles. PCA shows clear group separation among Healthy and Tumor samples (n = 3 per group). Ellipses represent 95% confidence intervals around group centroids.

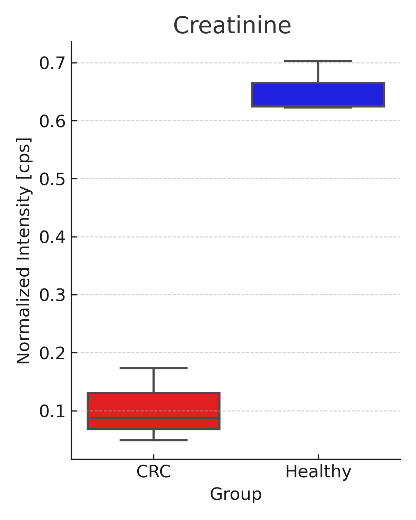

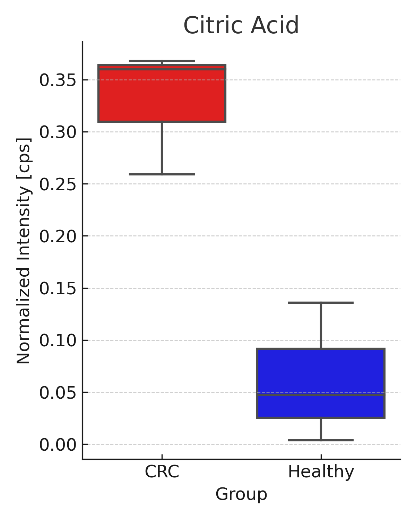

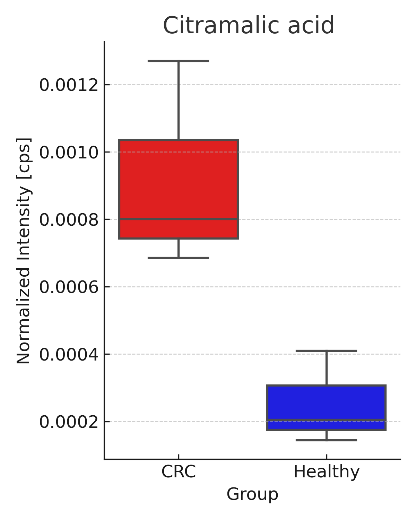

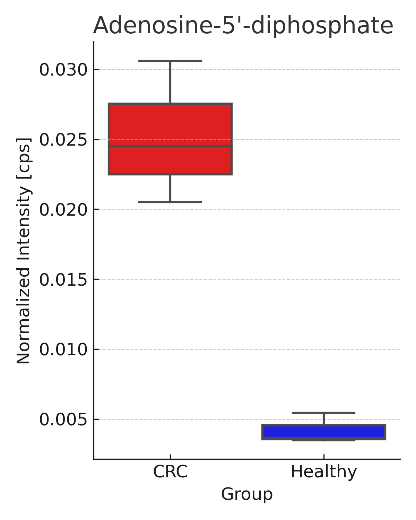

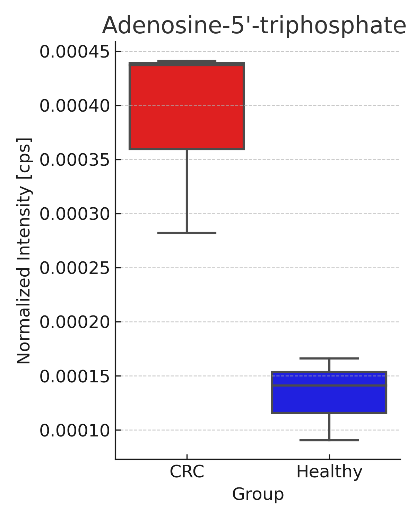

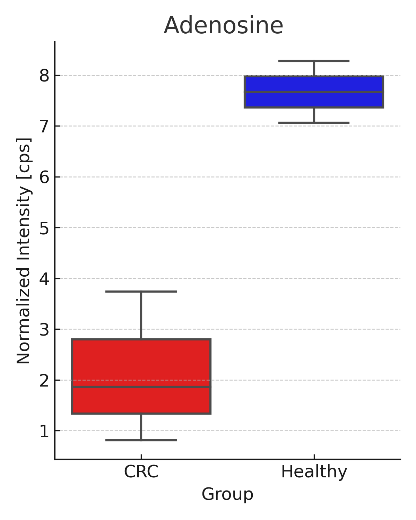

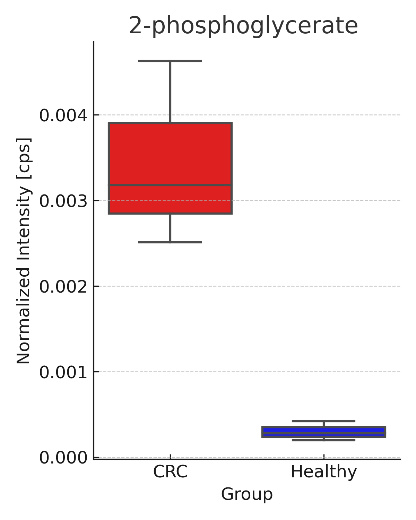

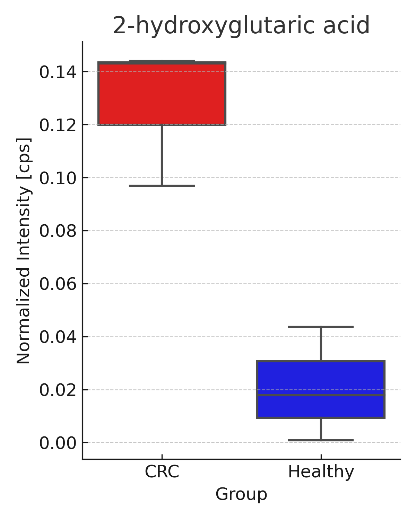

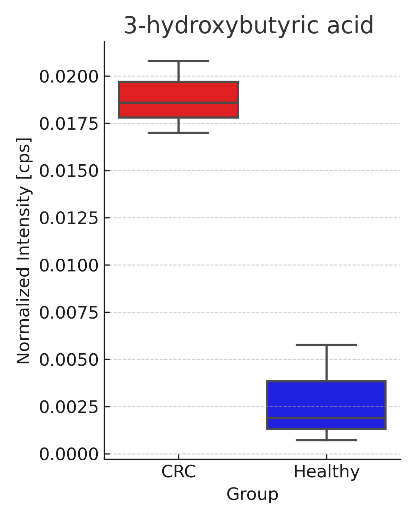

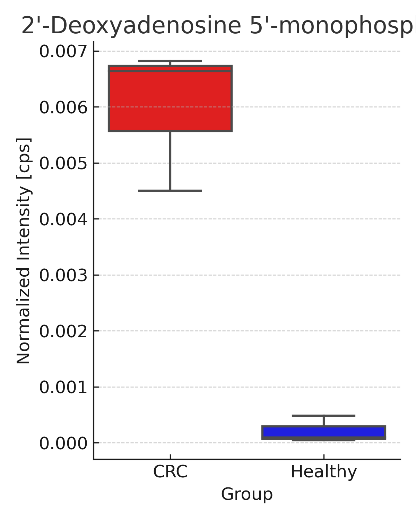

### **Figure S2** | Boxplots showing normalized intensities [cps] of the significantly regulated metabolites (p < 0.05, |log₂ fold change| > 1.0) between healthy and tumor-derived colorectal organoids (n = 3). Each plot displays relative abundance based on normalized signal intensities.

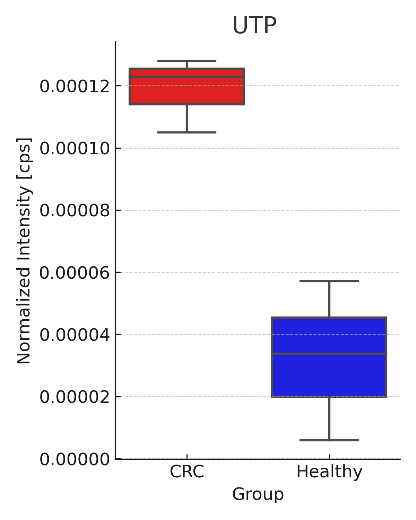

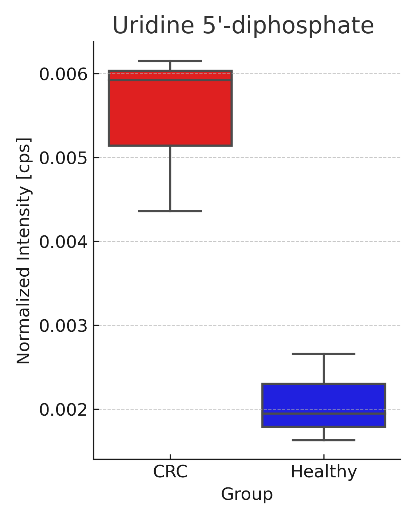

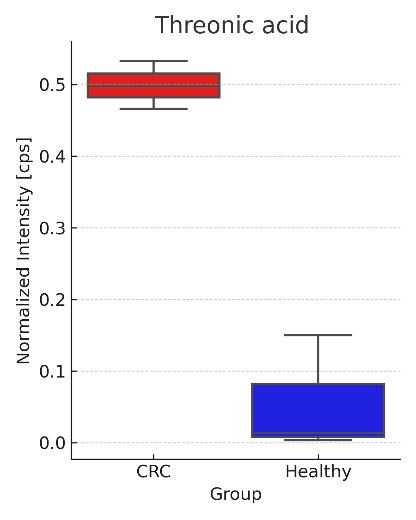

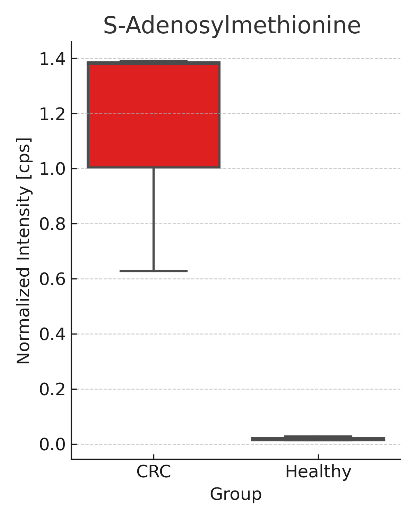

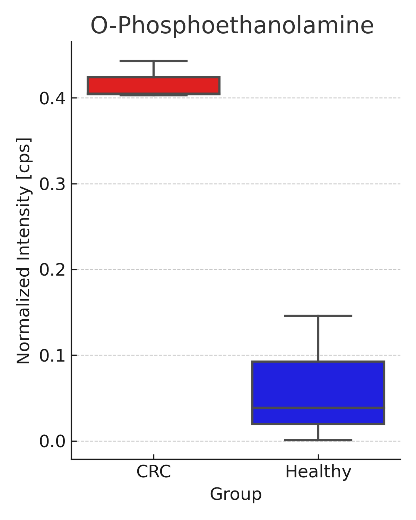

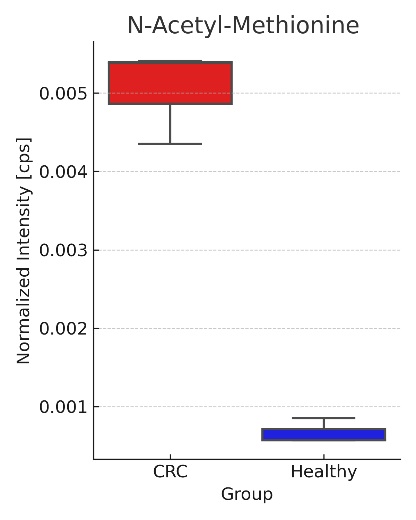

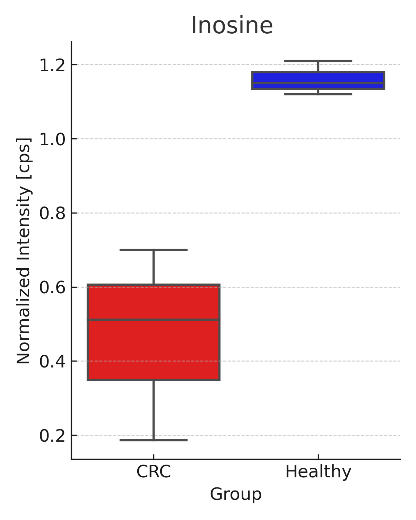

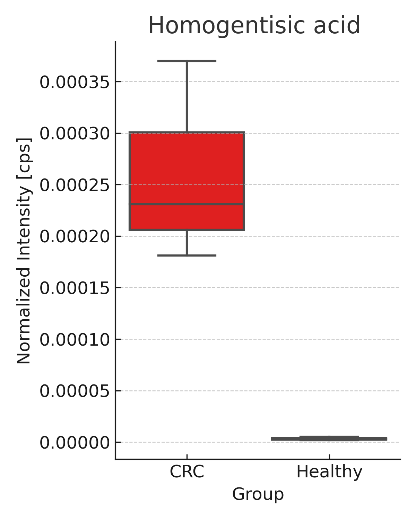

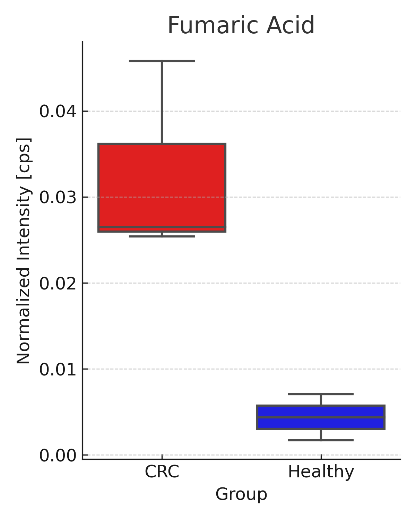
